# Proteolysis Targeting Chimeras With Reduced Off-targets

**DOI:** 10.1101/2021.11.18.468552

**Authors:** Tuan M. Nguyen, Vedagopuram Sreekanth, Arghya Deb, Praveen Kokkonda, Praveen K. Tiwari, Katherine A. Donovan, Veronika Shoba, Santosh K. Chaudhary, Jaron A. M. Mercer, Sophia Lai, Ananthan Sadagopan, Max Jan, Eric S. Fischer, David R. Liu, Benjamin L. Ebert, Amit Choudhary

## Abstract

Proteolysis Targeting Chimeras (PROTACs), a class of heterobifunctional molecules that recruit target proteins to E3 ligases, have gained traction for targeted protein degradation. However, pomalidomide, a widely used E3 ligase recruiter in PROTACs, can independently degrade other targets, such as zinc-finger (ZF) proteins, that hold key functions in normal development and disease progression. This off-target degradation of pomalidomide-based PROTACs raises concerns about their therapeutic applicability and long-term side effects. Therefore, there is a crucial need to develop rules for PROTAC design that minimize off-target degradation. In this study, we developed a high-throughput platform that interrogates the off-target degradation of ZF domains and discovered, using this platform, that PROTACs with the current design paradigm induce degradation of several ZF proteins. To identify new rules for PROTAC design, we generated a library of pomalidomide analogs that allowed systematic exploration of the impact of positional isomerism (e.g., C4 and C5 positions of the phthalimide ring), hydrogen bonding, steric and hydrophobic effects on propensities for ZF protein degradation. We found that modifications of appropriate size on the C5 position reduced off-target ZF degradation. We validated these results using immunoblotting, target engagement, and global mass spectrometric studies. We applied our newfound design principles on a previously developed ALK oncoprotein-targeting PROTAC and generated PROTACs with enhanced potency and minimal off-target degradation. We envision the reported off-target profiling platform and pomalidomide analogs will find utility in design of specific PROTACs.

## INTRODUCTION

Imide-based molecular glues (e.g., pomalidomide) induce proximity between cereblon (CRBN), the substrate receptor for an E3 ubiquitin ligase, and proteins with Zn-finger (ZF) motifs to trigger ubiquitination and degradation of the latter.^1–6^ Pomalidomide is often appended to target protein binders to generate CRBN-based Proteolysis Targeting Chimeras (PROTACs) that induce proximity-mediated target protein degradation.^7–9^ However, these pomalidomide-based PROTACs can also recruit other proteins with or without ZF motifs that serve key biological functions in normal development and disease progression.^10–14^ For example, tissue-specific deletion of pomalidomide-degradable ZF protein, ZFP91, in regulatory T cells (Tregs) leads to Treg dysfunction and increases the severity of inflammation-driven colorectal cancer.^15^ Furthermore, numerous other proteins with important roles in cellular function, such as transcription factors, harbor ZF domains.^16–17^ The off-target degradation of these key ZF-containing proteins may have long-term implications, such as the development of new cancers, dysregulation of lymphocyte development, and teratogenic effects.^18–22^ The ability of pomalidomide to degrade other proteins in a PROTAC-independent manner raises concerns about the precariousness of off-target ubiquitination and degradation of these compounds, several of which are already in clinical trials.^23–24^ Thus, there is an urgent need for robust, sensitive, and high-throughput methods that can determine off-target degradation of such PROTACs.^25^

Currently, off-target degradation can be assessed by mass–spectrometry-based methods^26–29^ that detect protein levels, but these techniques lack sensitivity for low abundant proteins.^30^ In addition to the expense, the implementation of mass spectrometry is technically challenging and requires examination PROTACs across multiple tissue types owing to lineage-specific protein expression.^31^ These analyses are further complicated by the need to perform the off-target assessments under various dose and temporal regimen of PROTAC administration.

Herein, we report the development of an inexpensive, sensitive, robust, and high-throughput imaging platform that profiles the off-target degradation propensities of ZF domains by measuring the decrease in fluorescence of a panel of GFP-tagged various ZF domains upon compound treatment. ^32^ We validated this platform using immunoblotting, target engagement, and global proteomics experiments. With this validated platform, we profiled off-target activities of literature-reported PROTACs with different linker chemistries. Surprisingly, nearly all the profiled PROTACs exhibited off-target degradation of ZF domains. To reduce this, we rationally designed and generated approximately 80 pomalidomide analogs and profiled them for off-target activity. We discovered and validated that substitution at specific positions (C5) on pomalidomide reduced degradation propensities. With these findings, we generated new PROTACs that target anaplastic lymphoma kinase (ALK) with reduced off-target ZF degradation and enhanced on-target potencies. Overall, we developed an off-target profiling platform and used that platform to identify pomalidomide analogs with lower off-target propensity and then deployed such analog to generate a specific PROTAC for ALK.

## RESULTS AND DISCUSSION

### Development and validation of an off-target profiling platform for PROTACs

To profile the ZF degradation propensity of pomalidomide and its analogs, we first developed an automated imaging assay (Figure 1A). For this platform, we selected 23–amino-acid ZF degrons of 11 ZF proteins that are reportedly degraded by pomalidomide and 3 ZFs that are not (Supplementary Table S1).^32^ We inserted these ZF degrons into a lentiviral degradation reporter vector (cilantro 2) ^32^ to compare the fluorescence of ZF-tagged enhanced green fluorescent protein (eGFP) to untagged mCherry (Figure 1A). With this assay, wherein compounds are tested against the 14 stable U2OS cell lines containing our tagged ZF-proteins, we reliably detected dose-dependent degradation of ZF degron-tagged eGFP by pomalidomide with a robust readout (Z’ =0.8) (Figure S1). As expected, pomalidomide degraded all 12 ZF degrons that are sensitive to it in a dose-dependent manner that ranged from 4.3 nM to 6 μM (Figure 1A). Unlike mass spectrometry, this reporter-based method is not limited by cell–type-specific expression levels of analyte proteins nor by the accessibility to ZFs in the context of full-length proteins that are engaged in protein complexes. Thus, this method may have enhanced sensitivity over mass spectrometry-based methods for the detection of pomalidomide-sensitive ZF protein degradation.

**Figure 1.**
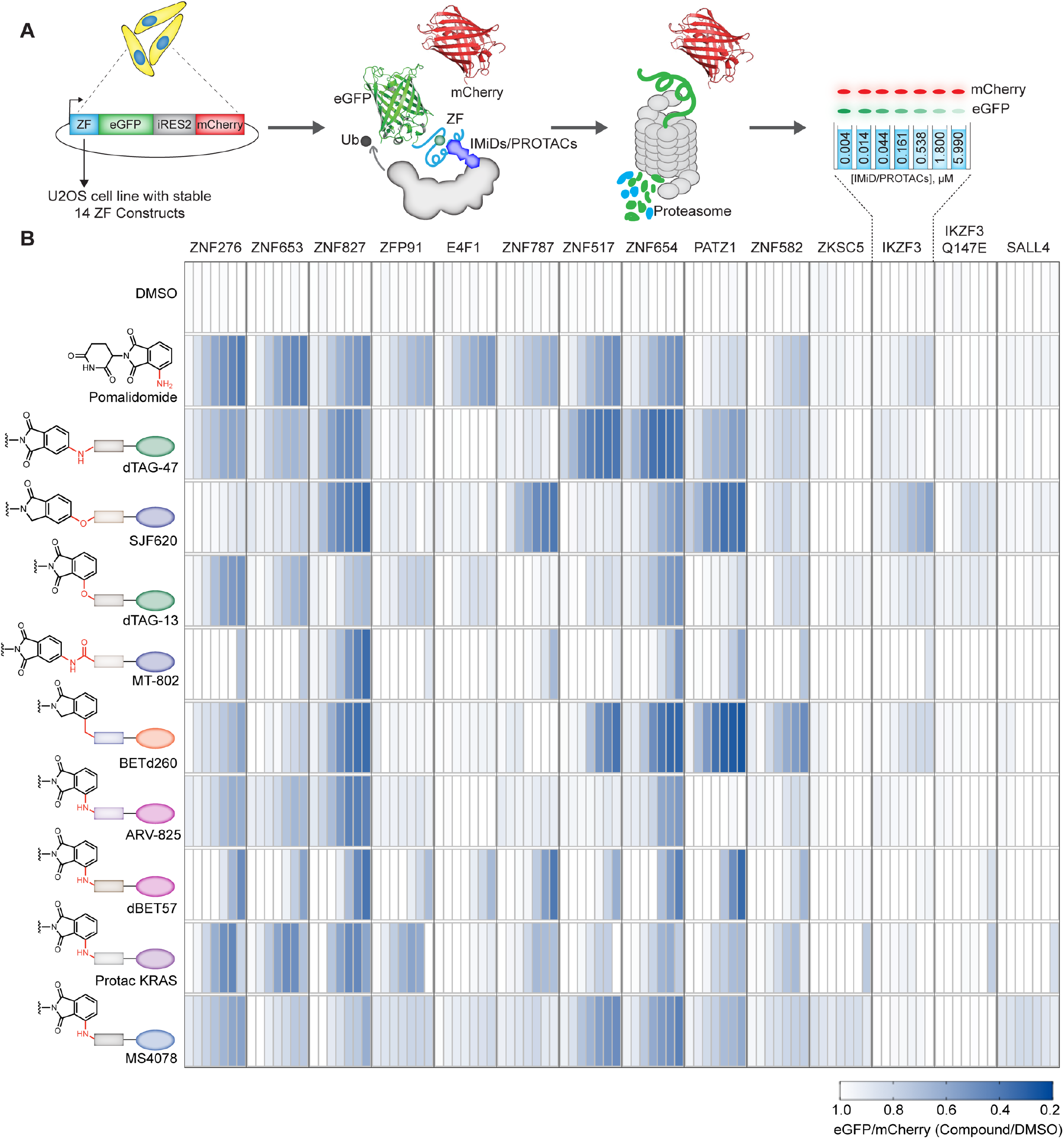
Development of a high-throughput assay for evaluating off-target ZF degradation of poma-lidomide-based PROTACs. **(A)** Schema of the automated imaging screen for the degradation of ZF degrons by pomalidomide analogs and pomalidomide-based PROTACs. Briefly, U2OS cells stably expressing 14 ZF degrons fused to eGFP were treated with PROTACs followed by imaging to assess ZF degradation. **(B)** Degradation of validated and pomalidomide-sensitive ZF degrons inside cells by reported PROTACs in a dosing range of 4.3 nM to 6 μM.

With this assay in hand, we profiled the off-target activity of 9 reported PROTACs with varying exit vectors from pomalidomide end and linker lengths (Figure 1B, Figure S2). We observed degradation of many ZF-domains with almost all the 9 PROTACs (Figure 1B). Notably, PROTACs with common exit vectors, such as arylamine, - ether, -carbon, and -amide, generally had greater ZF degradation capabilities in similar fashion to pomalidomide (Figure 1B).

To validate this imaging platform, we determined the relative stability of the ternary complex formed by the PROTACs with CRBN and the off-target ZFP91 using a reported nanoBRET assay (Figure 2A).^36^ Here, the PROTACs induce a bioluminescence resonance energy transfer (BRET) between luciferase (appended to ZFP91) and a fluorophore (appended to CRBN via a HaloTag domain). We generated a cell line that stably expresses ZFP91-nanoluciferase fusion and transfected these cells with the HaloTag-fused CRBN (HT-CRBN) followed by treatment with a fluorophore (i.e., NanoBRET-618 ligand) and PROTACs (Figure 2A). We observed a BRET signal for the PROTACs, suggesting ternary complex formation between CRBN, off-target ZFP91, and PROTACs (Figure 2B). Since ternary complex formation does not guarantee target degradation, we also confirmed the off-target degradation of endogenous ZF protein such as ZFP91 by reported PROTACs MS4078 (ALK PROTAC, Figure 2C, S3)^33^ and dTAG-13 (FKBP12^F36V^ PROTAC, Figure 2D)^34–35^, using immunoblotting and by global proteomic analysis (Figure 2C, 2D, and S3).

**Figure 2.**
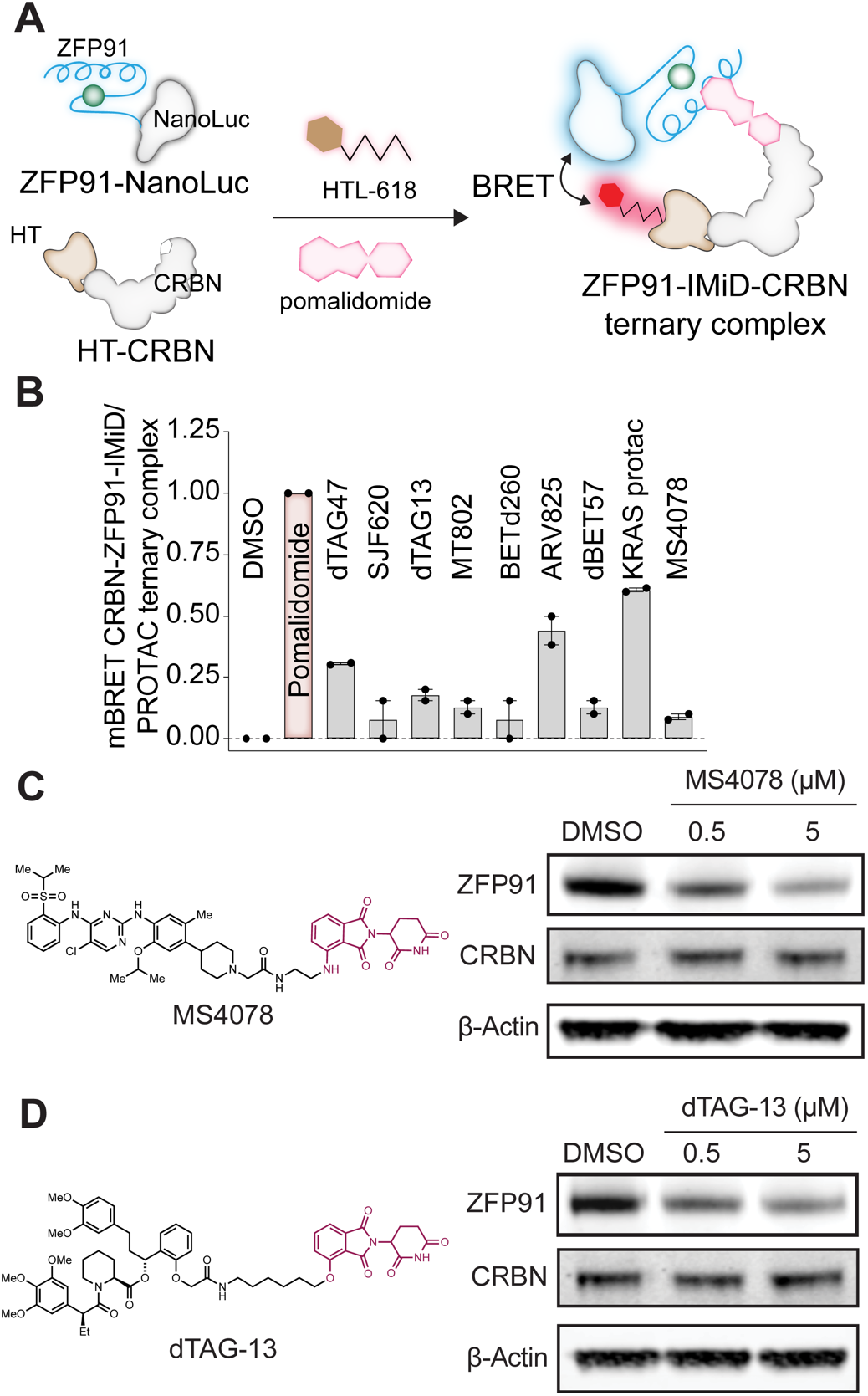
Validation of image-based results using NanoBRET and immunoblotting. **(A)** A nano-BRET assay to quantify ternary complex formation between ZFP91, CRBN, and imide analogs or PROTACs. **(B)** BRET ratio of reported PROTACs treated at 1 μM dose in a NanoBRET assay. **(C-D)** Immunoblots demonstrating off-target degradation of endogenous ZF protein ZFP91 by **(C)** MS4078 (ALK PROTAC) and **(D)** dTAG-13 (FKBP12^F36V^PROTAC) in a dose-dependent manner in Jurkat cells.

Next, we queried the influence of exit vectors on pomalidomide-based PROTACs and the degradation of ZFs. Towards this goal, we analyzed changes in endogenous ZF proteins from 124 proteomics datasets that were generated for cells treated with pomalidomide-based PROTACs.^31^ In Figure S4, we show the relative abundance of proteins that contained the ZF motif as previously described^32^ and were detectable in at least 1 proteomics dataset (i.e., 284 ZF proteins). We computed a ZF degradation score for every PROTAC dataset by taking the sum of ZF protein abundance. Analyzing the degradation score distribution confirmed that PROTACs had ZF protein degradation activity for amino acetamide and arylamine, -ether, and -carbon exit vectors (Figure S4). The agreement between our automated imaging assay and proteomics suggests that we can apply our methodology in a high-throughput manner to identify pomalidomide analogs that confer minimal ZF degradation. Furthermore, we can develop a new set of rules for PROTAC development to minimize off-target degradation of endogenous ZF proteins.

### Generation of a library of rationally designed pomalidomide analogs

We next created a library of rationally designed pomalidomide analogs that can provide insights into design of pomalidomide-based PROTACs with minimal off-target ZF degradation. We gained structural insight from the crystal structure of the DDB1-CRBN-pomalidomide complex bound to transcription factor IKZF1 (PDB: 6H0F). ^32^ In the crystal structure, the glutarimide ring of pomalidomide is deeply buried inside CRBN, while the phthalimide ring is accessible for modification. Q147 of IKZF1 forms a water-mediated hydrogen-bonding interaction with the C4 amino group of the compound, while the C5 position is proximal to the ZF domain (Figure 3A).^32^ Mutation of Q147, or equivalent residues, has been reported to abrogate IKZF degradation.^32^ We hypothesized that appropriate substitutions at the C4 and/or C5 positions could disrupt the ternary complex of the ZF domains with CRBN while maintaining its interaction with CRBN through glutarimide ring. To investigate this, we first synthesized a thalidomide analog, 5-aminothalidomide (8, Figure 3B). The treatment of MM1.S cells with this C5-amino analog revealed a decrease in overall degradation potency as compared to pomalidomide (**1**, Figure 3B, C), suggesting that modifications on the C5 position will “bump-off,” or eliminate, the endogenous ZFs. Therefore, we generated approximately 80 pomalidomide analogs (Schemes 1-4) with increasing size and diverse modifications on the C4 and C5 positions of the phthalimide ring.

**Figure 3.**
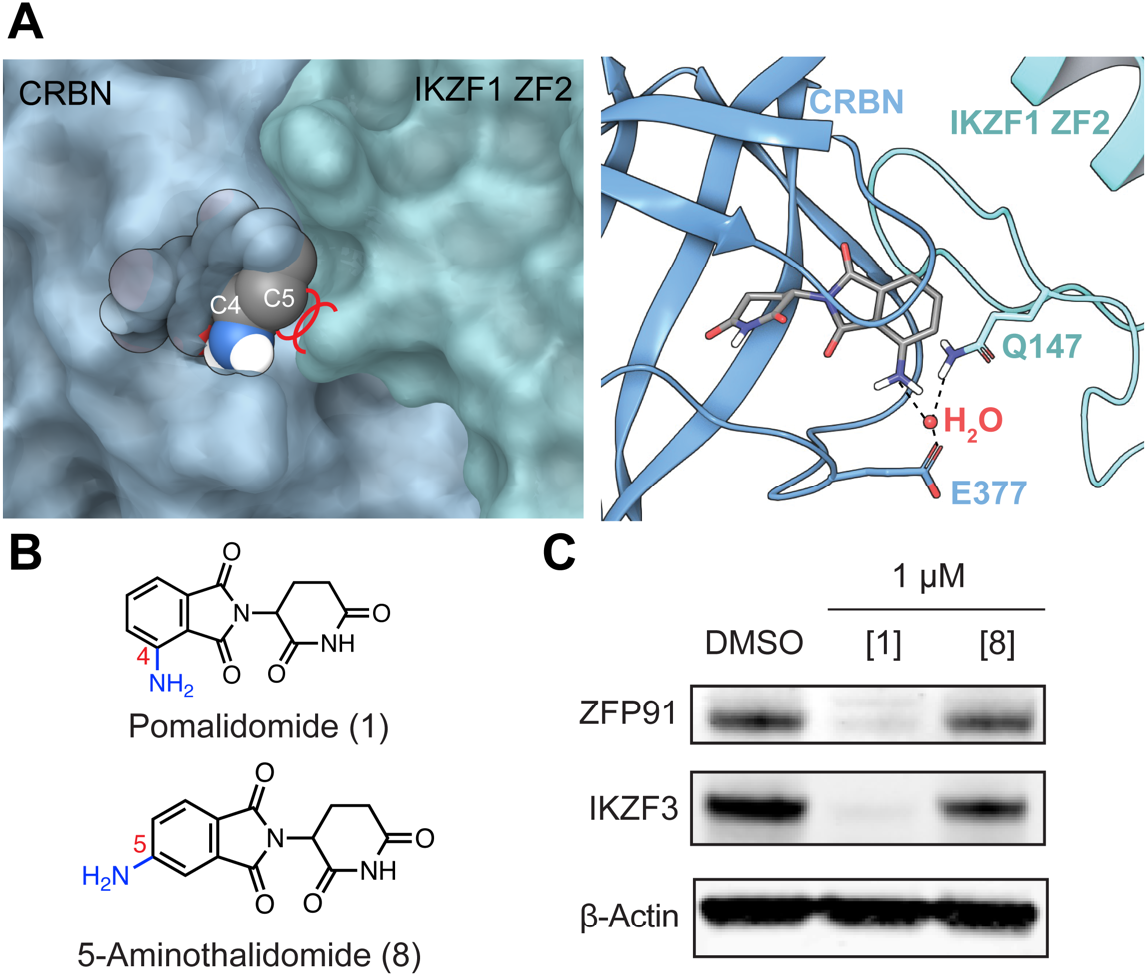
Structural insights and degradation potential of the thalidomide analogs with C4 and C5 amino groups on the phthalimide ring. **(A)** Crystal structure showing the glutarimide ring deeply buried in the CRBN exposing the 4-amino group of pomalidomide, which makes a crucial water-mediated hydrogen bond between CRBN (E377) and IKZF1 (Q147). Modification on C5 position would potentially bump-off the ZF degrons (PDB: 6H0F). **(B)** Structures of pomalidomide (**1**) and 5-aminothalidomide (**8**). **(C)** Immunoblots of endogenous ZF proteins ZFP91 and IKZF3 in MM1.S cells treated with pairs of imide-based analogs with C4 (pomalidomide) and C5 amino modifications on the phthalimide ring.

To rapidly and systematically construct the library of pomalidomide analogs, we leveraged robust reactions such as nucleophilic aromatic substitution (S_N_Ar), Suzuki, Sonogashira cross-couplings, and amidation with commercially available imide compounds for the facile and scalable incorporation of C4 and C5 substitutions on the phthalimide ring (Schemes 1–4 and Figure S5). We synthesized the pomalidomide analogs in pairs at C4 and C5 positions and generated a library that can be categorized into three main synthetic groups: C–N (S_N_Ar), C–C (Suzuki/Sonogashira cross-coupling), and N–C (acylation). We employed several aliphatic amines with variable sizes for high-yielding S_N_Ar reactions with 4- and 5-fluorothalidomides. For the S_N_Ar library, we incorporated *N*-Boc-piperazine and *N*-Boc diazaspiro[3.3]heptane, which can subsequently be used as handle to append target binder. In addition, we synthesized several pomalidomide analogs with a fluoro substitution at the 6-position of thalidomide (Scheme 2). We utilized Suzuki/Sonogashira cross-coupling reactions on 4- and 5-bromo thalidomides with heterocyclic boronic acids and phenylacetylene (Scheme 3) to prepare the imide analogs (**50**—**61**). We carried out amidation with a diverse class of acid chlorides varying from aliphatic to heterocyclic cores. We systematically varied the sizes from a small acetyl group (**62** and **72**) to the largest cubanoyl group (**71** and **81**) (Scheme 4). The physicochemical properties of the overall library encompass a reasonable distribution of druglike properties, such as partition coefficient, or cLogP (−2 to 4), molecular weights (250–550 g/mol), and topological polar surface area (tPSA, 80 to 140 Å^2^) (Figure S6D).

### Systematic evaluation of ZF protein degradation propensity of pomalidomide analogs

Using our developed off-target profiling platform, we tested the library of pomalidomide analogues to derive rules for the impact of exit vector modifications on pomalidomide and ZF protein degradation. First, we observed that analogs with C5 modifications on the phthalimide ring had reduced ZF degradation relative to identical modifications on the C4 position, particularly for S_N_Ar (C-N) analogs (Figures 4A, S7A). The trend was less for modifications made with amidation (N-C) and Suzuki/Sonogashira cross-couplings (C-C), partly due to having smaller sets of ana logs compared with the S_N_Ar group (Figure S7B-C). Our structure modeling and docking data also suggest that

**Scheme 1:**
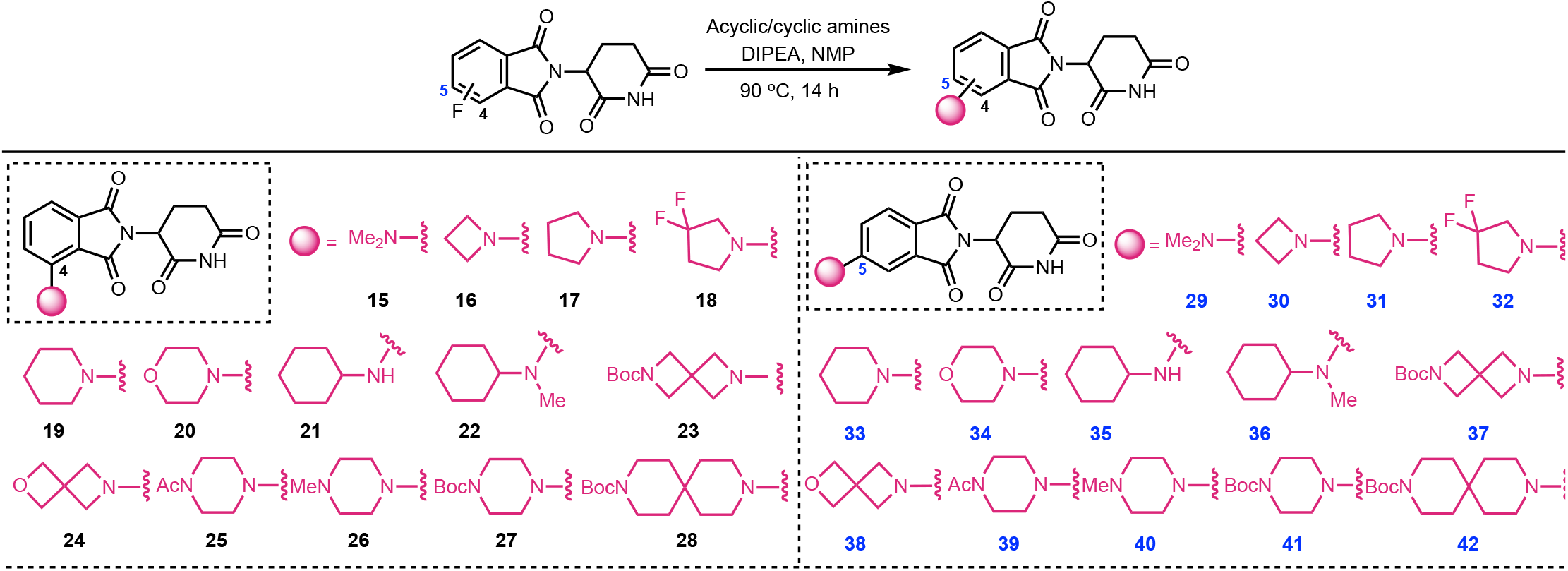
Synthesis of imide analogs *via* nucleophilic aromatic substitution (S_N_Ar)

**Scheme 2:**
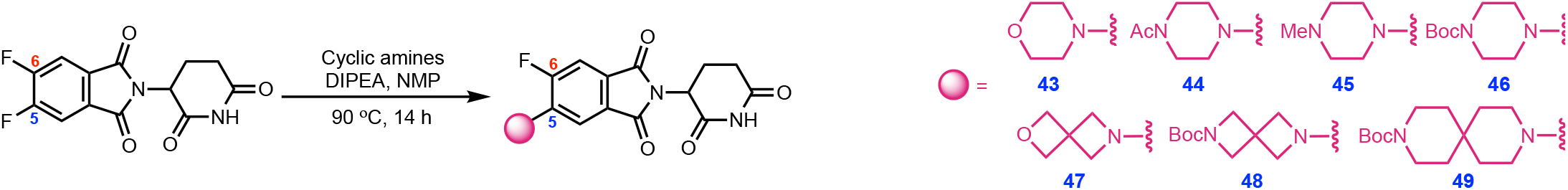
Synthesis of 6-Fluoro substituted imide analogs *via* nucleophilic aromatic substitution (S_N_Ar)

**Scheme 3:**
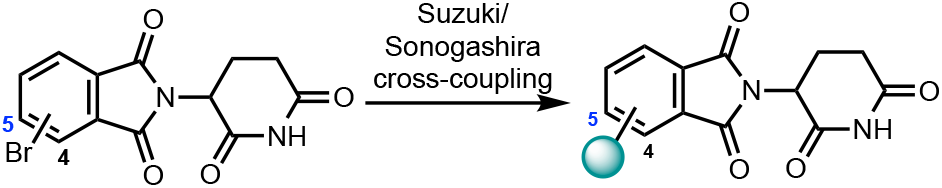
Synthesis of imide analogs *via* cross-coupling

**Scheme 4:**
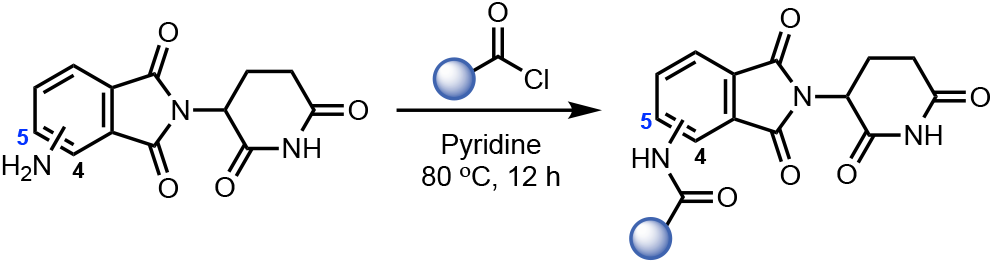
Synthesis of imide analogs *via* amidation

**Scheme 1-4:**
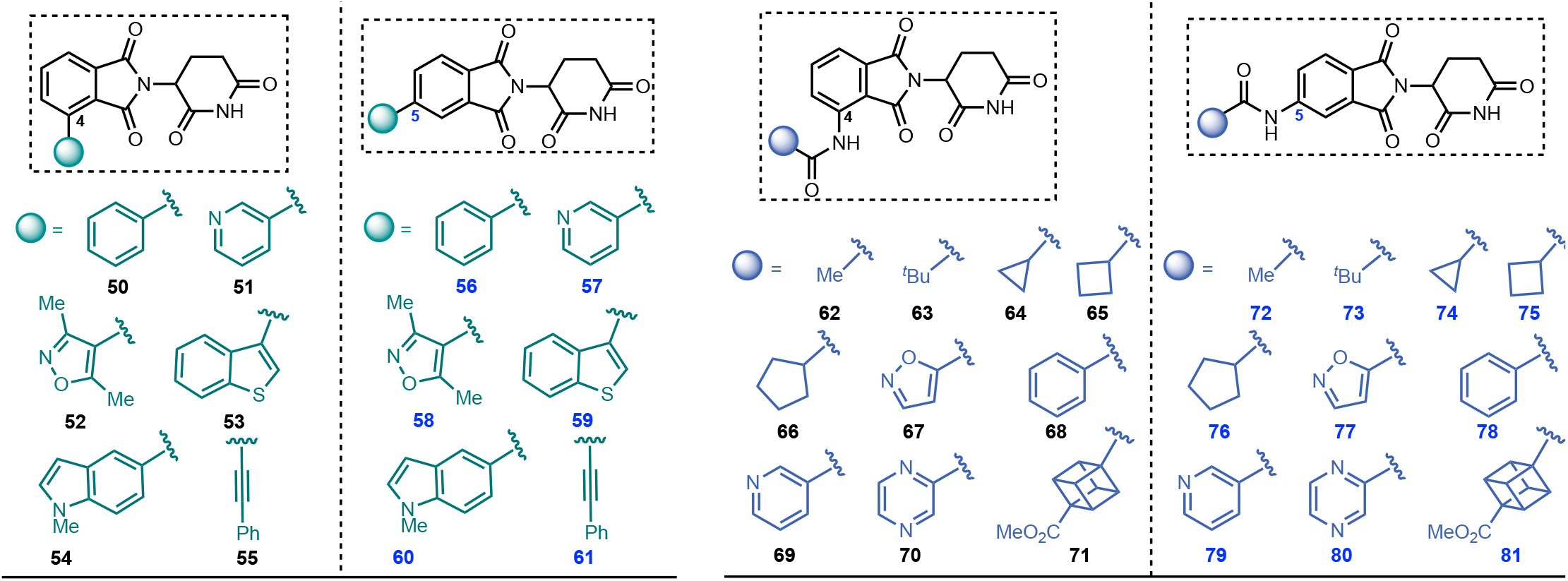
Synthesis of imide analogs through SNAr, cross-coupling, and amidation chemistries.

C5 modifications are more likely to create a steric clash between the S_N_Ar exit vectors and the ZF domain than C4 position modifications (Figure S6A-C). Second, analogs lacking hydrogen-bond (H-bond) donors immediately attached to the phthalimide ring had reduced ZF degradation compared to those with H-bond donors, regardless of the position of the modification relative to the ring (Figure 4B), in agreement with ZF degradation pattern observed with the reported PROTACs (Figure 1B). Furthermore, the immunoblotting studies using **21** and **22** (Figure 4C) are in agreement with the high-throughput imaging data regarding degradation of endogenous ZF proteins, ZFP91 and IKZF3. Taken together, these data reveal that pomalidomide-based PROTACs with an arylamine exit vector, where NH-is a H-bond donor, induced greater ZF degradation and suggest that PROTACs without H-bond donors will have lower off-target activity. This finding is consistent with reports that pomalidomide can stabilize the ternary complex between ZF and CRBN using H-bonds.^32^ We aimed to further minimize off-target ZF degradation in S_N_Ar analogs with C5 position modifications through the addition of a fluoro group at the C6 position (Schemes 1-4, Figure S5). The fluoro group reduced ZF degradation for most S_N_Ar exit vectors, such as -acetylpiperazine and morpholine, but not for diazaspiro[3.3]heptane (Figures S7A and S7D).

**Figure 4.**
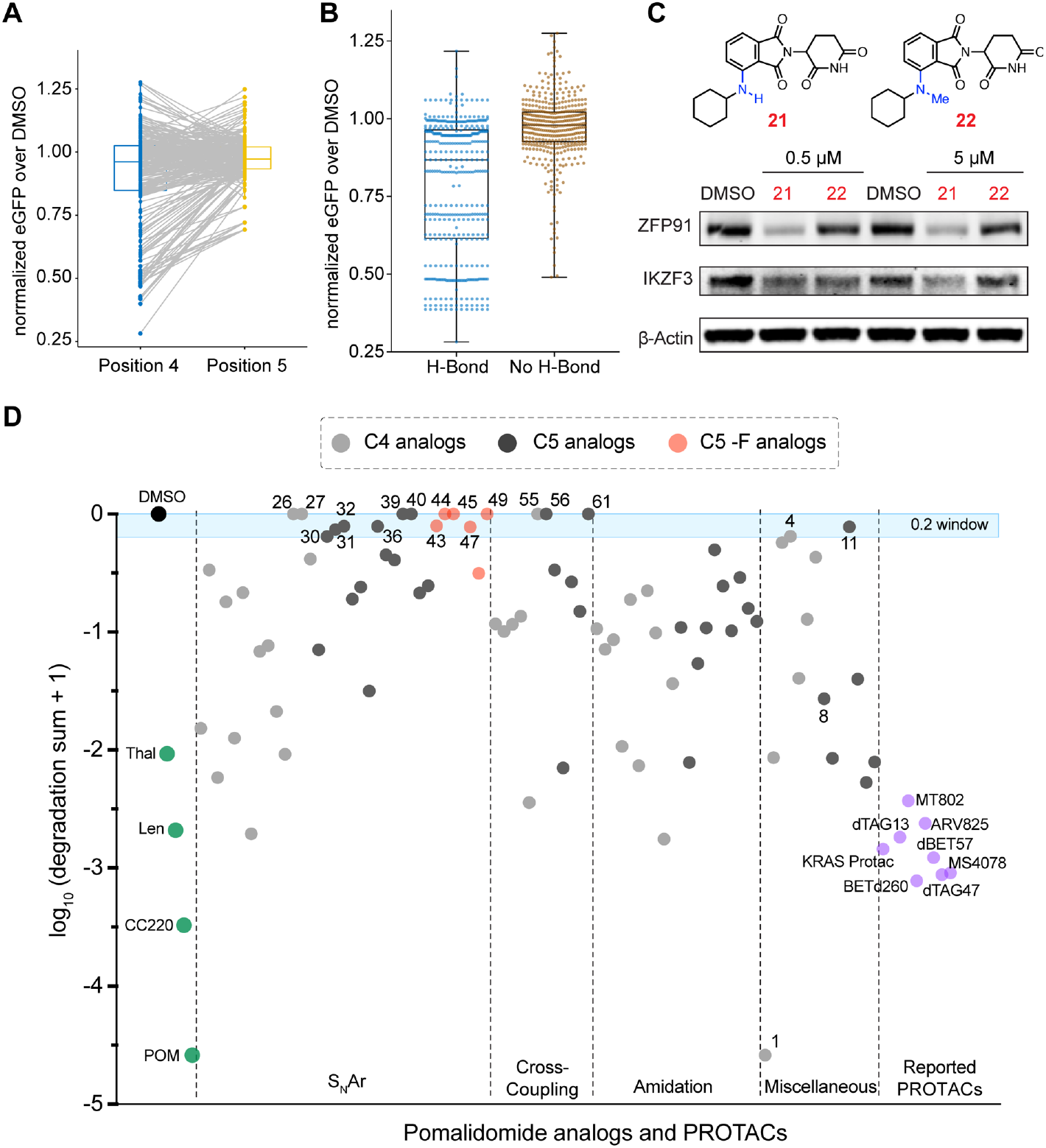
Degradation score as metric to nominate the imide analogs with reduced off-targets. **(A)** Plot showing pair-wise comparison of GFP degradation levels induced by pairs of S_N_Ar pomalidomide analogs with C4 and C5 modifications (Wilcoxon matched-pairs signed rank test; p = 0.0029) at the same dose. **(B)** Degradation of pomalidomide-sensitive ZF degrons inside cells by pomalidomide analogs with and without H-bond donor(s) immediately adjacent to the phthalimide ring. (two tailed Welch’s T-test; p < 0.0001) **(C)** Immunoblots for endogenous ZF proteins ZFP91 and IKZF3 in Jurkat cells treated with pomalidomide analogs (**21**, **22**) with and without a H-bond donor (NH) immediately attached to the phthalimide ring. **(D)** Scatter dot plot showing ZF degradation scores [i.e. -Log(degradation sum + 1)] for individual pomalidomide analogs and pomalidomide-based PROTACs investigated in this study.

Next, we aimed to identify exit vector modifications that confer the least off-target ZF degradation. Towards this goal, we derived a degradation score for each pomalidomide analog, including pomalidomide-based PROTACs, by taking the sum of eGFP degradation values for the ZF degrons at multiple doses for each analog (Figure 4D). Compounds with degradation scores close to zero induced the least ZF degradation, whereas compounds with the most negative scores induced the most ZF degradation (Figure 4D). Analogs with piperazine (**39** and **40**), and alkyne (**55** and **61**) exit vectors predominated the group of compounds with degradation scores of 0 (Figure 4D, S8). Other exit vectors with minimal degradation scores include phenyl (**56**), azetidine (**30**), pyrrolidine (**31**), 3,3-difluoropyrrolidine (**32**), *N*-methylcyclohexylamine (**36**), and *N*-Boc-diazaspiro[3.3]heptane (**37**) as well as 6-fluoro substituted analogs of morpholine (**43**), *N*-protected piperazines (**44** and **47**), 2-oxa-6-azaspiro[3.3]heptane (**45**), and diazaspiro[5.5]undecane (**49**) (Figure 4D, S8). From this study, we established two main rules for designing pomalidomide-based PROTACs to minimize off-target effects. First, exit vectors should predominantly have modifications on the C5 position. Second, the H-bond donors immediately adjacent to the phthalimide ring preferably should be masked.

### Imaging platform validation using CRBN engagement, ternary-complex stability, and global proteomics studies

We first demonstrated CRBN engagement by the imide analogs with close to zero degradation scores in cells using a NanoBRET-based CRBN binding assay. Here, the imide analogs compete with the fluorophore-labeled CRBN binder and the decrease the BRET signal (Figure 5A) correlates with the CRBN engagement. All the analogs showed a dose-dependent decrease in the BRET signal indicative of CRBN engagement (Figure 5B). We next evaluated the relative stability of the ternary complex formed by these imide analogs with CRBN and the off-target ZFP91 using a nanoBRET assay.^36^ In agreement with the results of our imaging platform, we observed reduced BRET signal for the analogs compared to that of pomalidomide, suggesting reduced ternary complex formation between CRBN, ZFP91, and analogs (Figure 5C, S9). Furthermore, imide analogs with H-bond donor group (e.g., analogs obtained using amidation chemistry) formed stronger ternary complex compared to analogs with C-N bond (S_N_Ar chemistry) or C-C (cross-coupling chemistries) bonds (Figure S9).

**Figure 5.**
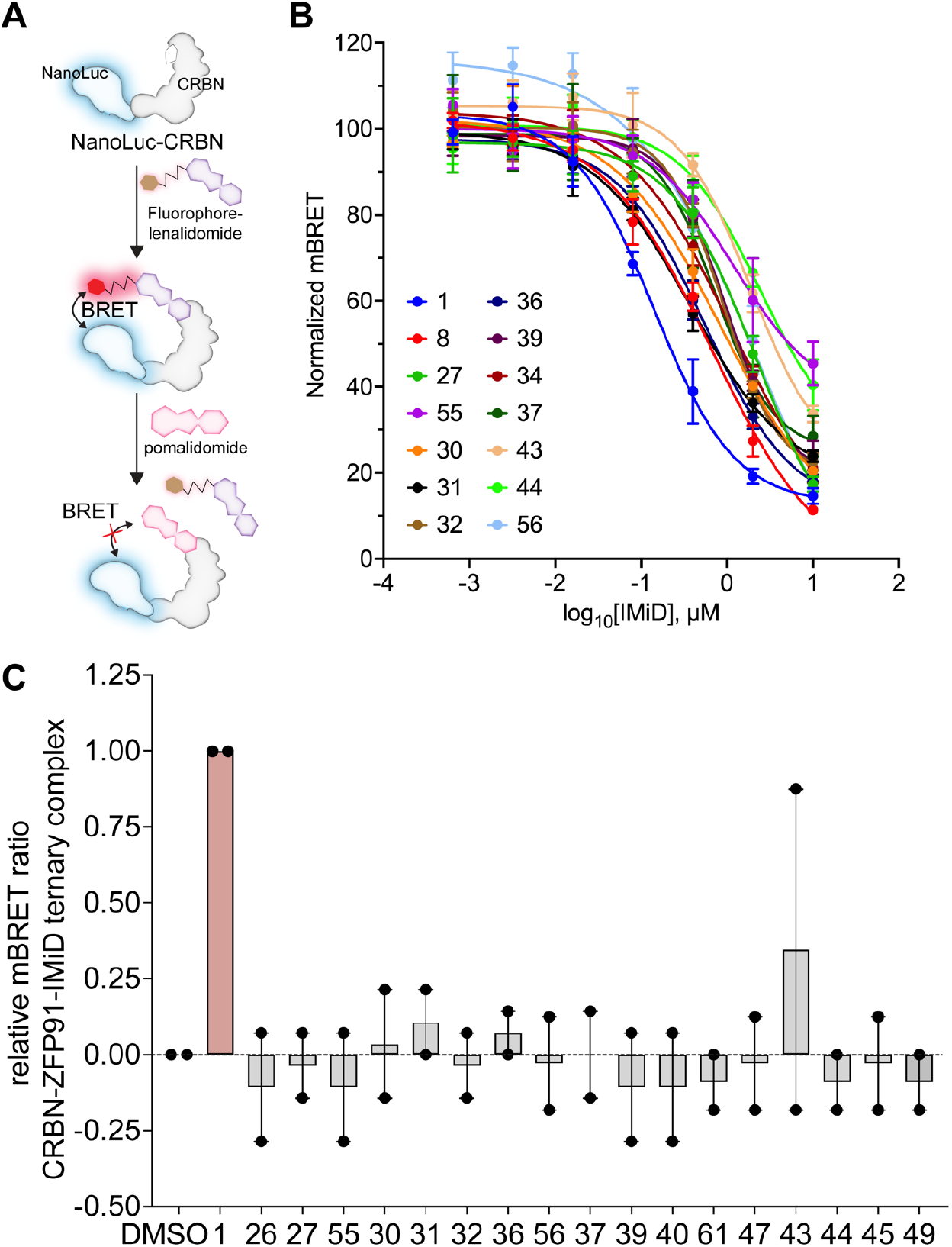
Cellular target engagement studies. **(A)** A competition based NanoBRET assay for CRBN engagement in cells by the pomalidomide analogs. **(B)** Dose-dependent CRBN binding curves of selected analogs determined in U2OS cells using NanoBRET-based intracellular CRBN binding assay. **(C)** BRET ratio for selected analogs treated at 1 μM dose in a NanoBRET based ternary complex formation assay (see Figure S9 for the ternary complex analysis of all the imide analogs generated in this study).

We next performed mass spectrometry-based global proteomics of MOLT4 (Figure 6) and KELLY cells (Figure S10) treated with imide analogs. These cells were treated with pomalidomide or structurally diverse imides with degradation scores close to zero and the lysates were analyzed by mass spectrometry. Pomalidomide (**1**) triggered the degradation of several ZF-proteins (e.g., IKZF1, ZFP91, and SALL4 in KELLY cells) and other off-targets (e.g., RAB28, DTWD1, CUTA, POLR2J, Figure 6A, S10A). Analogs with C5 substitutions with piperazine (**39**, Figure 6B, S10B), dia-zaspiro[3.3]heptane (**37**, Figure 6C, S10C) and phenyl (**56**, Figure 6D, S10D), 2-oxa-6-azaspiro[3.3]heptane (**38**, Figure 6E, S10E), 3,3-difluoropyrrolidine (**32**, Figure 6F, S10F), *N*-methylcyclohexanamine (**36**, Figure 6G, S10G) phenylacetylene (**61**, Figure 6H, S10H) did not trigger degradation of any common zinc finger proteins or other off-targets. Taken together, these studies validate the platform and has led to identification of a collection of exit vectors to guide the development of pomalidomide-based PROTACS with reduced l off-target ZF degradation.

**Figure 6.**
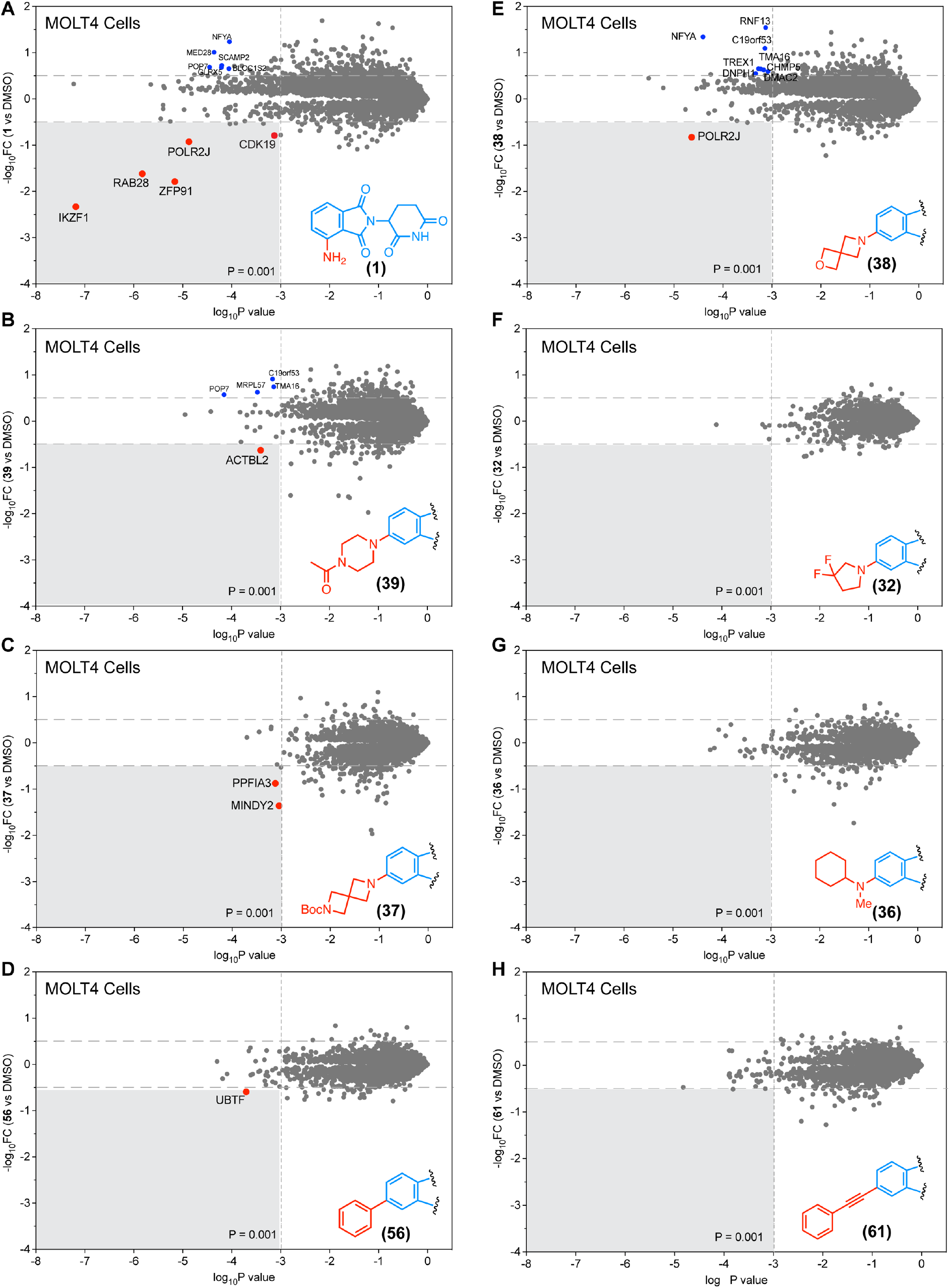
Validation of image-based platform by global proteomics. **(A-H)** Global proteomic analysis of MOLT4 cells treated with analogs **1** (A), **39** (B), **37** (C), **56** (D), **38** (E), **32** (F), **36** (G), and **61** (H).

### Development of PROTACs with reduced off-target degradation propensities

As an application of imaging platform and identified analogs with reduced off-targers, we re-engineered the reported ALK PROTAC (MS4078),^33^ which had a high level of off-target ZF degradation (Figure 1B, 2C), by altering the exit vectors on pomalidomide to reduce off-target ZF degradation while maintaining on-target degradation. We selected pomalidomide analogs, including alkyne, piperazine on the C5 positions, and 2,6-diazaspiro[3.3]heptane with a C6 fluoro modification, PROTAC development owing to their near-zero ZF degradation score (Figure 4D, S8). We rationally designed and synthesized 12 new ALK PROTACs with different exit vectors, such as an alkyne (C4: dALK-1 and C5: dALK-2), piperazine with varying alkyl (dALK-3 to dALK-6) and acyl linkers (dALK-7 to 10), and diazaspiro[3.3]heptane (dALK-11 and dALK-12) by Sonogashira cross-coupling, reductive amination and amidation chemistry, respectively (Figure S11). To investigate the effect of the fluoro group on the on- and off-target propensities, we also synthesized the corresponding fluoro pairs for non-alkyne dALK PROTACs (dALK-4, 6, 8, 10, and 12) (Figure 7A). We also synthesized (5)-MS4078 (Figure S11D), which has a C5 substitution (vs. C4 substitution) for a direct comparison and performed the off-target analysis of these ALK PROTACs. C5 substituted derivative of MS4078 (i.e., (5)-MS4078) reduced the off-targets but retained its ability to degrade ZNF276, ZNF827, and PATZ1 (Figure 7B). The re-engineered ALK PROTAC with C5 alkyne exit vector (dALK-2) dramatically reduced the off-target effects of the original PROTAC MS4078, reducing the affinity for proteins such as ZNF517, ZNF654, ZNF276, ZNF653, and PATZ1 (Figure 7B). New ALK PROTACs with piperazine exit vectors with and without the fluoro group exhibited minimized off-target effects (Figure 7B). In agreement with imagebased off-target analysis, these re-engineered dALK PROTACs have a reduced ternary complex formation with off-target ZFP91 (Figure S9) and so does ARV110, a PROTAC under clinical trials (Figure S9). In agreement with these findings, the global proteomics studies revealed that while both MS4078 and dALK-10 induced degradation of ALK (Figure S12A, B), MS4078 also induced degradation of SALL4 (Figure 8A), an off-target associated with teratogenicity, while dALK-10 did not (Figure 8B). Finally, we confirmed that the redesigned PROTACs reduced the viability of the ALK dependent SU-DHL-1 cells (Figure S13) with 7 of the redesigned PROTACs exhibiting lower EC_50_ values than that of MS4078 (Figure 8C).

**Figure 7.**
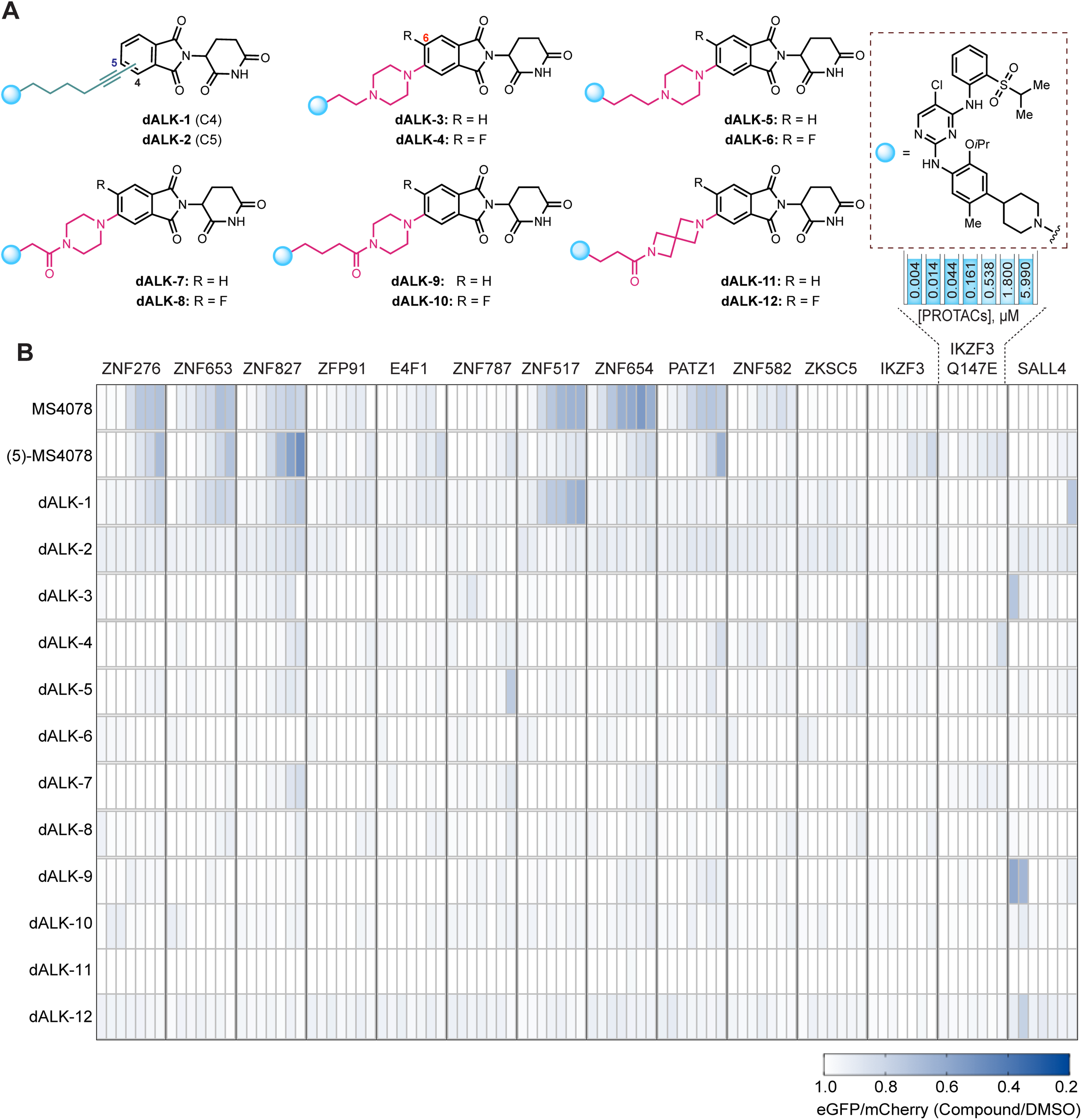
Re-engineering of ALK PROTACs based on the new design principles. **(A)** Structures of rationally redesigned ALK PROTACs to minimize off-target ZF degradation. **(B)** Degradation of validated and pomalidomide-sensitive ZF degrons inside cells by redesigned ALK PROTACs in a dosing range of 4.3 nM to 6 μM.

**Figure 8.**
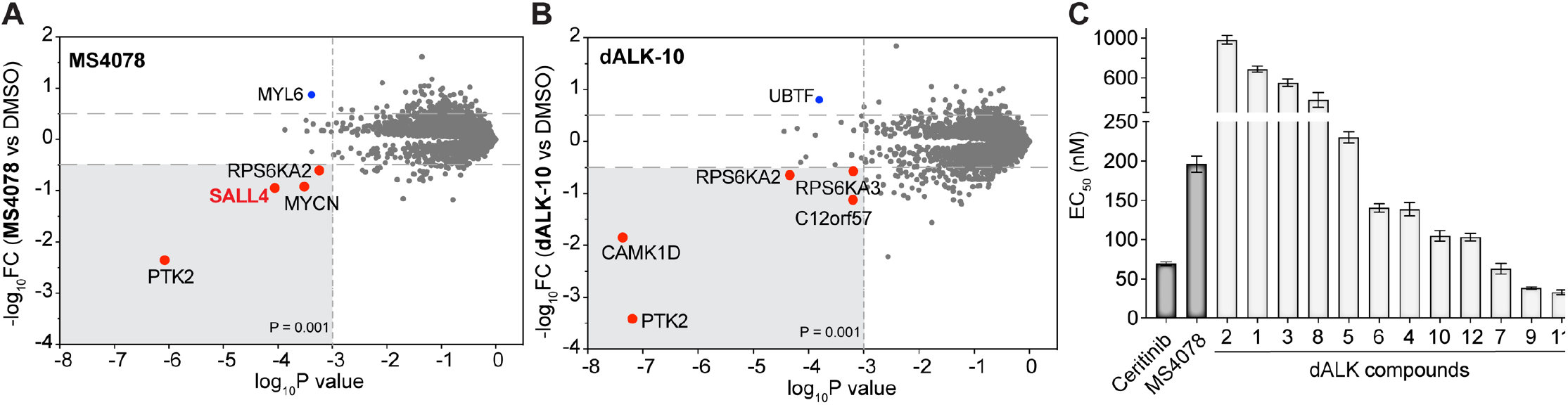
Validation of image-based degradation studies. **(A-B)** Global proteomic analysis of KELLY cells treated with 100 nM of MS4078 (A) or dALK-10 (B), respectively. **(C)** Determination of EC_50_ values (nM) of SU-DH-L1 cells upon treatment with redesigned ALK PROTACs.

## CONCLUSIONS

We have developed and validated an image-based high-throughput off-target profiling platform for the systematic evaluation of PROTAC-mediated off-target degradation of ZF proteins, which play crucial roles in biology and disease progression. We leveraged this platform to identify the rules for designing pomalidomide-based PROTACs that minimize off-target degradation of ZF proteins by designing and testing a library of pomalidomide analogs—this library allowed systematic exploration of impact of positional isomerism, non-covalent interactions, and steric and hydrophobic effects on off-target degradation. Here, the greatest reduction in off-target ZF degradation was achieved through modification of the exit vectors on the C5 position of the phthalimide ring via nucleophilic aromatic substitution (S_N_Ar). Guided by these new designed principles, we re-engineered a reported ALK PROTAC, MS4078 for enhanced potency and reduced off-target degradation. Our collection of pomalidomide analogs with minimal off-target ZF degradation can be widely adopted to generate safer and clinically relevant PROTACs. Notably, recently reported PROTACs in advanced clinical trials by Arvinas, Inc. (ARV-110 and ARV-471) and Foghorn therapeutics (FHD-609) employ piperazine and spirocyclic linkers (Figure S14) that have exhibit reduced off-targets. The new rules for pomalidomide-based PROTACs generated in this study can be readily applied to address the crucial need for PROTACs that do not indiscriminately degrade key ZF proteins. While our platform focusses on key off-targets (i.e., ZF finger proteins), future studies will expand our platform to other classes. Regardless, the findings from this study offer opportunities to develop new and safer PROTACs and improve on existent PROTACs with enhanced on-target potency.

## Supporting information

SI_Biology

SI_Chemistry

## ACKNOWLEDGEMENTS

We thank Patrick Byrne (Broad Institute) for assistance with automated imaging screening experiments. This work was supported by DARPA (N66001-17-2-4055) and NIH (R01 GM137606 and R01 GM132825 to A.C.; NIH CA214608 and CA218278 to E.S.F.; R01 EB031172, R01 EB027793, and R35 GM118062 to D.R.L. J.A.M.M. is a Ruth L. Kirchstein National Research Service Award Postdoctoral Fellow (F32 GM133088). D.R.L. and B.L.E. are investigators of the Howard Hughes Medical Institute.

