## Supplementary material for "Proteolysis Targeting Chimeras With Reduced Off-targets": SI_Biology

**Table of Contents**

**1. Supplementary table** 3
**Table S1.** Amino acid sequences of the zinc finger motifs cloned in cilantro 2 vector 3

**2. Supplementary figures** 4
**Figure S1.** Establishment of automated imaging assay for the degradation of ZFs 4
**Figure S2.** Structures of literature known PROTACs investigated in this study 5
**Figure S3.** Immunoblots and global proteomic analsysis for off-target degradation 6
**Figure S4.** Off-target ZF degradation assessed by mass spectrometry-based proteomics 7
**Figure S5.** Structures of miscellaneous IMiD analogs 8
**Figure S6.** Docking scores and topological polar surface area of C4 and C5 pomalidomide analogs 9
**Figure S7.** Degradation of validated pomalidomide-sensitive ZF degrons induced by the pomalidomide analogs 11
**Figure S8.** Structures of IMiD analogs with the least degradation score (close to 0) 13
**Figure S9.** NanoBRET based ternary complex analysis of all the IMiD analogs and PROTACs reported in this study 14
**Figure S10.** Global Proteomic analysis of selected pomalidomide analogs (1, 39, 37, 56, 38, 32, 36, and 61) in KELLY cells. 15
**Figure S11.** Schemes for **dALK-1** to **12** and **(5)-MS4078** synthesis 16
**Figure S12.** Global Proteomic analysis of selected PROTACs (MS4078 and dALK-10) in SU-DHL-1 cells 17
**Figure S13.** Viability dose curves for the redesigned ALK PROTACs in SU-DHL-1 cells 18
**Figure S14.** Structures of the PROTACs currently in clinical trials 19

**3. Methods** 20

**4. Nucleotide sequence of the luciferase plasmid generated in this study** 25

**5. References** 27

### 1. Supplementary table

**Table S1.** Amino acid sequences of the zinc finger motifs cloned in cilantro 2 vector

| Protein (residue numbers) | Function of the protein | Amino acid sequences |
|---|---|---|
| **ZNF276** (524-546 aa) | Transcriptional regulation | LQCEVCGFQCRQRASLKYHMTKH |
| **ZNF653** (556-578 aa) | Transcriptional repressor | LQCEICGYQCRQRASLNWHMKKH |
| **ZNF827** (374-396 aa) | Transcriptional regulation | FQCPICGLVIKRKSYWKRHMVIH |
| **ZFP91** (400-422 aa) | Atypical E3 ubiquitin-protein ligase, has key role in cell proliferation, anti-apoptosis | LQCEICGFTCRQKASLNWHMKKH |
| **E4F1** (220-242 aa) | Transcriptional regulation | HECKLCGASFRTKGSLIRHHRRH |
| **ZNF787** (178-200 aa) | Transcriptional regulation | FVCPRCGRGFSQPKSLARHLRLH |
| **ZNF517** (452-474 aa) | Transcriptional regulation | YRCRACGRACSRLSTLIQHQKVH |
| **ZNF654** (25-47 aa) | Transcriptional regulation | FACVICGRKFRNRGLMQKHLKNH |
| **PATZ1** (383-405 aa) | chromatin modeling and transcription regulation, has key functions in embryogenesis, senescence, T-cell development, neurogenesis | YSCPVCGLRFKRKDRMSYHVRSH |
| **ZNF582** (395-417 aa) | Transcriptional regulation | YQCKVCGRAFKRVSHLTVHYRIH |
| **ZKSC5** (430-452 aa) | Transcriptional regulation | YGCNECGKNFGRHSHLIEHLKRH |
| **IKZF3** (146-168 aa) | Transcription factor, plays key role in regulation of B-cell differentiation, proliferation and maturation | FQCNQCGASFTQKGNLLRHIKLH |
| **IKZF3 Q147E** (146-168 aa) | Inactive mutant of IKZF3, serves as negative control in image-based assay | FECNQCGASFTQKGNLLRHIKLH |
| **SALL4** (410-432) | Transcription factor, plays key role in the maintenance and self-renewal of embryonic, hematopoietic stem cells | FVCSVCGHRFTTKGNLKVHFHRH |

### 2. Supplementary figures

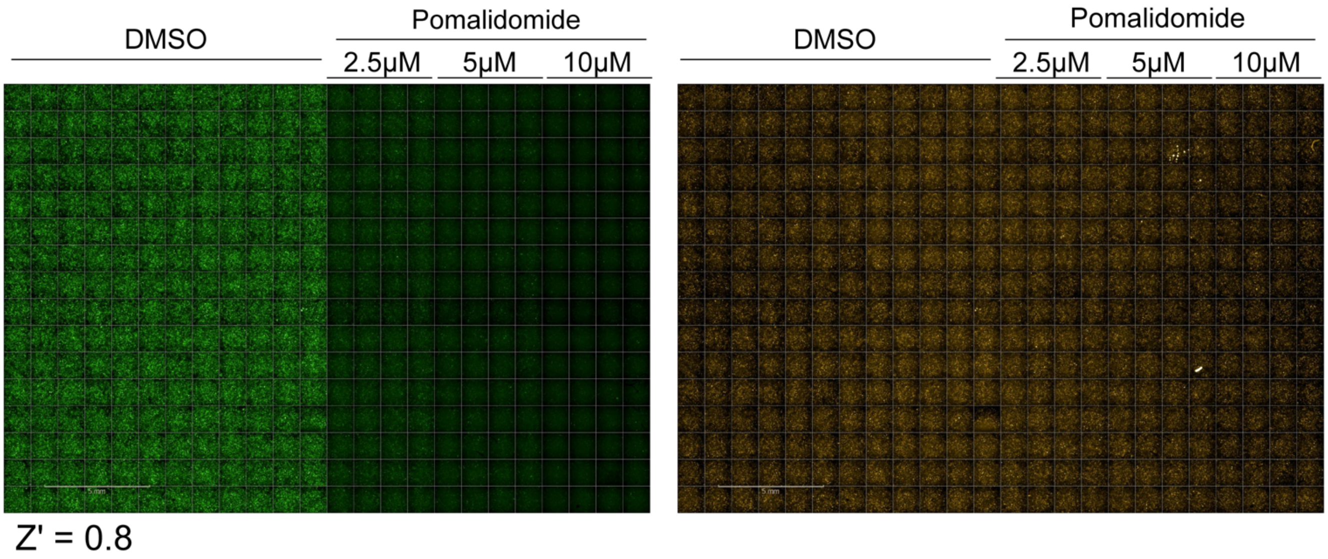


**Figure S1. Establishment of automated imaging assay for the degradation of ZFs.** Representative readouts of the imaging assay in 384-well plates, demonstrating robust detection of ZF-tagged eGFP degradation as induced by pomalidomide. Shown are images of U2OS cells with stable expression of pomalidomide-sensitive ZFP91 degron reporter. Z′ value was calculated using an in-built module in the Harmony software (See materials and methods).

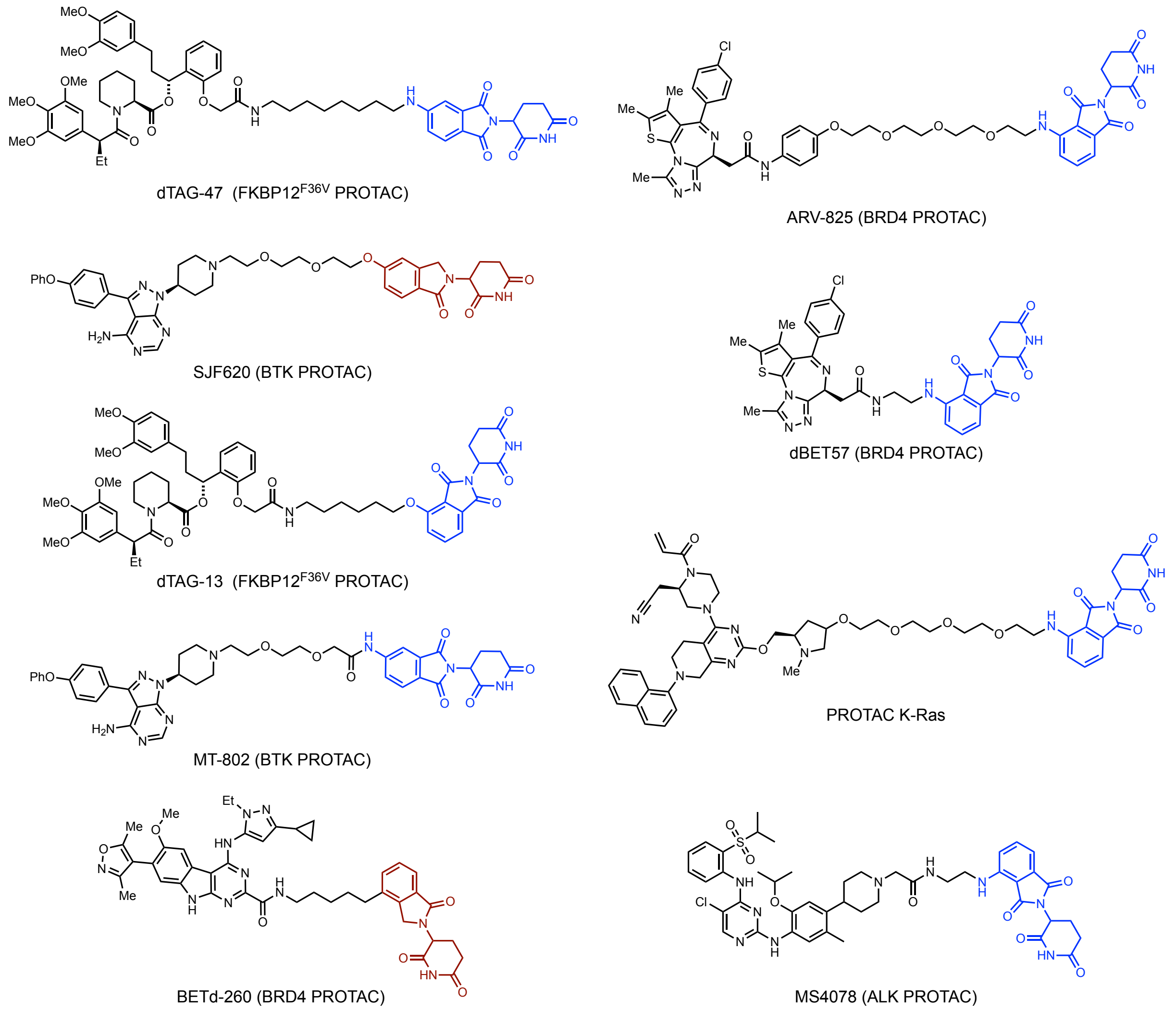


**Figure S2.** Structures of literature known PROTACs investigated in this study.

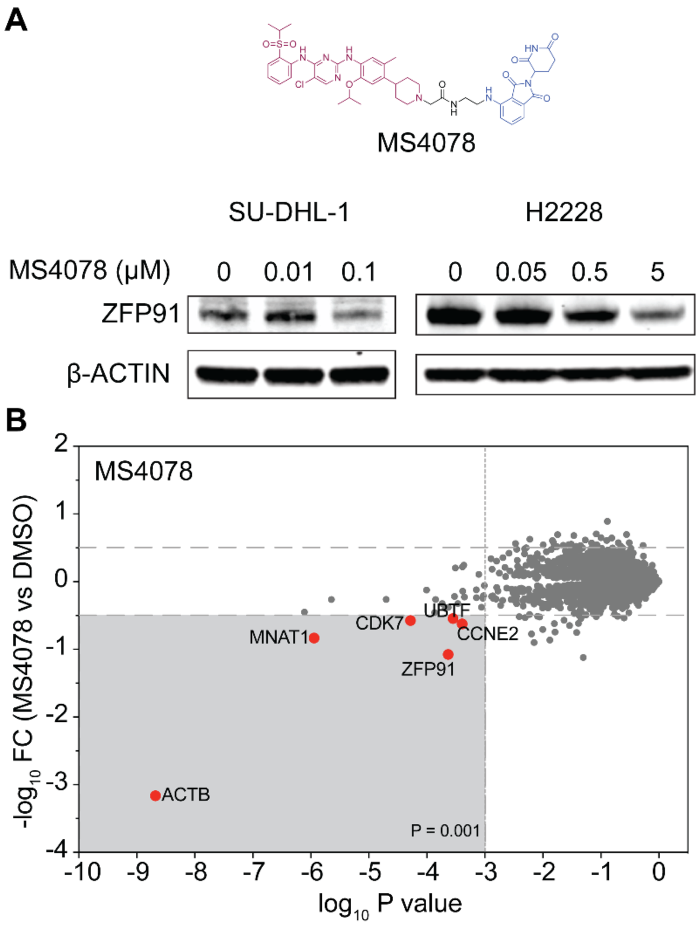


**Figure S3. Immunoblots and global proteomic analsysis for off-target degradation. A)** Immunoblots quantifying off-target degradation of endogenous ZF proteins ZFP91 by MS4078 (ALK PROTAC) in a dose-dependent manner across cell lines SU-DHL-1 and H2228. **B)** Label-free proteomic analysis of MS4078 in MOLT4 cells.

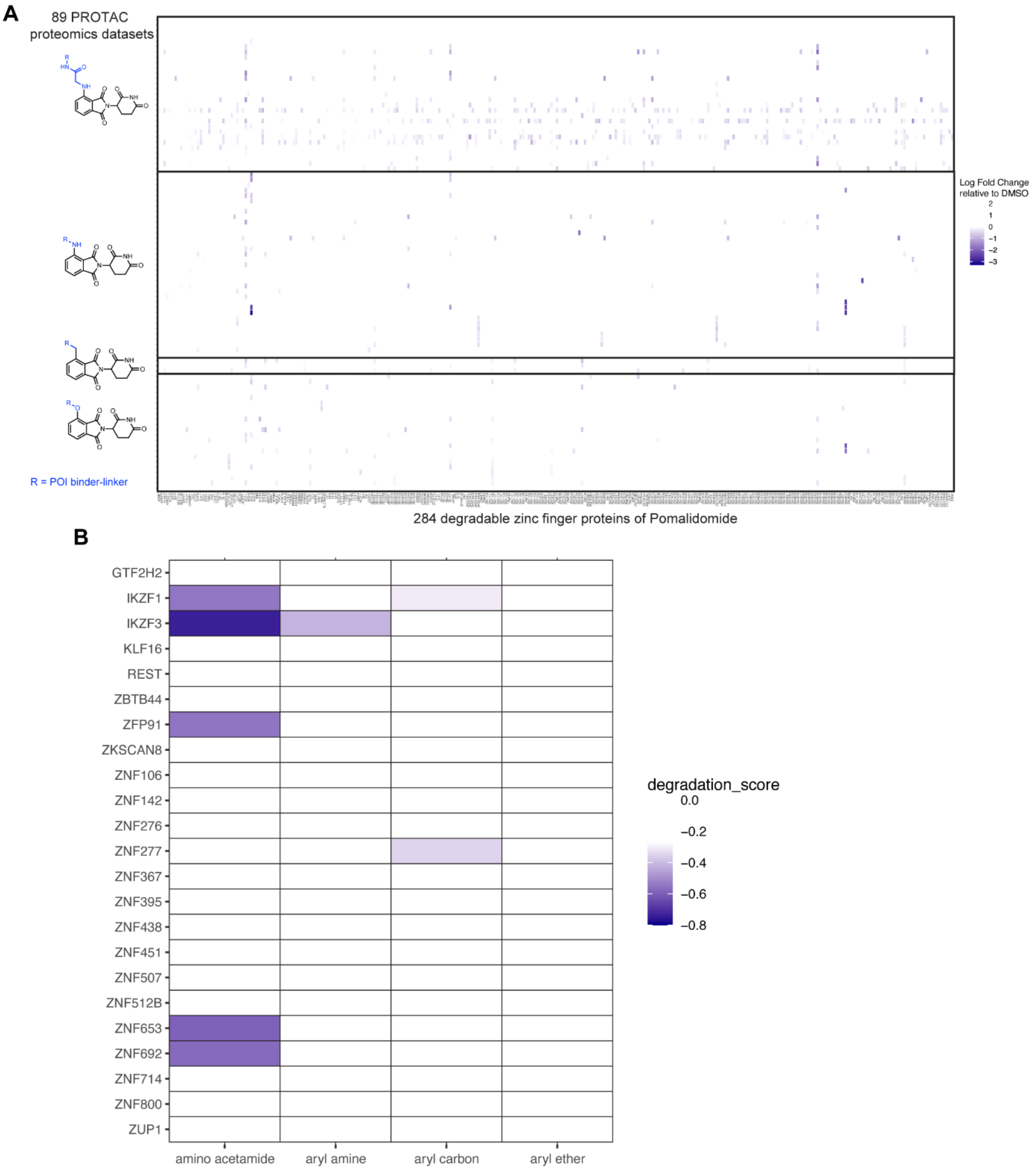


**Figure S4. Off-target ZF degradation assessed by mass spectrometry-based proteomics.** Identification of the same group of exit vectors on pomalidomide with minimal off-target ZF degradation assessed by mass spectrometry-based proteomics for entire data set (A) and top 10 most degraded proteins (B). Relative abundance of endogenous ZF proteins in cells treated with pomalidomide-based PROTACs as arranged based on pomalidomide's exit vector groups. Data were extracted from proteomics datasets published in Donovan et al., 2020.[1]

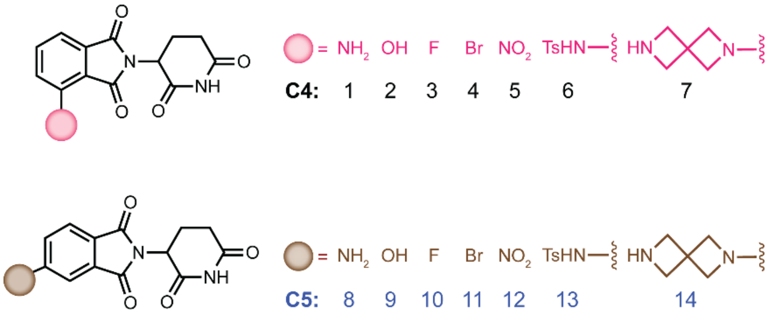


**Figure S5.** Structures of miscellaneous IMiD analogs.

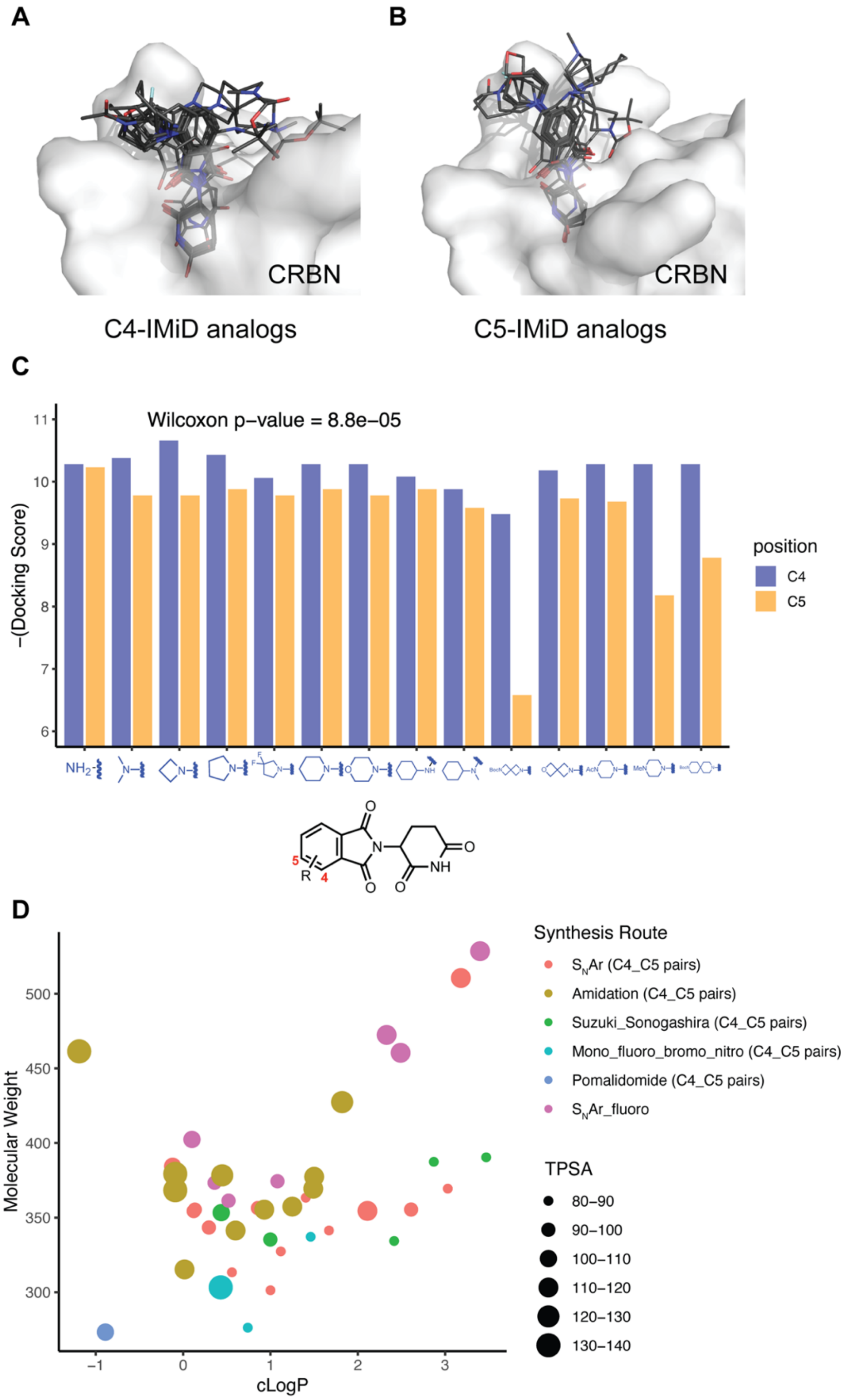


**Figure S6. Docking scores and topological polar surface area of C4 and C5 pomalidomide analogs. A-C)** Structural docking of pomalidomide analogs with C4 (A) and C5 (B) modifications on the phthalimide ring. Docking score (C) of each pair of modifications on C4 and C5 (paired Wilcoxon test, p = 8.8 x $e^{-05}$). **(D)** Distribution of the physicochemical properties of the pomalidomide analog library. The topological polar surface area (TPSA) of each molecule is indicated by color and size (see legend). Note that each dot in the scatter plot represents a pair of compounds with the same modification on C4 and C5 positions, except the $S_NAr$ fluoro group. Synthetic routes are represented by different shapes shown in the legend).

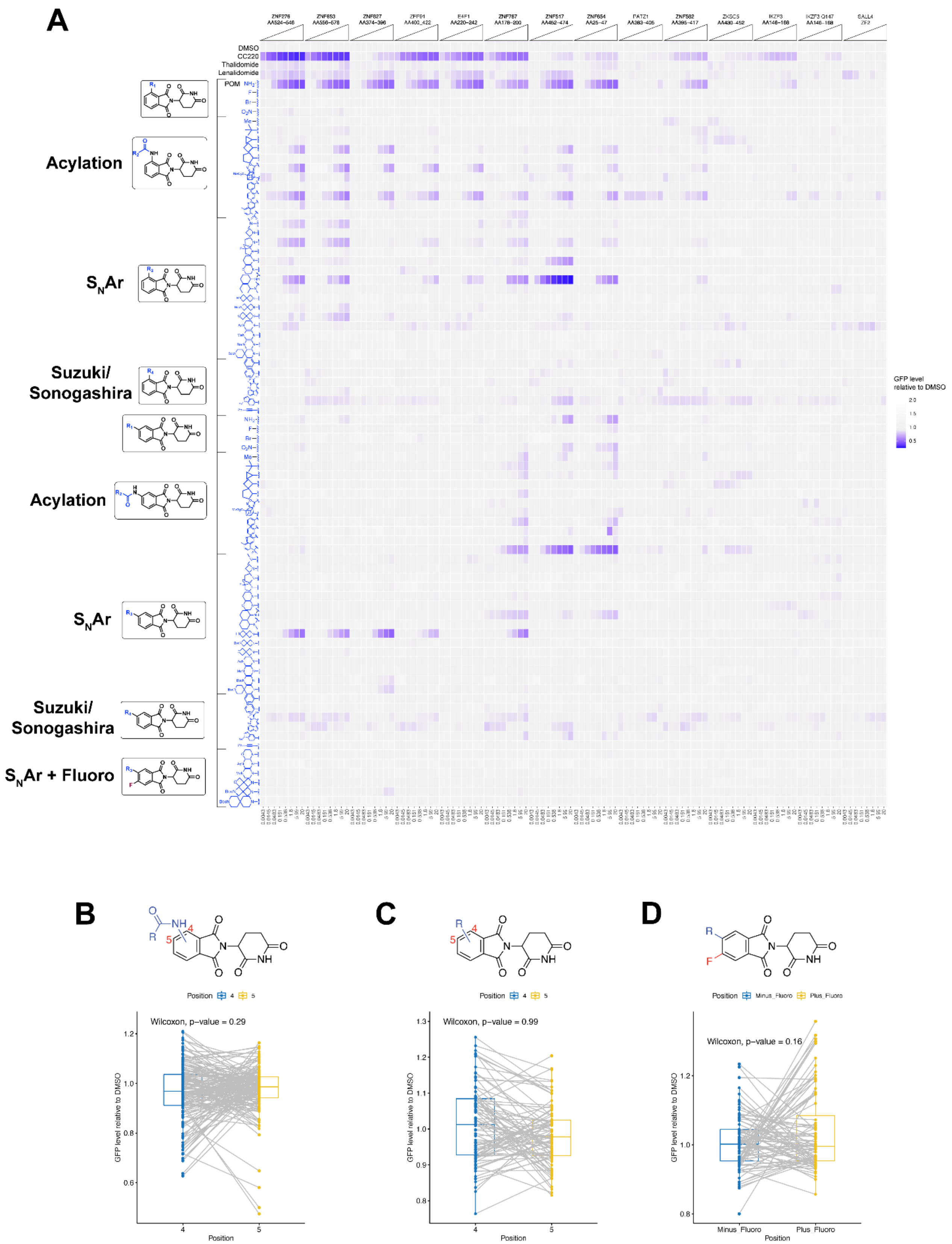


**Figure S7. Degradation of validated pomalidomide-sensitive ZF degrons induced by the pomalidomide analogs. (A)** Normalized eGFP intensity in 14 ZF reporter cell lines treated with different doses of 81 pomalidomide analogs ranging from 4.3 nM to 20 μM. Each block of 4.3 nM to 20 μM doses on the x axis represents one ZF reporter cell line. (**B-D)** Box-and-Whisker plots with statistical analysis for pomalidomide analogs arranged in pairs of C4 and C5 modifications such as acylation (B), Suzuki/Sonogashira coupling (C) and effect of -F group (D) on the phthalimide ring. Data are shown for cells treated with 5 μM of each compound (for acylated and Suzuki/Sonogashira couplings) and 20 μM data of each compound for deciphering the -F group effect.

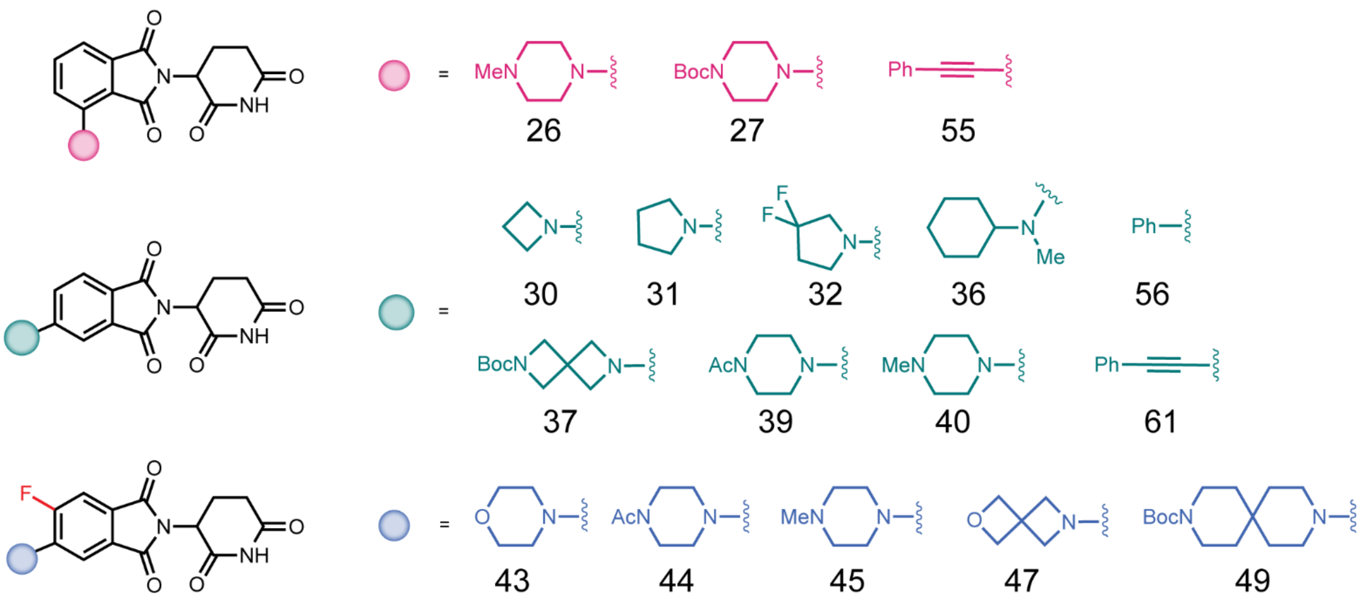


**Figure S8.** Structures of IMiD analogs with the least degradation score (close to 0).

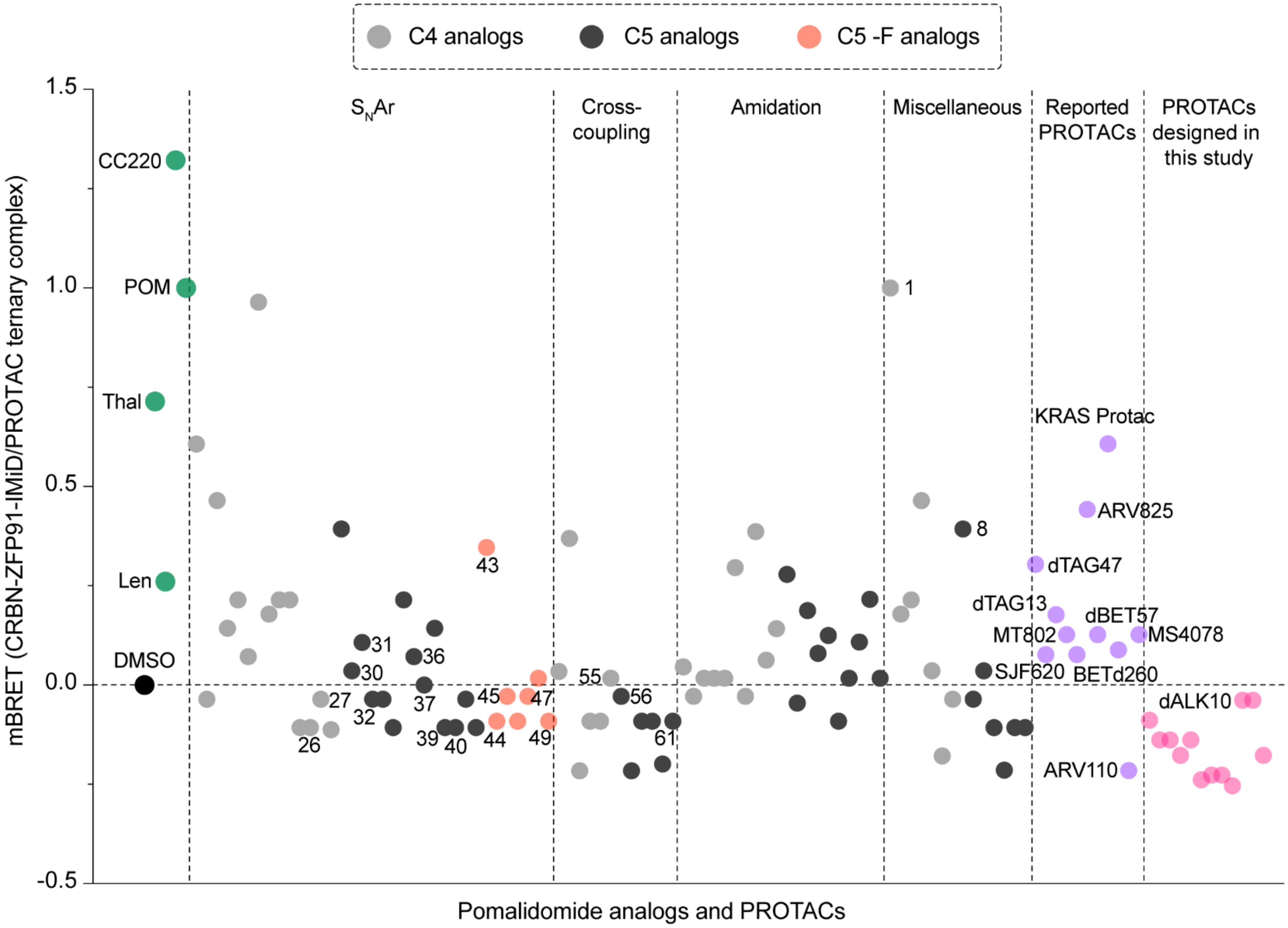


**Figure S9.** NanoBRET based ternary complex analysis of all the IMiD analogs and PROTACs reported in this study.

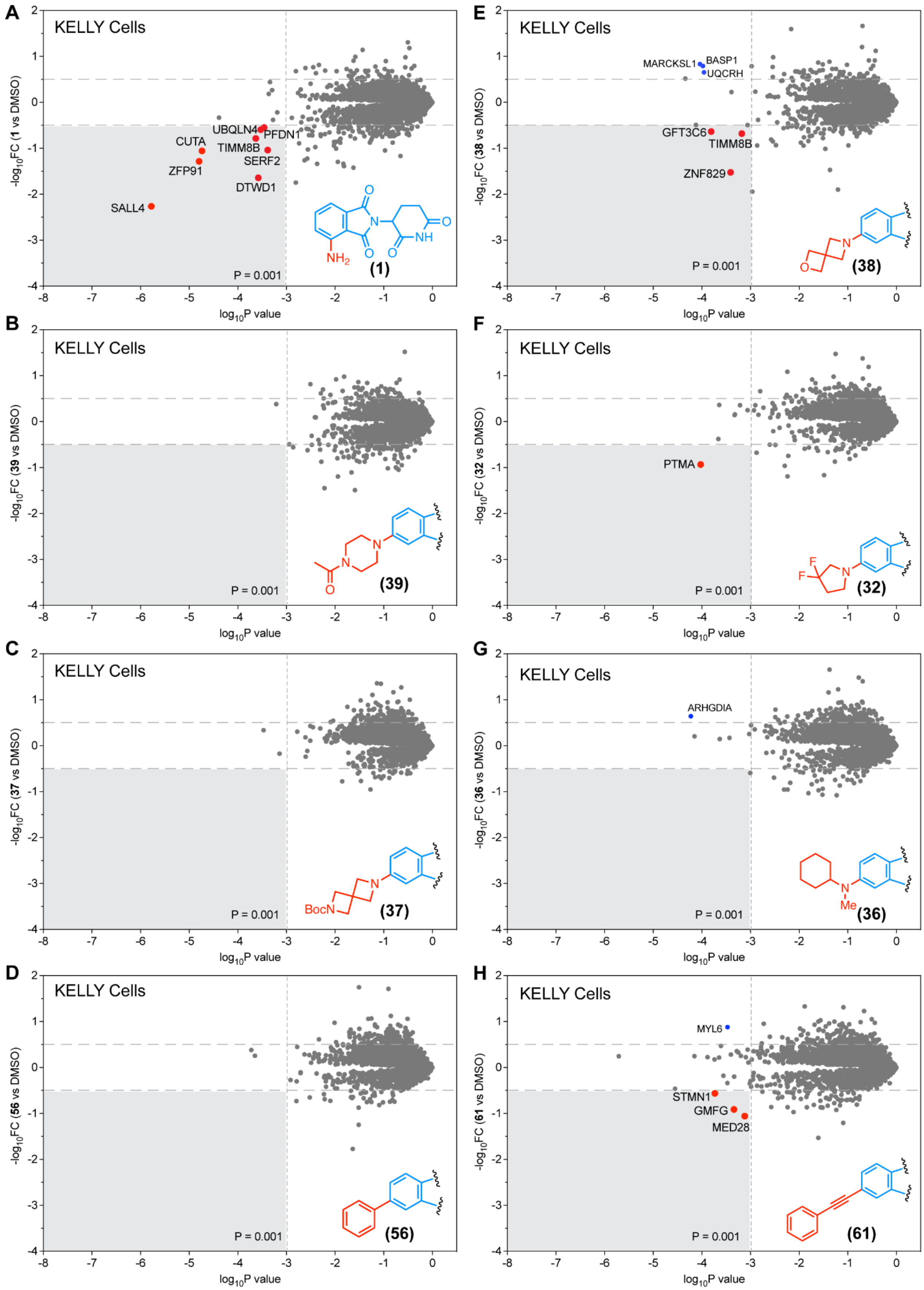


**Figure S10.** Global Proteomic analysis of selected pomalidomide analogs (1, 39, 37, 56, 38, 32, 36, and 61) in KELLY cells.

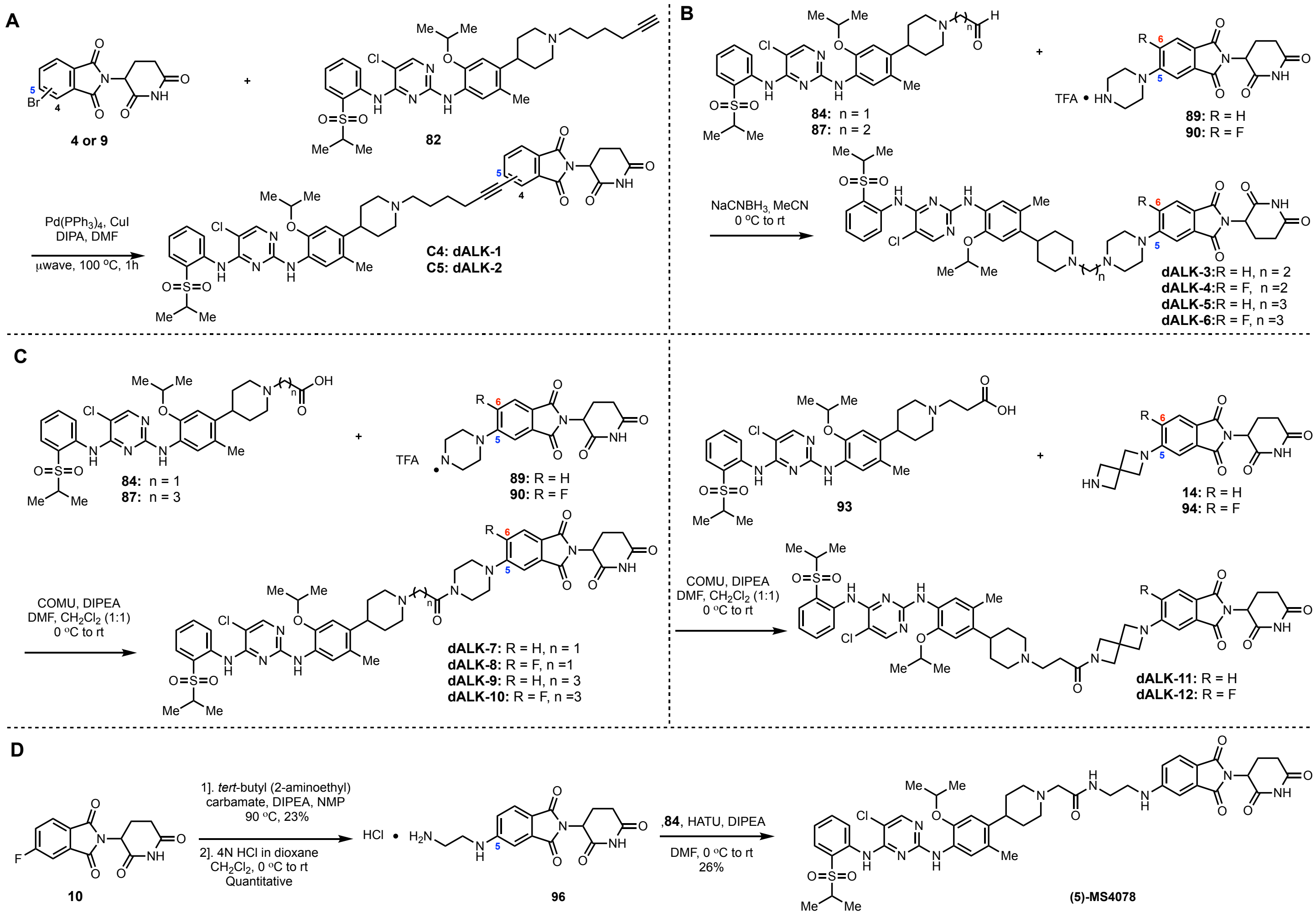


**Figure S11.** Schemes for **dALK-1** to **12** and **(5)-MS4078** synthesis.

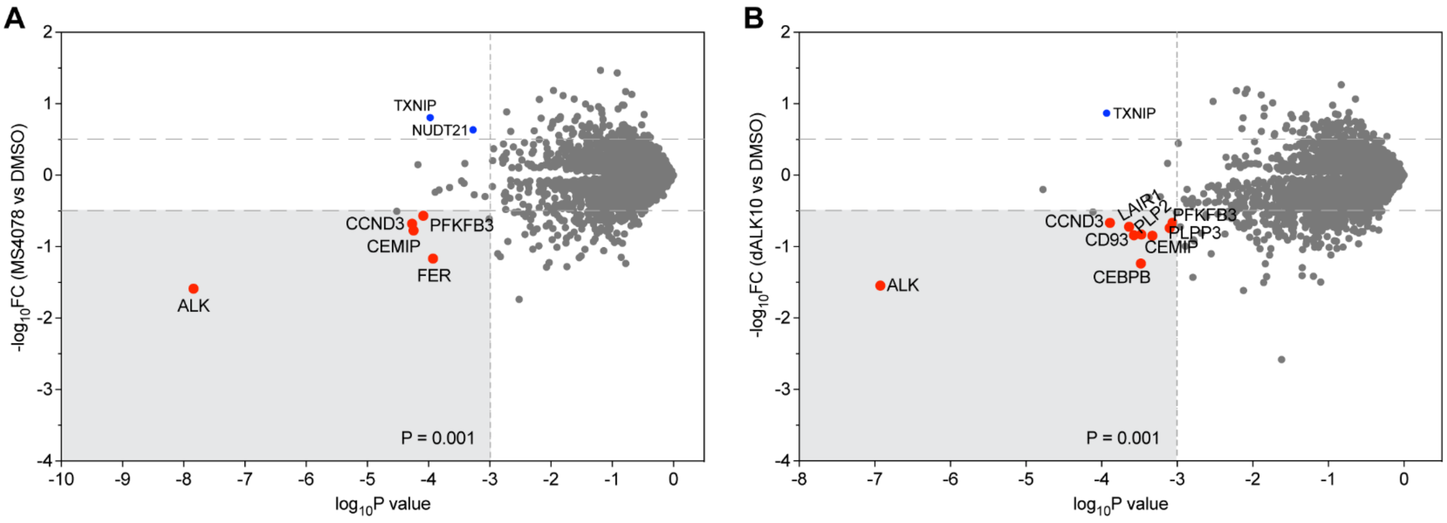


**Figure S12.** Global Proteomic analysis of selected PROTACs (MS4078 and dALK-10) in SU-DHL-1 cells.

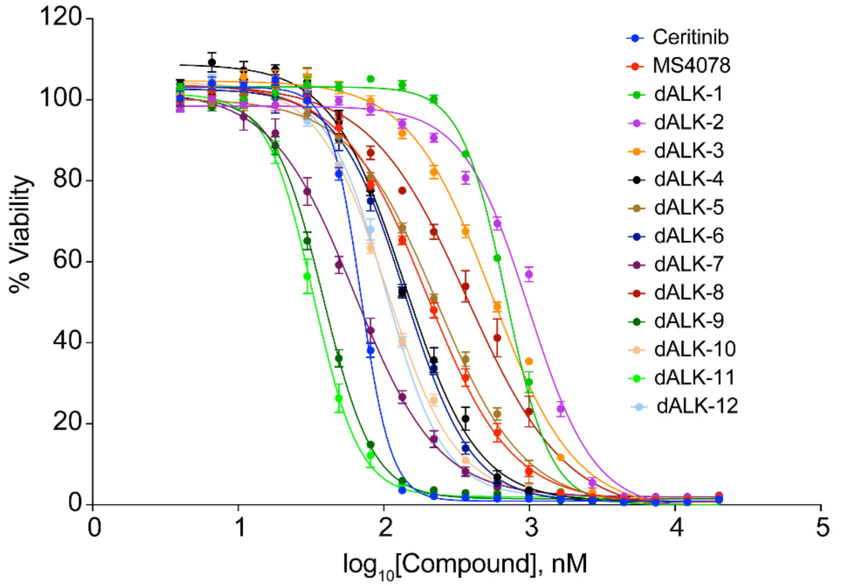


**Figure S13.** Viability dose curves for the redesigned ALK PROTACs in SU-DHL-1 cells.

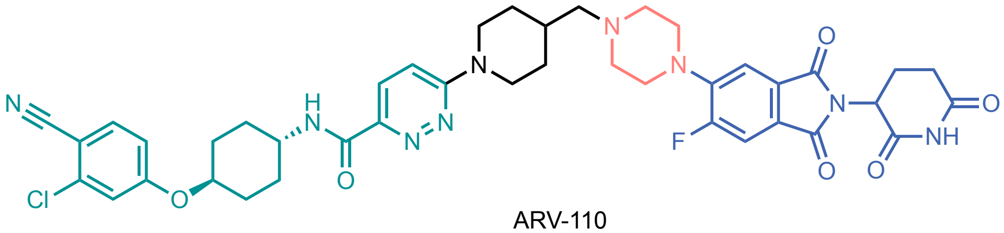

ARV-110

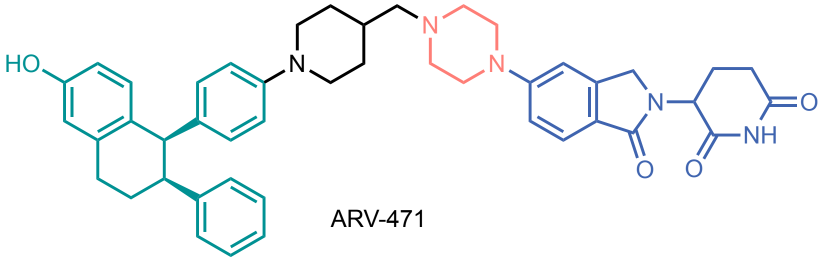

ARV-471

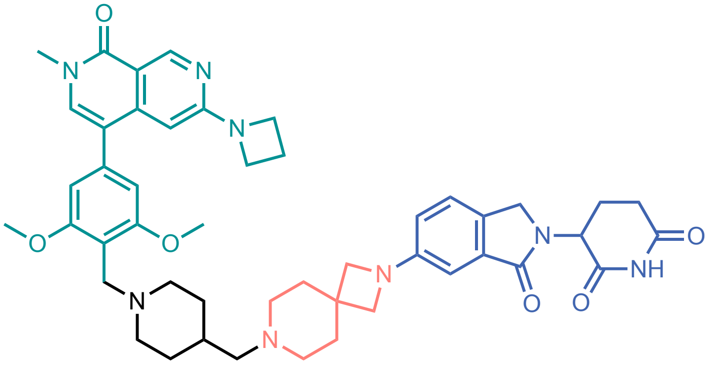

FHD-609

**Figure S14.** Structures of the PROTACs currently in clinical trials.

### 3. Methods

#### Cell lines

U2OS (ATCC, HTB-96) and 293T (ATCC, CRL-3216) cells were cultured in Dulbecco's modified Eagle's medium (DMEM) (Thermo Fisher Scientific, 12430062), 10% (v/v) fetal bovine serum (FBS) (Thermo Fisher Scientific, 16140071), and 100 U/ml Antibiotic-Antimycotic (ThermoFisher Scientific, 15240062). U2OS cells stably expressing the ZF constructures were maintained in full-DMEM media with 1 µg/mL puromycin (Thermo Fisher Scientific, A1113803). MM1.S (ATCC, CRL-2974), Jurkat Clone E6-1 (ATCC, TIB-152), SU-DHL-1 (ATCC, CRL-2955), and H2228 cells (ATCC, CRL-5935) were cultured in RPMI 1640 medium (Thermo Fisher Scientific, 11875119) with 10% (v/v) FBS, and 100 U/ml Antibiotic-Antimycotic.

#### Plasmids

The 14 lentiviral ZF plasmids were generated using the Cilantro 2 degradation reporter vector (Addgene, 74450) as previously described.[2] The luciferase version (pSVBV1, sequence map provided in section 4) of Cilantro 2 degradation reported vector generated by swapping the eGFP with nanoluciferase (NLuc) and mcherry with firefly luciferase (FLuc) followed by cloning the ZFP91 sequence in luciferase vector. The amino sequences of the 14 ZFs were mentioned in supplementary table S1.

#### Lentivirus production and transduction

Viral packaging plasmids psPAX2 (Addgene, 12260) and pMD2.G (Addgene, 12259) together with lentiviral ZF degron plasmids were transfected to 293T cells in a 2:1:3 ratio using TransIT®-LT1 transfection reagent (MirusBio, MIR2300) following the manufacturer's guidelines. Lentiviruses were collected 48 and 72 h after transfection and filtered with 0.45-µm Millex®-HP PES filters (Millipore, SLHPM33RS). For transduction in U2OS cells, the viral supernatant was mixed with U2OS culture media in a 1:1 ratio with 10 µg/ml Polybrene transfection reagent (Millipore Sigma, TR-1003-G), then selected with 2 µg/ml puromycin after at least 24 h of transduction. Finally, antibiotic-selected cells were maintained in 1 µg/mL puromycin for routine culture. We chose U2OS for the primary screening as they are large, flat, and adherent, making them easy to image. Furthermore, several zinc finger proteins have low expression in U2OS cells but are abundantly expressed in the myeloid cells, blood cancer cell lines, neuronal cell lines, and embryonic stem cells, which were chosen for subsequent validation.

#### Automated imaging screening and analysis

CellCarrier-384 Ultra Microplates (PerkinElmer, 6057302) were printed with 10 mM of stock compounds in varying volumes using Tecan D300e Digital Dispenser (Tecan) and HP T8+ dispensehead cassettes (F0L59A). U2OS cells stably expressing the ZF degron reporters were seeded in 30 µL of 3000 cells per well in compound preprinted 384 plates using Thermofisher combi instrument. After 24 hours of incubation, the 384-well plates were washed once with PBS then fixed in 4% paraformaldehyde and stained by HCS NuclearMask Blue stain (ThermoFisher Scientific, H10325) using ThermoFisher combi instrument. Imaging of eGFP and mCherry was performed for each plate using an Opera Phenix imaging system followed by analysis with Harmony Software v4.9 (PerkinElmer). In-built module was used to calculate the Z′ value using the following formula.

$$Z' = 1 - \frac{3(\boldsymbol{\sigma}pc + \boldsymbol{\sigma}nc)}{|\ \mu pc + \boldsymbol{\mu}nc\ |}$$

**σ**pc = standard deviation of positive controls

**σ**nc = standard deviation of negative controls

**µ**pc = Mean of positive controls

**µ**nc = Mean of negative controls

The fluorescence intensity of eGFP was normalized to that of mCherry for every cell. The mean normalized eGFP intensity of all cells in each well was then normalized by that in DMSO-treated wells to determine the GFP level relative to DMSO for each compound across doses. The GFP level relative to DMSO was used to generate a heatmap using R v4.0.2 or Graphpad Prism 9. Compounds that cause minimal ZF degradation have values close to 1 (i.e., DMSO value), whereas compounds that cause extensive ZF degradation have values close to 0. To compute the degradation score for each compound, all DMSO-normalized GFP values greater than 0.9 were excluded to ensure that only ZF degradation values contribute to the score but not those that were associated with increases in GFP or caused minimal to no change in GFP degradation. The remaining DMSO-normalized GFP values were subtracted from 1 to derive the fraction of GFP degradation caused by the compounds. The degradation score was then calculated by taking the sum of the GFP degradation fractions across all doses for each compound.

**Proteomics analysis of PROTAC degradation of endogenous ZF proteins**

The relative abundance of 811 unique pomalidomide-sensitive ZF proteins[2] were extracted from 124 publicly available pomalidomide-based PROTAC proteomics datasets reported previously.[1] Here, 644 out of 811 ZF proteins were detectable in at least one PROTAC proteomics dataset and are shown in Figure S4. To compute the ZF degradation score for each pomalidomide-based PROTAC dataset, all relative abundance values that are greater than 0 were excluded to ensure that only degradation values contribute to the score but not those that are associated with increases in protein abundance. The degradation score was then calculated by taking the sum of negative abundance values, representing ZF proteins that reduced in abundance when the cells were treated with PROTACs.

**Immunoblot analysis**

Lysates of cells treated with different compounds were collected with ice-cold M-PER$^{TM}$ Mammalian Protein Extraction Reagent (ThermoFisher Scientific, 78501) with freshly added phosphatase (Sigma-Aldrich, 04906837001) and protease (Sigma-Aldrich, 04693124001) inhibitors following the manufacturer's instructions.

Next, 20–50 µg of proteins were fractionated with NuPAGE™ 4–12% Bis-Tris gels (ThermoFisher Scientific, NP0335BOX) then transferred onto nitrocellulose membranes using iBlot™ Transfer Stacks (ThermoFisher Scientific, IB23002) following the manufacturer's instructions. Membranes were then stained with primary antibodies in 1:1000 dilutions and secondary fluorescent antibodies in 1:3000 dilutions using iBind™ Flex Fluorescent Detection Solution Kit (ThermoFisher Scientific, SLF2019) following the manufacturer's instructions.

IKZF3 and ZFP91 are one of the common and important off-targets of pomalidomide-based PROTACs. Furthermore, for these targets, we could easily access detection reagents and cell lines expressing them at high level–both choices allowed us to use western blotting for cross-validation. Primary antibodies used in this study include ZFP91 (Bethyl Laboratories, A303-245A), IKZF3/Aiolos (D1C1E) (Cell Signaling, 15103S), CRBN (D8H3S) (Cell Signaling, 71810S), ALK (D5F3) XP (Cell Signaling, 3633S), phospho-ALK (Tyr1507) (D6F1V) (Cell Signaling, 14678S), β-actin (8H10D10) Mouse mAb (Cell Signaling, 3700S). Fluorescent secondary antibodies used in this study include IRDye 680RD goat anti-mouse IgG (LI-COR Biosciences, 926-68070) and IRDye 800CW goat anti-rabbit IgG (LI-COR Biosciences, 926-32211). Immunoblot blot detection was performed using an Odyssey CLx Imaging System (LI-COR Biosciences). Quantification of the relative area and density values of western blot bands was carried out using ImageJ v2.1.0 following the ImageJ User Guide for gel analysis (https://imagej.nih.gov/ij/docs/guide/). Quantified values were normalized by values for loading controls such as β-actin. For phospho NPM-ALK, quantified values were normalized by the values for total ALK. All the blots were repeated at least twice.

**Global proteomic analysis of IMiDs and PROTACs**

Sample preparation LFQ quantitative mass spectrometry. MOLT4 or Kelly or SU-DHL-1 cells were treated with DMSO or IMiD analogues (1 µM) or PROTACs (0.1 µM) for 5 h. Cells were harvested by centrifugation and washed with phosphate-buffered saline (PBS) before snap freezing in liquid nitrogen. Cells were lysed by the addition of lysis buffer (8 M Urea, 50 mM NaCl, 50 mM 4-(2-hydroxyethyl)-1-piperazineethanesulfonic acid (EPPS) pH 8.5, Protease and Phosphatase inhibitors) and homogenization by bead beating (BioSpec) for three repeats of 30 seconds at 2400. Bradford assay was used to determine the final protein concentration in the clarified cell lysate. 50 µg of protein for each sample was reduced, alkylated, and precipitated using methanol/chloroform as previously described[3], and the resulting washed precipitated protein was allowed to air dry. Precipitated protein was resuspended in 4 M Urea, 50 mM HEPES pH 7.4, followed by dilution to 1 M urea with the addition of 200 mM EPPS, pH 8. Proteins were first digested with LysC (1:50; enzyme:protein) for 12 h at RT. The LysC digestion was diluted to 0.5 M Urea with 200 mM EPPS pH 8 followed by digestion with trypsin (1:50; enzyme:protein) for 6 h at 37 °C. Sample digests were acidified with formic acid to a pH of 2-3 prior to desalting using C18 solid phase extraction plates (SOLA, Thermo Fisher Scientific). Desalted peptides were dried in a vacuum-centrifuged and reconstituted in 0.1% formic acid for LC-MS analysis. Data were collected using a TimsTOF Pro2 (Bruker Daltonics, Bremen, Germany) coupled to a nanoElute LC pump (Bruker Daltonics, Bremen, Germany) via a CaptiveSpray nano-electrospray source. Peptides were separated on a reversed-phase

$C_{18}$ column (25 cm x 75 µm ID, 1.6 µM, IonOpticks, Australia) containing an integrated captive spray emitter. Peptides were separated using a 50 min gradient of 2 - 30% buffer B (acetonitrile in 0.1% formic acid) with a flow rate of 250 nL/min and column temperature maintained at 50 ºC.

DDA was performed in Parallel Accumulation-Serial Fragmentation (PASEF) mode to determine effective ion mobility windows for downstream diaPASEF data collection [4]. The ddaPASEF parameters included: 100% duty cycle using accumulation and ramp times of 50 ms each, 1 TIMS-MS scan and 10 PASEF ramps per acquisition cycle. The TIMS-MS survey scan was acquired between 100 – 1700 *m/z* and 1/k0 of 0.7 - 1.3 V.s/cm$^2$. Precursors with 1 – 5 charges were selected and those that reached an intensity threshold of 20,000 arbitrary units were actively excluded for 0.4 min. The quadrupole isolation width was set to 2 *m/z* for *m/z* <700 and 3 *m/z* for *m/z* >800, with the *m/z* between 700-800 *m/z* being interpolated linearly. The TIMS elution voltages were calibrated linearly with three points (Agilent ESI-L Tuning Mix Ions; 622, 922, 1,222 *m/z*) to determine the reduced ion mobility coefficients ($1/K_0$). To perform diaPASEF, the precursor distribution in the DDA *m/z*-ion mobility plane was used to design an acquisition scheme for DIA data collection which included two windows in each 50 ms diaPASEF scan. Data was acquired using sixteen of these 25 Da precursor double window scans (creating 32 windows) which covered the diagonal scan line for doubly and triply charged precursors, with singly charged precursors able to be excluded by their position in the m/z-ion mobility plane. These precursor isolation windows were defined between 400 - 1200 *m/z* and 1/k0 of 0.7 - 1.3 V.s/cm$^2$.

_LC-MS data analysis._ The diaPASEF raw file processing and controlling peptide and protein level false discovery rates, assembling proteins from peptides, and protein quantification from peptides was performed by analysis in DIA-NN 1.8 using library or library free methods. For library analysis: targeted cell line specific spectral libraries were generated by searching offline fractionated DDApasef data against a Swissprot human database (January 2021) For library free methods: an in silico digestion of a given protein sequence database is performed alongside deep learning-based predictions to extract the DIA precursor data into a collection of MS2 spectra. The Search results are then sued to generate a spectral library which is then employed for the targeted analysis of the DIA data searched against a Swissprot human database (January 2021).[5] Database search criteria largely followed the default settings for DIA including: tryptic with two missed cleavages, carbomidomethylation of cysteine, and oxidation of methionine and precursor Q-value (FDR) cut-off of 0.01. Precursor quantification strategy was set to Robust LC (high accuracy) with RT-dependent cross run normalization. Proteins with missing values in any of the treatments and with poor quality data were excluded from further analysis (summed abundance across channels of <100 and mean number of precursors used for quantification <2). Protein abundances were scaled using in-house scripts in the R framework [6] and statistical analysis was carried out using the limma package within the R framework [7].

**NanoBRET assays:**

**a) CRBN binding assay**

U2OS cells were transfected with a combination of NanoLuc-CRBN fusion vector (1 µg/ 100,000 cells) and DDB1 expression vector (4 µg/ 100,000 cells) encoded plasmid (Promega, N2910) using Lipofectamine 3000 reagent (Thermo Fischer Scientific, L3000015). After 24 hours of transfection, a 384-well white microplate (Corning, 3765) is prepared with doses at 10x concentration printed using 10 mM of stock compounds in varying volumes and Tecan D300e Digital Dispenser (Tecan). Then the transfected cells were washed with PBS, trypsinized, and plated in a 384-well white microplate (Corning, 3765) at 3000 cells in 34 µL Opti-MEM I reduced serum medium/well density. Then complete 20X NanoBRET™ Tracer Reagent (2 µL) and dosed of 10x IMiD compounds (4 µL) were dispensed in each well. Then the plates were incubated for 1 hour at 37 ˚C, 5% CO2. After incubation plates were brought to room temperature and incubated with 3X complete substrate plus inhibitor solution (20 µL) for 3 minutes at room temperature before taking the BRET signal by an EnVision multilabel plate reader with EnVision manager 1.13 (PerkinElmer).

**b) Ternary complex assay**

U2OS cells stably expressing ZFP91-Nanoluciferase were transfected (2 µg/ 100,000 cells) with HaloTag-CRBN encoded plasmid (Promega, N2691) using Lipofectamine 3000 reagent (Thermo Fischer Scientific, L3000015). 14 h following transfection, cells were trypsinized and plated at 3000 cells in 30 µL Opti-MEM I reduced serum medium (no phenol red + 4% FBS) per well in a 384-well white microplate (Corning, 3765). Then the cells were treated with 10 µL HaloTag NanoBRET 618 ligand (Promega, G9801) in Opti-MEM I reduced serum medium (no phenol red + 4% FBS) to a final concentration of 100 nM for 18 hours. No ligand control wells were added 10 µL above mentioned Opti-MEM I reduced serum media. Post HaloTag reaction, cells were incubated for 15 min with 10 µL of IMiD compound or PROTAC solution in above mentioned Opti-MEM I reduced serum media to the final concentration of 1 µM. Followed by compound incubation, 10 µL of nanoluciferase substrate, furimazine (Aobious, AOB36539) in Opti-MEM I reduced serum media was added at a final concentration of 10 µM and incubated for 5 minutes before taking the BRET signal by an EnVision multilabel plate reader with EnVision manager 1.13 (PerkinElmer).

NanoBRET values for the CRBN binding or ternary complex formation assay were calculated as follows:

1) Raw NanoBRET Ratio (mBU) = ($618nm_{Em}$/$460nm_{Em}$) × 1000
2) Mean corrected NanoBRET ratio mBU = Mean mBU experimental − Mean mBU no-ligand control

Note that at each liquid handling step, the plates were shaked on a Thermo Fisher combi instrument followed by centrifugation at 100 xg for 1 minute.

**Cell Viability Assay**

384 well white microplates (Corning, 8867BC) were printed with 10 mM of stock compounds in varying volumes using Tecan D300e Digital Dispenser (Tecan) and HP T8+ dispense head cassettes (F0L59A). SU-DHL-1 were seeded in 30 µL of 3000 cells per well in compound preprinted 384 plates using the Thermo Fisher combi instrument. After 24 hours, cell viability was determined using a CellTiter-Glo Luminescent Cell Viability Assay (Promega, G7571) following the manufacturer's instructions. Dose-response curve fitting and $EC_{50}$ quantification were determined with four-parameter nonlinear regression analysis using GraphPad Prism v8.4.2.

**Statistical analysis**

Statistical tests were conducted using suitable underlying assumptions on variance characteristics and data distribution. Unless otherwise noted, two-tailed Student's *t*-tests were used for comparisons between groups.

**4. Nucleotide sequence of the luciferase plasmid generated in this study.**

**pSVBV1 (Flexi-NLuc-IRES2-FLuc)**

cgaacgaccgagcgcagcgagtcagtgagcgaggaagcggaagagcgcccaatacgcaaaccgcctctccccgcgcgttggccgattcattaatgca
gctggcacgacaggtttcccgactggaaagcgggcagtgagcgcaacgcaattaatgtgagttagctcactcattaggcaccccaggctttacactttatgct
tccggctcgtatgttgtgtggaattgtgagcggataacaatttcacacaggaaacagctatgaccatgattacgccaagcgcgcaattaaccctcactaaag
ggaacaaaagctggagctgcaagcttaatgtagtcttatgcaatactcttgtagtcttgcaacatggtaacgatgagttagcaacatgccttacaaggagaga
aaaagcaccgtgcatgccgattggtggaagtaaggtggtacgatcgtgccttattaggaaggcaacagacgggtctgacatggattggacgaaccactga
attgccgcattgcagagatattgtatttaagtgcctagctcgatacataaacgggtctctctggttagaccagatctgagcctgggagctctctggctaactagg
gaacccactgcttaagcctcaataaagcttgccttgagtgcttcaagtagtgtgtgcccgtctgttgtgtgactctggtaactagagatccctcagacccttttagt
cagtgtggaaaatctctagcagtggcgcccgaacagggacttgaaagcgaaagggaaaccagaggagctctctcgacgcaggactcggcttgctgaag
cgcgcacggcaagaggcgaggggcggcgactggtgagtacgccaaaaattttgactagcggaggctagaaggagagagatgggtgcgagagcgtca
gtattaagcgggggagaattagatcgcgatgggaaaaaattcggttaaggccagggggaaagaaaaaatataaattaaaacatatagtatgggcaagc
agggagctagaacgattcgcagttaatcctggcctgttagaaacatcagaaggctgtagacaaatactgggacagctacaaccatcccttcagacaggat
cagaagaacttagatcattatataatacagtagcaaccctctattgtgtgcatcaaaggatagagataaaagacaccaaggaagctttagacaagataga
ggaagagcaaaacaaaagtaagaccaccgcacagcaagcggccgctgatcttcagacctggaggaggagatatgagggacaattggagaagtgaat
tatataaatataaagtagtaaaaattgaaccattaggagtagcacccaccaaggcaaagagaagagtggtgcagagagaaaaaagagcagtgggaat
aggagctttgttccttgggttcttgggagcagcaggaagcactatgggcgcagcgtcaatgacgctgacggtacaggccagacaattattgtctggtatagtg
cagcagcagaacaatttgctgagggctattgaggcgcaacagcatctgttgcaactcacagtctggggcatcaagcagctccaggcaagaatcctggctg
tggaaagatacctaaaggatcaacagctcctggggatttggggttgctctggaaaactcatttgcaccactgctgtgccttggaatgctagttggagtaataaa
tctctggaacagatttggaatcacacgacctggatggagtgggacagagaaattaacaattacacaagcttaatacactccttaattgaagaatcgcaaaa
ccagcaagaaaagaatgaacaagaattattggaattagataaatgggcaagtttgtggaattggtttaacataacaaattggctgtggtatataaaattattca
taatgatagtaggaggcttggtaggtttaagaatagtttttgctgtactttctatagtgaatagagttaggcagggatattcaccattatcgtttcagacccacctcc
caaccccgaggggacccttgcgccttttccaaggcagccctgggtttgcgcagggacgcggctgctctgggcgtggttccgggaaacgcagcggcgccg
accctgggtctcgcacattcttcacgtccgttcgcagcgtcacccggatcttcgccgctacccttgtgggcccccggcgacgcttcctgctccgcccctaagt
cgggaaggttccttgcggttcgcggcgtgccggacgtgacaaacggaagccgcacgactcactagtaccctcgcagacggacagcgccagggagcaa
tggcagcgcgccgaccgcgatgggctgtggccaatagcggctgctcagcagggcgcgccgagagcagcggccgggaaggggcggtgcgggaggc
ggggtgtggggcggtagtgtgggccctgttcctgcccgcgcggtgttccgcattctgcaagcctccggagcgcacgtcggcagtcggctccctcgttgaccg
aatcaccgacctctctccccagggatcgataccgtcgactccggaatagccaccATGGAGACGGACGTCTCAGGCGGCGGCGGCT
CCGGCGGCGGCGGctccATGGTCTTCACACTCGAAGATTTCGTTGGGGACTGGCGACAGACAGCCGGCTA
CAACCTGGACCAAGTCCTTGAACAGGGAGGTGTGTCCAGTTTGTTTCAGAATCTCGGGGTGTCCGTAACT
CCGATCCAAAGGATTGTCCTGAGCGGTGAAAATGGGCTGAAGATCGACATCCATGTCATCATCCCGTATG
AAGGTCTGAGCGGCGACCAAATGGGCCAGATCGAAAAAATTTTTAAGGTGGTGTACCCTGTGgatgatcatcac
tttaaggtgatcctgcactatggcacactggtaatcgacggggttacgccgaacatgatcgactatttcggacggccgtatgaaggcatcgccgtgttcgacg
gcaaaaagatcactgtaacagggaccctgtggaacggcaacaaaattatcgacgagcgcctgatcaaccccgacggctccctgctgttccgagtaacca

tcaacggagtgaccggctggcggctgtgcgaacgcattctggcgtaaggatccactagtccagtgtggtgagtcagggagggtcgagccattgaagggc
acaaaaggaaactcaccctaactgtaaagtaattgtgtgttttgagactataagtatcccttggagaaccaccttgttggcgggtacaccgaatgctgtttaa
gagctaagctggaaaaagtggcaccgagtcgcccctctccctcccccccccctaacgttactggccgaagccgcttggaataaggccggtgtgcgtttgtct
atatgttattttccaccatattgccgtcttttggcaatgtgagggcccggaaacctggccctgtcttcttgacgagcattcctaggggtctttcccctctcgccaaag
gaatgcaaggtctgttgaatgtcgtgaaggaagcagttcctctggaagcttcttgaagacaaacaacgtctgtagcgaccctttgcaggcagcggaacccc
ccacctggcgacaggtgcctctgcggccaaaagccacgtgtataagatacacctgcaaaggcggcacaaccccagtgccacgttgtgagttggatagttg
tggaaagagtcaaatggctctcctcaagcgtattcaacaaggggctgaaggatgcccagaaggtaccccattgtatgggatctgatctggggcctcggtgc
acatgctttacatgtgtttagtcgaggttaaaaaaacgtctaggccccccgaaccacggggacgtggttttcctttgaaaaacacgatgataatatggccaca
accATGGAAGATGCCAAAAACATTAAGAAGGGCCCAGCGCCATTCTACCCACTCGAAGACGGGACCGCCG
GCGAGCAGCTGCACAAAGCCATGAAGCGCTACGCCCTGGTGCCCGGCACCATCGCCTTTACCGACGCAC
ATATCGAGGTGGACATTACCTACGCCGAGTACTTCGAGATGAGCGTTCGGCTGGCAGAAGCTATGAAGC
GCTATGGGCTGAATACAAACCATCGGATCGTGGTGTGCAGCGAGAATAGCttgcagttcttcatgcccgtgttgggtgccct
gttcatcggtgtggctgtggccccagctaacgacatctacaacgagcgcgagctgctgaacagcatgggcatcagccagcccaccgtcgtattcgtgagc
aagaaagggctgcaaaagatcctcaacgtgcaaaagaagctaccgatcatacaaaagatcatcatcatggatagcaagaccgactaccagggcttcca
aagcatgtacaccttcgtgacttcccatttgccacccggcttcaacgagtacgacttcgtgcccgagagcttcgaccgggacaaaaccatcgccctgatcat
gaacagtagtggcagtaccggattgcccaagggcgtagccctaccgcaccgcaccgcttgtgtccgattcagtcatgcccgcgaccccatcttcggcaacc
agatcatccccgacaccgctatcctcagcgtggtgccatttcaccacggcttcggcatgttcaccacgctgggctacttgatctgcggctttcgggtcgtgctca
tgtaccgcttcgaggaggagctattcttgcgcagcttgcaagactataagattcaatctgccctgctggtgcccacactatttagcttcttcgctaagagcactct
catcgacaagtacgacctaagcaacttgcacgagatcgccagcggcggggcgccgctcagcaaggaggtaggtgaggccgtggccaaacgcttccac
ctaccaggcatccgccagggctacggcctgacagaaacaaccagcgccattctgatcacccccgaaggggacgacaagcctggcgcagtaggcaag
gtggtgcccttcttcgaggctaaggtggtggacttggacaccggtaagacactgggtgtgaaccagcgcggcgagctgtgcgtccgtggccccatgatcat
gagcggctacgttaacaaccccgaggctacaaacgctctcatcgacaaggacggctggctgcacagcggcgacatcgcctactgggacgaggacgag
cacttcttcatcgtggaccggctgaagagcctgatcaaatacaagggctaccaggtagccccagccgaactggagagcatcctgctgcaacaccccaac
atcttcgacgccggggtcgccggcctgcccgacgacgatgccggcgagctgcccgccgcagtcgtcgtgctggaacacggtaaaaccatgaccgagaa
ggagatcgtggactatgtggccagccaggttacaaccgccaagaagctgcgcggtggtgttgtgttcgtggacgaggtgcctaaaggactgaccggcaa
gttggacgcccgcaagatccgcgagattctcattaaggccaagaagggcggcaagatcgccgtgtgaccggcggcatggacgagctgtacaagtaaac
tagtaagcttggcgtaactagatcttgagacaaatggcagtattcatccacaattttaaaagaaaaggggggattggggggtacagtgcaggggaaagaa
tagtagacataatagcaacagacatacaaactaaagaattacaaaaacaaattacaaaaattcaaaattttcgggtttattacagggacagcagagatcc
actttgggctcgagggggcccgggtgcaaagatggataaagttttaaacagagaggaatctttgcagctaatggaccttctaggtcttgaaaggagtggga
attggctccggtgcccgtcagtgggcagagcgcacatcgcccacagtccccgagaagttggggggaggggtcggcaattgatccggtgcctagagaag
gtggcgcggggtaaactgggaaagtgatgtcgtgtactggctccgcctttttcccgagggtgggggagaaccgtatataagtgcagtagtcgccgtgaacgt
tctttttcgcaacgggtttgccgccagaacacaggtaagtgccgtgtgtggttcccgcgggcctggcctctttacgggttatggcccttgcgtgccttgaattactt
ccacctggctgcagtacgtgattcttgatcccgagcttcgggttggaagtgggtgggagagttcgaggccttgcgcttaaggagccccttcgcctcgtgcttga
gttgaggcctggcctgggcgctggggccgccgcgtgcgaatctggtggcaccttcgcgcctgtctcgctgctttcgataagtctctagccatttaaaatttttgat
gacctgctgcgacgctttttttctggcaagatagtcttgtaaatgcgggccaagatctgcacactggtatttcggtttttggggccgcgggcggcgacggggcc
cgtgcgtcccagcgcacatgttcggcgaggcggggcctgcgagcgcggccaccgagaatcggacgggggtagtctcaagctggccggcctgctctggt
gcctggcctcgcgccgccgtgtatcgccccgccctgggcggcaaggctggcccggtcggcaccagttgcgtgagcggaaagatggccgcttcccggccc
tgctgcagggagctcaaaatggaggacgcggcgctcgggagagcgggcgggtgagtcacccacacaaaggaaaagggcctttccgtcctcagccgtc
gcttcatgtgactccacggagtaccgggcgccgtccaggcacctcgattagttctcgagcttttggagtacgtcgtctttaggttggggggaggggttttatgcg
atggagtttccccacactgagtgggtggagactgaagttaggccagcttggcacttgatgtaattctccttggaatttgccctttttgagtttggatcttggttcattct
caagcctcagacagtggttcaaagtttttttcttccatttcaggtgtcgtgacgtacggccaccatgaccgagtacaagcccacggtgcgcctcgccacccgc
gacgacgtccccagggccgtacgcaccctcgccgccgcgttcgccgactaccccgccacgcgccacaccgtcgatccggaccgccacatcgagcggg
tcaccgagctgcaagaactcttcctcacgcgcgtcgggctcgacatcggcaaggtgtgggtcgcggacgacggcgccgccgtggcggtctggaccacgc
cggagagcgtcgaagcgggggcggtgttcgccgagatcggcccgcgcatggccgagttgagcggttcccggctggccgcgcagcaacagatggaagg
cctcctggcgccgcaccggcccaaggagcccgcgtggttcctggccaccgtcggagtctcgcccgaccaccagggcaagggtctgggcagcgccgtcg
tgctccccggagtggaggcggccgagcgcgccggggtgcccgccttcctggagacctccgcgccccgcaacctccccttctacgagcggctcggcttcac
cgtcaccgccgacgtcgaggtgcccgaaggaccgcgcacctggtgcatgacccgcaagcccggtgcctgaacgcgttaagtcgacaatcaacctctgg
attacaaaatttgtgaaagattgactggtattcttaactatgttgctccttttacgctatgtggatacgctgctttaatgcctttgtatcatgctattgcttcccgtatggct
ttcattttctcctccttgtataaatcctggttgctgtctctttatgaggagttgtggcccgttgtcaggcaacgtggcgtggtgtgcactgtgtttgctgacgcaacccc
cactggttggggcattgccaccacctgtcagctcctttccgggactttcgctttccccctccctattgccacggcggaactcatcgccgcctgccttgcccgctgc
tggacaggggctcggctgttgggcactgacaattccgtggtgttgtcggggaaatcatcgtcctttccttggctgctcgcctgtgttgccacctggattctgcgcg
ggacgtccttctgctacgtcccttcggccctcaatccagcggaccttccttcccgcggcctgctgccggctctgcggcctcttccgcgtcttcgccttcgccctca
gacgagtcggatctccctttgggccgcctccccgcgtcgactttaagaccaatgacttacaaggcagctgtagatcttagccactttttaaaagaaaagggggg
gactggaagggctaattcactcccaacgaagacaagatctgcttttgcttgtactgggtctctctggttagaccagatctgagcctgggagctctctggctaac

tagggaacccactgcttaagcctcaataaagcttgccttgagtgcttcaagtagtgtgtgcccgtctgttgtgtgactctggtaactagagatccctcagaccctttt
tagtcagtgtggaaaatctctagcagtacgtatagtagttcatgtcatcttattattcagtatttataacttgcaaagaaatgaatatcagagagtgagaggaactt
gtttattgcagcttataatggttacaaataaagcaatagcatcacaaatttcacaaataaagcatttttttcactgcattctagttgtggtttgtccaaactcatcaat
gtatcttatcatgtctggctctagctatcccgcccctaactccgcccatcccgcccctaactccgcccagttccgcccattctccgccccatggctgactaatttttt
ttatttatgcagaggccgaggccgcctcggcctctgagctattccagaagtagtgaggaggcttttttggaggcctagggacgtacccaattcgccctatagtg
agtcgtattacgcgcgctcactggccgtcgttttacaacgtcgtgactgggaaaaccctggcgttacccaacttaatcgccttgcagcacatccccctttcgcc
agctggcgtaatagcgaagaggcccgcaccgatcgcccttcccaacagttgcgcagcctgaatggcgaatggacgcgccctgtagcggcgcattaag
cgcggcgggtgtggtggttacgcgcagcgtgaccgctacacttgccagcgccctagcgcccgctcctttcgctttcttcccttcctttctcgccacgttcgccggc
tttccccgtcaagctctaaatcgggggctccctttagggttccgatttagtgctttacggcacctcgaccccaaaaaacttgattagggtgatggttcacgtagtg
ggccatcgccctgatagacggtttttcgccctttgacgttggagtccacgttctttaatagtggactcttgttccaaactggaacaacactcaaccctatctcggtc
tattcttttgatttataagggattttgccgatttcggcctattggttaaaaaatgagctgatttaacaaaaatttaacgcgaattttaacaaaatattaacgcttacaat
ttaggtggcacttttcggggaaatgtgcgcggaacccctatttgtttatttttctaaatacattcaaatatgtatccgctcatgagacaataaccctgataaatgcttc
aataatattgaaaaaggaagagtatgagtattcaacatttccgtgtcgcccttattcccttttttgcggcattttgccttcctgtttttgctcacccagaaacgctggtg
aaagtaaaagatgctgaagatcagttgggtgcacgagtgggttacatcgaactggatctcaacagcggtaagatccttgagagttttcgccccgaagaacg
ttttccaatgatgagcacttttaaagttctgctatgtggcgcggtattatcccgtattgacgccgggcaagagcaactcggtcgccgcatacactattctcagaat
gacttggttgagtactcaccagtcacagaaaagcatcttacggatggcatgacagtaagagaattatgcagtgctgccataaccatgagtgataacactgc
ggccaacttacttctgacaacgatcggaggaccgaaggagctaaccgcttttttgcacaacatgggggatcatgtaactcgccttgatcgttgggaaccgga
gctgaatgaagccataccaaacgacgagcgtgacaccacgatgcctgtagcaatggcaacaacgttgcgcaaactattaactggcgaactacttactcta
gcttcccggcaacaattaatagactggatggaggcggataaagttgcaggaccacttctgcgctcggcccttccggctggctggtttattgctgataaatctgg
agccggtgagcgtgggtctcgcggtatcattgcagcactggggccagatggtaagccctcccgtatcgtagttatctacacgacggggagtcaggcaacta
tggatgaacgaaatagacagatcgctgagataggtgcctcactgattaagcattggtaactgtcagaccaagtttactcatatatactttagattgatttaaaac
ttcatttttaatttaaaaggatctaggtgaagatcctttttgataatctcatgaccaaaatcccttaacgtgagttttcgttccactgagcgtcagaccccgtagaaa
agatcaaaggatcttcttgagatcctttttttctgcgcgtaatctgctgcttgcaaacaaaaaaaccaccgctaccagcggtggtttgtttgccggatcaagagct
accaactctttttccgaaggtaactggcttcagcagagcgcagataccaaatactgttcttctagtgtagccgtagttaggccaccacttcaagaactctgtagc
accgcctacatacctcgctctgctaatcctgttaccagtggctgctgccagtggcgataagtcgtgtcttaccgggttggactcaagacgatagttaccggata
aggcgcagcggtcgggctgaacggggggttcgtgcacacagcccagcttggagcgaacgacctacaccgaactgagatacctacagcgtgagctatga
gaaagcgccacgcttcccgaagggagaaaggcggacaggtatccggtaagcggcagggtcggaacaggagagcgcacgagggagcttccagggg
gaaacgcctggtatctttatagtcctgtcgggtttcgccacctctgacttgagcgtcgatttttgtgatgctcgtcaggggggcggagcctatggaaaaacgcca
gcaacgcggcctttttacggttcctggccttttgctggccttttgctcacatgttctttcctgcgttatcccctgattctgtggataaccgtattaccgcctttgagtgagc
tgataccgctcgccgcagc
