## Supplementary material for "Proteolysis Targeting Chimeras With Reduced Off-targets": SI_Chemistry

$^{1}H$ and $^{13}C$ NMR of Entry **49** ... 87
$^{19}F$ NMR of Entry **49** ... 88
$^{1}H$ and $^{13}C$ NMR of Entry **51** ... 89
$^{1}H$ and $^{13}C$ NMR of Entry **52** ... 90
$^{1}H$ and $^{13}C$ NMR of Entry **53** ... 91
$^{1}H$ and $^{13}C$ NMR of Entry **54** ... 92
$^{1}H$ and $^{13}C$ NMR of Entry **55** ... 93
$^{1}H$ and $^{13}C$ NMR of Entry **57** ... 94
$^{1}H$ and $^{13}C$ NMR of Entry **58** ... 95
$^{1}H$ and $^{13}C$ NMR of Entry **59** ... 96
$^{1}H$ and $^{13}C$ NMR of Entry **60** ... 97
$^{1}H$ and $^{13}C$ NMR of Entry **61** ... 98
$^{1}H$ and $^{13}C$ NMR of Entry **62** ... 99
$^{1}H$ and $^{13}C$ NMR of Entry **63** ... 100
$^{1}H$ and $^{13}C$ NMR of Entry **64** ... 101
$^{1}H$ and $^{13}C$ NMR of Entry **65** ... 102
$^{1}H$ and $^{13}C$ NMR of Entry **66** ... 103
$^{1}H$ and $^{13}C$ NMR of Entry **67** ... 104
$^{1}H$ and $^{13}C$ NMR of Entry **68** ... 105
$^{1}H$ and $^{13}C$ NMR of Entry **69** ... 106
$^{1}H$ and $^{13}C$ NMR of Entry **70** ... 107
$^{1}H$ and $^{13}C$ NMR of Entry **71** ... 108
$^{1}H$ and $^{13}C$ NMR of Entry **6** ... 109
$^{1}H$ and $^{13}C$ NMR of Entry **72** ... 110
$^{1}H$ and $^{13}C$ NMR of Entry **73** ... 112
$^{1}H$ and $^{13}C$ NMR of Entry **74** ... 113
$^{1}H$ and $^{13}C$ NMR of Entry **75** ... 114
$^{1}H$ and $^{13}C$ NMR of Entry **76** ... 115
$^{1}H$ and $^{13}C$ NMR of Entry **77** ... 116
$^{1}H$ and $^{13}C$ NMR of Entry **78** ... 118
$^{1}H$ and $^{13}C$ NMR of Entry **79** ... 119
$^{1}H$ and $^{13}C$ NMR of Entry **80** ... 120
$^{1}H$ and $^{13}C$ NMR of Entry **81** ... 121
$^{1}H$ and $^{13}C$ NMR of Entry **16** ... 122
$^{1}H$ and $^{13}C$ NMR of Entry **7** ... 123
$^{1}H$ and $^{13}C$ NMR of Entry **14** ... 124
$^{1}H$ and $^{13}C$ NMR of **dALK-1** ... 125
$^{1}H$ and $^{13}C$ NMR of **dALK-2** ... 126
$^{1}H$ and $^{13}C$ NMR of **dALK-3** ... 127
$^{1}H$ and $^{13}C$ NMR of **dALK-4** ... 128
$^{19}F$ NMR of **dALK-4** ... 129
$^{1}H$ and $^{13}C$ NMR of **dALK-5** ... 130
$^{1}H$ and $^{13}C$ NMR of **dALK-6** ... 131
$^{19}F$ NMR of **dALK-6** ... 132

#### General methods and materials

All reactions containing water or air-sensitive reagents were performed in oven-dried glassware under nitrogen or argon. All reagents were purchased and used as received from commercial sources without further purification. Reactions were implemented in round-bottom flasks or vials stirred with Teflon®-coated magnetic stir bars. Moisture and air-sensitive reactions were performed under a dry nitrogen/argon atmosphere. Moisture and air-sensitive liquids or solutions were transferred via nitrogen-flushed syringes. As necessary, organic solvents were degassed by bubbling nitrogen/argon through the liquid. The reaction progress was monitored by thin-layer chromatography (TLC) and ultra-performance liquid chromatography-mass spectrometry (UPLC-MS). Flash column chromatography was performed using silica gel (60 Å mesh, 20–40 µm) on a Teledyne ISCO CombiFlash Rf system. Pomalidomide analogs and ALK-PROTACs (**dALK-1** to **12**) were further purified by RP-Preparative HPLC using a Teledyne ISCO ACCQPrep HP150 instrument equipped with an XBridge C18 column (19 x 250 mm, 5 µm with a gradient of 10-100% MeCN – $H_2O$ (both solvents contain 0.1% Formic acid) before testing them in biological assays. Analytical TLC was performed using Merck Silica gel 60 F254 pre-coated plates (0.25 mm); illumination at 254 nm allowed the visualization of UV-active material. UPLC-MS was performed on a Waters ACQUITY UPLC I-Class PLUS System with an ACQUITY SQ Detector 2. Nuclear magnetic resonance (NMR) spectra were recorded on a Bruker 400 Spectrometer ($^{1}$H NMR, 400 MHz; $^{13}$C, 101 MHz) at the Broad Institute of MIT and Harvard. $^{1}$H and $^{13}$C chemical shifts are indicated in parts per million (ppm) relative to $SiMe_4$ (δ = 0.00 ppm) and are internally referenced to residual solvent signals. NMR solvents were purchased from Cambridge Isotope Laboratories, Inc., and NMR data were obtained in DMSO-$d_6$. Data for $^{1}$H NMR are reported as follows: chemical shift value in ppm, multiplicity (s = singlet, d = doublet, t = triplet, dd = doublet of doublets, and m = multiplet), integration value, and coupling constant value in Hz. High-resolution mass spectra were recorded on a Thermo Q Exactive Plus mass spectrometer system equipped with a HESI-II electrospray ionization source at Harvard Center for Mass Spectrometry at the Harvard FAS Division of Science Core Facility.

**General Procedure A**: Preparation of 4-substituted IMiD analogs through $S_NAr$

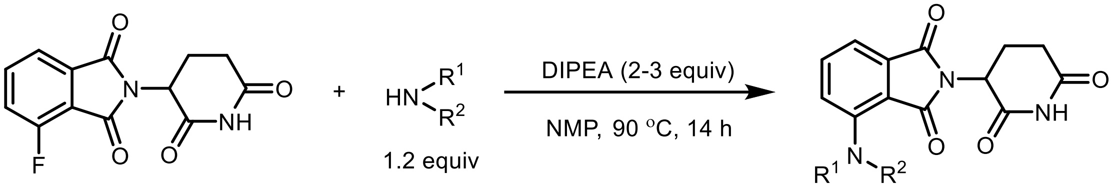

In an oven-dried 5 mL vial with a stir-bar, DIPEA (0.34 mmol, 2.5 equiv.) was added to a solution of 2-(2,6-dioxopiperidin-3-yl)-4-fluoroisoindoline-1,3-dione (54 mg, 0.2 mmol), and 1° or 2° amines (0.26 mmol 1.3 equiv.) in dry *N*-methyl pyrrolidine (1.5 mL) at rt. The reaction mixture was stirred at 90 °C for 14 h. The reaction mixture was diluted with ethyl acetate (60 mL), washed with water (10 mL x3), followed by Brine (10 mL), dried over anhydrous $Na_2SO_4$, filtered, and concentrated under reduced pressure. The crude residue was purified by Combiflash ISCO to obtain the pure product.

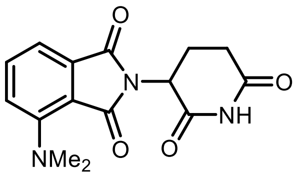

**4-(dimethylamino)-2-(2,6-dioxopiperidin-3-yl)isoindoline-1,3-dione (15)** was prepared by following the general procedure **A** using dimethylamine (0.26 mmol). The final product was isolated by flash column chromatography ($CH_2Cl_2$: MeOH 90:10) in 53% yield (32 mg; yellow solid). $^1$H and $^{13}$C NMR spectral data were matched with the literature reported values.[1]

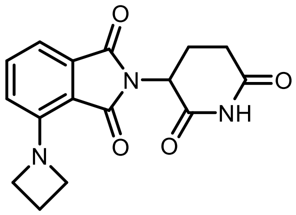

**4-(azetidin-1-yl)-2-(2,6-dioxopiperidin-3-yl)isoindoline-1,3-dione (16)** was prepared by following the general procedure **A** using azetidine. HCl (0.26 mmol). Final product was isolated by flash column chromatography ($CH_2Cl_2$: MeOH 90:10) in 61% yield (38 mg; yellow solid). $^1$H NMR (400 MHz, DMSO-$d_6$) δ 11.05 (s, 1H), 7.53 (dd, $J$ = 8.5, 7.0 Hz, 1H), 7.08 (d, $J$ = 6.9 Hz, 1H), 6.72 (d, $J$ = 8.4 Hz, 1H), 5.05 (dd, $J$ = 12.9, 5.4 Hz, 1H), 4.17 (t, $J$ = 7.5 Hz, 4H), 2.96 – 2.79 (m, 1H), 2.63 – 2.47 (m, 2H), 2.34 – 2.25 (m, 2H), 2.07 – 1.96 (m, 1H). $^{13}$C NMR (101 MHz, DMSO-$d_6$) δ 172.7, 170.0,167.2, 166.4, 148.1, 134.8, 133.3, 119.7, 111.5, 109.8, 54.1, 48.6, 30.9, 22.1, 16.0. HRMS (ESI) calc'd for $C_{16}H_{15}N_3O_4$ + H = 314.1135, found 314.1135.

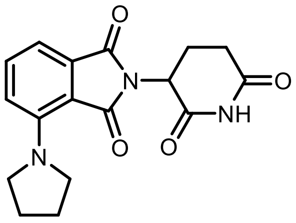

**2-(2,6-dioxopiperidin-3-yl)-4-(pyrrolidin-1-yl)isoindoline-1,3-dione (17)** was prepared by following the general procedure **A** using pyrrolidine (0.26 mmol). Final product was isolated by flash column chromatography ($CH_2Cl_2$: MeOH 90:10) in 67% yield (44 mg; yellow solid). $^1$H NMR (400 MHz, DMSO-$d_6$) δ 10.84 (s, 1H), 7.35 (dd, $J$ = 8.6, 6.9 Hz, 1H), 6.96 – 6.82 (m, 2H), 4.86 (dd, $J$ = 12.8, 5.4 Hz, 1H), 3.38 – 3.27 (m, 3H), 3.10 (d, $J$ = 7.0 Hz, 1H), 2.79 – 2.56 (m, 1H), 2.44 – 2.29 (m, 3H), 1.98 (t, $J$ = 8.1 Hz, 1H), 1.87 – 1.77 (m, 1H), 1.73 (d, $J$ = 3.4 Hz, 3H). $^{13}$C NMR (101 MHz, DMSO-$d_6$) δ 172.7, 170.0, 167.1, 166.5, 146.0, 134.7, 133.9, 121.2, 111.2, 109.9, 51.3, 48.7, 48.5, 30.9, 30.1, 29.0, 25.2, 22.1, 17.2. HRMS (ESI) calc'd for $C_{17}H_{17}N_3O_4$ + H = 328.1292, found 328.1291.

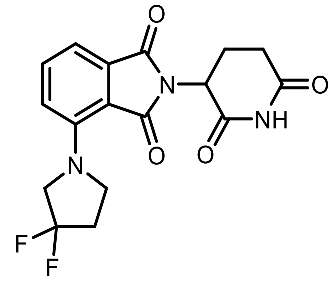

**4-(3,3-difluoropyrrolidin-1-yl)-2-(2,6-dioxopiperidin-3-yl)isoindoline-1,3-dione (18)** was prepared by following the general procedure **A** using 3,3-difluoropyrrolidine (0.26 mmol). Final product was isolated by flash column chromatography ($CH_2Cl_2$: MeOH 90:10) in 61% yield (44 mg; yellow solid). $^1$H NMR (400 MHz, DMSO-$d_6$) δ 11.10 (s, 1H), 7.66 (dd, *J* = 8.6, 7.0 Hz, 1H), 7.27 (d, *J* = 7.0 Hz, 1H), 7.19 (d, *J* = 8.6 Hz, 1H), 5.09 (dd, *J* = 12.9, 5.4 Hz, 1H), 4.06 (t, *J* = 13.4 Hz, 2H), 3.75 (t, *J* = 7.2 Hz, 2H), 2.88 (ddd, *J* = 17.4, 14.1, 5.4 Hz, 1H), 2.68 – 2.51 (m, 4H), 2.13 – 1.91 (m, 1H). $^{19}$F NMR (376 MHz, DMSO-$d_6$) δ -100.35. $^{13}$C NMR (101 MHz, DMSO-$d_6$) δ 172.8, 169.9, 166.9, 166.6, 145.3, 135.3, 133.8, 128.1 (t, *J* = 247.5 Hz), 121.5, 113.1, 112.0, 57. 5 (t, *J* = 35.3 Hz), 48.9, 48.2 (t, *J* = 5.05 Hz), 33.0 (t, *J* = 25.3 Hz), 30.9, 22.1. HRMS (ESI) calc'd for $C_{17}H_{15}F_2N_3O_4$ + H = 364.1103, found 364.1101.

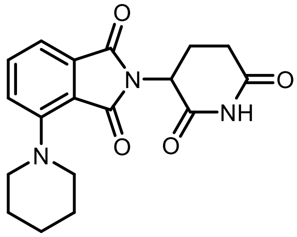

**2-(2,6-dioxopiperidin-3-yl)-4-(piperidin-1-yl)isoindoline-1,3-dione (19)** was prepared by following the general procedure **A** using piperidine (0.26 mmol). Final product was isolated by flash column chromatography ($CH_2Cl_2$: MeOH 95:5) in 71% yield (48 mg; yellow solid). $^1$H NMR (400 MHz, DMSO-$d_6$) δ 11.10 (s, 1H), 7.67 (dd, *J* = 8.5, 7.1 Hz, 1H), 7.31 (dd, *J* = 7.8, 2.7 Hz, 2H), 5.09 (dd, *J* = 12.9, 5.4 Hz, 1H), 3.23 (t, *J* = 5.2 Hz, 4H), 2.87 (ddd, *J* = 17.4, 14.1, 5.5 Hz, 1H), 2.65 – 2.51 (m, 2H), 2.02 (dtd, *J* = 10.7, 6.0, 3.3 Hz, 1H), 1.67 (q, *J* = 5.6 Hz, 4H), 1.57 (q, *J* = 5.8 Hz, 2H). $^{13}$C NMR (101 MHz, DMSO-$d_6$) δ 172.9, 170.1, 167.1, 166.3, 150.5, 135.8, 133.7, 123.9, 116.4, 114.4, 51.9, 48.8, 31.0, 25.6, 23.6, 22.1. HRMS (ESI) calc'd for $C_{18}H_{19}N_3O_4$ + H = 342.1448, found 342.1447.

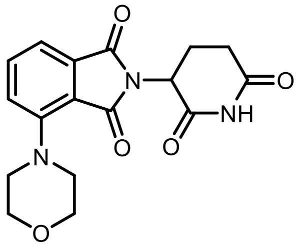

**2-(2,6-dioxopiperidin-3-yl)-4-morpholinoisoindoline-1,3-dione (20)** was prepared by following the general procedure **A** using morpholine (0.26 mmol). Final product was isolated by flash column chromatography ($CH_2Cl_2$: MeOH 90:10) in 71% yield (44 mg; orange solid). $^1$H NMR (400 MHz, DMSO-$d_6$) δ 11.07 (s, 1H), 7.72 (dd, *J* = 8.4, 7.2 Hz, 1H), 7.39 (d, *J* = 7.1 Hz, 1H), 7.35 (d, *J* = 8.4 Hz, 1H), 5.10 (dd, *J* = 12.9, 5.4 Hz, 1H), 3.77 (dd, *J* = 5.7, 3.5 Hz, 4H), 3.28 (t, *J* = 4.6 Hz, 4H), 2.87 (ddd, *J* = 17.4, 14.0, 5.4 Hz, 1H), 2.65 – 2.51 (m, 2H), 2.08 – 1.97 (m, 1H). $^{13}$C NMR (101 MHz, DMSO-$d_6$) δ 172.7, 169.9, 167.0, 166.3, 149.7, 135.9, 133.6, 123.5, 116.7, 115.1, 66.1, 50.8, 48.8, 39.5, 30.9, 22.0. HRMS (ESI) calc'd for $C_{17}H_{17}N_3O_5$ + H = 344.1241, found 344.1240.

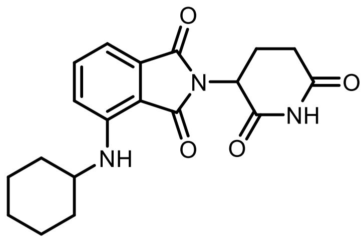

**4-(cyclohexylamino)-2-(2,6-dioxopiperidin-3-yl) isoindoline-1,3-dione (21)** was prepared by following the general procedure **A** using cyclohexylamine (0.26 mmol). Final product was isolated by flash column chromatography ($CH_2Cl_2$: MeOH 95:5) in 57% yield (40 mg; yellow solid). $^1$H NMR (400 MHz, DMSO-$d_6$) δ 11.08 (s, 1H), 7.57 (dd, *J* = 8.5, 7.1 Hz, 1H), 7.13 (d, *J* = 8.6 Hz, 1H), 7.02 (d, *J* = 7.0 Hz, 1H), 6.23 (d, *J* = 8.3 Hz, 1H), 5.05 (dd, *J* = 12.9, 5.4 Hz, 1H), 3.55 (tq, *J* = 9.5, 5.7, 4.9 Hz, 1H), 2.88 (ddd, *J* = 17.0, 13.9, 5.2 Hz, 1H), 2.61 – 2.50 (m, 1H), 2.02 (ddt, *J* = 13.1, 5.6, 2.7 Hz, 1H), 1.98 – 1.87 (m, 2H), 1.68 (dq, *J* = 11.9, 3.9 Hz, 2H), 1.58 (dt, *J* = 12.6, 4.1 Hz, 1H), 1.45 – 1.34 (m, 2H), 1.35 – 1.17 (m, 3H).$^{13}$C NMR (101 MHz, DMSO-$d_6$) δ 172.7, 170.0, 169.2, 167.2, 145.5, 136.3, 132.2, 117.6, 110.5,

109.0, 49.8, 48.6, 32.2, 31.0, 25.1, 24.0, 22.1. HRMS (ESI) calc'd for $C_{19}H_{21}N_3O_4$ + H = 356.1605, found 356.1603.

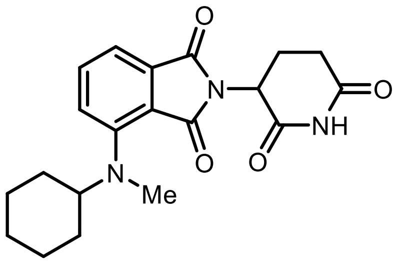

**4-(cyclohexyl(methyl)amino)-2-(2,6-dioxopiperidin-3-yl) isoindoline-1,3-dione (22)** was prepared by following the general procedure **A** using *N*-Methyl cyclohexylamine (0.26 mmol). Final product was isolated by flash column chromatography ($CH_2Cl_2$: MeOH 95:5) in 72% yield (53 mg; yellow solid). $^1$H NMR (400 MHz, DMSO-$d_6$) δ 11.09 (s, 1H), 7.61 (dd, *J* = 8.6, 7.0 Hz, 1H), 7.27 (d, *J* = 8.6 Hz, 1H), 7.22 (d, *J* = 7.0 Hz, 1H), 5.10 (dd, *J* = 12.9, 5.4 Hz, 1H), 3.66 (td, *J* = 10.4, 9.4, 5.6 Hz, 1H), 2.96 – 2.77 (m, 4H), 2.70 – 2.51 (m, 2H), 2.11 – 1.94 (m, 1H), 1.76 (d, *J* = 11.3 Hz, 4H), 1.63 – 1.48 (m, 3H), 1.37 – 1.19 (m, 2H), 1.09 (dtd, *J* = 13.3, 9.8, 3.2 Hz, 1H). $^{13}$C NMR (101 MHz, DMSO-$d_6$) δ 172.9, 170.1, 167.1, 166.4, 149.7, 135.2, 134.1, 123.6, 114.0, 113.0, 62.6, 48.8, 33.1, 31.0, 29.7, 29.7, 25.6, 25.6, 25.2, 22.1. HRMS (ESI) calc'd for $C_{20}H_{23}N_3O_4$ + H = 370.1761, found 370.1758.

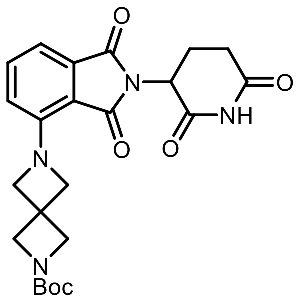

**2,6-dioxopiperidin-3-yl)-1,3-dioxoisoindolin-4-yl)-2,6-diazaspiro[3.3]heptane-2-carboxylate (23)** was prepared by following the general procedure **A** using Boc-2,6-diazaspiro[3,3]heptane HCl (0.71 mmol, 1.3 equiv.) The pure product was isolated by ISCO and the column ran with $CH_2Cl_2$, then grading to 5% MeOH (contains 1% $NH_4OH$)-$CH_2Cl_2$ in 69% yield (170 mg; yellow solid).$^1$H NMR (400 MHz, DMSO-$d_6$) δ 11.05 (s, 1H), 7.57 (dd, *J* = 8.4, 7.1 Hz, 1H), 7.13 (d, *J* = 7.0 Hz, 1H), 6.77 (d, *J* = 8.5 Hz, 1H), 4.31 (s, 4H), 4.03 (s, 4H), 2.87 (ddd, *J* = 16.7, 13.6, 5.4 Hz, 1H), 2.64 – 2.50 (m, 2H), 2.06 – 1.94 (m, 1H), 1.38 (s, 9H). $^{13}$C NMR (101 MHz, DMSO-$d_6$) δ 173.2, 170.4, 167.6, 166.9, 155.9, 148.1, 135.5, 133.7, 120.6, 112.4, 110.9, 79.1, 64.1, 59.7, 49.1, 33.0, 31.4, 28.6, 22.6. HRMS (ESI) calc'd for $C_{23}H_{26}N_4O_6$ + H = 455.1925, found 455.1923.

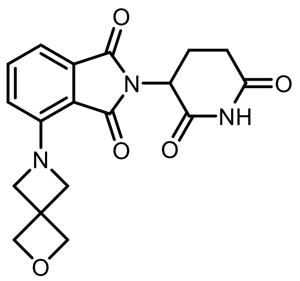

**2-(2,6-dioxopiperidin-3-yl)-4-(2-oxa-6-azaspiro[3.3]heptan-6-l)isoindoline-1,3-dione (24)** was prepared by following the general procedure **A** using 2-Oxa-6-azaspiro[3.3]heptane (0.26 mmol). The pure product was isolated by ISCO and the column ran with $CH_2Cl_2$, then grading to 5% MeOH (contains 1% $NH_4OH$)-$CH_2Cl_2$ in 65% yield (25 mg; yellow solid).$^1$H NMR (400 MHz, DMSO-$d_6$) δ 11.05 (s, 1H), 7.57 (dd, *J* = 8.5, 7.0 Hz, 1H), 7.12 (d, *J* = 7.0 Hz, 1H), 6.79 (d, *J* = 8.5 Hz, 1H), 5.05 (dd, *J* = 12.7, 5.4 Hz, 1H), 4.72 (s, 4H), 4.36 (s, 4H), 2.88 (ddd, *J* = 16.8, 13.8, 5.2 Hz, 1H), 2.64 – 2.50 (m, 3H), 2.06 – 1.94 (m, 1H). $^{13}$C NMR (101 MHz, DMSO-$d_6$) δ 172.7, 169.9, 167.2, 166.5, 147.7, 134.9, 133.2, 120.1, 111.9, 110.4, 79.8, 63.1, 54.9, 48.6, 38.0, 30.9, 22.1. HRMS (ESI) calc'd for $C_{18}H_{17}N_3O_5$ + H = 356.1241, found 356.1241.

**4-(4-acetylpiperazin-1-yl)-2-(2,6-dioxopiperidin-3-yl)isoindoline-1,3-dione (25)** was prepared by following the general procedure **A** using 1-(piperazin-1-yl)ethan-1-one (0.26 mmol). Final product was isolated by flash column chromatography (DCM: MeOH 90:10) in 78% yield (60 mg; yellow solid). $^{1}$H NMR (400 MHz, DMSO-$d_6$) δ 11.11 (s, 1H), 7.78 – 7.67 (m, 1H), 7.39 (d, $J$ = 7.1 Hz, 1H), 7.34 (d, $J$ = 8.4 Hz, 1H), 5.11 (dd, $J$ = 12.9, 5.4 Hz, 1H), 3.62 (t, $J$ = 5.0 Hz, 4H), 3.40 – 3.19 (m, 6H), 2.88 (ddd, $J$ = 17.4, 14.0, 5.4 Hz, 1H), 2.63 – 2.51 (m, 2H), 2.04 (s, 4H).$^{13}$C NMR (101 MHz, DMSO-$d_6$) δ 172.9,170.0, 168.5, 167.0, 166.4, 149.5, 136.0, 133.6, 123.9, 116.9, 115.3, 50.9, 50.2, 48.8, 45.7, 40.8, 31.0, 22.1, 21.3. HRMS (ESI) calc'd for $C_{19}H_{20}N_4O_5$ + H = 385.1506, found 385.1500.

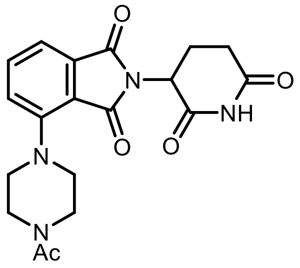

**2-(2,6-dioxopiperidin-3-yl)-4-(4-methylpiperazin-1-yl)isoindoline-1,3-dione (26)** was prepared by following the general procedure **A** using 1-(piperazin-1-yl)ethan-1-one (0.26 mmol). The pure product was isolated by ISCO and the column ran with $CH_2Cl_2$, then grading to 10% MeOH (contains 1% $NH_4OH$)-$CH_2Cl_2$ in 56% yield (40 mg; yellow solid). $^{1}$H NMR (400 MHz, DMSO-$d_6$) δ 11.08 (s, 1H), 7.69 (dd, $J$ = 8.4, 7.2 Hz, 1H), 7.34 (t, $J$ = 7.3 Hz, 2H), 5.09 (dd, $J$ = 12.9, 5.4 Hz, 1H), 3.33 (s, 4H), 2.88 (ddd, $J$ = 17.4, 14.0, 5.4 Hz, 1H), 2.70 – 2.50 (m, 6H), 2.23 (s, 3H), 2.02 (ddq, $J$ = 9.9, 5.6, 2.7 Hz, 1H). $^{13}$C NMR (101 MHz, DMSO-$d_6$) δ 172.8, 170.0, 167.0, 166.3, 149.7, 135.8, 133.7, 123.7, 116.5, 114.8, 54.5, 50.4, 48.8, 45.7, 30.9, 22.1. HRMS (ESI) calc'd for $C_{18}H_{20}N_4O_4$ + H = 357.1557, found 357.1552.

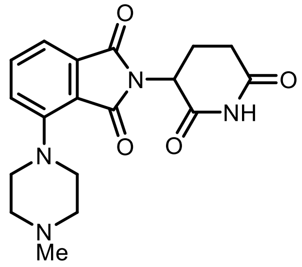

***tert*-butyl 4-(2-(2,6-dioxopiperidin-3-yl)-1,3-dioxoisoindolin-4-yl)piperazine-1-carboxylate (27)** was prepared by following the general procedure **A** using 1-Boc-piperazine (0.26 mmol). The pure product was isolated by ISCO and the column ran with $CH_2Cl_2$, then grading to 5% MeOH (contains 1% $NH_4OH$)-$CH_2Cl_2$ in 77% yield (67 mg; yellow solid). $^{1}$H NMR (400 MHz, DMSO-$d_6$) δ 11.08 (s, 1H), 7.72 (dd, $J$ = 8.4, 7.1 Hz, 1H), 7.37 (dd, $J$ = 15.5, 7.8 Hz, 2H), 5.10 (dd, $J$ = 12.9, 5.4 Hz, 1H), 3.51 (t, $J$ = 5.0 Hz, 4H), 3.25 (t, $J$ = 5.1 Hz, 4H), 2.88 (ddd, $J$ = 17.5, 14.1, 5.4 Hz, 1H), 2.65 – 2.52 (m, 2H), 2.11 – 1.96 (m, 1H), 1.42 (s, 9H).$^{13}$C NMR (101 MHz, DMSO-$d_6$) δ 172.8, 170.0, 167.0, 166.3, 153.9, 149.6, 135.9, 133.6, 123.9, 117.0, 115.3, 79.1, 50.4, 48.8, 31.0, 28.1, 22.0. HRMS (ESI) calc'd for $C_{22}H_{26}N_4O_6$ + H = 443.1925, found 443.1921.

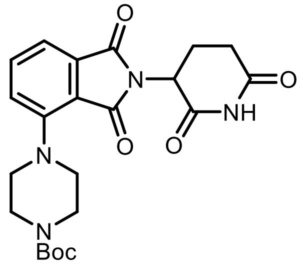

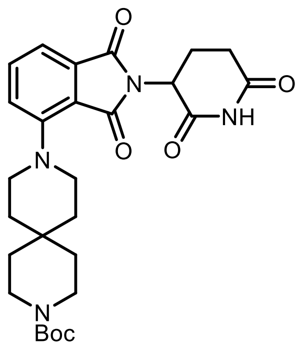

***tert*-butyl 9-(2-(2,6-dioxopiperidin-3-yl)-1,3-dioxoisoindolin-4-yl)-3,9-diazaspiro[5.5]undecane-3-carboxylate (28)** was prepared by following the general procedure **A** using *tert*-butyl 3,9-diazaspiro[5.5]undecane-3-carboxylate (0.26 mmol). The pure product was isolated by ISCO and the column ran with $CH_2Cl_2$, then grading to 5% MeOH (contains 1% $NH_4OH$)-$CH_2Cl_2$ in 55% yield (56 mg; yellow solid). $^1$H NMR (400 MHz, DMSO-$d_6$) δ 10.87 (s, 1H), 7.47 (dd, $J$ = 8.5, 7.1 Hz, 1H), 7.12 (dd, $J$ = 7.8, 6.0 Hz, 2H), 4.89 (dd, $J$ = 12.8, 5.4 Hz, 1H), 3.14 (d, $J$ = 7.0 Hz, 4H), 3.08 (d, $J$ = 6.5 Hz, 4H), 2.69 (ddd, $J$ = 17.1, 13.9, 5.4 Hz, 1H), 2.44 – 2.31 (m, 2H), 1.97 (d, $J$ = 8.2 Hz, 2H), 1.83 (dqd, $J$ = 11.6, 6.2, 3.5 Hz, 1H), 1.74 – 1.68 (m, 2H), 1.43 (t, $J$ = 5.4 Hz, 2H), 1.25 – 1.22 (m, 2H), 1.20 (s, 9H). $^{13}$C NMR (101 MHz, DMSO-$d_6$) δ 173.7, 172.7, 169.9, 167.0, 166.3, 153.9, 150.0, 135.6, 133.7, 123.7, 116.1, 114.3, 78.4, 48.8, 48.5, 46.4, 34.9, 31.0, 30.1, 29.2, 28.9, 28.1, 22.1, 17.2. HRMS (ESI) calc'd for $C_{27}H_{34}N_4O_6$ + H = 511.2551, found 511.2549.

**General Procedure B**: Preparation of 5-substituted IMiD analogs through $S_NAr$

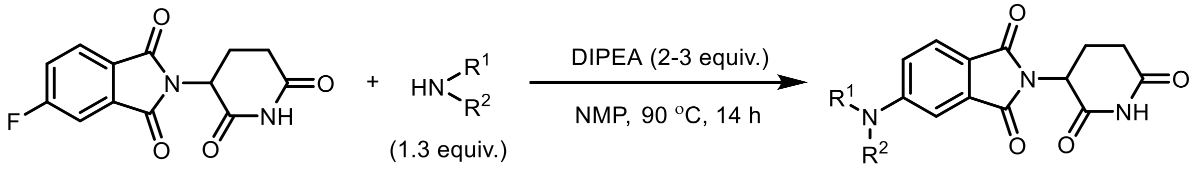

In an oven-dried 5 mL vial with a stir-bar, DIPEA (0.34 mmol, 2.5 equiv.) was added to a solution of 2-(2,6-dioxopiperidin-3-yl)-5-fluoroisoindoline-1,3-dione (54 mg, 0.2 mmol), and 1° or 2° amines (0.26 mmol 1.3 equiv.) in dry *N*-methyl pyrrolidine (1.5 mL) at rt. The reaction mixture was stirred at 90 °C for 14 h. The reaction mixture was diluted with ethylacetate (60 mL), washed with water (10 mL x3), followed by Brine (10 mL), dried over anhydrous $Na_2SO_4$, filtered, and concentrated under reduced pressure. The crude residue was purified by Combiflash ISCO to obtain the pure product.

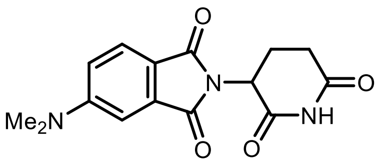

**5-(dimethylamino)-2-(2,6-dioxopiperidin-3-yl)isoindoline-1,3-dione (29)** was prepared by following the general procedure **B** using dimethylamine (0.26 mmol, 1.3 equiv.). The pure product was isolated by ISCO, and the column ran with $CH_2Cl_2$, then grading to 5% MeOH (contains 1% $NH_4OH$)-$CH_2Cl_2$ in 59% yield (35 mg; yellow solid). $^1$H and $^{13}$C NMR spectral data were matched with the literature reported values.[2]

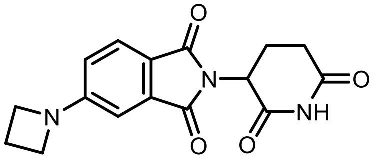

**5-(azetidin-1-yl)-2-(2,6-dioxopiperidin-3-yl)isoindoline-1,3-dione (30)** was prepared by following the general procedure **B** using azetidine. HCl (0.26 mmol). The pure product was isolated by ISCO and the column ran with $CH_2Cl_2$, then grading to 5% MeOH (contains 1% $NH_4OH$)-$CH_2Cl_2$ in in 77% yield (35 mg; yellow solid). $^1$H NMR (400 MHz, DMSO-$d_6$) δ 11.05 (s, 1H), 7.63 (d, $J$ = 8.3 Hz, 1H), 6.75 (d, $J$ = 2.1 Hz, 1H), 6.62 (dd, $J$ = 8.3, 2.1 Hz, 1H), 5.05 (dd, $J$ = 12.9, 5.4 Hz, 1H), 4.03 (t, $J$ = 7.4 Hz, 4H), 2.88 (ddd, $J$ = 17.4, 14.0, 5.5 Hz, 1H), 2.63 – 2.51 (m, 1H), 2.39 (p, $J$ = 7.4 Hz, 2H), 2.06 – 1.95 (m, 1H).$^{13}$C NMR (101 MHz, DMSO-$d_6$) δ 172.8, 170.1, 167.5, 167.2, 155.3, 133.8, 124.8, 116.6, 113.9, 104.2, 51.5, 48.7, 31.0, 22.2, 15.9. HRMS (ESI) calc'd for $C_{16}H_{15}N_3O_4$+ H = 314.1135, found 314.1134.

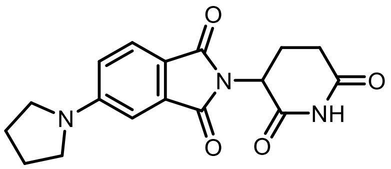

**2-(2,6-dioxopiperidin-3-yl)-5-(pyrrolidin-1-yl)isoindoline-1,3-dione (31)** was prepared by following the general procedure **B** using pyrrolidine (0.47 mmol, 1.3 equiv). The pure product was isolated by ISCO and the column ran with $CH_2Cl_2$, then grading to 5% MeOH (contains 1% $NH_4OH$)-$CH_2Cl_2$ in) in 76% yield (90 mg; yellow solid). $^1$H NMR (400 MHz, DMSO-$d_6$) δ 11.05 (s, 1H), 7.63 (d, $J$ = 8.4 Hz, 1H), 6.90 (d, $J$ = 2.2 Hz, 1H), 6.81 (dd, $J$ = 8.5, 2.3 Hz, 1H), 5.05 (dd, $J$ = 12.9, 5.4 Hz, 1H), 3.44 – 3.35 (m, 4H), 2.88 (ddd, $J$ = 17.2, 14.0, 5.5 Hz, 1H), 2.63 – 2.51 (m, 2H), 2.00 (dq, $J$ = 6.7, 4.5, 3.7 Hz, 5H). $^{13}$C NMR (101 MHz, DMSO-$d_6$) δ 172.8, 170.1, 167.7, 167.2, 151.8, 134.0, 124.9, 115.4, 115.3, 105.5, 48.7, 47.8, 39.5, 31.0, 24.9, 22.2. HRMS (ESI) calc'd for $C_{17}H_{17}N_3O_4$ + H = 328.1292, found 328.1289.

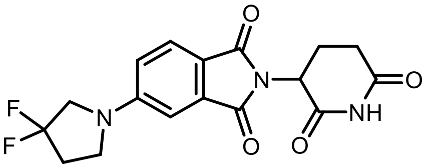

**5-(3,3-difluoropyrrolidin-1-yl)-2-(2,6-dioxopiperidin-3-yl)isoindoline-1,3-dione (32)** was prepared by following the general procedure **B** using 3,3-difluoropyrrolidine (0.26 mmol, 1.3 equiv). The pure product was isolated by ISCO and the column ran with $CH_2Cl_2$, then grading to 5% MeOH (contains 1% $NH_4OH$)-$CH_2Cl_2$ in in 40% yield (21 mg; yellow solid).$^1$H NMR (400 MHz, DMSO-$d_6$) δ 11.07 (s, 1H), 7.71 (d, $J$ = 8.4 Hz, 1H), 7.03 (d, $J$ = 2.3 Hz, 1H), 6.91 (dd, $J$ = 8.5, 2.3 Hz, 1H), 5.07 (dd, $J$ = 12.9, 5.4 Hz, 1H), 3.92 (t, $J$ = 13.1 Hz, 2H), 2.89 (ddd, $J$ = 17.3, 14.0, 5.5 Hz, 1H), 2.67 – 2.52 (m, 3H), 2.02 (dtd, $J$ = 10.3, 6.1, 2.8 Hz, 1H). $^{19}$F NMR (376 MHz, DMSO-d6) δ -99.48. $^{13}$C NMR (101 MHz, DMSO-$d_6$) δ 172.8, 170.1, 167.5, 167.2, 151.5, 133.9, 128.2 (t, $J$ = 246.1 Hz), 117.4, 115.8, 106.0, 54.9, 54.4 (t, $J$ = 31.6 Hz), 48.8, 45.6, 31.0, 22.2. HRMS (ESI) calc'd for $C_{17}H_{15}F_2N_3O_4$+ H = 364.1103, found 364.1100.

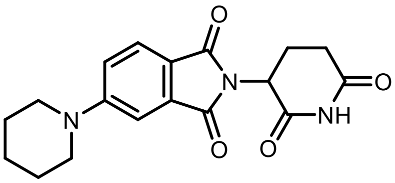

**2-(2,6-dioxopiperidin-3-yl)-5-(piperidin-1-yl)isoindoline-1,3-dione (33)** was prepared by following the general procedure **B** using piperidine (0.26 mmol, 1.3 equiv). The pure product was isolated by ISCO and the column ran with 0-10% MeOH (contains 1% $NH_4OH$)-$CH_2Cl_2$ 63% yield (39 mg; yellow solid). $^1$H NMR (400 MHz, DMSO-$d_6$) δ 11.06 (s, 1H), 7.64 (d, $J$ = 8.6 Hz, 1H), 7.29 (d, $J$ = 2.3 Hz, 1H), 7.21 (dd, $J$ = 8.7, 2.3 Hz, 1H), 5.06 (dd, $J$ = 12.9, 5.4 Hz, 1H), 3.47 (t, $J$ = 4.9 Hz, 4H), 2.88 (ddd, $J$ = 17.3, 14.0, 5.4 Hz, 1H), 2.69 – 2.51 (m, 2H), 2.08 – 1.91 (m, 1H), 1.58 (dd, $J$ = 9.3, 5.0 Hz, 6H). $^{13}$C NMR (101 MHz, DMSO-$d_6$) δ 172.8, 170.1, 167.6, 166.9, 155.0, 134.0, 125.0, 117.5, 117.2, 107.6, 48.7, 48.1, 39.5, 31.0, 24.7, 23.9, 22.2. HRMS (ESI) calc'd for $C_{18}H_{19}N_3O_4$ + H = 342.1448, found 342.1447.

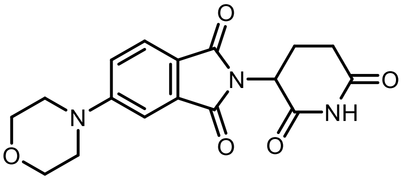

**2-(2,6-dioxopiperidin-3-yl)-5-morpholinoisoindoline-1,3-dione (34)** was prepared by following the general procedure **B** using morpholine (1.14 mmol, 1.3 equiv. The pure product was isolated by ISCO and the column ran with 0-10% MeOH (contains 1% $NH_4OH$)-$CH_2Cl_2$ in 68% yield (255 mg; yellow solid). $^1$H NMR (400 MHz, DMSO-$d_6$) δ 11.12 (s, 1H), 7.70 (d, $J$ = 8.5 Hz, 1H), 7.43 – 7.32 (m, 1H), 7.26 (dd, $J$ = 8.5, 2.3 Hz, 1H), 5.08 (dd, $J$ = 13.0, 5.4 Hz, 1H), 3.74 (t, $J$ = 4.7 Hz, 4H), 3.39 (t, $J$ = 5.3 Hz, 4H), 2.88 (ddd, $J$ = 18.8, 14.3, 5.4 Hz, 1H), 2.66 – 2.51 (m, 2H), 2.11 – 1.92 (m, 1H). $^{13}$C NMR (101 MHz, DMSO-$d_6$) δ 172.9, 170.2, 167.6, 167.0, 155.6, 133.8, 124.9, 118.9, 117.8, 108.0, 65.8, 48.8, 47.0, 39.5, 31.0, 22.2. HRMS (ESI) calc'd for $C_{17}H_{17}N_3O_5$ + H = 344.1241, found 344.1239.

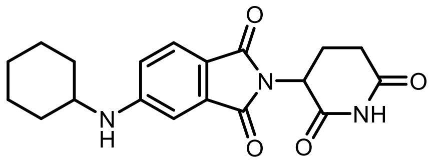

**5-(cyclohexylamino)-2-(2,6-dioxopiperidin-3-yl)isoindoline-1,3-dione (35)** was prepared by following the general procedure **B** using cyclohexylamine (0.26 mmol). The pure product was isolated by ISCO and the column ran with 0-10% MeOH (contains 1% $NH_4OH$)-$CH_2Cl_2$ in 31% yield (16 mg; yellow solid).$^1$H NMR (400 MHz, DMSO-$d_6$) δ 11.04 (s, 1H), 7.54 (d, $J$ = 8.4 Hz, 1H), 7.00 – 6.93 (m, 2H), 6.85 (dd, $J$ = 8.4, 2.1 Hz, 1H), 5.02 (dd, $J$ = 12.9, 5.3 Hz, 1H), 2.87 (ddd, $J$ = 17.3, 14.0, 5.4 Hz, 1H), 2.62 – 2.50 (m, 2H), 2.08 – 1.86 (m, 3H), 1.72 (d, $J$ = 13.0 Hz, 2H), 1.61 (d, $J$ = 12.7 Hz, 1H), 1.38 (q, $J$ = 12.3 Hz, 2H), 1.26 – 1.13 (m, 3H). $^{13}$C NMR (101 MHz, DMSO-$d_6$) δ 172.87, 170.2, 167.8, 167.2, 153.5, 134.3, 125.2, 115.6, 105.6, 105.6, 50.5, 48.6, 32.1, 31.0, 25.4, 24.3, 22.3. HRMS (ESI) calc'd for $C_{19}H_{21}N_3O_4$ + H = 356.1605, found 356.1602.

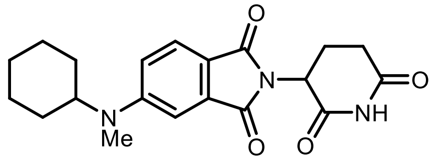

**5-(cyclohexyl(methyl)amino)-2-(2,6-dioxopiperidin-3-yl)isoindoline-1,3-dione (36)** was prepared by following the general procedure **B** using *N*-Methyl cyclohexylamine (0.26 mmol). The pure product was isolated by ISCO and the column ran with 0-10% MeOH (contains 1% $NH_4OH$)-$CH_2Cl_2$ in 48% yield (26 mg; yellow solid).$^1$H NMR (400 MHz, DMSO-$d_6$) δ 11.05 (s, 1H), 7.63 (d, $J$ = 8.6 Hz, 1H), 7.13 (d, $J$ = 2.4 Hz, 1H), 7.05 (dd, $J$ = 8.7, 2.4 Hz, 1H), 5.05 (dd, $J$ = 12.9, 5.4 Hz, 1H), 3.85 (tt, $J$ = 11.4, 3.7 Hz, 1H), 2.89 (s, 4H), 2.63 – 2.52 (m, 2H), 2.00 (dtd, $J$ = 10.2, 6.1, 2.8 Hz, 1H), 1.83 – 1.73 (m, 2H), 1.73 – 1.58 (m, 4H), 1.58 – 1.40 (m, 4H), 1.25 – 1.08 (m, 1H). $^{13}$C NMR (101 MHz, DMSO-$d_6$) δ 173.3, 170.6, 168.3, 167.6, 154.6, 134.6, 125.5, 116.3, 116.0, 106.2, 57.4, 49.2, 31.9, 31.5, 29.9, 25.6, 25.6, 22.7. HRMS (ESI) calc'd for $C_{20}H_{23}N_3O_4$ + H = 370.1761, found 370.1755.

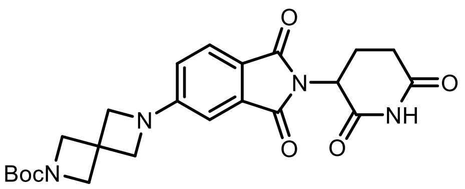

***tert*-butyl 6-(2-(2,6-dioxopiperidin-3-yl)-1,3-dioxoisoindolin-5-yl)-2,6-diazaspiro[3.3]heptane-2-carboxylate (63)** was prepared by following the general procedure **B** using Boc-2,6-diazaspiro[3,3]heptane hydrochloride (0.71 mmol, 1.3 equiv.) The pure product was isolated by ISCO and the column ran with $CH_2Cl_2$, then grading to 5% MeOH (contains 1% $NH_4OH$)-$CH_2Cl_2$ in 73% yield (180 mg; yellow solid). $^1$H NMR (400 MHz, DMSO-$d_6$) δ 11.05 (s, 1H), 7.64 (d, $J$ = 8.2 Hz, 1H), 6.78 (d, $J$ = 2.1 Hz, 1H), 6.65 (dd, $J$ = 8.4, 2.1 Hz, 1H), 5.05 (dd, $J$ = 12.9, 5.4 Hz, 1H), 4.17 (s, 4H), 4.05 (s, 4H), 2.87 (ddd, $J$ = 17.4, 14.0, 5.4 Hz, 1H), 2.62 – 2.50 (m, 4H), 2.06 – 1.95 (m, 1H), 1.38 (s, 9H). $^{13}$C NMR (101 MHz, DMSO-$d_6$) δ 172.7, 170.0, 167.4, 167.1, 155.3, 154.8, 133.7, 124.8, 117.2, 114.5, 104.7, 78.7, 61.2, 58.9, 48.7, 32.8, 30.9, 28.0, 22.2. HRMS (ESI) calc'd for $C_{23}H_{26}N_4O_6$ + H = 455.1925, found 455.1918.

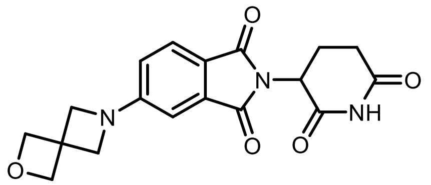

**2-(2,6-dioxopiperidin-3-yl)-5-(2-oxa-6-azaspiro[3.3]heptan-6-yl)isoindoline-1,3-dione (38)** was prepared by following the general procedure **B** using 2-Oxa-6-azaspiro[3.3]heptane (0.26 mmol). The pure product was isolated by ISCO and the column ran with $CH_2Cl_2$, then grading to 5% MeOH (contains 1% $NH_4OH$)-$CH_2Cl_2$ in 62% yield (24 mg; yellow solid). $^1$H NMR (400 MHz, DMSO-$d_6$) δ 11.05 (s, 1H), 7.64 (d, $J$ = 8.2 Hz, 1H), 6.79 (d, $J$ = 2.1 Hz, 1H), 6.66 (dd, $J$ = 8.3, 2.1 Hz, 1H), 5.05 (dd, $J$ = 12.9, 5.4 Hz, 1H), 4.74 (s, 4H), 4.21 (s, 4H), 2.87 (ddd, $J$ = 17.4, 14.1, 5.4 Hz, 1H), 2.66 – 2.50 (m, 3H), 2.12 – 1.94 (m, 1H). $^{13}$C NMR (101 MHz, DMSO-$d_6$) δ 172.7, 170.0,

167.4, 167.1, 154.9, 133.7, 124.8, 117.2, 114.5, 104.7, 79.7, 60.6, 48.7, 38.3, 31.0, 22.2. HRMS (ESI) calc'd for $C_{18}H_{17}N_3O_5$ + H = 356.1241, found 356.1240.

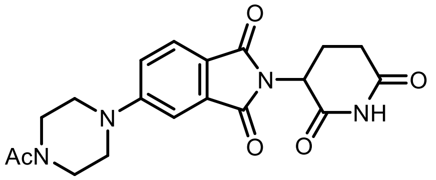

**5-(4-acetylpiperazin-1-yl)-2-(2,6-dioxopiperidin-3-yl)isoindoline-1,3-dione (39)** was prepared by following the general procedure **B** using 1-(piperazin-1-yl)ethan-1-one (0.26 mmol). The pure product was isolated by ISCO and the column ran with 0-10% MeOH (contains 1% $NH_4OH$)-$CH_2Cl_2$ in 76% yield (58 mg; yellow solid). $^1$H NMR (400 MHz, DMSO-$d_6$) δ 11.10 (s, 1H), 7.69 (d, *J* = 8.5 Hz, 1H), 7.34 (d, *J* = 2.3 Hz, 1H), 7.24 (dd, *J* = 8.6, 2.4 Hz, 1H), 5.07 (dd, *J* = 12.9, 5.4 Hz, 1H), 3.64 – 3.42 (m, 8H), 2.88 (ddd, *J* = 17.4, 14.1, 5.4 Hz, 1H), 2.62 – 2.52 (m, 2H), 2.04 (s, 3H). $^{13}$C NMR (101 MHz, DMSO-$d_6$) δ 172.9, 170.2, 168.6, 167.6, 167.1, 154.9, 133.9, 125.0, 118.5, 117.9, 108.0, 48.8, 46.8, 46.6, 44.9, 40.3, 31.1, 22.3, 21.1. HRMS (ESI) calc'd for $C_{19}H_{20}N_4O_5$ + H = 385.1506, found 385.1498.

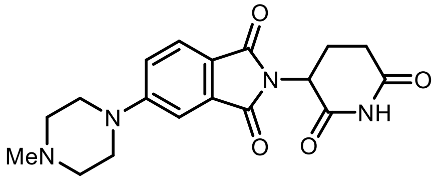

**2-(2,6-dioxopiperidin-3-yl)-5-(4-methylpiperazin-1-yl)isoindoline-1,3-dione (40)** was prepared by following the general procedure **B** using 1-(piperazin-1-yl)ethan-1-one (0.26 mmol). The pure product was isolated by ISCO and the column ran with 0-10% MeOH (contains 1% $NH_4OH$)-$CH_2Cl_2$ in 61% yield (43 mg; yellow solid). $^1$H NMR (400 MHz, DMSO-$d_6$) δ 11.08 (s, 1H), 7.66 (dd, *J* = 8.5, 1.6 Hz, 1H), 7.33 (t, *J* = 2.3 Hz, 1H), 7.24 (dd, *J* = 8.6, 2.4 Hz, 1H), 5.07 (dd, *J* = 12.9, 5.4 Hz, 1H), 3.43 (t, *J* = 5.0 Hz, 4H), 2.88 (ddd, *J* = 17.4, 14.0, 5.5 Hz, 1H), 2.73 – 2.53 (m, 2H), 2.43 (t, *J* = 5.0 Hz, 4H), 2.22 (s, 3H), 2.02 (dtd, *J* = 10.0, 5.8, 2.6 Hz, 1H). $^{13}$C NMR (101 MHz, DMSO-$d_6$) δ 172.8, 170.0, 167.5, 166.9, 155.2, 133.8, 124.8, 118.3, 117.8, 107.9, 54.1, 48.8, 46.8, 45.6, 31.0, 22.2. HRMS (ESI) calc'd for $C_{18}H_{20}N_4O_4$ + H = 357.1557, found 357.1552.

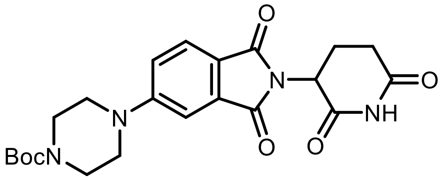

***tert*-butyl 4-(2-(2,6-dioxopiperidin-3-yl)-1,3-dioxoisoindolin-5-yl)piperazine-1-carboxylate (41)** was prepared by following the general procedure **B** using 1-Boc-piperazine (0.26 mmol). The pure product was isolated by ISCO and the column ran with $CH_2Cl_2$, then grading to 5% MeOH (contains 1% $NH_4OH$)-$CH_2Cl_2$ in 70% yield (65 mg; yellow solid).$^1$H NMR (400 MHz, DMSO-$d_6$) δ 11.07 (s, 1H), 7.69 (d, *J* = 8.5 Hz, 1H), 7.34 (d, *J* = 2.3 Hz, 1H), 7.24 (dd, *J* = 8.6, 2.3 Hz, 1H), 5.08 (dd, *J* = 12.9, 5.4 Hz, 1H), 3.47 (s, 8H), 2.89 (ddd, *J* = 17.3, 14.0, 5.5 Hz, 1H), 2.65 – 2.46 (m, 2H), 2.02 (ddd, *J* = 12.4, 6.4, 4.1 Hz, 1H), 1.43 (s, 9H).$^{13}$C NMR (101 MHz, DMSO-$d_6$) δ 172.8, 170.0, 167.5, 166.9, 155.0, 153.8, 133.8, 124.9, 118.5, 117.9, 108.0, 79.1, 48.8, 46.6, 31.0, 28.0, 22.2. HRMS (ESI) calc'd for [$C_{22}H_{26}N_4O_6$ + $NH_4^+$] = 460.2191, found 460.2188.

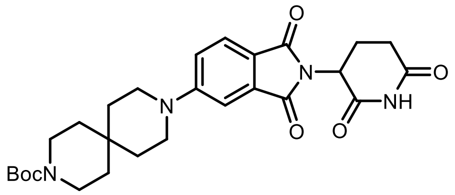

***tert*-butyl 9-(2-(2,6-dioxopiperidin-3-yl)-1,3-dioxoisoindolin-5-yl)-3,9-diazaspiro[5.5]undecane-3-carboxylate (42)** was prepared by following the general procedure **B** using 3,9-diazaspiro[5.5]undecane-3-carboxylate (0.26 mmol). The pure product was isolated by ISCO and the column ran with 0-10% MeOH (contains1% $NH_4OH$)-$CH_2Cl_2$ in 63% yield (64 mg; yellow solid). $^1$H NMR (400 MHz, DMSO-$d_6$) δ 11.05 (s, 1H), 7.64 (d, *J* = 8.6 Hz, 1H), 7.29 (d, *J* = 2.3 Hz, 1H), 7.21 (dd, *J* = 8.7, 2.4 Hz, 1H), 5.06 (dd, *J* = 12.8, 5.4 Hz, 1H), 3.50 – 3.42 (m, 4H), 3.32 (s, 4H), 2.88 (ddd, *J* = 17.3, 14.0, 5.5 Hz, 1H), 2.65 – 2.46 (m, 2H), 2.01 (dtd, *J* = 10.3, 6.1, 2.8 Hz, 1H), 1.58 – 1.50 (m, 4H), 1.41 (m, 4H), 1.39 (s, 9H). $^{13}$C NMR (101 MHz, DMSO-$d_6$) δ 172.7, 170.5, 167.1, 166.4, 154.9, 154.0, 134.0, 124.9, 117.3, 107.4, 78.2, 48.3, 42.8, 34.7, 34.0, 31.0, 29.6, 28.1, 22.2. HRMS (ESI) calc'd for $C_{27}H_{34}N_4O_6$ + H = 511.2551, found 511.2553.

**General Procedure C**: Preparation of 6-Fluoro substituted IMiD analogs through $S_NAr$

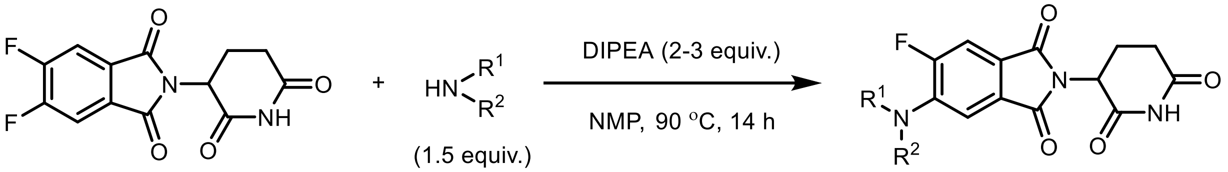

In an oven-dried 5 mL vial with a stir-bar, DIPEA (0.34 mmol, 2.5 equiv.) was added to a solution of 2-(2,6-dioxopiperidin-3-yl)-5,6-difluoroisoindoline-1,3-dione (40 mg, 0.14 mmol), cyclic amines (0.20 mmol, 1.5 equiv.) in dry *N*-methyl pyrrolidine (1.5 mL) at rt. The reaction mixture was stirred at 90 °C for 14 h. The reaction mixture was diluted with ethyl acetate (50 mL), washed with water (10 mL x3), followed by Brine (10 mL), dried over anhydrous $Na_2SO_4$, filtered, and concentrated under reduced pressure. The crude residue was purified by Combiflash ISCO to obtain the pure product.

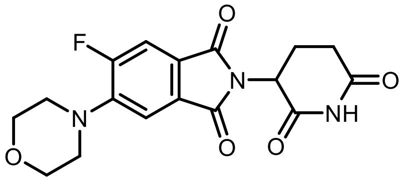

**2-(2,6-dioxopiperidin-3-yl)-5-fluoro-6-morpholinoisoindoline-1,3-dione dione (43)** was prepared by following the general procedure **C** using morpholine (0.20 mmol). The pure product was isolated by ISCO and the column ran with 0-10% MeOH (contains1% $NH_4OH$)-$CH_2Cl_2$ in 39% yield (20 mg; yellow solid). $^1$H NMR (400 MHz, DMSO-$d_6$) δ 11.12 (s, 1H), 7.76 (d, *J* = 11.5 Hz, 1H), 7.48 (d, *J* = 7.4 Hz, 1H), 5.11 (dd, *J* = 12.8, 5.4 Hz, 1H), 3.79 – 3.72 (m, 4H), 3.26 – 3.19 (m, 4H), 2.88 (ddd, *J* = 16.7, 13.7, 5.3 Hz, 1H), 2.63 – 2.50 (m, 2H), 2.03 (ddd, *J* = 10.6, 5.7, 3.4 Hz, 1H). $^{19}$F NMR (376 MHz, DMSO-$d_6$) δ -112.13.$^{13}$C NMR (101 MHz, DMSO-$d_6$) δ 172.7, 169.9, 166.6, 166.16, 166.1 (d, *J* = 2.4 Hz), 157.4 (d, *J* = 253.5 Hz), , 145.3 (d, *J* = 8.7 Hz),, 128.7 (d, *J* = 2.4 Hz), , 123.7 (d, *J* = 9.9 Hz), , 113.6 (d, *J* = 4.7 Hz), 112.0 (d, *J* = 25.2 Hz), 65.9, 49.8, 49.8, 49.1, 30.9, 22.1. HRMS (ESI) calc'd for $C_{17}H_{16}FN_3O_5$ + H = 362.1147, found 362.1142.

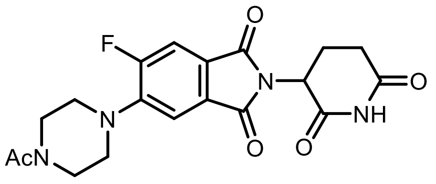

**5-(4-acetylpiperazin-1-yl)-2-(2,6-dioxopiperidin-3-yl)-6-fluoroisoindoline-1,3-dione (44)** was prepared by following the general procedure **C** using 1-Acetylpiperazine (0.20 mmol). The pure product was isolated by ISCO and the column ran with 0-10% MeOH (contains1% $NH_4OH$)-$CH_2Cl_2$ in 36% yield (20 mg; yellow solid). $^{1}$H NMR (400 MHz, DMSO-$d_6$) δ 11.10 (s, 1H), 7.76 (d, *J* = 11.3 Hz, 1H), 7.48 (d, *J* = 7.3 Hz, 1H), 5.11 (dd, *J* = 12.8, 5.4 Hz, 1H), 3.65 – 3.57 (m, 4H), 3.31 – 3.17 (m, 4H), 2.95 – 2.82 (m,m1H), 2.64 – 2.51 (m, 1H), 2.05 (s, 3H). $^{19}$F NMR (376 MHz, DMSO-$d_6$) δ –112.15.$^{13}$C NMR (101 MHz, DMSO-$d_6$) δ 173.2,170.3, 168.9, 167.1, 166.6 (d, *J* = 2.4 Hz), 159.1, 156.6, 145. 6 (d, *J* = 8.9 Hz), 129.2 (d, *J* = 2.5 Hz), 124.2 (d, *J* = 9.8 Hz), 114.5 (d, *J* = 4.8 Hz), 112.5 (d, *J* = 25.1 Hz), 55.4, 49.6, 45.8, 41.0, 31.4, 22.5, 21.6. HRMS (ESI) calc'd for $C_{19}H_{19}FN_4O_5$ + H = 403.1412, found 403.1407.

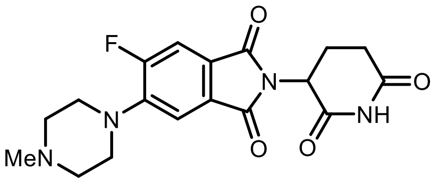

**2-(2,6-dioxopiperidin-3-yl)-5-fluoro-6-(4-methylpiperazin-1-yl)isoindoline-1,3-dione (45)** was prepared by following the general procedure **C** using 1-methylpiperazine (0.20 mmol). The pure product was isolated by ISCO and the column ran with 0-10% MeOH (contains1% $NH_4OH$)-$CH_2Cl_2$ in 37% yield (19 mg; yellow solid). $^{1}$H NMR (400 MHz, DMSO-$d_6$) δ 11.12 (s, 1H), 7.73 (d, *J* = 11.5 Hz, 1H), 7.46 (d, *J* = 7.5 Hz, 1H), 5.11 (dd, *J* = 12.8, 5.4 Hz, 1H), 3.24 (t, *J* = 4.9 Hz, 4H), 2.88 (ddd, *J* = 16.7, 13.7, 5.3 Hz, 1H), 2.64 – 2.51 (m, 2H), 2.23 (s, 3H), 2.03 (ddd, *J* = 13.1, 5.8, 3.4 Hz, 1H). $^{19}$F NMR (376 MHz, DMSO-$d_6$) δ -111.95.$^{13}$C NMR (101 MHz, DMSO-$d_6$) δ 172.8, 170.0, 166.7, 166.2 (d, *J* = 2.4 Hz) 157.4 (d, *J* = 253.3 Hz), 145.4 (d, *J* = 8.9 Hz), 128.8 (d, *J* = 2.5 Hz), 123.4 (d, *J* = 9.9 Hz), 113.8 (d, *J* = 4.9 Hz), 111.9 (d, *J* = 25.3 Hz), 54.3, 49.4 (d, J = 4.6 Hz), 49.1, 45.7, 31.0, 22.1. HRMS (ESI) calc'd for $C_{18}H_{19}FN_4O_4$ + H = 375.1463, found 375.1457.

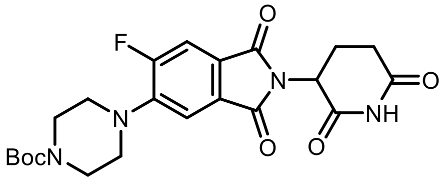

***tert*-butyl 4-(2-(2,6-dioxopiperidin-3-yl)-6-fluoro-1,3-dioxoisoindolin-5-yl)piperazine-1-carboxylate (46)** was prepared by following the general procedure **C** using 1-Boc piperazine (0.20 mmol). The pure product was isolated by ISCO and the column ran with 0-10% MeOH (contains1% $NH_4OH$)-$CH_2Cl_2$ in 31% yield (20 mg; yellow solid). $^{1}$H NMR (400 MHz, DMSO-$d_6$) δ 11.10 (s, 1H), 7.74 (d, *J* = 11.3 Hz, 1H), 7.48 (d, *J* = 7.4 Hz, 1H), 5.11 (dd, *J* = 12.9, 5.4 Hz, 1H), 3.50 (t, *J* = 5.0 Hz, 4H), 3.21 (dd, *J* = 6.3, 3.8 Hz, 4H), 2.89 (ddd, *J* = 17.1, 13.8, 5.4 Hz, 1H), 2.66 – 2.50 (m, 1H), 2.04 (ddd, *J* = 10.5, 5.5, 3.2 Hz, 1H), 1.43 (s, 9H). $^{19}$F NMR (376 MHz, DMSO-$d_6$) δ -112.03. $^{13}$C NMR (101 MHz, DMSO-$d_6$) δ 173.2, 170.3, 167.05, 166.6 (d, *J* = 2.4 Hz), 159.1, 157.8 (d, *J* = 253.5 Hz), 154.3, 145.7 (d, *J* = 9.0 Hz), 124.3 (d, *J* = 9.6 Hz), 114.5 (d, *J* = 4.8 Hz), 112.4 (d, *J* = 25.1 Hz), 79.6, 49.9, 49.8, 49.6, 31.4, 28.5, 22.5. HRMS (ESI) calc'd for $C_{18}H_{19}FN_4O_4$ + H = 461.1831, found 461.1822.

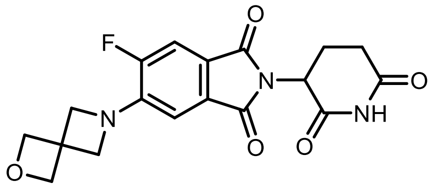

**2-(2,6-dioxopiperidin-3-yl)-5-fluoro-6-(2-oxa-6-azaspiro[3.3]heptan-6-yl)isoindoline-1,3-dione (47)** was prepared by following the general procedure **C** using 1 using 2-Oxa-6-azaspiro[3.3]heptane (0.20 mmol). The pure product was isolated by ISCO and the column ran with 0-10% MeOH (contains1% $NH_4OH$)-$CH_2Cl_2$ in 47% yield (24 mg; yellow solid). $^1$H NMR (400 MHz, DMSO-$d_6$) δ 11.10 (s, 1H), 7.61 (d, *J* = 11.2 Hz, 1H), 6.93 (d, *J* = 7.7 Hz, 1H), 5.07 (dd, *J* = 12.8, 5.4 Hz, 1H), 4.73 (s, 5H), 4.34 (d, *J* = 2.4 Hz, 4H), 2.88 (ddd, *J* = 16.6, 13.7, 5.3 Hz, 1H), 2.63 – 2.52 (m, 2H), 2.08 – 1.96 (m, 1H). $^{19}$F NMR (376 MHz, DMSO-$d_6$) δ -127.64. $^{13}$C NMR (101 MHz, DMSO-$d_6$) δ 173.3, 170.5, 167.3, 166.9 (d, *J* = 2.9 Hz), 154.3 (d, *J* = 247.6 Hz), 144.3 (d, *J* = 12.1 Hz), 129.7 (d, *J* = 1.9 Hz), 119.3 (d, *J* = 8.4 Hz), 111.8 (d, *J* = 22.2 Hz), 108.7 (d, *J* = 6.8 Hz), 80.1, 62.9, 48.9, 39.4, 31.4, 22.6. HRMS (ESI) calc'd for $C_{18}H_{16}FN_3O_5$ + H = 374.1147, found 374.1140.

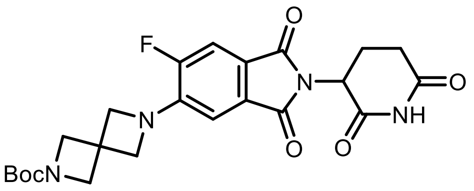

***tert*-butyl 6-(2-(2,6-dioxopiperidin-3-yl)-6-fluoro-1,3-dioxoisoindolin-5-yl)-2,6-diazaspiro[3.3]heptane-2-carboxylate (48)** was prepared by following the general procedure **C** using Boc-2,6-diazaspiro[3,3]heptane hydrochloride (0.66 mmol). The pure product was isolated by ISCO and the column ran with 0-10% MeOH (contains 1% $NH_4OH$)-$CH_2Cl_2$ in 78% yield (150 mg; yellow solid). $^1$H NMR (400 MHz, DMSO-$d_6$) δ 11.07 (s, 1H), 7.60 (d, *J* = 11.1 Hz, 1H), 6.90 (d, *J* = 7.7 Hz, 1H), 5.06 (dd, *J* = 12.8, 5.4 Hz, 1H), 4.29 (d, *J* = 2.3 Hz, 4H), 4.04 (s, 4H), 2.87 (ddd, *J* = 16.5, 13.7, 5.4 Hz, 1H), 2.63 – 2.51 (m, 2H), 2.08 – 1.96 (m, 1H), 1.38 (s, 9H). $^{19}$F NMR (376 MHz, DMSO-$d_6$) δ -127.55. $^{13}$C NMR (101 MHz, DMSO-$d_6$) δ 172.7, 169.9, 166.7, 166.4 (d, *J* = 2.6 Hz), 155.31, 153.8 (d, *J* = 247.9 Hz), 143.7 (d, *J* = 12.1 Hz), 129.2 (d, *J* = 2.0 Hz), 118.8 (d, *J* = 8.3 Hz), 111.3 (d, *J* = 22.2 Hz), 108.2 (d, *J* = 6.7 Hz), 78.6, 63.0, 58.8, 48.9, 33.4 (d, *J* = 2.7 Hz), 30.9, 28.0, 22.1. HRMS (ESI) calc'd for $C_{23}H_{25}FN_4O_6$ + H = 473.1831, found 473.1824.

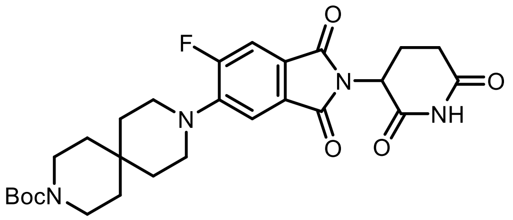

***tert*-butyl 9-(2-(2,6-dioxopiperidin-3-yl)-6-fluoro-1,3-dioxoisoindolin-5-yl)-3,9-diazaspiro[5.5]undecane-3-carboxylate (49)** was prepared by following the general procedure **C** using 3,9-diazaspiro[5.5]undecane-3-carboxylate (1.27 mmol) The pure product was isolated by ISCO and the column ran with 0-10% MeOH (contains 1% $NH_4OH$)-$CH_2Cl_2$ in 56% yield (150 mg; yellow solid). $^1$H NMR (400 MHz, DMSO-$d_6$) δ 11.12 (s, 1H), 7.70 (d, *J* = 11.5 Hz, 1H), 7.44 (d, *J* = 7.4 Hz, 1H), 5.10 (dd, *J* = 12.8, 5.4 Hz, 1H), 3.33 – 3.13 (m, 9H), 2.63 – 2.51 (m, 2H), 2.03 (ddd, *J* = 13.3, 5.8, 3.4 Hz, 1H), 1.60 (t, *J* = 5.7 Hz, 4H), 1.39 (s, 13H). $^{19}$F NMR (376 MHz, DMSO-$d_6$) δ -112.14. $^{13}$C NMR (101 MHz, DMSO-$d_6$) δ 172.8, 170.0, 166.7, 166.3 (d, *J* = 2.5 Hz), 157.2 (d, *J* = 252.9 Hz), 154.0, 145.8 (d, *J* = 8.8 Hz), 128.8 (d, *J* = 2.5 Hz), 122.8 (d, *J* = 9.7 Hz), 113.7 (d, *J* = 4.9 Hz), 111.9 (d, *J* = 25.4 Hz), 78.5, 55.0, 49.1, 45.5, 45.4, 34.6, 31.0, 29.3, 28.1, 22.1. HRMS (ESI) calc'd for $C_{27}H_{33}FN_4O_6$ + H = 529.2457, found 529.2443.

**General Procedure D**: Suzuki coupling of 4-bromothalidoimide with boronic acids

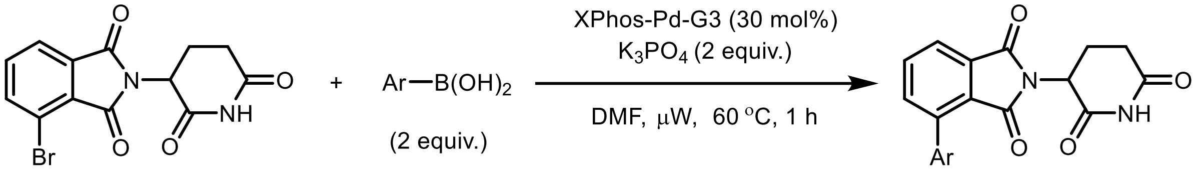

In an oven-dried 5 mL microwave vial with previously placed stir-bar was charged with 4-bromo-2-(2,6-dioxopiperidin-3-yl)isoindoline-1,3-dione (66 mg, 0.2 mmol), aryl boronic acid (0.4 mmol), XPhos-Pd-G3 (51 mg, 0.06 mmol) and dry $K_3PO_4$ (84 mg, 0.4 mmol). To this mixture, DMF (2 mL) and water (0.1 mL) were added, and the mixture was deaerated under Argon at rt for 5 min. The reaction vial was sealed with a microwave-supported Cap with Septum. The reaction mixture was stirred at 60 °C for 1 h in the microwave reactor. The solvent was evaporated under the reduced pressure, the residue was diluted with ethyl acetate (50 mL) and washed with water (10 mL x2), brine (20 mL), dried over anhydrous $Na_2SO_4$, filtered, and concentrated under reduced pressure. The crude residue was purified by Combiflash ISCO to obtain the pure product.

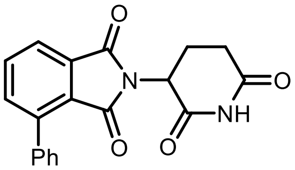

**2-(2,6-dioxopiperidin-3-yl)-4-phenylisoindoline-1,3-dione (50)** was prepared by following the general procedure **D** using phenylboronic acid (0.4 mmol). The final product was isolated by flash column chromatography ($CH_2Cl_2$: MeOH 85:15) in a 61% yield (41 mg; white solid). $^1$H and $^{13}$C NMR data were matched with the literature reported values.[3]

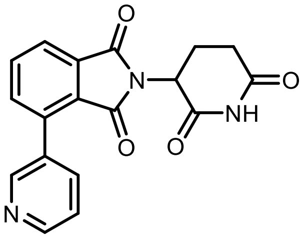

**2-(2,6-dioxopiperidin-3-yl)-4-(pyridin-3-yl)isoindoline-1,3-dione (51)** was prepared by following the general procedure **D** using pyridine-3-boronic acid (0.4 mmol). Final product was isolated by flash column chromatography ($CH_2Cl_2$: MeOH 95:5) in 67% yield (45 mg; white solid). $^1$H NMR (400 MHz, DMSO-$d_6$) δ 11.17 (s, 1H), 8.06 (d, $J$ = 8.0 Hz, 1H), 7.94 (d, $J$ = 7.3 Hz, 1H), 7.78 (t, $J$ = 7.7 Hz, 1H), 5.17 (dd, $J$ = 12.8, 5.4 Hz, 1H), 2.89 (ddd, $J$ = 17.0, 13.8, 5.4 Hz, 1H), 2.69 – 2.50 (m, 2H), 2.07 (dtt, $J$ = 11.0, 5.4, 2.2 Hz, 1H). $^{13}$C NMR (101 MHz, DMSO-$d_6$) δ 172.8,169.8, 165.6, 165.2, 139.2, 136.3, 133.7, 128.8, 122.9, 117.7, 49.2, 30.9, 21.9. HRMS (ESI) calc'd for $C_{18}H_{13}N_3O_4$ + H = 336.0979, found 336.0979.

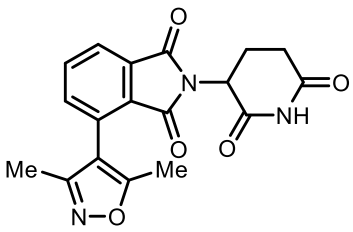

**4-(3,5-dimethylisoxazol-4-yl)-2-(2,6-dioxopiperidin-3-yl)isoindoline-1,3-dione (52)** was prepared by following the general procedure **D** using (3,5-dimethylisoxazol-4-yl)boronic acid (0.222 mmol, 1.5 equiv). Final product was isolated by flash column chromatography ($CH_2Cl_2$: MeOH 90:10) in 38% yield (20 mg; white solid). $^1$H NMR (400 MHz, DMSO-$d_6$) δ 11.10 (s, 1H), 8.02 – 7.91 (m, 2H), 7.83 – 7.76 (m, 1H), 5.13 (dd, $J$ = 12.8, 5.4 Hz, 1H), 2.88 (ddd, $J$ = 17.0, 13.8, 5.4 Hz, 1H), 2.64 – 2.51 (m, 2H), 2.10 (d, $J$ = 9.3 Hz, 3H), 2.07 – 2.00 (m, 1H). $^{13}$C NMR (101 MHz, DMSO- $d_6$) δ 173.2, 170.3, 167.3, 167.21, 167.1, 167.0, 167.0, 159.1, 159.0, 137.7, 137.7, 135.5, 135.5, 132.9, 128.9, 128.9, 128.1, 123.6, 123.6, 112.3, 112.2, 49.5, 31.4, 22.4, 12.0 11.8, 10.6. HRMS (ESI) calc'd for $C_{18}H_{15}N_3O_5$ + H = 354.1084, found 354.1081.

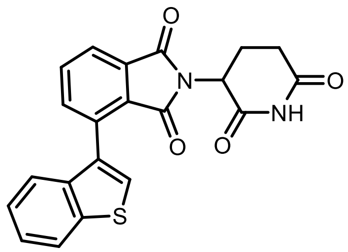

**4-(benzo[b]thiophen-3-yl)-2-(2,6-dioxopiperidin-3-yl)isoindoline-1,3-dione (53)** was prepared by following the general procedure **D** using thiophene-3-boronic acid (0.4 mmol). Final product was isolated by flash column chromatography ($CH_2Cl_2$: MeOH 95:5) in 72% yield (56 mg; white solid). $^1H$ NMR (400 MHz, DMSO-$d_6$) δ 11.06 (s, 1H), 8.13 – 8.08 (m, 1H), 8.02 – 7.95 (m, 3H), 7.90 (dd, $J$ = 6.9, 1.9 Hz, 1H), 7.49 – 7.29 (m, 2H), 5.09 (dd, $J$ = 12.8, 5.4 Hz, 1H), 2.85 (ddd, $J$ = 17.0, 13.8, 5.4 Hz, 1H), 2.61 – 2.36 (m, 3H), 2.05 (ddd, $J$ = 9.9, 6.3, 2.1 Hz, 1H). $^{13}C$ NMR (101 MHz, DMSO-$d_6$) δ 172.7, 169.7, 166.7, 166.2, 139.1, 137.7, 136.8, 134.8, 133.2, 132.5, 131.3, 128.2, 127.8, 124.5, 123.0, 122.8, 122.4, 48.9, 30.9, 21.9. HRMS (ESI) calc'd for $C_{21}H_{14}N_2O_4S$ + H = 391.0747, found 391.0743.

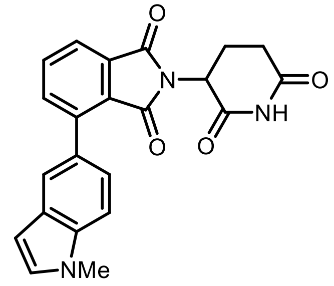

**2-(2,6-dioxopiperidin-3-yl)-4-(1-methyl-1H-indol-5-yl)isoindoline-1,3-dione (54)** was prepared by following the general procedure **D** 1-Methylindole-5-boronic acid pinacol ester (0.222 mmol, 1.5 equiv). Final product was isolated by flash column chromatography ($CH_2Cl_2$: MeOH 90:10) in 30% yield (22 mg; yellow solid).$^1H$ NMR (400 MHz, DMSO-$d_6$) δ 11.08 (s, 1H), 7.94 – 7.81 (m, 3H), 7.78 (d, $J$ = 1.6 Hz, 1H), 7.51 (d, $J$ = 8.5 Hz, 1H), 7.41 – 7.33 (m, 2H), 6.50 (dd, $J$ = 3.1, 0.7 Hz, 1H), 5.11 (dd, $J$ = 12.9, 5.4 Hz, 1H), 3.83 (s, 3H), 2.87 (ddd, $J$ = 17.4, 14.0, 5.4 Hz, 1H), 2.64 – 2.51 (m, 2H), 2.06 (dtd, $J$ = 12.6, 5.6, 2.6 Hz, 1H).$^{13}C$ NMR (101 MHz, DMSO-$d_6$) δ 172.8, 169.9, 166.9, 166.7, 142.1, 136.9, 136.4, 134.4, 132.6, 130.5, 127.7, 126.7, 126.4, 122.9, 121.6, 121.4, 109.2, 101.0, 48.9, 32.6, 30.9, 22.0. HRMS (ESI) calc'd for $C_{22}H_{17}N_3O_4$ + H = 388.1292, found 388.1286.

**General Procedure E**: Suzuki coupling of 5-bromothalidoimide with boronic acids

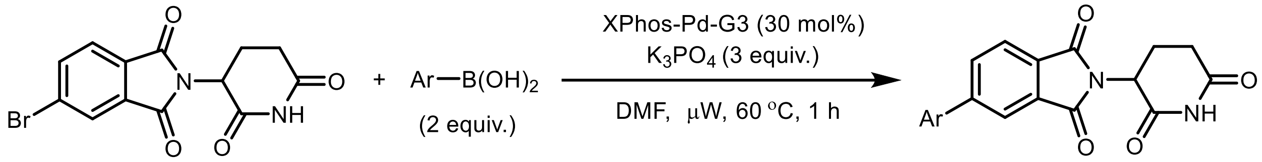

In an oven-dried 5 mL microwave vial with a stir-bar was charged with 5-bromo-2-(2,6-dioxopiperidin-3-yl)isoindoline-1,3-dione (66 mg, 0.2 mmol), aryl boronic acid (0.4 mmol), XPhos-Pd-G3 (51 mg, 0.06 mmol) and dry $K_3PO_4$ (84 mg, 0.4 mmol). DMF (2 mL) and water (0.1 mL) were added to this mixture, and the mixture was deaerated under Argon at rt for 5 min. The reaction vial was sealed with a microwave-supported Cap with Septum. The reaction mixture was stirred at 60 °C for 1 h in the microwave reactor. The solvent was evaporated under the reduced pressure, the residue was diluted with ethyl acetate (50 mL) and washed with water (10 mL x2), brine (20 mL), dried over anhydrous $Na_2SO_4$, filtered, and concentrated under reduced pressure. The crude residue was purified by Combiflash ISCO to obtain the pure product.

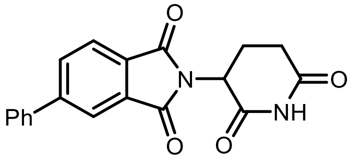

**2-(2,6-dioxopiperidin-3-yl)-5-phenylisoindoline-1,3-dione (56)** was prepared by following the general procedure **E** using phenylboronic acid (0.4 mmol). The pure product was isolated by ISCO and the column ran with $CH_2Cl_2$, then grading to 15%MeOH (contains 1% $NH_4OH$)-$CH_2Cl_2$ in 67% yield (45 mg; white solid). $^1$H and $^{13}$C NMR data were matched with the literature reported values.[3]

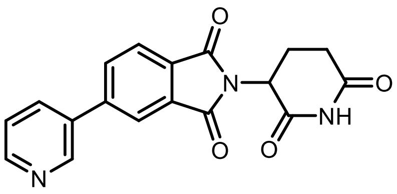

**2-(2,6-dioxopiperidin-3-yl)-5-(pyridin-3-yl)isoindoline-1,3-dione (57)** was prepared by following the general procedure **E** using pyridine-3-boronic acid (0.4 mmol). The pure product was isolated by ISCO and the column ran with $CH_2Cl_2$, then grading to 5%MeOH (contains 1% $NH_4OH$)-$CH_2Cl_2$ in 70% yield (35 mg, Yellow solid). $^1$H NMR (400 MHz, DMSO-$d_6$) δ 11.13 (s, 1H), 9.06 (d, $J$ = 2.3 Hz, 1H), 8.68 (dd, $J$ = 4.8, 1.5 Hz, 1H), 8.38 – 8.17 (m, 3H), 8.04 (d, $J$ = 7.8 Hz, 1H), 7.56 (dd, $J$ = 8.0, 4.8 Hz, 1H), 5.20 (dd, $J$ = 12.9, 5.4 Hz, 1H), 3.03 – 2.82 (m, 1H), 2.73 – 2.55 (m, 2H), and 2.10 ppm (ddd, $J$ = 9.4, 4.9, 2.7 Hz, 1H). $^{13}$C NMR (101 MHz, DMSO-$d_6$) δ 172.7, 169.7, 166.8, 149.8, 148.1, 143.6, 134.9, 133.7, 133.2, 132.3, 130.4, 124.1, 124.0, 121.8, 49.1, 30.9, 22.0. HRMS (ESI) calc'd for $C_{18}H_{13}N_3O_4$ + H = 336.0979, found 336.0976.

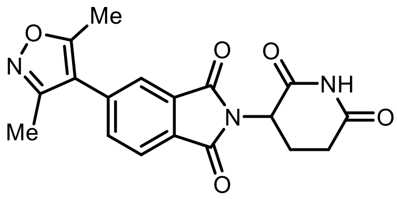

**5-(3,5-dimethylisoxazol-4-yl)-2-(2,6-dioxopiperidin-3-yl)isoindoline-1,3-dione (58)** was prepared by following the general procedure **E** using (3,5-dimethylisoxazol-4-yl)boronic acid (0.222 mmol, 1.5 equiv). The pure product was isolated by ISCO and the column ran with $CH_2Cl_2$, then grading to 10% MeOH (contains 1% $NH_4OH$)-$CH_2Cl_2$ in 38% yield (20 mg; white solid). $^1$H NMR (400 MHz, DMSO-$d_6$) δ 11.13 (s, 1H), 8.01 (dd, $J$ = 7.7, 0.7 Hz, 1H), 7.95 (dd, $J$ = 1.5, 0.7 Hz, 1H), 7.89 (dd, $J$ = 7.7, 1.5 Hz, 1H), 5.19 (dd, $J$ = 12.9, 5.4 Hz, 1H), 2.91 (ddd, $J$ = 17.3, 13.9, 5.4 Hz, 1H), 2.68 – 2.53 (m, 2H), 2.46 (s, 3H), 2.28 (s, 3H), 2.09 (ddt, $J$ = 10.6, 5.5, 3.2 Hz, 1H). $^{13}$C NMR (101 MHz, DMSO-$d_6$) δ 172.7, 169.8, 166.8, 166.8, 166.5, 158.0, 136.8, 135.1, 132.1, 130.0, 123.9, 123.6, 114.9, 49.1, 39.5, 30.9, 21.9, 11.4, 10.3. HRMS (ESI) calc'd for $C_{18}H_{15}N_3O_5$ + H = 354.1084, found 354.1083.

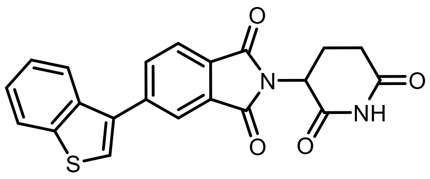

**5-(benzo[b]thiophen-3-yl)-2-(2,6-dioxopiperidin-3-yl)isoindoline-1,3-dione (59)** was prepared by following the general procedure **E** using thiophene-3-boronic acid (0.4 mmol). Final product was isolated by flash column chromatography ($CH_2Cl_2$: MeOH 95:5) in 76% yield (59 mg; white solid). $^1$H NMR (400 MHz, DMSO-$d_6$) δ 11.18 (s, 1H), 8.17 (s, 1H), 8.14 – 8.08 (m, 3H), 8.06 (d, $J$ = 7.7 Hz, 1H), 7.95 – 7.91 (m, 1H), 7.48 (td, $J$ = 7.5, 6.8, 3.7 Hz, 2H), 5.22 (dd, $J$ = 12.9, 5.4 Hz, 1H), 2.92 (ddd, $J$ = 17.1, 13.8, 5.4 Hz, 1H), 2.70 – 2.54 (m, 2H), 2.11 (dtd, $J$ = 10.8, 5.8, 5.4, 2.9 Hz, 1H). $^{13}$C NMR (101 MHz, DMSO-$d_6$) δ 172.9, 169.9, 167.0, 167.0, 141.6, 140.2, 136.5, 135.0, 134.4, 132.2, 129.9, 127.7, 125.2, 125.0, 124.2, 123.5, 123.0, 122.1, 49.1, 31.0, 22.1. HRMS (ESI) calc'd for $C_{21}H_{14}N_2O_4S$ + H = 391.0747, found 391.0743.

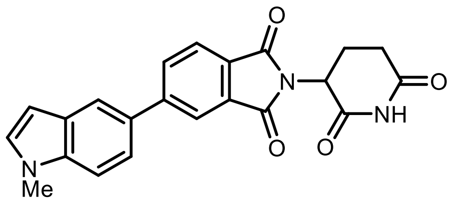

**2-(2,6-dioxopiperidin-3-yl)-5-(1-methyl-1H-indol-5-yl)isoindoline-1,3-dione (60)** was prepared by following the general procedure **G** using 1-Methylindole-5-boronic acid pinacol ester (0.222 mmol, 2.0 equiv). The pure product was isolated by ISCO and the column ran with $CH_2Cl_2$, then grading to 5% MeOH (contains 1% $NH_4OH$)-$CH_2Cl_2$ in in 30% yield (22 mg; yellow solid).[1]H NMR (400 MHz, DMSO-$d_6$) δ 11.13 (s, 1H), 8.23 – 8.14 (m, 1H), 8.05 (d, $J$ = 1.7 Hz, 1H), 7.96 (d, $J$ = 7.8 Hz, 1H), 7.68 – 7.55 (m, 1H), 7.41 (d, $J$ = 3.1 Hz, 1H), 6.54 (d, $J$ = 3.0 Hz, 1H), 5.18 (dd, $J$ = 12.8, 5.4 Hz, 1H), 3.84 (s, 3H), 2.92 (ddd, $J$ = 17.1, 13.8, 5.4 Hz, 1H), 2.68 – 2.51 (m, 2H), 2.09 (ddq, $J$ = 10.7, 5.7, 2.9 Hz, 1H). $^{13}$C NMR (101 MHz, DMSO-$d_6$) δ 172.8, 169.9, 167.2, 167.1, 148.4, 136.8, 132.5, 132.3, 131.0, 129.2, 128.7, 128.6, 124.0, 121.1, 120.6, 119.6, 110.5, 101.3, 49.0, 32.6, 31.0, 22.1. HRMS (ESI) calc'd for $C_{22}H_{17}N_3O_4$ + H = 388.1292, found 388.1290.

**General Procedure F**: Sonogashira coupling of 4-bromothalidomide with phenylacetylene

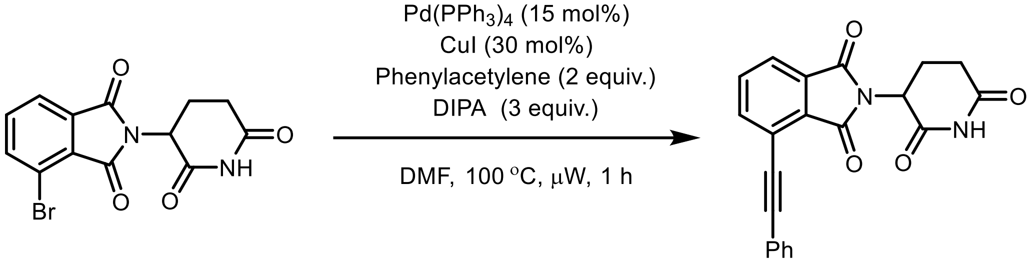

In an oven-dried 5 mL microwave reaction vial with previously placed stir-bar was charged with 4-bromo-2-(2,6-dioxopiperidin-3-yl)isoindoline-1,3-dione (66 mg, 0.2 mmol), $Pd(PPh_3)_4$ (35 mg, 0.03 mmol), CuI (12 mg, 0.06 mmol). To this mixture, phenylacetylene (0.4 mmol), DIPA (0.6 mmol), and DMF (2 mL) were added and deaerated under Argon at rt for 5 min. The reaction vial was sealed with a microwave-supported Cap with Septum. The reaction mixture was stirred at 100 °C for 1 h in the microwave reactor. The solvent was evaporated under the reduced pressure, the residue was diluted with ethyl acetate (50 mL) and washed with water (10 mL x2), brine (20 mL), dried over anhydrous $Na_2SO_4$, filtered, and concentrated under reduced pressure. The crude residue was purified by Combiflash ISCO to obtain the pure product.

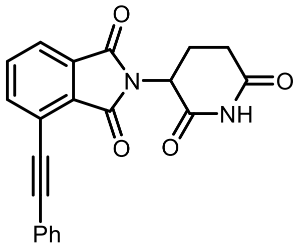

**2-(2,6-dioxopiperidin-3-yl)-4-(phenylethynyl)isoindoline-1,3-dione (54)** was prepared by following the general procedure **F**. The pure product was isolated by ISCO and the column ran with $CH_2Cl_2$, then grading to 5%MeOH (contains 1% $NH_4OH$)-$CH_2Cl_2$ in 81% yield (58 mg; white solid). $^{1}$H and $^{13}$C NMR data was matched with the literature reported values. $^{1}$H NMR (400 MHz, DMSO-$d_6$) δ 11.15 (s, 1H), 8.05 – 7.77 (m, 3H), 7.64 (dd, $J$ = 6.7, 3.0 Hz, 2H), 7.48 (p, $J$ = 3.8 Hz, 3H), 5.19 (dd, $J$ = 12.9, 5.4 Hz, 1H), 2.91 (ddd, $J$ = 17.2, 13.9, 5.5 Hz, 1H), 2.73 – 2.42 (m, 2H), 2.28 – 1.93 (m, 1H). $^{13}$C NMR (101 MHz, DMSO-$d_6$) δ 172.7, 169.8, 166.2, 165.7, 137.8, 134.8, 132.1, 131.7, 130.3, 129.7, 128.9, 123.3, 121.5, 118.9, 96.0, 84.7, 49.0, 30.9, 21.9. HRMS (ESI) calc'd for $C_{21}H_{14}N_2O_4$ + H = 359.1026, found 359.1022.

**General Procedure G**: Sonogashira coupling of 5-bromothalidomide with phenylacetylene

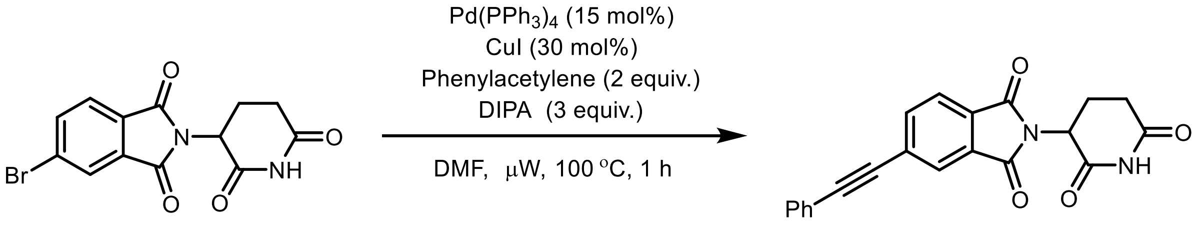

In an oven-dried 5 mL microwave reaction vial with a stir-bar was charged with 5-bromo-2-(2,6-dioxopiperidin-3-yl)isoindoline-1,3-dione (50 mg, 0.15 mmol), $Pd(PPh_3)_4$ (25 mg, 0.02 mmol), CuI (8.3 mg, 0.06 mmol). To this mixture, phenylacetylene (30.2 mg, 0.3 mmol), DIPA (45 mg, 0.4 mmol), and DMF (1.5 mL) were added and deaerated under Argon at rt for 5 min. The reaction vial was sealed with a microwave-supported Cap with Septum. The reaction mixture was stirred at 100 °C for 1 h in the microwave reactor. The solvent was evaporated under the reduced pressure, the residue was diluted with ethyl acetate (50 mL) and washed with water (10 mL x2), brine (20 mL), dried over anhydrous $Na_2SO_4$, filtered, and concentrated under reduced pressure. The crude residue was purified by Combiflash ISCO to obtain the pure product.

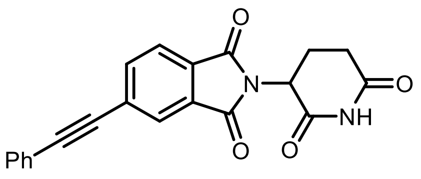

**2-(2,6-dioxopiperidin-3-yl)-5-(phenylethynyl)isoindoline-1,3-dionedione (61)** was prepared by following the general procedure **G**. The pure product was isolated by ISCO and the column ran with $CH_2Cl_2$, then grading to 5%MeOH (contains 1% $NH_4OH$)-$CH_2Cl_2$ in 66% yield (35 mg; white solid). $^{1}$H NMR (400 MHz, DMSO-$d_6$) δ 11.14 (s, 1H), 8.09 – 8.00 (m, 2H), 7.97 (d, $J$ = 7.7 Hz, 1H), 7.65 (dd, $J$ = 6.5, 3.0 Hz, 2H), 7.48 (dd, $J$ = 4.9, 1.9 Hz, 3H), 5.18 (dd, $J$ = 12.8, 5.4 Hz, 1H), 2.67 – 2.51 (m, 2H), 2.08 (ddd, $J$ = 10.6, 5.7, 3.3 Hz, 1H). $^{13}$C NMR (101 MHz, DMSO-$d_6$) δ 172.7, 169.7, 166.5, 166.4, 137.4, 131.9, 131.7, 130.4, 129.6, 128.9, 128.7, 125.8, 123.8, 121.4, 93.6, 87.9, 49.1, 30.9, 21.9. HRMS (ESI) calc'd for $C_{21}H_{14}N_2O_4$ + H = 359.1026, found 359.1024.

**General Procedure H**: Preparation of 4-substituted IMiD analogs through amidation with acid chlorides

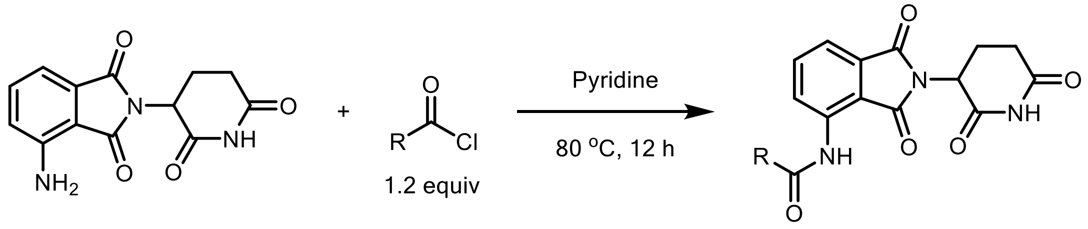

In an oven-dried 5 mL vial, acid chloride (0.26 mmol, 1.3 equiv.) was added to a solution of pomalidomide (54 mg, 0.2 mmol) in dry pyridine (3 mL) at 0 ˚C. The mixture was warmed to RT and stirred at 80 °C for 12 h. The solvent was evaporated under reduced pressure, and the residue was purified by the Teledyne ISCO CombiFlash system with $CH_2Cl_2$-MeOH to obtain the pure product.

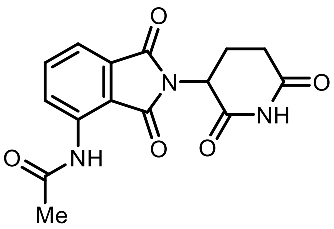

***N*-(2-(2,6-dioxopiperidin-3-yl)-1,3-dioxoisoindolin-4-yl)acetamide (62)** was prepared by following the general procedure **H** using acetyl chloride (1.19 mmol, 1.3 equiv). Final product was isolated by flash column chromatography ($CH_2Cl_2$: MeOH 90:10) in 69% yield (200 mg; light yellow solid). $^{1}H$ NMR (400 MHz, DMSO-$d_6$) δ 11.14 (s, 1H), 9.72 (s, 1H), 8.45 (dd, *J* = 8.4, 0.7 Hz, 1H), 7.82 (dd, *J* = 8.4, 7.3 Hz, 1H), 7.61 (dd, *J* = 7.3, 0.8 Hz, 1H), 5.14 (dd, *J* = 12.7, 5.4 Hz, 1H), 2.90 (ddd, *J* = 16.8, 13.7, 5.4 Hz, 1H), 2.67 – 2.52 (m, 2H), 2.13 – 1.99 (m, 1H). $^{13}C$ NMR (101 MHz, DMSO-$d_6$) δ 172.7, 169.7, 169.2, 167.5, 166.6, 136.5, 136.0, 131.5, 126.4, 118.3, 117.0, 48.9, 39.5, 30.9, 24.1, 22.0. HRMS (ESI) calc'd for $C_{15}H_{13}N_3O_5$ + H = 316.0928, found 316.0928.

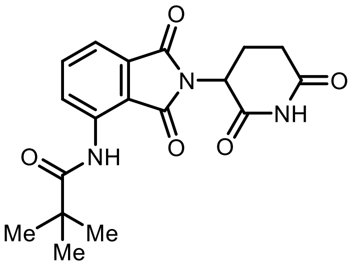

***N*-(2-(2,6-dioxopiperidin-3-yl)-1,3-dioxoisoindolin-4-yl)pivalamide (63)** was prepared by following the general procedure **H** using pivaloyl chloride (31 mg, 0.26 mmol). The crude product was purified by ISCO with a 4 g column. The column ran with $CH_2Cl_2$ then grading to 5% MeOH-$CH_2Cl_2$ to afford 14% yield (10 mg; white solid). $^{1}H$ NMR (400 MHz, DMSO-d6) δ 11.13 (s, 1H), 9.77 (s, 1H), 8.60 (dd, *J* = 8.4, 0.7 Hz, 1H), 7.84 (dd, *J* = 8.5, 7.3 Hz, 1H), 7.61 (dd, *J* = 7.3, 0.7 Hz, 1H), 5.16 (dd, *J* = 12.8, 5.3 Hz, 1H), 2.89 (tdd, *J* = 15.9, 4.4, 2.9 Hz, 1H), 2.67 – 2.51 (m, 2H), 2.13 – 2.02 (m, 1H), 1.28 (s, 9H). $^{13}C$ NMR (101 MHz, DMSO-$d_6$) δ 176.9, 172.7, 169.7, 168.6, 166.7,137.0, 136.4, 131.1, 124.9, 118.1, 116.5, 48.9, 39.6, 30.9, 26.9, 21.9. HRMS (ESI) calc'd for $C_{18}H_{19}N_3O_5$ + H = 358.1397, found 358.1395.

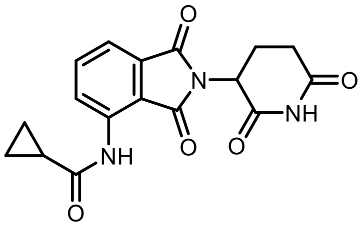

***N*-(2-(2,6-dioxopiperidin-3-yl)-1,3-dioxoisoindolin-4-yl)cyclopropanecarboxamide (64)** was prepared by following the general procedure **H** using cyclopropanecarbonyl chloride (0.24 mmol, 1.3 equiv). Final product was isolated by flash column chromatography ($CH_2Cl_2$: MeOH 90:10) in 71% yield (44 mg; white solid). $^{1}H$ NMR (400 MHz, DMSO-$d_6$) δ 11.14 (s, 1H), 9.98 (s, 1H), 8.41 (dd, *J* = 8.5, 0.8 Hz, 1H), 7.81 (dd, *J* = 8.5, 7.3 Hz, 1H), 7.61 (dd, *J* = 7.3, 0.8 Hz, 1H), 5.15 (dd, *J* = 12.8, 5.4 Hz, 1H), 2.90 (ddd, *J* = 16.7, 13.7, 5.2 Hz, 1H), 2.67 – 2.51 (m, 2H), 2.24 – 2.03 (m, 1H), 2.02 – 1.90 (m, 1H), 0.99 – 0.81 (m, 4H). $^{13}C$ NMR (101 MHz, DMSO-$d_6$) δ 172.7, 172.5, 169.8, 167.5, 166.6, 136.4, 135.9, 131.5, 126.7, 118.3, 117.0, 48.9, 39.5, 30.9, 22.0, 14.9, 8.1. HRMS (ESI) calc'd for $C_{17}H_{15}N_3O_5$ + H = 342.1084, found 342.1082.

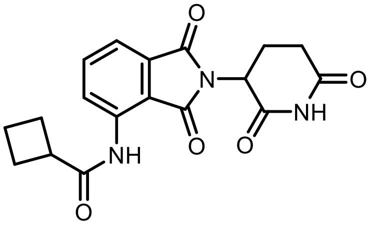

***N*-(2-(2,6-dioxopiperidin-3-yl)-1,3-dioxoisoindolin-4-yl)cyclobutanecarboxamide (65)** was prepared by following the general procedure **H** using cyclobutanecarbonyl chloride (0.24 mmol, 1.3 equiv). Final product was isolated by flash column chromatography ($CH_2Cl_2$: MeOH 90:10) in 68% yield (92 mg; light yellow solid). $^{1}H$ NMR (400 MHz, DMSO-$d_6$) δ 11.14 (s, 1H), 9.54 (s, 1H), 8.51 (dd, *J* = 8.4, 0.7 Hz, 1H), 7.83 (dd, *J* = 8.5, 7.3 Hz, 1H), 7.60 (dd, *J* = 7.3, 0.8 Hz, 1H), 5.14 (dd, *J* = 12.8, 5.4 Hz, 1H), 3.39 (pd, *J* = 8.5, 1.1 Hz, 1H), 2.89 (ddd, *J* = 16.8,13.7, 5.4 Hz, 1H), 2.67 – 2.51 (m, 2H), 2.33 – 2.14 (m, 4H), 2.11 – 2.02 (m, 1H), 2.01 – 1.90 (m, 1H), 1.88 – 1.77 (m,

1H). $^{13}C$ NMR (101 MHz, DMSO-$d_6$) δ 173.6, 172.7, 169.7, 167.9, 166.6, 136.7, 136.2, 131.4, 125.9, 118.2, 116.8, 48.9, 39.5, 30.9, 24.7, 24.6, 22.0, 17.5. HRMS (ESI) calc'd for $C_{18}H_{17}N_3O_5$ + H = 356.1241, found 356.1238.

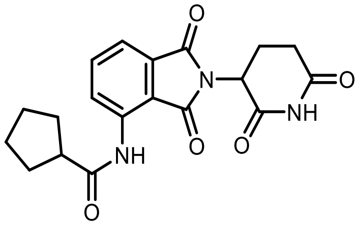

***N*-(2-(2,6-dioxopiperidin-3-yl)-1,3-dioxoisoindolin-4-yl)cyclopentanecarboxamide (66)** was prepared by following the general procedure **H** using cyclopentanecarbonyl chloride (0.24 mmol, 1.3 equiv). Final product was isolated by flash column chromatography ($CH_2Cl_2$: MeOH 90:10) in 60% yield (40 mg; white solid). $^{1}H$ NMR (400 MHz, DMSO-$d_6$) δ 11.18 (s, 1H), 9.70 (s, 1H), 8.46 (d, *J* = 8.4 Hz, 1H), 7.83 (dd, *J* = 8.4, 7.3 Hz, 1H), 7.61 (d, *J* = 7.2 Hz, 1H), 5.15 (dd, *J* = 12.8, 5.4 Hz, 1H), 3.08 – 2.80 (m, 2H), 2.69 – 2.51 (m, 2H), 2.06 (ddd, *J* = 13.2, 6.3, 3.9 Hz, 1H), 1.91 (dt, *J* = 12.4, 8.0 Hz, 2H), 1.77 (dtd, *J* = 12.7, 10.3, 9.0, 6.4 Hz, 2H), 1.71 – 1.52 (m, 4H). $^{13}C$ NMR (101 MHz, DMSO-$d_6$) δ 174.8, 172.7, 169.7, 167.9, 166.7, 136.7, 136.1, 131.4, 126.1, 118.2, 116.9, 48.9, 45.7, 39.5, 30.9, 29.7, 29.7, 25.5, 22.0. HRMS (ESI) calc'd for $C_{19}H_{19}N_3O_5$ + H = 370.1397, found 370.1394.

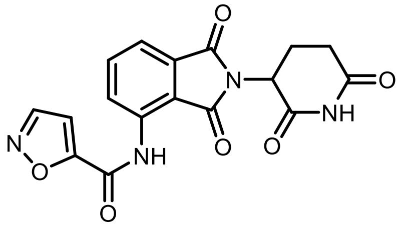

***N*-(2-(2,6-dioxopiperidin-3-yl)-1,3-dioxoisoindolin-4-yl)isoxazole-5-carboxamide (67)** was prepared by following the general procedure **H** using isoxazole-5-carbonyl chloride (0.24 mmol). After reaction completion, the solvent was removed under reduced pressure. Water was added and solid residue isolated by filtration, washed with diethyl ether, and dried on air affording the desired product (24 mg, 33% yield). $^{1}H$ NMR (400 MHz, DMSO-$d_6$) δ 11.15 (s, 1H), 10.70 (s, 1H), 8.89 (d, *J* = 2.0 Hz, 1H), 8.46 (d, *J* = 8.3 Hz, 1H), 7.94 (t, *J* = 7.8 Hz, 1H), 7.75 (d, *J* = 7.3 Hz, 1H), 7.34 (d, *J* = 2.0 Hz, 1H), 5.17 (dd, *J* = 13.0, 5.4 Hz, 1H), 2.90 (ddd, *J* = 18.7, 14.3, 5.4 Hz, 1H), 2.59 (dd, *J* = 23.9, 15.0 Hz, 2H), 2.24 – 1.97 (m, 1H). $^{13}C$ NMR (101 MHz, DMSO-$d_6$) δ 172.7, 169.6, 167.5, 166.5, 161.5, 153.9, 152.3, 136.4, 134.8, 131.5, 126.9, 119.8, 118.9, 107.9, 49.0, 30.9, 21.9. HRMS (ESI) calc'd for $C_{17}H_{12}N_4O_6$ + H = 369.0830, found 369.0826.

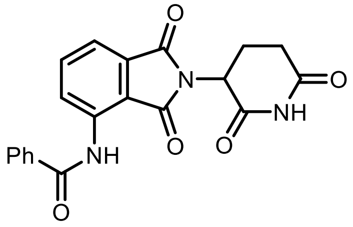

***N*-(2-(2,6-dioxopiperidin-3-yl)-1,3-dioxoisoindolin-4-yl)benzamide (67)** was prepared by following the general procedure **H** using benzoyl chloride (0.24 mmol, 1.3 equiv). Final product was isolated by flash column chromatography (DCM: MeOH 90:10) in 80% yield (55 mg; white solid). $^{1}H$ NMR (400 MHz, DMSO-$d_6$) δ 11.15 (s, 1H), 10.42 (s, 1H), 8.61 (dd, *J* = 8.4, 0.7 Hz, 1H), 8.10 – 7.96 (m, 2H), 7.91 (dd, *J* = 8.3, 7.4 Hz, 1H), 7.71 – 7.66 (m, 2H), 7.65 – 7.59 (m, 2H), 5.18 (dd, *J* = 13.0, 5.4 Hz, 1H), 2.90 (ddd, *J* = 17.4, 14.0, 5.4 Hz, 1H), 2.67 – 2.53 (m, 2H), 2.18 – 2.02 (m, 1H). $^{13}C$ NMR (101 MHz, DMSO-$d_6$) δ 172.7, 169.7, 168.1, 166.6, 165.0, 136.6, 136.3, 133.3, 132.6, 131.4, 129.0, 127.3, 126.2, 118.7, 117.9, 49.0, 39.5, 30.9, 22.0. HRMS (ESI) calc'd for $C_{20}H_{15}N_3O_5$ + H = 378.1084, found 378.1080.

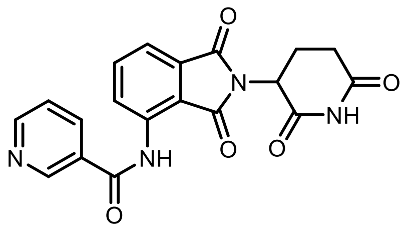

***N*-(2-(2,6-dioxopiperidin-3-yl)-1,3-dioxoisoindolin-4-yl)nicotinamide (68)** was prepared by following the general procedure **H** using nicotinyl chloride (0.26 mmol). Final product was isolated by flash column chromatography (DCM: MeOH 95:5) in 46% yield (35 mg; white solid). $^1$H NMR (400 MHz, DMSO-$d_6$) δ 11.15 (s, 1H), 10.54 (s, 1H), 9.15 (dd, J = 2.3, 0.9 Hz, 1H), 8.83 (dd, J = 4.8, 1.6 Hz, 1H), 8.47 (dd, J = 8.3, 0.8 Hz, 1H), 8.32 (dt, J = 8.0, 1.9 Hz, 1H), 7.92 (dd, J = 8.3, 7.4 Hz, 1H), 7.72 (dd, J = 7.3, 0.7 Hz, 1H), 7.64 (ddd, J = 8.0, 4.8, 0.8 Hz, 1H), 5.18 (dd, J = 12.9, 5.4 Hz, 1H), 2.91 (ddd, J = 17.3, 13.9, 5.4 Hz, 1H), 2.69 – 2.53 (m, 2H), 2.20 – 1.97 (m, 1H).$^{13}$C NMR (101 MHz, DMSO-$d_6$) δ 172.7, 169 7, 167.5, 166.6, 163.8, 152.9, 148.5, 136.2, 135.9, 135.2, 131.5, 129.1, 127.3, 123.9, 119.3, 119.0, 49.0, 30.9, 22.0. HRMS (ESI) calc'd for $C_{19}H_{14}N_4O_5$ + H = 379.1037, found 379.1033.

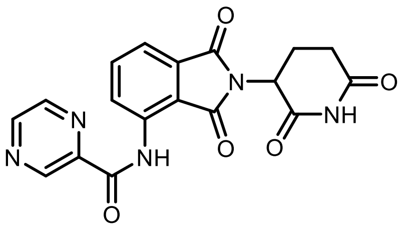

***N*-(2-(2,6-dioxopiperidin-3-yl)-1,3-dioxoisoindolin-4-yl)pyrazine-2-carboxamide (68)** was prepared by following the general procedure **H** using pyrazine-2-carbonyl chloride (0.26 mmol). Final product was isolated by flash column chromatography (DCM: MeOH 90:10) in 23% yield (19 mg; white solid). $^1$H NMR (400 MHz, DMSO-$d_6$) δ 11.63 (s, 1H), 11.17 (s, 1H), 9.40 (s, 1H), 9.03 (d, *J* = 2.4 Hz, 1H), 8.92 (d, *J* = 7.5 Hz, 2H), 7.97 (t, *J* = 7.9 Hz, 1H), 7.71 (d, *J* = 7.3 Hz, 1H), 5.20 (dd, *J* = 12.9, 5.5 Hz, 1H), 2.92 (t, *J* = 16.7 Hz, 1H), 2.77 – 2.54 (m, 2H), 2.10 (dd, *J* = 13.0, 6.2 Hz, 1H).$^{13}$C NMR (101 MHz, DMSO-$d_6$) δ 172.7, 169.6, 168.3, 166.6, 161.7, 148.7, 144.0, 143.6, 143.1, 136.6, 135.9, 131.4, 124.4, 118.3, 116.7, 49.0, 30.9, 22.0. HRMS (ESI) calc'd for $C_{18}H_{13}N_5O_5$ + H = 380.0989, found 380.0989.

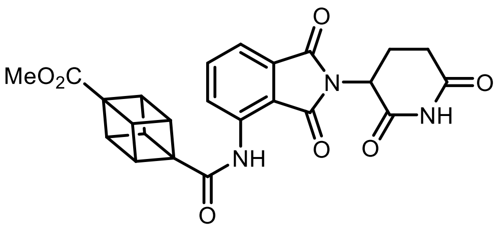

**Methyl 4-((2-(2,6-dioxopiperidin-3-yl)-1,3dioxoisoindolin-4-yl)carbamoyl)cubane-1-carboxylate (69)** was prepared by following the general procedure **H** methyl (1r,2R,3r,8S)-4-(chlorocarbonyl)cubane-1-carboxylate (0.26 mmol). Final product was isolated by flash column chromatography (DCM: MeOH 95:5) in 42% yield (39 mg; yellow solid). $^1$H NMR (400 MHz, DMSO-$d_6$) δ 11.16 (s, 1H), 9.69 (s, 1H), 8.52 (d, *J* = 8.4 Hz, 1H), 7.85 (t, *J* = 7.9 Hz, 1H), 7.63 (d, *J* = 7.3 Hz, 1H), 5.17 (dd, *J* = 12.8, 5.4 Hz, 1H), 4.37 – 4.09 (m, 6H), 3.65 (s, 3H), 2.89 (ddd, *J* = 16.7, 13.6, 5.3 Hz, 1H), 2.74 – 2.54 (m, 2H), 2.06 (dd, *J* = 11.2, 5.9 Hz, 1H). $^{13}$C NMR (101 MHz, DMSO-$d_6$) δ172.8, 171.0, 169.7, 168.1, 166.7, 136.4, 136.3, 131.3, 125.6, 118.5, 117.0, 57.9, 55.2, 51.4, 49.0, 46.6, 46.1, 46.1, 46.1, 31.0, 22.0. HRMS (ESI) calc'd for $C_{24}H_{19}N_3O_7$ + H = 462.1296, found 462.1294.

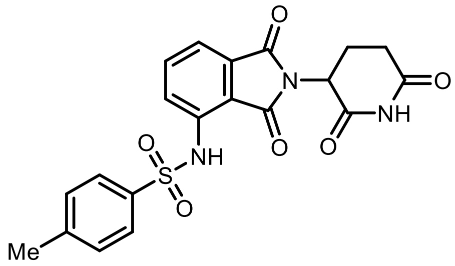

***N*-(2-(2,6-dioxopiperidin-3-yl)-1,3-dioxoisoindolin-4-yl)-4-methylbenzene sulfonamide (6)** was prepared by following the general procedure **H** using tosyl chloride (0.24 mmol). Final product was isolated by flash column chromatography (DCM: MeOH 90:10) in 51% yield (44 mg; white solid). $^1$H NMR (400 MHz, DMSO-$d_6$) δ 11.09 (s, 1H), 9.84 (s, 1H), 7.84 – 7.78 (m, 2H), 7.73 (dd, *J* = 8.4, 7.3 Hz, 1H), 7.56 (dd, *J* = 10.3, 7.8 Hz, 2H), 7.35 (d, *J* = 8.1 Hz, 2H), 5.07 (dd, *J* = 12.9, 5.4 Hz, 1H), 2.85 (ddd, *J* = 17.1, 13.8, 5.4 Hz, 1H), 2.57 (dt, *J* = 17.1, 2.9 Hz, 1H), 2.45 (dd, *J* = 13.2, 4.4 Hz, 1H), 2.31 (s, 3H), 2.00 (ddd, *J* = 9.9, 8.6, 4.6 Hz, 1H). $^{13}$C NMR (101 MHz, DMSO-$d_6$) δ 172.7, 169.6, 166.6, 166.3, 144.2, 136.4, 136.3, 135.1, 132.1, 129.9, 127.0, 125.1, 118.8, 118.4, 48.9, 30.9, 21.9, 21.0. HRMS (ESI) calc'd for $C_{20}H_{17}N_3O_6S$ + H = 428.0911, found 428.0909.

**General Procedure I**: Preparation of 5-substituted IMiD analogs through amidation with acid chlorides

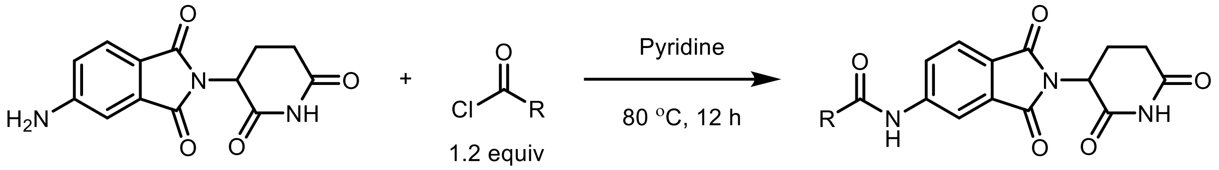

In an oven-dried 5 mL vial with a stir-bar was charged with 5-amino-2-(2,6-dioxopiperidin-3-yl)isoindoline-1,3-dione (54 mg, 0.2 mmol, 1 equiv.). The flask was capped with a rubber septum and was evacuated and back-filled with nitrogen. This process was repeated for additional three times. Dry pyridine (3 mL) followed by acid chloride (0.26 mmol, 1.3 equiv.) were sequentially added into the vial by using a syringe. The mixture was warmed to room temperature and stirred at 80 °C for 12 h. The solvent was evaporated under reduced pressure and the residue was purified by Teledyne ISCO Chromatography on silica gel ($CH_2Cl_2$/MeOH) to obtain the pure product.

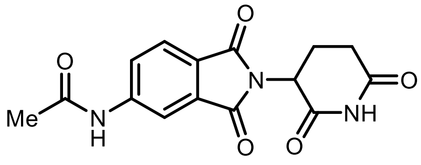

***N*-(2-(2,6-dioxopiperidin-3-yl)-1,3-dioxoisoindolin-5-yl)acetamide (72)** was prepared by following the general procedure **I** using acetyl chloride (0.26 mmol). Final product was isolated by flash column chromatography (DCM: MeOH 90:10) in 72% yield (45 mg; yellow solid). $^1$H NMR (400 MHz, DMSO-$d_6$) δ 11.10 (s, 1H), 10.57 (d, *J* = 3.5 Hz, 1H), 8.22 (d, *J* = 2.2 Hz, 1H), 7.90 – 7.75 (m, 2H), 5.12 (dd, *J* = 12.9, 5.4 Hz, 1H), 2.89 (ddd, *J* = 17.4, 14.0, 5.4 Hz, 1H), 2.71 – 2.46 (m, 2H), 2.13 (s, 3H), 2.09 – 1.99 (m, 1H). $^{13}$C NMR (101 MHz, DMSO-$d_6$) δ 172.8, 169.9, 169.4, 167.0, 166.8, 145.1, 132.8, 124.8, 124.6, 123.4, 112.8, 49.0, 31.0, 24.3, 22.1. HRMS (ESI) calc'd for $C_{15}H_{13}N_3O_5$ + H = 316.0928, found 316.092.

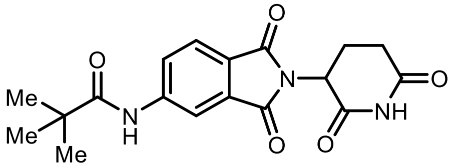

***N*-(2-(2,6-dioxopiperidin-3-yl)-1,3-dioxoisoindolin-5-yl)pivalamide (73)** was prepared by following the general procedure **H** using pivaloyl chloride (0.26 mmol). Final product was isolated by flash column chromatography (DCM: MeOH 90:10) in 68% yield (48 mg; yellow solid). $^1$H NMR (400 MHz, DMSO-$d_6$) δ 11.11 (s, 1H), 9.80 (s, 1H), 8.30 (d, *J* = 1.8 Hz, 1H), 8.09 (dd, *J* = 8.3, 1.9 Hz, 1H), 7.86 (d, *J* = 8.3 Hz, 1H), 5.12 (dd, *J* = 12.9, 5.4 Hz, 1H), 2.89 (ddd, *J* = 17.4, 14.0, 5.4 Hz,

1H), 2.70 – 2.51 (m, 2H), 2.06 (ddd, *J* = 10.4, 5.3, 2.9 Hz, 1H), 1.26 (s, 9H). $^{13}$C NMR (101 MHz, DMSO-$d_6$) δ 177.4, 172.7, 169.9, 167.1, 166.8, 145.3, 132.5, 124.8, 124.4, 124.3, 113.9, 48.9, 39.5, 30.9, 26.9, 22.1. HRMS (ESI) calc'd for $C_{18}H_{19}N_3O_5$ + H = 358.1397, found 358.1395

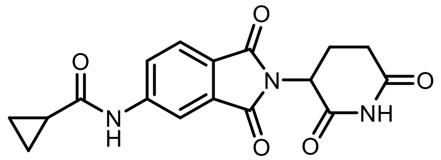

***N*-(2-(2,6-dioxopiperidin-3-yl)-1,3-dioxoisoindolin-5-yl) cyclopropane carboxamide (74)** was prepared by following the general procedure **I** using cyclopropanecarbonyl chloride (0.26 mmol). Final product was isolated by flash column chromatography ($CH_2Cl_2$: MeOH 90:10) in 61% yield (42 mg). $^1$H NMR (400 MHz, DMSO-$d_6$) δ 11.10 (s, 1H), 10.86 (s, 1H), 8.23 (d, *J* = 1.7 Hz, 1H), 8.04 – 7.77 (m, 2H), 5.11 (dd, *J* = 12.9, 5.4 Hz, 1H), 3.00 – 2.75 (m, 1H), 2.63 – 2.40 (m, 2H), 2.13 – 1.98 (m, 1H), 1.83 (p, *J* = 6.2 Hz, 1H), 0.88 (d, *J* = 6.1 Hz, 4H). $^{13}$C NMR (101 MHz, DMSO-$d_6$) δ 172.7, 172.7, 169.9, 167.0, 166.7, 145.0, 132.8, 124.7, 124.6, 123.4, 112.8, 48.9, 30.9, 22.0, 14.9, 8.0. HRMS (ESI) calc'd for $C_{17}H_{15}N_3O_5$ + H = 342.1084, found 342.1084.

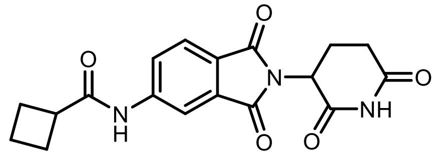

***N*-(2-(2,6-dioxopiperidin-3-yl)-1,3-dioxoisoindolin-5-yl) cyclobutane carboxamide (75)** was prepared by following the general procedure **I** using cyclobutanecarbonyl chloride (0.26 mmol). Final product was isolated by flash column chromatography ($CH_2Cl_2$: MeOH 90:10) in 67% yield (47 mg; yellow solid). $^1$H NMR (400 MHz, DMSO-$d_6$) δ 11.10 (s, 1H), 10.39 (s, 1H), 8.27 (d, *J* = 1.7 Hz, 1H), 7.93 (dd, *J* = 8.2, 1.8 Hz, 1H), 7.85 (d, *J* = 8.2 Hz, 1H), 5.12 (dd, *J* = 12.9, 5.4 Hz, 1H), 3.28 (d, *J* = 8.4 Hz, 1H), 2.89 (ddd, *J* = 17.4, 14.0, 5.5 Hz, 1H), 2.68 – 2.51 (m, 2H), 2.36 – 2.10 (m, 4H), 2.09 – 1.77 (m, 3H). $^{13}$C NMR (101 MHz, DMSO-$d_6$) δ 173.9, 172.7, 169.9, 167.0, 166.7, 145.2, 132.7,124.7, 124.6, 123.6, 112.9, 49.0, 30.9, 24.5, 22.1, 17.7. HRMS (ESI) calc'd for $C_{18}H_{17}N_3O_5$ + H = 356.1241, found 356.1240.

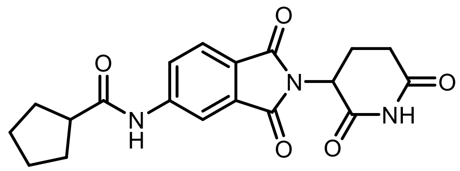

***N*-(2-(2,6-dioxopiperidin-3-yl)-1,3-dioxoisoindolin-5-yl) cyclopentane carboxamide (76)** was prepared by following the general procedure **I** using cyclopentanecarbonyl chloride (0.26 mmol). Final product was isolated by flash column chromatography ($CH_2Cl_2$: MeOH 90:10) in 58% yield (43 mg; Yellow solid). $^1$H NMR (400 MHz, DMSO-$d_6$) δ 11.10 (s, 1H), 10.51 (s,1H), 8.26 (d, J = 1.7 Hz, 1H), 7.92 (dd, J = 8.2, 1.9 Hz,1H), 7.85 (d, J = 8.2 Hz, 1H), 5.12 (dd, J = 12.9, 5.4 Hz, 1H), 3.17 (d, J = 5.1 Hz, 1H), 2.97 – 2.75 (m, 2H), 2.65 – 2.48 (m, 2H), 2.05 (ddt, J = 13.2, 5.8, 2.6 Hz, 1H), 1.93 – 1.82 (m, 2H), 1.82 – 1.63 (m, 3H), 1.57 (dt, J = 8.3, 4.3 Hz, 2H). $^{13}$C NMR (101 MHz, DMSO-$d_6$) δ 175.4, 172.7, 169.9, 167.0, 166.8, 145.3, 132.8, 124.7, 124.6, 123.5, 112.9, 49.0, 45.5, 31.0, 30.0, 25.7, 22.1. HRMS (ESI) calc'd for $C_{19}H_{19}N_3O_5$ + H = 370.1397, found 370.1395.

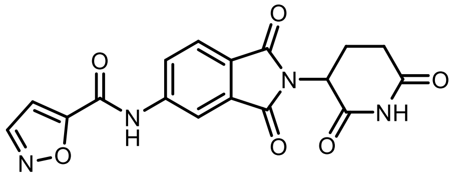

***N*-(2-(2,6-dioxopiperidin-3-yl)-1,3-dioxoisoindolin-5-yl)isoxazole-5-carboxamide (77)** was prepared by following the general procedure **I** using Isoxazole-5-carbonyl chloride (0.26 mmol). Final product was isolated by flash column chromatography (DCM: MeOH 90:10) in 43% yield (32 mg; yellow solid). $^1$H NMR (400 MHz, DMSO-$d_6$) δ 11.33 (s, 1H), 11.13 (s, 1H), 8.86 (s, 1H), 8.38 (s, 1H), 8.20 (d, *J* = 8.2 Hz, 1H), 7.96 (d, *J* = 8.2 Hz, 1H), 7.35 (s, 1H), 5.15 (dd, *J* = 12.8, 5.4 Hz, 1H), 2.89 (dd, *J* = 10.9, 6.7 Hz, 1H), 2.59 (dd, *J* = 26.4, 15.4 Hz, 2H), 2.08 ppm (dd, *J* = 8.6, 4.6 Hz, 1H). $^{13}$C NMR (101 MHz, DMSO-$d_6$) δ 173.2, 170.3, 167.3, 167.1, 162.3, 155.1, 152.5, 144.1, 133.1, 126.8, 125.8, 125.1, 115.1, 108.2, 49.5, 31.4, 22.5. HRMS (ESI) calc'd for $C_{17}H_{12}N_4O_6$ + H = 369.0830, found 369.0827.

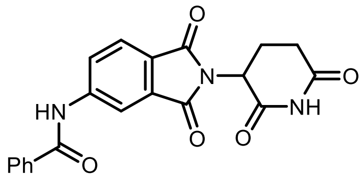

***N*-(2-(2,6-dioxopiperidin-3-yl)-1,3-dioxoisoindolin-5-yl)benzamide (78)** was prepared by following the general procedure **I** using benzoyl chloride (0.26 mmol). Final product was isolated by flash column chromatography (DCM: MeOH 90:10) in 64% yield (48.2 mg; white solid). $^1$H NMR (400 MHz, DMSO-$d_6$) δ 11.12 (s, 1H), 10.84 (s, 1H), 8.42 (d, *J* = 1.9 Hz, 1H), 8.20 (dd, *J* = 8.2, 1.9 Hz, 1H), 8.05 – 7.98 (m, 2H), 7.93 (d, *J* = 8.2 Hz, 1H), 7.70 – 7.61 (m, 1H), 7.57 (dd, *J* = 8.2, 6.6 Hz, 2H), 5.14 (dd, *J* = 12.9, 5.4 Hz, 1H), 2.90 (ddd, *J* = 17.3, 14.0, 5.4 Hz, 1H), 2.67, 2.52 (m, 2H), 2.08 (ddp, *J* = 10.3, 5.7, 2.7 Hz, 1H). $^{13}$C NMR (101 MHz, DMSO-$d_6$) δ 172.7, 169.9, 167.0, 166.8, 166.3, 145.1,134.1,132.6, 132.2, 128.5, 127.9, 125.3, 124.7, 124.5, 114.1, 49.0, 39.5, 31.0, 22.1. HRMS (ESI) calc'd for $C_{20}H_{15}N_3O_5$ + H = 378.1084, found 378.1082.

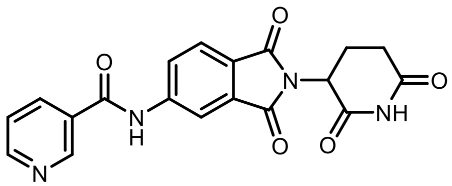

***N*-(2-(2,6-dioxopiperidin-3-yl)-1,3-dioxoisoindolin-5-yl) nicotinamide (79)** was prepared by following the general procedure **I** using nicotinyl chloride (0.26 mmol). Final product was isolated by flash column chromatography ($CH_2Cl_2$: MeOH 95:5) in 46% yield (35 mg; white solid). $^1$H NMR (400 MHz, DMSO-$d_6$) δ 11.13 (s, 1H), 11.01 (s, 1H), 9.15 (d, *J* = 2.2 Hz, 1H), 8.80 (dd, *J* = 4.8, 1.7 Hz, 1H), 8.40 (d, *J* = 1.8 Hz, 1H), 8.34 (dt, *J* = 8.0, 2.0 Hz, 1H), 8.18 (dd, *J* = 8.2, 1.9 Hz, 1H), 7.94 (d, *J* = 8.2 Hz, 1H), 7.61 (dd, *J* = 7.9, 4.8 Hz, 1H), 5.15 (dd, *J* = 12.9, 5.4 Hz, 1H), 2.91 (ddd, *J* = 17.4, 14.0, 5.4 Hz, 1H), 2.71 – 2.44 (m, 3H), 2.20 – 1.98 ppm (m, 1H). $^{13}$C NMR (101 MHz, DMSO-$d_6$) δ 173.2, 170.4, 167.4, 167.2, 165.3, 153.1, 149.3, 145.2, 136.2, 133.1, 130.4, 126.1, 125.3, 125.0, 124.1, 114.6, 49.5, 31.4, 22.5. HRMS (ESI) calc'd for $C_{19}H_{14}N_4O_5$ + H = 379.1037, found 379.1032.

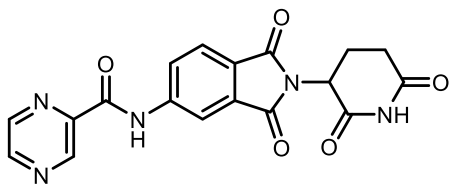

***N*-(2-(2,6-dioxopiperidin-3-yl)-1,3-dioxoisoindolin-5-yl)pyrazine-2-carboxamide (80)** was prepared by following the general procedure **I** using pyrazine-2-carbonyl chloride (0.26 mmol). Final product was isolated by flash column chromatography (DCM: MeOH 90:10) in 21% yield (16 mg; white solid). $^1$H NMR (400 MHz, DMSO-$d_6$) δ 11.41 (s, 1H), 11.12 (s, 1H), 9.36 (s, 1H), 8.99 (s, 1H), 8.87 (s, 1H), 8.55 (s, 1H), 8.38 (d, *J* = 8.2 Hz, 1H), 7.96 (d, *J* = 8.2 Hz, 1H), 5.16 (dd, *J* = 13.3, 5.5 Hz, 1H), 2.97– 2.74 (m, 1H), 2.62 (d, *J* = 16.8 Hz, 2H), 2.09 (d, *J* = 9.7 Hz, 1H). $^{13}$C NMR (101 MHz, DMSO-$d_6$) δ 172.8, 169.9, 167.0, 166.7, 162.7, 148.2, 144.5, 144.4, 144.2, 143.3, 132.6, 126.0, 125.4, 124.5, 114.6, 49.0, 31.0, 22.0. HRMS (ESI) calc'd for $C_{18}H_{13}N_5O_5$ + H = 380.0989, found 380.0989.

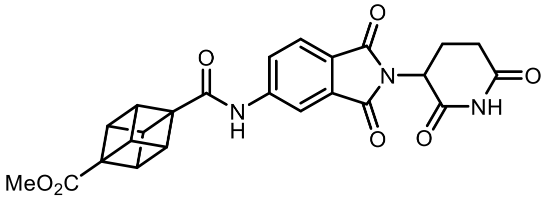

**Methyl4-((2-(2,6-dioxopiperidin-3-yl)-1,3- dioxoisoindolin-5-yl)carbamoyl)cubane-1-carboxylate (81)** was prepared by following the general procedure **I** methyl (1r,2R,3r,8S)-4-(chlorocarbonyl)cubane-1-carboxylate (0.26 mmol). Final product was isolated by flash column chromatography (DCM: MeOH 95:5) in 35% yield (32 mg; yellow solid). $^{1}$H NMR (400 MHz, DMSO-$d_6$) δ 11.13 (s, 1H), 10.34 (s, 1H), 8.37 – 8.11 (m, 1H), 8.04 (dd, $J$ = 8.4, 1.9 Hz, 1H), 7.88 (d, $J$ = 8.2 Hz, 1H), 5.12 (dd, $J$ = 12.9, 5.3 Hz, 1H), 4.31 (s, 3H), 4.19 (s, 3H), 3.64 (s, 3H), 2.99 – 2.75 (m, 1H), 2.69 – 2.51 (m, 2H), 2.13 – 1.82 (m, 1H). $^{13}$C NMR (101 MHz, DMSO-$d_6$) δ 172.8, 171.2, 170.3, 170.0, 167.0, 166.8, 145.0, 132.7, 125.0, 124.6, 124.0, 113.4, 57.8, 54.9, 51.3, 49.0, 46.7, 46.1, 31.0, 22.1. HRMS (ESI) calc'd for $C_{24}H_{19}N_3O_7$ + H = 462.1296, found 462.1296.

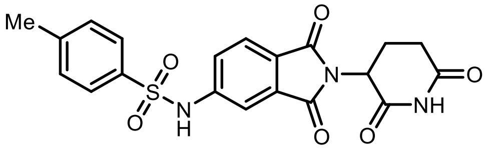

***N*-(2-(2,6-dioxopiperidin-3-yl)-1,3-dioxoisoindolin-5-yl)-4-methyl benzene sulfonamide (13)** was prepared by following the general procedure **I** using tosyl chloride (0.26 mmol). Final product was isolated by flash column chromatography ($CH_2Cl_2$: MeOH 90:10) in 53% yield (45 mg; white solid). $^{1}$H NMR (400 MHz, DMSO-$d_6$) δ 11.04 (s, 1H), 11.08 (s, 1H), 7.88 – 7.62 (m, 3H), 7.54 – 7.41 (m, 2H), 7.38 (d, $J$ = 8.1 Hz, 2H), 5.07 (dd, $J$ = 12.8, 5.4 Hz, 1H), 2.86 (ddd, $J$ = 17.1, 13.8, 5.4 Hz, 1H), 2.63 – 2.46 (m, 2H), 2.33 (s, 3H), 2.15 – 1.91 (m, 1H). $^{13}$C NMR (101 MHz, DMSO-$d_6$) δ 172.7, 169.8, 166.7, 166.5, 145.1, 143.7, 136.6, 133.1, 130.0, 126.6, 125.0, 124.6, 123.4, 112.1, 48.9, 30.9, 22.0, 21.0. HRMS (ESI) calc'd for $C_{20}H_{17}N_3O_6S$ + H = 428.0911, found 428.0909.

Preparation of compounds **7** and **14**

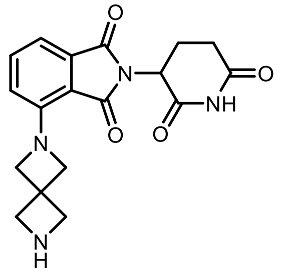

**2-(2,6-dioxopiperidin-3-yl)-4-(2,6-diazaspiro[3.3]heptan-2-yl)isoindoline-1,3-dione) (7):** In an oven-dried 7 mL vial with a stir-bar, TMSOTf (0.36 g, 1.65 mmol) was added to the solution of 2,6-dioxopiperidin-3-yl)-1,3-dioxoisoindolin-4-yl)-2,6-diazaspiro[3.3]heptane-2-carboxylate (**26**) (0.15 g, 0.33 mmol) and 2,6-lutidine (0.21 g, 1.98 mmol) in $CH_2Cl_2$ (4 mL) at 0 °C. The reaction was slowly warmed to rt and at stirred for 2 h. At this time MeOH (0.5 mL) was added and stirred at rt for 10 min. The reaction was cooled to rt and the solvent was evaporated. The residue was purified by Combiflash ISCO, and the column ran with 0-15% MeOH (contains 1% $NH_4OH$)-$CH_2Cl_2$ to afford 90 mg of the pure product as a yellow-colored solid in 76% yield. $^{1}$H NMR (400 MHz, DMSO-$d_6$) δ 11.05 (s, 1H), 8.43 (s, 1H), 7.59 (dd, $J$ = 8.5, 7.1 Hz, 1H), 7.15 (d, $J$ = 7.0 Hz, 1H), 6.81 (d, $J$ = 8.5 Hz, 1H), 5.05 (dd, $J$ = 12.7, 5.5 Hz, 1H), 4.36 (s, 4H), 4.17 (t, $J$ = 5.8 Hz, 4H), 2.88 (ddd, $J$ = 16.9, 13.8, 5.4 Hz, 1H), 2.64 – 2.50 (m, 3H), 2.01 (dtd, $J$ = 13.0, 5.3, 2.2 Hz, 1H). $^{13}$C NMR (101 MHz, DMSO-$d_6$) δ 172.8, 169.9, 167.1, 166.5, 147.4, 135.0, 133.2, 120.1, 112.1, 110.5, 63.1, 55.0, 48.7, 35.6, 30.9, 22.1. HRMS (ESI) calc'd for $C_{18}H_{18}N_4O_4$ + H = 355.1401, found 355.1400.

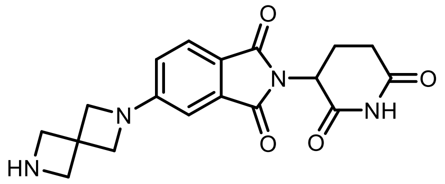

**2-(2,6-dioxopiperidin-3-yl)-5-(2,6-diazaspiro[3.3]heptan-2-yl)isoindoline-1,3-dione (14):** In an oven-dried 7 mL vial with a stir-bar, TMSOTf (0.36 g, 1.65 mmol) was added to the solution of *tert*-butyl (2-(2,6-dioxopiperidin-3-yl)-1,3-dioxoisoindolin-5-yl)-2,6-diazaspiro[3.3]heptane-2-carboxylate (0.15 g, 0.33 mmol) and 2,6-lutidine (0.21 g, 1.98 mmol) in $CH_2Cl_2$ (4 mL) at 0 °C. The reaction was slowly warmed to rt and at stirred for 2 h. At this time MeOH (0.5 mL) was added and stirred at rt for 10 min. The reaction was cooled to rt and the solvent was evaporated. The residue was purified by Combiflash ISCO, and the column ran with 0-15% MeOH (contains 1% $NH_4OH$)-$CH_2Cl_2$ to afford 95 mg of the pure product as a yellow-colored solid in 81% yield. $^1$H NMR (400 MHz, DMSO-d6) δ 11.06 (s, 1H), 8.48 (s, 1H), 7.67 (d, *J* = 8.2 Hz, 1H), 6.84 (d, *J* = 2.1 Hz, 1H), 6.70 (dd, *J* = 8.3, 2.1 Hz, 1H), 5.06 (dd, *J* = 12.9, 5.4 Hz, 1H), 4.21 (d, *J* = 9.6 Hz, 8H), 2.88 (ddd, *J* = 17.4, 14.1, 5.4 Hz, 1H), 2.64 – 2.51 (m, 2H), 2.01 (ddt, *J* = 13.1, 5.7, 2.6 Hz, 1H). $^{13}$C NMR (101 MHz, DMSO-d6) δ 172.7, 170.0, 167.4, 167.1, 154.6, 133.7, 124.8, 117.4, 114.7, 104.8, 60.7, 54.7, 48.6, 35.8, 30.9, 22.2. HRMS (ESI) calc'd for $C_{18}H_{18}N_4O_4$ + H = 355.1401, found 355.1400.

Preparation of 5-chloro-N2-(4-(1-(hex-5-yn-1-yl)piperidin-4-yl)-2-isopropoxy-5-methylphenyl)-N4-(2-(isopropylsulfonyl)phenyl)pyrimidine-2,4-diamine (**82**)

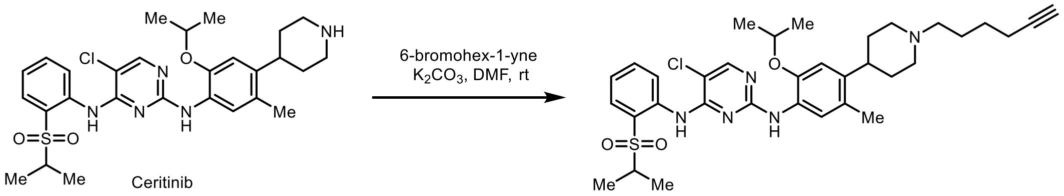

In an oven-dried 5 mL flask with a stir-bar, a mixture of ceritinib (0.11 g, 0.2 mmol), dry $K_2CO_3$ (0.040 g, 0.3 mmol), and 6-bromohex-1-yne (0.064 mg, 0.4 mmol) in DMF (3 mL) was stirred at rt for overnight. The reaction was cooled to rt and the solvent was evaporated under reduced pressure and then extracted with EtOAc (60 mL), washed with water (10 mL × 2), followed by brine (10 mL). The organic extracts were dried over anhydrous $Na_2SO_4$, filtered, and concentrated under reduced pressure. The residue was purified by Combiflash ISCO and the column ran with 0-10% MeOH (contains 1% $NH_4OH$)-$CH_2Cl_2$ to afford 0.1 g of 5-chloro-N2-(4-(1-(hex-5-yn-1-yl)piperidin-4-yl)-2-isopropoxy-5-methylphenyl)-N4-(2-(isopropylsulfonyl)phenyl)pyrimidine-2,4-diamine (**82**) as an off-white foam in 81% yield.

$^1$H NMR (400 MHz, DMSO-$d_6$) δ 9.45 (s, 1H), 8.46 (d, *J* = 8.4 Hz, 1H), 8.24 (s, 1H), 8.02 (s, 1H), 7.84 (d, *J* = 1.6 Hz, 0H), 7.67 – 7.57 (m, 1H), 7.51 (s, 1H), 7.40 – 7.31 (m, 1H), 6.84 (s, 1H), 4.57 (p, *J* = 6.0 Hz, 1H), 3.44 (q, *J* = 6.6 Hz, 1H), 3.09 – 2.91 (m, 2H), 2.74 (t, *J* = 2.6 Hz, 1H), 2.65 – 2.52 (m, 1H), 2.31 (t, *J* = 6.9 Hz, 2H), 2.18 (td, *J* = 6.8, 2.6 Hz, 2H), 2.11 (s, 3H), 1.98 (ddd, *J* = 11.0, 8.5, 5.9 Hz, 2H), 1.65 (h, *J* = 3.6 Hz, 4H), 1.58 – 1.38 (m, 4H), 1.21 (d, *J* = 6.0 Hz, 6H), 1.16 (d, *J* = 6.8 Hz, 6H). $^{13}$C NMR (101 MHz, $CDCl_3$) δ 162.6, 157.6, 155.4, 155.4, 144.9, 138.6, 134.7, 131.3 127.6, 126.7, 125.0, 123.8, 123.2, 120.6, 110.9, 105.8, 84.1, 71.5, 68.8, 58.3, 55.5, 54.5, 37.7, 36.5, 32.3, 31.5, 26.5, 25.6, 22.4, 19.0, 18.4, 15.4. HRMS (ESI) calc'd for $C_{34}H_{44}ClN_5O_3S$ + H = 638.2926, found 638.2923.

Preparation of **dALK-1**

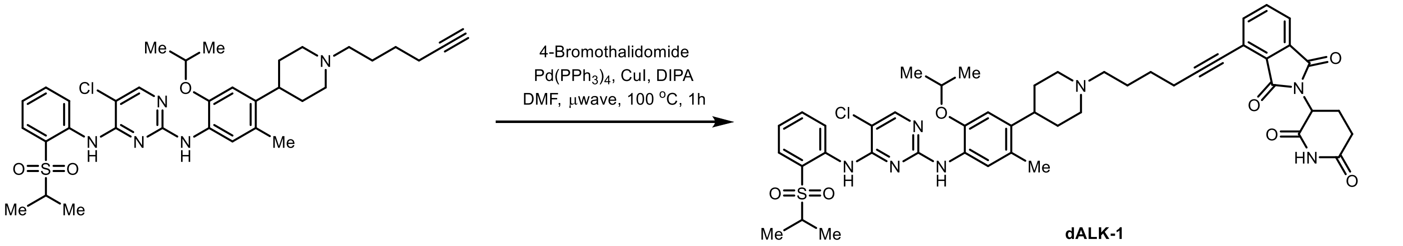

**dALK-1**

In an oven-dried 5 mL microwave vial with a stir-bar, a mixture of 4-bromothalidomide (33 mg, 0.1 mmol), $Pd(PPh_3)_4$ (18 mg, 0.015 mmol), CuI (6 mg, 0.03 mmol), 5-chloro-N2-(4-(1-(hex-5-yn-1-yl)piperidin-4-yl)-2-isopropoxy-5-methylphenyl)-*N*4-(2-(isopropylsulfonyl)phenyl)pyrimidine-2,4-diamine **82** (0.1 mmol), and DIPA (0.4 mmol) in DMF (2 mL) was deaerated under Argon at rt for 5 min. The vial was capped with a microwave supported rubber-septum. The reaction mixture was stirred in the microwave reactor at 100 °C for 1 h. The reaction was cooled to rt and the solvent was evaporated under reduced pressure and then extracted with EtOAc (60 mL), washed with water (10 mL × 2), followed by brine (10 mL). The organic extracts were dried over anhydrous $Na_2SO_4$, filtered, and concentrated under reduced pressure. The residue was purified by Combiflash ISCO, and the column ran with 0-10% MeOH (contains 1% $NH_4OH$)-$CH_2Cl_2$ to afford the pure product as a yellow oil. The final product was further purified by using Teledyne ISCO ACCQPrep HP150 with a gradient of 10-100% MeCN – $H_2O$ (both solvents contain 0.1% Formic acid) to furnish 23 mg of **dALK-1** as an off-white solid in 49% yield.

$^1H$ NMR (400 MHz, DMSO-$d_6$) δ 11.15 (s, 1H), 9.45 (s, 1H), 8.46 (d, *J* = 8.4 Hz, 1H), 8.24 (s, 1H), 8.06 (s, 1H), 7.89 – 7.81 (m, 3H), 7.62 (t, *J* = 8.0 Hz, 1H), 7.51 (s, 1H), 7.35 (t, *J* = 7.6 Hz, 1H), 6.83 (s, 1H), 5.14 (dd, *J* = 13.0, 5.4 Hz, 1H), 4.57 (p, *J* = 6.1 Hz, 1H), 3.06 (d, *J* = 10.8 Hz, 2H), 2.94 – 2.82 (m, 2H), 2.57 (p, *J* = 5.8, 4.8 Hz, 5H), 2.42 (d, *J* = 7.3 Hz, 2H), 2.10 (d, *J* = 11.0 Hz, 6H), 1.67 (dq, *J* = 15.6, 8.2 Hz, 8H), 1.20 (d, *J* = 6.0 Hz, 6H), 1.15 (d, *J* = 6.8 Hz, 6H). $^{13}C$ NMR (101 MHz, DMSO-$d_6$) δ 172.8, 169.9, 166.3, 165.8, 158.0, 155.4, 154.9, 146.6, 139.4, 138.3, 138.0, 134.9, 134.7, 132.1, 131.0, 130.2, 126.7, 126.5, 124.4, 123.8, 123.7, 123.6, 122.7, 120.0, 111.6, 104.2, 98.8, 76.3, 70.7, 57.5, 54.8, 53.9, 49.0, 37.6, 32.1, 31.0, 25.9, 25.3, 21.9, 18.9, 18.4, 14.9. HRMS (ESI) calc'd for $C_{47}H_{52}ClN_7O_7S$ + H = 894.3410, found 894.3397.

Preparation of **dALK-2**

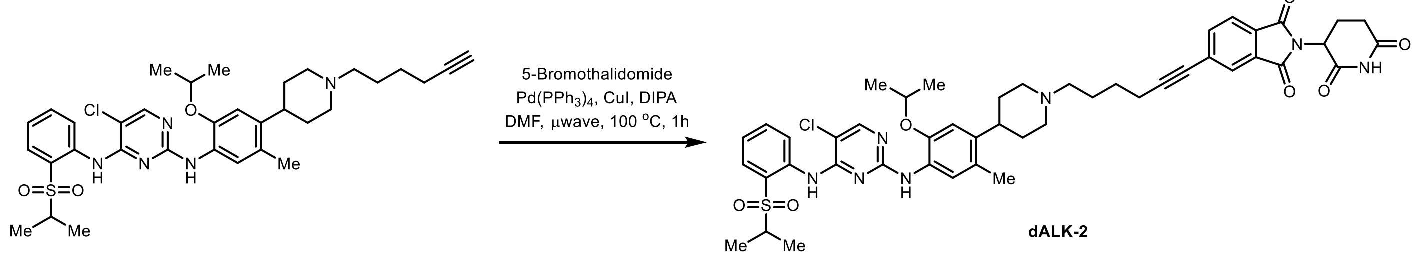

**dALK-2**

In an oven-dried 5 mL microwave vial with a stir-bar, a mixture of 5-bromothalidomide (18 mg, 0.054 mmol), $Pd(PPh_3)_4$ (6.5 mg, 0.0054 mmol), CuI (2 mg, 0.03 mmol), 5-chloro-N2-(4-(1-(hex-5-yn-1-yl)piperidin-4-yl)-2-isopropoxy-5-methylphenyl)-N4-(2-(isopropylsulfonyl)phenyl)pyrimidine-2,4-diamine **82** (45 mg, 0.07 mmol), and DIPA (16 mg, 0.16 mmol) in (2 mL) was deaerated under Argon at rt for 5 min. The vial was capped with a microwave supported rubber-septum. The reaction mixture was stirred in the microwave reactor at 100 °C for 1 h. The solvent was evaporated under reduced pressure and then extracted with EtOAc (60 mL), washed with water (10 mL × 2), followed by brine (10 mL). The organic extracts were dried over anhydrous $Na_2SO_4$, filtered, and concentrated under reduced pressure. The residue was purified by Combiflash ISCO, and the column ran with 0-10% MeOH (contains 1% $NH_4OH$)-$CH_2Cl_2$ to afford the pure product as a yellow oil. The final product was further purified by using Teledyne ISCO ACCQPrep HP150 with a gradient of 10-100% MeCN – $H_2O$ (both solvents contain 0.1% Formic acid) to furnish 26 mg of **dALK-2** as an off-white solid in 54% yield.

$^1$H NMR (400 MHz, DMSO-d6) δ 11.13 (s, 1H), 8.46 (d, *J* = 8.4 Hz, 1H), 8.23 (s, 1H), 8.02 (s, 1H), 7.61 (td, *J* = 7.9, 1.7 Hz, 1H), 7.52 (s, 1H), 7.38 – 7.30 (m, 1H), 6.83 (s, 1H), 5.15 (dd, *J* = 12.8, 5.4 Hz, 1H), 4.55 (p, *J* = 6.1 Hz, 1H), 3.43 (dd, *J* = 14.6, 7.8 Hz, 3H), 3.01 (d, *J* = 10.6 Hz, 2H), 2.89 (ddd, *J* = 16.6, 13.6, 5.3 Hz, 1H), 2.67 – 2.51 (m, 4H), 2.11 (s, 3H), 1.64 (dt, *J* = 17.6, 5.8 Hz, 8H), 1.20 (d, *J* = 6.0 Hz, 6H), 1.15 (d, *J* = 6.7 Hz, 6H). $^{13}$C NMR (101 MHz, DMSO-d6) δ 172.7, 169.8, 166.6, 166.5, 158.0 155.4, 154.8, 146.6, 139.4, 138.0, 137.4, 134.8, 131.8, 130.9, 129.8, 129.7, 126.7, 126.5, 125.6, 124.4, 123.8, 123.7, 123.6, 123.6, 111.6, 104.3, 96.1, 79.6, 70.7, 57.6, 54.9, 54.1, 49.1, 37.8, 32.3, 30.9, 25.9, 25.6, 22.0, 21.9, 18.7, 18.4, 14.9. HRMS (ESI) calc'd for $C_{47}H_{52}ClN_7O_7S$ + H = 894.3410, found 894.3396.

Preparation of 2-(4-(4-((5-chloro-4-((2-(isopropylsulfonyl)phenyl)amino)pyrimidin-2-yl)amino)-5-isopropoxy-2-methylphenyl)piperidin-1-yl)-*N*-methoxy-*N*-methylacetamide (**83**)

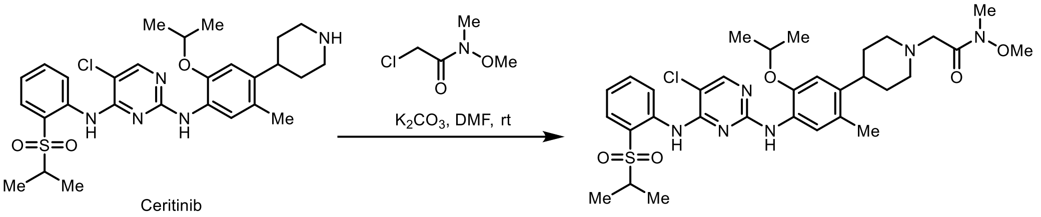

In an oven-dried 7 mL vial with a stir-bar, a mixture of ceritinib (110 mg, 0.2 mmol), dry $K_2CO_3$ (56 mg, 0.4 mmol), 2-chloro-*N*-methoxy-*N*-methylacetamide (0.4 mmol) and DMF (3 mL) were stirred at rt for overnight. The reaction mixture was evaporated under reduced pressure to remove DMF and then extracted with ethyl acetate (50 mL) water (10 mL × 2) and finally washed with brine (10 mL). Combined organic extracts were dried over anhydrous $Na_2SO_4$, filtered, and concentrated under reduced pressure. The residue was purified by Combiflash ISCO on silica gel using 5-10% MeOH in $CH_2Cl_2$ as eluent to obtain 88 mg of 2-(4-(4-((5-chloro-4-((2-(isopropylsulfonyl)phenyl)amino)pyrimidin-2-yl)amino)-5-isopropoxy-2-methylphenyl)piperidin-1-yl)-*N*-methoxy-*N*-methylacetamide (**83**) as an off-white solid in 67% yield.

$^1$H NMR (400 MHz, DMSO-$d_6$) δ 9.45 (s, 1H), 8.46 (d, *J* = 8.3 Hz, 1H), 8.24 (s, 1H), 8.02 (s, 1H), 7.83 (dd, *J* = 8.0, 1.6 Hz, 1H), 7.62 (ddd, *J* = 8.6, 7.4, 1.7 Hz, 1H), 7.51 (s, 1H), 7.35 (ddd, *J* = 8.2, 7.4, 1.1 Hz, 1H), 6.83 (s, 1H), 4.57 (p, *J* = 6.0 Hz, 1H), 3.61 (s, 3H), 3.47 – 3.41 (m, 4H), 2.96 (dt, *J* = 11.7, 3.1 Hz, 2H), 2.60 (qt, *J* = 7.5, 3.5 Hz, 3H), 2.11 (s, 3H), 2.09 – 1.95 (m, 2H), 1.64 (dd, *J* = 8.1, 3.3 Hz, 4H), 1.21 (d, *J* = 6.0 Hz, 6H), 1.16 (d, *J* = 6.8 Hz, 6H). $^{13}$C NMR (101 MHz, DMSO-d6) δ 172.5, 156.0, 155.4, 154.8, 146.6, 139.3, 138.0, 134.8, 130.9, 126.6, 126.4, 124.4, 123.7, 123.6, 123.5, 111.6, 104.2, 70.6, 54.8, 53.7, 53.5, 51.2, 39.5, 37.6, 32.3, 31.7, 21.9, 18.3, 14.8. HRMS (ESI) calc'd for $C_{32}H_{43}ClN_6O_5S$ + H = 659.2777, found 659.2769.

Preparation of 2-(4-(4-((5-chloro-4-((2-(isopropylsulfonyl)phenyl)amino)pyrimidin-2-yl)amino)-5-isopropoxy-2-methylphenyl)piperidin-1-yl)acetaldehyde (**84**)

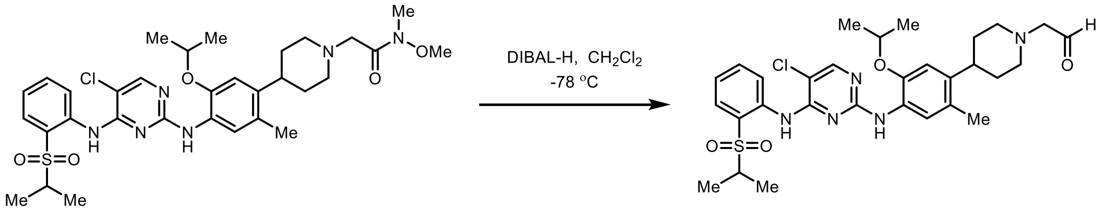

In an oven-dried 100 mL flask with a stir-bar, DIBAL-H (1.0 M in $CH_2Cl_2$, 0.57 mL, 0.57 mmol) was added dropwise to a solution of 22-(4-(4-((5-chloro-4-((2-(isopropylsulfonyl)phenyl)amino)pyrimidin-2-yl)amino)-5-isopropoxy-2-methylphenyl)piperidin-1-yl)-*N*-methoxy-*N*-methylacetamide (0.25 g, 0.38 mmol) in $CH_2Cl_2$ (10 mL) was stirred at -78 °C. The reaction was stirred at the same temperature for 30 min. At this time second portion of DIBAL-H ((1.0 M in $CH_2Cl_2$, 0.57 mL, 0.57 mmol) was added at -78 °C and the reaction was stirred at the same temperature for 1 h. The reaction mixture was quenched with Methanol (1 mL) at -78 °C followed by addition of saturated aq. Rochelle salt solution (15 mL). The resulting mixture was stirred at rt for 4 to 5 h. The separated aq. layer was extracted with $CH_2Cl_2$ (60 mL × 2). The separated combined organic layers were dried over anhydrous $Na_2SO_4$, filtered, and concentrated under reduced pressure. The 2-(4-(4-((5-chloro-4-((2-(isopropylsulfonyl)phenyl)amino)pyrimidin-2-yl)amino)-5-isopropoxy-2-methylphenyl)piperidin-1-yl)acetaldehyde (**84**) was used in the next step without any further purification.

Preparation of **dALK-3**

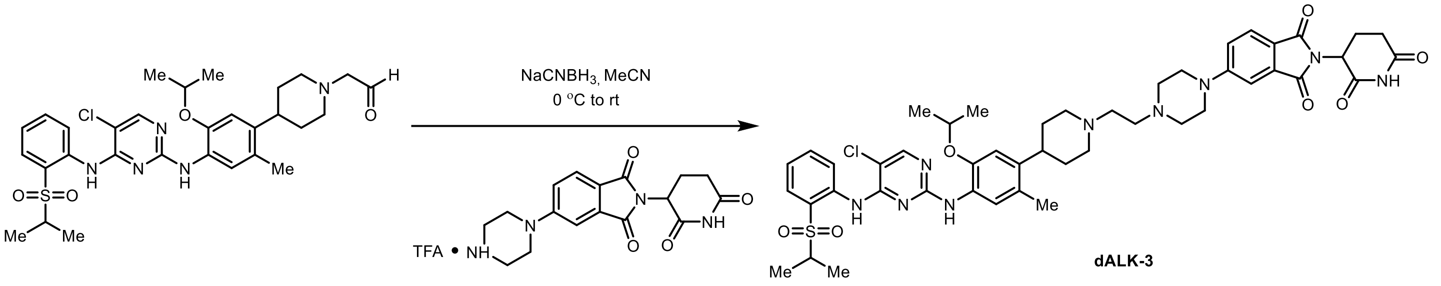

In an oven-dried 7 mL vial with a stir-bar, a solution of 2-(4-(4-((5-chloro-4-((2-(isopropylsulfonyl)phenyl)amino)pyrimidin-2-yl)amino)-5-isopropoxy-2-methylphenyl)piperidin-1-yl)acetaldehyde **84** (0.020 g, 0.033 mmol) and trfluoroacetic acid salt of 2-(2,6-dioxopiperidin-3-yl)-5-(piperazin-1-yl)isoindoline-1,3-dione **89** (0.022g, 0.04 mmol) in MeCN (3 mL) was stirred at rt for 15 min. At this time $NaCNBH_3$ (0.008 g, 0.05 mmol) was added to the reaction mixture at 0 °C and the progress of the reaction was monitored by LCMS. After completion of starting material, the reaction was quenched by adding saturated $NaHCO_3$ (2 mL) 0 °C, and then extracted with EtOAc (50 mL) and finally washed sat. $NaHCO_3$ (10 mL). The organic extracts were dried over anhydrous $Na_2SO_4$, filtered, and concentrated under reduced pressure. The residue was purified by Combiflash ISCO on silica gel using 5-10% MeOH (1% $NH_4OH$) in $CH_2Cl_2$ as eluent to obtain the pure product. The final product was further purified by using Teledyne ISCO ACCQPrep HP150 with a gradient of 10-100% MeCN – $H_2O$ (both solvents contain 0.1% Formic acid) to furnish 7 mg of **dALK-3** as a yellow colored solid in 25 % yield.

$^1H$ NMR (400 MHz, DMSO-$d_6$) δ 11.07 (s, 1H), 9.46 (s, 1H), 8.46 (d, $J$ = 8.4 Hz, 1H), 8.24 (s, 1H), 8.03 (s, 1H), 7.83 (dd, $J$ = 7.9, 1.6 Hz, 1H), 7.68 (d, $J$ = 8.5 Hz, 1H), 7.64 – 7.57 (m, 1H), 7.52 (s, 1H), 7.36 (d, $J$ = 8.0 Hz, 1H), 7.26 (dd, $J$ = 8.6, 2.3 Hz, 1H), 6.84 (s, 1H), 5.07 (dd, $J$ = 12.9, 5.4 Hz, 1H), 4.55 (q, $J$ = 6.0 Hz, 1H), 3.55 – 3.36 (m, 10H), 3.08 (s, 2H), 2.93 – 2.79 (m, 1H), 2.68 – 2.51 (m, 13H), 2.36 – 2.16 (m, 2H), 2.12 (s, 3H), 2.06 – 1.96 (m, 1H), 1.67 (d, $J$ = 8.7 Hz, 4H), 1.22 (d, $J$ = 6.0 Hz, 7H), 1.16 (d, $J$ = 6.8 Hz, 6H). $^{13}C$ NMR (101 MHz, DMSO) δ 172.7, 170.0, 167.5, 166.9, 158.0, 155.4, 155.2, 154.8, 146.6, 139.2, 138.0, 134.8, 133.8, 130.9, 126.7, 126.5, 124.8, 124.4, 123.7, 123.6, 123.5, 118.3, 117.8, 111.6, 107.8, 104.2, 79.1, 70.7, 69.8, 55.1, 54.8, 54.2, 52.6, 48.8, 46.9, 32.0, 31.0, 22.2, 21.9, 18.3, 14.8. HRMS (ESI) calc'd for $C_{47}H_{56}ClN_9O_7S$ + H = 926.3785, found 926.3777.

Preparation of **dALK-4**

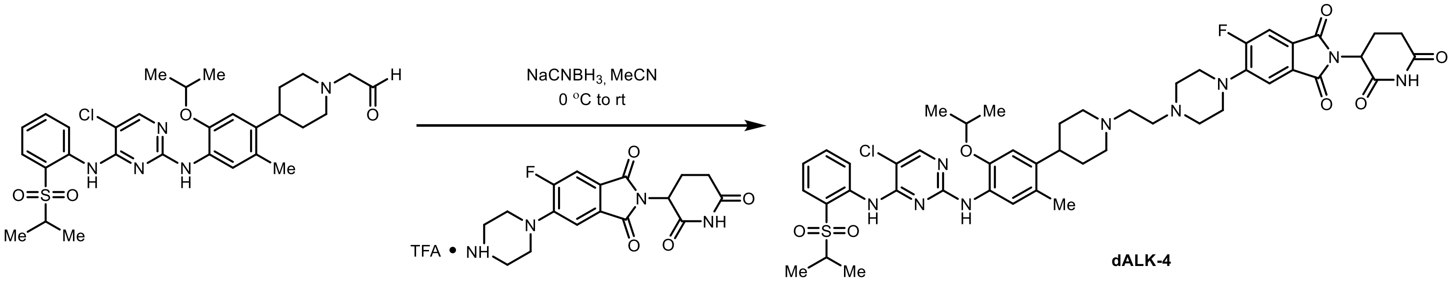

In an oven-dried 7 mL vial with a stir-bar 2-(4-(4-((5-chloro-4-((2-(isopropylsulfonyl)phenyl)amino)pyrimidin-2-yl)amino)-5-isopropoxy-2-methylphenyl)piperidin-1-yl)acetaldehyde **84** (0.020 g, 0.033 mmol) and trfluoroacetic acid salt of 2-(2,6-dioxopiperidin-3-yl)-5-fluoro-6-(piperazin-1-yl)isoindoline-1,3-dione **90** (0.022g, 0.04 mmol) in MeCN (3 mL) was stirred at rt for 15 min. At this time $NaCNBH_3$ (0.008 g, 0.05 mmol) was added to the reaction mixture at 0 °C and the progress of the reaction was monitored by LCMS. After completion of starting material, the reaction was quenched by adding saturated $NaHCO_3$ (2 mL) 0 °C. The reaction mixture was evaporated under reduced pressure to remove MeCN and then extracted with ethyl acetate (30 mL) water (5 mL × 3) and finally washed with brine (5 mL). Combined organic extracts were dried over anhydrous $Na_2SO_4$, filtered, and concentrated under reduced pressure. The residue was purified by combiflash ISCO on silica gel using 5-10% MeOH (1% $NH_4OH$) in $CH_2Cl_2$ as eluent to obtain the pure product. The final product was further purified by using Teledyne ISCO ACCQPrep HP150 with a gradient of 10-100% MeCN – $H_2O$ (both solvents contain 0.1% Formic acid) to furnish 7 mg of **dALK-4** as a yellow-colored solid in 22% yield.

$^1$H NMR (400 MHz, DMSO-d6) δ 11.10 (s, 1H), 9.46 (s, 1H), 8.47 (d, *J* = 8.4 Hz, 1H), 8.24 (s, 1H), 8.03 (s, 1H), 7.83 (dd, *J* = 8.0, 1.6 Hz, 1H), 7.72 (d, *J* = 11.4 Hz, 1H), 7.64 – 7.60 (m, 1H), 7.52 (s, 1H), 7.45 (d, *J* = 7.3 Hz, 1H), 7.39 – 7.31 (m, 1H), 6.84 (s, 1H), 5.11 (dd, *J* = 12.8, 5.4 Hz, 1H), 4.57 (p, *J* = 6.0 Hz, 1H), 3.43 (p, *J* = 6.9 Hz, 1H), 3.25 (t, *J* = 4.8 Hz, 4H), 3.05 (d, *J* = 10.5 Hz, 2H), 2.89 (ddd, *J* = 16.8, 13.7, 5.3 Hz, 1H), 2.64 – 2.57 (m, 7H), 2.52 (s, 3H), 2.12 (s, 3H), 2.10 – 1.99 (m, 2H), 1.67 (q, *J* = 8.1 Hz, 4H), 1.22 (d, *J* = 6.0 Hz, 6H), 1.16 (d, *J* = 6.7 Hz, 6H). $^{13}$C NMR (101 MHz, DMSO-d6) δ 173.2, 170.4, 167.1, 166.7, 166.6 (d, *J* = 2.4 Hz), 158.5, 157.8 (d, *J* = 260.0 Hz), 156.6, 155.9, 155.3, 147.1, 145.9, 145.8 (d, *J* = 8.9 Hz, 139.9, 138.5, 135.3, 131.4, 129.2 (d, *J* = 2.5 Hz), , 127.2, 126.9, 124.9, 124.2, 124.1 (d, *J* = 7.0 Hz), 124.0, 123.8 (d, *J* = 9.6 Hz), 114.2 (d, *J* = 2.5 Hz), 112.5, 112.3, 112.1, 104.7, 71.1, 55.9, 55.4, 55.3, 54.9, 53.3, 50.0, 50.0, 49.5, 38.1, 32.7, 31.4, 22.5, 22.4, 18.8, 15.3. HRMS (ESI) calc'd for $C_{47}H_5ClFN_9O_7S$ + H = 944.3690, found 944.3677.

Preparation of methyl 3-(4-(4-((5-chloro-4-((2-(isopropylsulfonyl)phenyl)amino)pyrimidin-2-yl)amino)-5-isopropoxy-2-methylphenyl)piperidin-1-yl)propanoate (**85**)

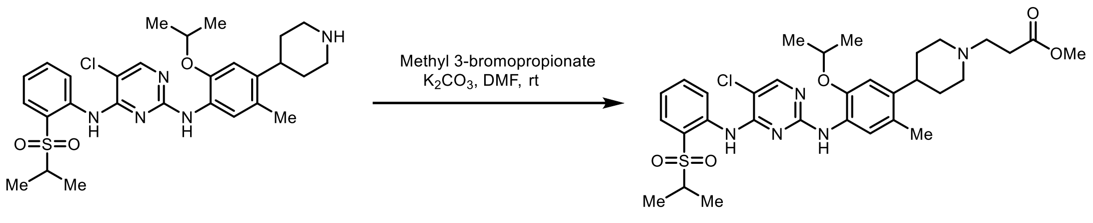

In an oven-dried 25 mL flask with a stir-bar, a mixture of ceritinib (400 mg, 0.7 mmol), dry $K_2CO_3$ (120 mg, 0.8 mmol) and methyl-3-bromopropionate (0.5 mmol) in DMF (6 mL) was stirred at room temperature for overnight, and the progress of the reaction was monitored by LCMS. The reaction mixture was evaporated under reduced pressure to remove DMF and then extracted with EtOAc (200 mL) water (40 mL × 2) and finally washed with brine (40 mL). Combined organic extracts were dried over anhydrous $Na_2SO_4$, filtered, and concentrated under reduced pressure. The residue was purified by Combiflash ISCO on silica gel using 5-10% MeOH (contains 1% $NH_4OH$) in $CH_2Cl_2$ as eluent to obtain the pure methyl 3-(4-(4-((5-chloro-4-((2-(isopropylsulfonyl)phenyl)amino)pyrimidin-2-yl)amino)-5-isopropoxy-2-methylphenyl)piperidin-1-yl)propanoate (**85**) as an off-white solid (400 mg, 86%).

$^1$H NMR (400 MHz, DMSO-d6) δ 9.45 (s, 1H), 8.46 (d, *J* = 8.3 Hz, 1H), 8.24 (s, 1H), 8.02 (s, 1H), 7.83 (dd, *J* = 8.0, 1.6 Hz, 1H), 7.51 (s, 1H), 7.39 – 7.31 (m, 1H), 6.83 (s, 1H), 4.57 (hept, *J* = 6.0 Hz, 1H), 3.61 (s, 3H), 3.43 (p, *J* = 6.8 Hz, 1H), 2.95 (d, *J* = 10.8 Hz, 2H), 2.61 (t, *J* = 7.0 Hz, 3H), 2.11 (s, 3H), 2.05 (td, *J* = 10.8, 4.7 Hz, 2H), 1.64 (dd, *J* = 8.3, 3.3 Hz, 4H), 1.21 (d, *J* = 6.0 Hz, 6H), 1.16 (d, *J* = 6.8 Hz, 6H). $^{13}$C NMR (101 MHz, DMSO-d6) δ 172.5, 158.0, 155.4, 154.8, 146.6, 139.4, 138.0, 134.8, 130.9, 126.6, 126.4, 124.4, 123.7, 123.6, 123.5, 111.6, 104.2, 70.6, 54.8, 53.7, 53.5, 51.2, 37.7, 32.3, 31.7, 21.9, 18.3, 14.8. HRMS (ESI) calc'd for $C_{32}H_{42}ClN_5O_5S$ + H = 644.2668, found 644.2670.

Preparation of 3-(4-(4-((5-chloro-4-((2-(isopropylsulfonyl)phenyl)amino)pyrimidin-2-yl)amino)-5-isopropoxy-2-methylphenyl)piperidin-1-yl)-*N*-methoxy-*N*-methylpropanamide (**86**)

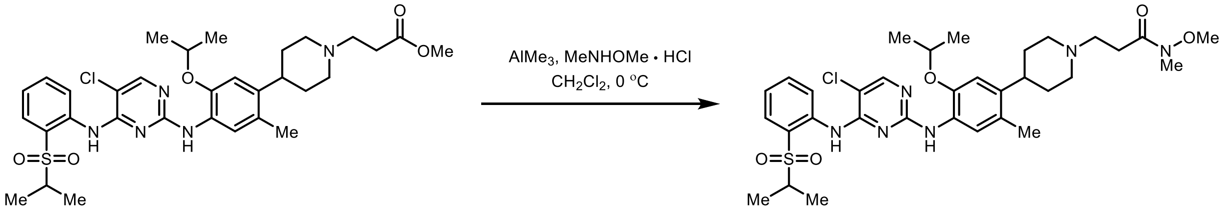

In an oven-dried 100 mL RBF with a stir-bar, $AlMe_3$ (2.0 M in hexanes, 1.74 mL, 3.4 mmol) was added dropwise to a mixture of *N*,*O*-dimethyl hydroxyl amine hydrochloride (0.34 g, 3.4 mmol) in $CH_2Cl_2$ (12 mL) and stirred at 0 °C for 1 h. At this time, a solution of methyl 3-(4-(4-((5-chloro-4-((2-(isopropylsulfonyl)phenyl)amino)pyrimidin-2-yl)amino)-5-isopropoxy-2-methylphenyl)piperidin-1-yl)propanoate **85** (0.28 g, 4.4 mmol) in $CH_2Cl_2$ (20 mL) was cannulated at 0 °C, and the reaction mixture stirred at 0 °C for 1 h, and the progress of the reaction was monitored by LCMS. After complete consumption of starting material, the reaction mixture was slowly poured in to sat'd aq. $NaHCO_3$ (60 mL) previously cooled to 0 °C, and the resulting suspension was filtered through a short pad of Celite. The filter cake was washed with $CH_2Cl_2$ (100 mL × 2), and the combined organic layers were dried over anhydrous $Na_2SO_4$, filtered, and concentrated under reduced pressure. The residue was purified by Combiflash ISCO by eluting with 5-10% MeOH (contains 1% $NH_4OH$) in $CH_2Cl_2$ as eluent to afford 0.2 g of pure 3-(4-(4-((5-chloro-4-((2-(isopropylsulfonyl)phenyl)amino)pyrimidin-2-yl)amino)-5-isopropoxy-2-methylphenyl)piperidin-1-yl)-*N*-methoxy-*N*-methylpropanamide (**86**) as an off-white foam in 68% yield.

$^1$H NMR (400 MHz, DMSO-d6) δ 9.45 (s, 1H), 8.46 (d, *J* = 8.4 Hz, 1H), 8.23 (s, 1H), 8.03 (s, 1H), 7.83 (dd, *J* = 8.0, 1.6 Hz, 1H), 7.65 – 7.57 (m, 1H), 7.51 (s, 1H), 7.34 (t, *J* = 7.6 Hz, 1H), 6.83 (s, 1H), 4.61 – 4.50 (m, 1H), 3.68 (s, 3H), 3.43 (dd, *J* = 14.3, 7.5 Hz, 2H), 3.09 (s, 3H), 2.98 (d, *J* = 10.8 Hz, 2H), 2.59 (hept, *J* = 5.4 Hz, 5H), 2.11 (s, 3H), 1.65 (q, *J* = 6.1 Hz, 4H), 1.21 (d, *J* = 5.9 Hz, 6H), 1.15 (d, *J* = 6.8 Hz, 6H). $^{13}$C NMR (101 MHz, DMSO-d6) δ 172.7, 158.5, 155.9, 155.3, 147.1, 139.9, 138.5, 135.3, 131.4, 127.1, 126.9, 124.9, 124.2, 124.1, 124.0, 112.1, 104.7, 71.1, 61.5, 55.4, 55.3, 54.4, 54.1, 38.1, 32.8, 29.6, 22.4, 18.8, 15.3. HRMS (ESI) calc'd for $C_{33}H_{45}ClN_6O_5S$ + H = 673.2933, found 673.2930.

Preparation of 3-(4-(4-((5-chloro-4-((2-(isopropylsulfonyl)phenyl)amino)pyrimidin-2-yl)amino)-5-isopropoxy-2-methylphenyl)piperidin-1-yl)propanal (**87**)

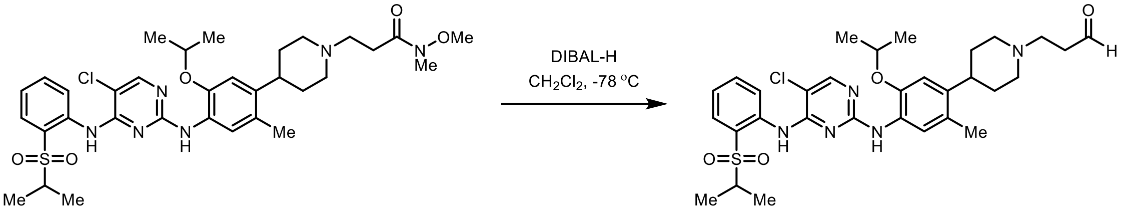

In an oven-dried 100 mL flask with a stir-bar, DIBAL-H (1.0 M in $CH_2Cl_2$, 0.65 mL, 0.65 mmol) was added dropwise to a solution of 2-(4-(4-((5-chloro-4-((2-(isopropylsulfonyl)phenyl)amino)pyrimidin-2-yl)amino)-5-isopropoxy-2-methylphenyl)piperidin-1-yl)-*N*-methoxy-*N*-methylacetamide **86** (0.2 g, 0.29 mmol) in $CH_2Cl_2$ (10 mL) was stirred at -78 °C. The reaction was stirred at the same temperature for 30 min. At this time second portion of DIBAL-H (1.0 M in $CH_2Cl_2$, 0.65 mL, 0.65 mmol) was added at -78 °C and the reaction was stirred at the same temperature for 1 h. The reaction mixture was quenched with Methanol (1 mL) at -78 °C followed by addition of saturated aq. Rochelle salt solution (10 mL). The resulting mixture was stirred at rt for 4 to 5 h. The separated aq. layer was extracted with $CH_2Cl_2$ (50 mL × 2). The separated combined organic layers were dried over anhydrous $Na_2SO_4$, filtered, and concentrated under reduced pressure. The 3-(4-(4-((5-chloro-4-((2-(isopropylsulfonyl)phenyl)amino)pyrimidin-2-yl)amino)-5-isopropoxy-2-methylphenyl)piperidin-1-yl)propanal (**87**) was used in the next step without any further purification.

Preparation of **dALK-5**

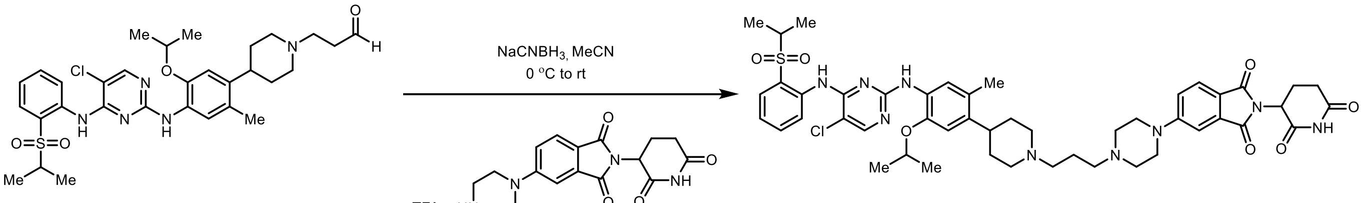

**dALK-5**

In an oven-dried 5 mL vial with stir-bar, a solution of 3-(4-(4-((5-chloro-4-((2-(isopropylsulfonyl)phenyl)amino)pyrimidin-2-yl)amino)-5-isopropoxy-2-methylphenyl)piperidin-1-yl)propanal **87** (0.017 g, 0.027 mmol) and trfluoroacetic acid salt of 2-(2,6-dioxopiperidin-3-yl)-5-(piperazin-1-yl)isoindoline-1,3-dione **89** (0.014 g, 0.033 mmol) in MeCN (2 mL) was stirred at rt for 15 min. At this time $NaCNBH_3$ (0.008 g, 0.054 mmol) was added to the reaction mixture at 0 °C and the progress of the reaction was monitored by LCMS. After completion of starting material, the reaction was quenched by adding saturated $NaHCO_3$ (2 mL) 0 °C, and then extracted with EtOAc (50 mL) and finally washed sat. $NaHCO_3$ (10 mL). The organic extracts were dried over anhydrous $Na_2SO_4$, filtered, and concentrated under reduced pressure. The residue was purified by Combiflash ISCO on silica gel using 5-10% MeOH (1% $NH_4OH$) in $CH_2Cl_2$ as eluent to obtain the pure product. The final product was further purified by using Teledyne ISCO ACCQPrep HP150 with a gradient of 10-100% MeCN – $H_2O$ (both solvents contain 0.1% Formic acid) to furnish 8 mg of **dALK-5** as a yellow-colored solid in 31 % yield.

$^1$H NMR (400 MHz, DMSO-$d_6$) δ 11.07 (s, 1H), 9.45 (s, 1H), 8.46 (d, *J* = 8.4 Hz, 1H), 8.24 (d, *J* = 1.0 Hz, 1H), 8.04 (d, *J* = 2.8 Hz, 1H), 7.83 (dt, *J* = 7.9, 1.7 Hz, 1H), 7.67 (dd, *J* = 8.6, 4.4 Hz, 1H), 7.62 (t, *J* = 8.0 Hz, 1H),

7.51 (s, 1H), 7.38 (d, $J$ = 3.9 Hz, 1H), 7.34 (dd, $J$ = 5.8, 2.8 Hz, 1H), 6.83 (d, $J$ = 9.9 Hz, 1H), 5.10 – 5.03 (m, 1H), 4.70 – 4.44 (m, 1H), 3.43 (s, 6H), 2.91 – 2.83 (m, 2H), 2.39 – 2.28 (m, 4H), 2.12 (s, 3H), 2.07 – 1.97 (m, 3H), 1.66 (m, 4H), 1.34 – 1.21 (m, 5H), 1.16 (dd, $J$ = 6.8, 1.5 Hz, 3H), 0.86 (d, $J$ = 6.9 Hz, 6H), 0.81 (d, $J$ = 6.8 Hz, 6H). $^{13}$C NMR (101 MHz, DMSO-d6) δ 172.7, 170.0, 167.5, 166.9, 158.0, 155.4, 155.2, 154.8, 146.6, 139.2, 138.0, 134.8, 133.8, 130.9, 126.7, 126.4, 124.8, 124.4, 123.7, 123.5, 118.3, 117.7, 111.6, 107.9, 104.2, 70.7, 69.8, 54.8, 54.2, 52.6, 48.8, 46.8, 32.0, 31.0, 22.2, 21.9, 18.4, 14.8. HRMS (ESI) calc'd for $C_{48}H_{58}ClN_9O_7S$ + H = 940.3577, found 940.3573.

Preparation of **dALK-6**

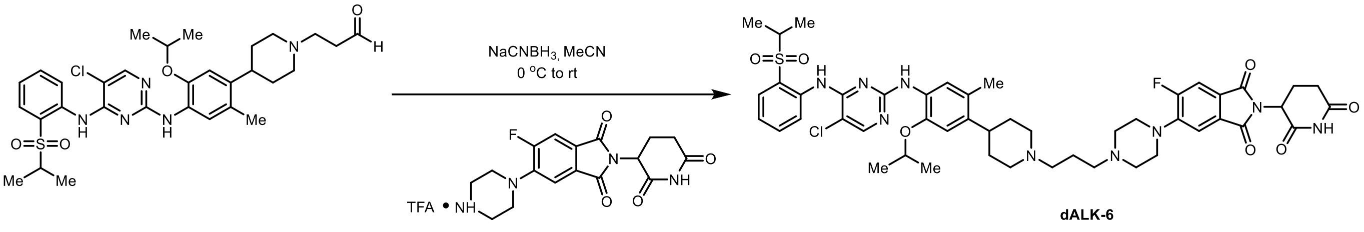

In an oven-dried 5 mL vial with stir-bar, a solution of 3-(4-(4-((5-chloro-4-((2-(isopropylsulfonyl)phenyl)amino)pyrimidin-2-yl)amino)-5-isopropoxy-2-methylphenyl)piperidin-1-yl)propanal **87** (0.017 g, 0.027 mmol) and trfluoroacetic acid salt of 2-(2,6-dioxopiperidin-3-yl)-5-fluoro-6-(piperazin-1-yl)isoindoline-1,3-dione **90** (0.015 g, 0.033 mmol) in MeCN (2 mL) was stirred at rt for 15 min. At this time $NaCNBH_3$ (0.004 g, 0.054 mmol) was added to the reaction mixture at 0 °C and the progress of the reaction was monitored by LCMS. After completion of starting material, the reaction was quenched by adding saturated $NaHCO_3$ (2 mL) 0 °C, and then extracted with EtOAc (50 mL) and finally washed sat. $NaHCO_3$ (10 mL). The organic extracts were dried over anhydrous $Na_2SO_4$, filtered, and concentrated under reduced pressure. The residue was purified by Combiflash ISCO on silica gel using 5-10% MeOH (1% $NH_4OH$) in $CH_2Cl_2$ as eluent to obtain the pure product. The final product was further purified by using Teledyne ISCO ACCQPrep HP150 with a gradient of 10-100% MeCN – $H_2O$ (both solvents contain 0.1% Formic acid) to furnish 10 mg of **dALK-6** as a yellow colored solid in 38% yield.

$^{1}$H NMR (400 MHz, DMSO) δ 11.11 (s, 1H), 9.46 (s, 1H), 8.47 (d, $J$ = 8.4 Hz, 1H), 8.25 (s, 2H), 8.05 (s, 1H), 7.84 (dd, $J$ = 8.0, 1.6 Hz, 1H), 7.73 (d, $J$ = 11.5 Hz, 1H), 7.67 – 7.58 (m, 1H), 7.52 (s, 1H), 7.46 (d, $J$ = 7.4 Hz, 1H), 7.40 – 7.32 (m, 1H), 6.85 (s, 1H), 5.11 (dd, $J$ = 12.8, 5.4 Hz, 1H), 4.58 (p, $J$ = 6.1 Hz, 1H), 3.26 (d, $J$ = 5.1 Hz, 5H), 3.04 (d, $J$ = 10.7 Hz, 2H), 2.89 (ddd, $J$ = 16.5, 13.6, 5.3 Hz, 1H), 2.64 – 2.52 (m, 7H), 2.39 (t, $J$ = 7.1 Hz, 4H), 2.13 (s, 3H), 2.05 (d, $J$ = 10.6 Hz, 4H), 1.68 (s, 6H), 1.22 (d, $J$ = 6.0 Hz, 6H), 1.17 (d, $J$ = 6.8 Hz, 6H). $^{19}$F NMR (376 MHz, DMSO-d6) δ -111.93. $^{13}$C NMR (101 MHz, DMSO-d6) δ 173.2, 170.4, 167.1, 166.7, 164.5, 159.1, 158.5, 156.5, 155.9, 155.3, 147.1, 145.9, 145.8, 139.9, 138.5, 135.3, 131.4, 129.3, 127.2, 126.9, 124.9, 124.3, 124.1, 124.1, 123.8, 112.5, 112.3, 112.1, 104.7, 71.2, 56.8, 56.5, 55.3, 54.5, 53.0, 50.0, 49.5, 38.2, 32.7 (d, $J$ = 2.8 Hz), 31.4, 24.2, 22.5, 22.4, 18.8, 15.3. HRMS (ESI) calc'd for $C_{48}H_{57}ClFN_9O_7S$ + H = 958.3483, found 958.3477.

Preparation of 2-(4-(4-((5-chloro-4-((2-(isopropylsulfonyl)phenyl)amino)pyrimidin-2-yl)amino)-5-isopropoxy-2-methylphenyl)piperidin-1-yl)acetic acid (**88**)

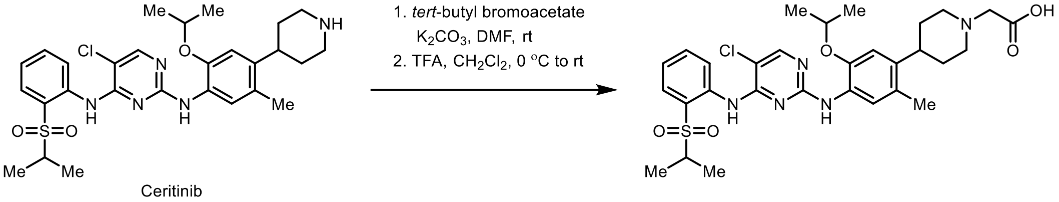

In an oven-dried 5 mL flask with a stir-bar, *tert*-butyl bromoacetate (0.078 g, 0.4 mmol) was added to a mixture of ceritinib (0.11 g, 0.2 mmol), and dry $K_2CO_3$ (0.040 mg, 0.3 mmol) in DMF (3 mL) at rt and stirred for overnight. The solvent was evaporated under reduced pressure to remove DMF and then extracted with EtOAc (60 mL), washed with water (10 mL × 2), followed by brine (10 mL). The organic extracts were dried over anhydrous $Na_2SO_4$, filtered, and concentrated under reduced pressure. The residue was purified by Combiflash ISCO, and the column ran with 0-10% MeOH (contains 1% $NH_4OH$)-$CH_2Cl_2$ to afford the pure product as an off-white foam in yield. The isolated *tert*-butyl 2-(4-(4-((5-chloro-4-((2-(isopropylsulfonyl)phenyl)amino)pyrimidin-2-yl)amino)-5-isopropoxy-2-methylphenyl)piperidin-1-yl)acetate was dissolved in dry $CH_2Cl_2$ (4 mL) and trifluoroacetic acid (0.3 mmol) was added in dropwise at 0 °C. The reaction mixture was kept stirring in room temperature and was monitored by recording LCMS. After completion of ester deprotection, the solvent and TFA were evaporated under reduced pressure to obtain the desired 2-(4-(4-((5-chloro-4-((2-(isopropylsulfonyl)phenyl)amino)pyrimidin-2-yl)amino)-5-isopropoxy-2-methylphenyl)piperidin-1-yl)acetic acid (**88**). $^1$H and $^{13}$C NMR spectral data was matched with the literature reported values.[4]

Preparation of TFA salt of 2-(2,6-dioxopiperidin-3-yl)-5-(piperazin-1-yl)isoindoline-1,3-dione (**89**)

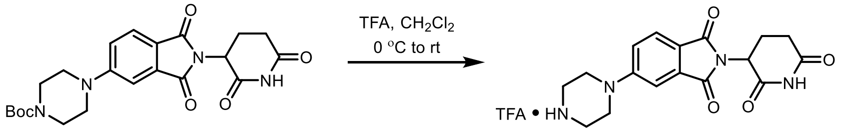

In an oven-dried 20 mL vial with a stir-bar, TFA (1.5 mL) was added dropwise to a solution of *tert*-butyl 4-(2-(2,6-dioxopiperidin-3-yl)-1,3-dioxoisoindolin-5-yl)piperazine-1-carboxylate (0.3 g, 0.68 mmol) in dry $CH_2Cl_2$ (6 mL) at 0 ˚C. The reaction was slowly warmed to rt, stirred at rt for 2 h and the progress of the reaction was monitored by LCMS. After the completion of reaction, the solvent and TFA were removed under the reduced pressure and dried under the vacuum. The resulting TFA salt of 2-(2,6-dioxopiperidin-3-yl)-5-(piperazin-1-yl)isoindoline-1,3-dione (**89**) was used in the next step without any further purification.

Preparation of **dALK-7**

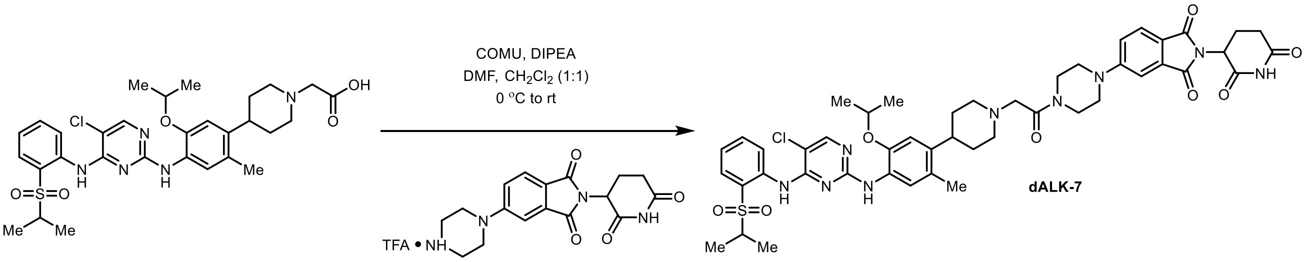

In an oven-dried 7mL vial with a stir-bar, DIPEA (0.019 g, 0.15 mmol) was added to a solution of 2-(4-(4-((5-chloro-4-((2-(isopropylsulfonyl)phenyl)amino)pyrimidin-2-yl)amino)-5-isopropoxy-2-methylphenyl)piperidin-1-yl)acetic acid **88** (0.03 g, 0.049 mmol), trfluoroacetic acid salt of 2-(2,6-dioxopiperidin-3-yl)-5-(piperazin-1-yl)isoindoline-1,3-dione **89** (0.022 g, 0.058 mmol), and COMU (0.023 g, 0.053 mmol) in 1:1 mixture of $CH_2Cl_2$ (1 mL) DMF (1 mL) at 0 °C. The mixture was kept stirring at 0 °C for 30 min. The solvent was evaporated under reduced pressure to remove DMF and then extracted with $CHCl_3$ (40 mL), washed with an ice cold sat. $NaHCO_3$ solution (10 mL). The organic extracts were dried over anhydrous $Na_2SO_4$, filtered, and concentrated under reduced pressure. The residue was purified by Combiflash ISCO, and the column ran with 0-10% MeOH (contains 1% $NH_4OH$)-$CH_2Cl_2$ to afford the pure product as a yellow colored solid. The final product was further purified by using Teledyne ISCO ACCQPrep HP150 with a gradient of 10-100% MeCN – $H_2O$ (both solvents contain 0.1% Formic acid) to furnish 11 mg of **dALK-7** as a yellow solid in 24% yield.

$^1$H NMR (400 MHz, DMSO-$d_6$) δ 11.06 (s, 1H), 9.45 (s, 1H), 8.45 (d, $J$ = 8.4 Hz, 1H), 8.24 (s, 1H), 8.04 (s, 1H), 7.82 (dd, $J$ = 8.0, 1.6 Hz, 1H), 7.70 (d, $J$ = 8.5 Hz, 1H), 7.65 – 7.55 (m, 1H), 7.51 (s, 1H), 7.34 (t, $J$ = 7.6 Hz, 1H), 7.28 (dd, $J$ = 8.5, 2.3 Hz, 1H), 6.81 (s, 1H), 5.07 (dd, $J$ = 12.9, 5.4 Hz, 1H), 4.53 (p, $J$ = 6.0 Hz, 1H), 3.76 (m, 2H), 3.68 – 3.53 (m, 4H), 3.52 – 3.39 (m, 7H), 3.06 – 2.78 (m, 3H), 2.58 (m, 4H), 2.26 – 2.13 (m, 2H), 2.12 (s, 3H), 2.06 – 1.95 (m, 1H), 1.67 (m, 3H), 1.20 (d, $J$ = 6.0 Hz, 6H), 1.15 (d, $J$ = 6.8 Hz, 6H). $^{13}$C NMR (101 MHz, DMSO-$d_6$) δ 172.8, 170.0, 168.0, 167.5, 166.9, 158.0, 155.4, 155.0, 154.8, 146.6, 139.3, 138.0, 134.8, 133.8, 130.9, 126.8, 126.6, 124.9, 124.4, 123.8, 123.7, 123.6, 118.5, 117.8, 108.0, 104.2, 70.8, 61.0, 54.8, 53.8, 48.8, 37.4, 32.4, 31.0, 22.2, 21.9, 18.4, 14.8. HRMS (ESI) calc'd for $C_{47}H_{54}ClN_9O_8S$ + H = 940.3577, found 940.3573.

Preparation of TFA salt of 2-(2,6-dioxopiperidin-3-yl)-5-fluoro-6-(piperazin-1-yl)isoindoline-1,3-dione (**90**)

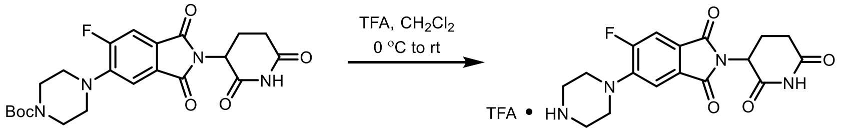

In an oven-dried 20 mL vial with a stir-bar, TFA (1 mL) was added to a solution of *tert*-butyl 4-(2-(2,6-dioxopiperidin-3-yl)-6-fluoro-1,3-dioxoisoindolin-5-yl)piperazine-1-carboxylate (0.18 g, 0.40 mmol) in dry $CH_2Cl_2$ (4 mL) at 0 °C. The reaction was slowly warmed to rt, stirred at rt for 2 h and the progress of the reaction was monitored by LCMS. After the completion of reaction, $CH_2Cl_2$ and TFA were removed under the reduced pressure and dried under the vacuum. The resulting TFA salt of 2-(2,6-dioxopiperidin-3-yl)-5-fluoro-6-(piperazin-1-yl)isoindoline-1,3-dione (**90**) was used in the next step without any further purification.

Preparation of **dALK-8**

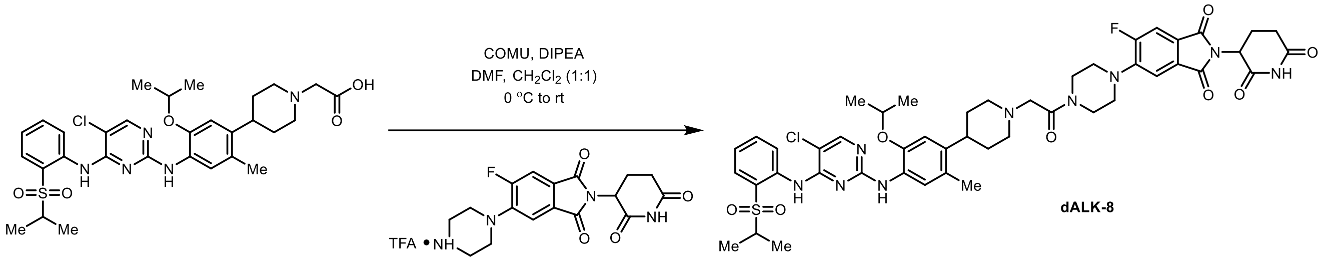

In an oven-dried 7 mL vial with a stir-bar, DIPEA (0.019 g, 0.15 mmol) was added to a solution of 2-(4-(4-((5-chloro-4-((2-(isopropylsulfonyl)phenyl)amino)pyrimidin-2-yl)amino)-5-isopropoxy-2-methylphenyl)piperidin-1-yl)acetic acid **88** (0.03 g, 0.049 mmol), TFA salt of 2-(2,6-dioxopiperidin-3-yl)-5-fluoro-6-(piperazin-1-yl)isoindoline-1,3-dione **90** (0.027g, 0.058 mmol), and COMU (0.023 g, 0.053 mmol) in 1:1 mixture of $CH_2Cl_2$ (1 mL) DMF (1 mL) at 0 °C. The mixture was kept stirring at 0 °C for 30 min. The solvent was evaporated under reduced pressure to remove DMF and then extracted with $CHCl_3$ (40 mL), washed with an ice cold sat. $NaHCO_3$ solution (10 mL). The organic extracts were dried over anhydrous $Na_2SO_4$, filtered, and concentrated under reduced pressure. The residue was purified by Combiflash ISCO, and the column ran with 0-10% MeOH (contains 1% $NH_4OH$)-$CH_2Cl_2$ to afford the pure product as a yellow-colored solid as a yellow oil. The final product was further purified by using Teledyne ISCO ACCQPrep HP150 with a gradient of 10-100% MeCN – $H_2O$ (both solvents contain 0.1% Formic acid) to furnish 10 mg of **dALK-8** as a yellow-colored solid in 20% yield.

$^1$H NMR (400 MHz, DMSO-d6) δ 11.11 (s, 1H), 9.46 (s, 1H), 8.46 (d, *J* = 8.4 Hz, 1H), 8.25 (s, 1H), 8.04 (s, 1H), 7.83 (dd, *J* = 8.0, 1.6 Hz, 1H), 7.77 (d, *J* = 11.2 Hz, 1H), 7.67 – 7.58 (m, 1H), 7.55 – 7.47 (m, 2H), 7.39 – 7.31 (m, 1H), 6.83 (s, 1H), 5.11 (dd, *J* = 12.8, 5.4 Hz, 1H), 4.54 (p, *J* = 6.0 Hz, 1H), 3.79 (s, 2H), 3.65 (s, 2H), 3.44 (p, *J* = 6.8 Hz, 1H), 3.25 (s, 4H), 2.98 (d, *J* = 10.6 Hz, 2H), 2.89 (ddd, *J* = 16.5, 13.6, 5.3 Hz, 1H), 2.65 – 2.51 (m, 2H), 2.18 (s, 3H), 2.13 (s, 3H), 2.08 – 1.97 (m, 0H), 1.70 – 1.65 (m, 4H), 1.21 (d, *J* = 6.0 Hz, 6H), 1.16 (d, *J* = 6.8 Hz, 6H). $^{19}$F NMR (376 MHz, DMSO-d6) δ -112.07. $^{13}$C NMR (101 MHz, DMSO-d6) δ 173.2, 170.3, 168.4, 167.1, 166.6 (d, *J* = 2.5 Hz), 159.1, 158.5, 156.6, 155.9, 155.3, 147.0, 145.6 (d, *J* = 8.8 Hz), 139.8, 138.5, 135.3, 131.4, 129.2 (d, *J* = 2.6 Hz), 127.3, 127.1, 124.9, 124.3, 124.2, 124.2, 124.1, 124.1, 114.5, 112.6, 112.4, 112.2, 104.7, 71.3, 61.5, 55.3, 54.3, 49.9, 49.6, 45.5, 41.5, 37.9, 32.8, 31.4, 22.3, 18.8, 15.3. HRMS (ESI) calc'd for $C_{47}H_{54}ClFN_9O_8S$ + H = 958.3483, found 958.3477.

Preparation of *tert*-butyl 4-(4-(4-((5-chloro-4-((2-(isopropylsulfonyl)phenyl)amino)pyrimidin-2-yl)amino)-5-isopropoxy-2-methylphenyl)piperidin-1-yl)butanoate (**91**)

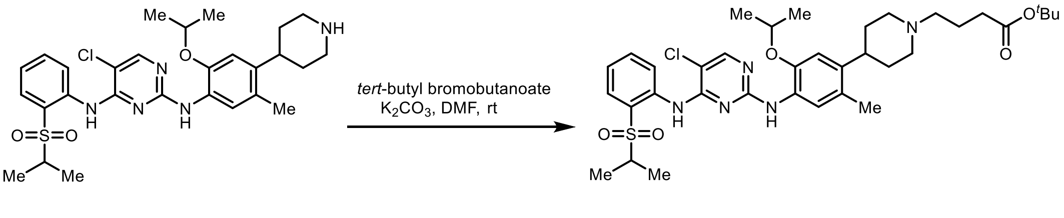

In an oven-dried 5 mL flask with a stir-bar, *tert*-butyl bromobutanoate (0.4 mmol) was added to a mixture of ceritinib (110 mg, 0.2 mmol), and dry $K_2CO_3$ (56 mg, 0.4 mmol) in DMF (3 mL). The mixture was stirred at room temperature for overnight. The solvent was evaporated under reduced pressure to remove DMF and then extracted with EtOAc (60 mL), washed with water (3x10 mL), followed by brine (10 mL). The organic extracts were dried over anhydrous $Na_2SO_4$, filtered, and concentrated under reduced pressure. The residue was purified by Combiflash ISCO, and the column ran with 0-10% MeOH (contains 1% $NH_4OH$)-$CH_2Cl_2$ to afford the pure 100 mg of *tert*-butyl 4-(4-(4-((5-chloro-4-((2-(isopropylsulfonyl)phenyl)amino)pyrimidin-2-yl)amino)-5-isopropoxy-2-methylphenyl)piperidin-1-yl)butanoate (**91**) as an off-white foam in 71% yield.

$^1$H NMR (400 MHz, DMSO-*d*$_6$) δ 9.46 (s, 1H), 8.47 (d, *J* = 8.4 Hz, 1H), 8.23 (d, *J* = 1.4 Hz, 1H), 8.01 (s, 1H), 7.83 (dd, *J* = 8.0, 1.6 Hz, 1H), 7.61 (ddd, *J* = 8.7, 7.3, 1.6 Hz, 1H), 7.53 (s, 1H), 7.34 (td, *J* = 7.6, 1.2 Hz, 1H), 6.82 (s, 1H), 4.55 (p, *J* = 6.0 Hz, 1H), 3.42 (p, *J* = 6.7 Hz, 1H), 2.97 (dt, *J* = 11.5, 3.1 Hz, 2H), 2.61 (qd, *J* = 8.0, 5.6 Hz, 1H), 2.32 (t, *J* = 7.2 Hz, 2H), 2.22 (t, *J* = 7.2 Hz, 2H), 2.11 (s, 3H), 2.03 (dt, *J* = 10.8, 7.1 Hz, 2H), 1.74 – 1.58 (m, 6H), 1.40 (d, *J* = 1.2 Hz, 9H), 1.21 (d, *J* = 6.0 Hz, 6H), 1.16 (d, *J* = 6.8 Hz, 6H). $^{13}$C NMR (101 MHz, DMSO-*d*$_6$) δ 172.7, 162.4, 157.4, 155.3, 155.2, 144.6, 138.4, 137.8, 134.6, 131.1, 127.3, 126.8, 124.7, 123.6, 123.0, 120.6, 110.8, 105.5, 80.0, 77.4, 71.3, 58.1, 55.4, 54.5, 38.1, 36.3, 33.6, 32.8, 31.3, 28.1, 22.4, 22.2, 18.8, 15.3. HRMS (ESI) calc'd for $C_{36}H_{50}ClN_5O_5S$ + H = 700.3294, found 700.3292.

Preparation of 4-(4-(4-((5-chloro-4-((2-(isopropylsulfonyl)phenyl)amino)pyrimidin-2-yl)amino)-5-isopropoxy-2-methylphenyl)piperidin-1-yl)butanoic acid (**92**)

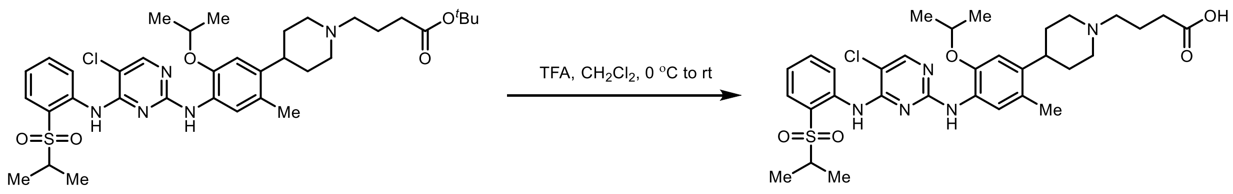

In an oven-dried 5 mL flask with a stir-bar, trifluoroacetic acid (0.3 mmol) was added dropwise to a solution of *tert*-butyl 4-(4-(4-((5-chloro-4-((2-(isopropylsulfonyl)phenyl)amino)pyrimidin-2-yl)amino)-5-isoprop oxy-2-methylphenyl)piperidin-1-yl)butanoate **91** (100 mg, 0.14 mmol) in dry $CH_2Cl_2$ (3 mL) at 0 °C. The reaction was slowly warmed to rt, stirred at rt and the progress of the reaction was monitored by LCMS. After completion of ester deprotection, the solvent and TFA were evaporated under reduced pressure to obtain the desired 4-(4-(4-((5-chloro-4-((2-(isopropylsulfonyl)phenyl)amino)pyrimidin-2-yl)amino)-5-isopropoxy-2-ethylphenyl)piperidin-1-yl)butanoic acid (**92**), and used in the next step without any further purification.

Preparation of **dALK-9**

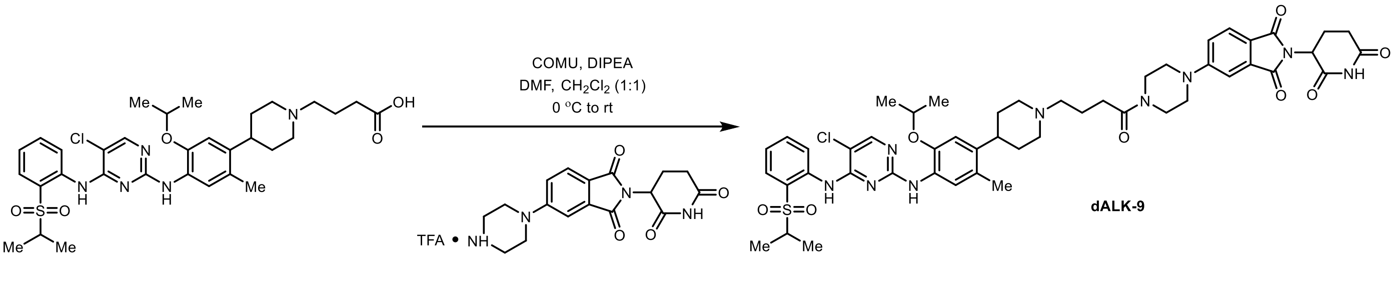

In an oven-dried 7mL vial with a stir-bar, DIPEA (0.019 g, 0.15 mmol) was added to a solution of 4-(4-(4-((5-chloro-4-((2-(isopropylsulfonyl)phenyl)amino)pyrimidin-2-yl)amino)-5-isopropoxy-2-methylphenyl)piperidin-1-yl)butanoic acid **92** (0.03g, 0.049 mmol), trfluoroacetic acid salt of 2-(2,6-dioxopiperidin-3-yl)-5-(piperazin-1-yl)isoindoline-1,3-dione **89** (0.022 g, 0.058 mmol), and COMU (0.023 g, 0.054 mmol) in 1:1 mixture of $CH_2Cl_2$ (1 mL) DMF (1 mL) at 0 °C. The mixture was kept stirring at 0 °C for 30 min. The solvent was evaporated under reduced pressure to remove DMF and then extracted with $CHCl_3$ (40 mL), washed with an ice cold sat. $NaHCO_3$ solution (10 mL). The organic extracts were dried over anhydrous $Na_2SO_4$, filtered, and concentrated under reduced pressure. The residue was purified by Combiflash ISCO, and the column ran with 0-10% MeOH (contains 1% $NH_4OH$)-$CH_2Cl_2$ to afford the pure product as a yellow colored solid. The final product was further purified by using Teledyne ISCO ACCQPrep HP150 with a gradient of 10-100% MeCN – $H_2O$ (both solvents contain 0.1% Formic acid) to furnish 10 mg of **dALK-9** as a yellow solid in 21% yield.

$^1$H NMR (400 MHz, DMSO-*d*$_6$) δ 11.08 (s, 1H), 9.45 (s, 1H), 8.45 (d, *J* = 8.4 Hz, 1H), 8.23 (s, 1H), 8.03 (s, 1H), 7.82 (dd, *J* = 8.0, 1.6 Hz, 1H), 7.69 (d, *J* = 8.5 Hz, 1H), 7.61 (ddd, *J* = 8.6, 7.3, 1.7 Hz, 1H), 7.51 (s, 1H), 7.40 – 7.29 (m, 2H), 7.23 (dd, *J* = 8.6, 2.3 Hz, 1H), 6.83 (s, 1H), 5.07 (dd, *J* = 12.9, 5.4 Hz, 1H), 4.54 (p, *J* = 6.0 Hz, 1H), 3.71 – 3.50 (m, 6H), 3.00 (d, *J* = 10.7 Hz, 1H), 2.88 (ddd, *J* = 17.3, 14.0, 5.5 Hz, 1H), 2.69 – 2.53 (m, 2H), 2.38 (q, *J* = 8.3, 7.9 Hz, 3H), 2.11 (s, 3H), 2.06 – 1.96 (m, 2H), 1.73 (t, *J* = 7.0 Hz, 1H), 1.71 – 1.62 (m, 3H), 1.20 (d, *J* = 6.0 Hz, 6H), 1.15 (d, *J* = 6.8 Hz, 6H). $^{13}$C NMR (101 MHz, DMSO-*d*$_6$) δ 172.8, 170.9, 170.1, 167.5, 167.0, 158.0, 155.4, 154.9, 146.6, 139.4, 138.0, 134.8, 133.9, 130.9, 126.7, 126.5, 124.9, 124.4, 123.8, 123.6, 123.6, 118.5, 117.7, 111.6, 107.9, 104.3, 70.7, 57.6, 54.9, 54.0, 48.8, 46.8, 46.6, 44.2, 37.7, 32.3, 31.0, 30.1, 22.2, 22.1, 21.9, 18.4, 14.9. HRMS (ESI) calc'd for $C_{49}H_{58}ClN_9O_8S$ + H = 968.3890, found 968.3879.

Preparation of **dALK-10**

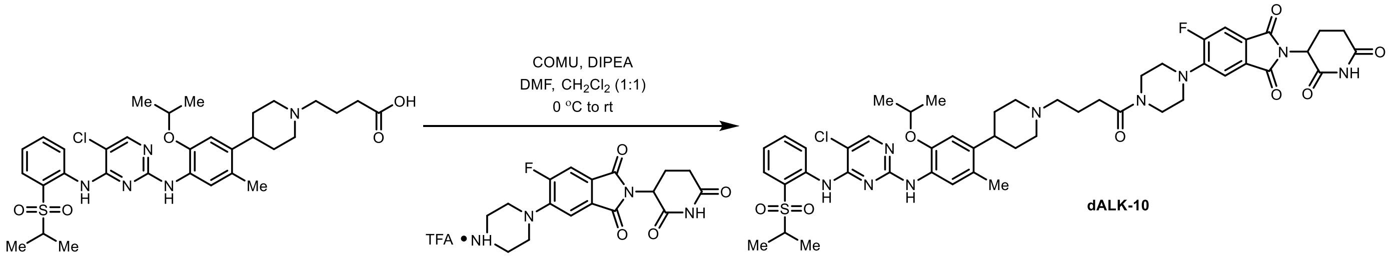

In an oven-dried 7mL vial with a stir-bar, DIPEA (0.019 g, 0.15 mmol) was added to a solution of 4-(4-(4-((5-chloro-4-((2-(isopropylsulfonyl)phenyl)amino)pyrimidin-2-yl)amino)-5-isopropoxy-2-methylphenyl)piperidin-1-yl)butanoic acid **92** (0.025 g, 0.039 mmol), trfluoroacetic acid salt of 2-(2,6-dioxopiperidin-3-yl)-5-fluoro-6-(piperazin-1-yl)isoindoline-1,3-dione **90** ( (0.022 g, 0.046 mmol), and COMU (0.018 g, 0.043 mmol) in 1:1 mixture of $CH_2Cl_2$ (1 mL) DMF (1 mL) at 0 °C. The mixture was kept stirring at 0 °C for 30 min. The solvent was evaporated under reduced pressure to remove DMF and then extracted with $CHCl_3$ (40 mL), washed with an ice cold sat. $NaHCO_3$ solution (10 mL). The organic extracts were dried over anhydrous $Na_2SO_4$, filtered, and concentrated under reduced pressure. The residue was purified by Combiflash ISCO, and the column ran with 0-10% MeOH (contains 1% $NH_4OH$)-$CH_2Cl_2$ to afford the pure product as a yellow colored solid. The final product was further purified by using Teledyne ISCO ACCQPrep HP150 with a gradient of 10-100% MeCN – $H_2O$ (both solvents contain 0.1% Formic acid) to furnish 7 mg of **dALK-10** as a yellow solid in 27% yield.

$^1$H NMR (400 MHz, DMSO) δ 11.11 (s, 1H), 9.46 (s, 1H), 8.47 (d, *J* = 8.3 Hz, 1H), 8.26 (d, *J* = 6.6 Hz, 1H), 8.05 (s, 1H), 7.84 (dd, *J* = 7.9, 1.6 Hz, 1H), 7.77 (d, *J* = 11.3 Hz, 1H), 7.62 (s, 0H), 7.51 (s, 1H), 7.49 (d, *J* = 7.3 Hz, 1H), 7.37 (d, *J* = 7.7 Hz, 1H), 6.84 (s, 1H), 5.11 (dd, *J* = 12.8, 5.4 Hz, 1H), 4.56 (p, *J* = 6.0 Hz, 1H), 3.66 (s, 4H), 3.50 – 3.40 (m, 1H), 3.24 (s, 4H), 2.99 (d, *J* = 10.7 Hz, 2H), 2.92 – 2.78 (m, 1H), 2.64 – 2.53 (m, 2H), 2.44 – 2.33 (m, 3H), 2.12 (s, 3H), 2.04 (d, *J* = 12.9 Hz, 3H), 1.73 (t, *J* = 7.1 Hz, 1H), 1.68 – 1.64 (m, 4H), 1.21 (d, *J* = 6.0 Hz, 5H), 1.16 (d, *J* = 6.8 Hz, 6H). $^{19}$F NMR (376 MHz, DMSO-d6) δ -112.16. $^{13}$C NMR (101 MHz, DMSO-d6) δ 172.7, 169.9,169.8, 166.6, 166.1 (d, *J* = 2.6 Hz), 158.0, 157.4 (d, *J* = 247.9 Hz), 156.1, 155.4, 154.8, 146.8, 145.1, 145.0, 138.0, 134.8, 130.9, 128.7 (d, *J* = 2.0 Hz), 127.2, 126.7, 124.4, 124.1, 123.8 (d, *J* = 9.8 Hz), 123.6, 114.0

(d, $J$ = 2.0 Hz), 112.0 (d, $J$ = 25.2 Hz), 111.6, 104.3, 70.9, 54.8, 53.3, 52.2, 49.5, 49.1, 44.5, 40.8, 38.3, 30.9, 29.5, 22.1, 21.9, 18.3, 18.0, 16.7, 14.8. HRMS (ESI) calc'd for $C_{49}H_{57}ClFN_9O_8S$ + H = 986.3796, found 986.3778.

Preparation of 3-(4-(4-((5-chloro-4-((2-(isopropylsulfonyl)phenyl)amino)pyrimidin-2-yl)amino)-5-isopropoxy-2-methylphenyl)piperidin-1-yl)propanoic acid (**93**)

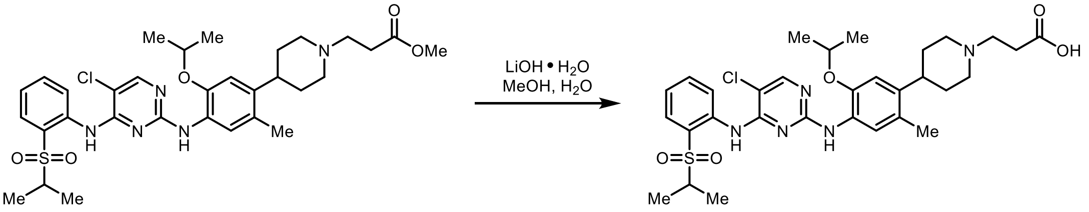

In a 5 mL flask with a stir-bar, a mixture of methyl 3-(4-(4-((5-chloro-4-((2-(isopropylsulfonyl)phenyl)amino)pyrimidin-2-yl)amino)-5-isopropoxy-2-methylphenyl)piperidin-1-yl)propanoate (80 mg, 0.12 mmol), LiOH.$H_2O$ (7 mg, 0.15 mmol), THF (2 mL), and water (0.2 mL) was stirred at 40 °C for 4 h and the progress of the reaction was monitored by LCMS. After complete consumption of starting material, the reaction was neutralized with 1N HCl (pH = 5) at 0 °C. The resulting white solids were filtered and dried under vacuum and used in the next step without any further purification.

$^1$H NMR (400 MHz, DMSO-d6) δ 9.46 (s, 1H), 8.47 (d, $J$ = 8.4 Hz, 1H), 8.29 – 8.21 (m, 2H), 8.04 (s, 1H), 7.84 (dd, $J$ = 7.9, 1.6 Hz, 1H), 7.67 – 7.58 (m, 1H), 7.53 (s, 1H), 7.36 (t, $J$ = 7.6 Hz, 1H), 6.85 (s, 1H), 4.58 (p, $J$ = 6.1 Hz, 1H), 3.44 (p, $J$ = 6.7 Hz, 1H), 3.10 (d, $J$ = 10.5 Hz, 2H), 2.71 (dt, $J$ = 15.7, 7.4 Hz, 3H), 2.42 (q, $J$ = 7.1 Hz, 2H), 2.27 (d, $J$ = 9.9 Hz, 2H), 2.13 (s, 3H), 1.70 (dt, $J$ = 11.7, 6.1 Hz, 4H), 1.22 (d, $J$ = 6.0 Hz, 6H), 1.16 (d, $J$ = 6.8 Hz, 6H). $^{13}$C NMR (101 MHz, DMSO-d6) δ 173.5, 164.1, 158.0, 155.4, 154.8, 146.6, 138.9, 138.0, 134.8, 130.9, 126.8, 126.5, 124.4, 123.7, 123.6, 123.6, 111.6, 104.3, 70.7, 54.8, 53.3, 53.1, 37.1, 31.7, 31.2, 21.9, 18.4, 14.8. HRMS (ESI) calc'd for $C_{31}H_{40}ClN_5O_5S$ + H = 630.2511, found 630.2510.

Preparation of **dALK-11**

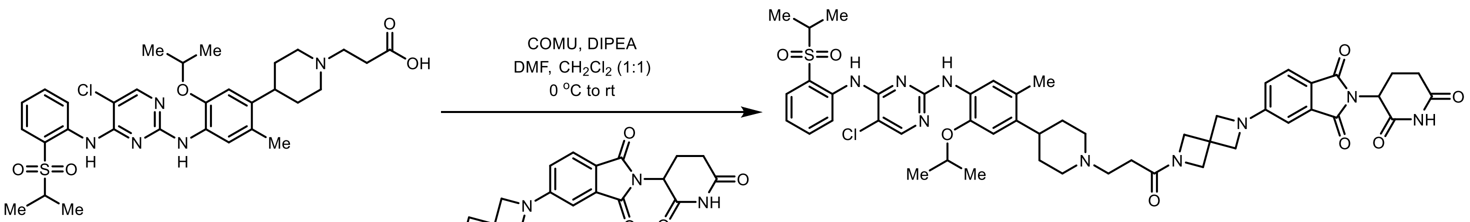

**dALK-11**

In an oven-dried 7mL vial with a stir-bar, DIPEA (0.014 g, 0.103 mmol) was added to a solution of 3-(4-(4-((5-chloro-4-((2-(isopropylsulfonyl)phenyl)amino)pyrimidin-2-yl)amino)-5-isopropoxy-2-methylphenyl)piperidin-1-yl)propanoic acid **93** (0.022 g, 0.034 mmol), 2-(2,6-dioxopiperidin-3-yl)-5-(2,6-diazaspiro[3.3]heptan-2-yl)isoindoline-1,3-dione **14** (0.015 g, 0.041 mmol), and COMU (0.016 g, 0.038 mmol) in 1:1 mixture of $CH_2Cl_2$ (1 mL) DMF (1 mL) at 0 °C. The mixture was kept stirring 0 °C for 30 min. The solvent was evaporated under reduced pressure to remove DMF and then extracted with $CHCl_3$ (40 mL), washed with an ice cold sat. $NaHCO_3$ solution (6 mL). The organic extracts were dried over anhydrous $Na_2SO_4$, filtered, and concentrated under reduced pressure. The residue was purified by Combiflash ISCO, and the column ran with 0-10% MeOH (contains 1% $NH_4OH$)-$CH_2Cl_2$ to afford the pure product as a yellow colored solid. The final product was further purified by using Teledyne ISCO ACCQPrep HP150 with a gradient of 10-100% MeCN – $H_2O$ (both solvents contain 0.1% Formic acid) to furnish 10 mg of **dALK-11** as a yellow solid in 30% yield.

$^{1}H$ NMR (400 MHz, DMSO-d6) δ 11.06 (s, 1H), 9.46 (s, 1H), 8.46 (d, *J* = 8.4 Hz, 1H), 8.24 (s, 1H), 8.05 (s, 1H), 7.83 (dd, *J* = 7.9, 1.6 Hz, 1H), 7.64 (dd, *J* = 11.5, 7.7 Hz, 2H), 7.51 (s, 1H), 7.39 – 7.31 (m, 1H), 6.86 – 6.78 (m, 2H), 6.67 (dd, *J* = 8.3, 2.1 Hz, 1H), 5.06 (dd, *J* = 12.8, 5.4 Hz, 1H), 4.57 (p, *J* = 6.0 Hz, 1H), 4.20 (s, 4H), 4.07 (s, 2H), 3.43 (p, *J* = 6.7 Hz, 1H), 2.99 (s, 1H), 2.94 – 2.79 (m, 1H), 2.57 (td, *J* = 9.6, 3.8 Hz, 4H), 2.25 (t, *J* = 7.4 Hz, 2H), 2.12 (s, 3H), 2.11 – 1.97 (m, 1H), 1.66 (t, *J* = 5.1 Hz, 5H), 1.22 (d, *J* = 6.0 Hz, 6H), 1.16 (d, *J* = 6.7 Hz, 6H). $^{13}C$ NMR (101 MHz, DMSO-d6) δ 172.8, 171.3, 170.1, 167.4, 167.1, 158.0, 155.4, 154.8, 154.8, 146.7, 139.4, 138.0, 134.8, 133.8, 130.9, 126.6, 126.4, 124.8, 124.4, 123.8, 123.6, 123.5, 117.3, 114.6, 111.6, 104.8, 104.2, 70.6, 61.3, 59.5, 57.5, 54.9, 53.9, 48.7, 37.6, 32.6, 32.3, 31.0 28.9, 22.2, 21.9, 18.3, 14.8. HRMS (ESI) calc'd for $C_{49}H_{56}ClN_9O8S$ + H = 966.3734, found 966.3724.

Preparation of 2-(2,6-dioxopiperidin-3-yl)-5-fluoro-6-(2,6-diazaspiro[3.3]heptan-2-yl)isoindoline-1,3-dione (**94**)

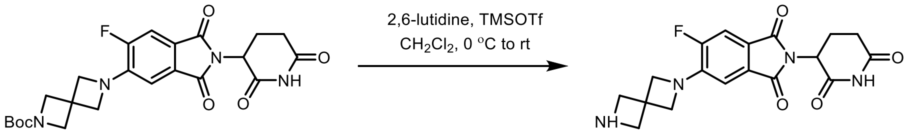

In an oven-dried 7 mL vial with a stir-bar, TMSOTf (0.14 g, 0.63 mmol) was added to the solution of *tert*-butyl 6-(2-(2,6-dioxopiperidin-3-yl)-6-fluoro-1,3-dioxoisoindolin-5-yl)-2,6-diazaspiro[3.3]heptane-2-carboxylate **48** (0.15 g, 0.13 mmol) and 2,6-lutidine (0.081 g, 0.76 mmol) in $CH_2Cl_2$ (4 mL) at 0 °C. The reaction was slowly warmed to rt and at stirred for 2 h. At this time MeOH (0.5 mL) was added and stirred at rt for 10 min. The reaction was cooled to rt and the solvent was evaporated. The mixture was kept stirring at room temperature and the progress of the reaction was monitored by LCMS. After complete consumption of starting material, the reaction was evaporated under reduced pressure and the material was used in the next step without any further purification.

Preparation of **dALK-12**

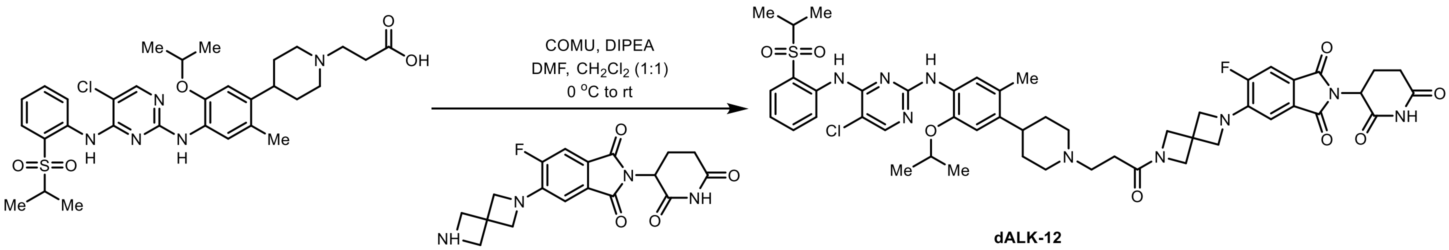

**dALK-12**

In an oven-dried 7mL vial with a stir-bar, DIPEA (0.014 g, 0.109 mmol) was added to a solution of 3-(4-(4-((5-chloro-4-((2-(isopropylsulfonyl)phenyl)amino)pyrimidin-2-yl)amino)-5-isopropoxy-2-methylphenyl)piperidin-1-yl)propanoic acid **93** (0.022 g, 0.034 mmol), 2-(2,6-dioxopiperidin-3-yl)-5-fluoro-6-(2,6-diazaspiro[3.3]heptan-2-yl)isoindoline-1,3-dione **94** (0.015 g, 0.041 mmol), and COMU (0.016 g, 0.038 mmol) in 1:1 mixture of $CH_2Cl_2$ (1 mL) DMF (1 mL) at 0 °C. The mixture was kept stirring at 0 °C for 30 min. The solvent was evaporated under reduced pressure to remove DMF and then extracted with $CHCl_3$ (40 mL), washed with an ice cold sat. $NaHCO_3$ solution (6 mL). The organic extracts were dried over anhydrous $Na_2SO_4$, filtered, and concentrated under reduced pressure. The residue was purified by Combiflash ISCO, and the column ran with 0-10% MeOH (contains 1% $NH_4OH$)-$CH_2Cl_2$ to afford the pure product as a yellow colored solid. The final product was further purified by using Teledyne ISCO ACCQPrep HP150 with a gradient of 10-100% MeCN – $H_2O$ (both solvents contain 0.1% Formic acid) to furnish 12 mg of **dALK-12** as a yellow solid in 35% yield.

$^{1}$H NMR (400 MHz, DMSO-d6) δ 11.08 (s, 1H), 9.46 (s, 1H), 8.47 (d, *J* = 8.4 Hz, 1H), 8.25 (s, 1H), 8.05 (s, 1H), 7.84 (dd, *J* = 8.0, 1.6 Hz, 1H), 7.63 (dd, *J* = 9.2, 5.8 Hz, 2H), 7.52 (s, 1H), 7.36 (t, *J* = 7.6 Hz, 1H), 6.94 (d, *J* = 7.6 Hz, 1H), 6.84 (s, 1H), 5.07 (dd, *J* = 12.8, 5.4 Hz, 1H), 4.57 (p, *J* = 6.0 Hz, 1H), 4.38 – 4.31 (m, 6H), 4.06 (s, 2H), 3.44 (p, *J* = 6.8 Hz, 1H), 2.99 (d, *J* = 10.5 Hz, 2H), 2.88 (ddd, *J* = 16.7, 13.6, 5.3 Hz, 1H), 2.63 – 2.52 (m, 3H), 2.26 (t, *J* = 7.2 Hz, 2H), 2.13 (s, 3H), 2.05 – 2.03 (m, 1H), 1.71 – 1.61 (m, 5H), 1.22 (d, *J* = 6.1 Hz, 6H), 1.17 (d, *J* = 6.8 Hz, 6H).$^{19}$F NMR (376 MHz, DMSO-d6) δ -127.49. $^{13}$C NMR (101 MHz, DMSO-d6) δ 173.2, 171.7, 170.4, 167.2, 166.8 (d, *J* = 2.7 Hz), 158.5, 155.9, 155.3 154.4 (d, *J* = 252.5 Hz), 147.1, 144.2 (d, *J* = 11.8 Hz), 139.9, 138.5, 135.3, 131.4, 129.7, 129.7, 127.1, 126.9, 124.9, 124.3, 124.1(d, *J* = 7.4 Hz),119.4 (d, *J* = 8.1 Hz), 112.1, 111.8 (d, *J* = 21.9 Hz), 108.8 (d, *J* = 6.1 Hz), 104.7, 71.1, 63.5, 59.9, 57.9, 55.3, 54.3, 54.1, 49.4, 38.1, 33.7 (d, *J* = 2.8 Hz), 32.7, 31.4, 29.3, 22.4, 18.8, 15.3. HRMS (ESI) calc'd for $C_{49}H_{55}ClFN_9O8S$ + H = 986.3796, found 986.3778.

Preparation of *tert*-butyl (2-((2-(2,6-dioxopiperidin-3-yl)-1,3-dioxoisoindolin-5-yl)amino)ethyl)carbamate **(95)**

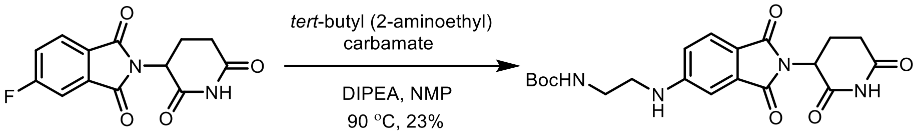

In an oven-dried 7 mL vial with a stir-bar, DIPEA (0.31 mmol, 2.5 equiv.) was added to a solution of 2-(2,6-dioxopiperidin-3-yl)-5-fluoroisoindoline-1,3-dione (65 mg, 0.24 mmol, 1.0 equiv.), and *tert*-butyl (2-aminoethyl)carbamate (0.26 mmol 1.3 equiv.) in dry *N*-methyl pyrrolidine (1.5 mL) at rt. The reaction mixture was stirred at 90 °C for 14 h. The reaction mixture was diluted with ethyl acetate (60 mL), washed with water (10 mL x3), followed by Brine (10 mL), dried over anhydrous $Na_2SO_4$, filtered, and concentrated under reduced pressure. The crude residue was purified by Combiflash ISCO to obtain the pure product and the column ran with $CH_2Cl_2$, then grading to 5% MeOH (contains 1% $NH_4OH$)-$CH_2Cl_2$ to afford *tert*-butyl (2-((2-(2,6-dioxopiperidin-3-yl)-1,3-dioxoisoindolin-5-yl)amino)ethyl)carbamate **(95)** in 23% yield (23 mg; yellow solid).

$^{1}$H NMR (400 MHz, DMSO-*d*$_6$) δ 11.07 (s, 1H), 7.56 (d, *J* = 8.3 Hz, 1H), 7.15 (t, *J* = 5.9 Hz, 1H), 6.97 (d, *J* = 2.1 Hz, 1H), 6.93 (t, *J* = 5.8 Hz, 1H), 6.86 (dd, *J* = 8.4, 2.1 Hz, 1H), 5.03 (dd, *J* = 12.9, 5.4 Hz, 1H), 3.23 (q, *J* = 6.4 Hz, 2H), 3.10 (q, *J* = 6.4 Hz, 2H), 2.87 (ddd, *J* = 17.5, 14.0, 5.4 Hz, 1H), 2.63 – 2.51 (m, 2H), 2.05 – 1.93 (m, 1H), 1.37 (s, 9H). $^{13}$C NMR (100 MHz, DMSO-d6) δ 172.9, 170.2, 167. 7, 167.1, 155.7, 154.4 134.2, 125.1, 116.2, 77.8, 54.9, 48.6, 42.2, 31.0, 28.2, 22.3. HRMS (ESI) calc'd for $C_{20}H_{24}N_4O_6$ + Na = 439.1594, found 439.1586.

Preparation of HCl salt of 5-((2-aminoethyl)amino)-2-(2,6-dioxopiperidin-3-yl)isoindoline-1,3-dione (**96**)

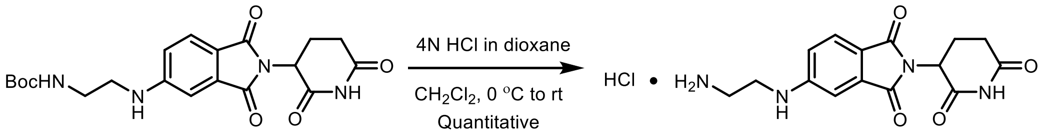

In an oven-dried 7 mL vial with a stir-bar, 4N HCl in dioxane (1.5 mL) was added dropwise to a solution of *tert*-butyl 4-(2-(2,6-dioxopiperidin-3-yl)-1,3-dioxoisoindolin-5-yl)piperazine-1-carboxylate **95** (12 mg, 0.029 mmol) in dry $CH_2Cl_2$ (3 mL) at 0 ˚C. The reaction was slowly warmed to rt, stirred at rt for overnight and the progress of the reaction was monitored by LCMS. After the completion of reaction, the solvents were removed under the reduced pressure, and dried under the vacuum. The resulting HCl salt of 5-((2-aminoethyl)amino)-2-(2,6-dioxopiperidin-3-yl)isoindoline-1,3-dione (**96**) was used in the next step without any further purification.

Preparation of **(5)-MS4078**

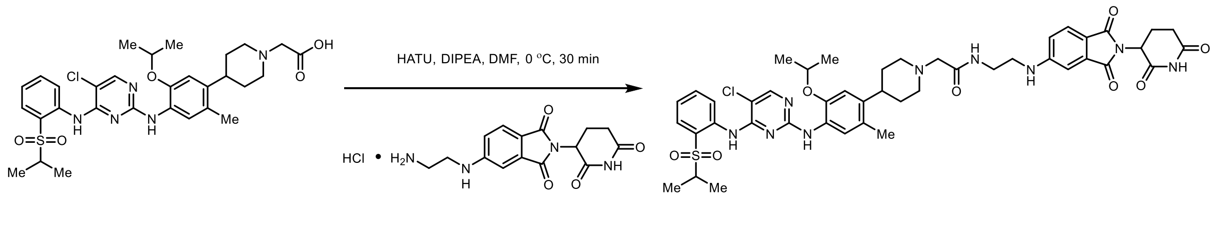

In an oven-dried 7 mL vial with a stir-bar, DIPEA (19 mg, 0.15 mmol, 5.0 equiv.) was added to a solution of 2-(4-(4-((5-chloro-4-((2-(isopropylsulfonyl)phenyl)amino)pyrimidin-2-yl)amino)-5-isopropoxy-2-methylphenyl)piperidin-1-yl)acetic acid **88** (13.5 mg, 0.029 mmol, 1.0 equiv.), HCl salt of 5-((2-aminoethyl)amino)-2-(2,6-dioxopiperidin-3-yl)isoindoline-1,3-dione **96** ( 0.029 mmol, 1.0 equiv.), and HATU (13.2 mg, 0.035 mmol, 1.2 equiv.) in DMF (2 mL) at 0 °C, and the mixture was stirred at 0 °C for 30 min. The solvent was evaporated under reduced pressure to remove DMF and then residue was purified by using Teledyne ISCO ACCQPrep HP150 with a gradient of 10-100% MeCN – $H_2O$ (both solvents contain 0.1% Formic acid) to furnish 6.8 mg of **(5)-MS4078** as an off-white colored solid in 26% yield.

$^1H$ NMR (400 MHz, DMSO-d6) δ 11.04 (s, 1H), 9.46 (s, 1H), 8.46 (d, J = 8.4 Hz, 1H), 8.25 (s, 1H), 8.06 (s, 1H), 8.01 – 7.89 (m, 1H), 7.83 (dd, J = 8.0, 1.6 Hz, 1H), 7.62 (ddd, J = 8.6, 7.3, 1.6 Hz, 1H), 7.55 (d, J = 8.4 Hz, 1H), 7.52 (s, 1H), 7.38 – 7.31 (m, 1H), 7.20 (t, J = 5.3 Hz, 1H), 7.02 (d, J = 2.1 Hz, 1H), 6.90 (dd, J = 8.4, 2.1 Hz, 1H), 6.83 (s, 1H), 5.01 (dd, J = 12.9, 5.4 Hz, 1H), 4.52 (p, J = 6.1 Hz, 1H), 3.54 – 3.42 (m, 1H), 2.94 (s, 2H), 2.93 – 2.77 (m, 3H), 2.71 – 2.54 (m, 2H), 2.16 (ddd, J = 11.5, 9.1, 2.6 Hz, 2H), 2.11 (s, 3H), 1.93 (dp, J = 11.1, 3.7, 3.3 Hz, 1H), 1.82 – 1.56 (m, 3H), 1.24 (d, J = 6.1 Hz, 6H), 1.16 (d, J = 6.8 Hz, 6H). $^{13}C$ NMR (101 MHz, DMSO-d6) δ 172.7, 170.1, 169.9, 167.6, 167.1, 158.0, 155.4, 154.8, 154.4, 146.5, 139.3, 138.0, 134.8, 134.2, 130.9, 126.8, 126.7, 125.0, 124.5, 123.8, 123.6, 123.6, 116.2, 111.7, 104.3, 94.7, 70.9, 70.5, 66.3, 61.8, 54.8, 54.3, 48.6, 41.9, 39.5, 37.3, 32.2, 30.9, 21.9, 18.3, 14.8. HRMS (ESI) calc'd for $C_{45}H_{52}ClN_9O_8S$ + H = 914.3426, found 914.3421.

[1]H and [13]C NMR of Entry **16**

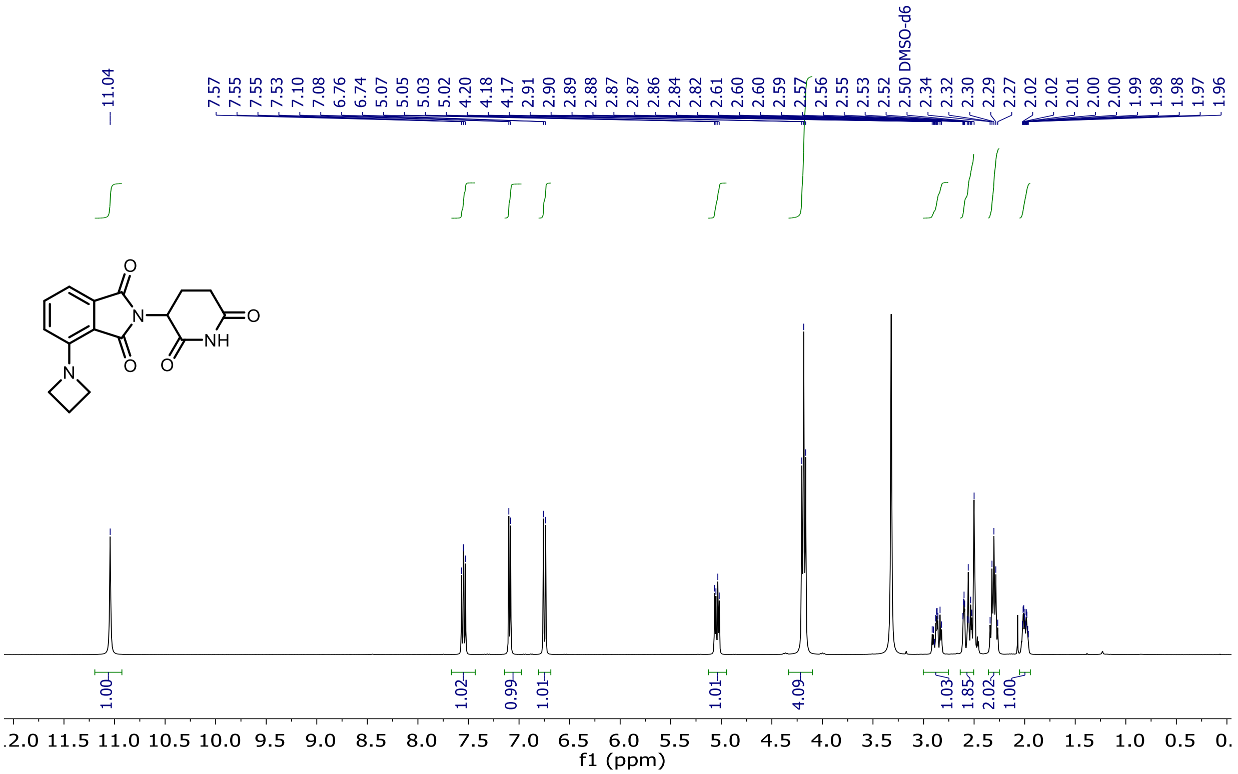

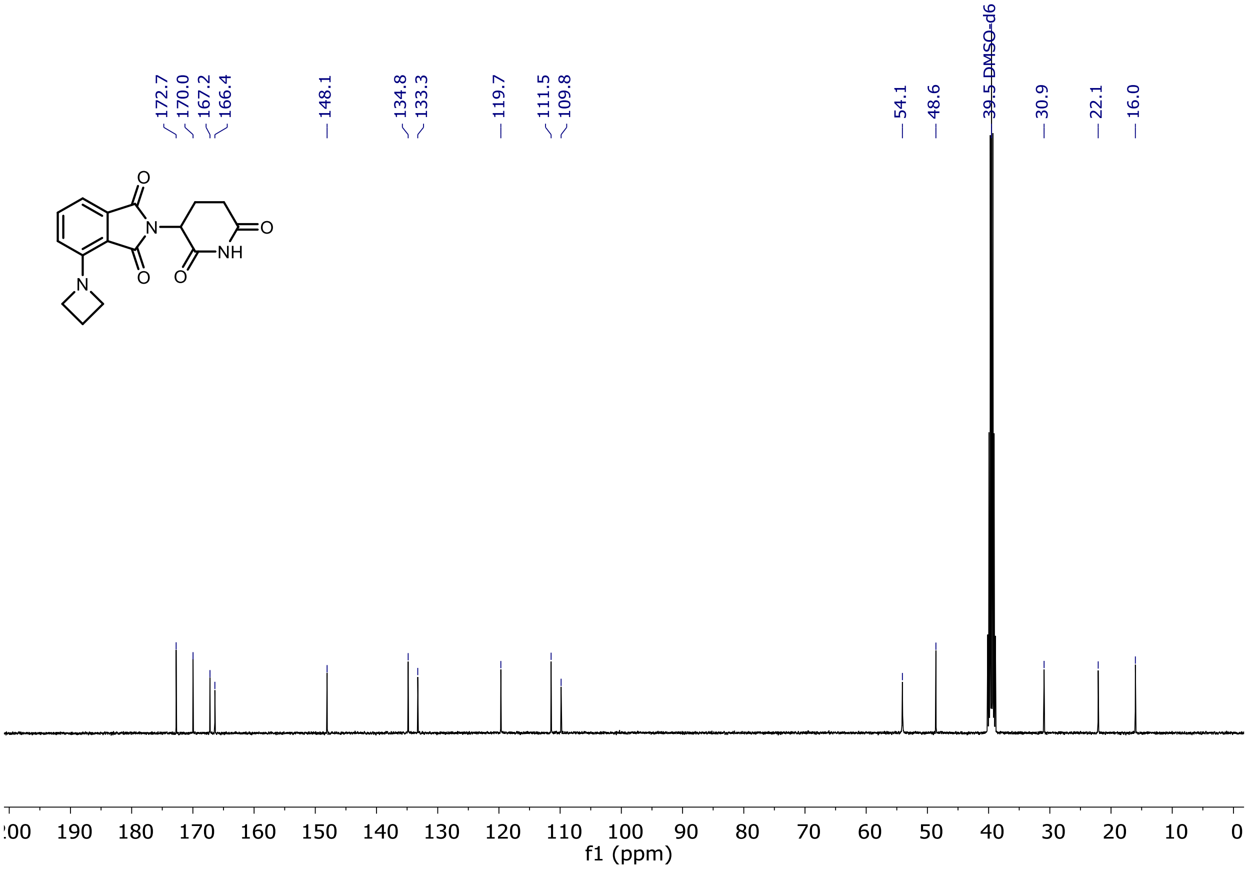

$^{1}$H and $^{13}$C NMR of Entry **17**

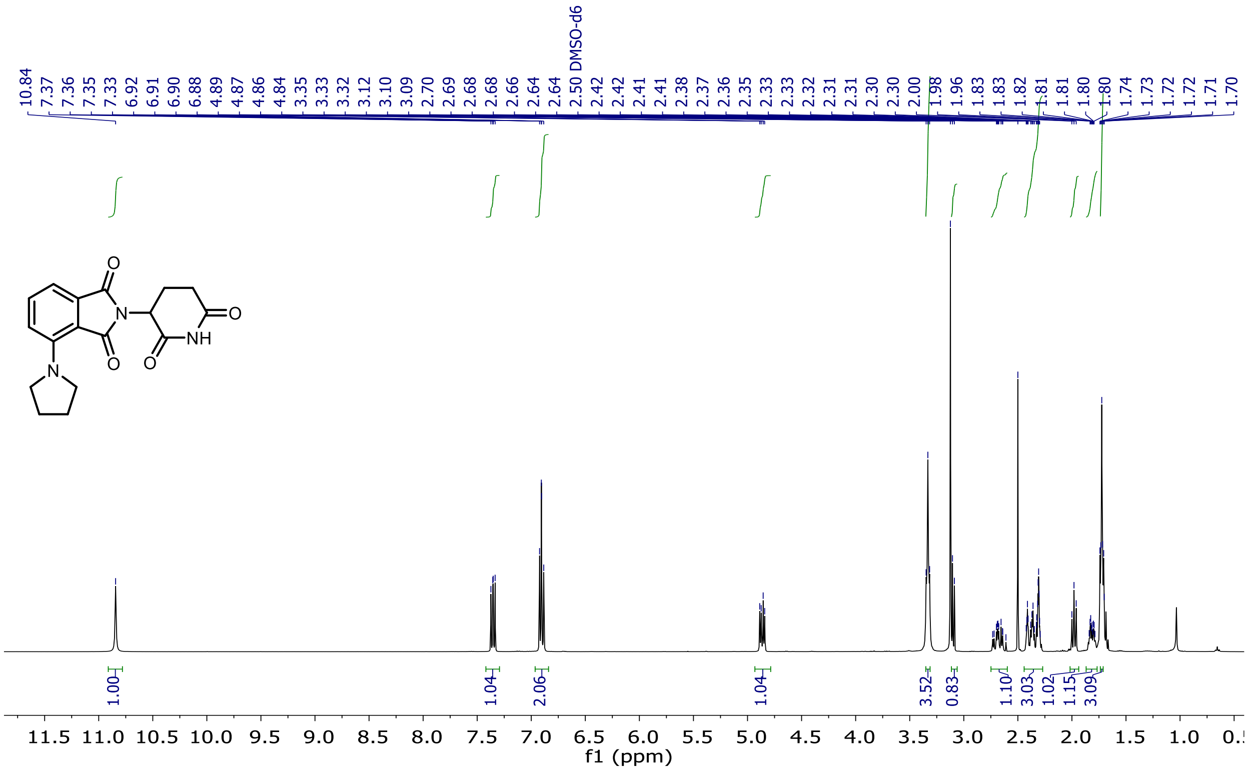

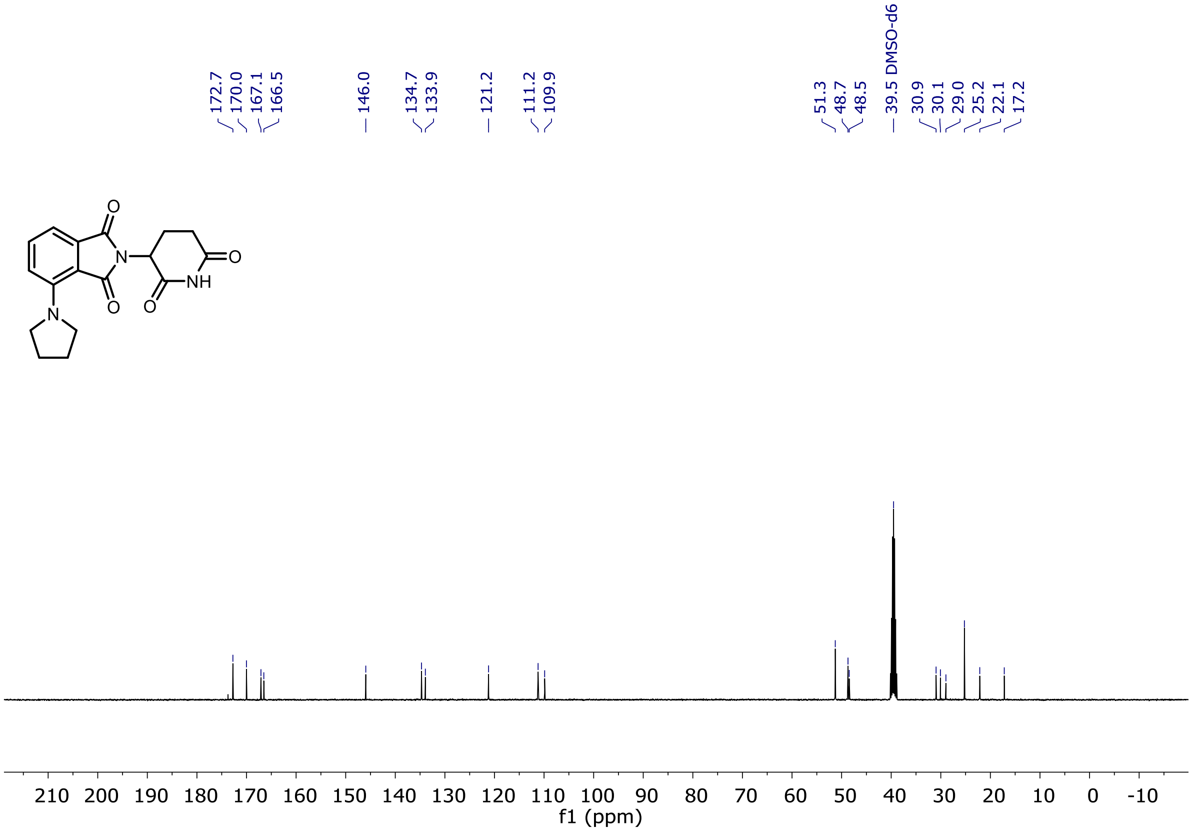

[1]H and [13]C NMR of Entry **18**

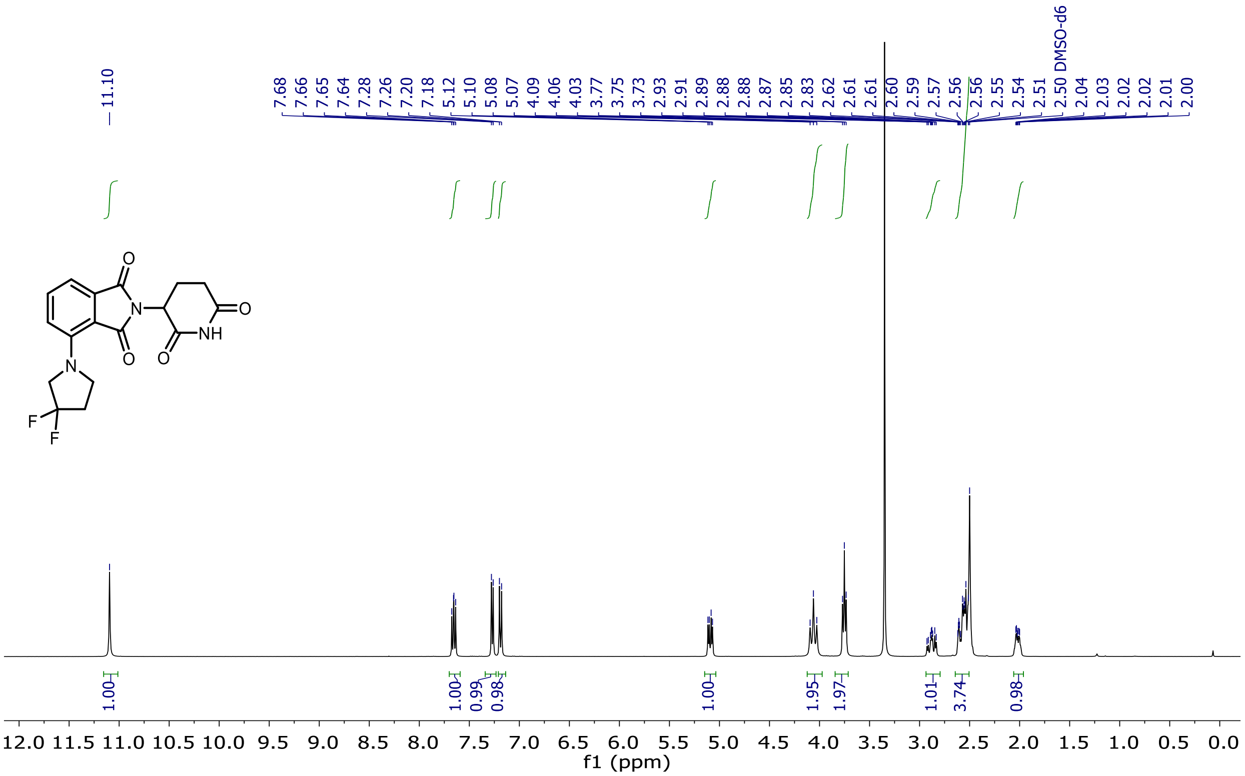

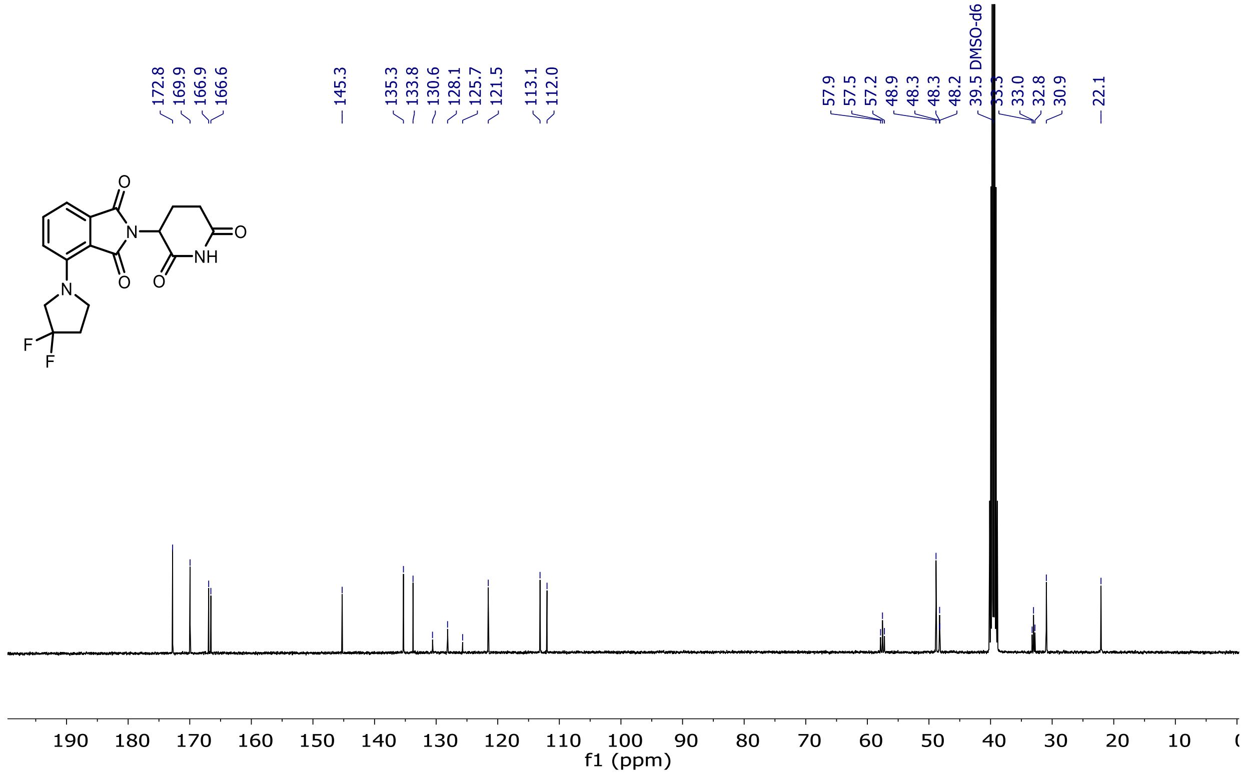

$^{19}$F NMR of Entry **18**

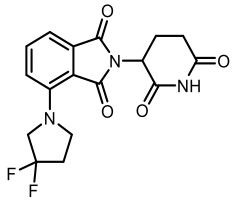

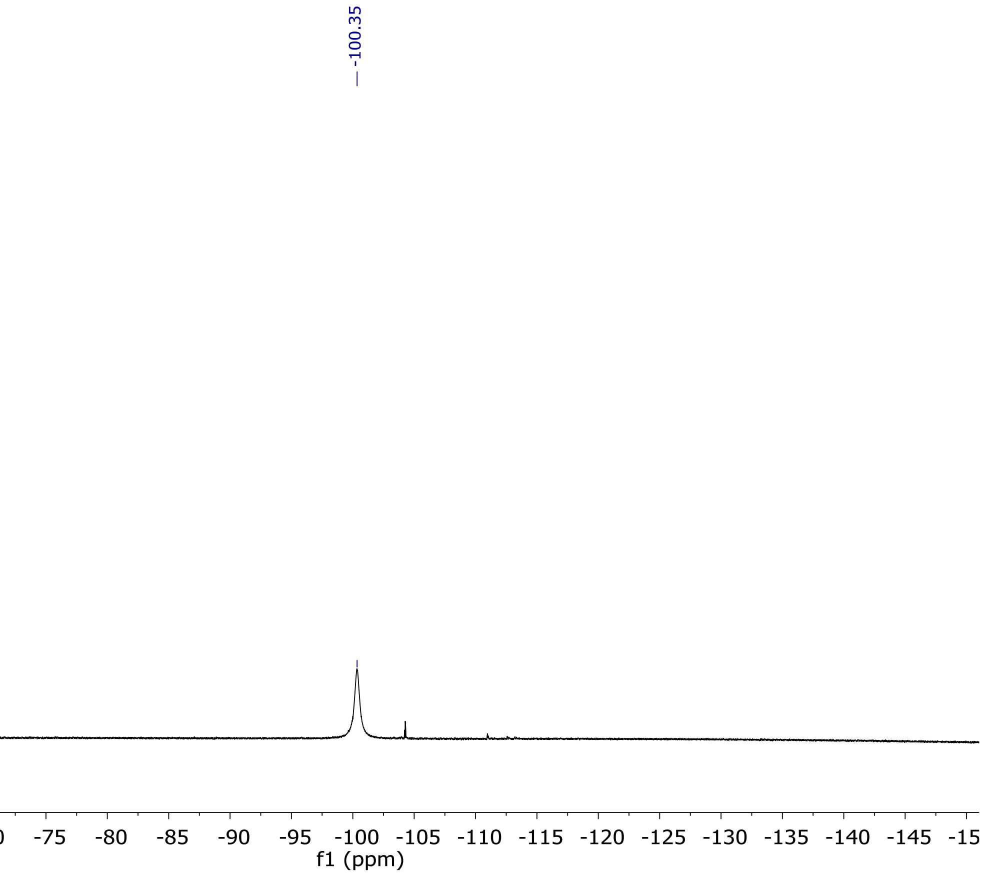

[1]H and [13]C NMR of Entry **19**

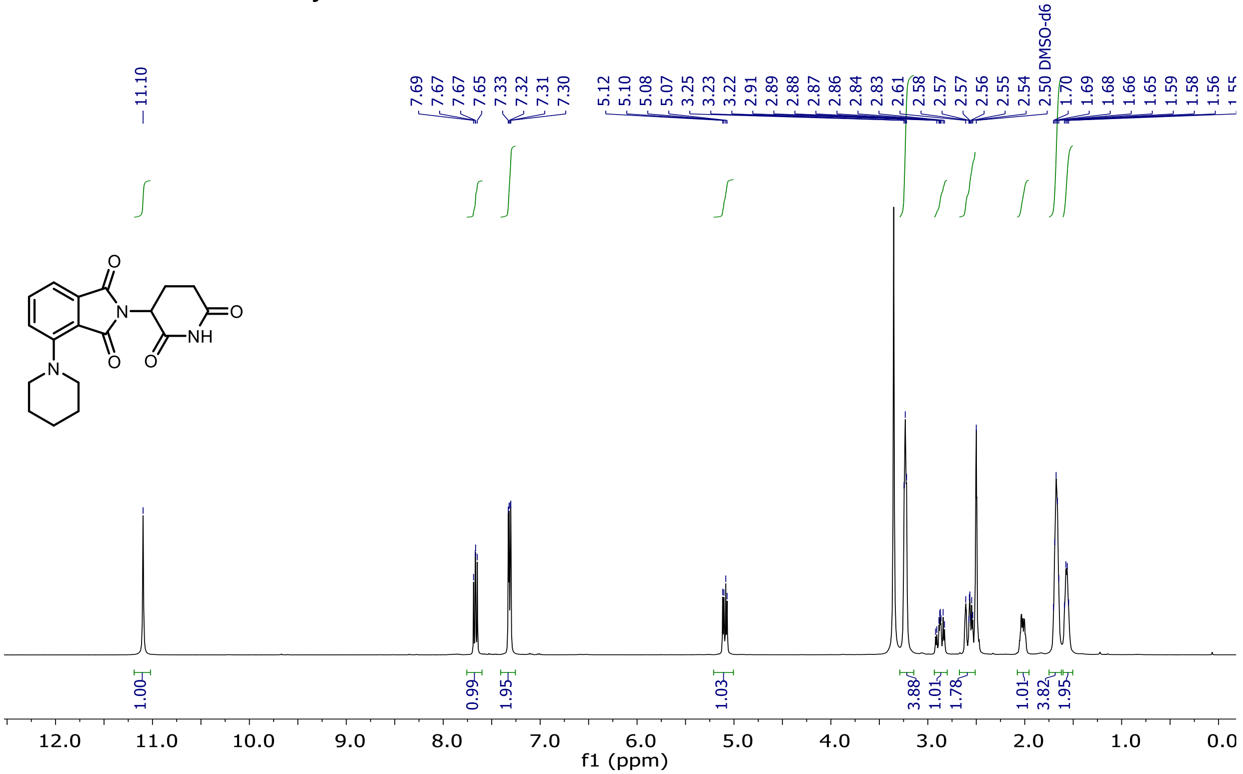

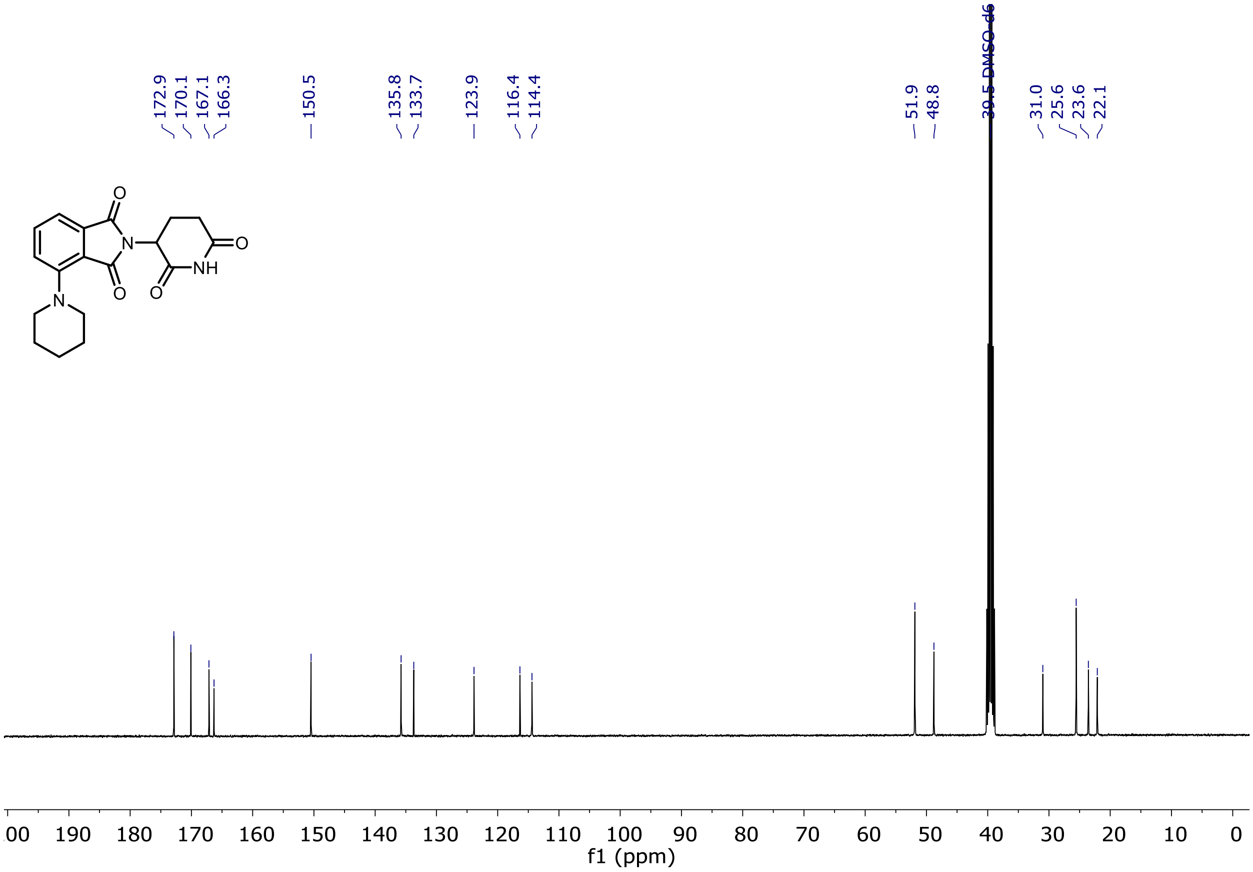

$^{1}H$ and $^{13}C$ NMR of Entry **20**

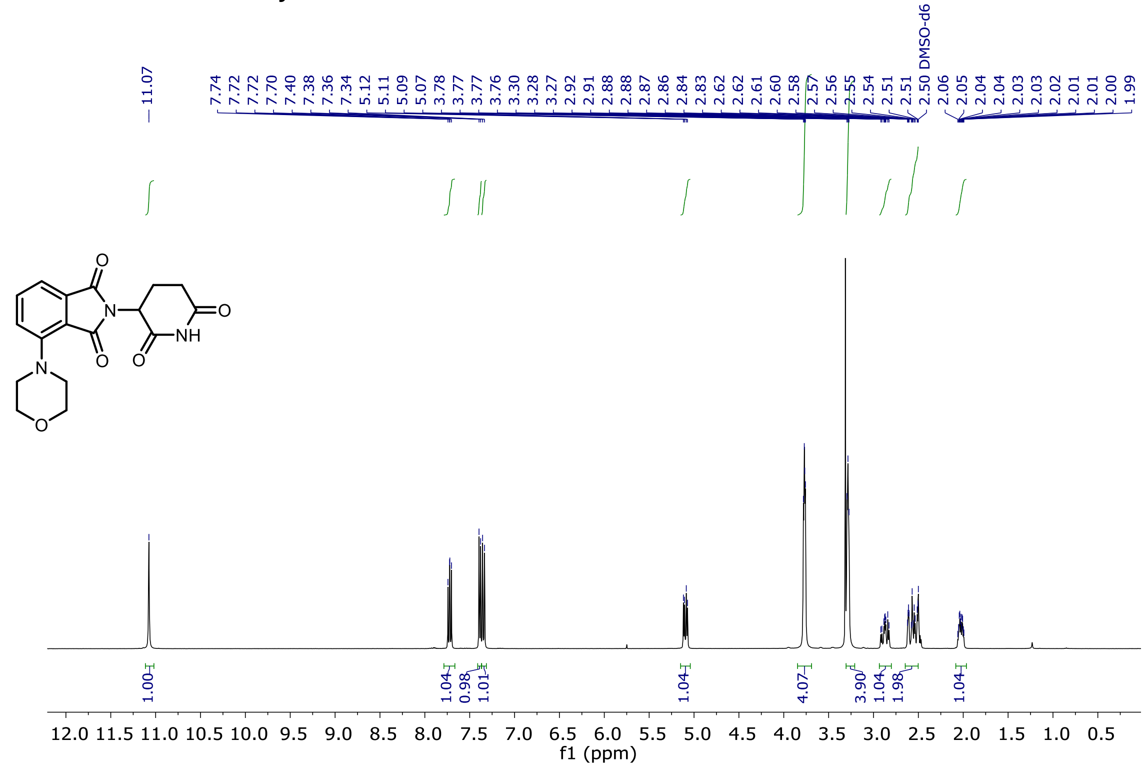

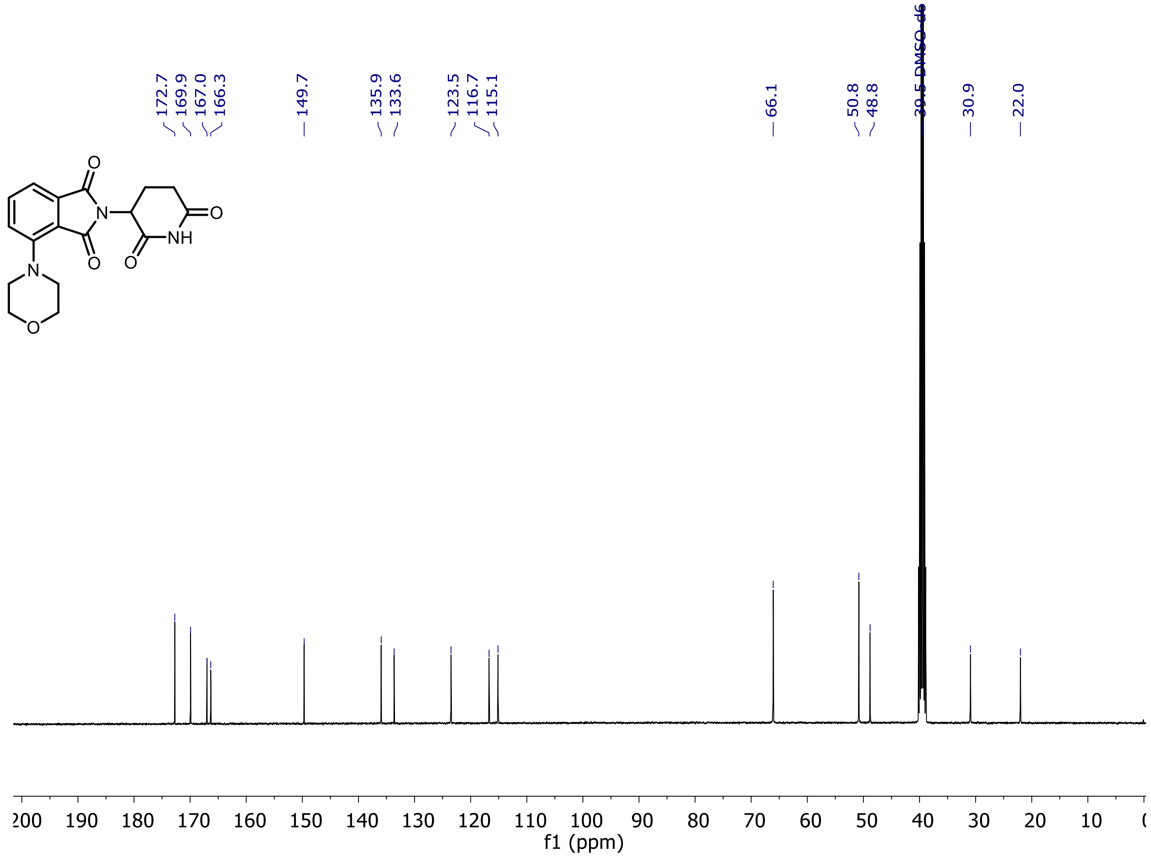

$^{1}H$ and $^{13}C$ NMR of Entry **21**

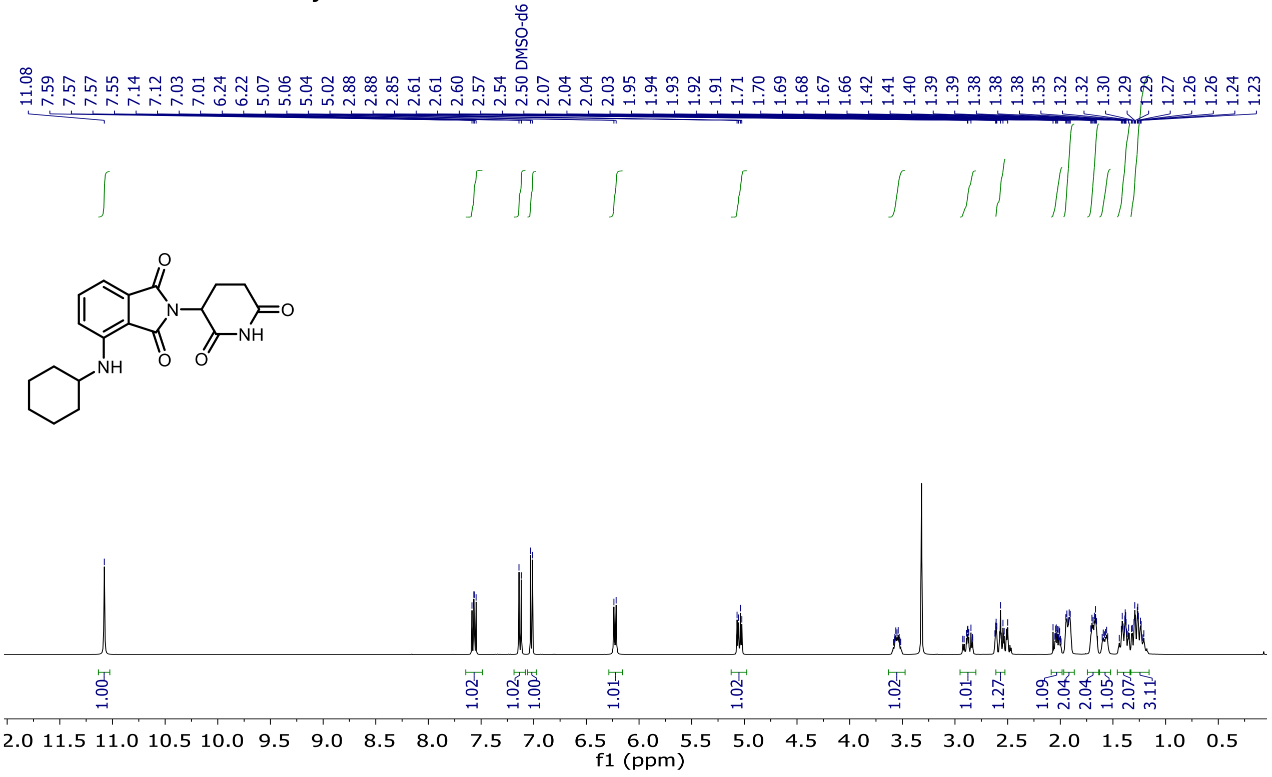

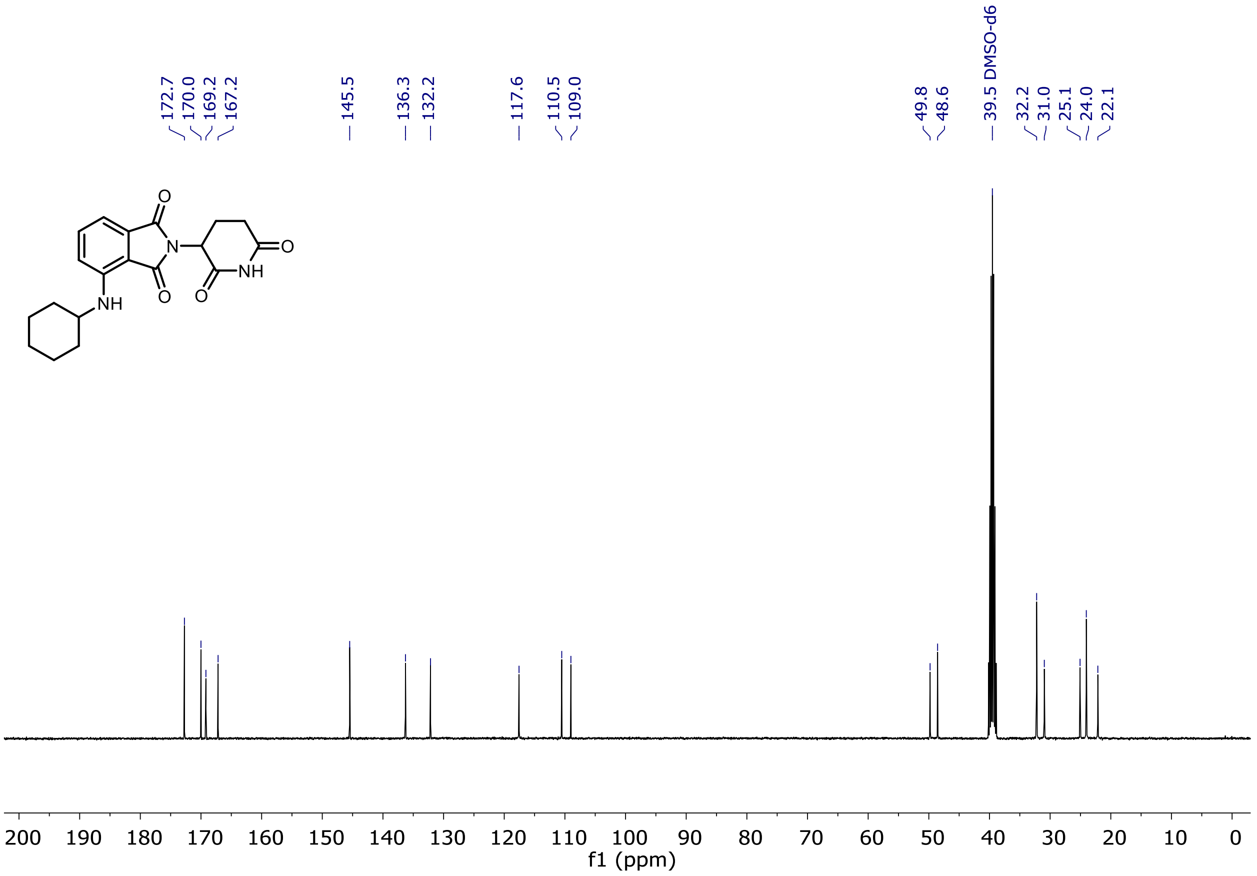

[1]H and [13]C NMR of Entry **22**

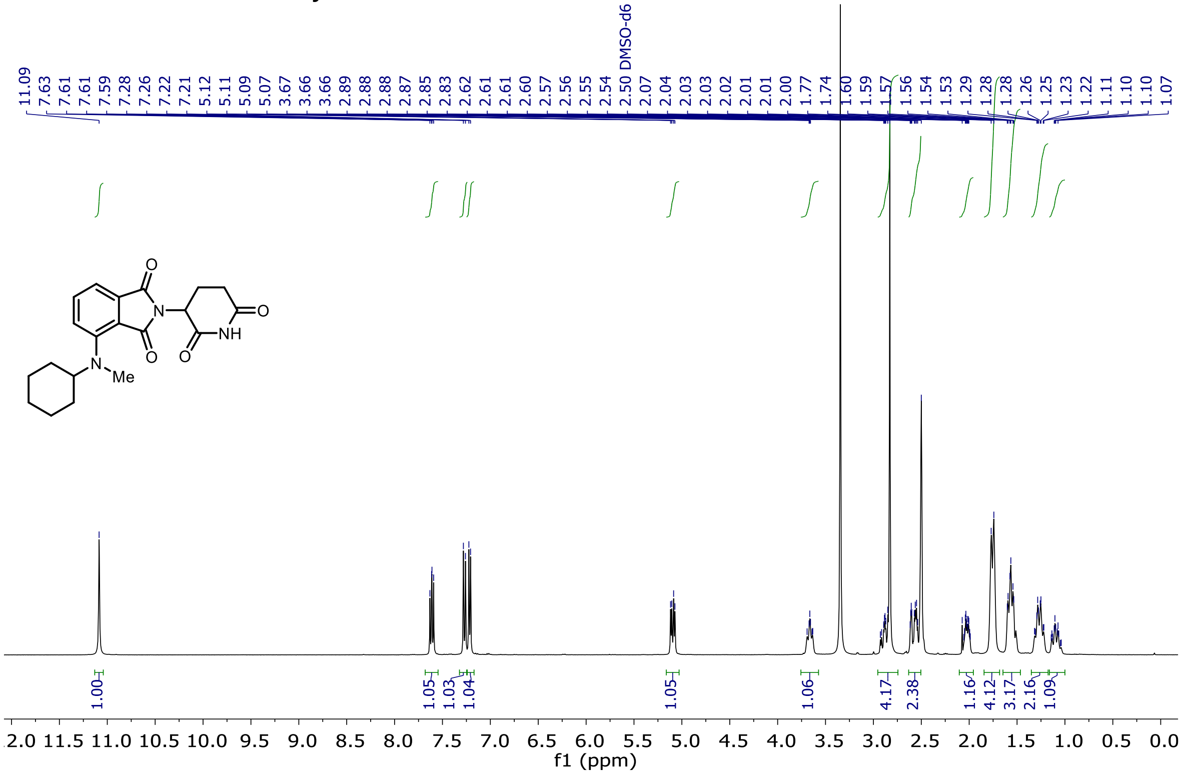

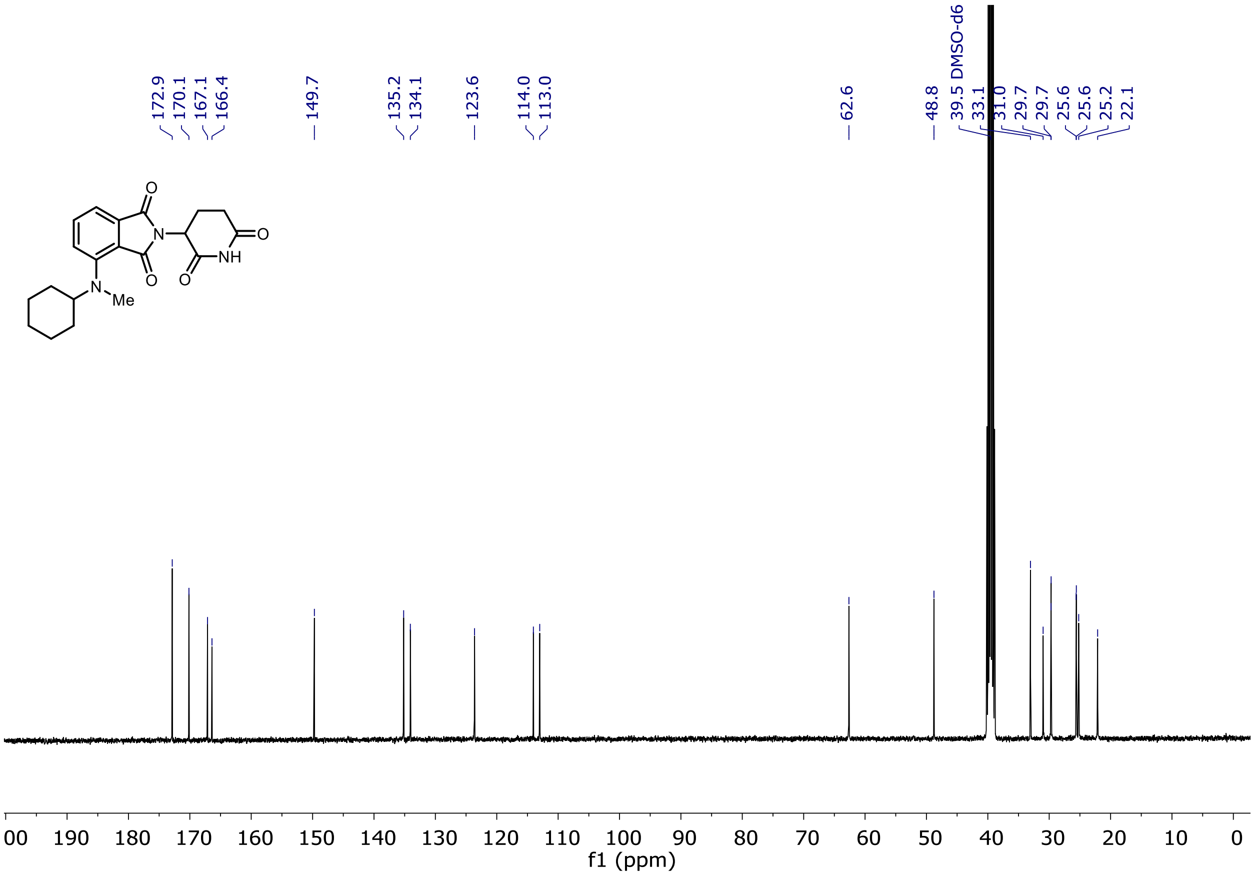

$^{1}H$ and $^{13}C$ NMR of Entry **23**

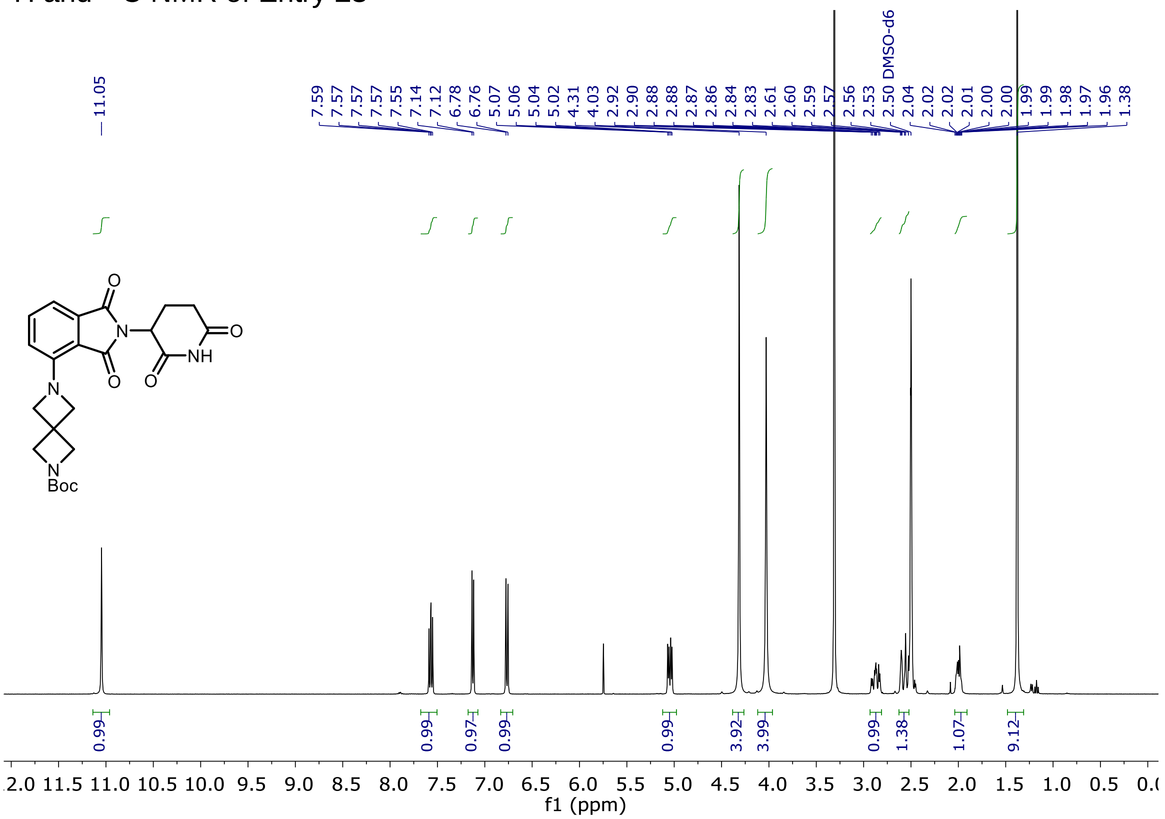

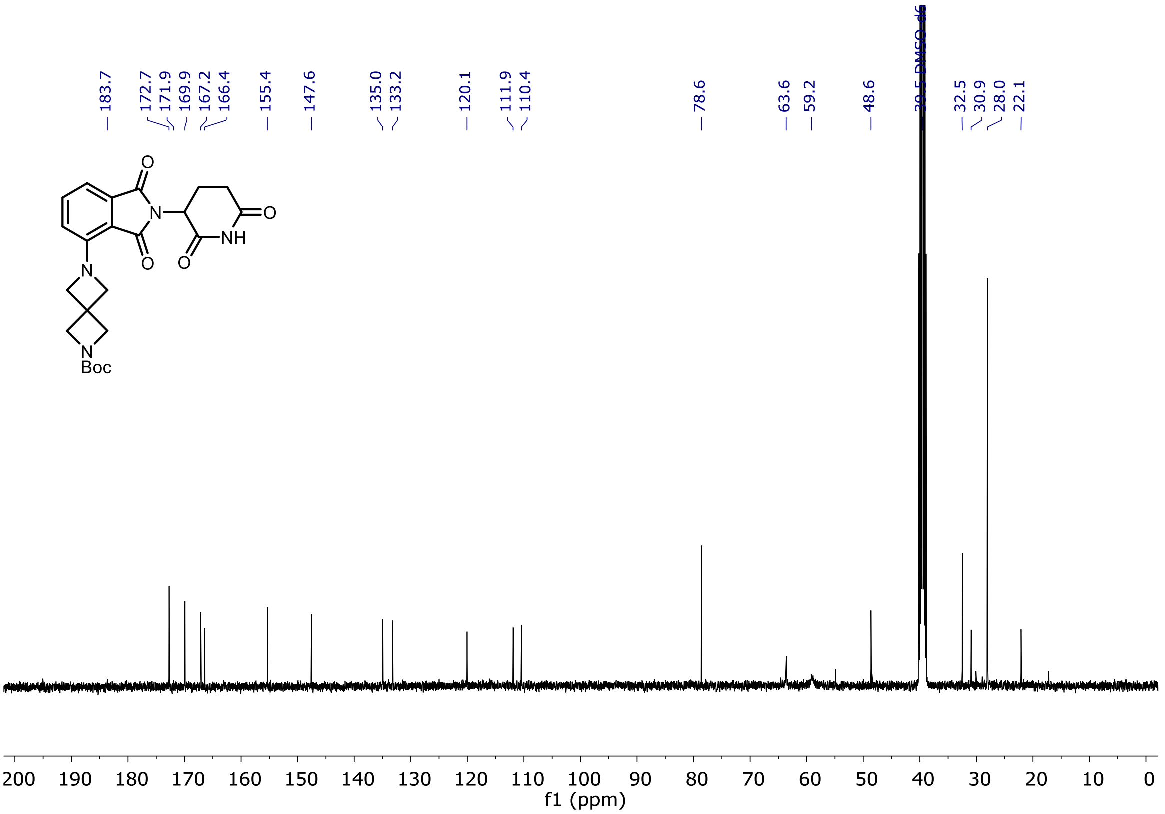

<sup></sup>

$^{1}H$ and $^{13}C$ NMR of Entry **24**

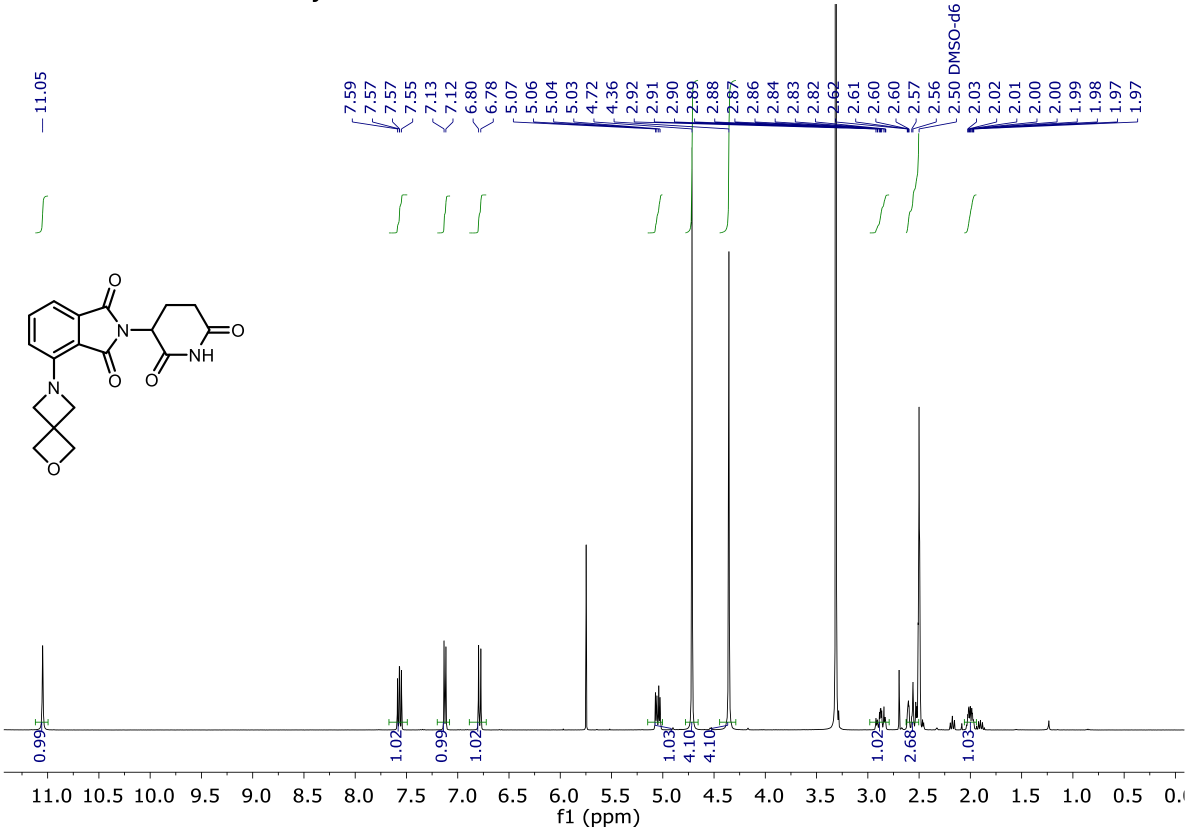

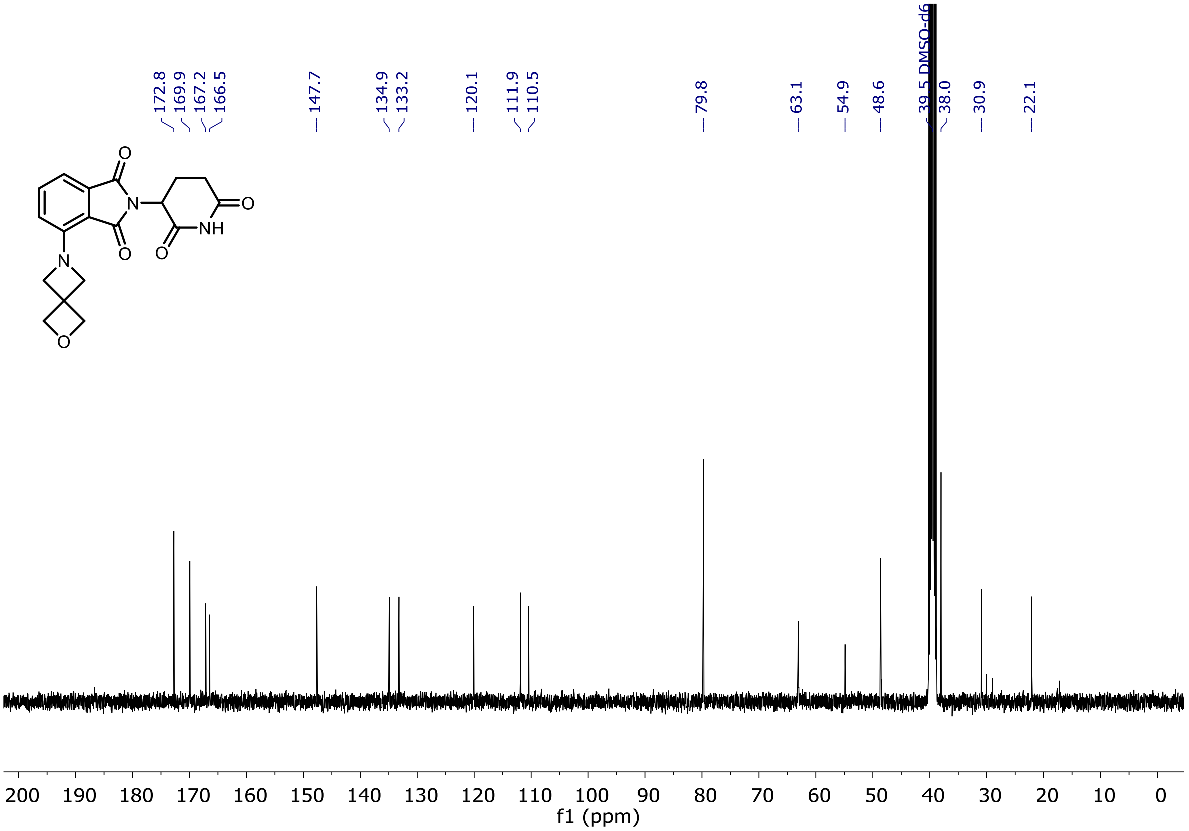

<sup></sup>

[1]H and [13]C NMR of Entry **25**

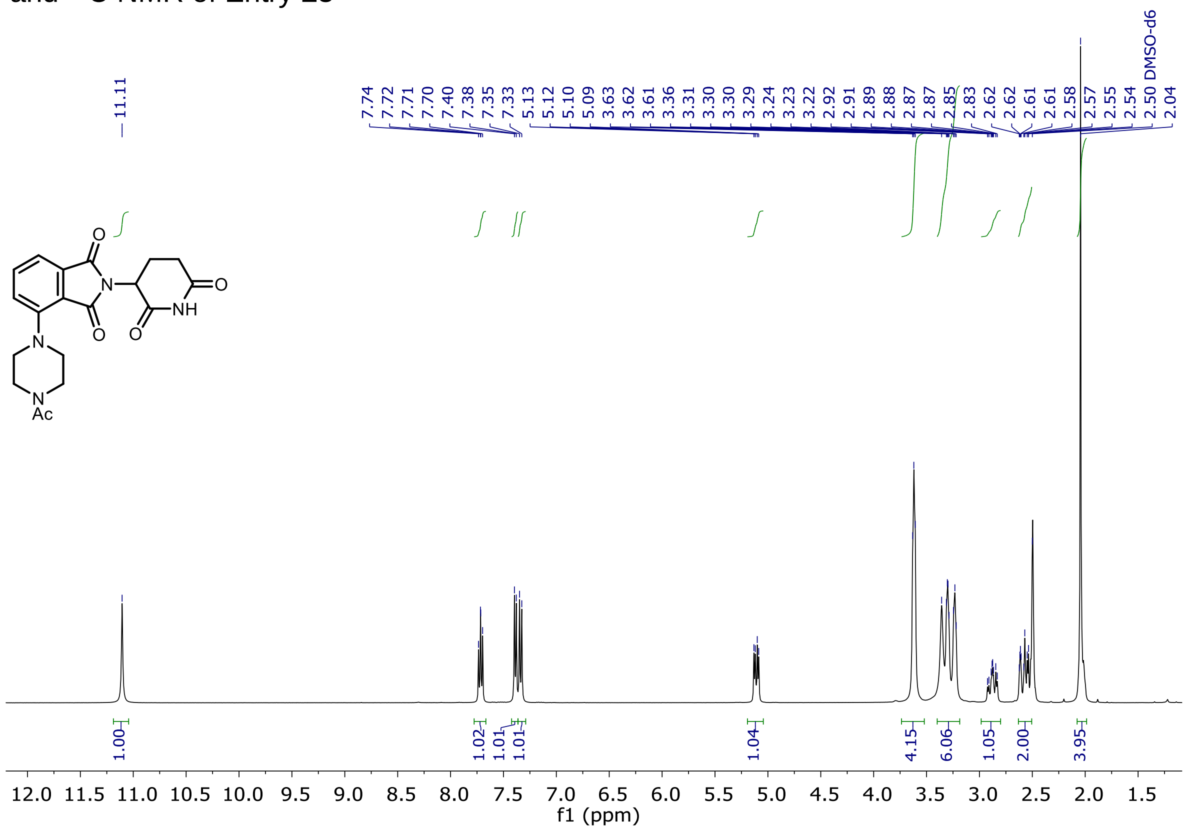

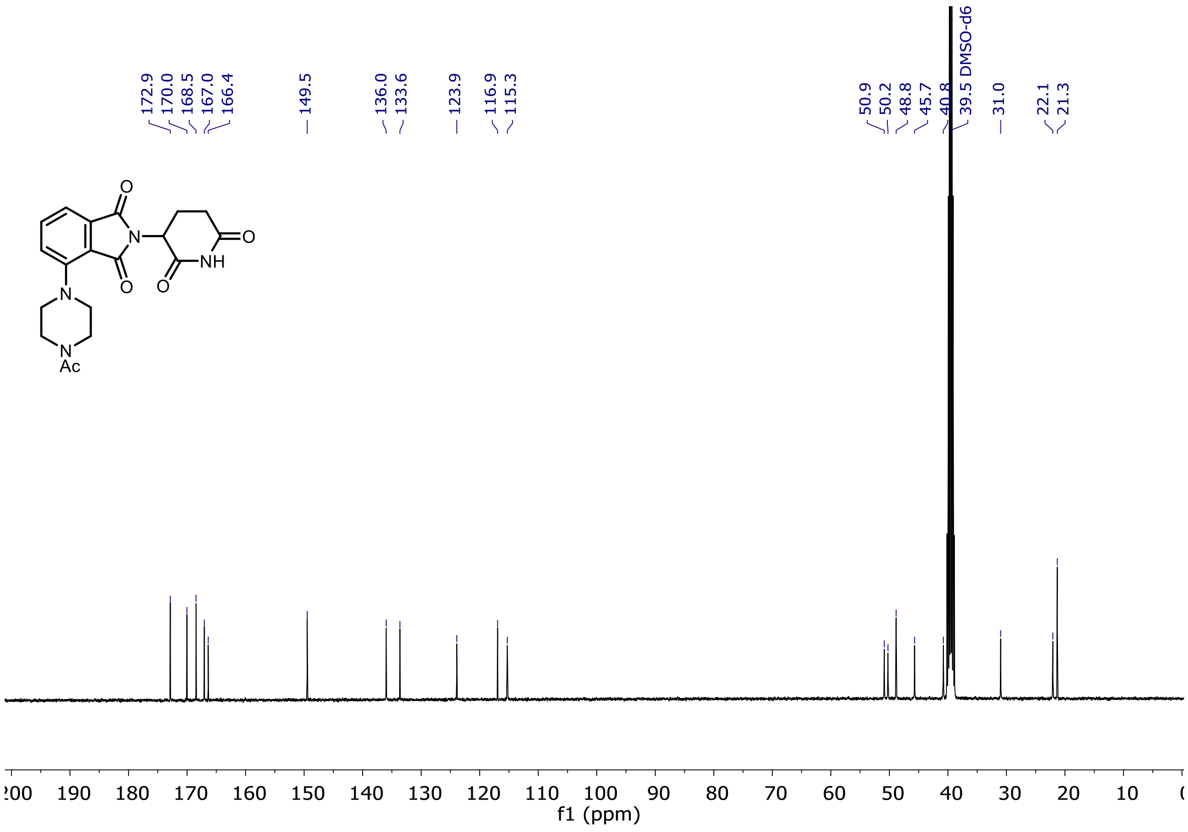

$^1$H and $^{13}$C NMR of Entry **26**

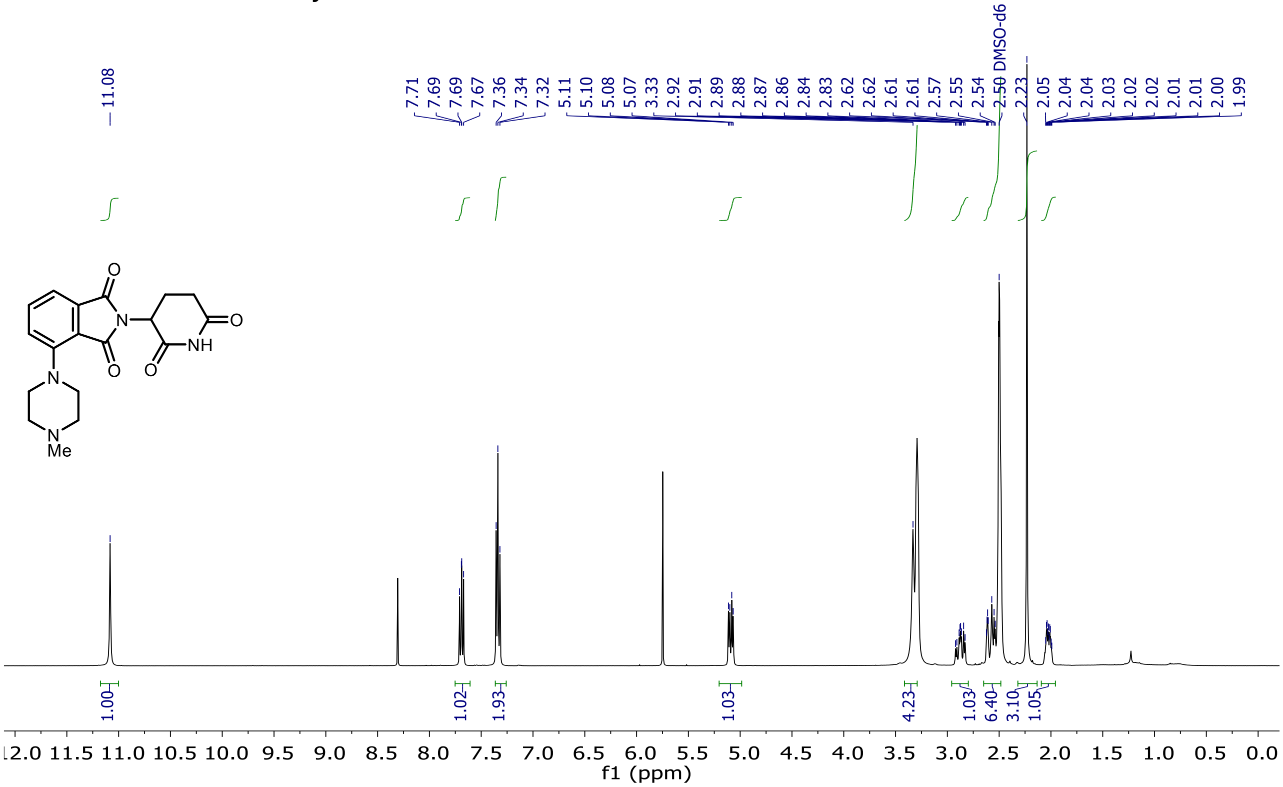

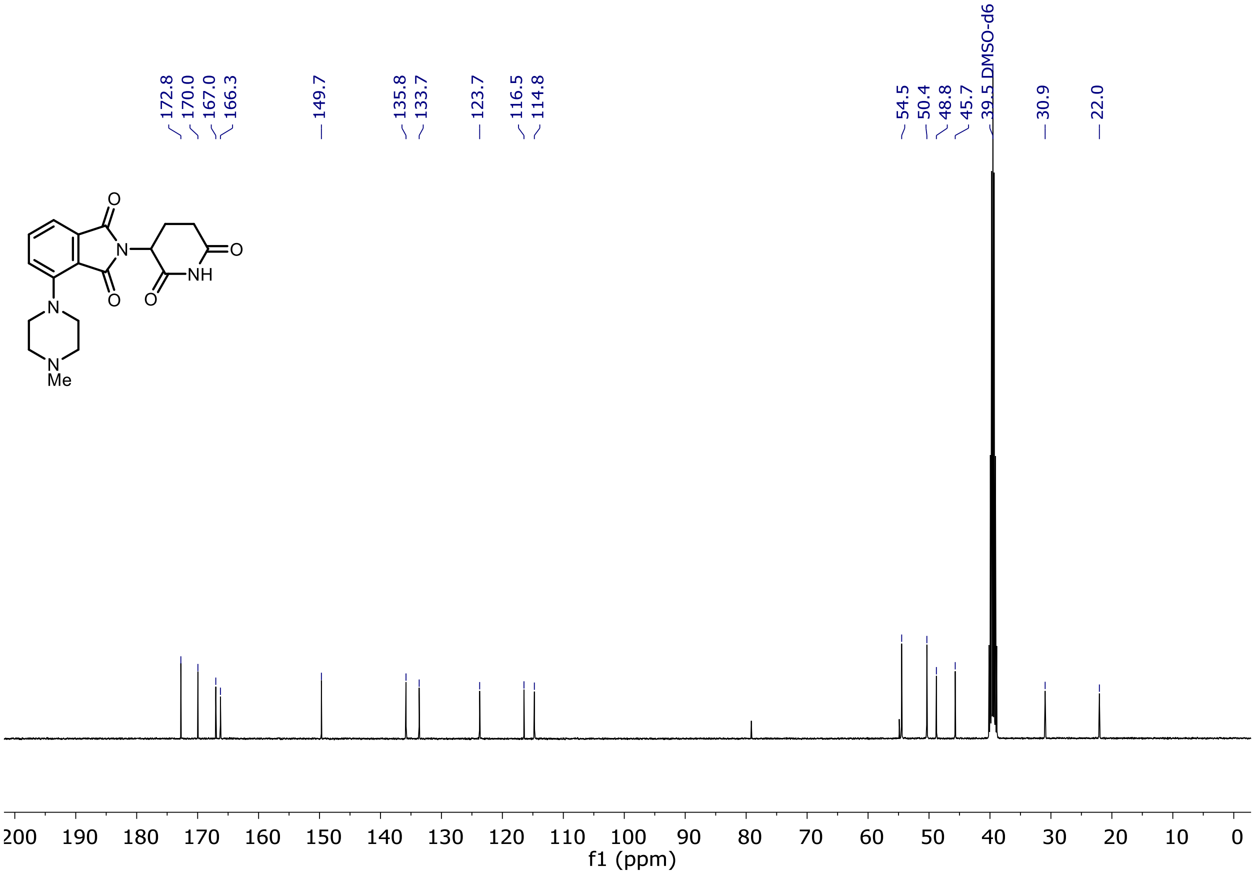

$^{1}H$ and $^{13}C$ NMR of Entry **27**

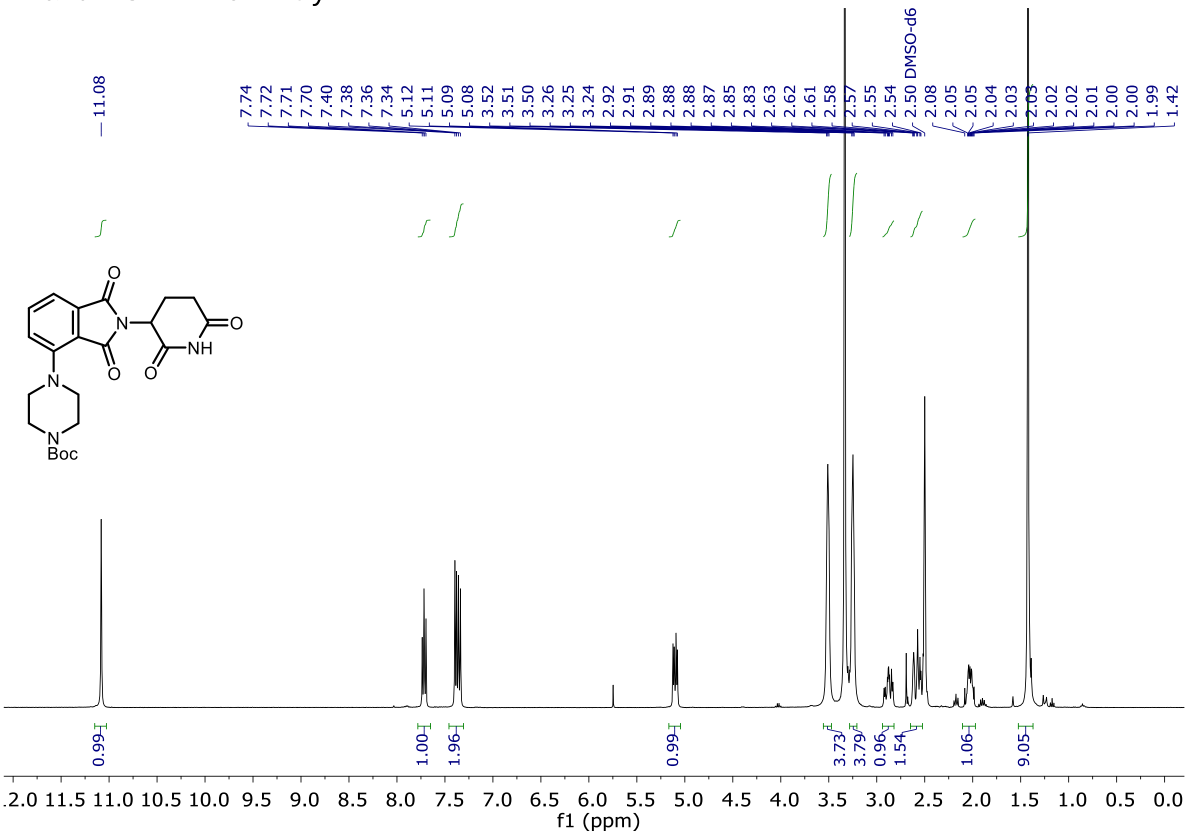

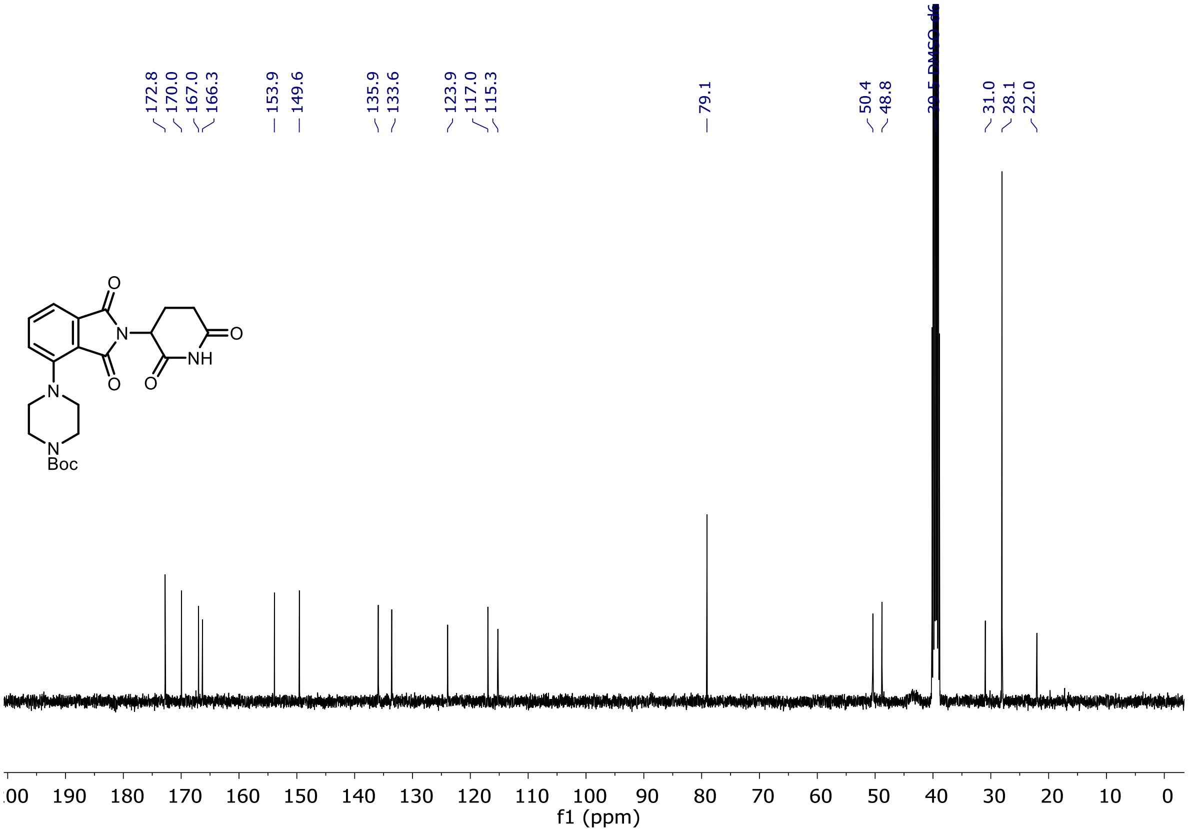

[1]H and [13]C NMR of Entry 28

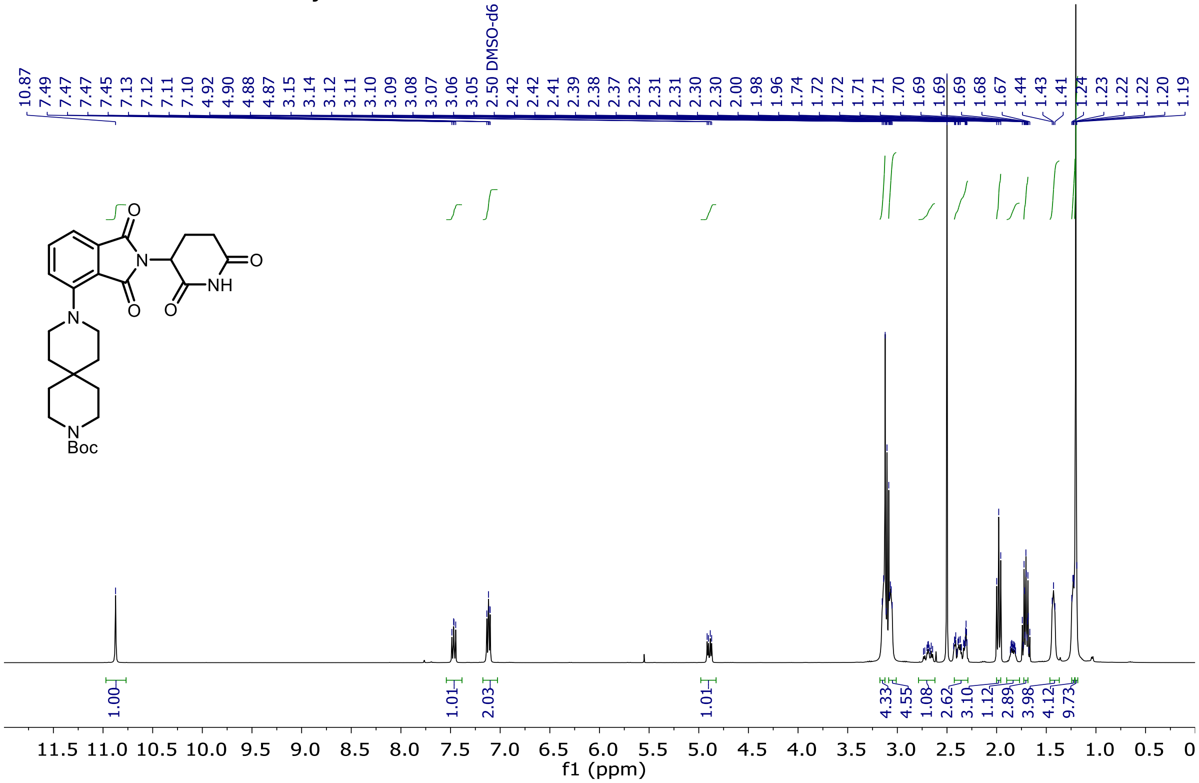

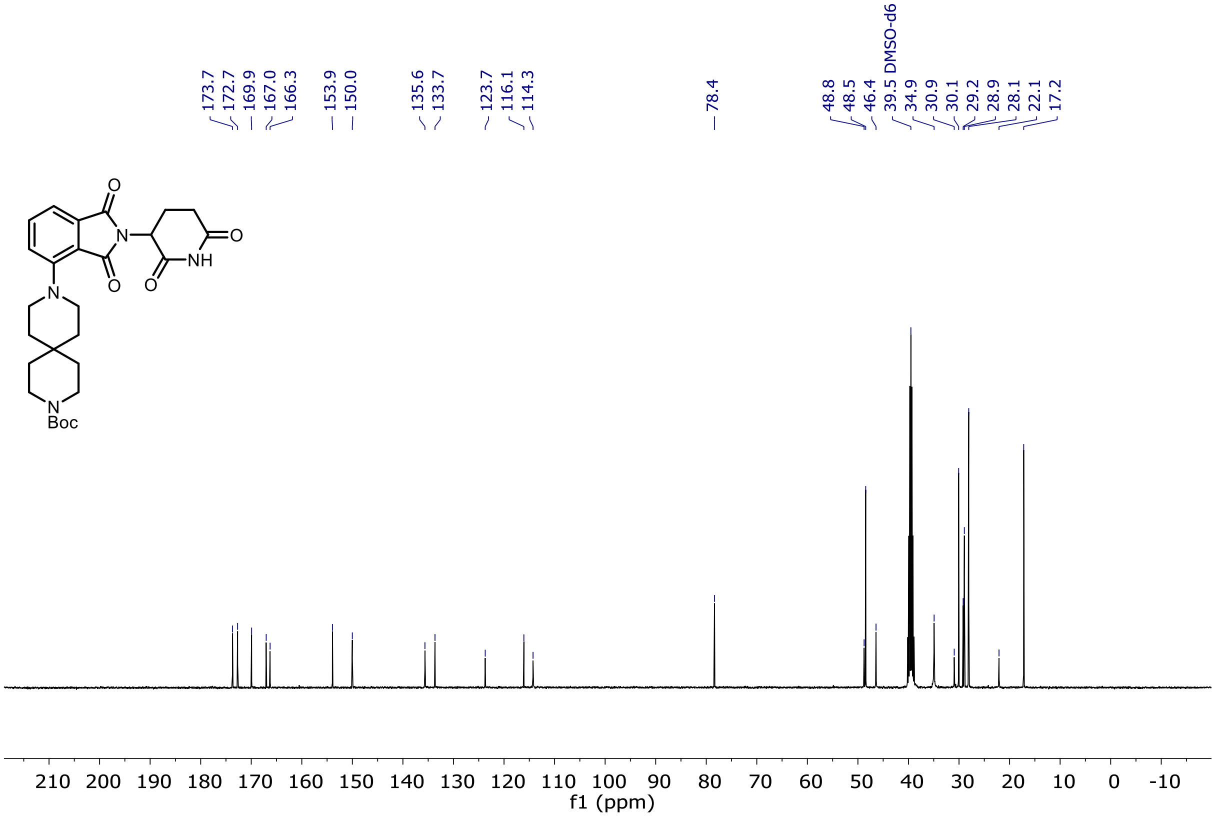

[1]H and [13]C NMR of Entry **30**

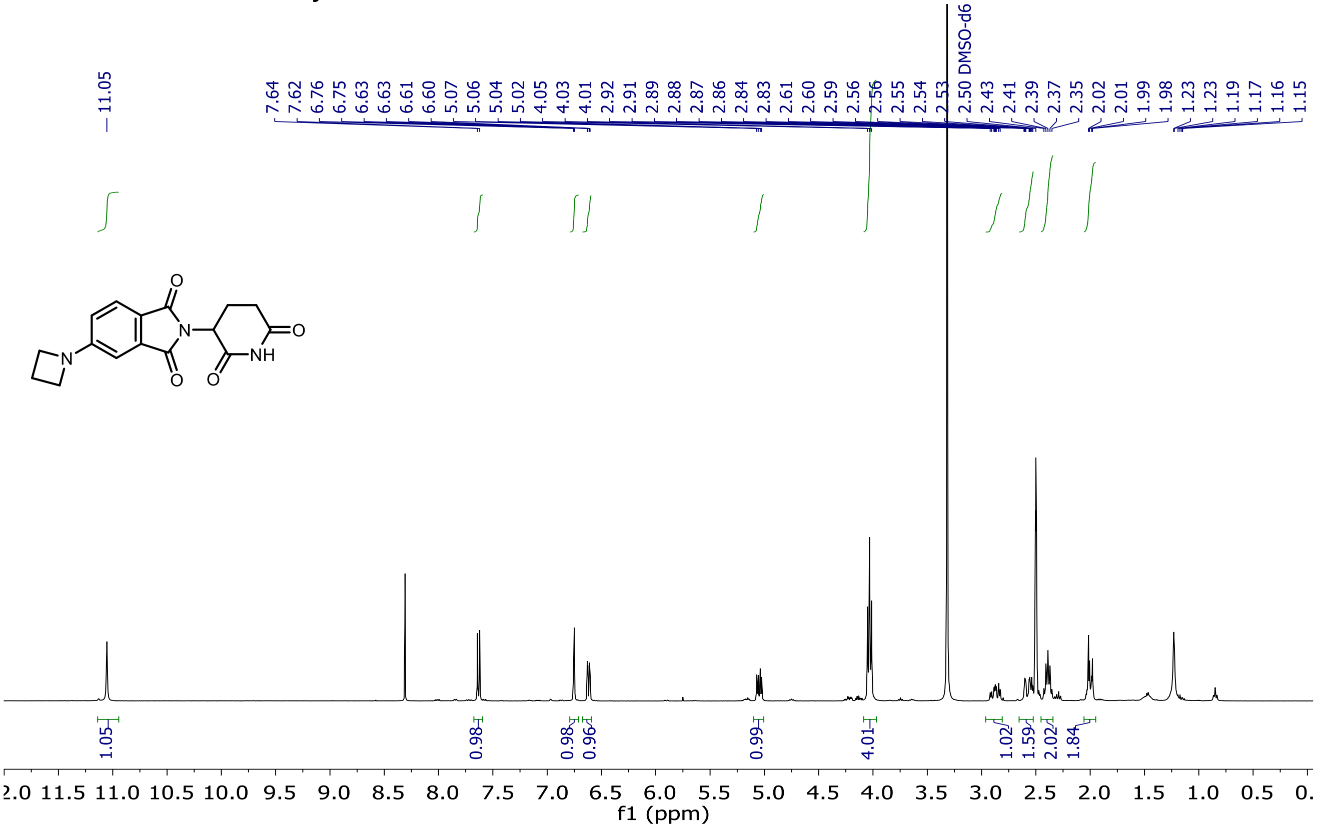

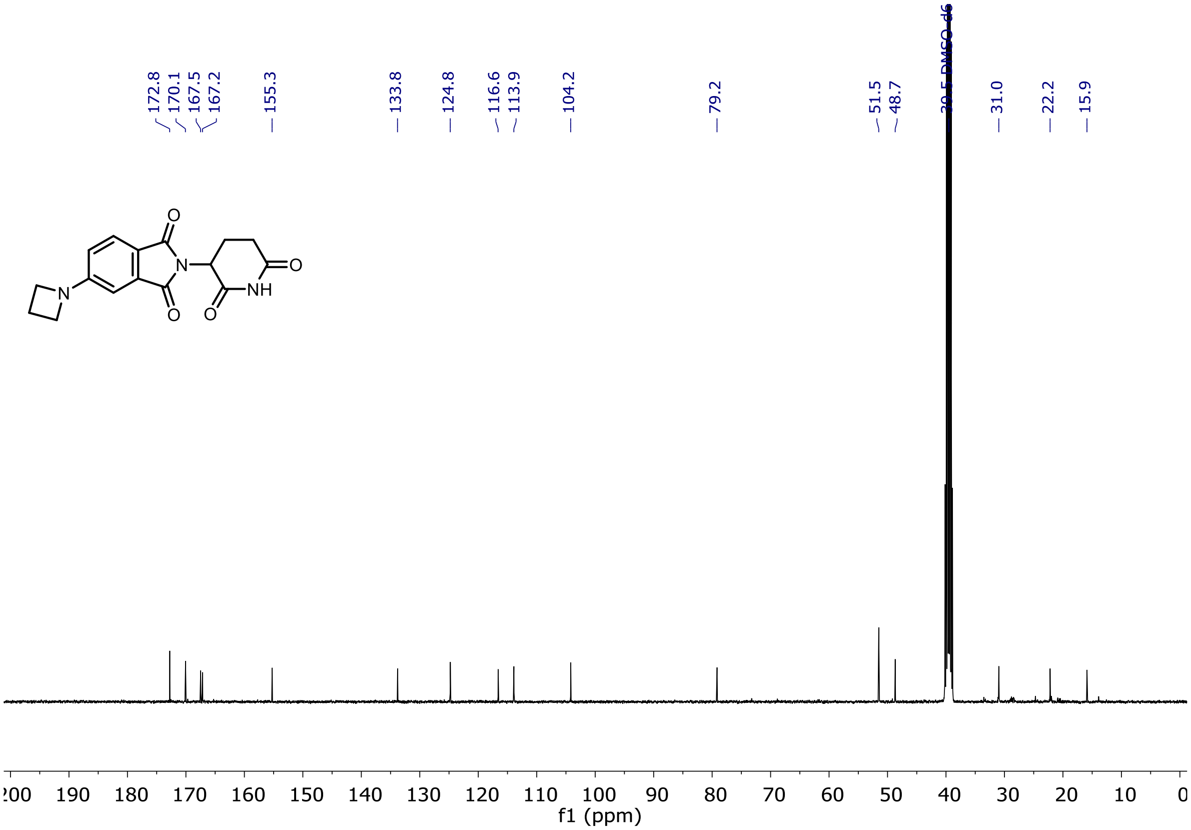

$^{1}H$ and $^{13}C$ NMR of Entry **31**

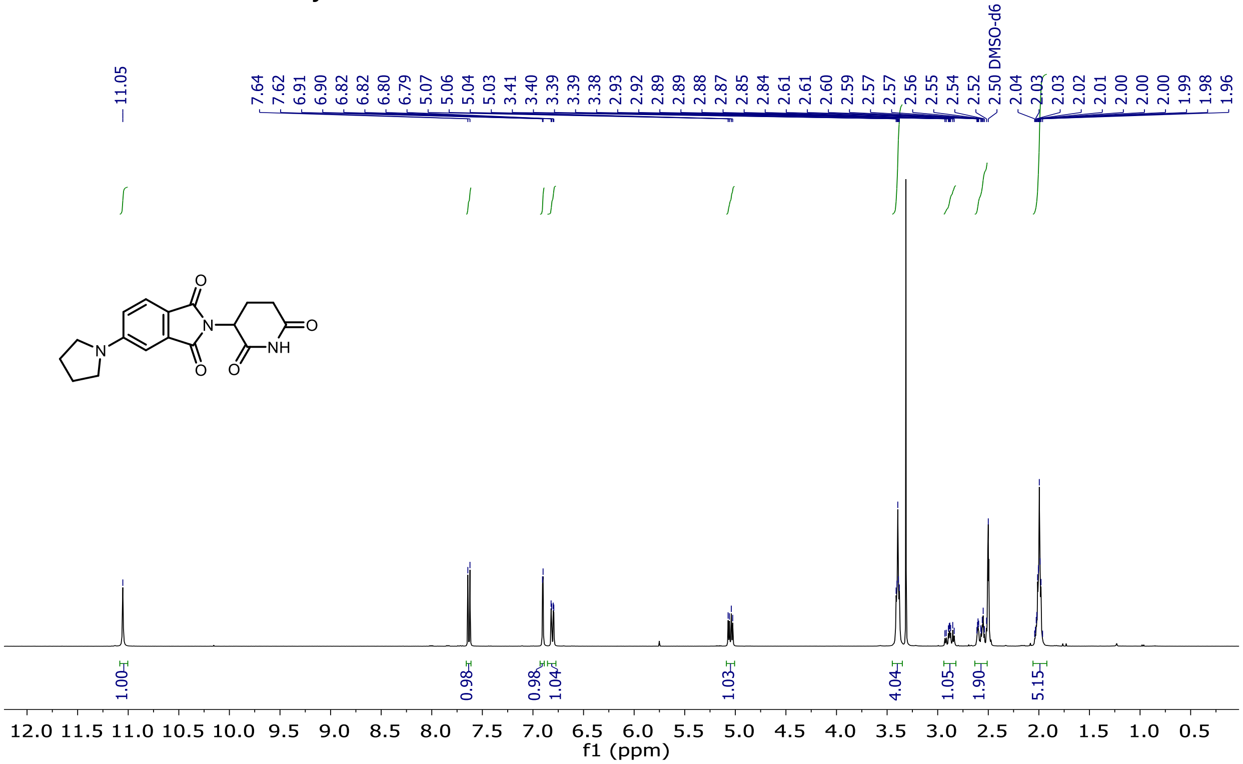

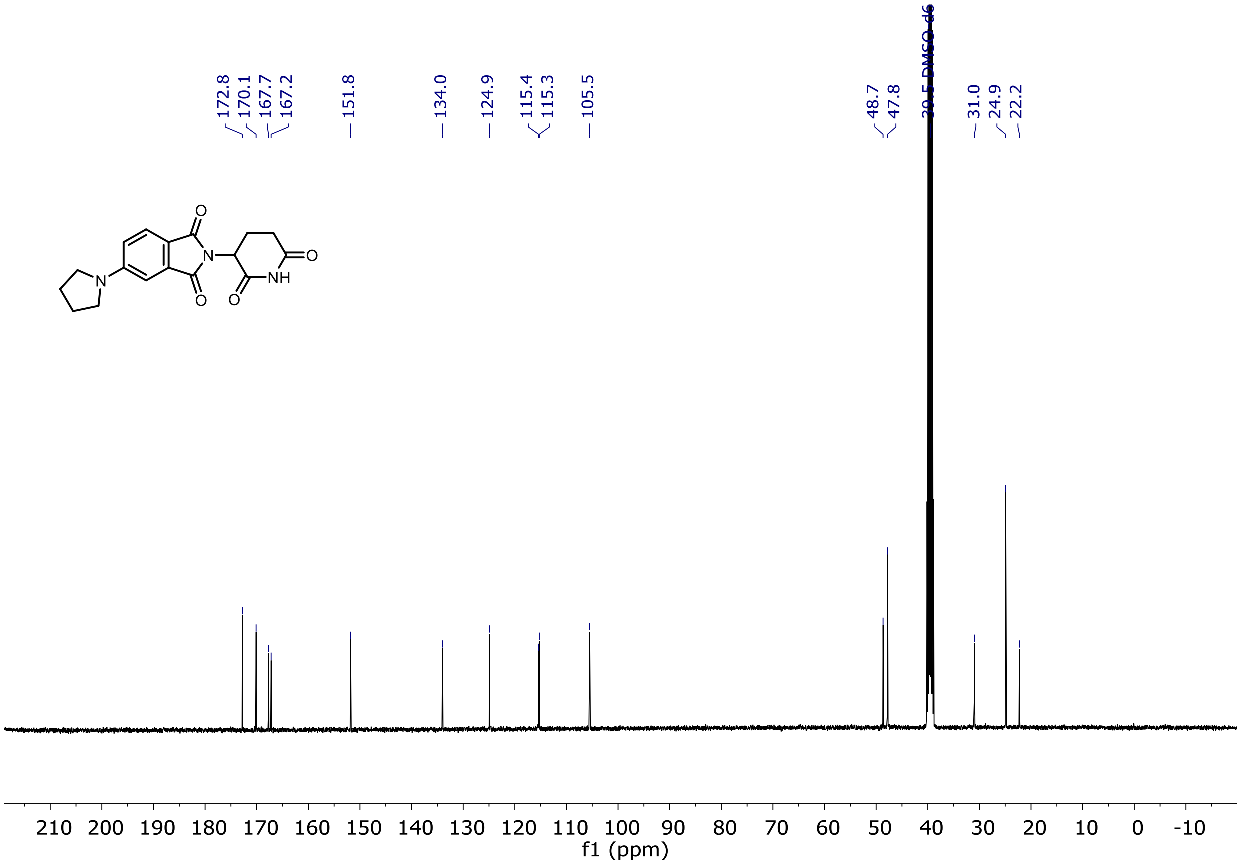

$^1H$ and $^{13}C$ NMR of Entry **32**

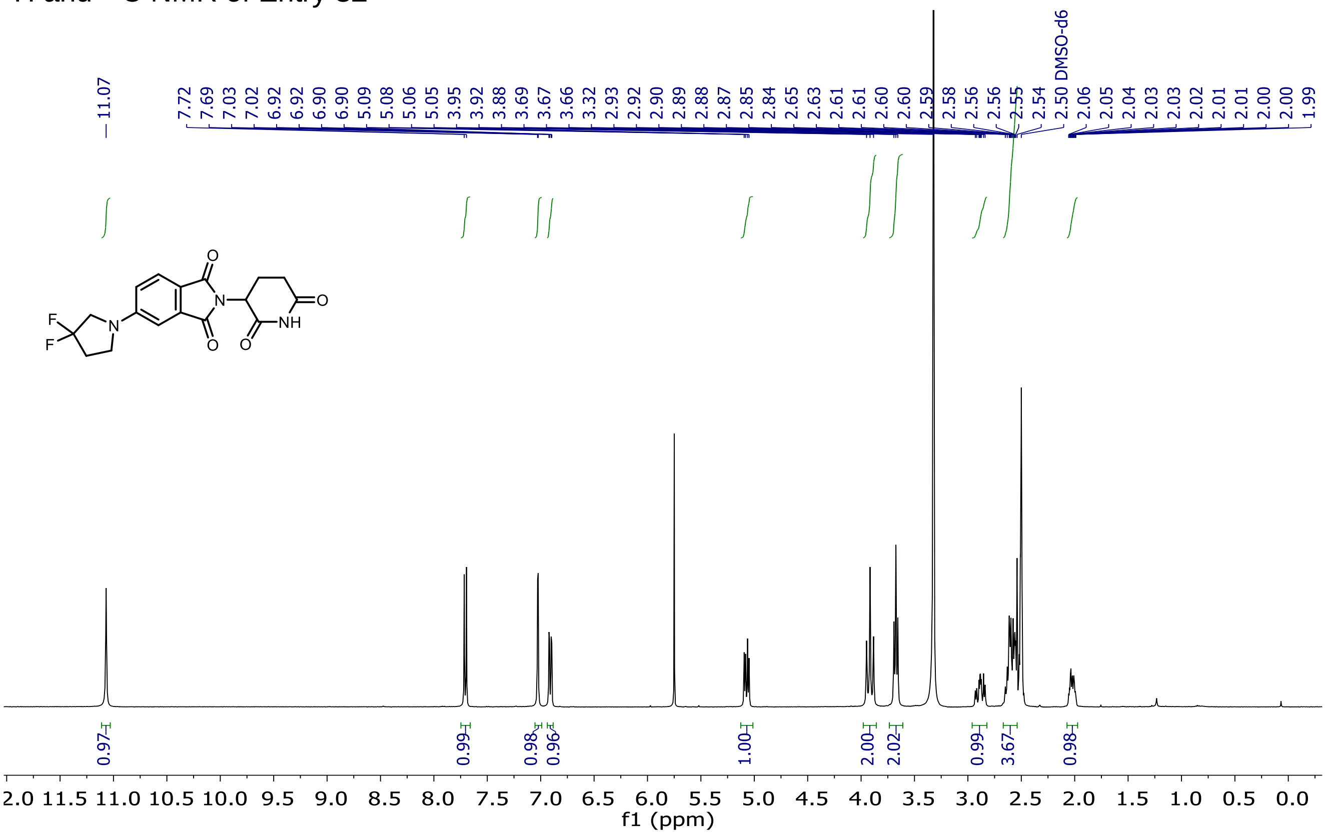

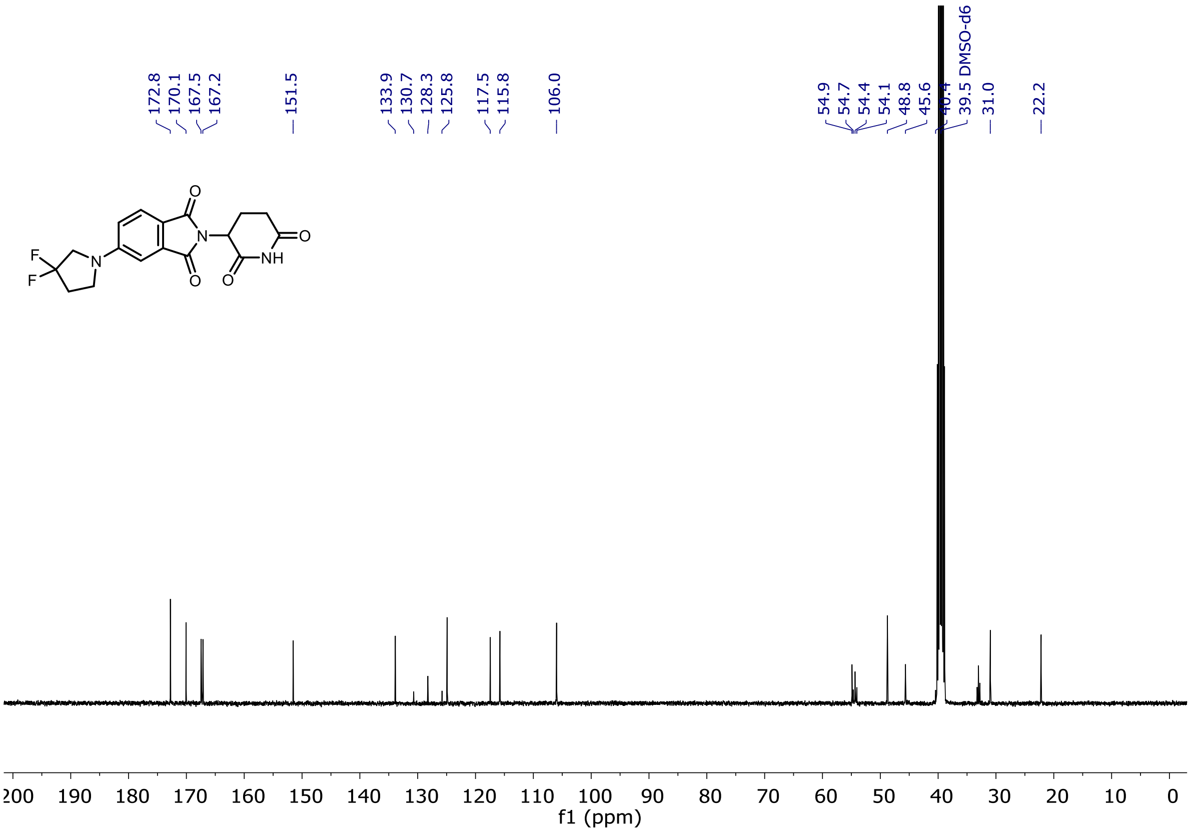

$^{19}$F NMR of Entry **32**

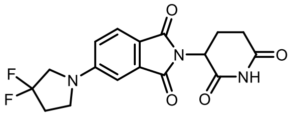

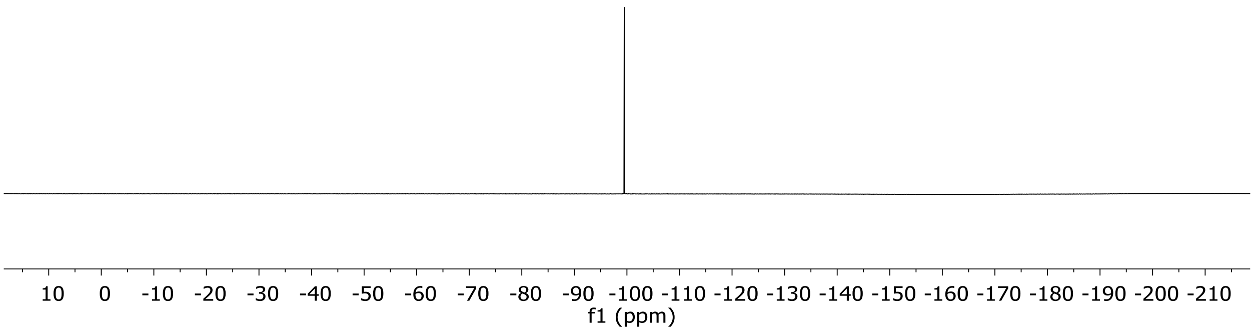

[1]H and [13]C NMR of Entry **33**

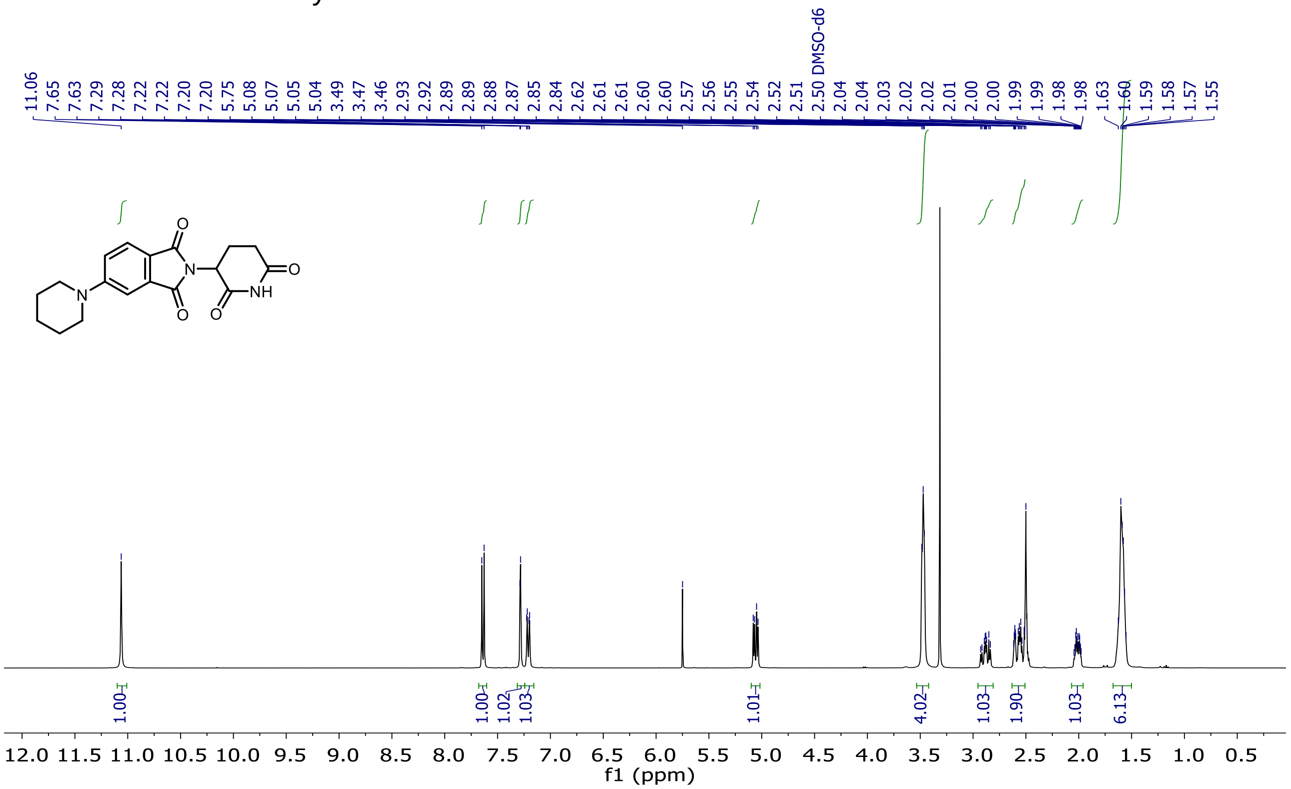

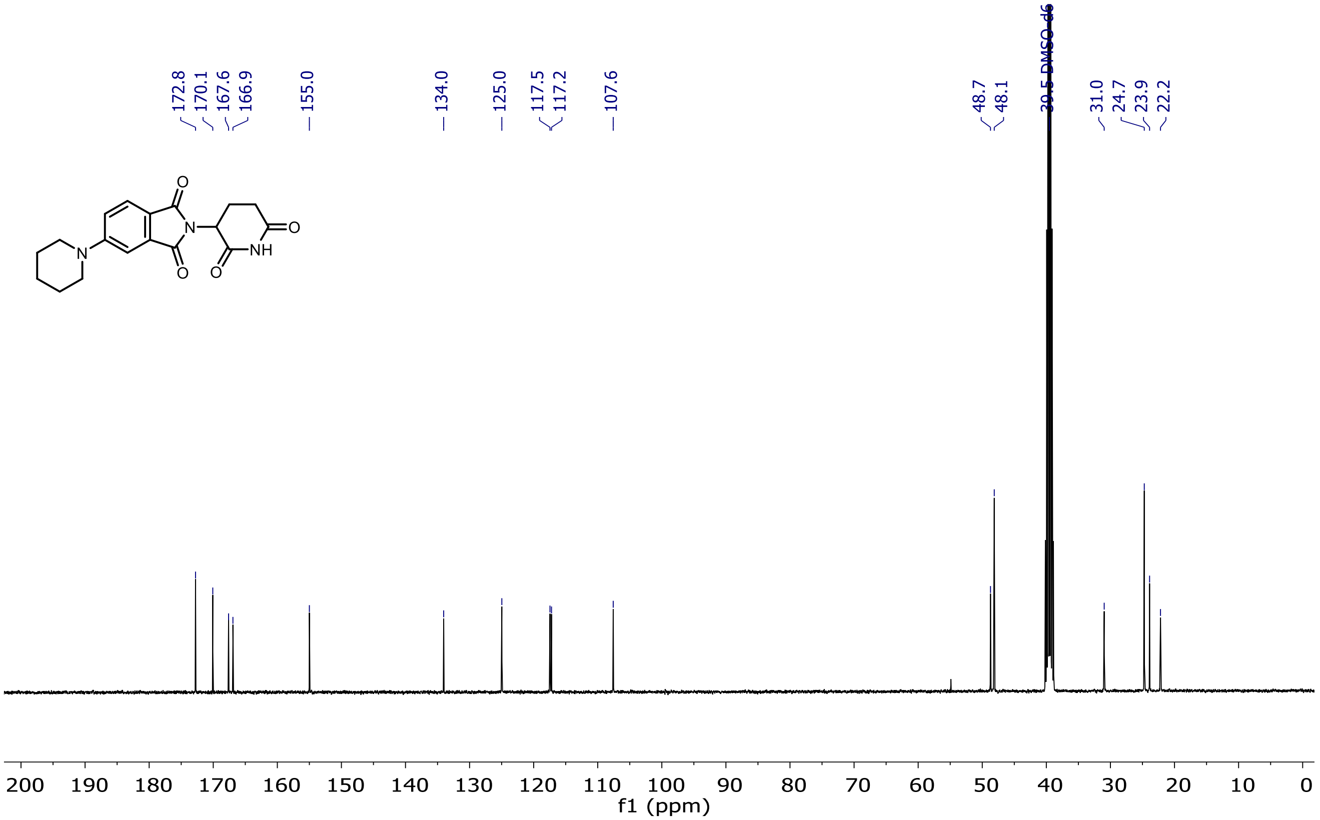

<sup></sup>

$^{1}$H and $^{13}$C NMR of Entry **34**

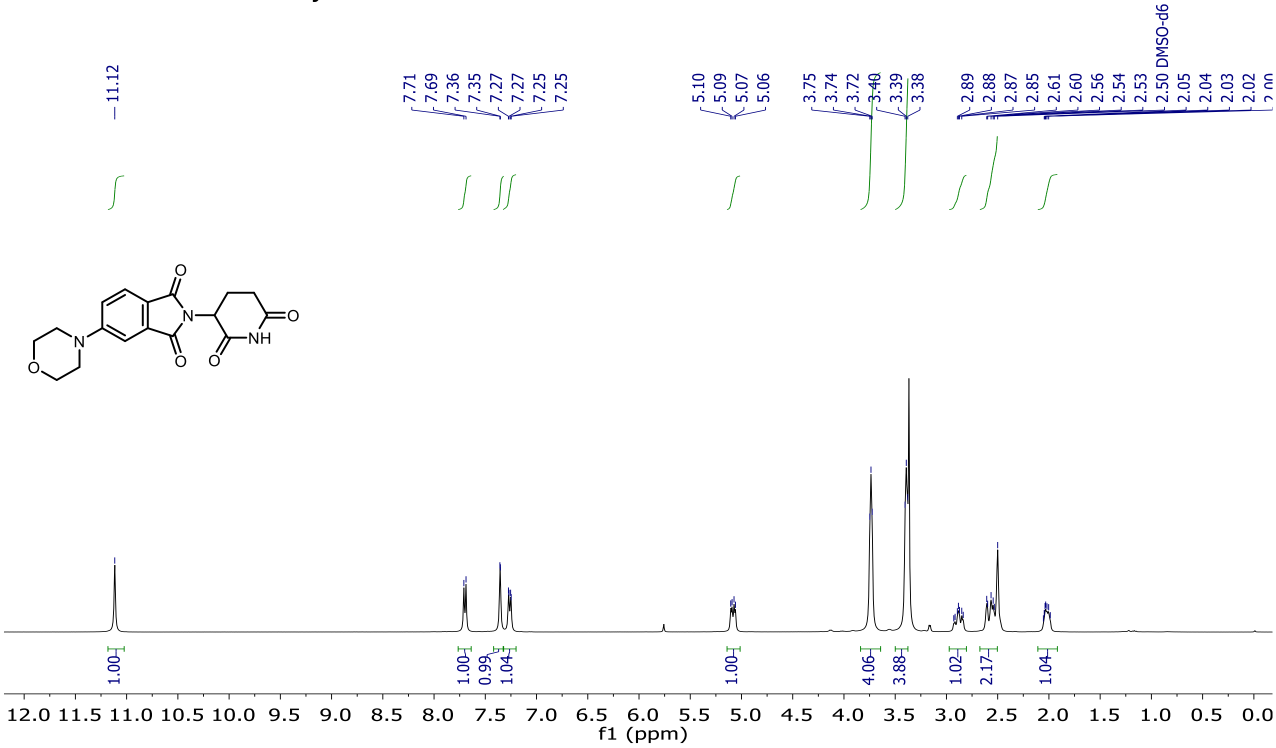

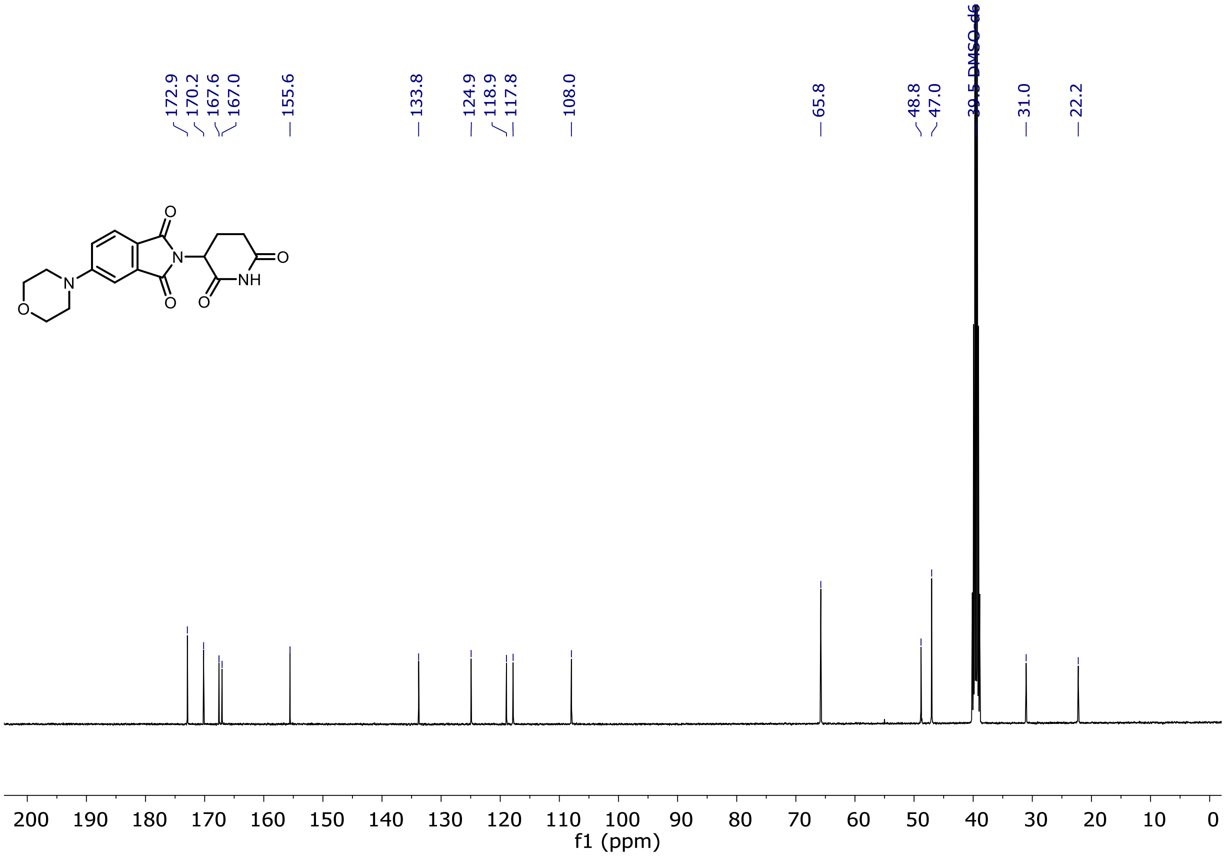

[1]H and [13]C NMR of Entry **35**

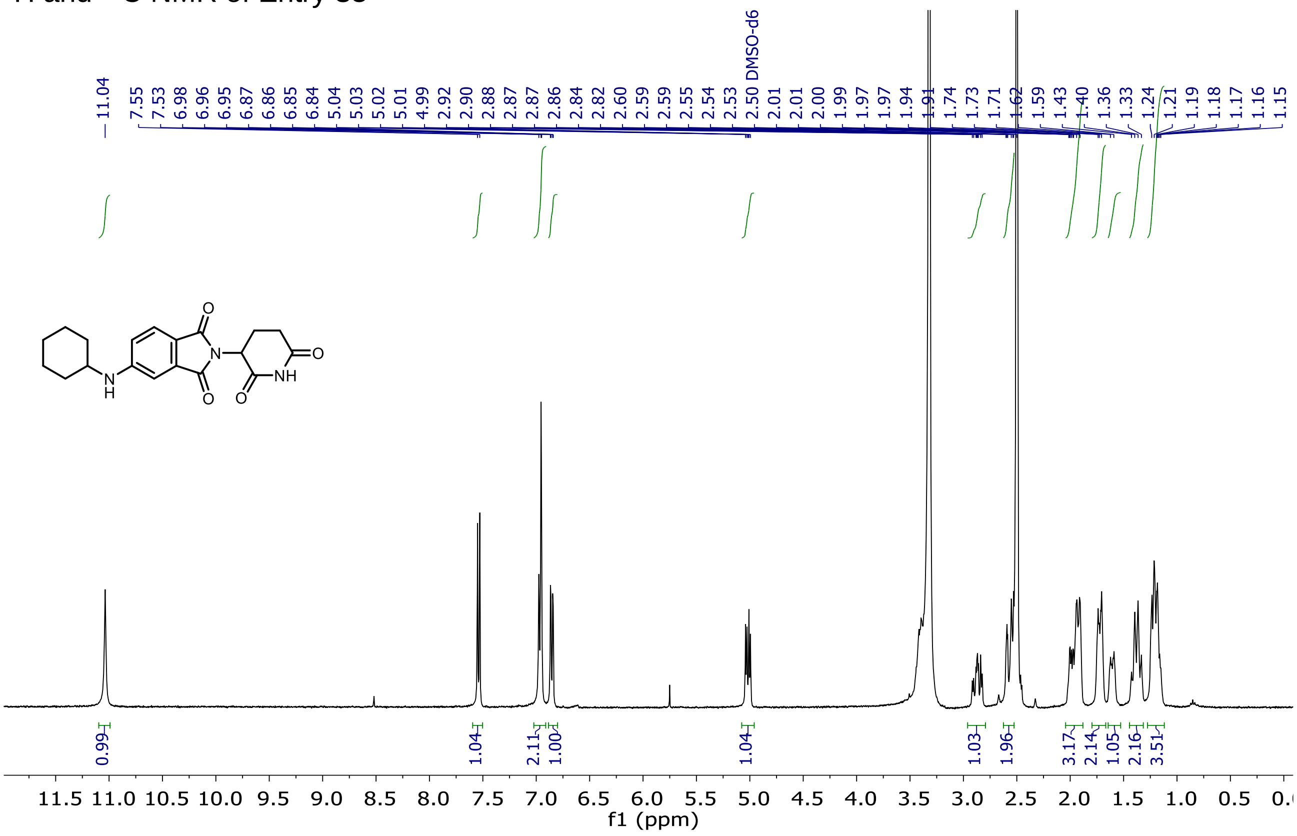

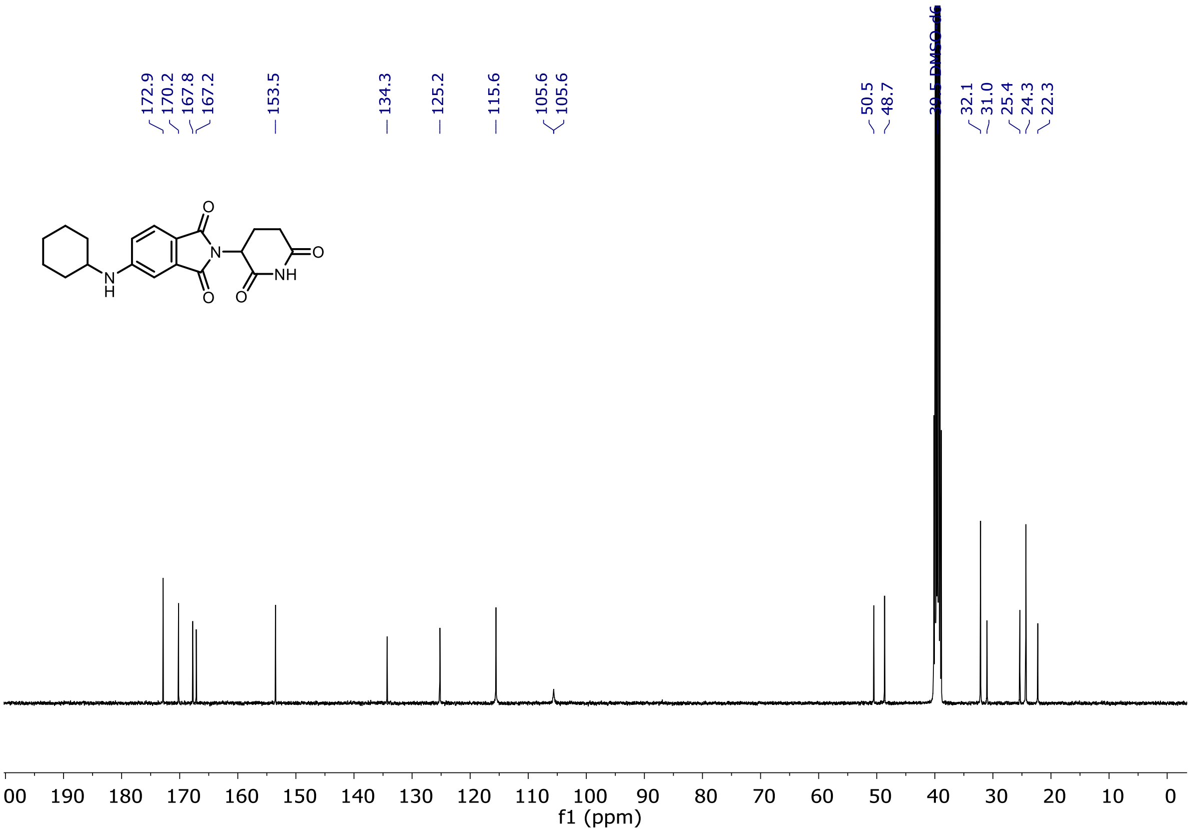

[1]H and [13]C NMR of Entry **36**

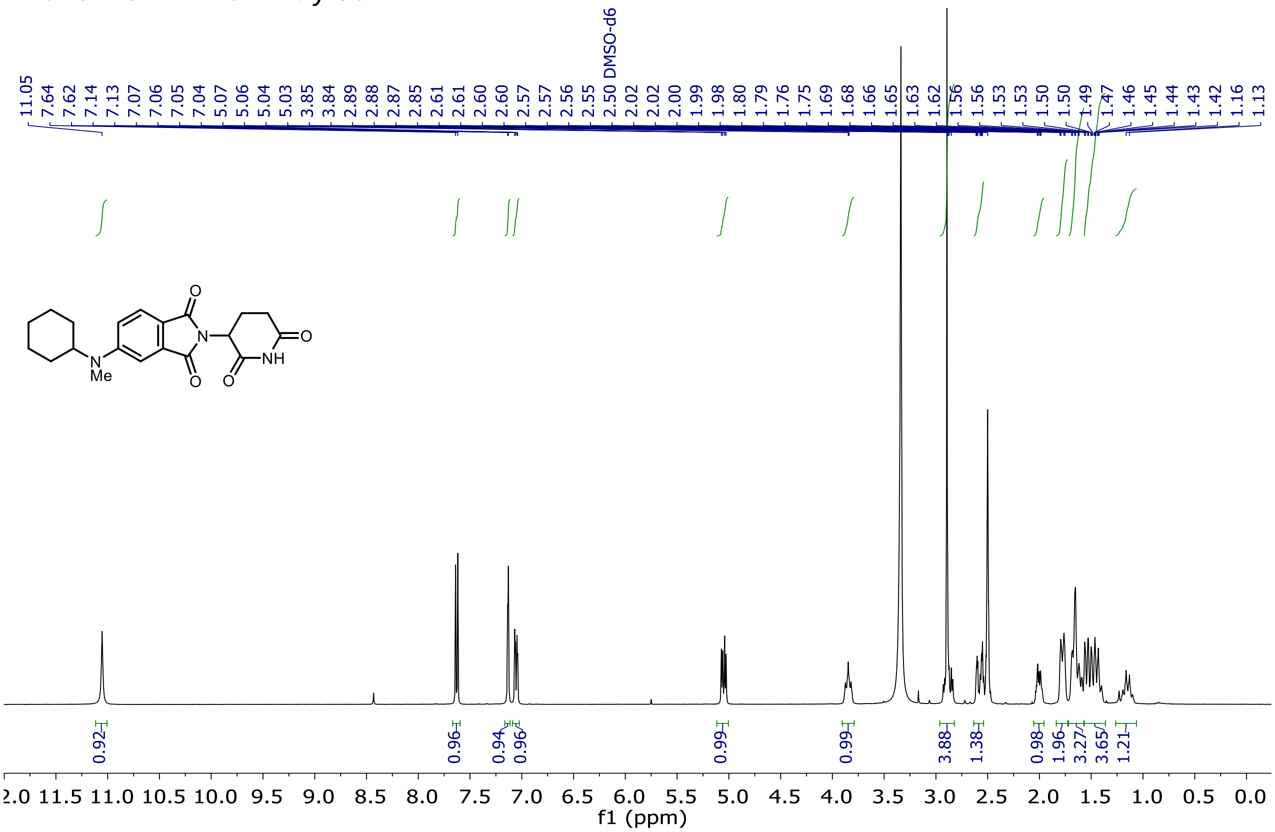

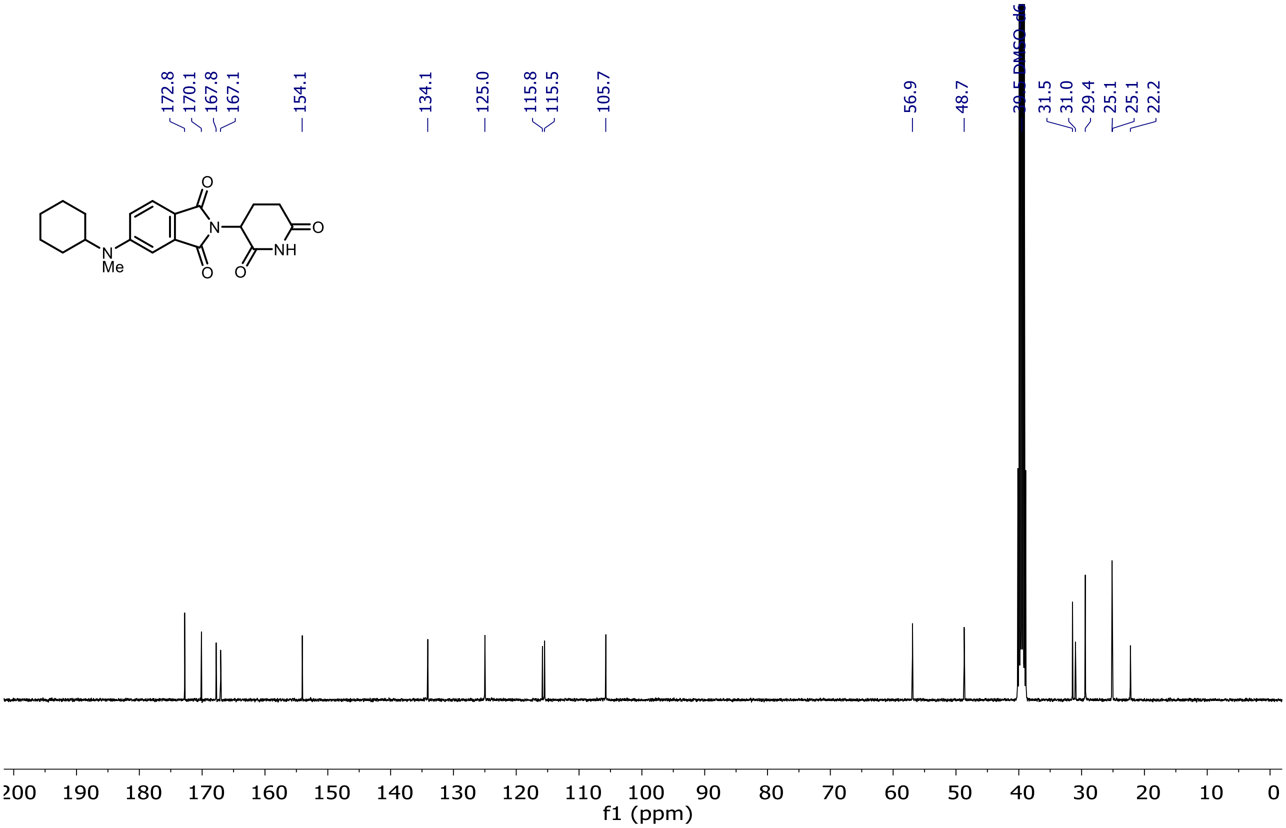

$^{1}H$ and $^{13}C$ NMR of Entry **37**

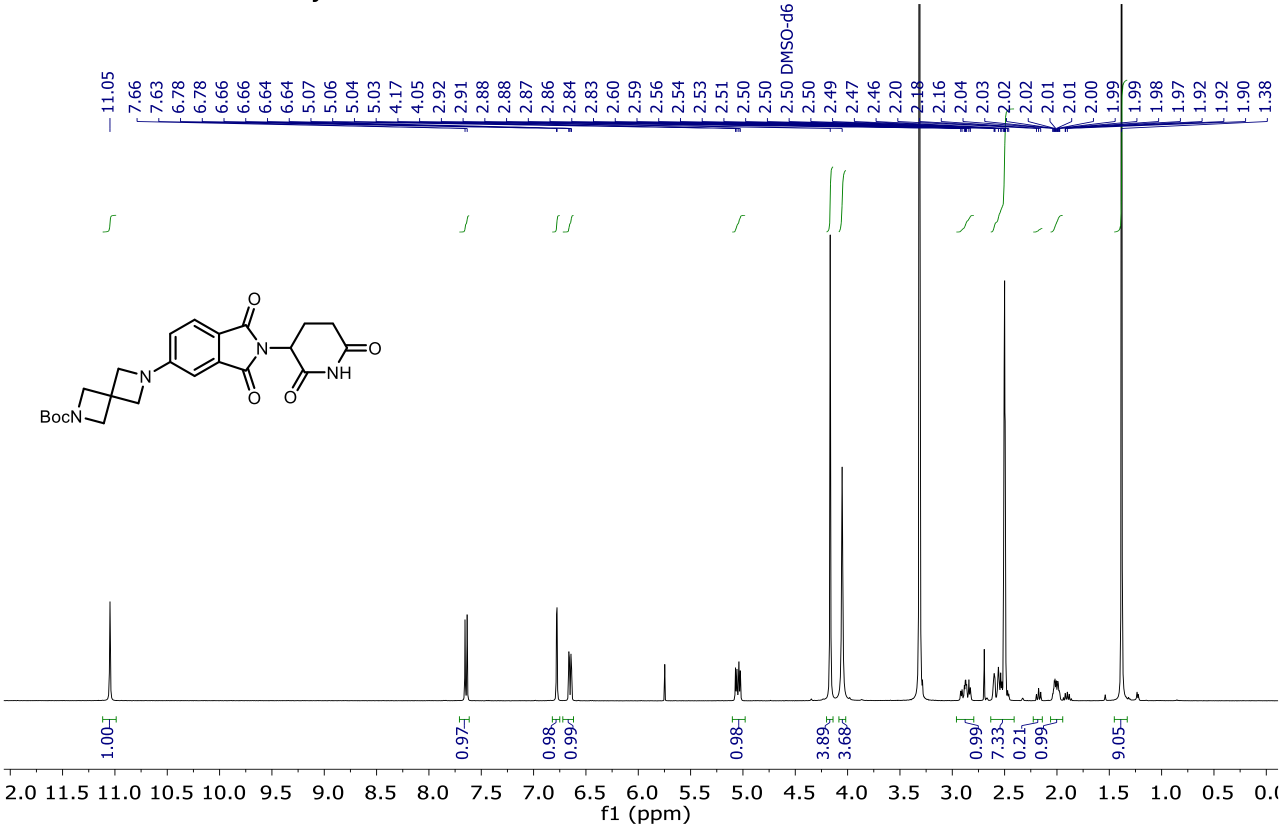

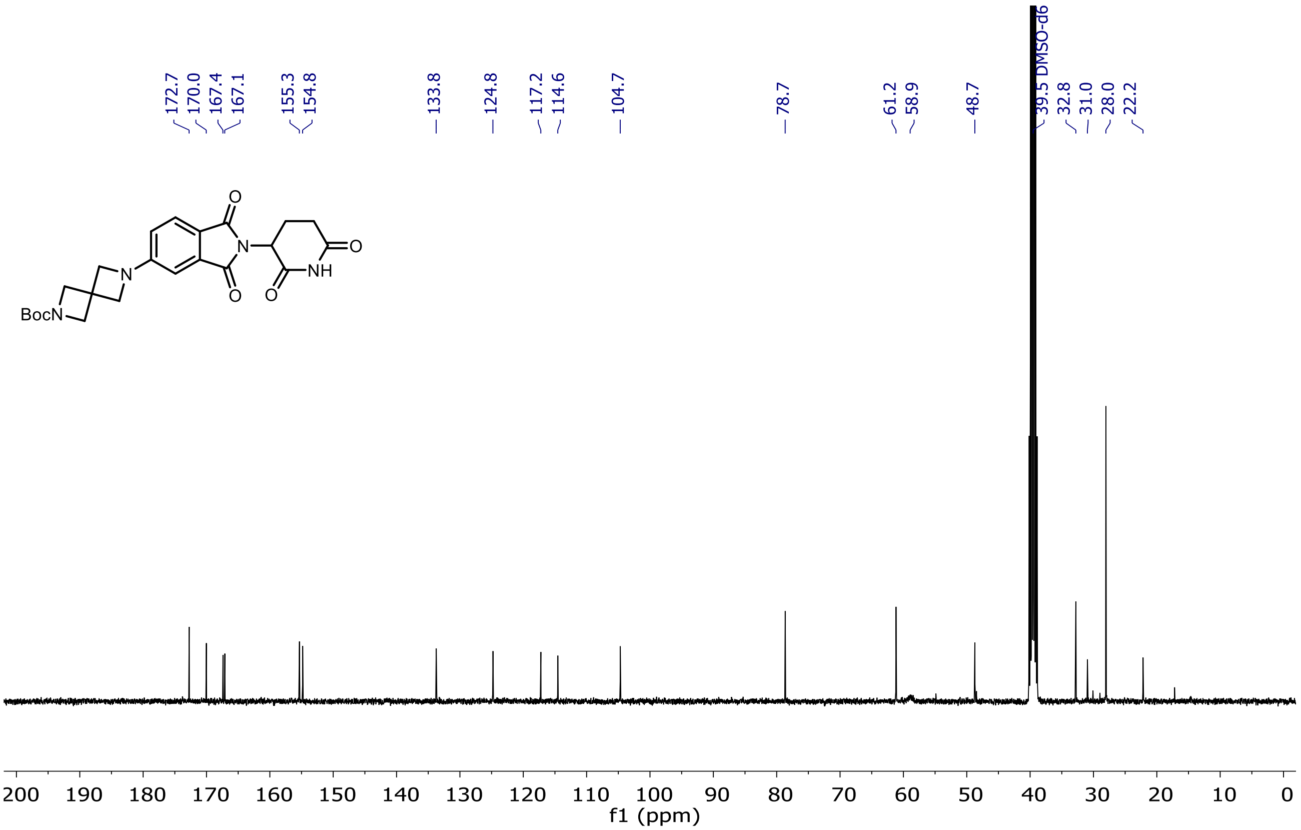

$^{1}H$ and $^{13}C$ NMR of Entry **38**

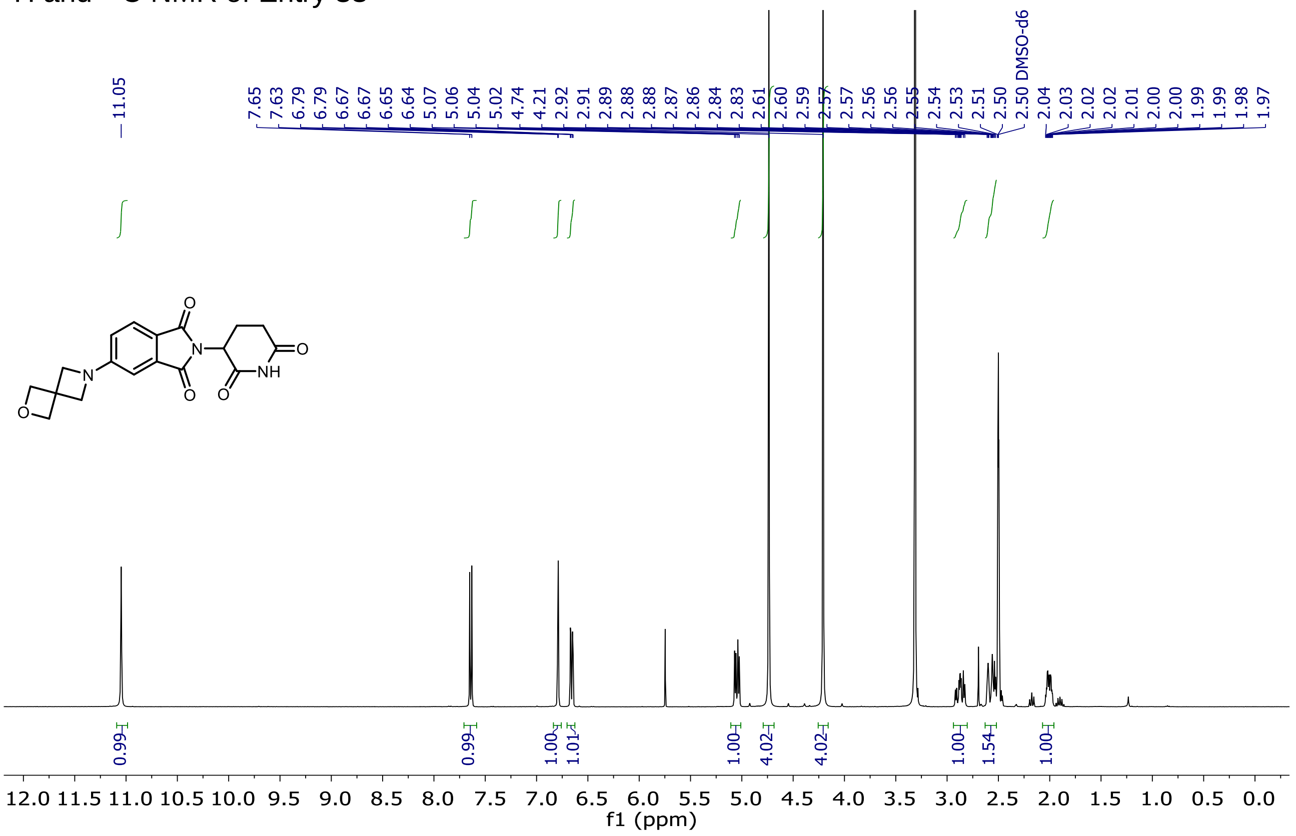

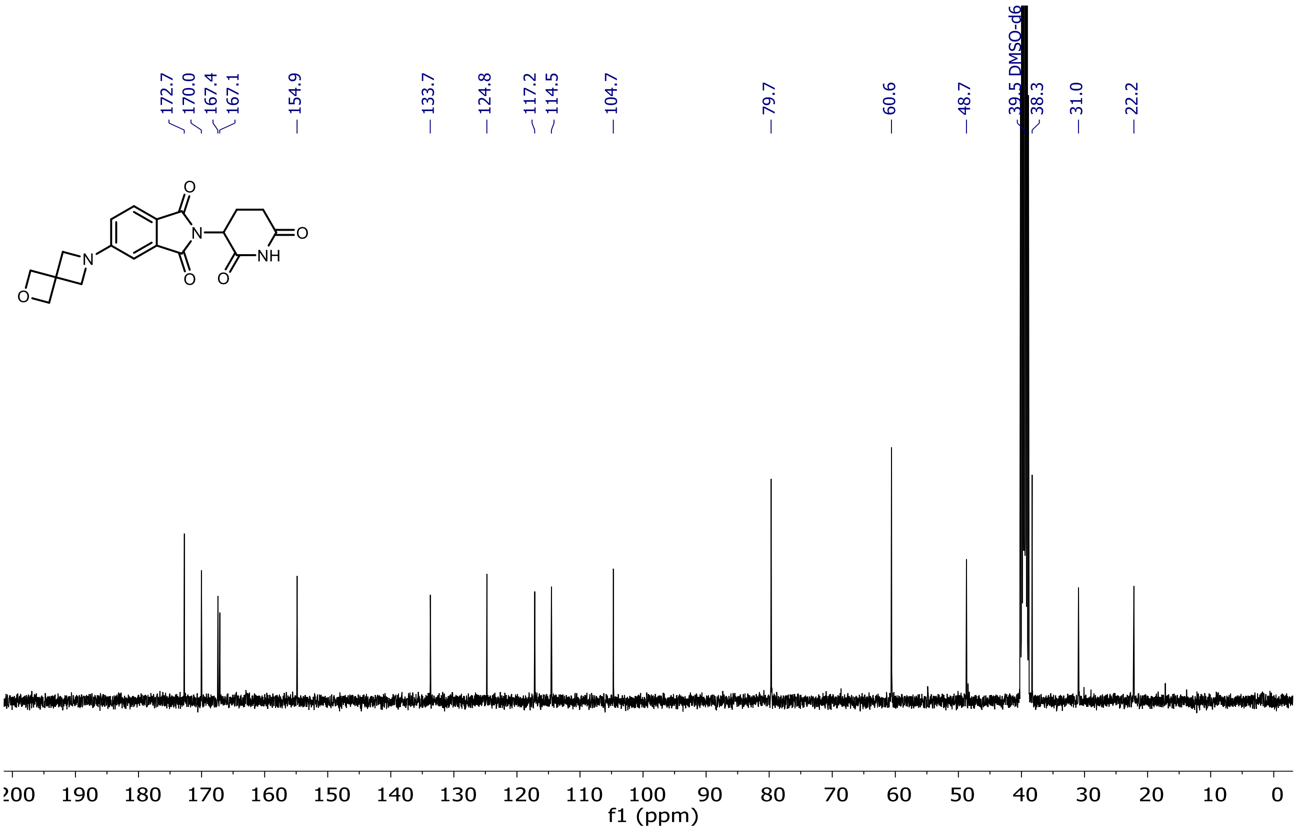

[1]H and [13]C NMR of Entry **39**

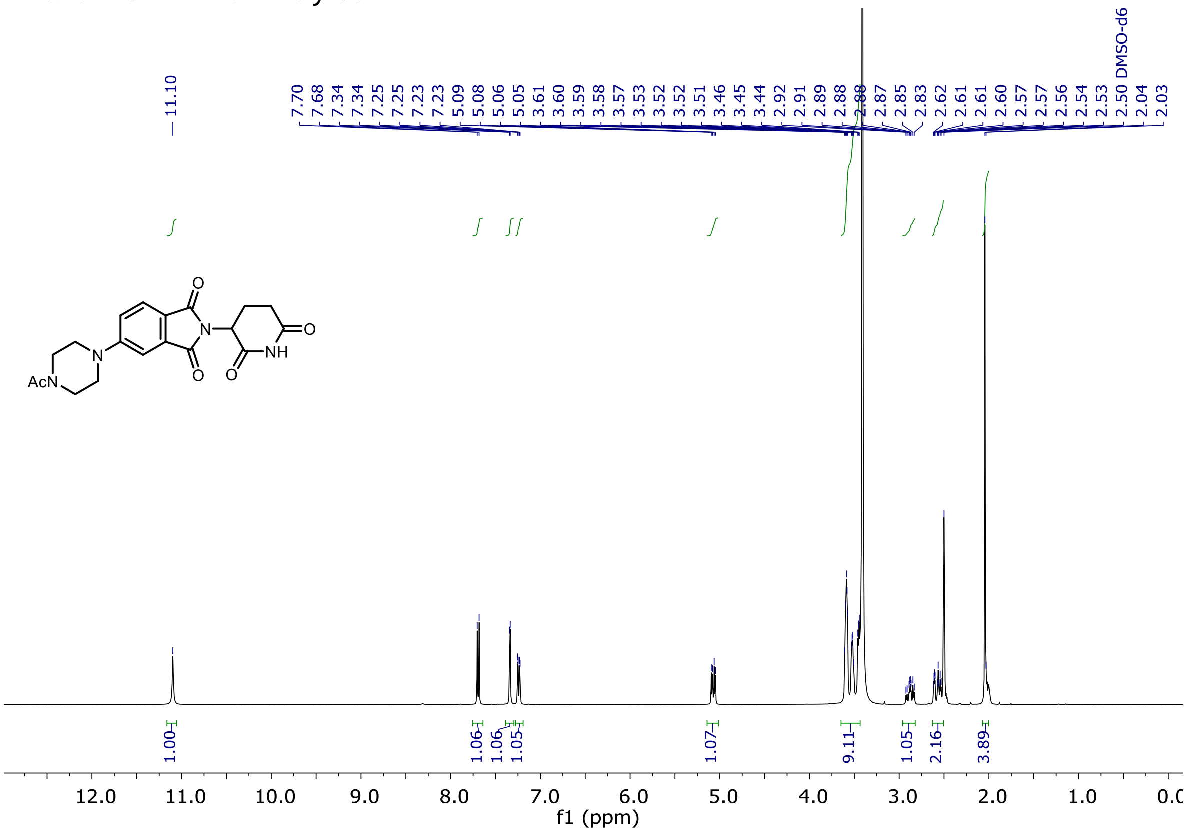

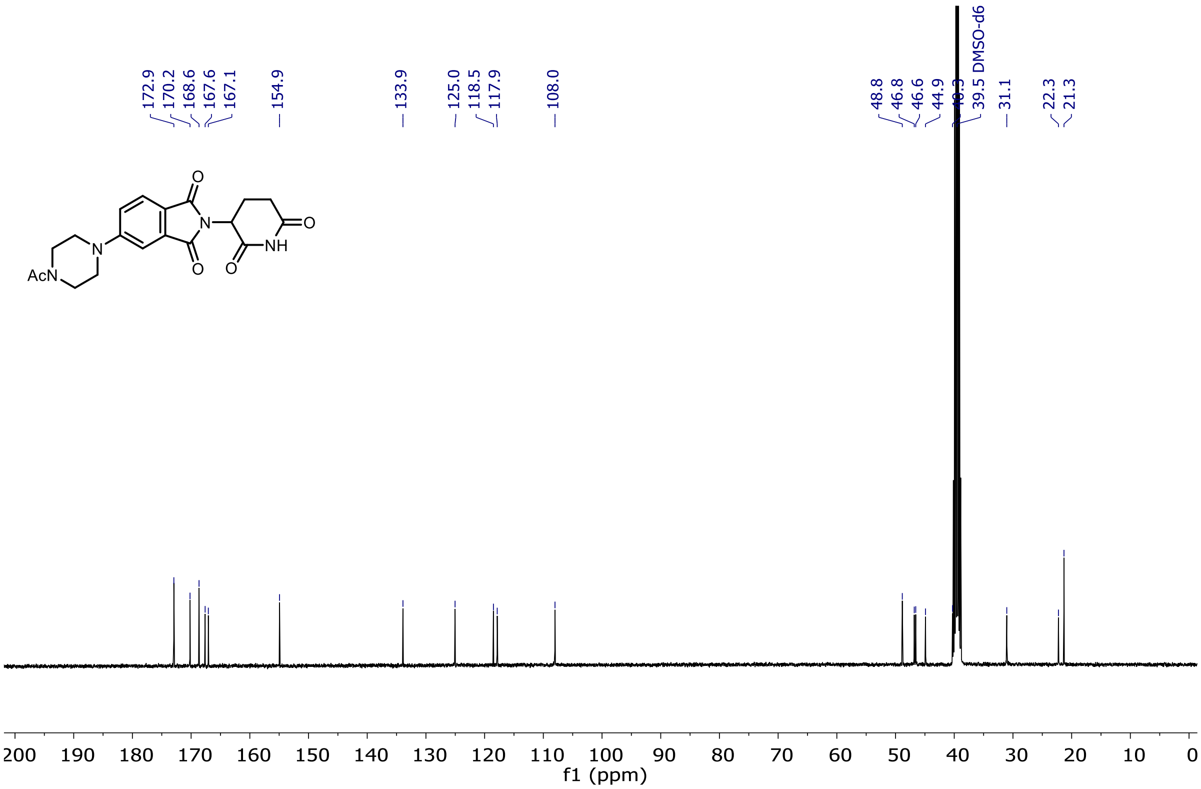

$^{1}$H and $^{13}$C NMR of Entry **40**

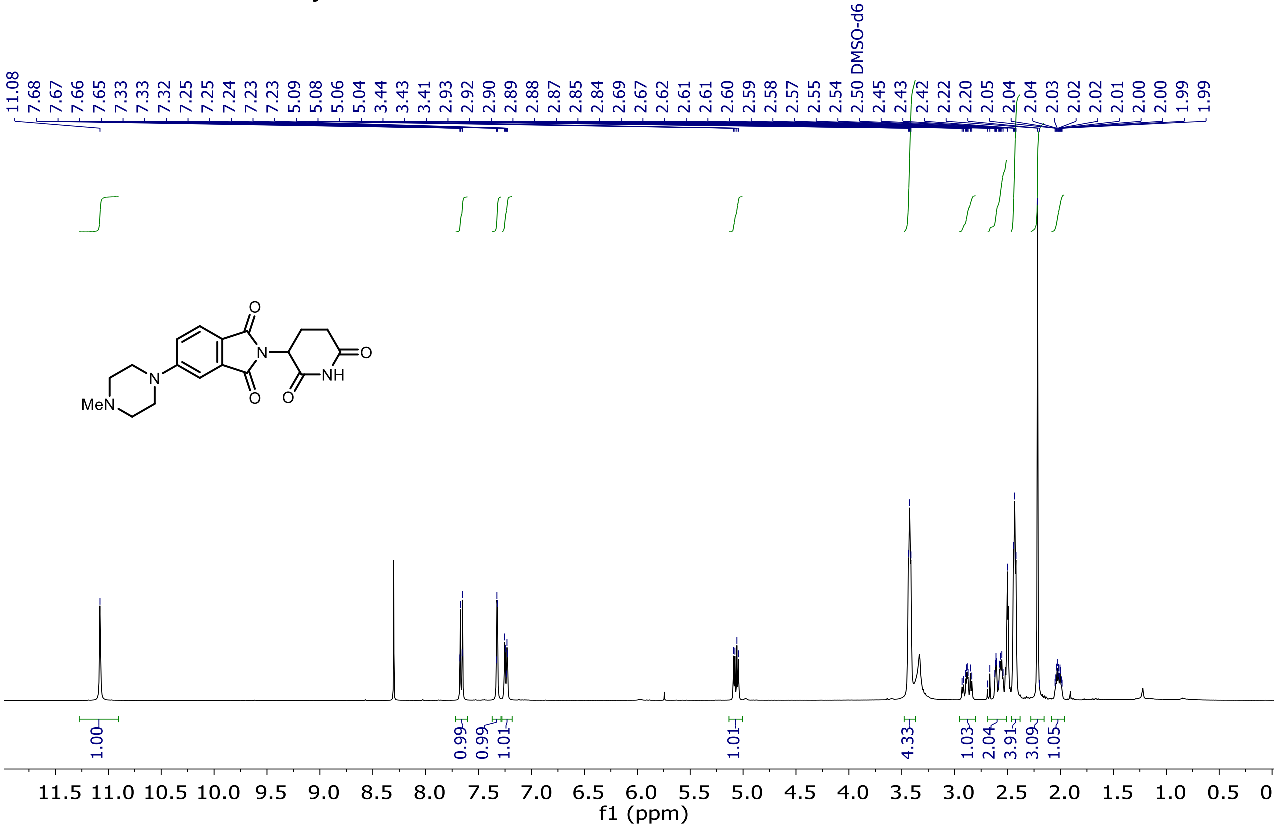

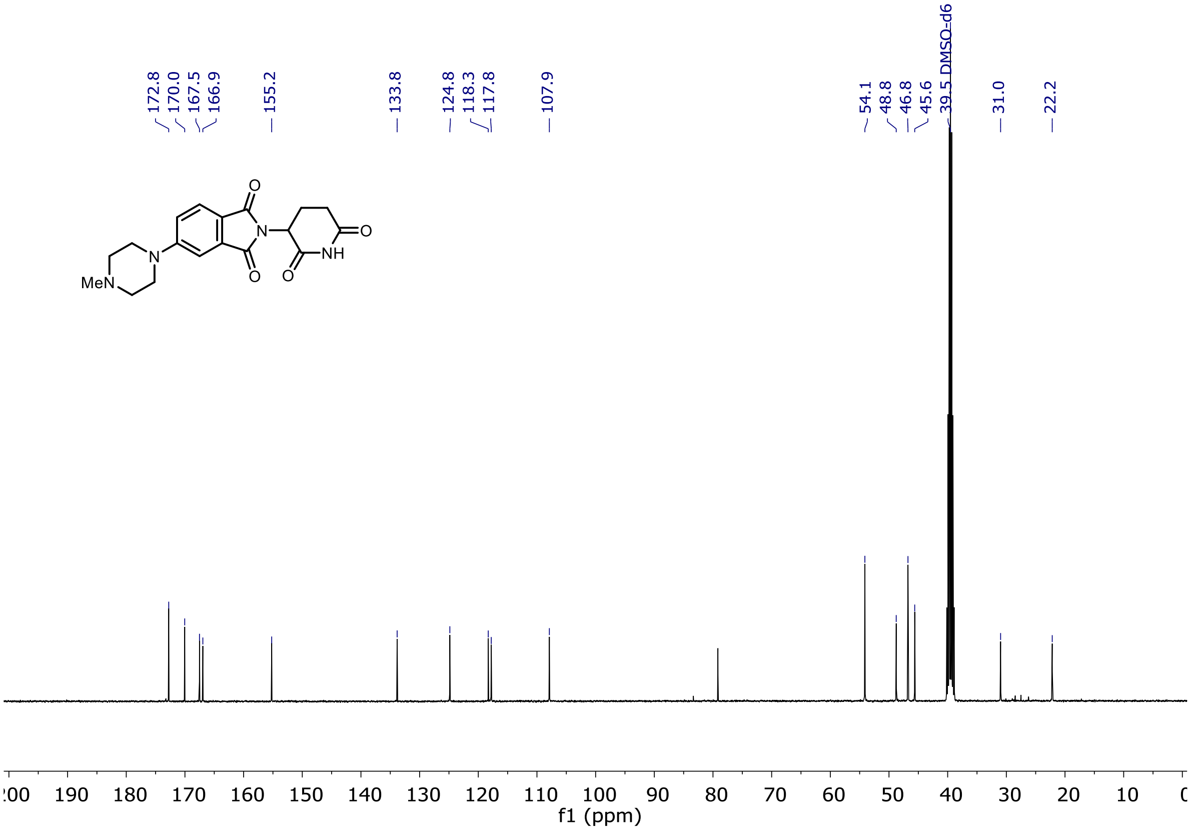

$^{1}H$ and $^{13}C$ NMR of Entry **41**

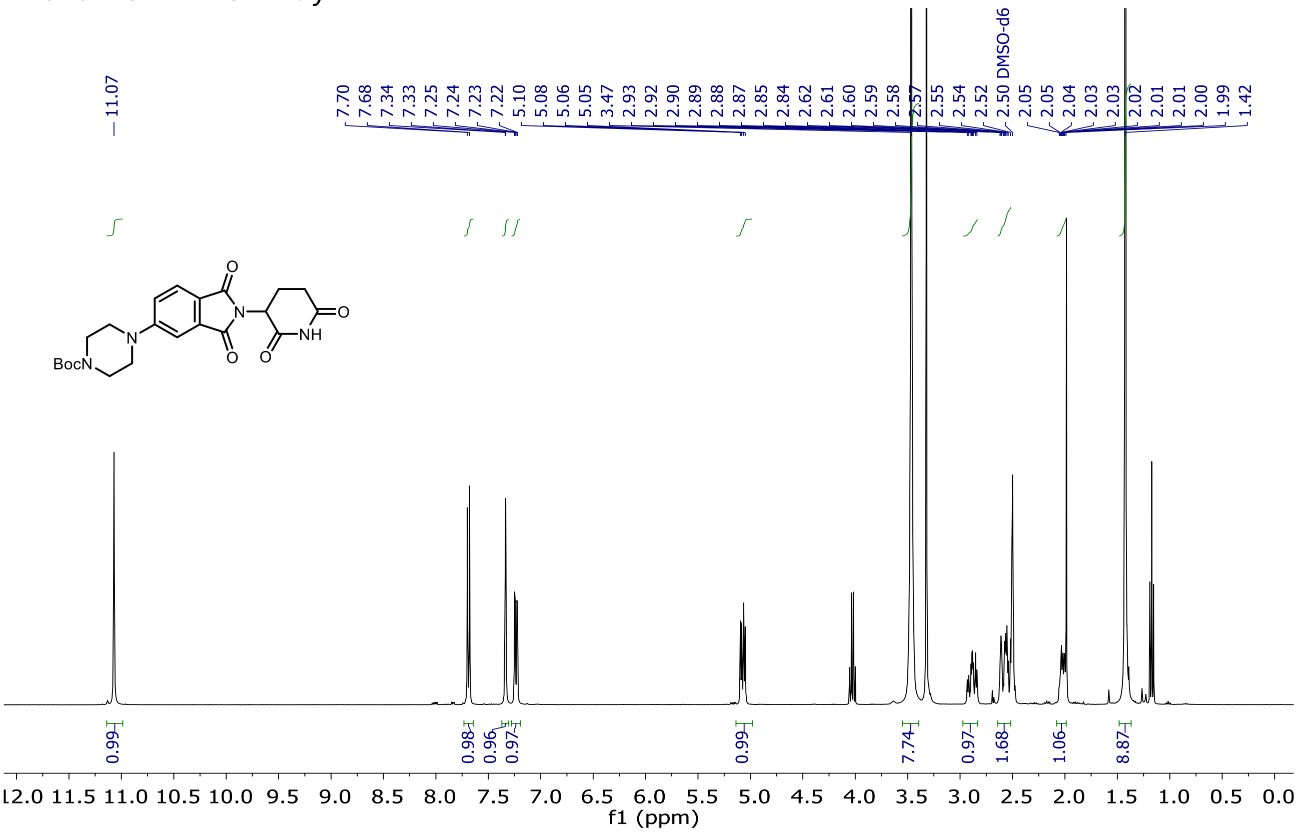

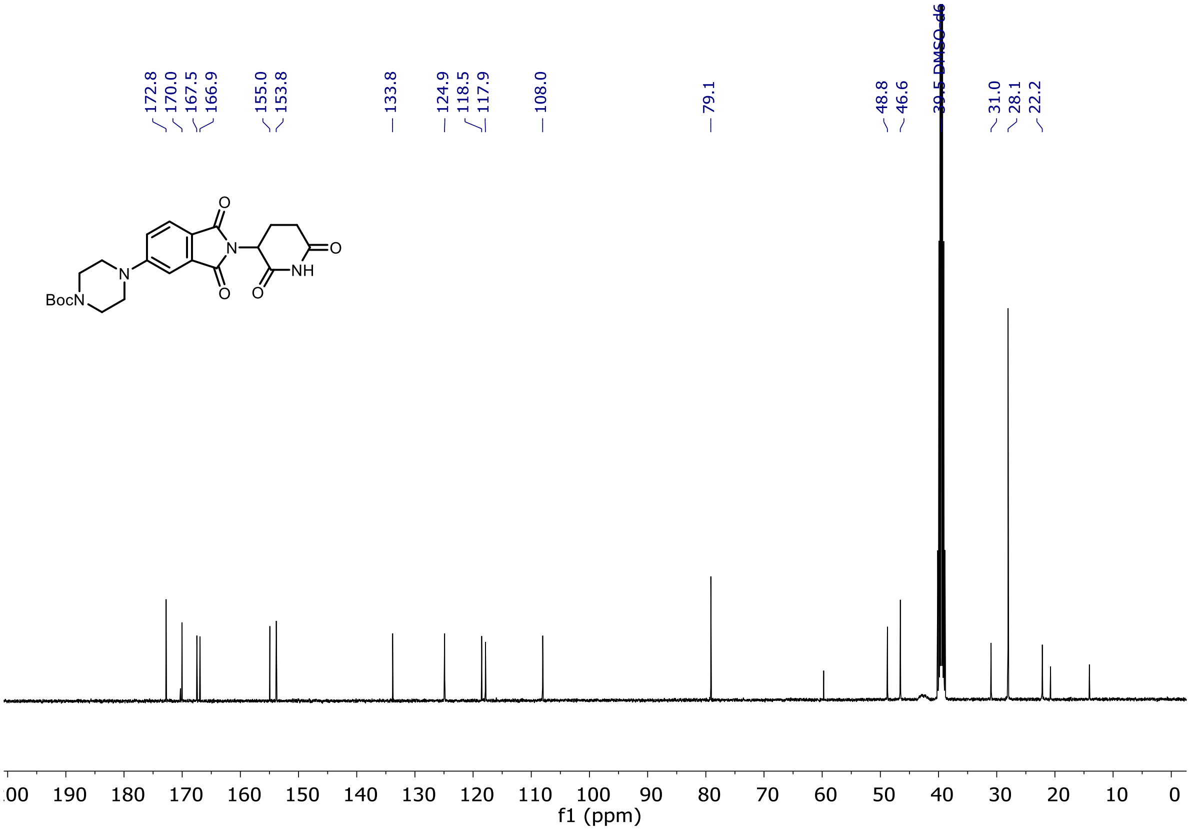

$^{1}H$ and $^{13}C$ NMR of Entry **42**

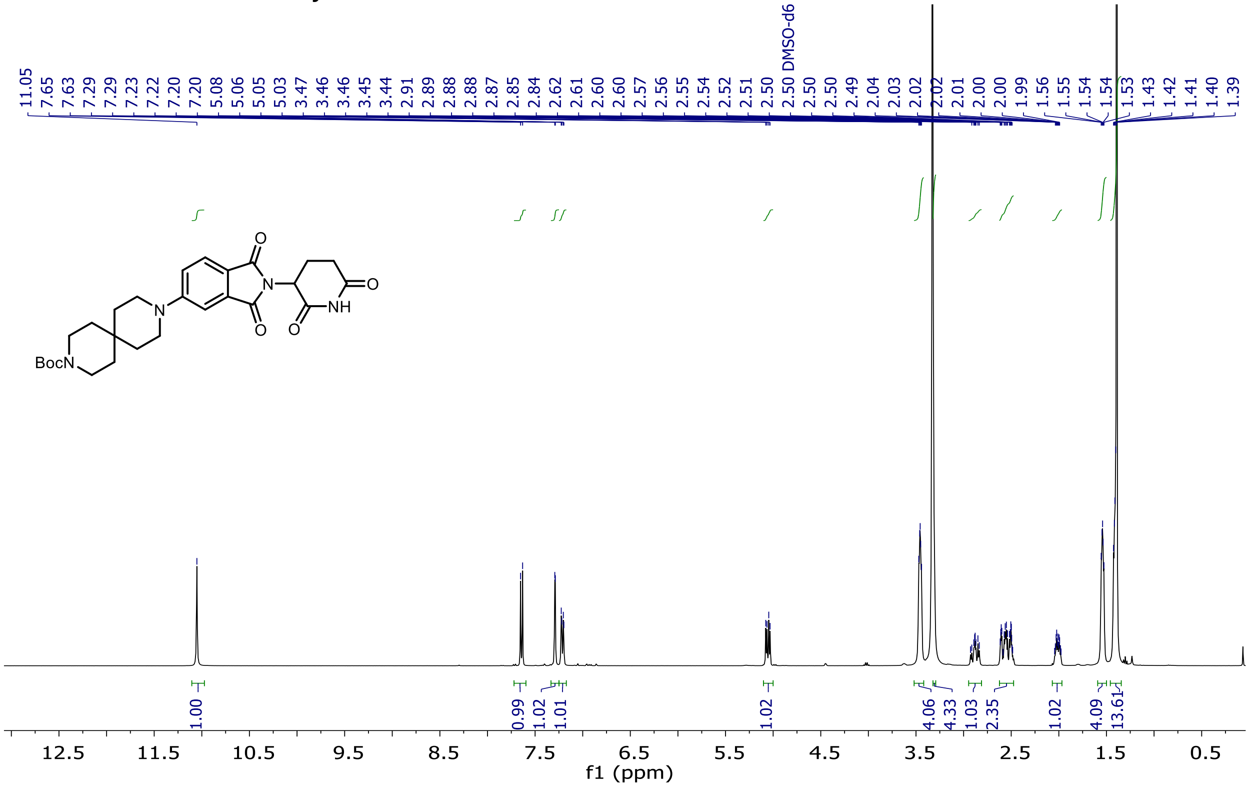

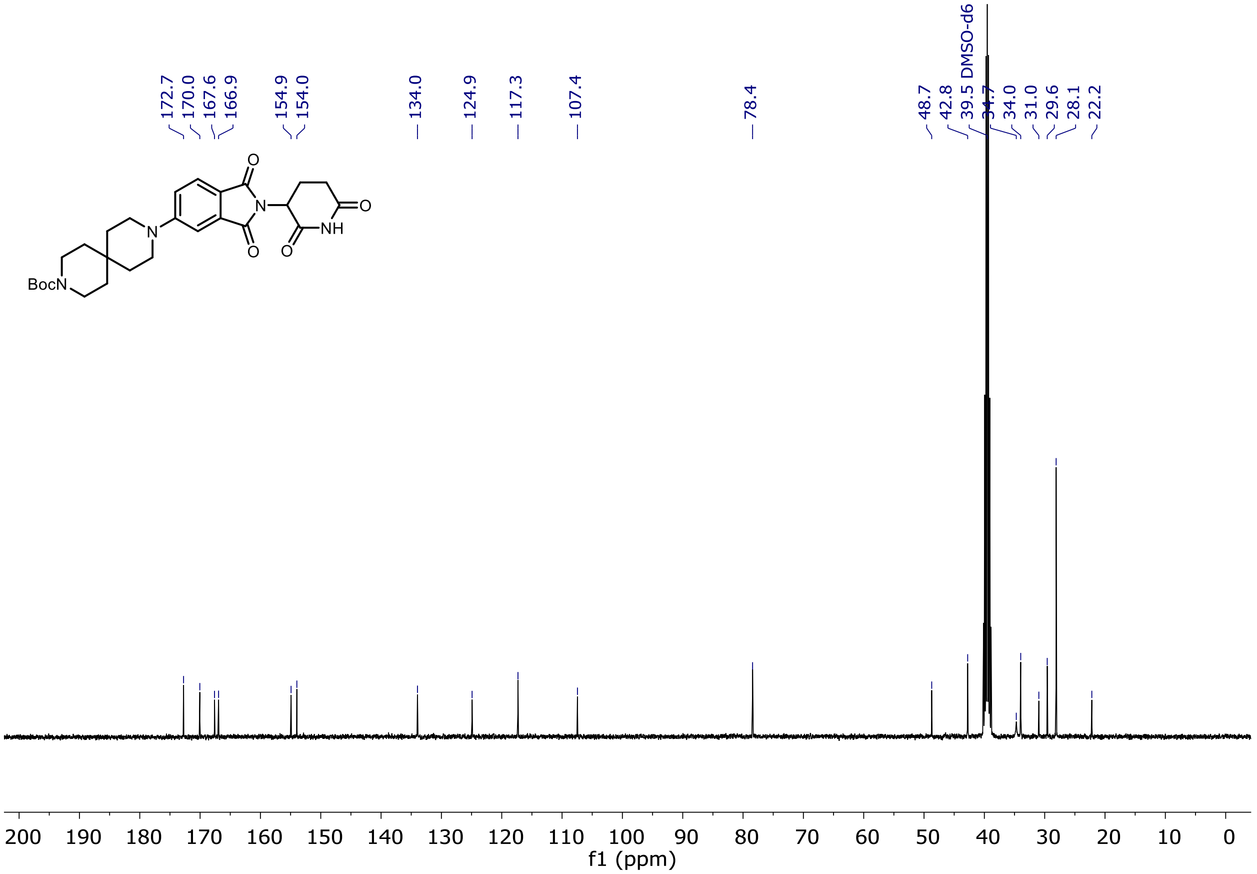

$^{1}$H and $^{13}$C NMR of Entry **43**

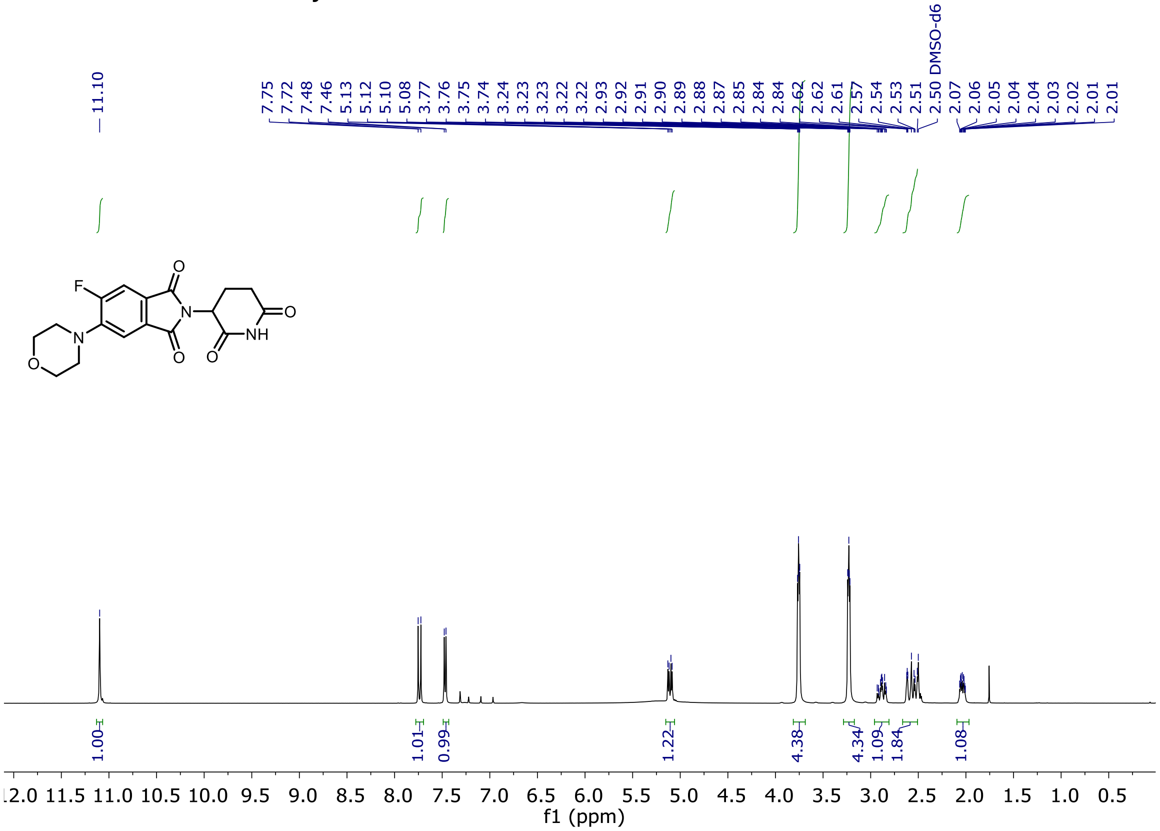

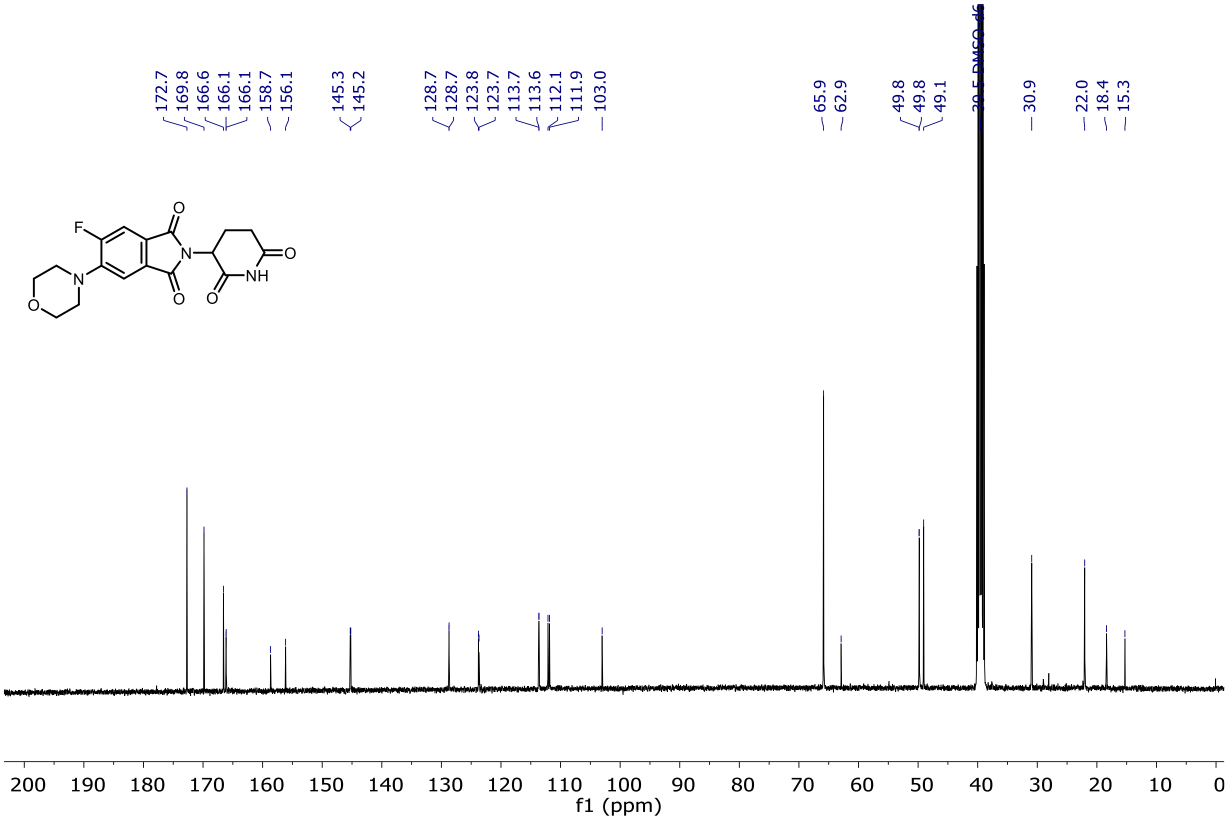

$^{19}$F NMR of Entry **43**

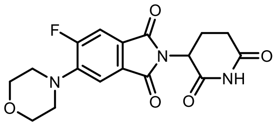

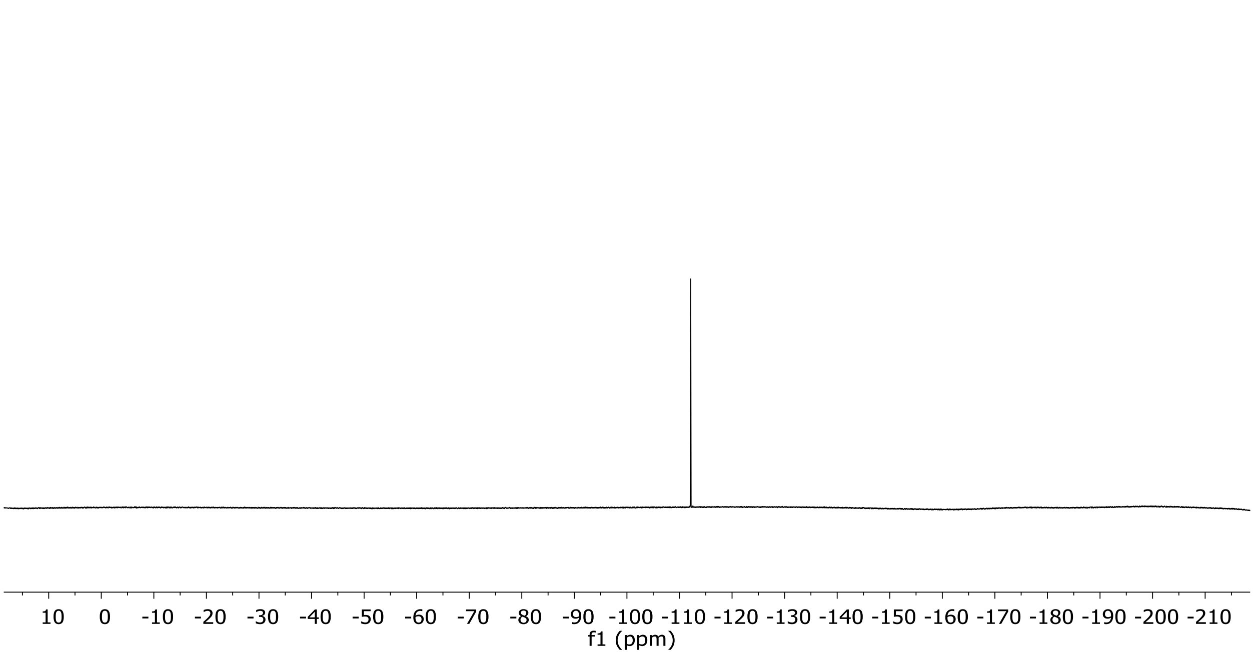

[1]H and [13]C NMR of Entry **44**

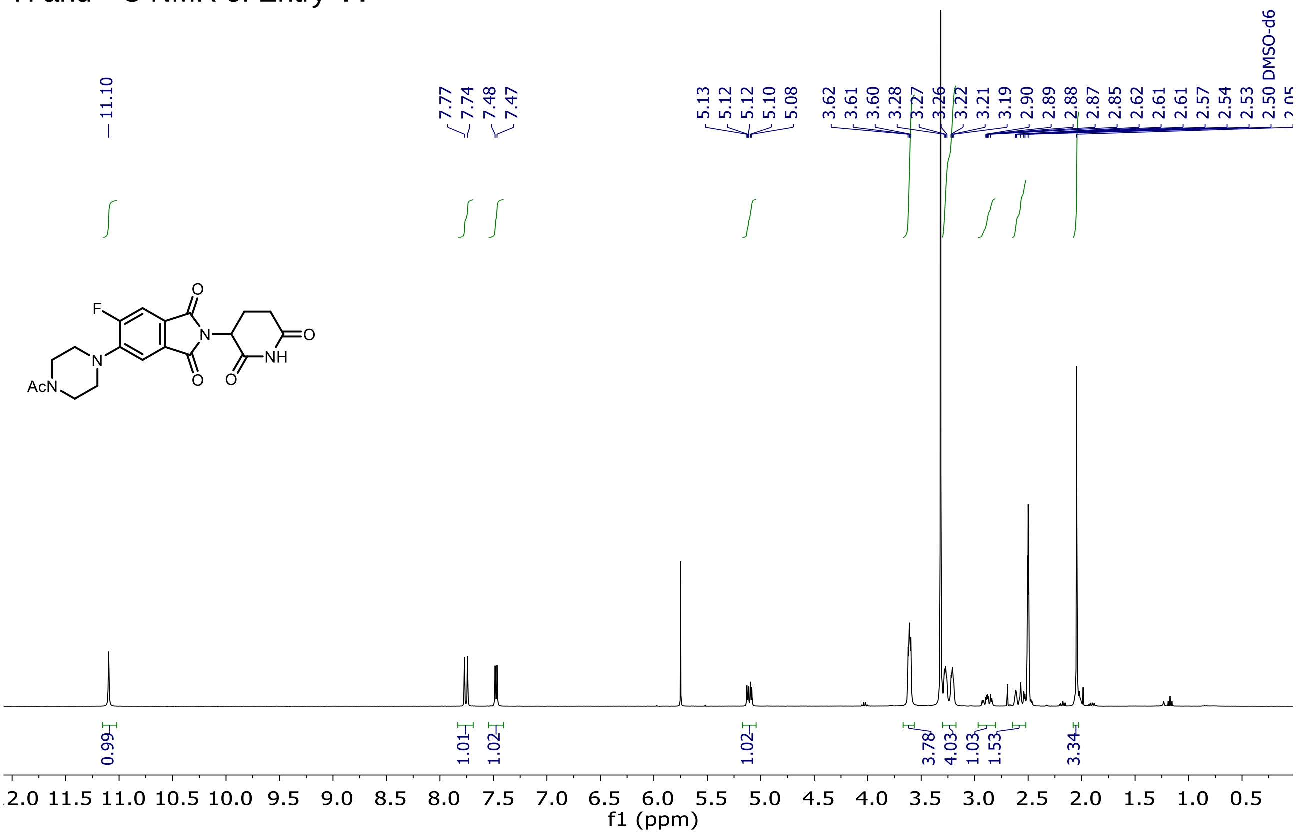

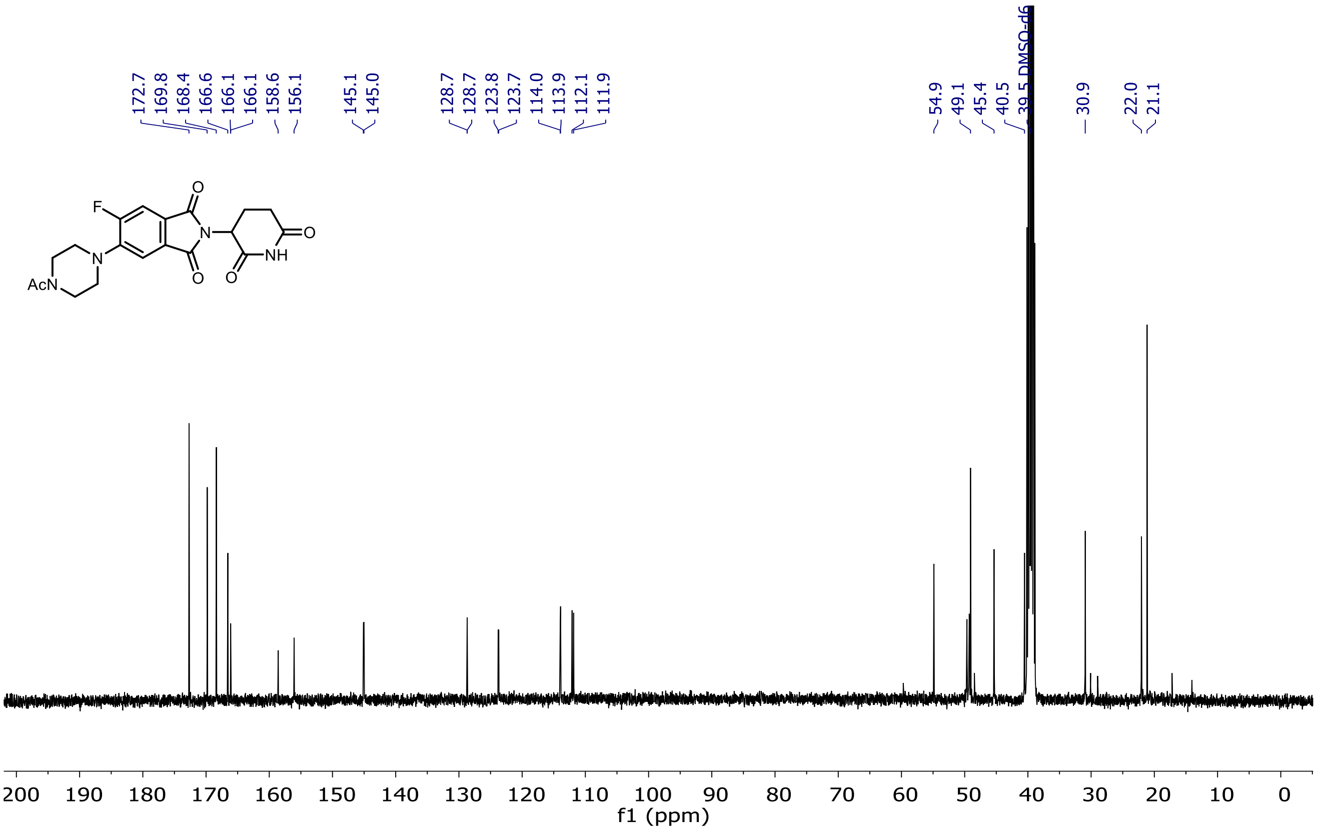

$^{19}$F NMR of Entry **44**

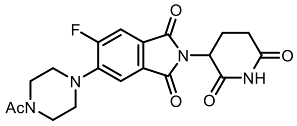

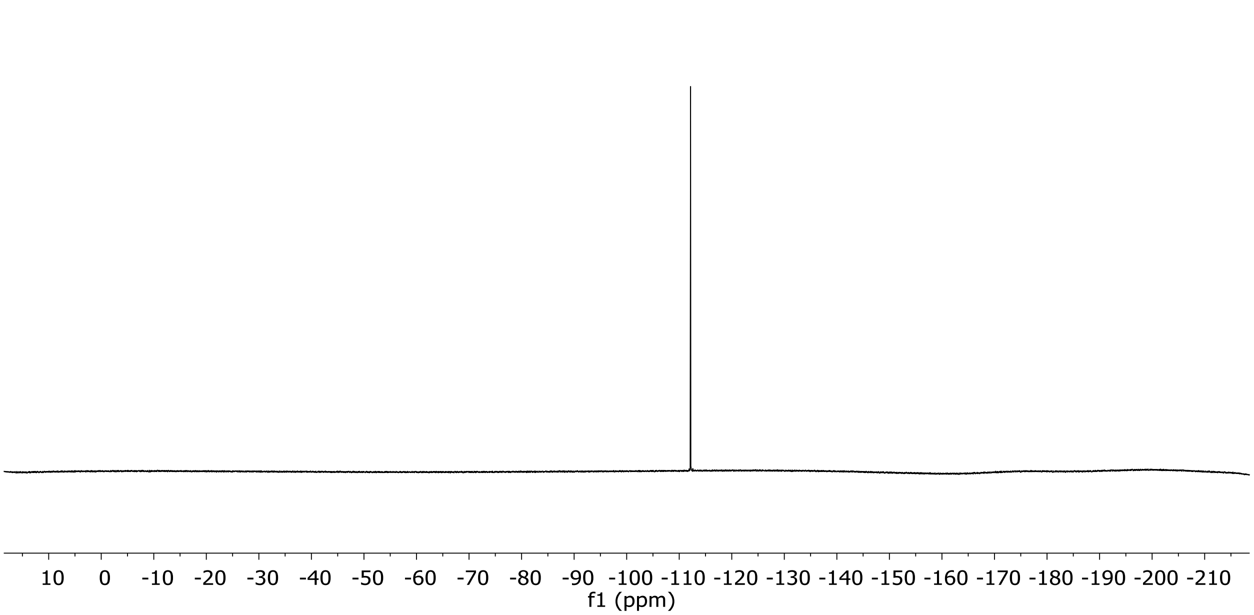

<sup></sup>

$^{1}$H and $^{13}$C NMR of Entry **45**

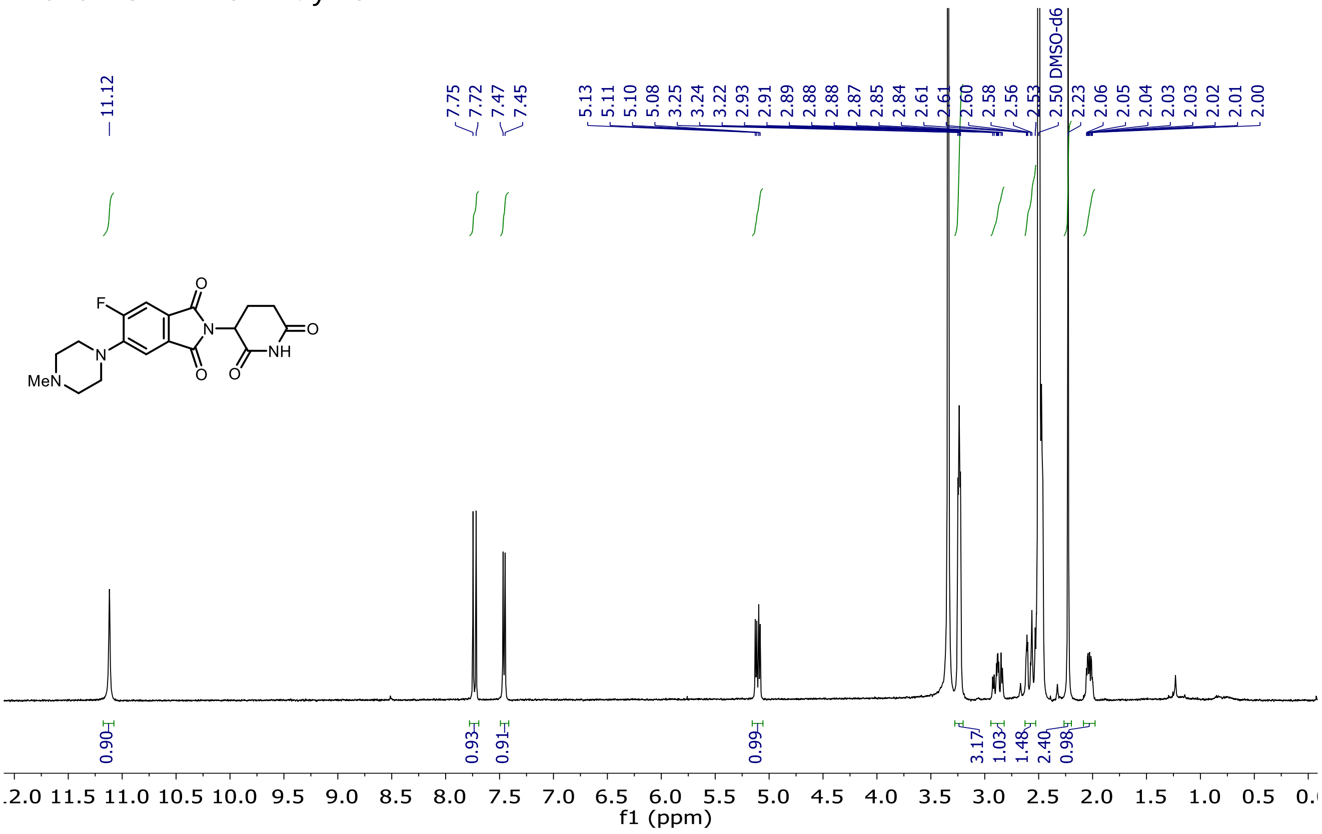

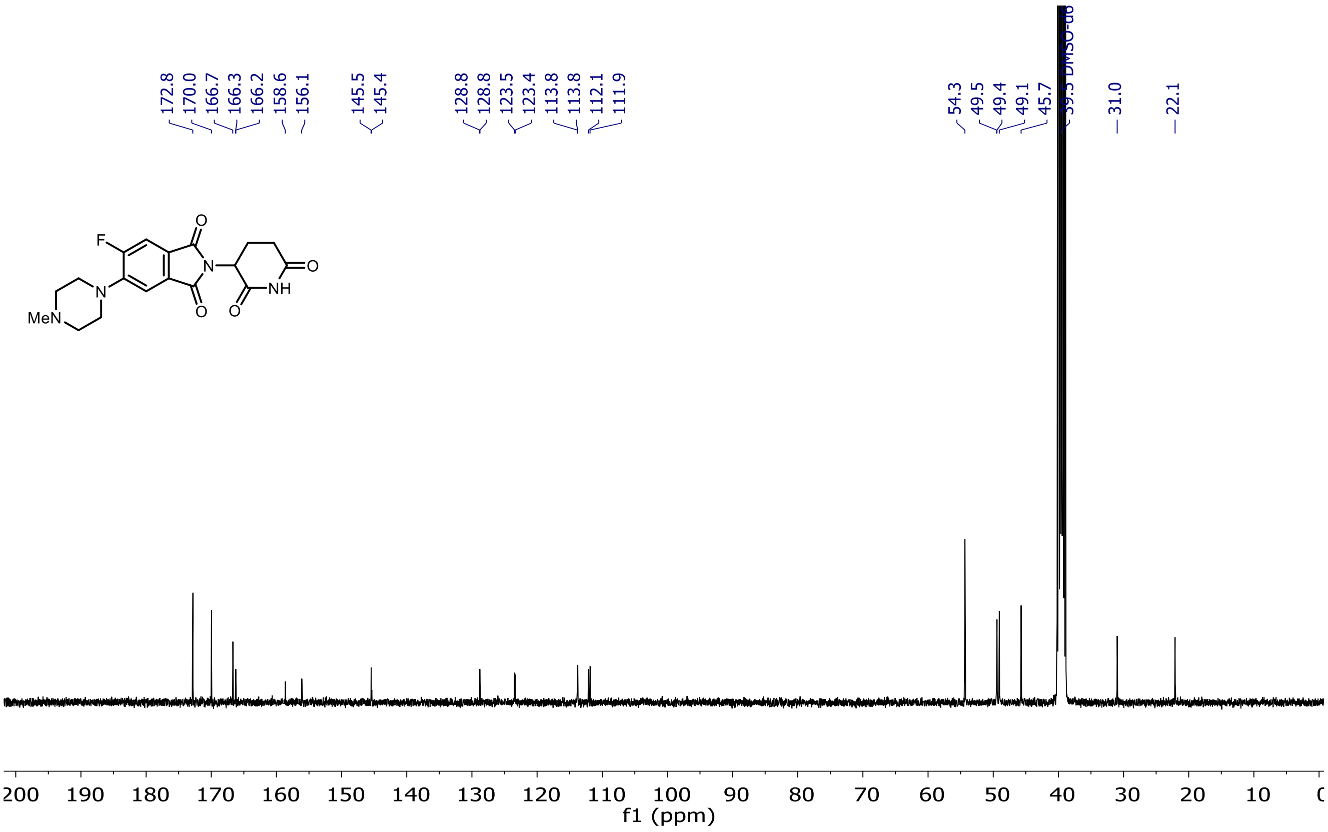

$^{19}$F NMR of Entry **45**

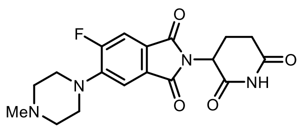

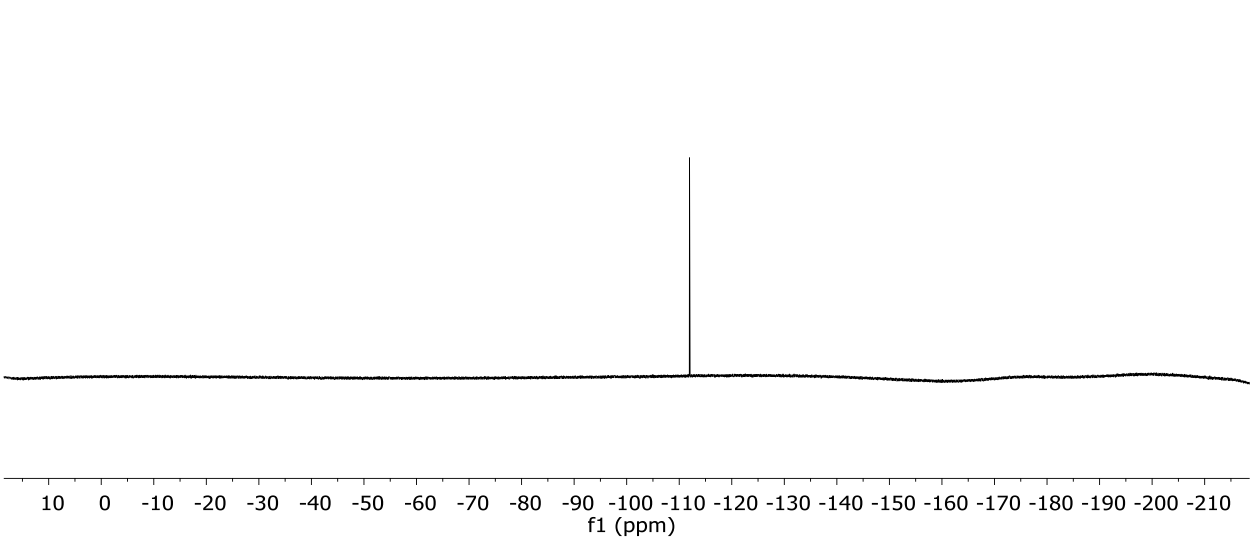

<sup></sup>

$^{1}$H and $^{13}$C NMR of Entry **46**

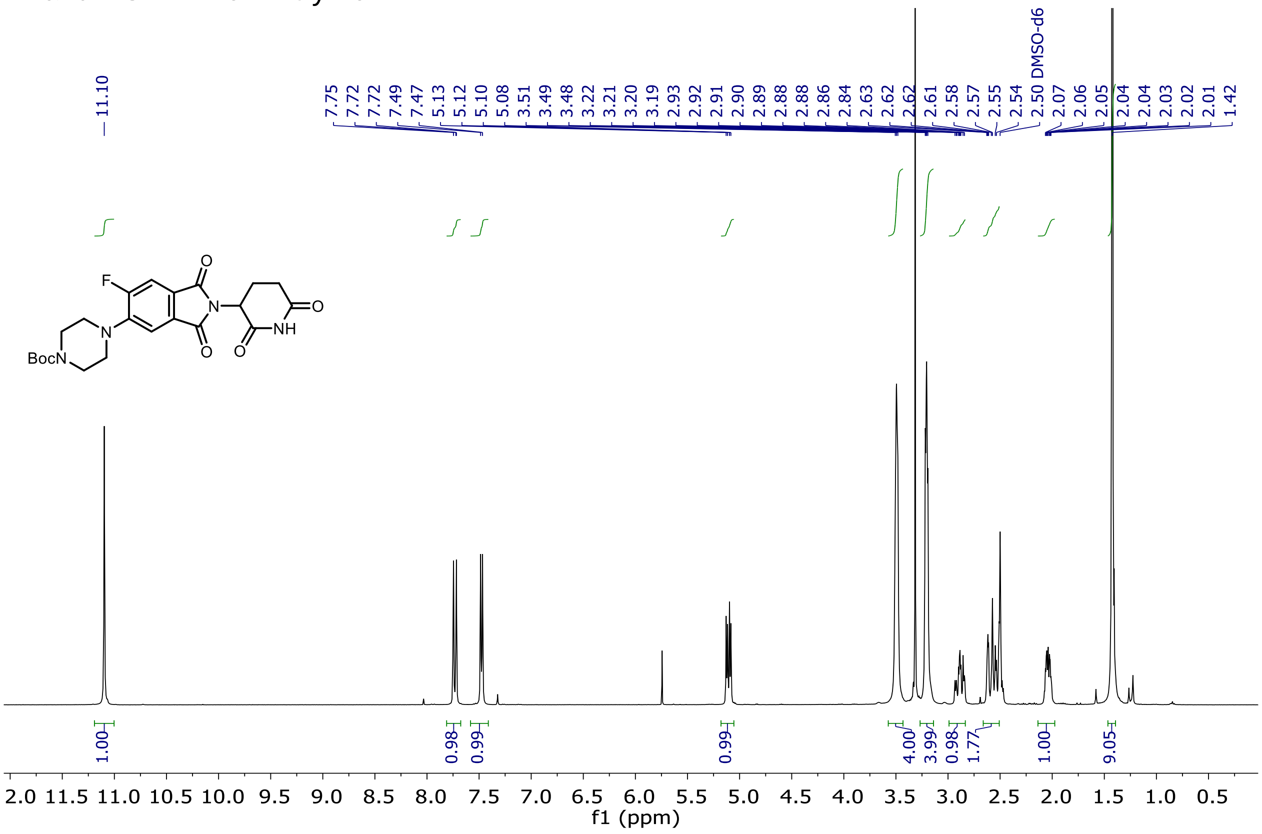

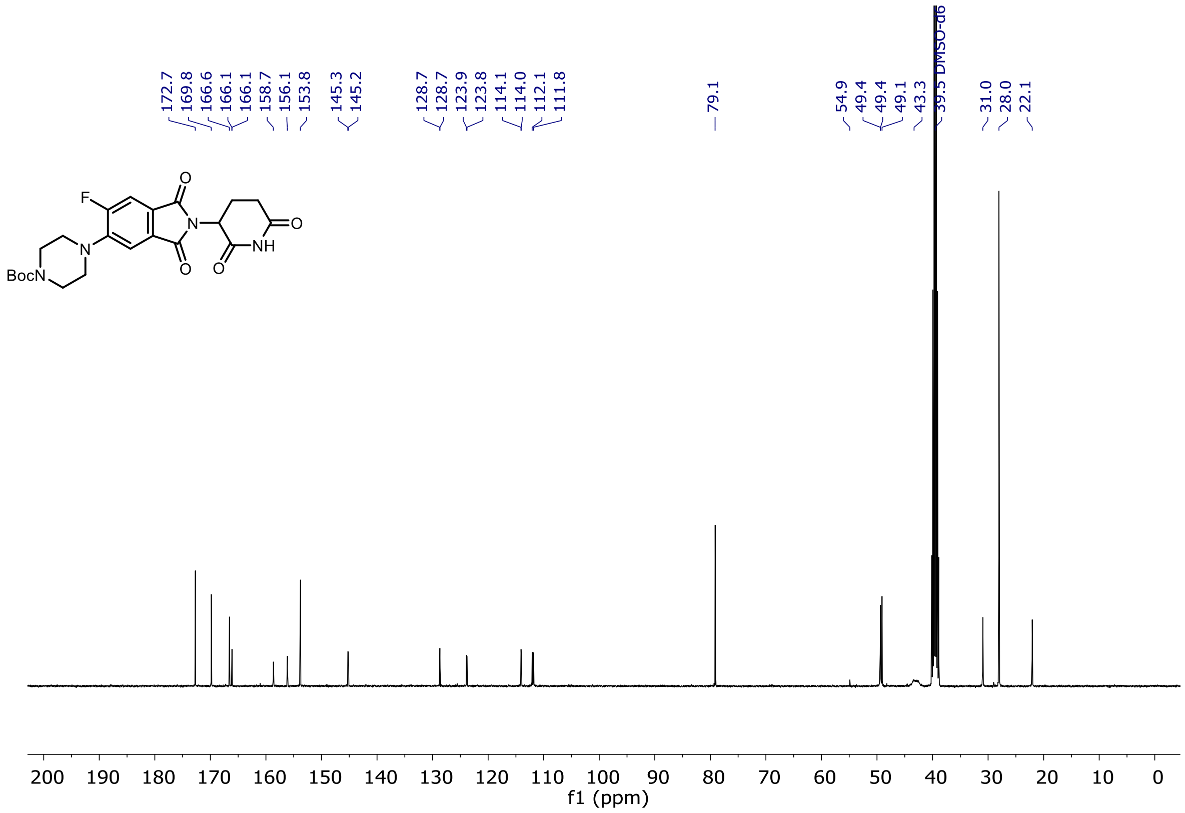

$^{19}F$ NMR of Entry **46**

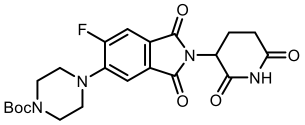

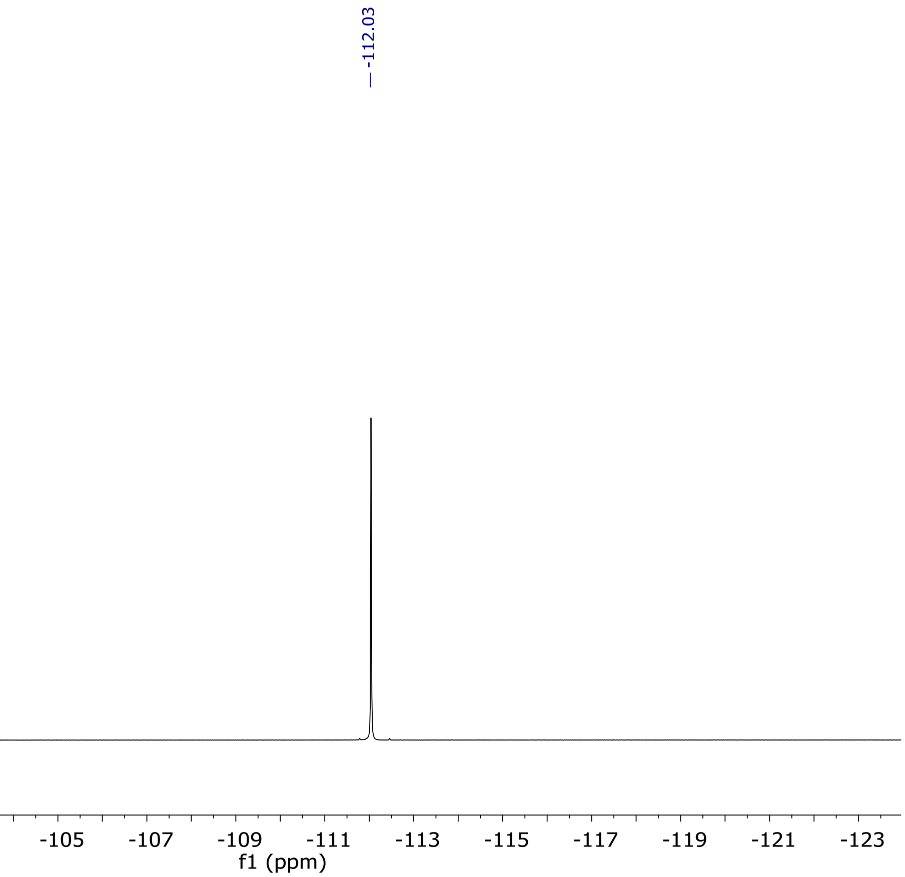

¹H and ¹³C NMR of Entry **47**

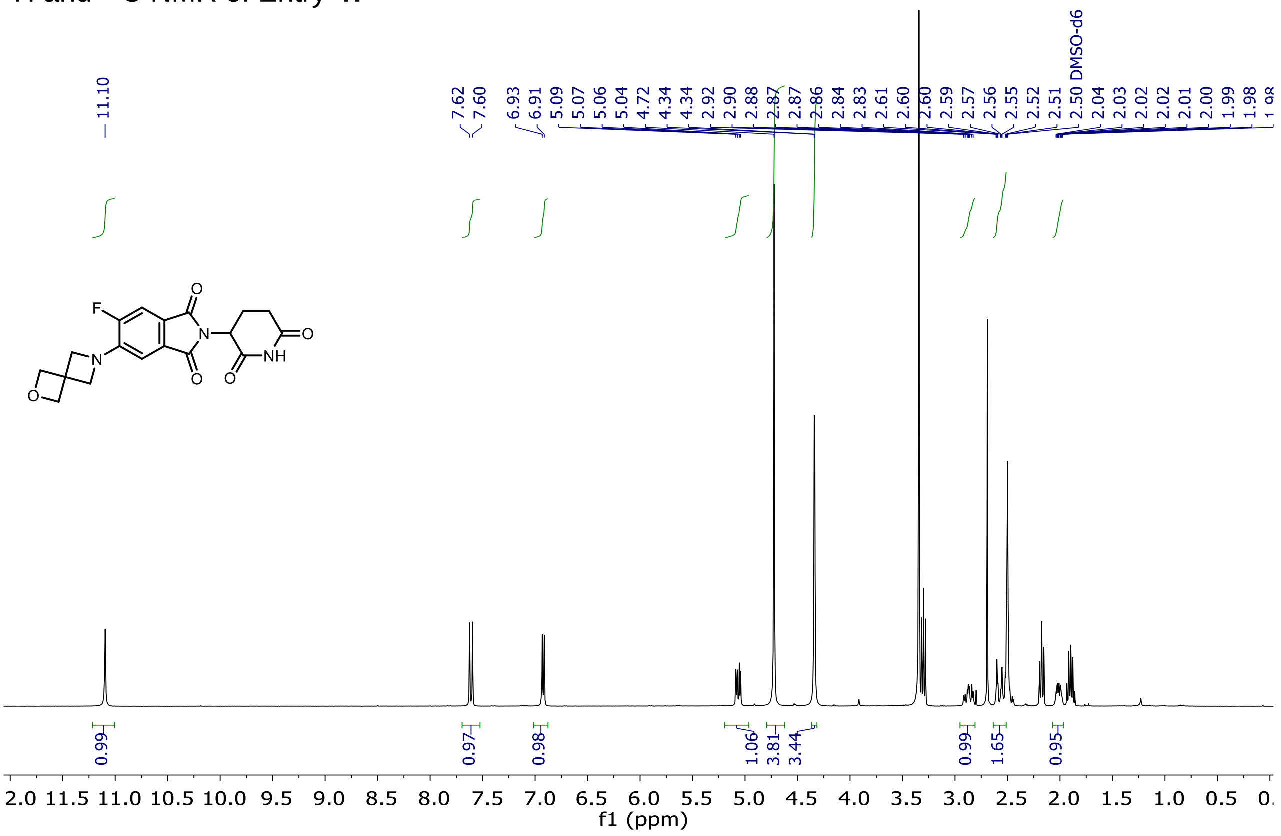

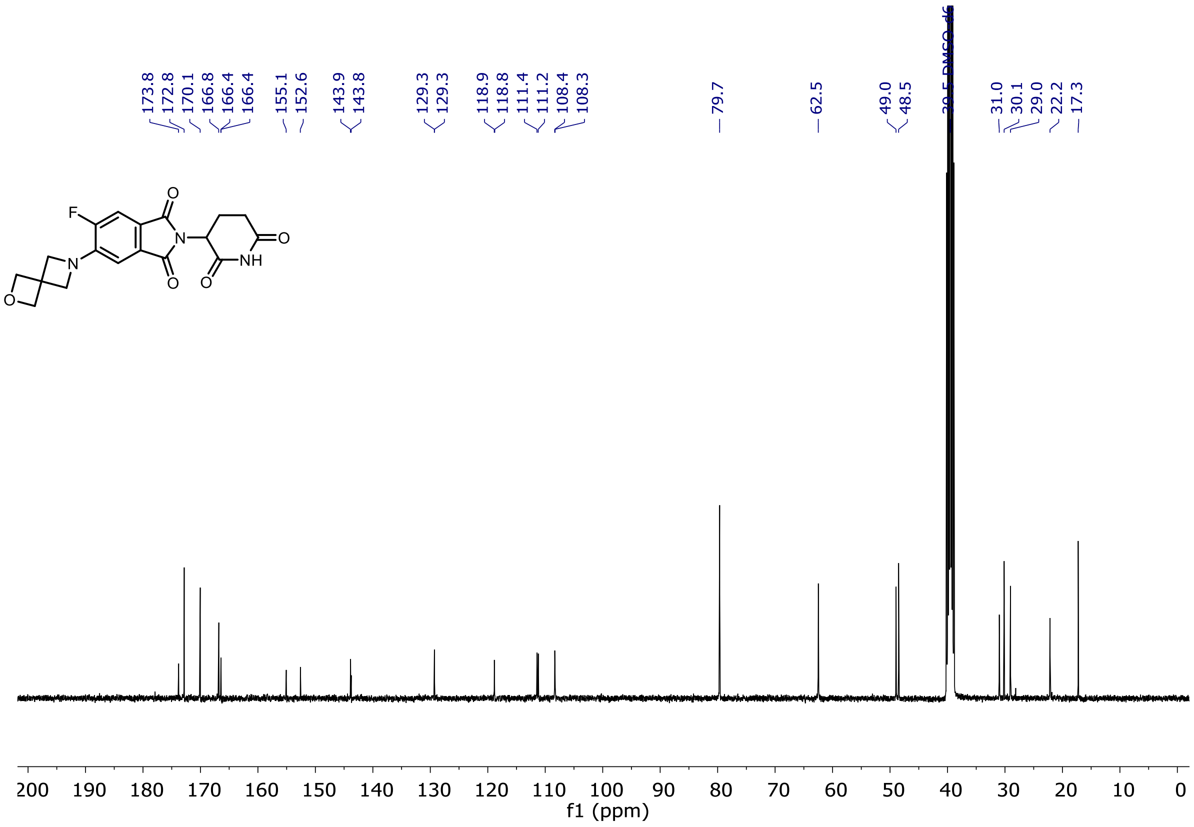

$^{19}$F NMR of Entry **47**

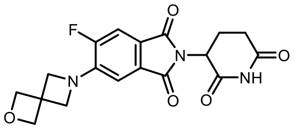

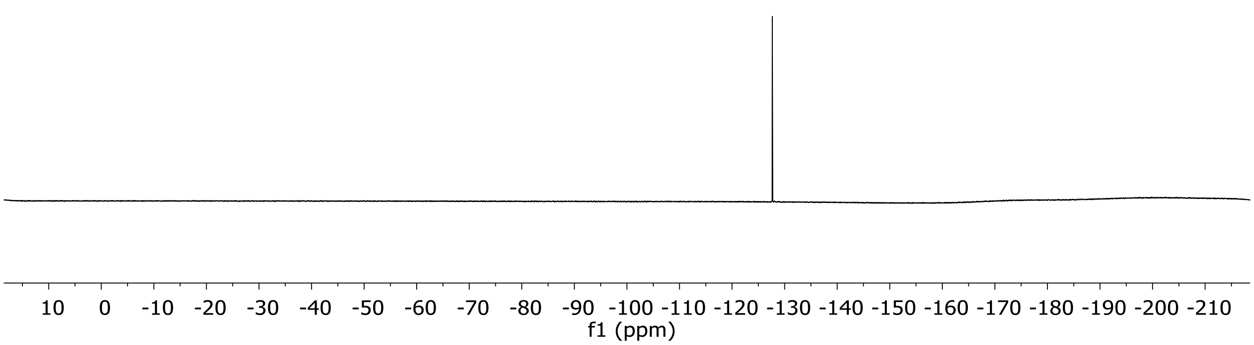

$^{1}H$ and $^{13}C$ NMR of Entry **48**

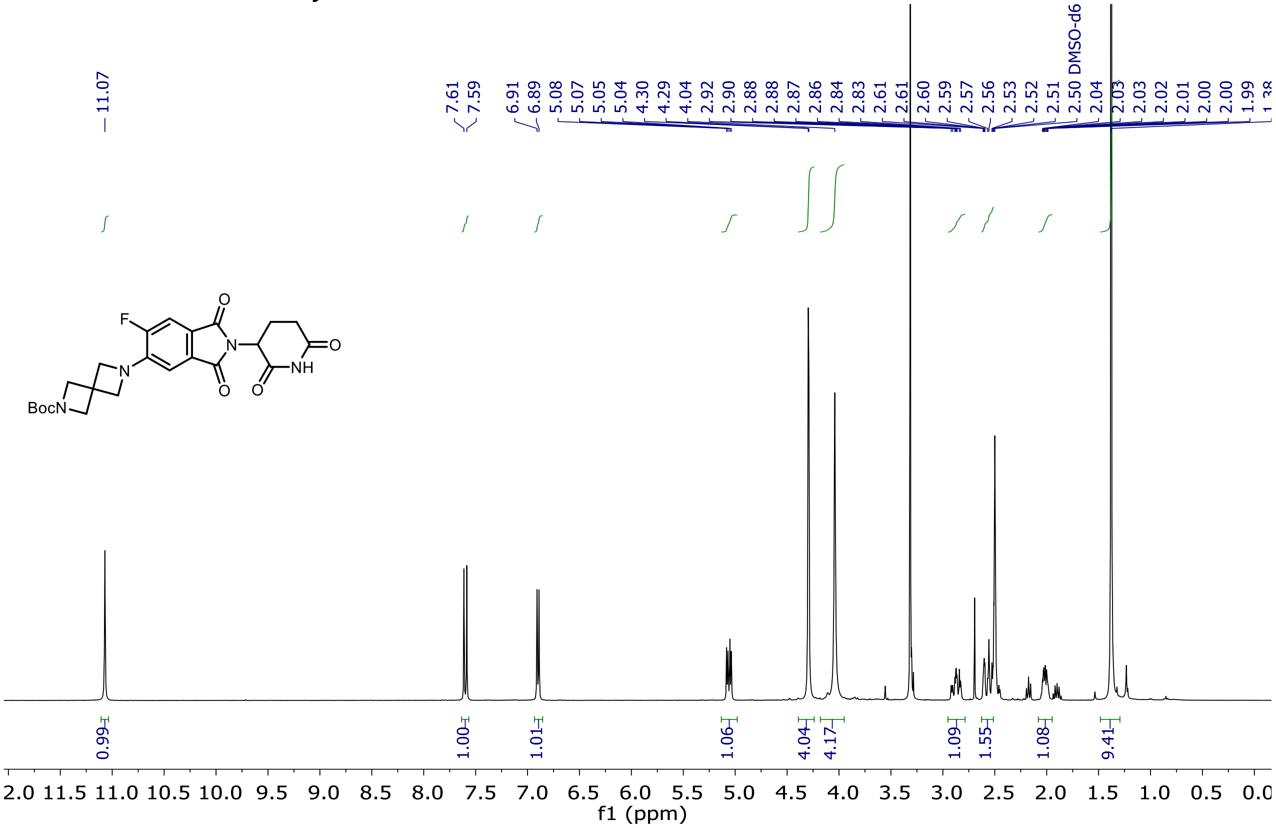

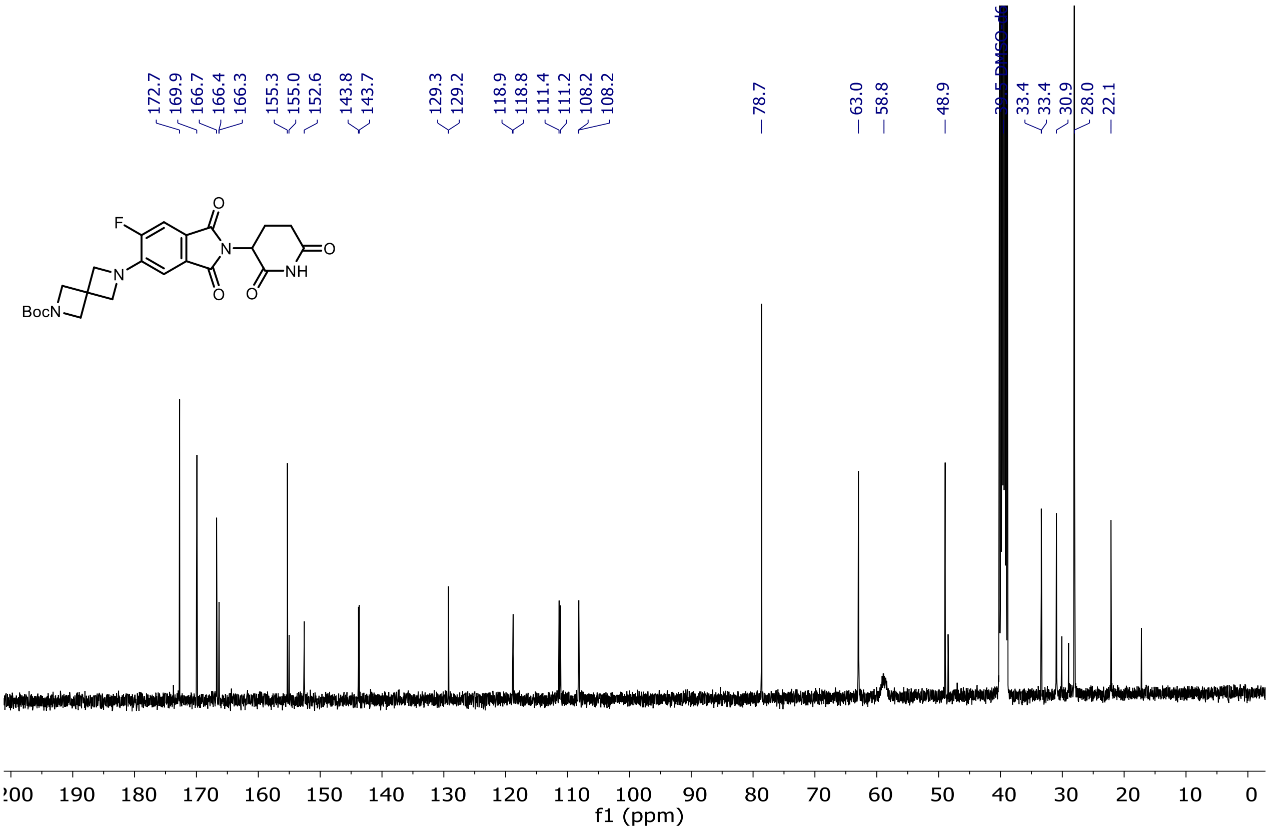

$^{19}$F NMR of Entry **48**

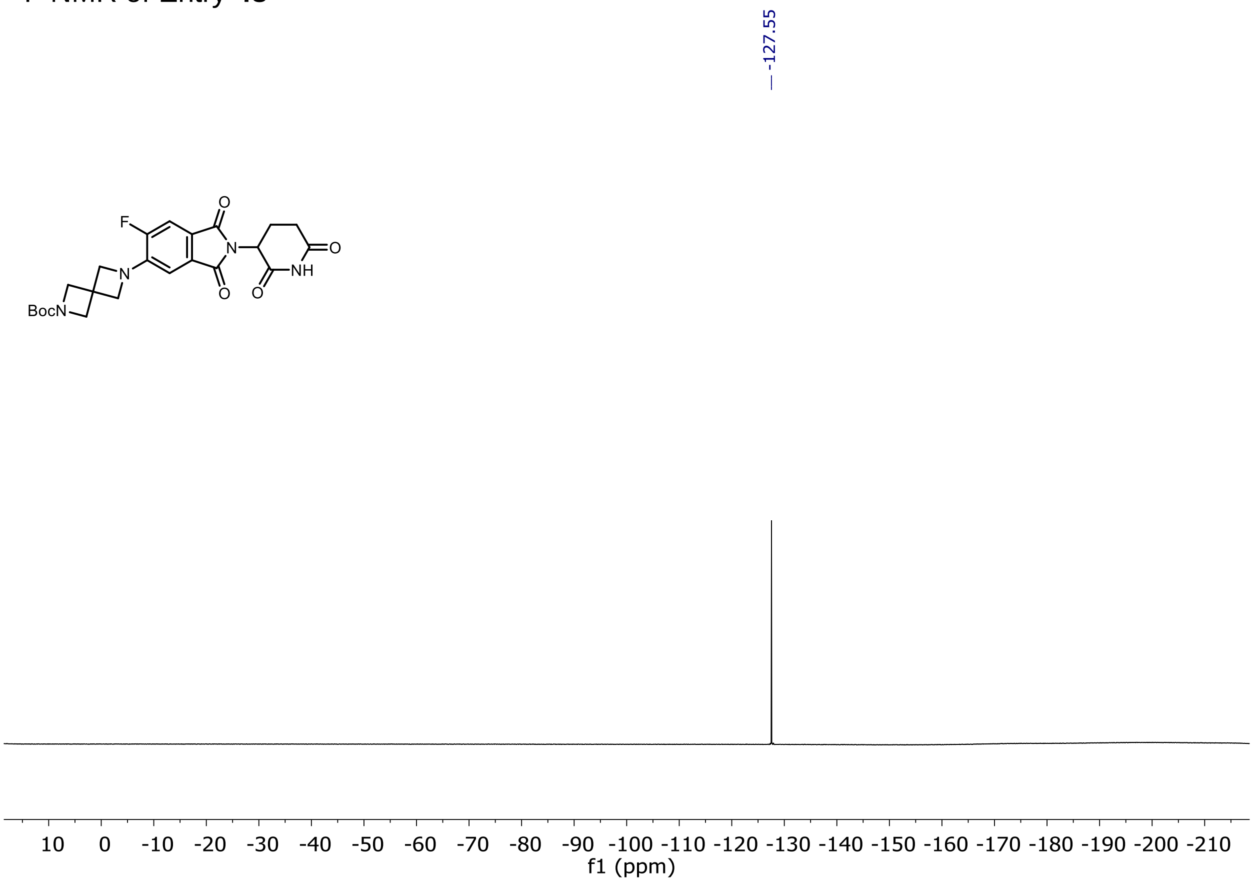

$^{1}H$ and $^{13}C$ NMR of Entry **49**

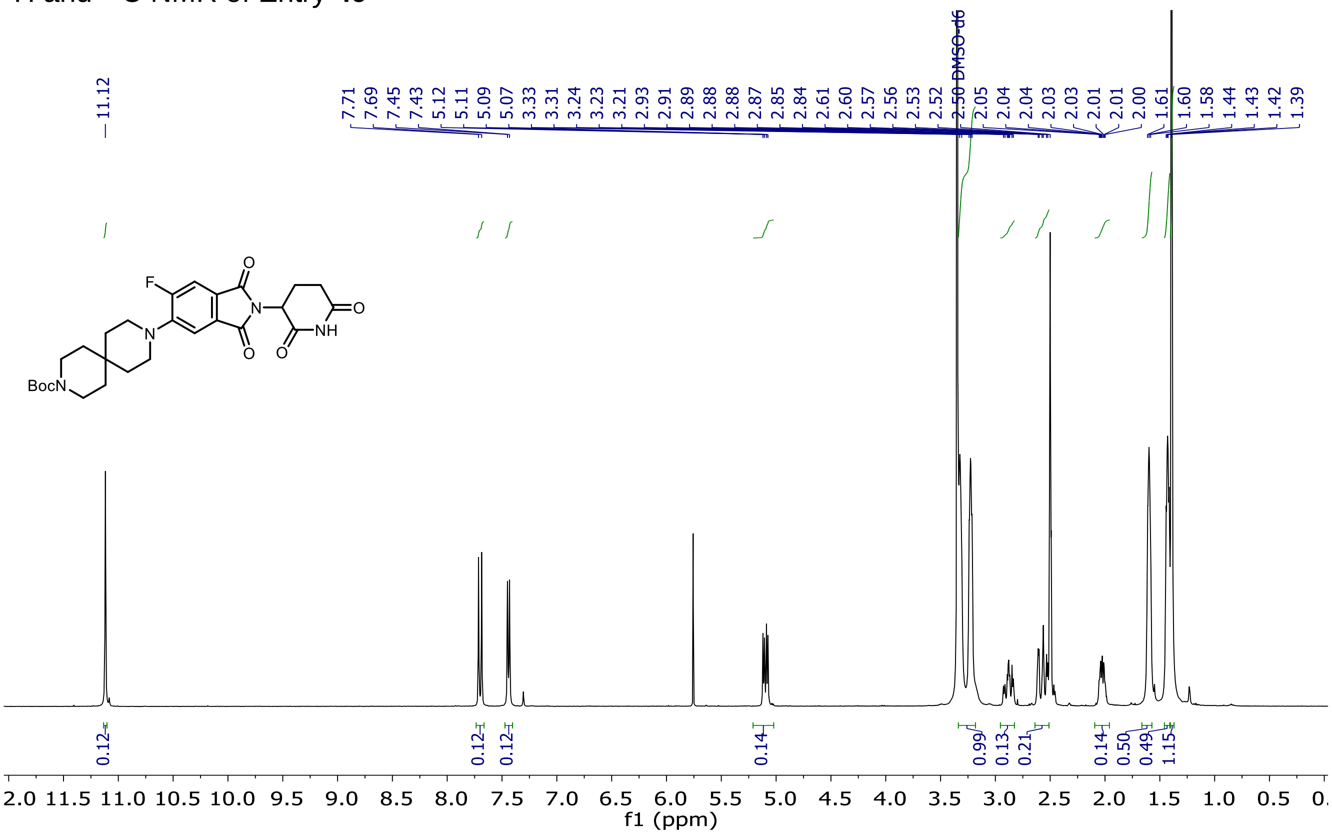

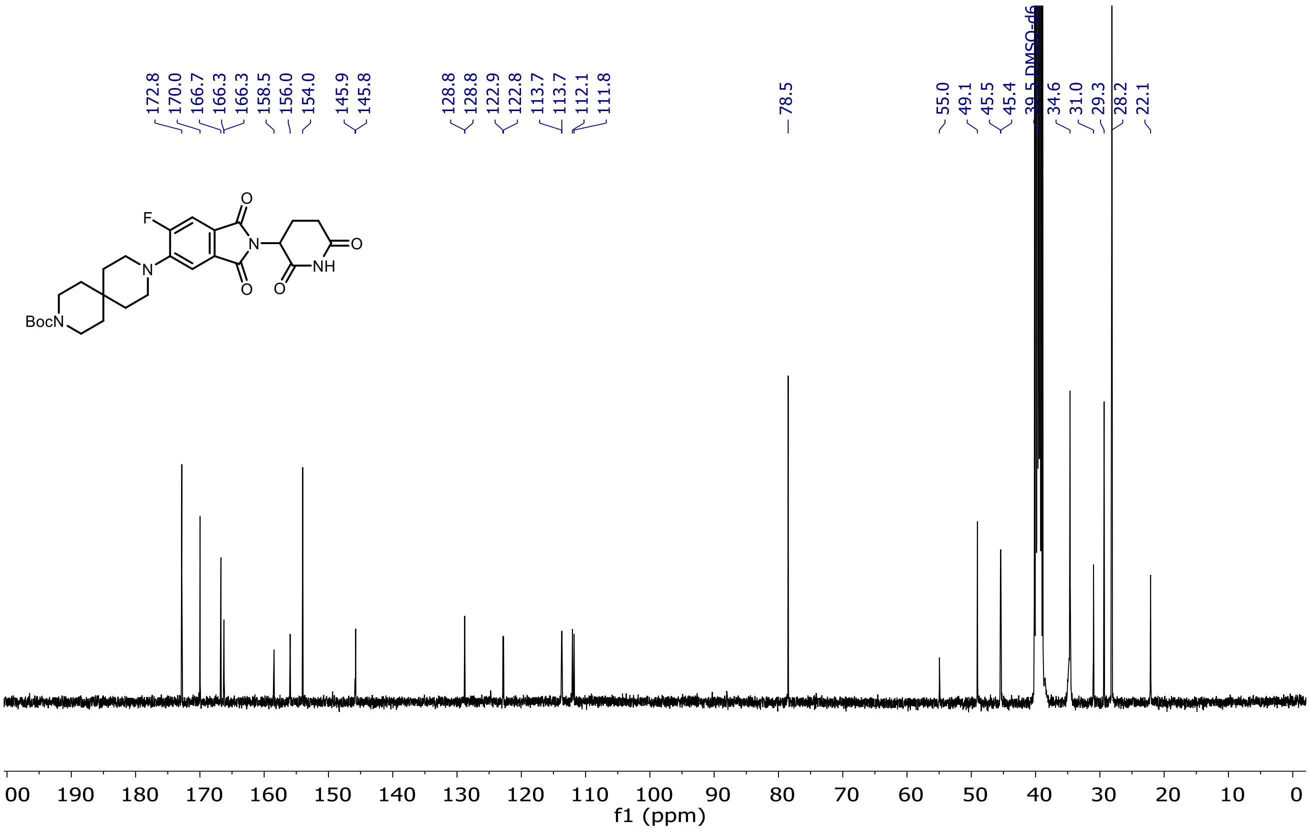

[19]F NMR of Entry **49**

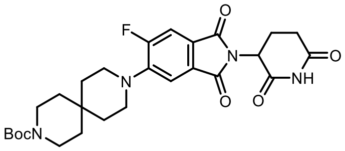

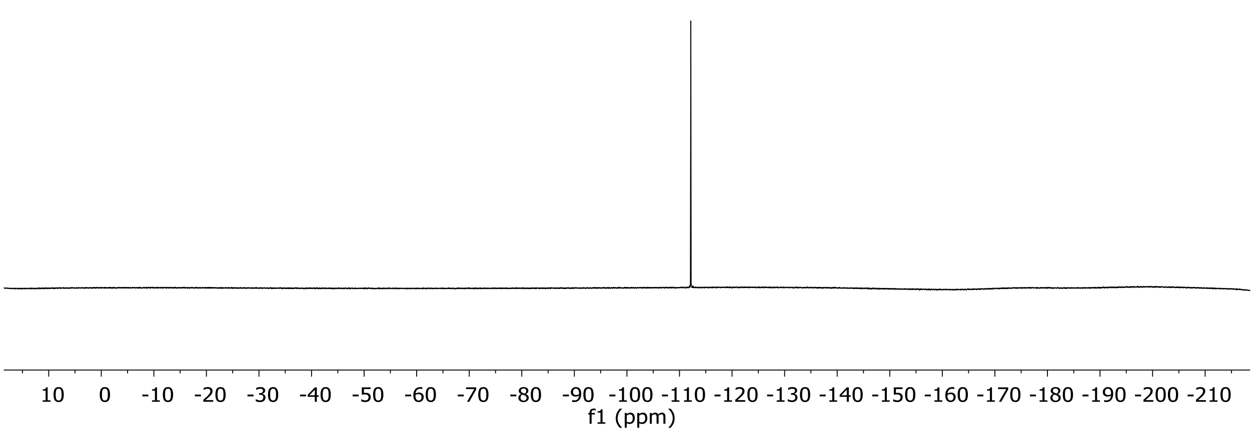

$^{1}H$ and $^{13}C$ NMR of Entry **51**

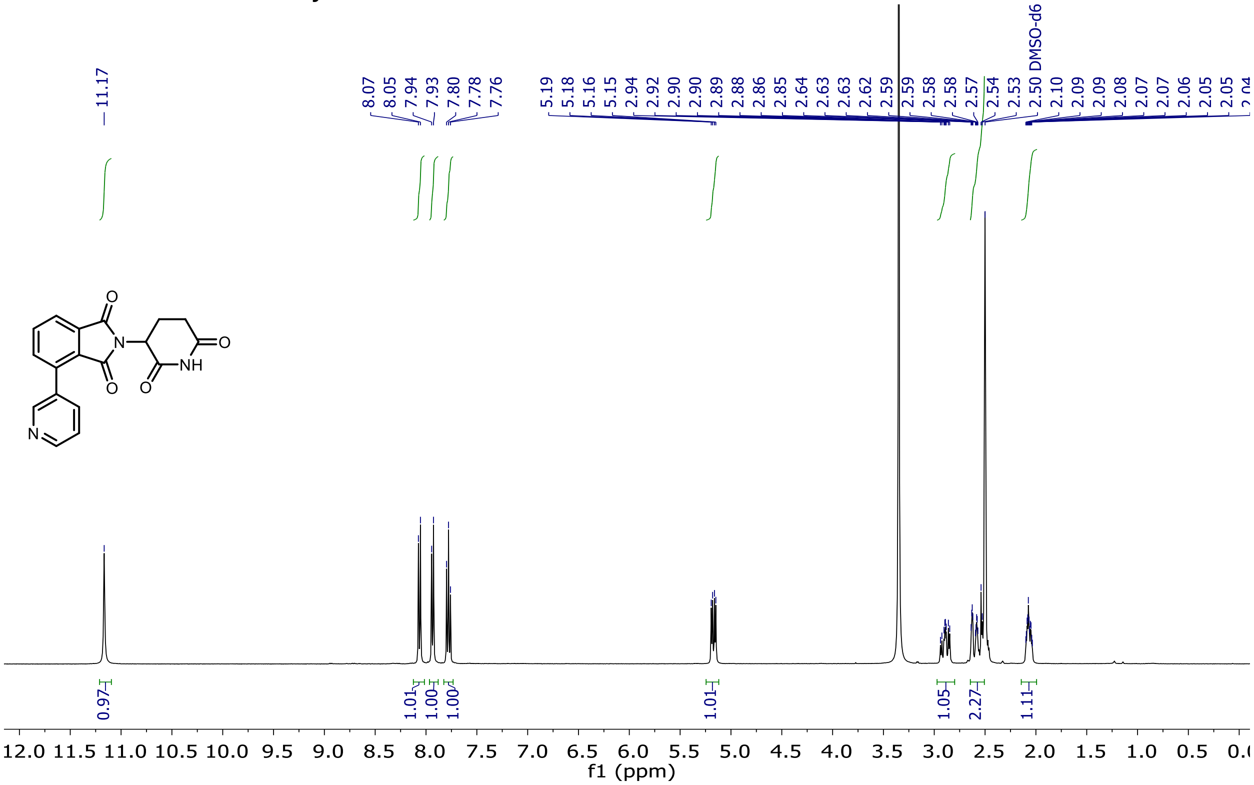

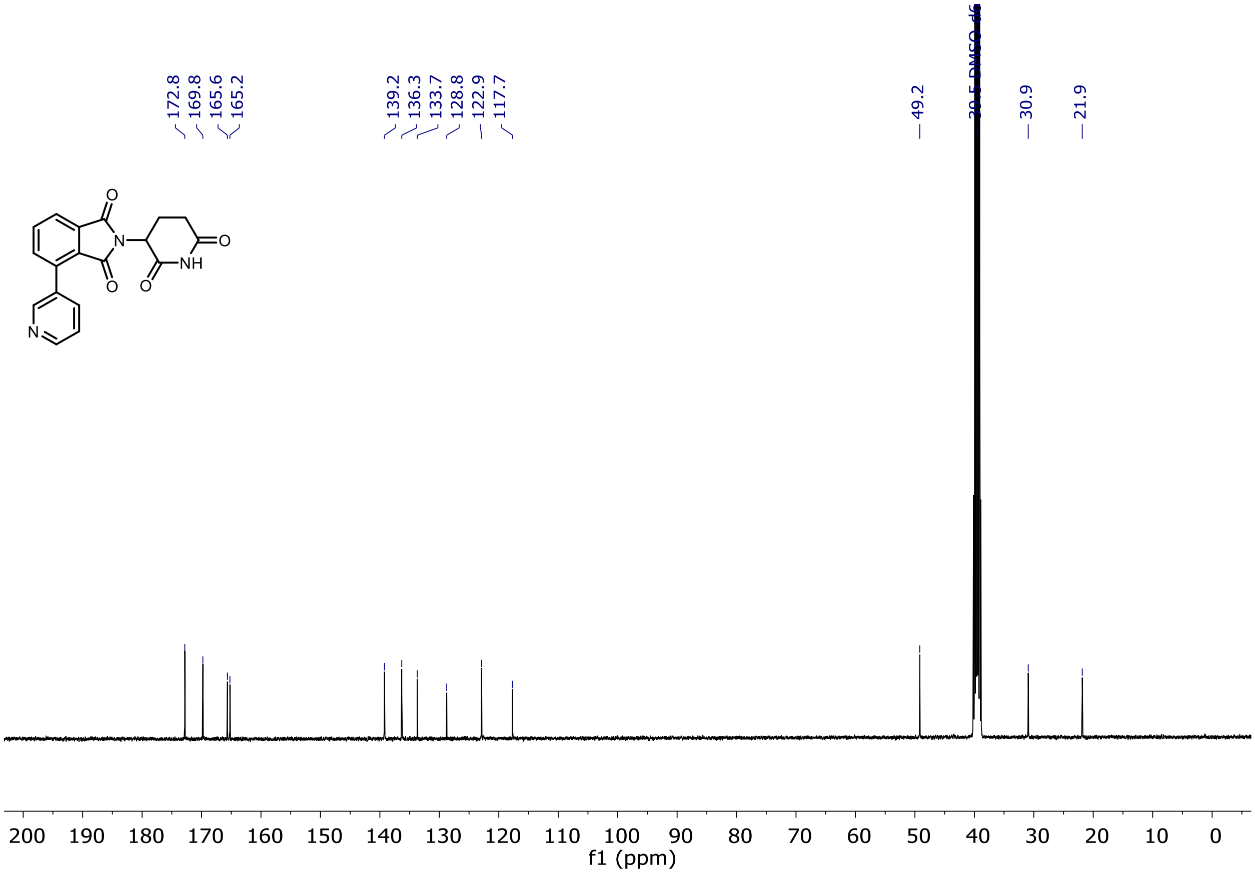

$^{1}H$ and $^{13}C$ NMR of Entry **52**

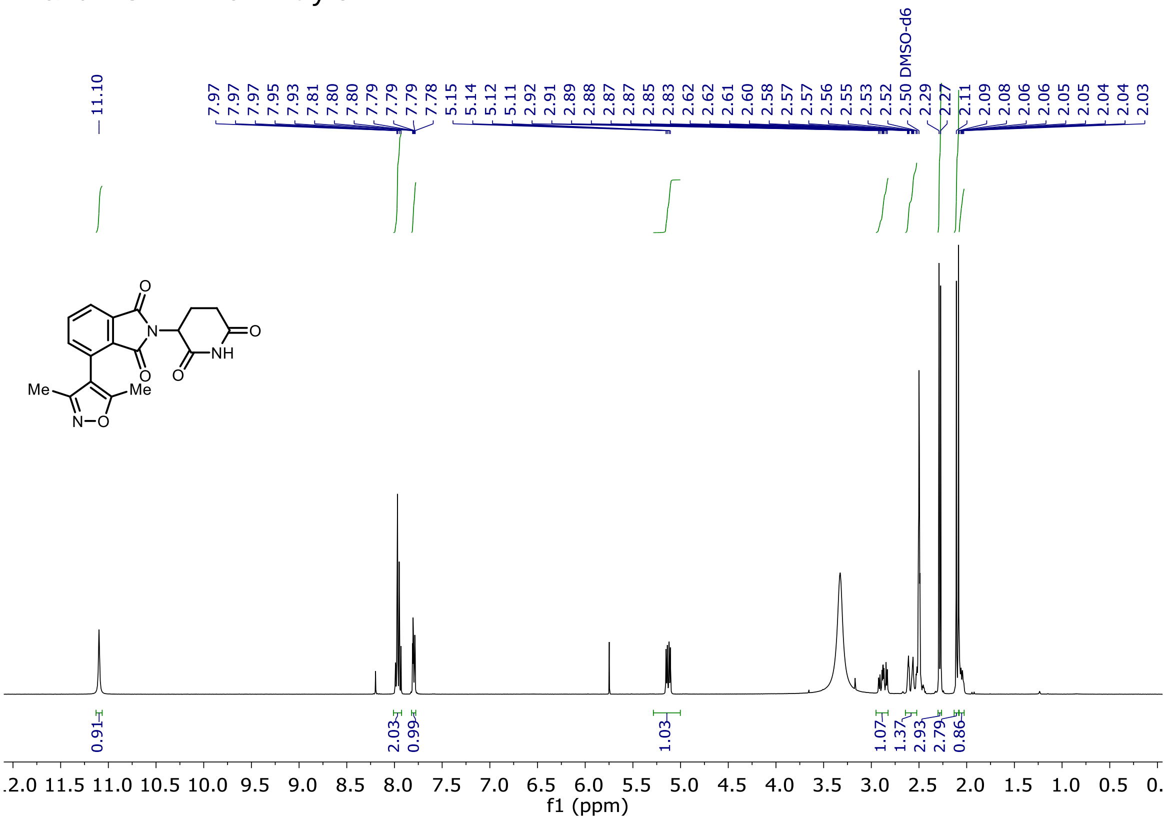

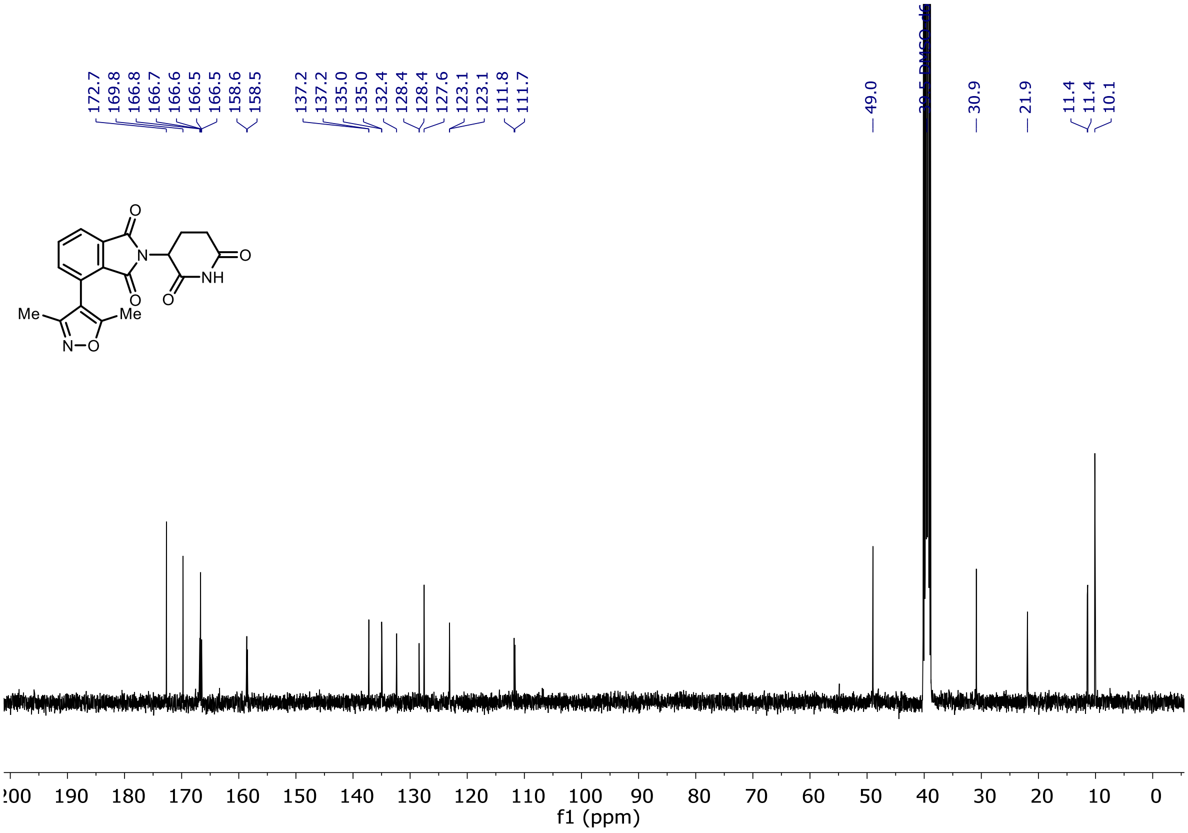

$^{1}H$ and $^{13}C$ NMR of Entry **53**

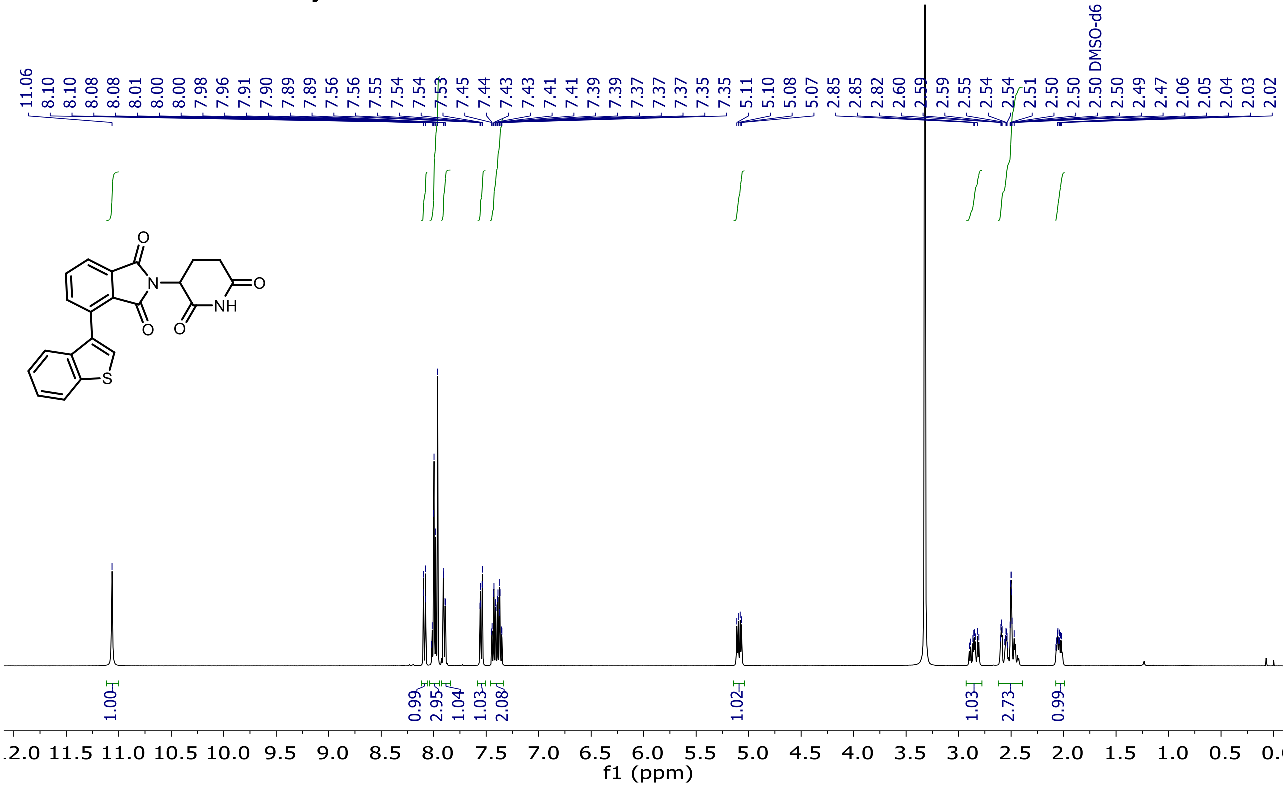

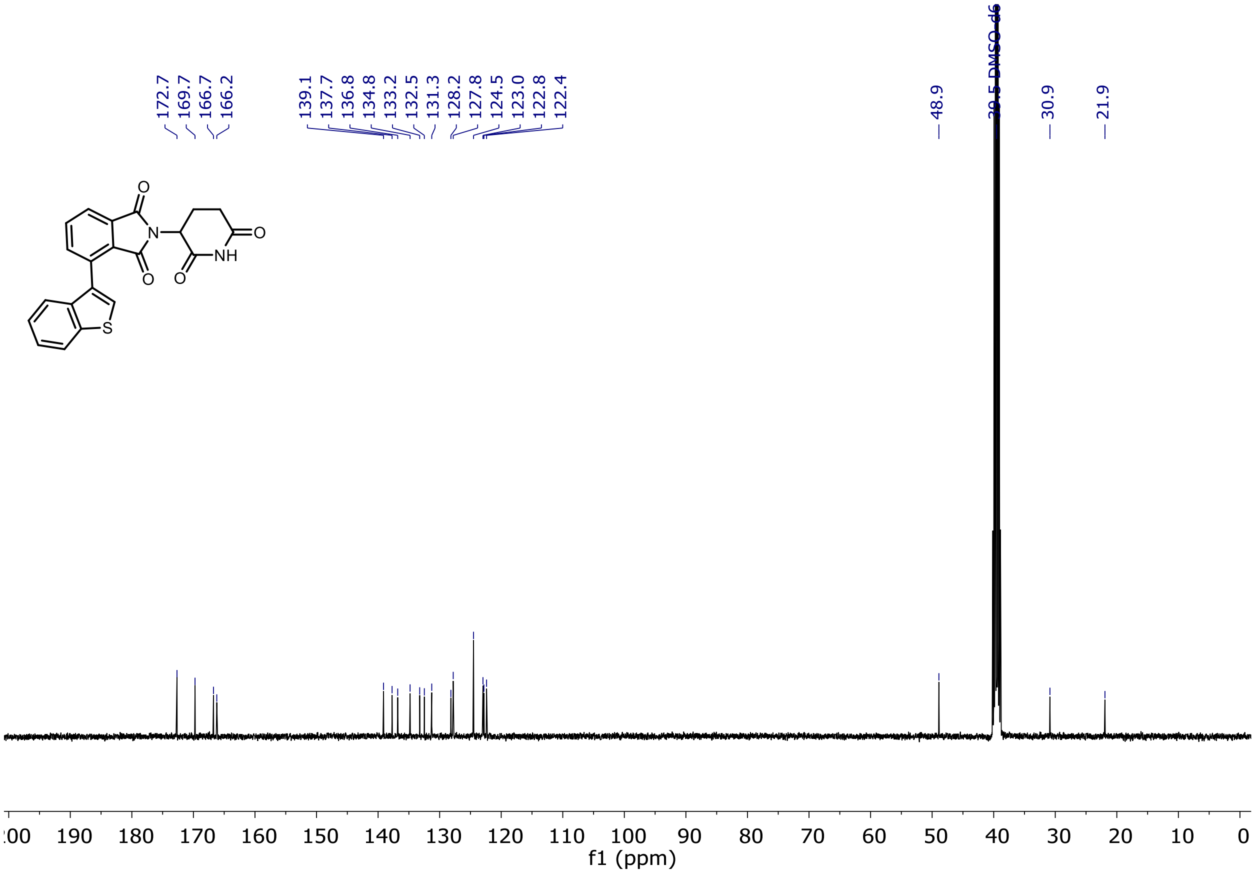

$^{1}H$ and $^{13}C$ NMR of Entry **54**

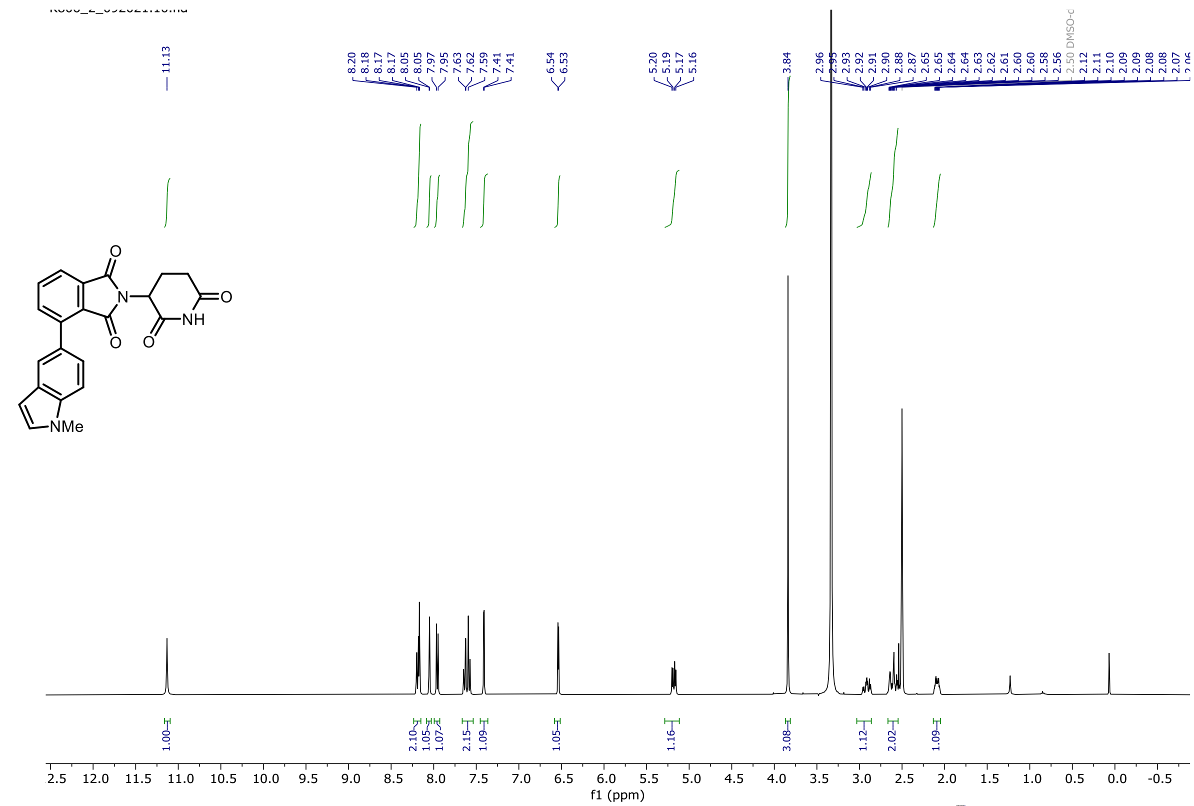

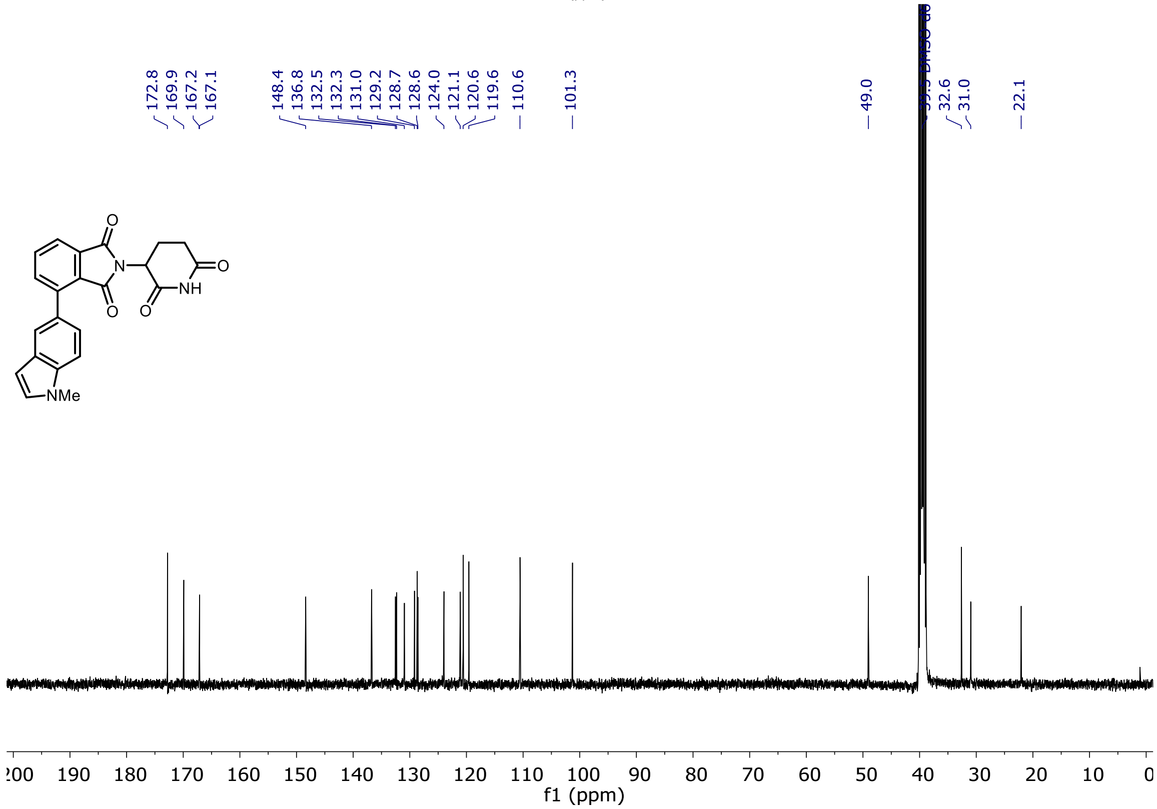

[1]H and [13]C NMR of Entry **55**

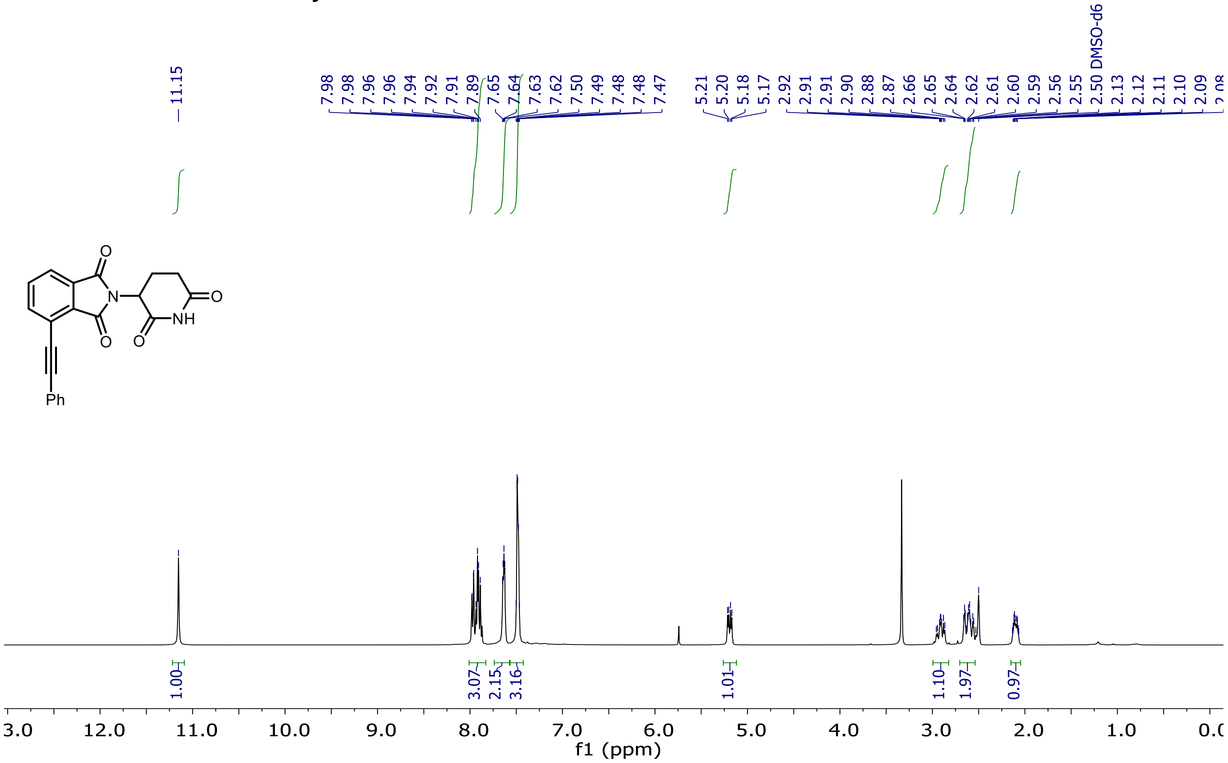

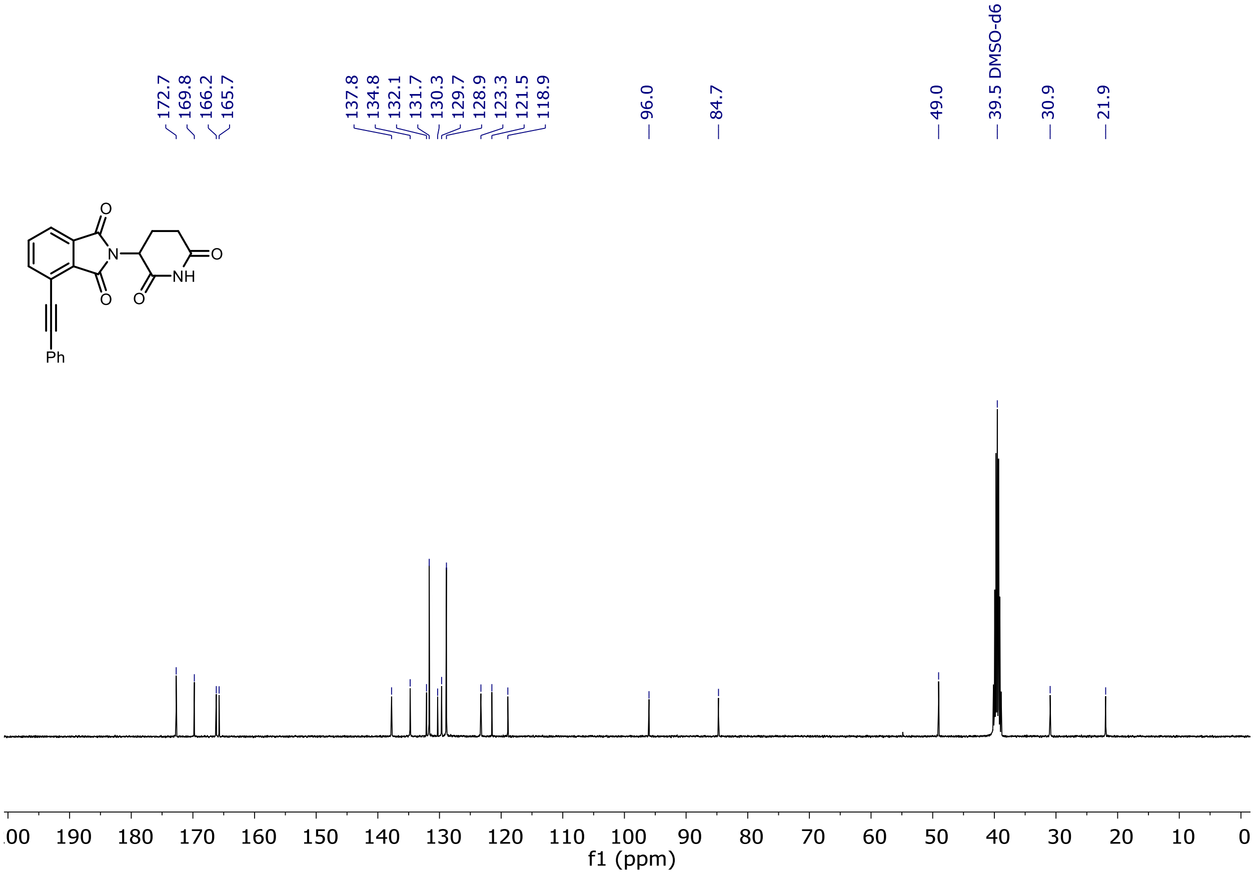

$^{1}H$ and $^{13}C$ NMR of Entry **57**

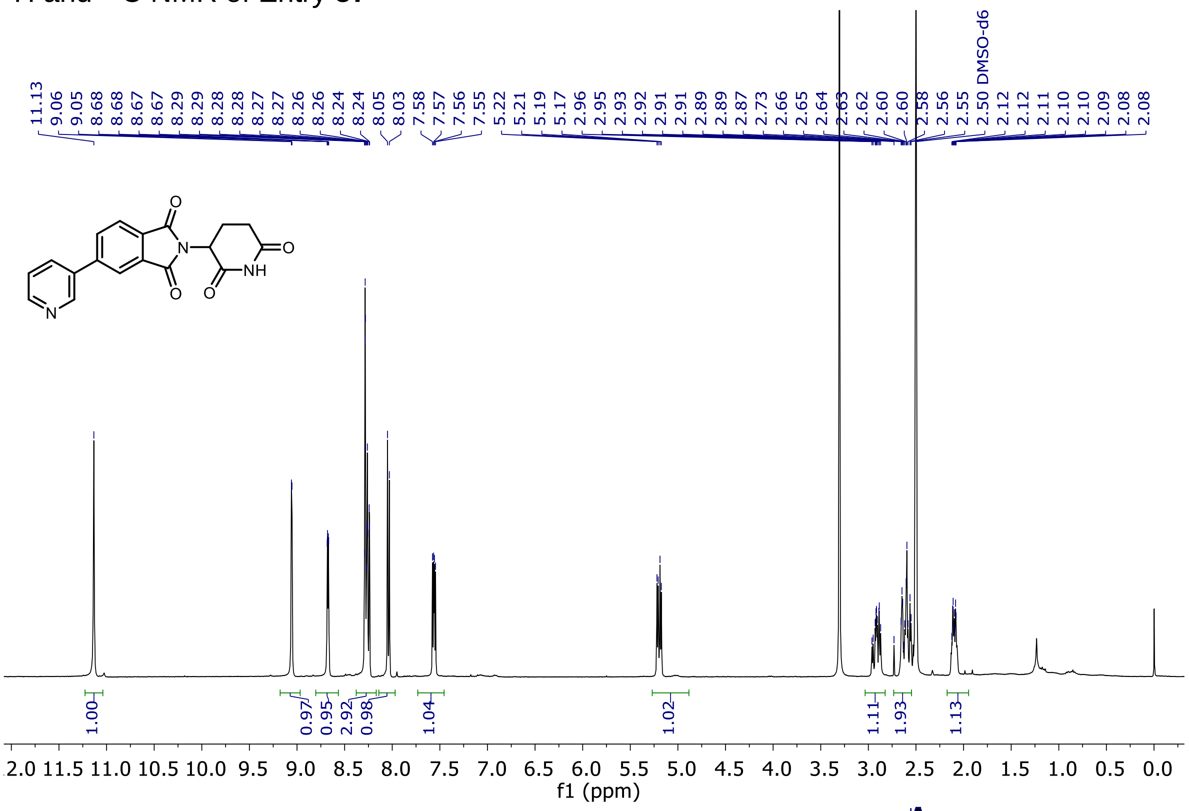

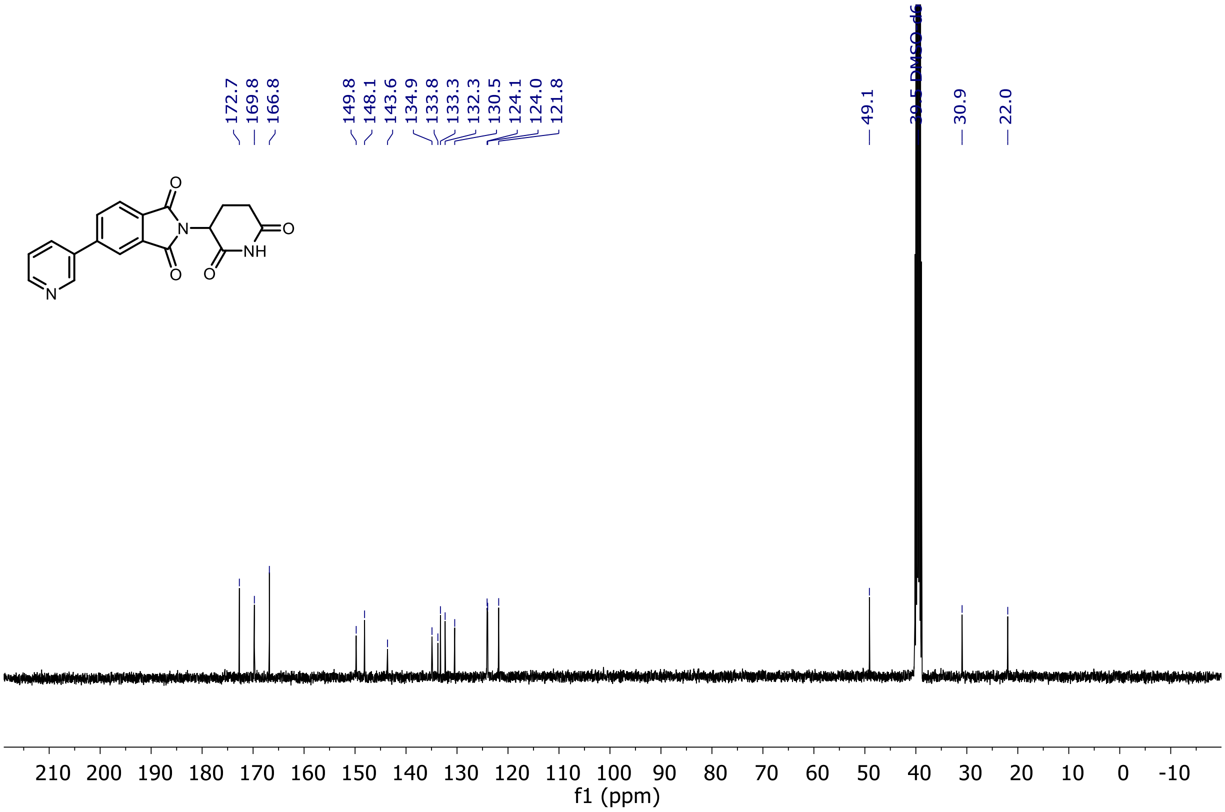

$^{1}H$ and $^{13}C$ NMR of Entry **58**

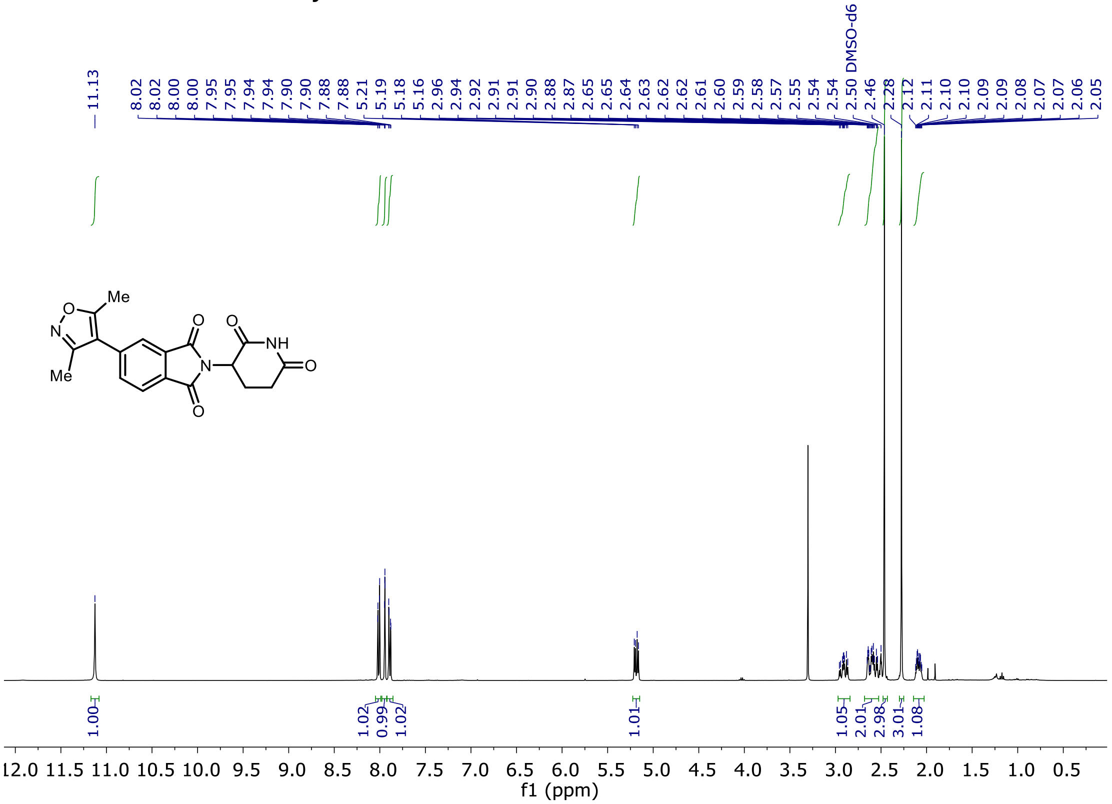

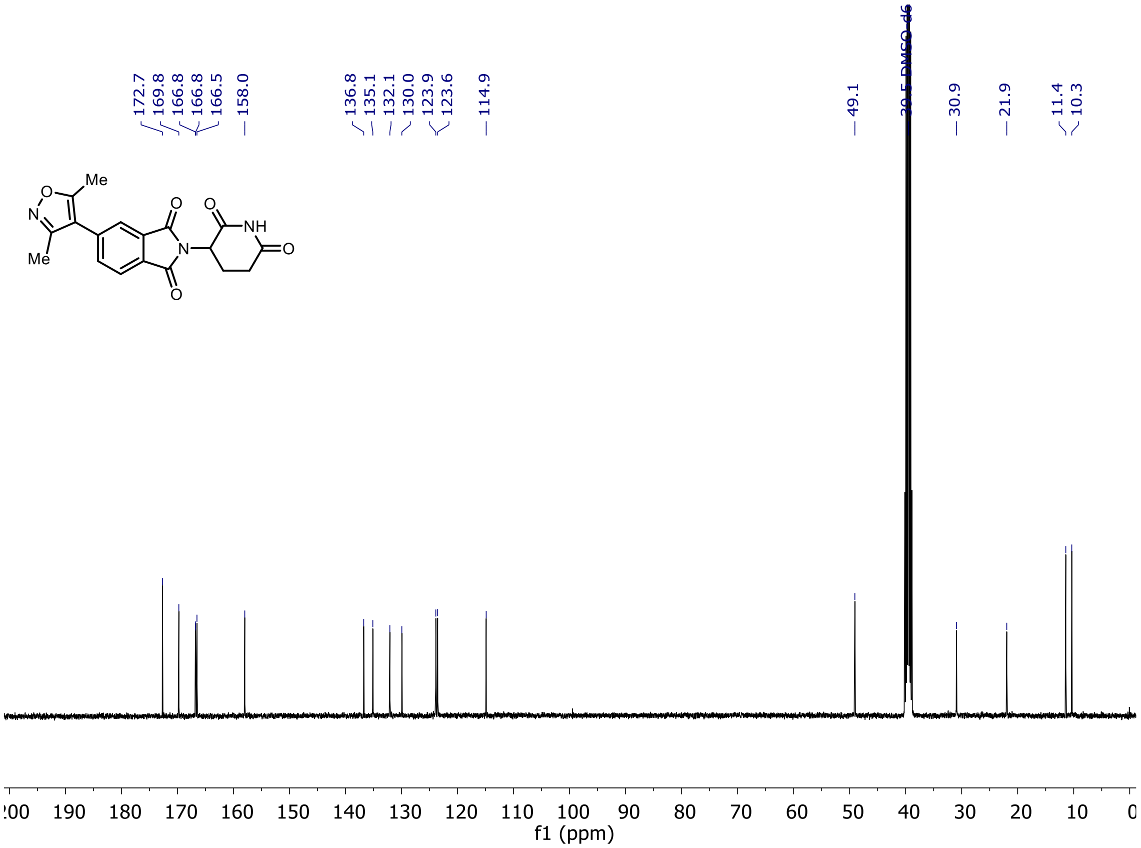

[1]H and [13]C NMR of Entry **59**

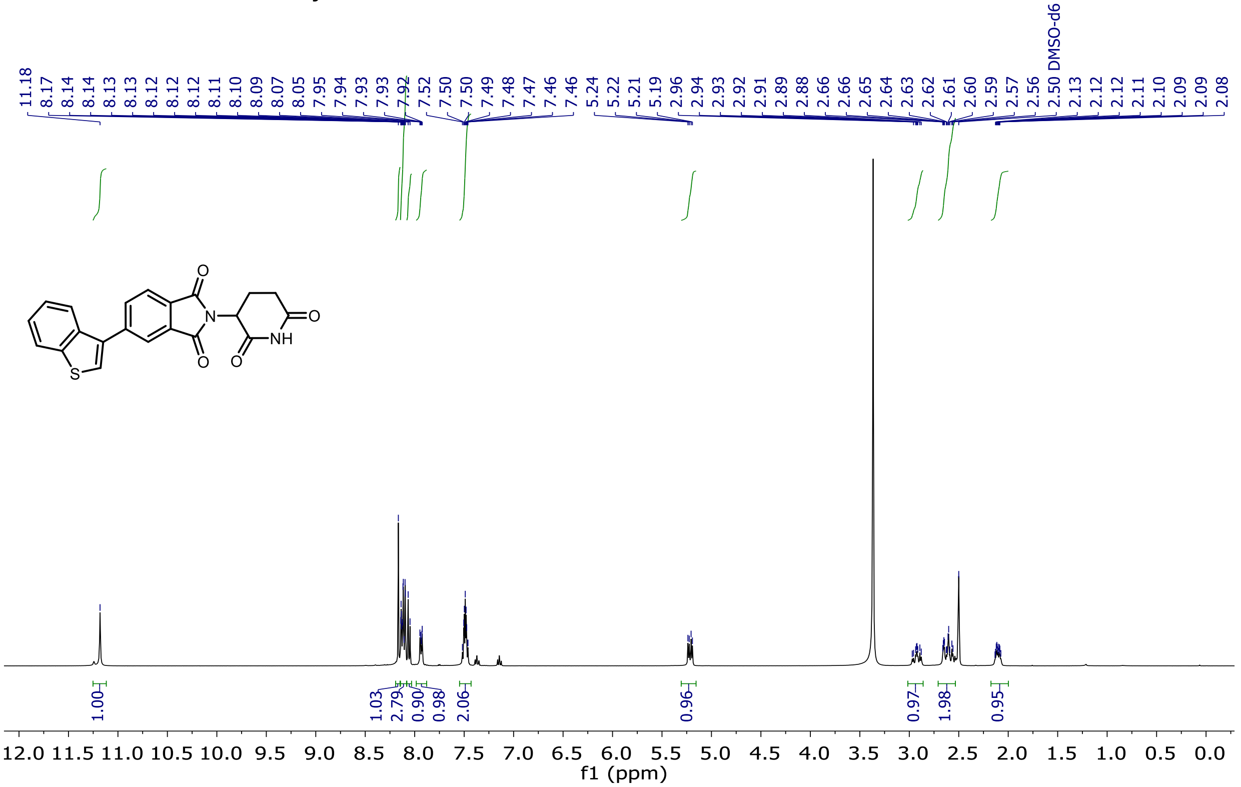

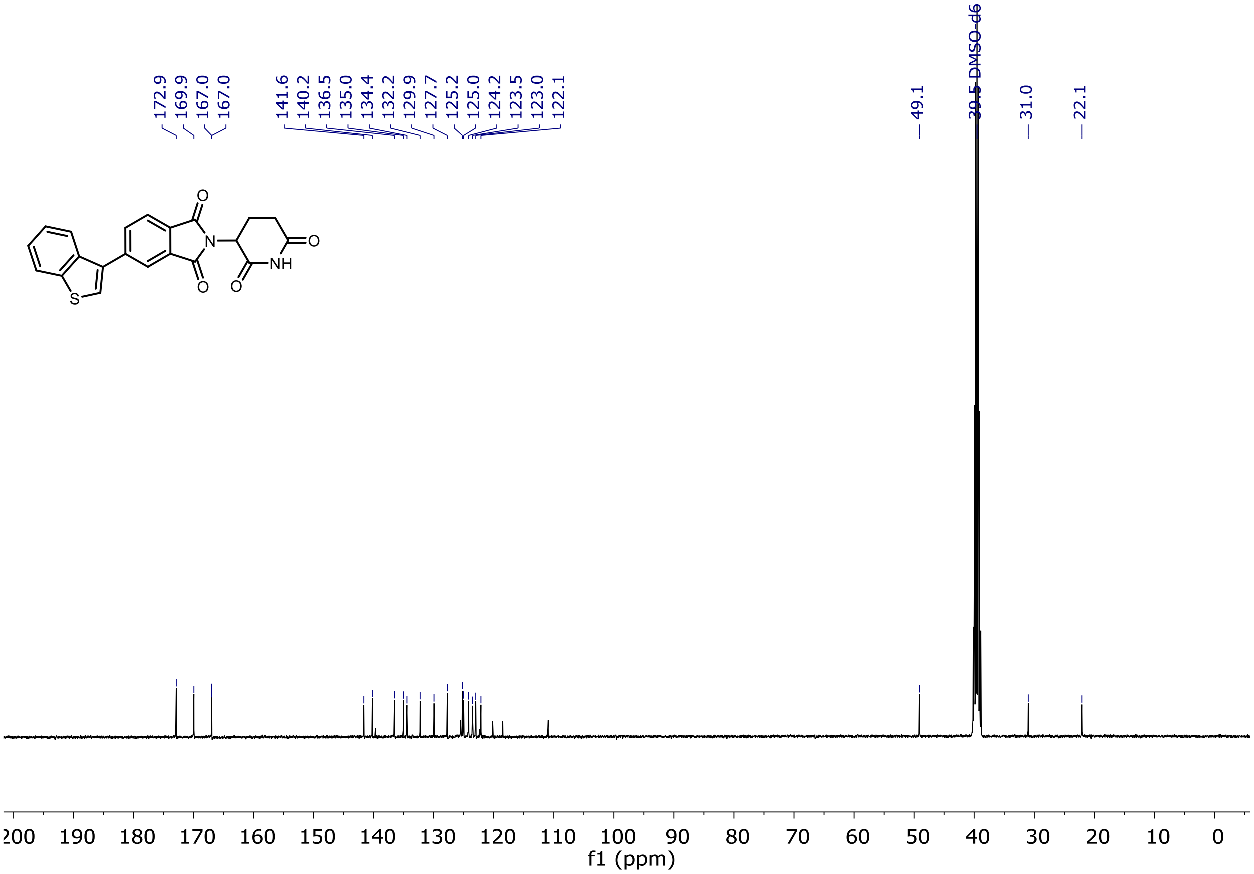

$^1H$ and $^{13}C$ NMR of Entry **60**

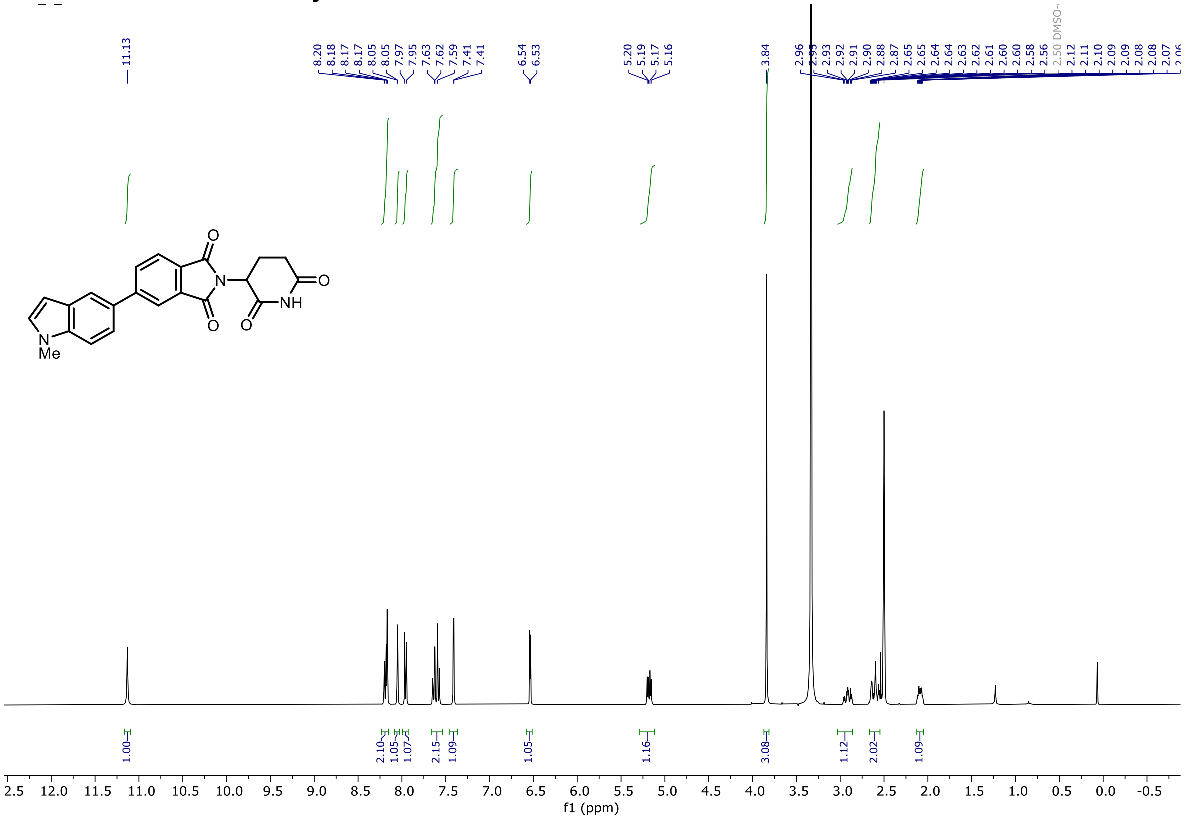

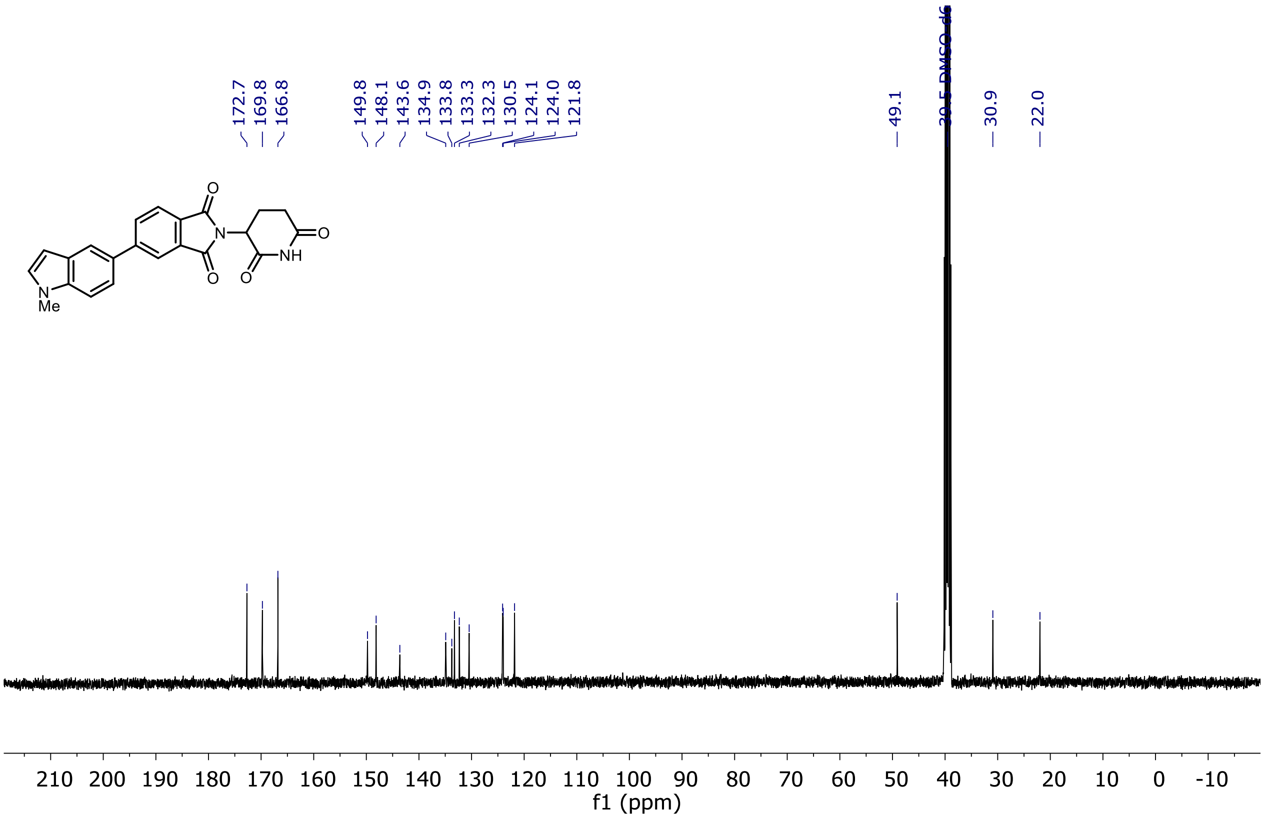

[1]H and [13]C NMR of Entry **61**

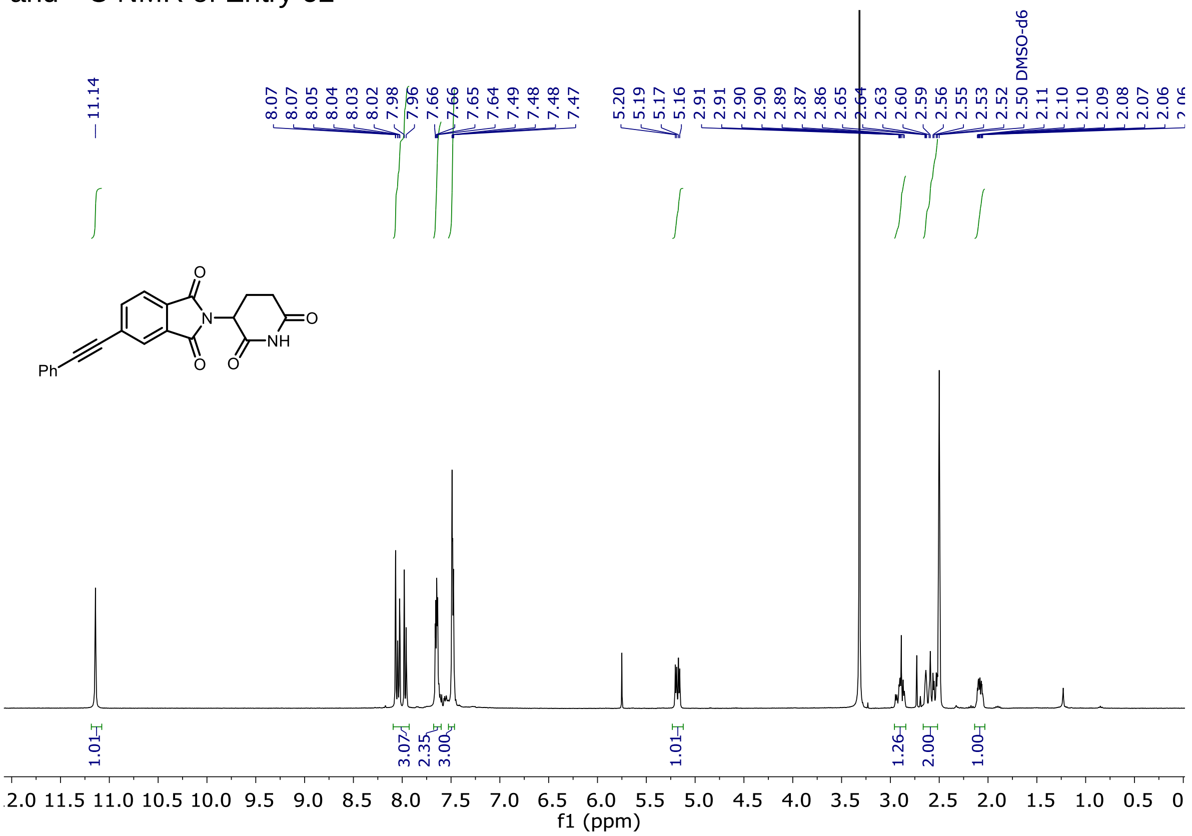

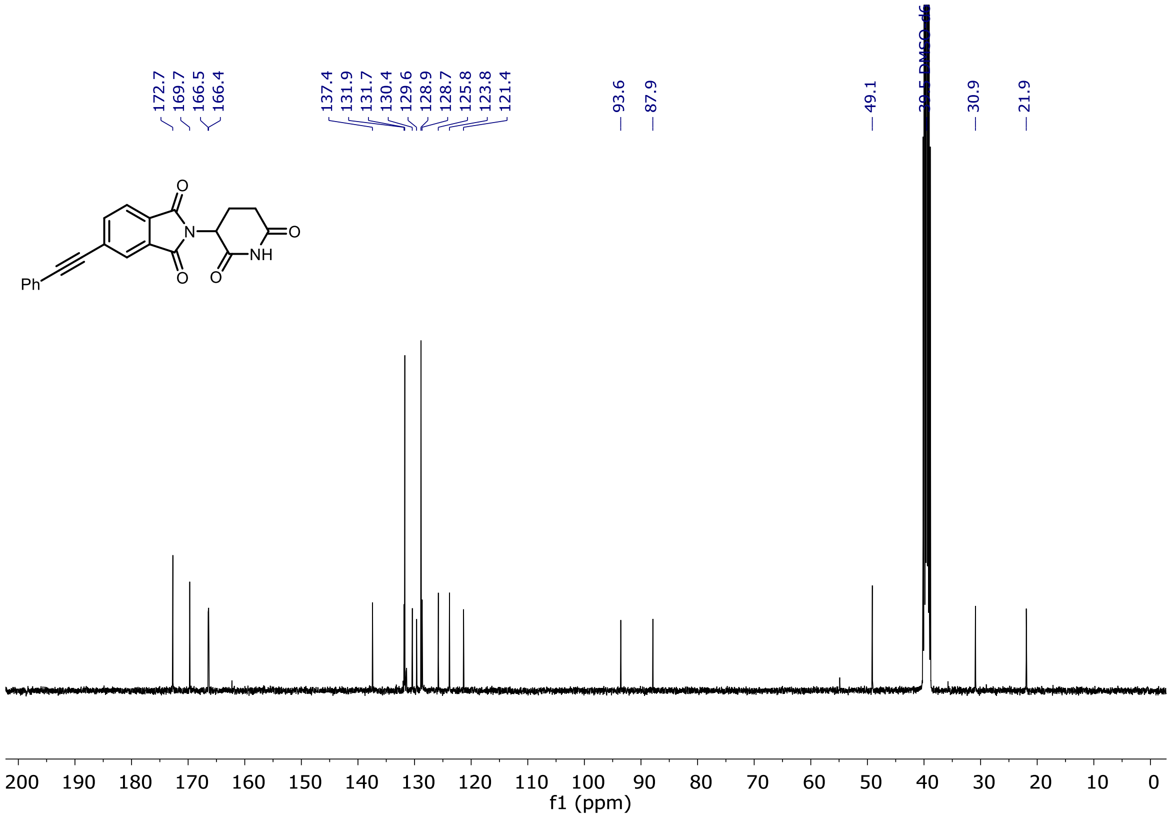

$^{1}H$ and $^{13}C$ NMR of Entry **62**

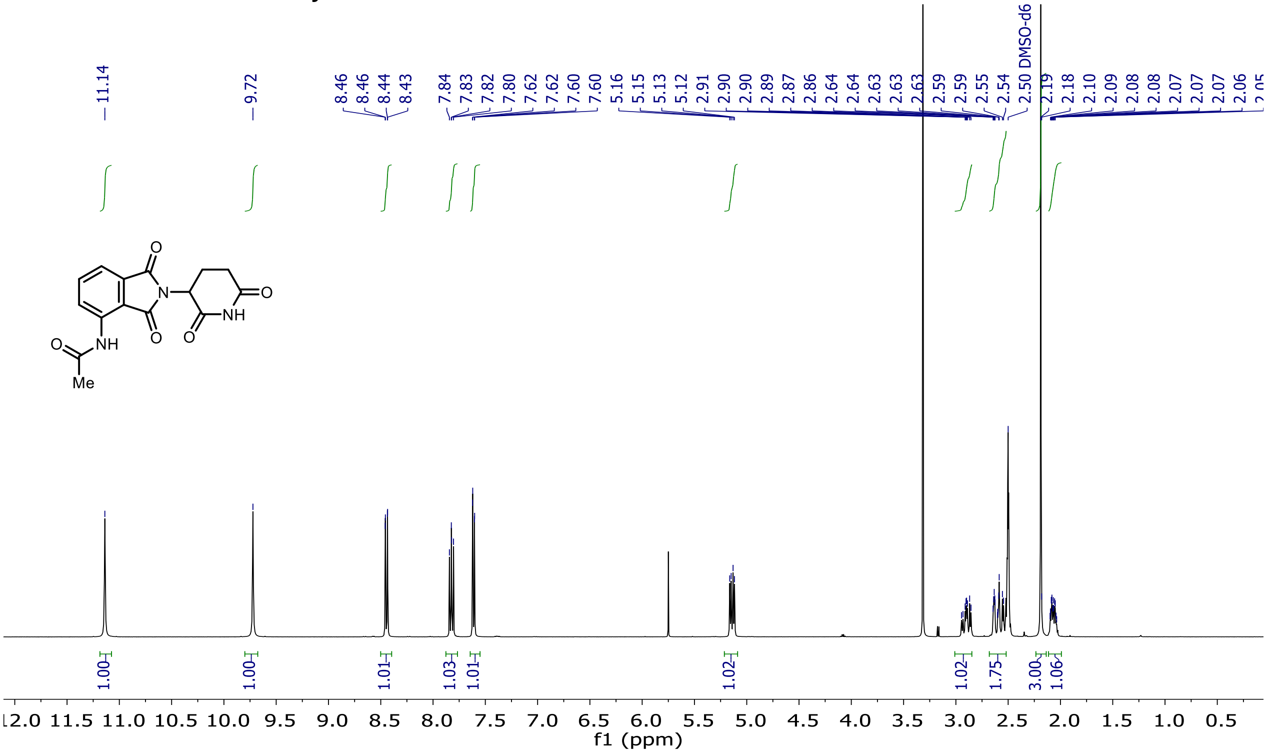

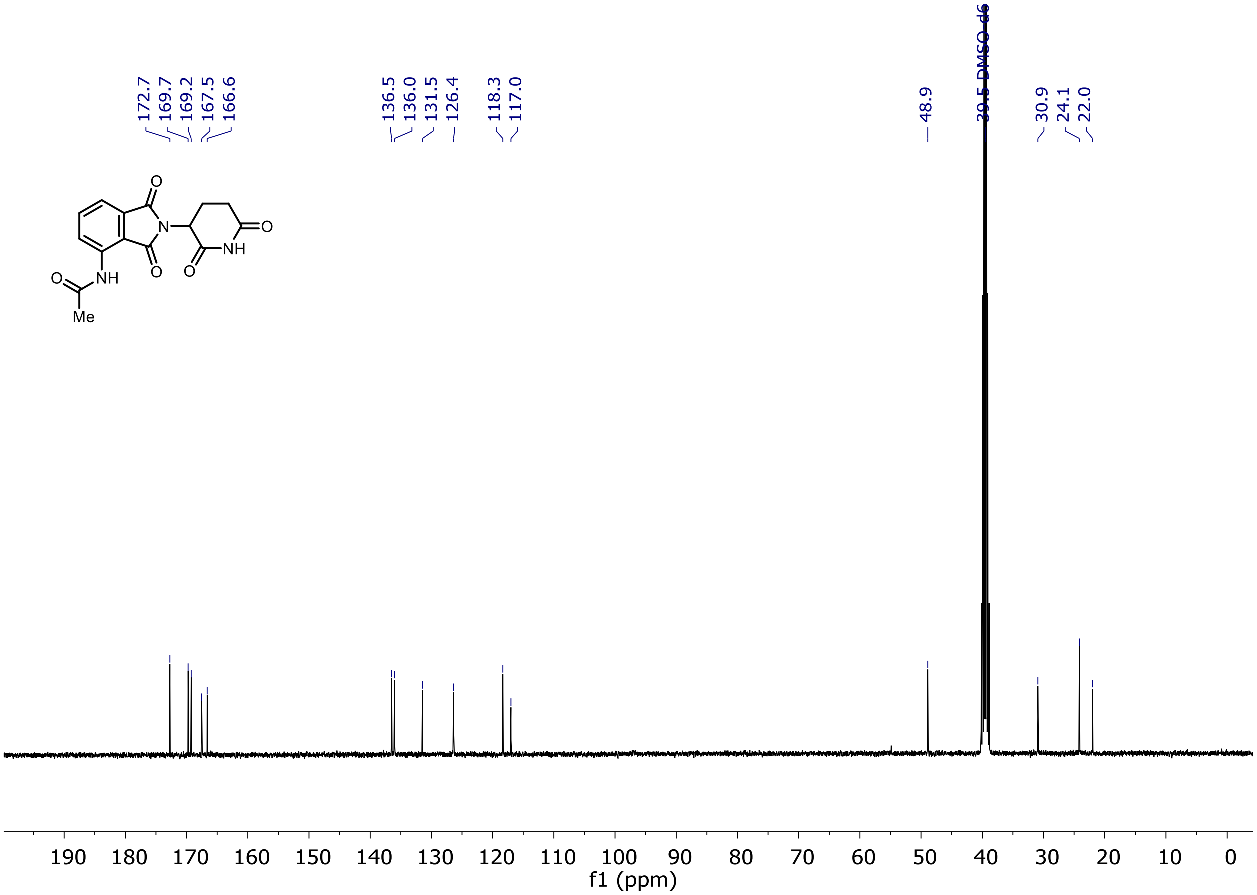

$^1$H and $^{13}$C NMR of Entry **63**

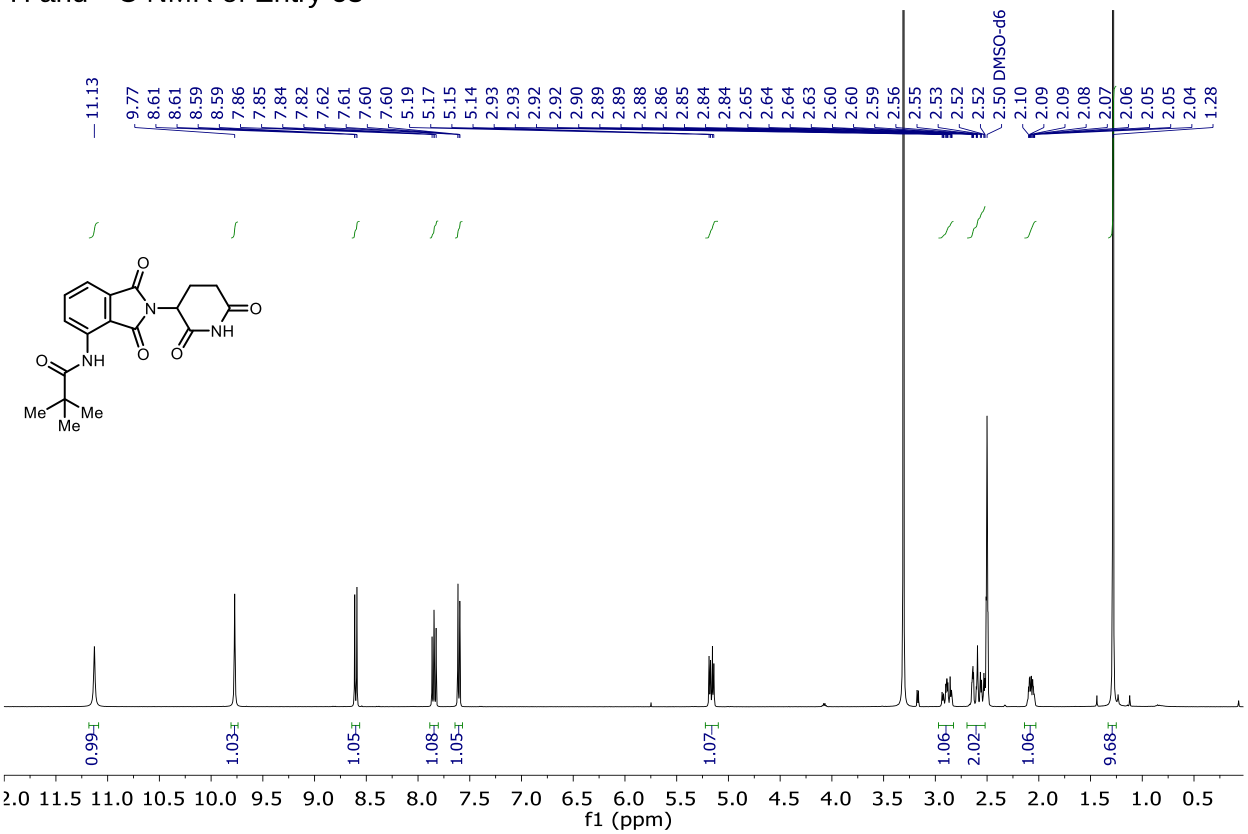

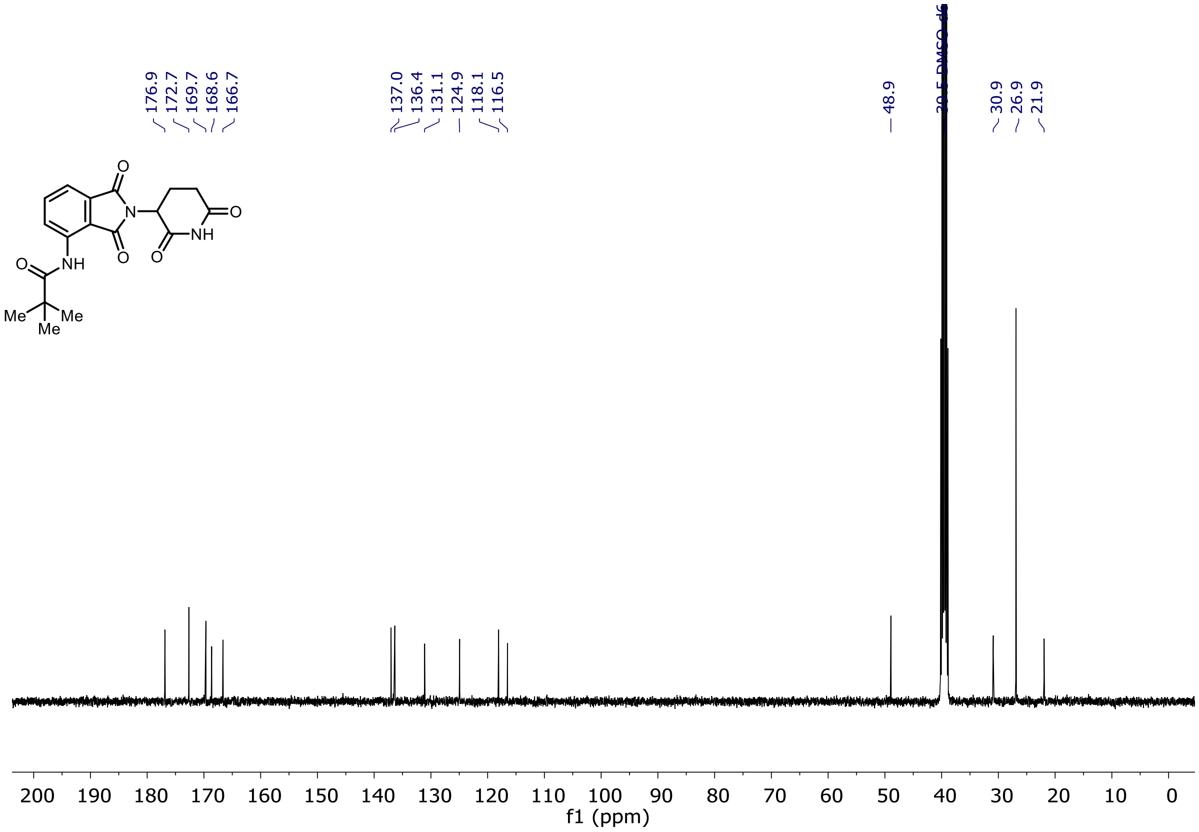

<sup></sup>$^{1}$H and $^{13}$C NMR of Entry **64**

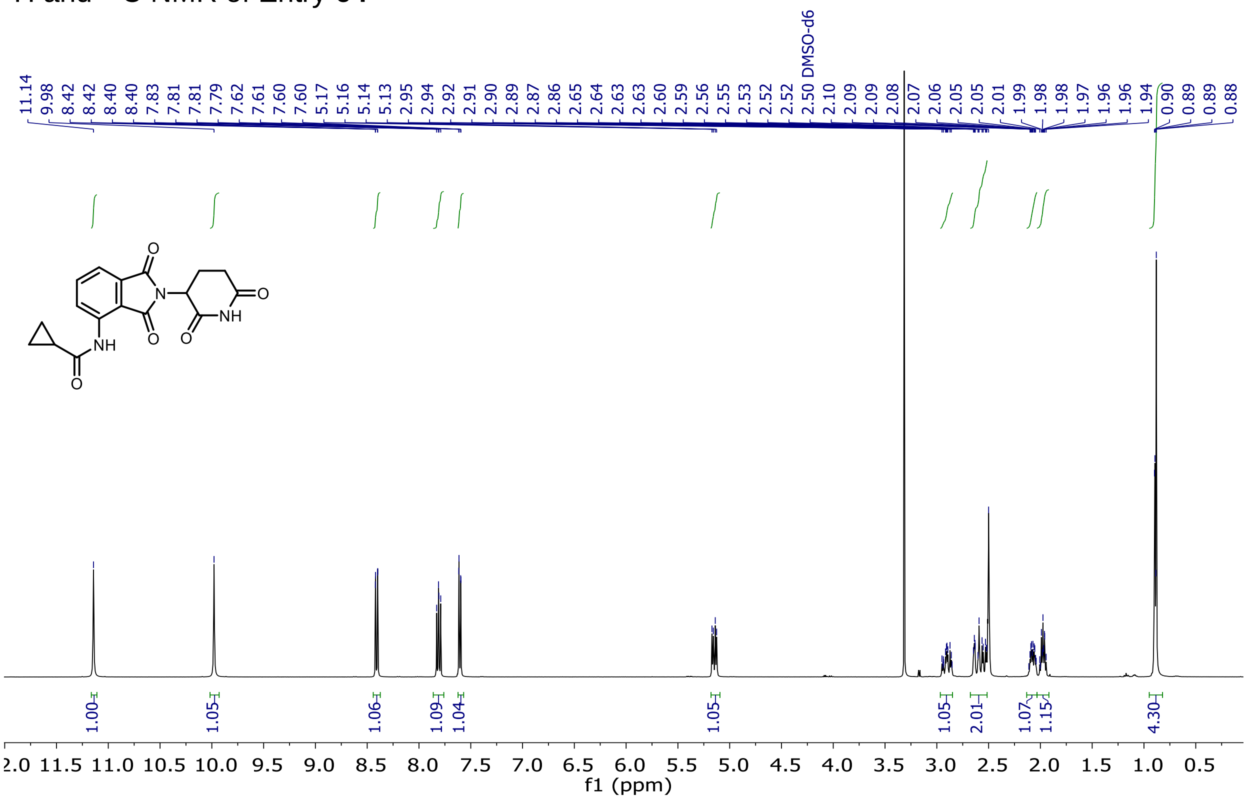

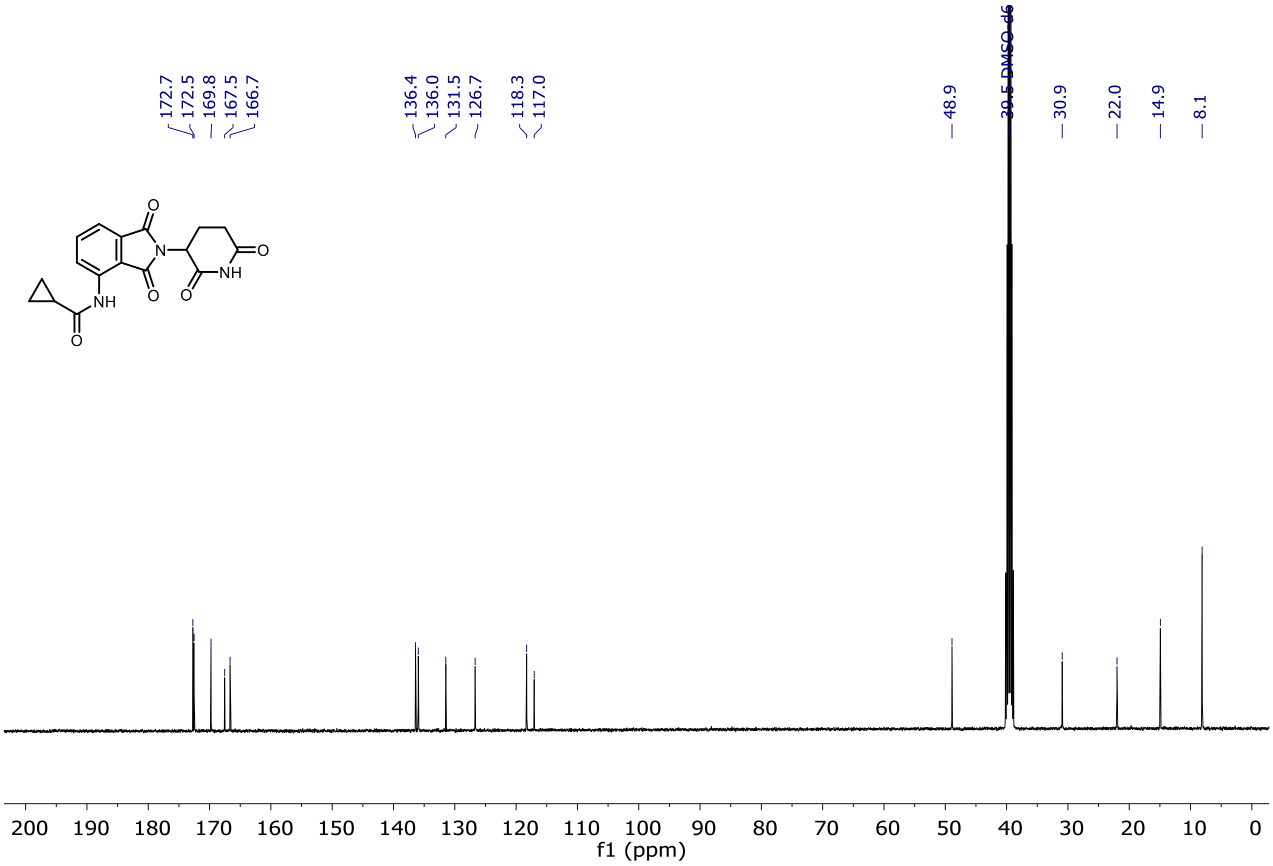

$^{1}H$ and $^{13}C$ NMR of Entry **65**

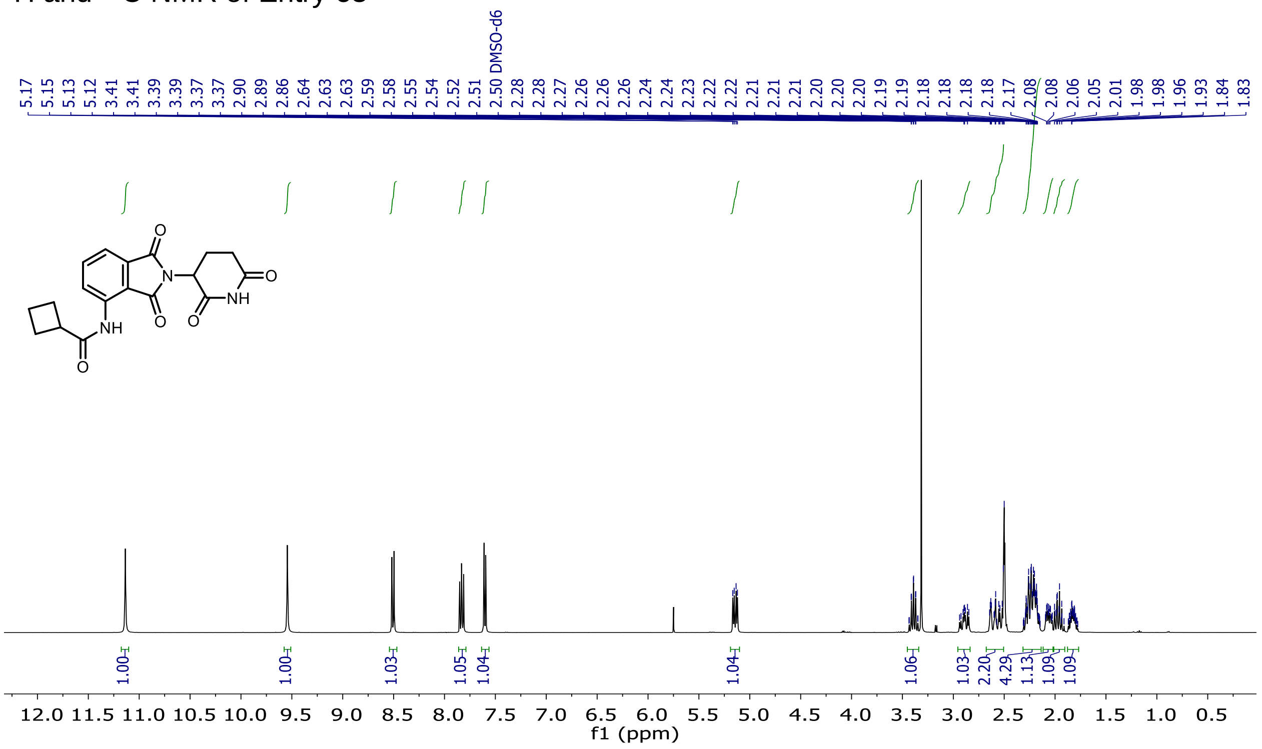

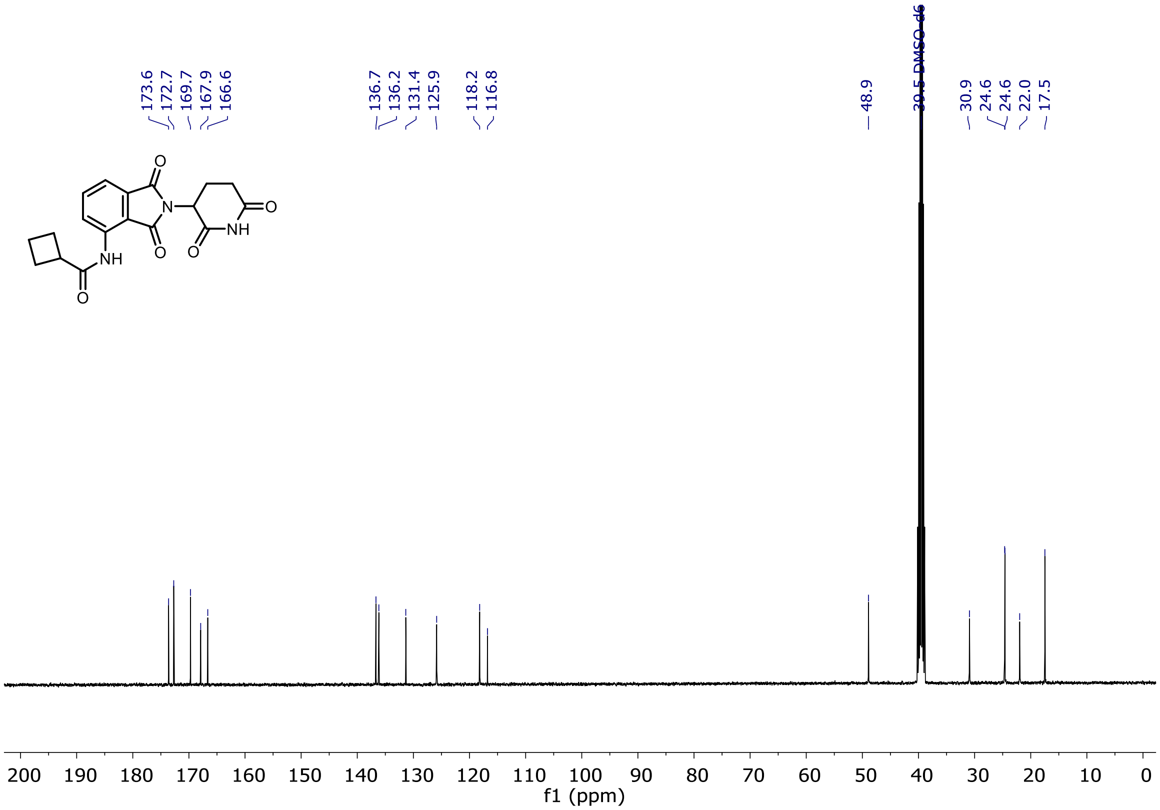

$^{1}H$ and $^{13}C$ NMR of Entry **66**

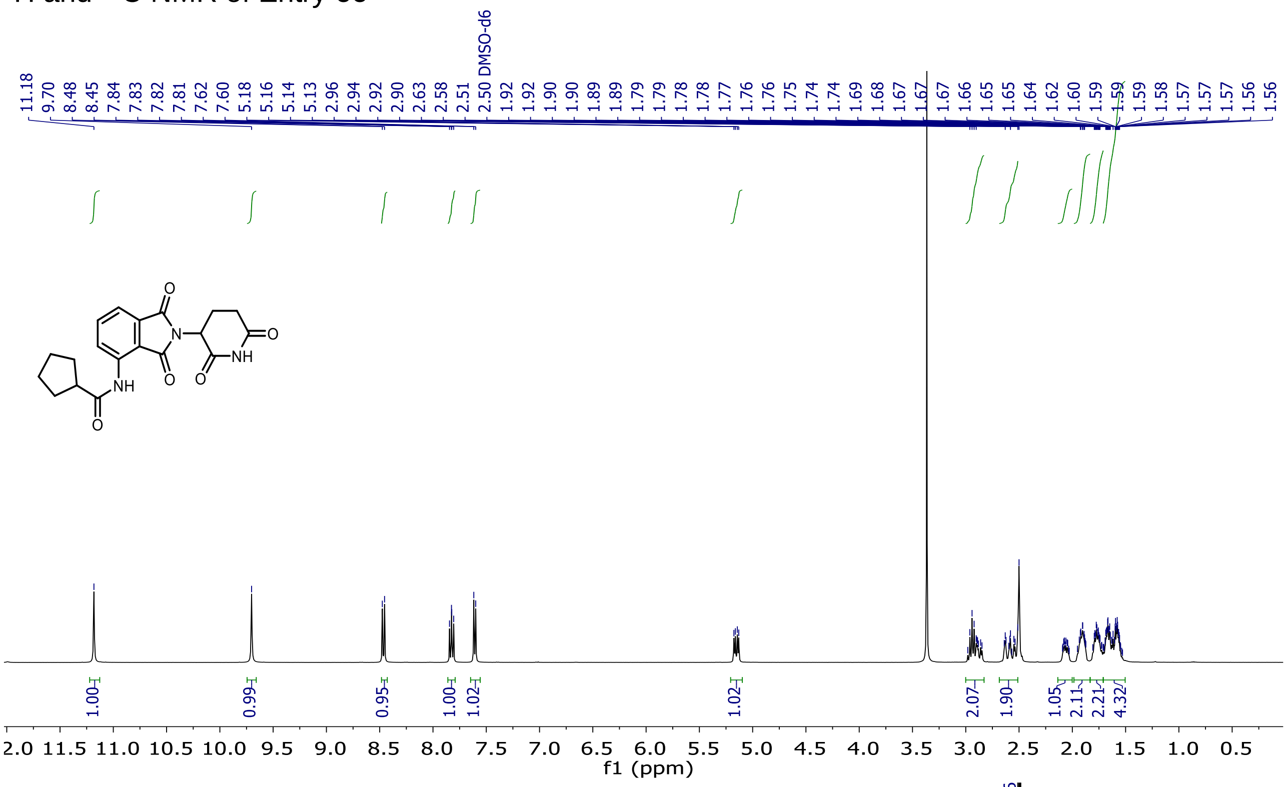

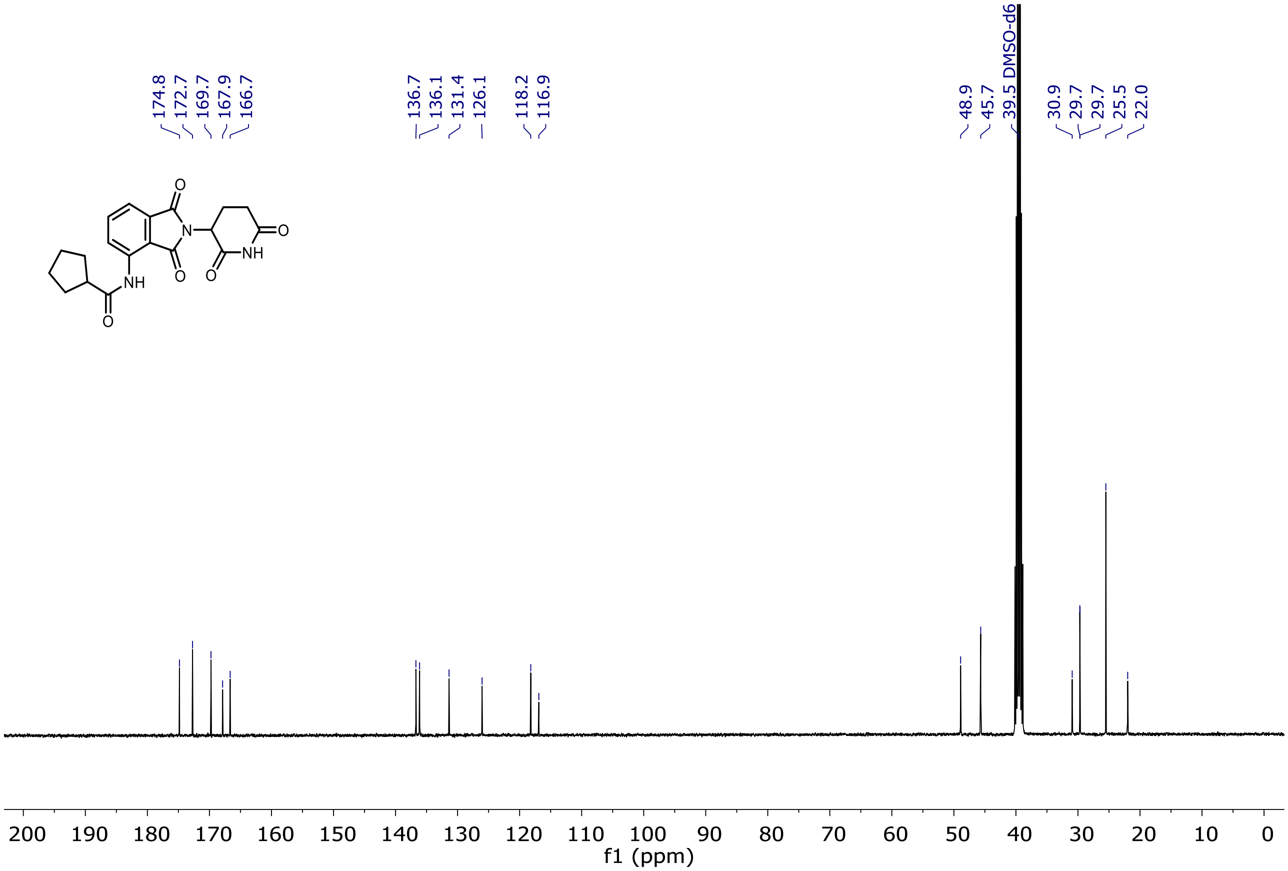

[1]H and [13]C NMR of Entry **67**

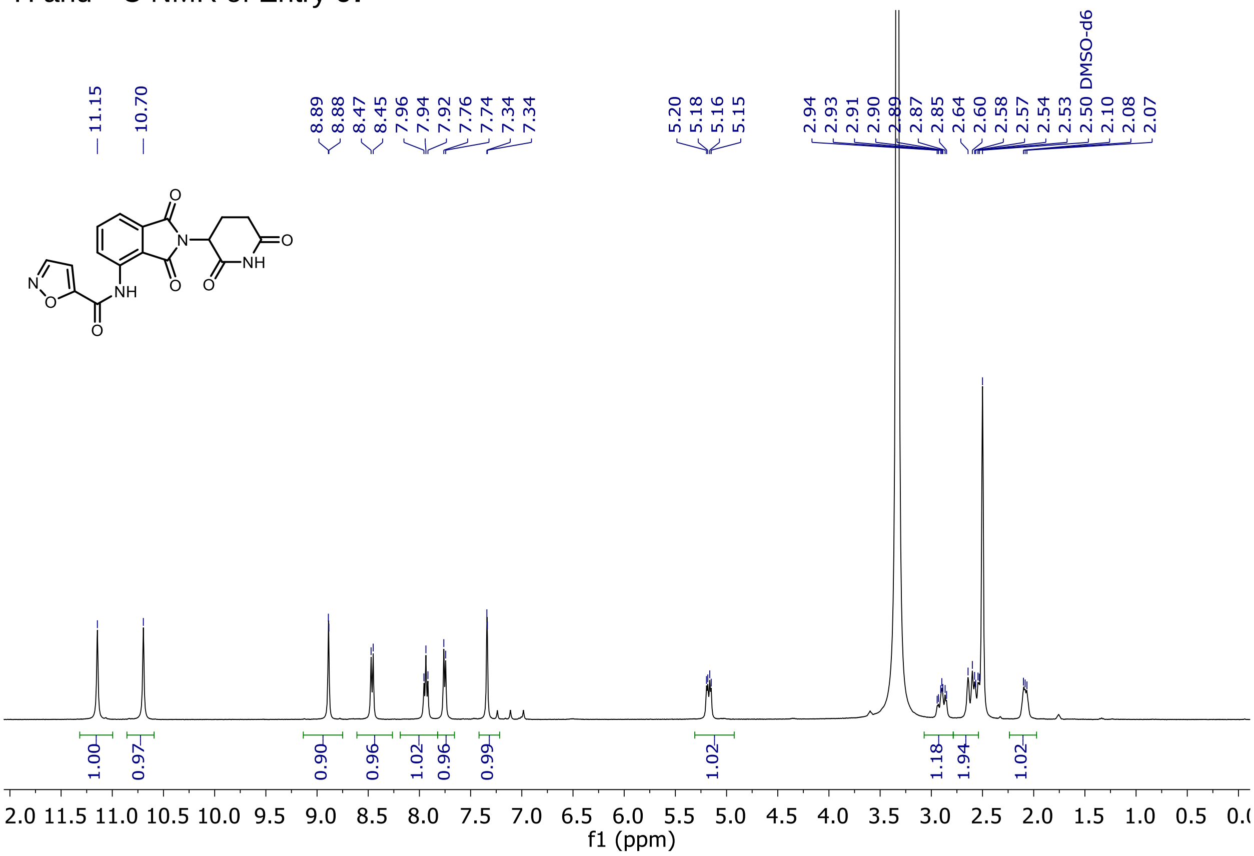

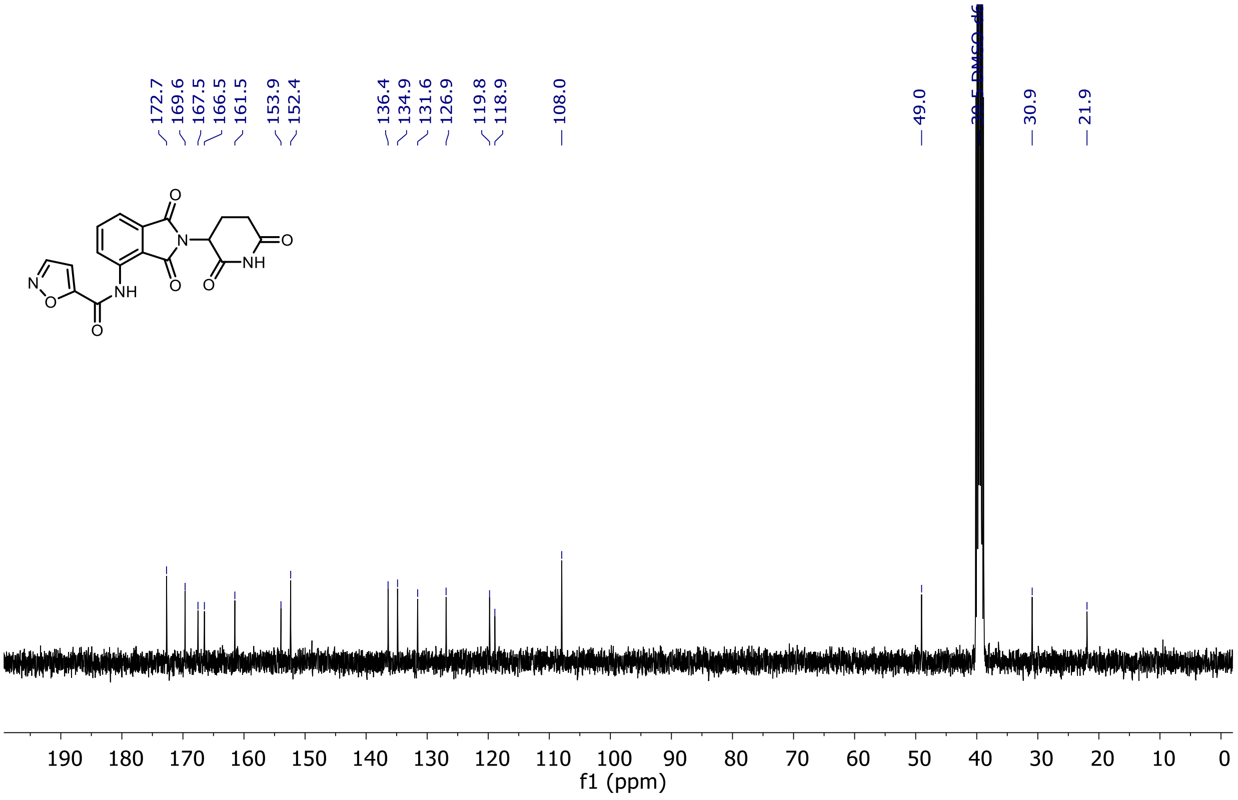

$^{1}$H and $^{13}$C NMR of Entry **68**

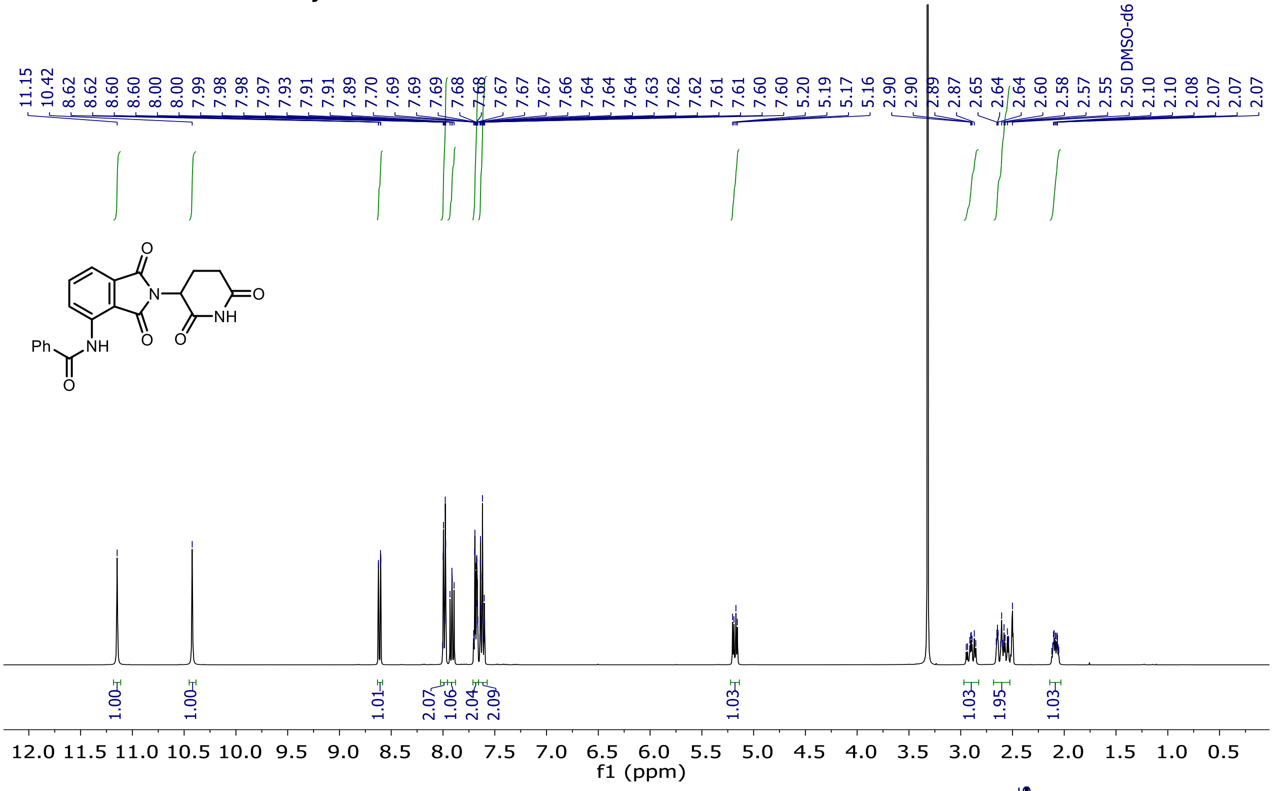

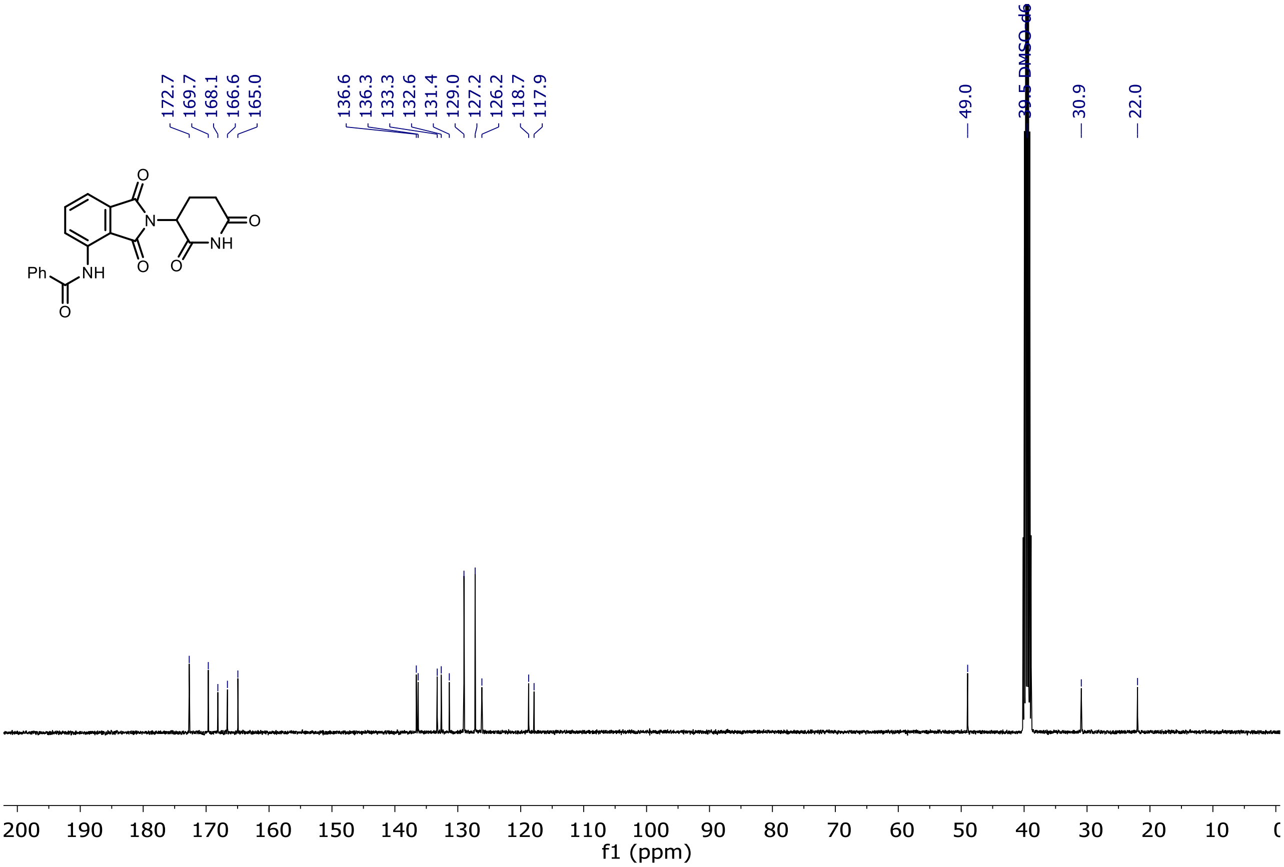

### $^{1}H$ and $^{13}C$ NMR of Entry **69**

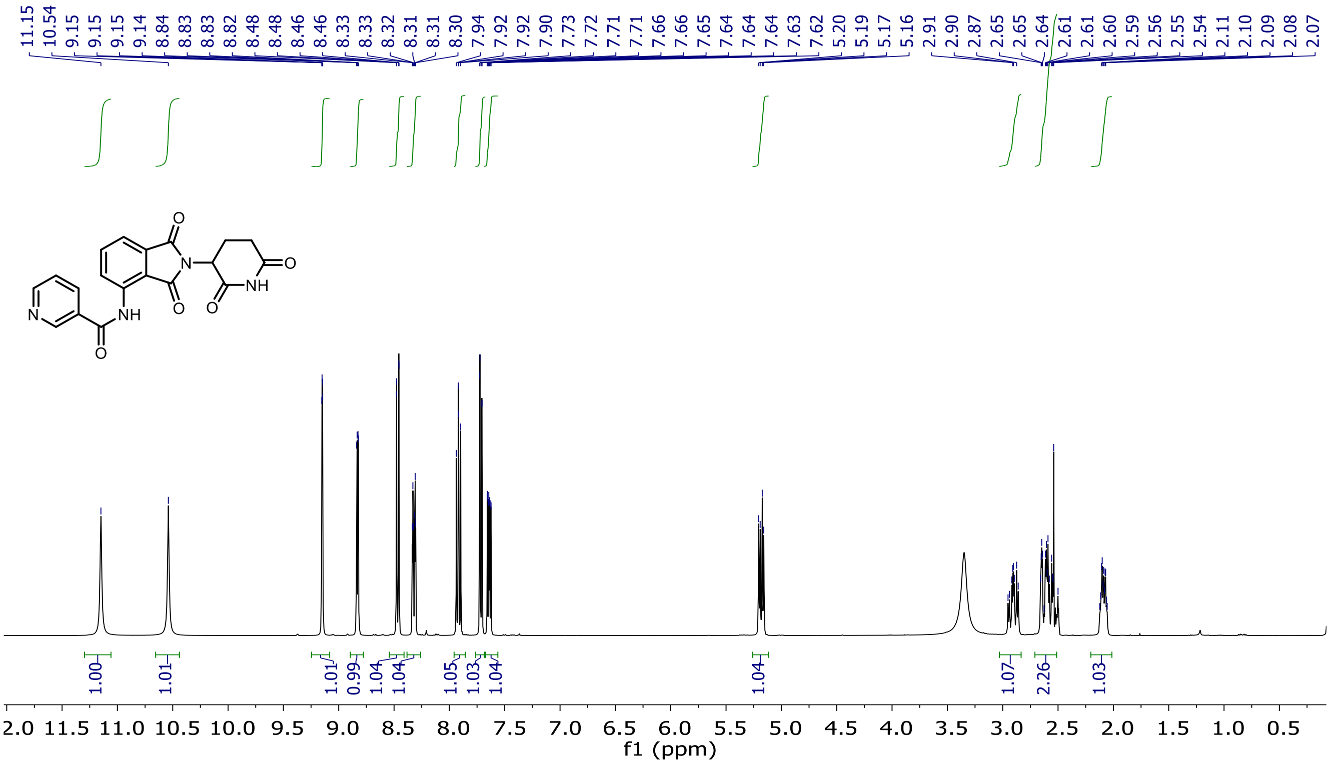

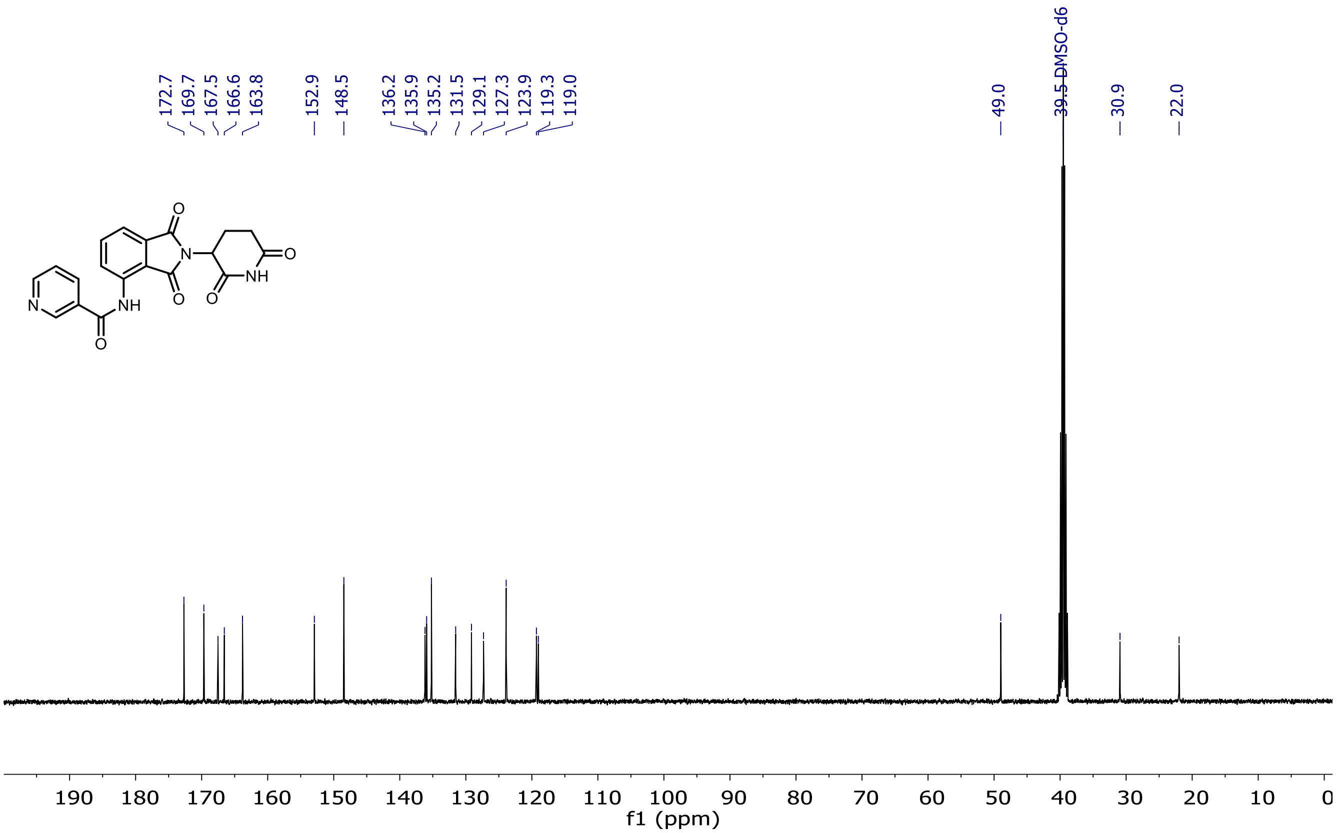

$^{1}H$ and $^{13}C$ NMR of Entry **70**

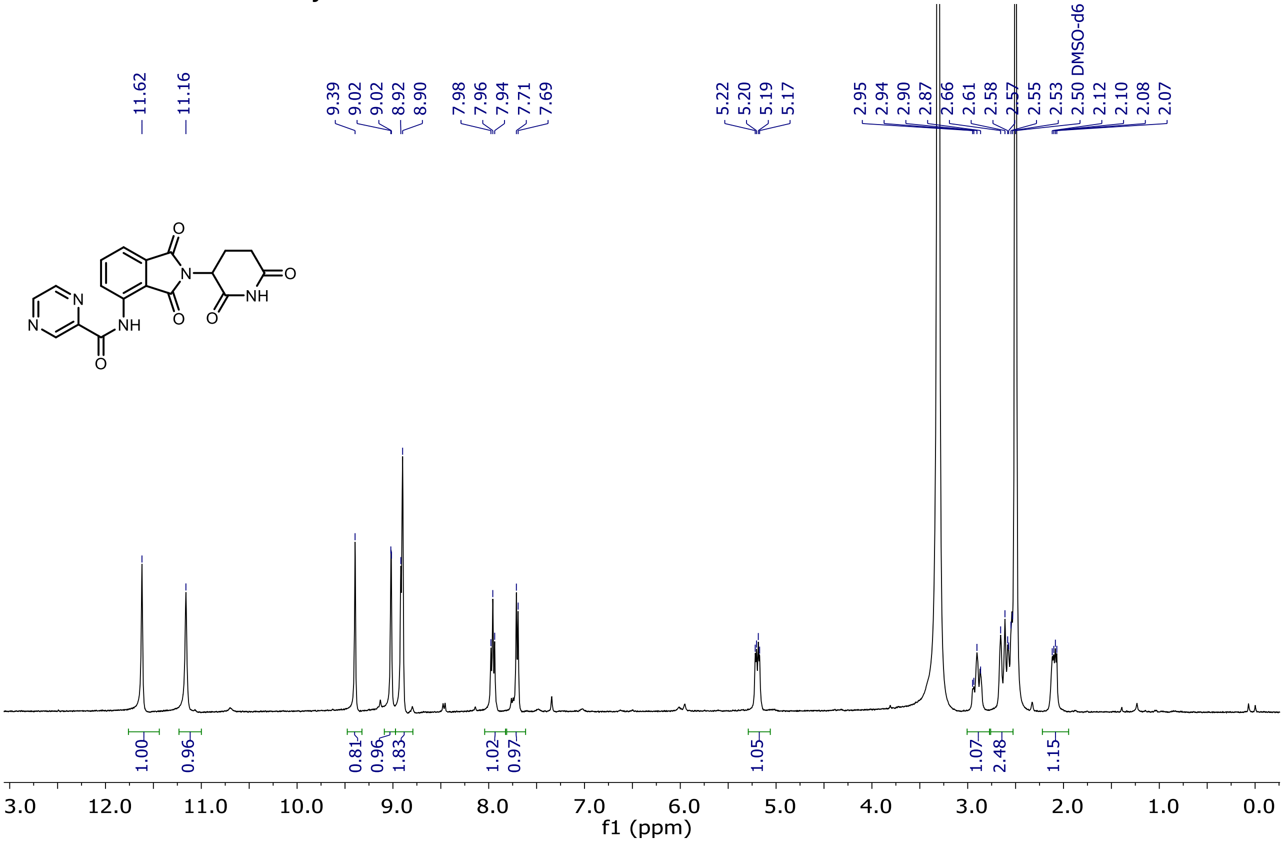

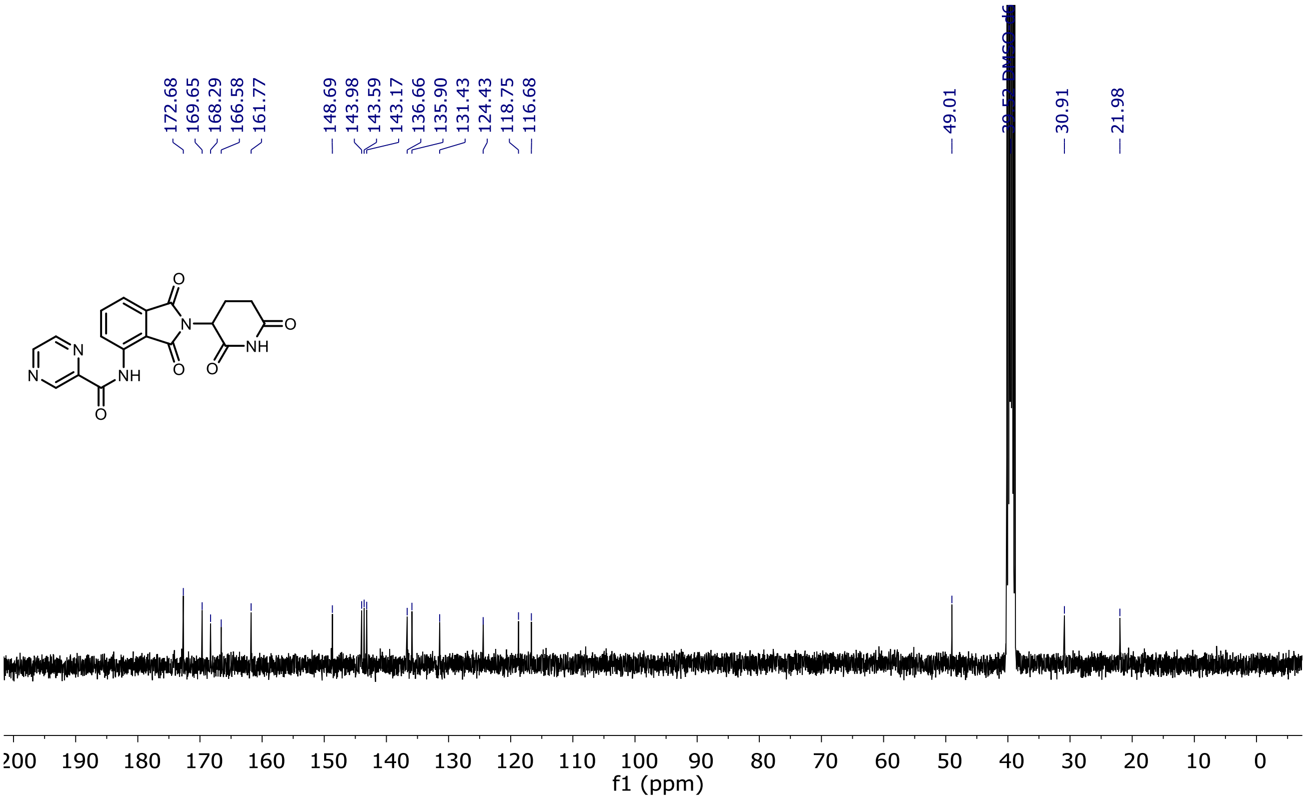

#### $^{1}H$ and $^{13}C$ NMR of Entry **71**

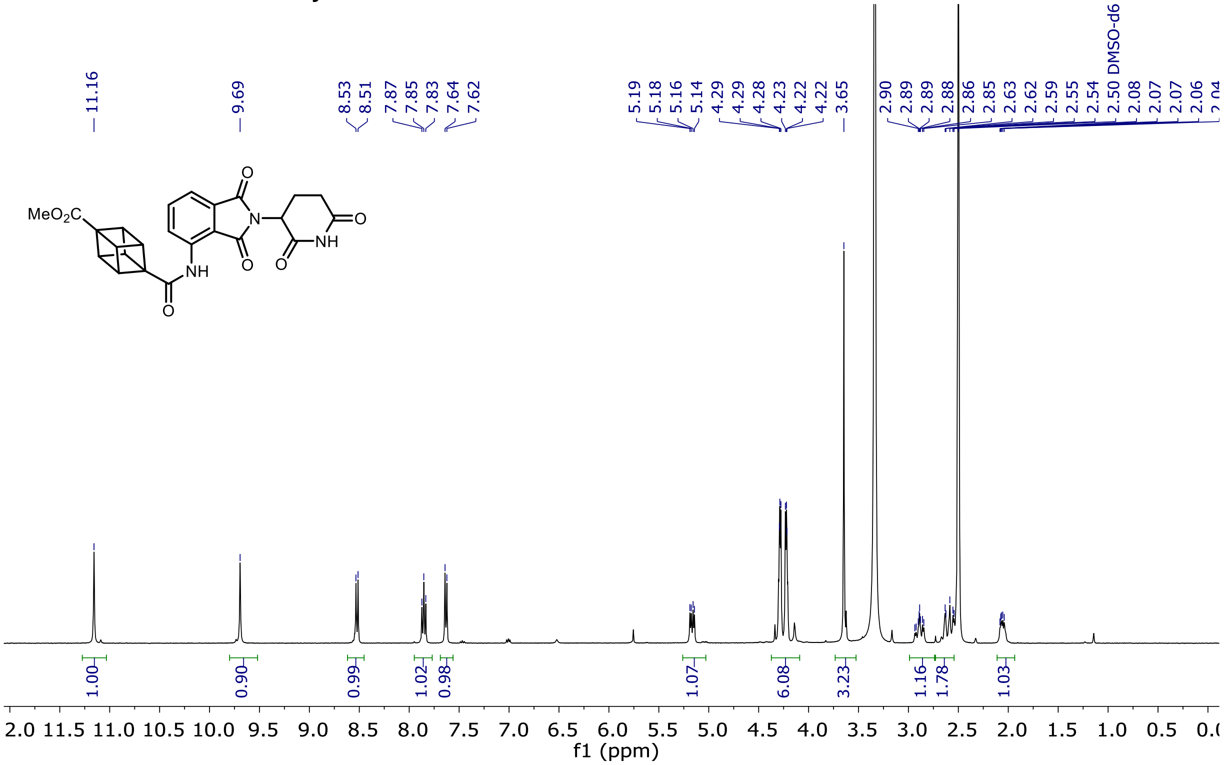

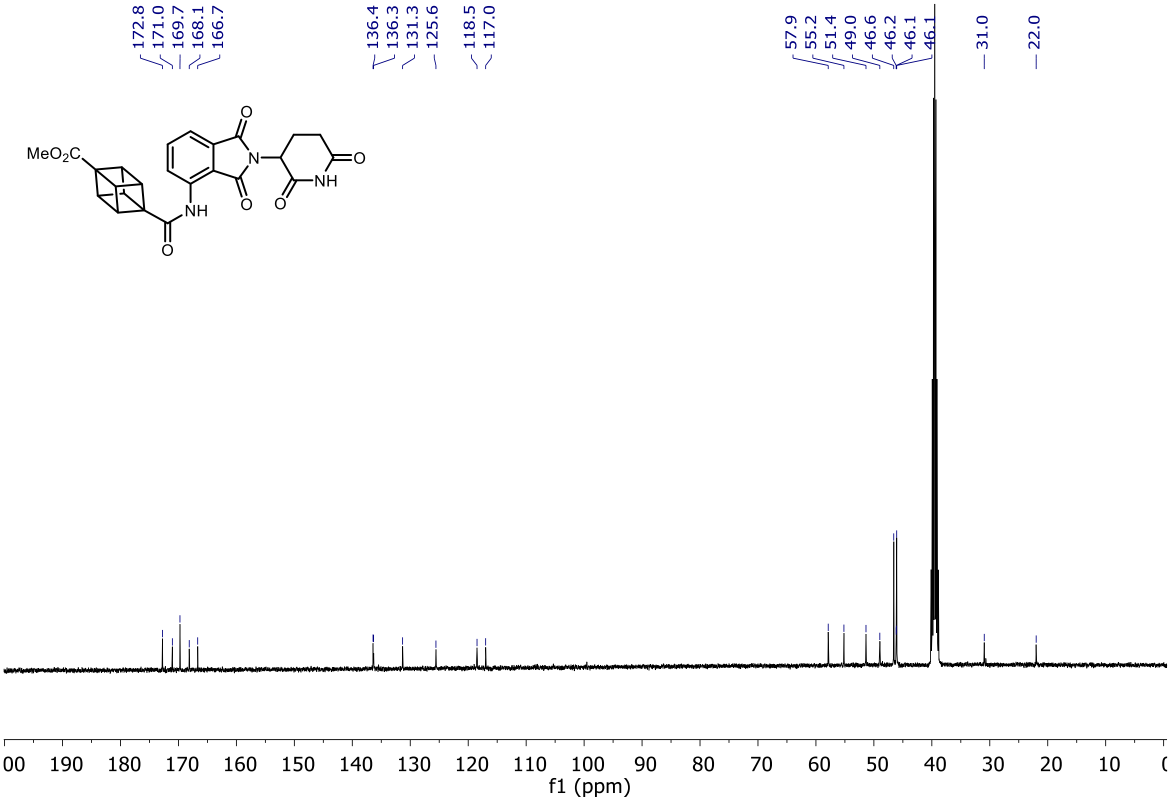

$^{1}H$ and $^{13}C$ NMR of Entry **6**

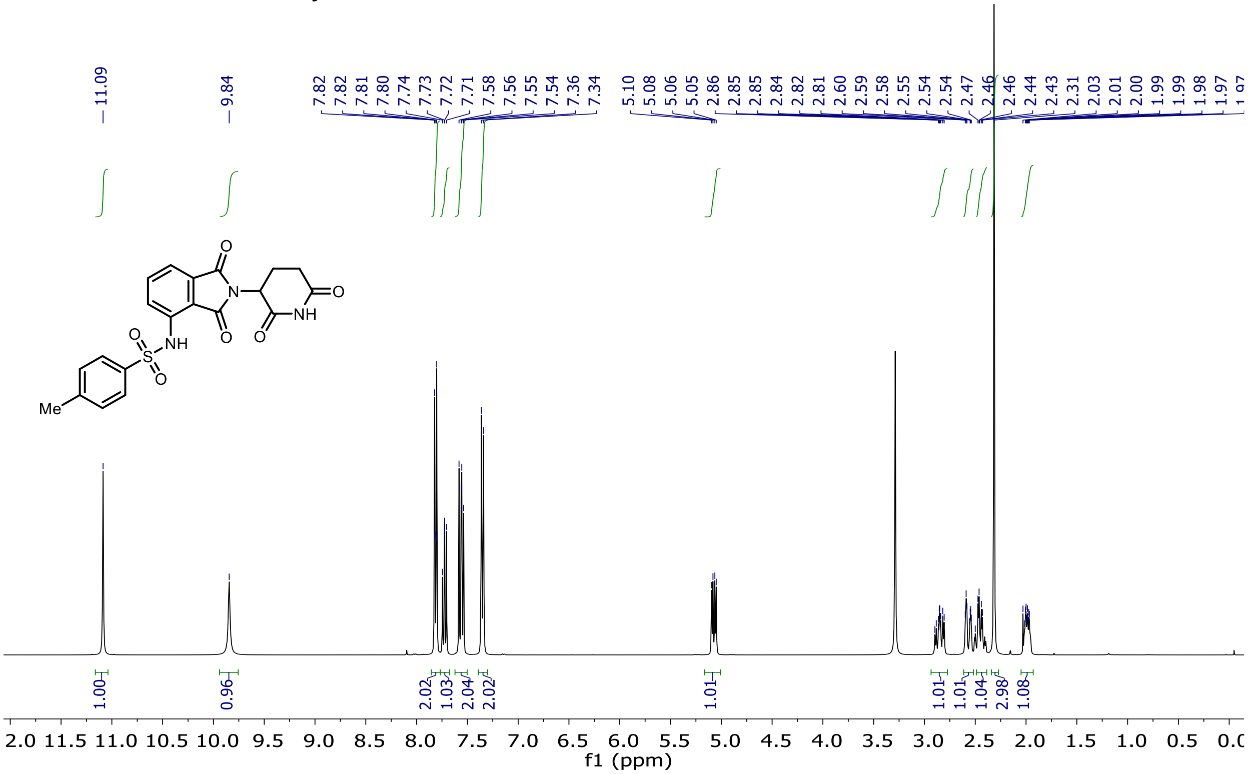

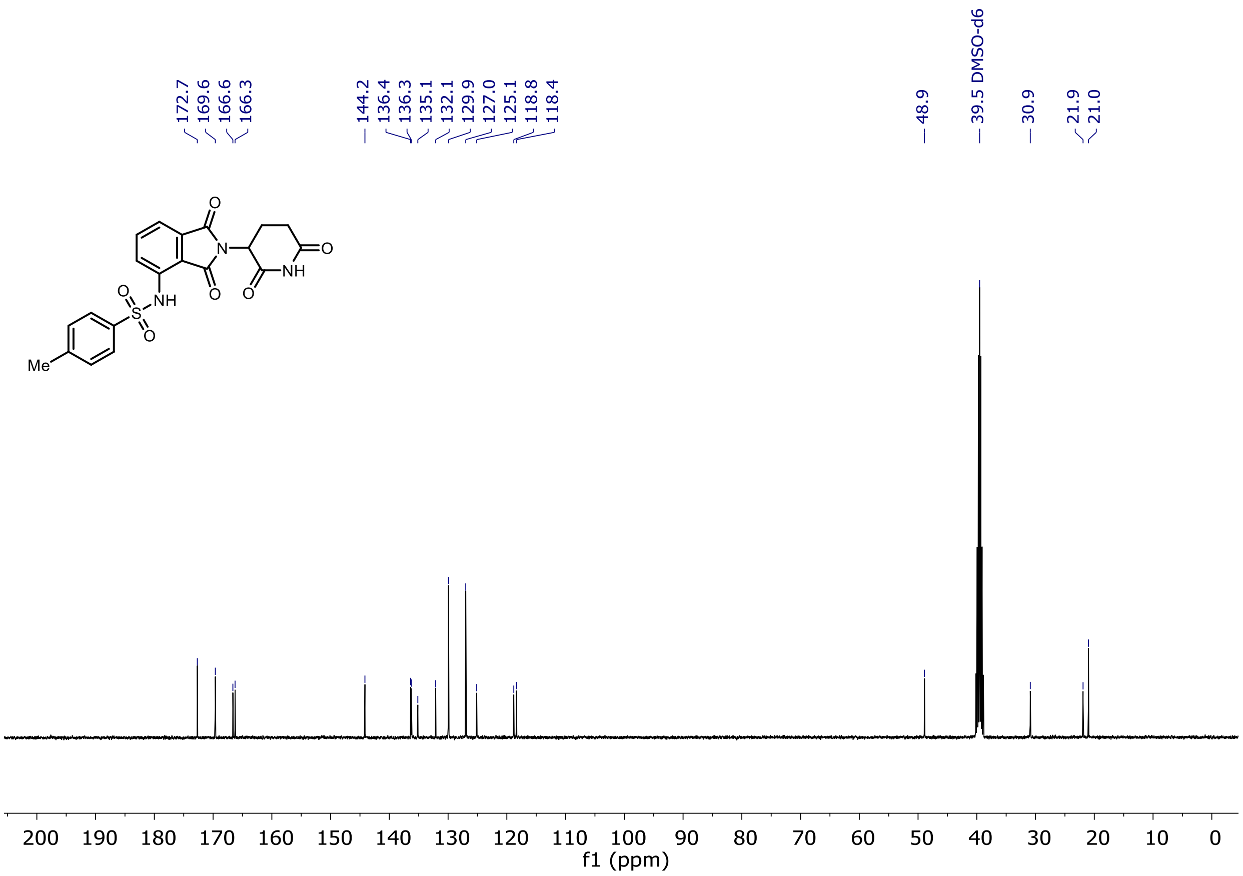

$^{1}$H and $^{13}$C NMR of Entry **72**

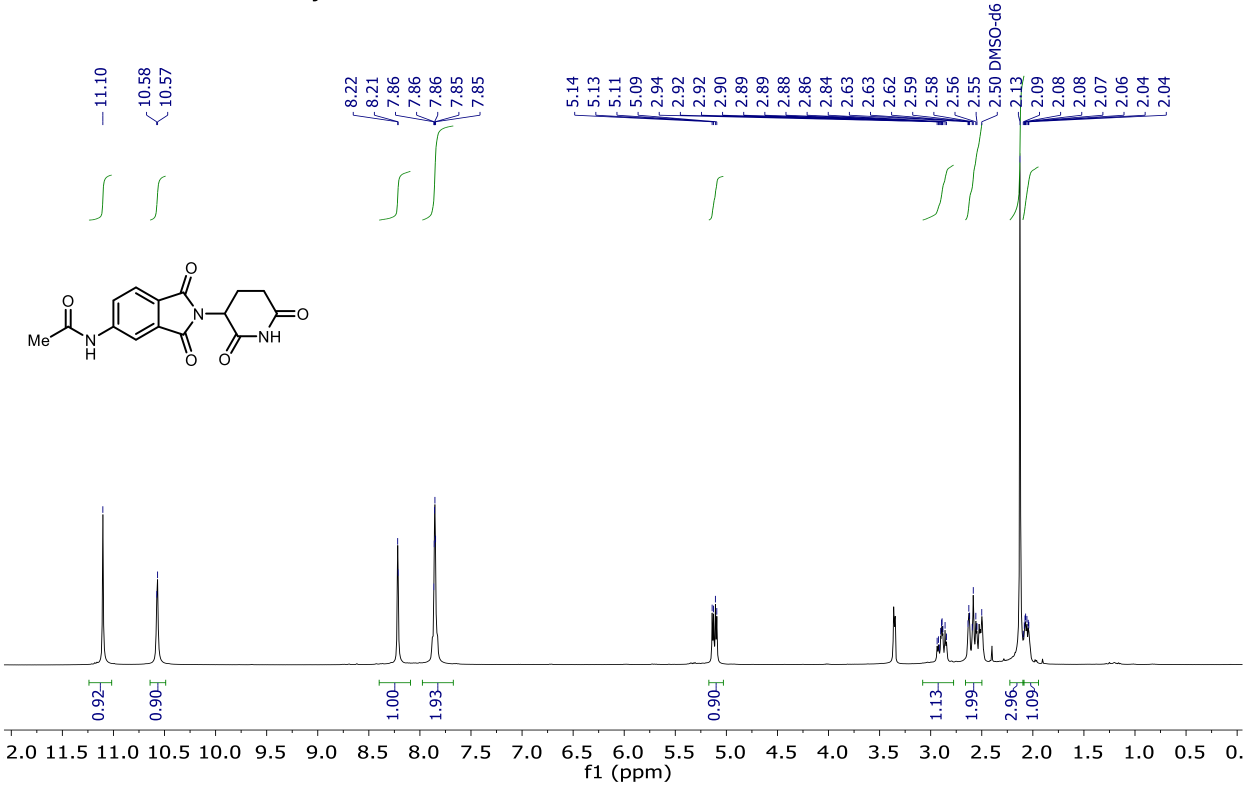

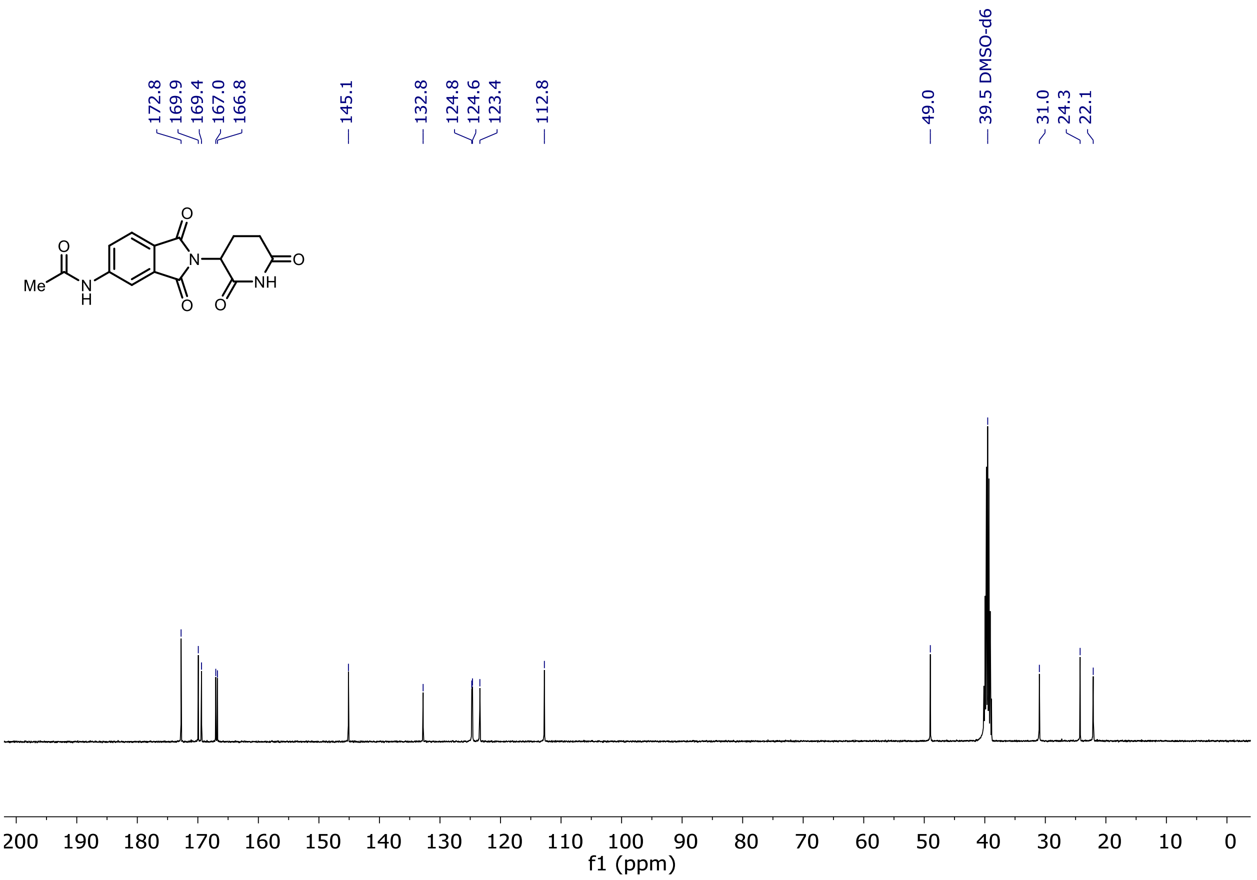

172.8
169.9
169.4
167.0
166.8
145.1
132.8
124.8
124.6
123.4
112.8
49.0
39.5 DMSO-d6
31.0
24.3
22.1
Me
O
N
H
O
O
N
NH
O
O
200 190 180 170 160 150 140 130 120 110 100 90 80 70 60 50 40 30 20 10 0
f1 (ppm)

$^{1}H$ and $^{13}C$ NMR of Entry **73**

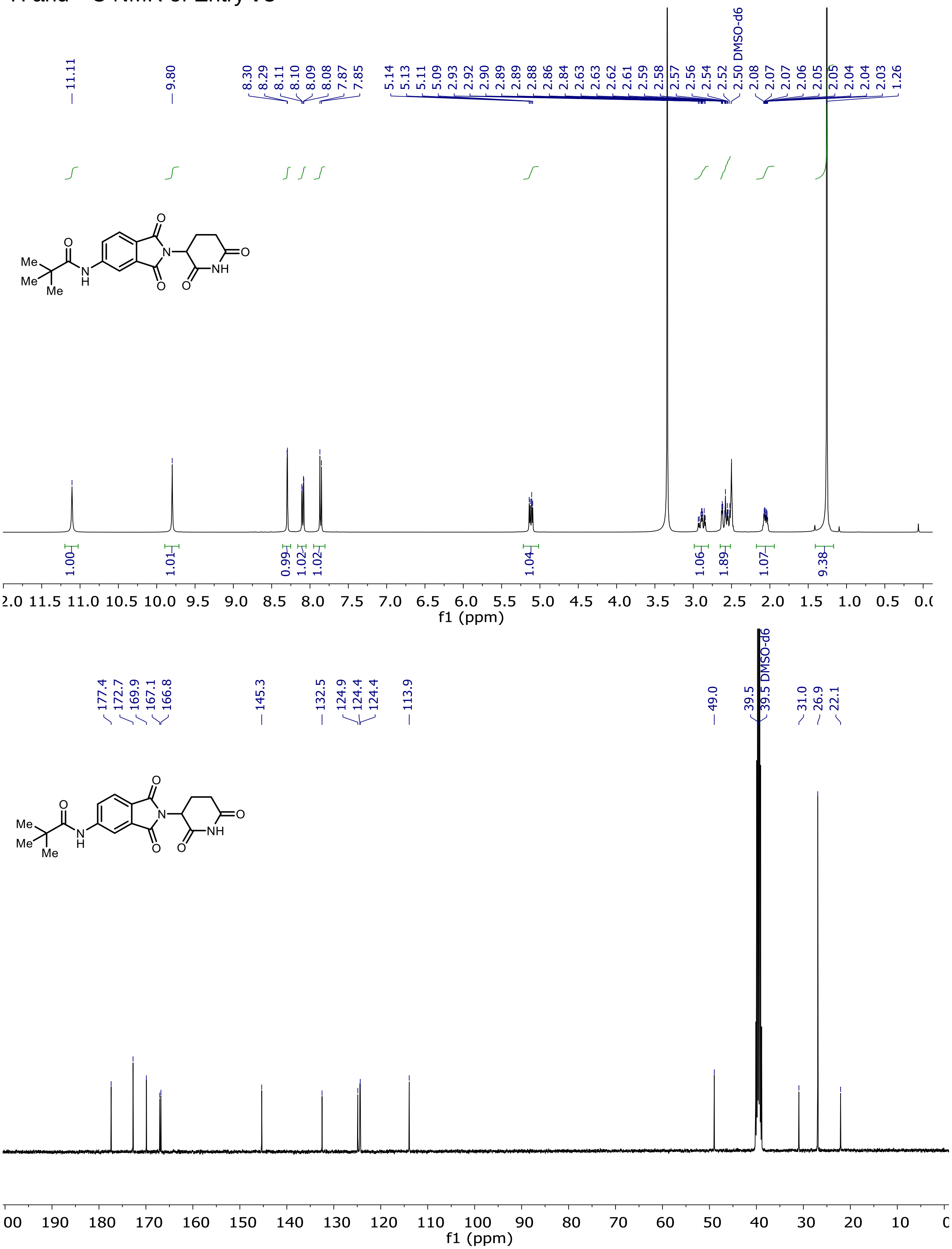

$^{1}H$ and $^{13}C$ NMR of Entry **74**

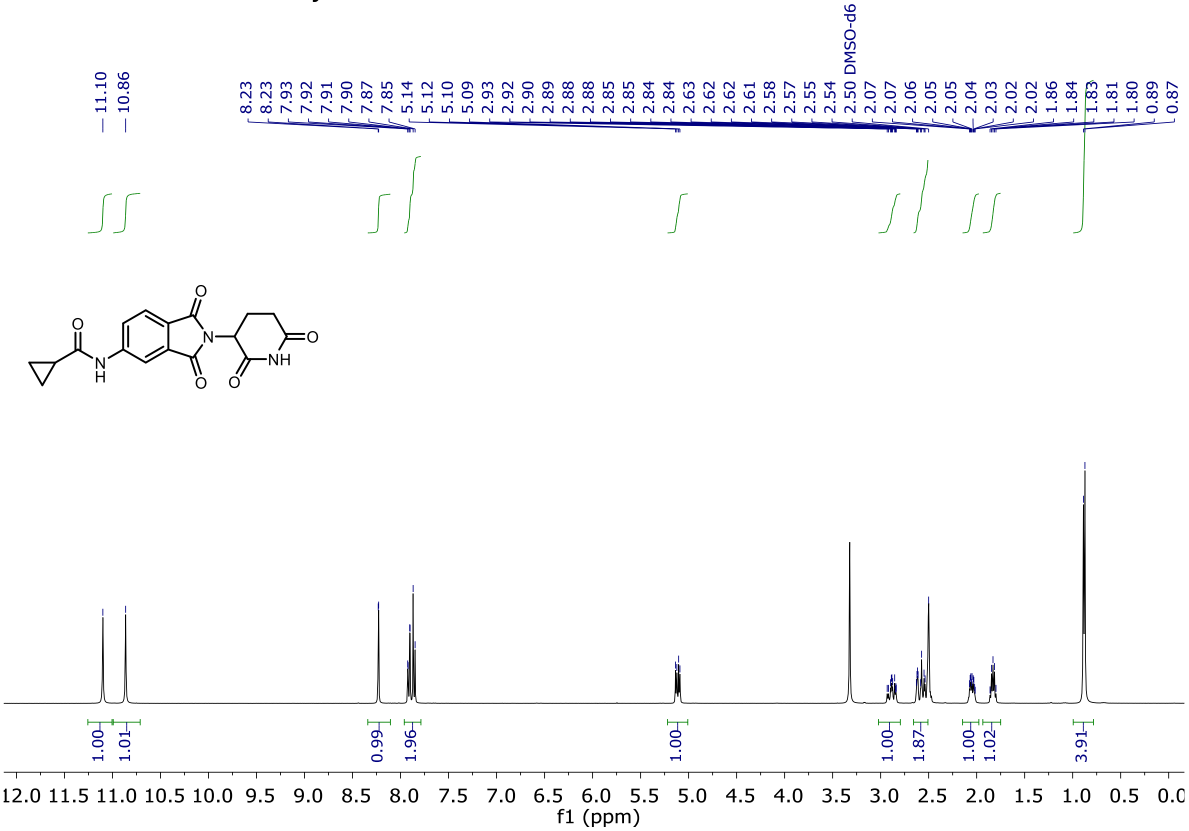

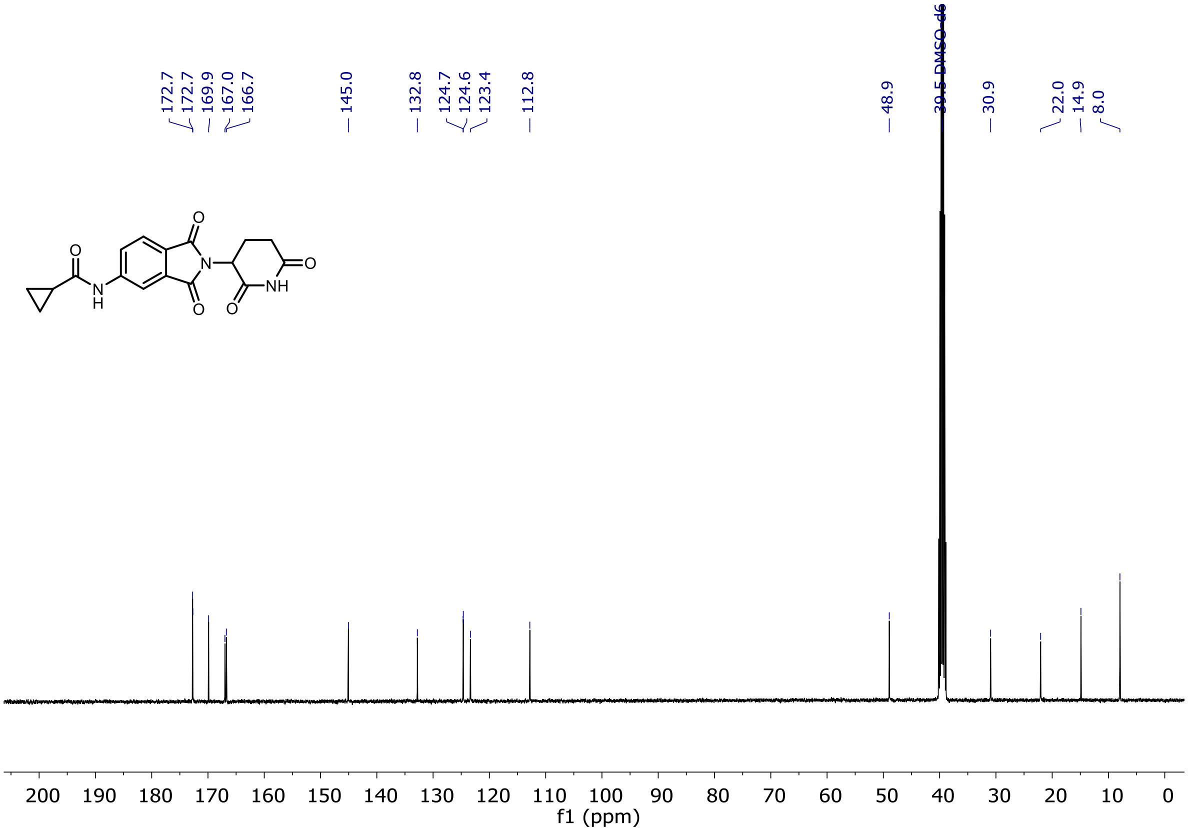

$^{1}H$ and $^{13}C$ NMR of Entry **75**

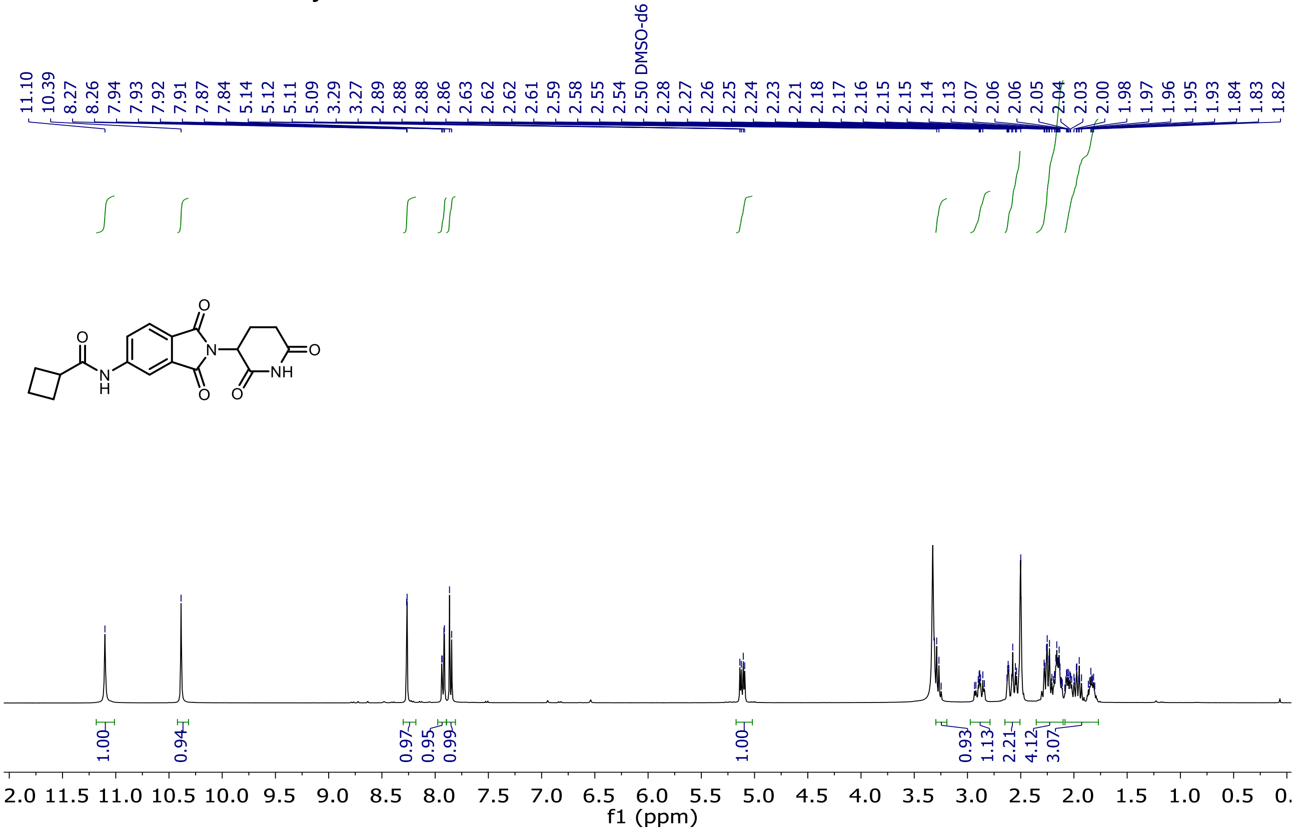

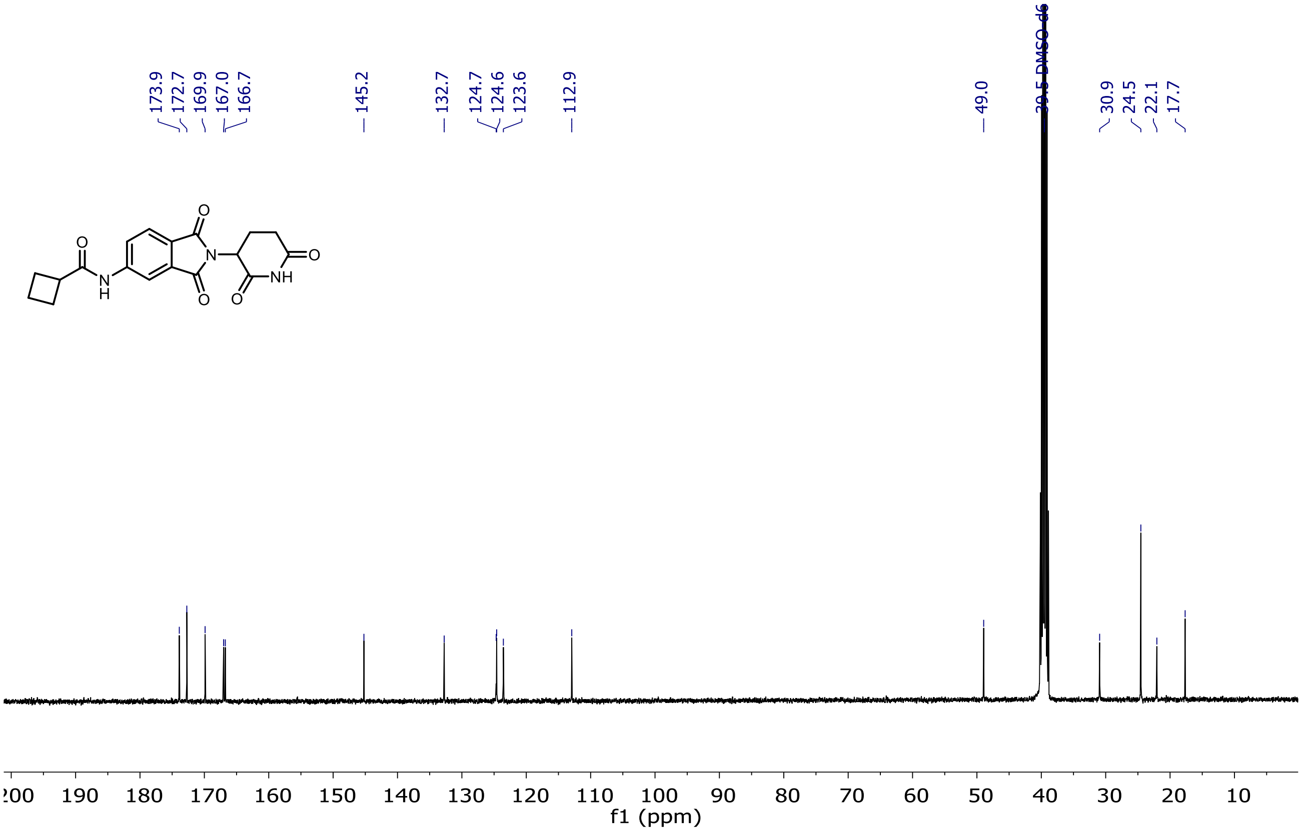

¹H and ¹³C NMR of Entry **76**

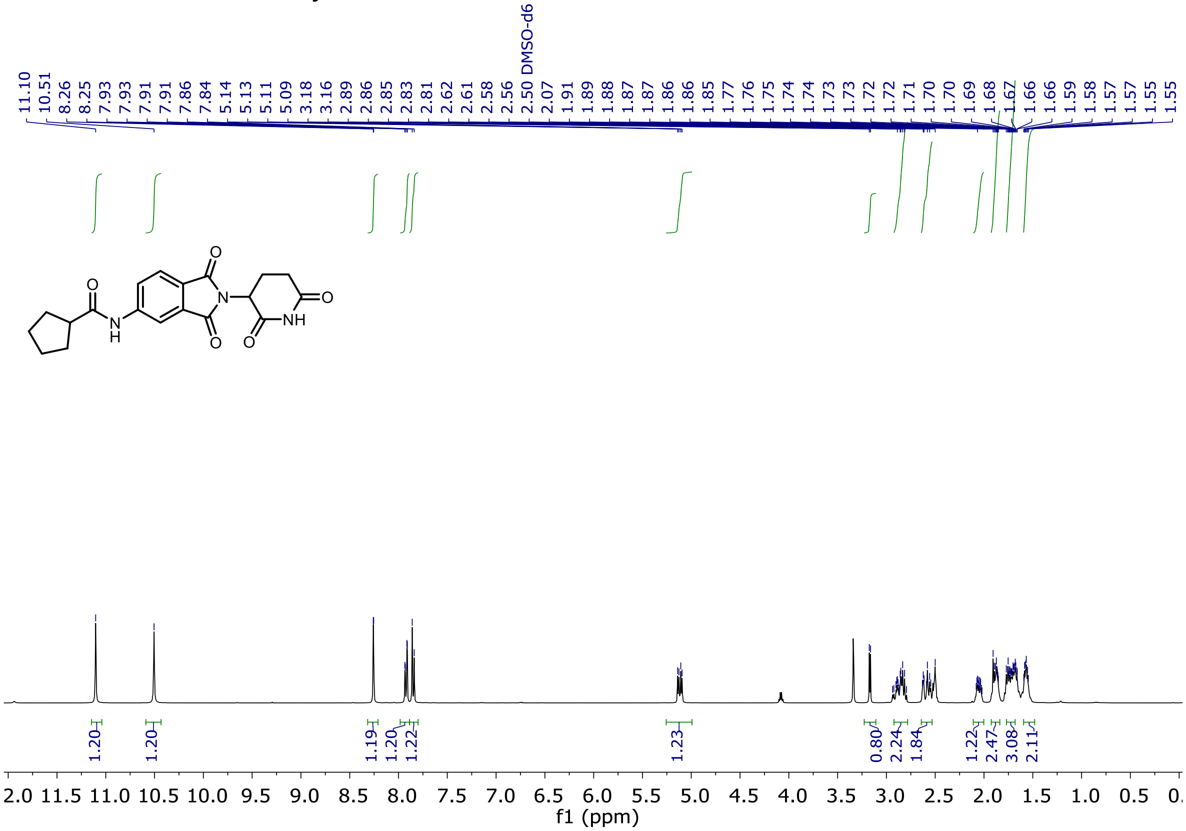

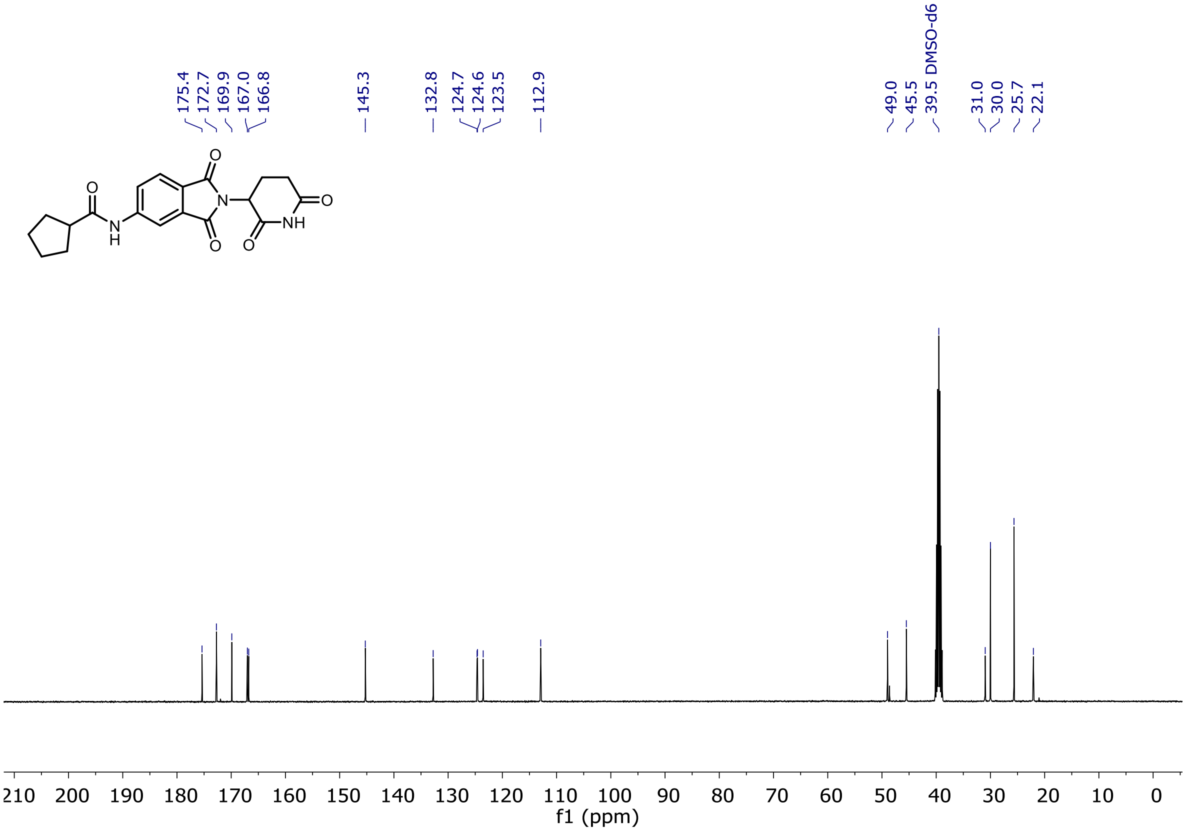

<sup></sup>

[1]H and [13]C NMR of Entry **77**

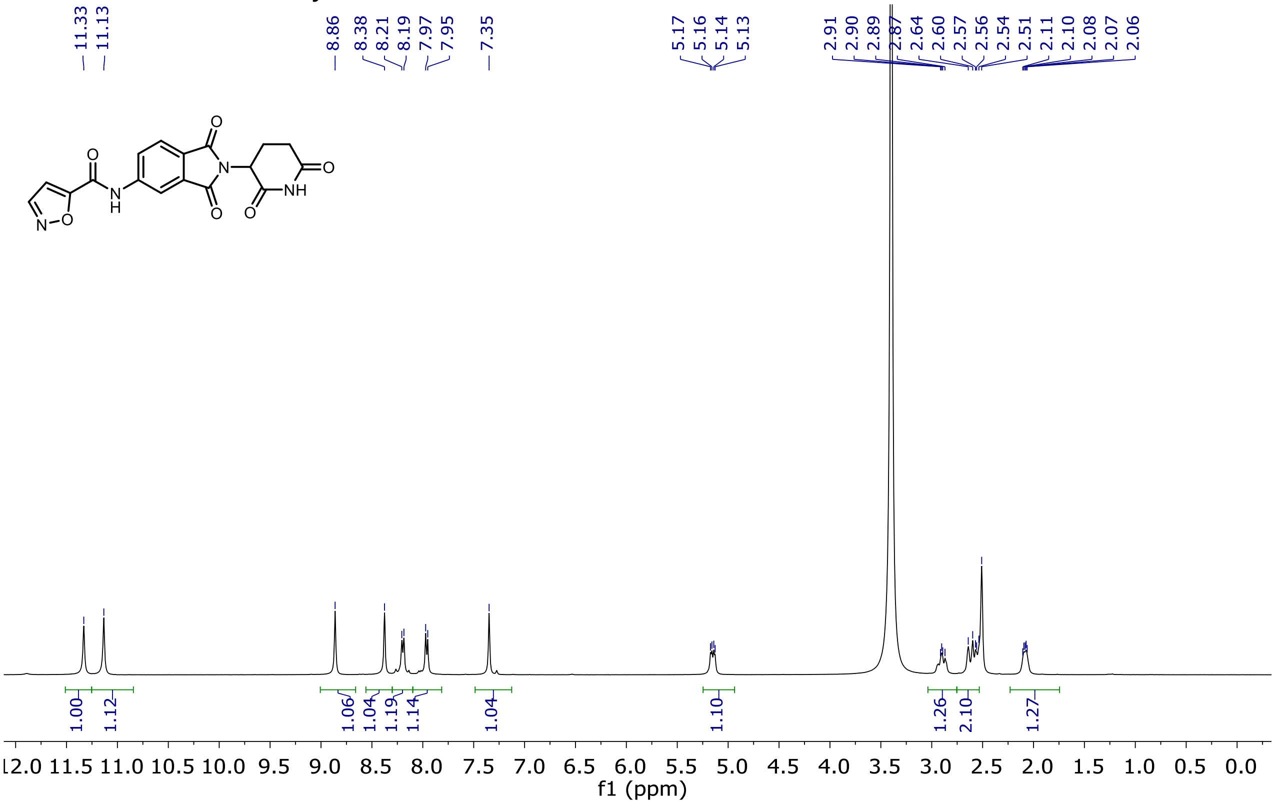

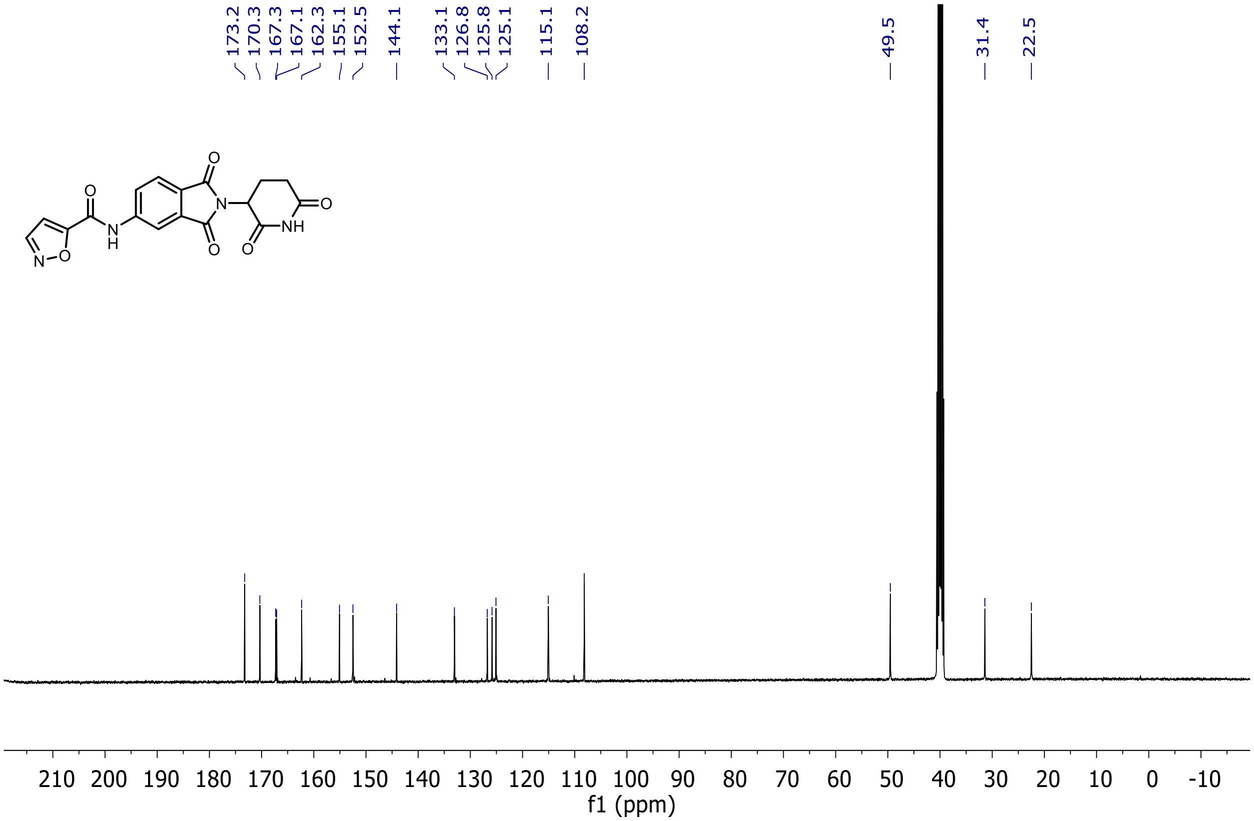
173.2
170.3
167.3
167.1
162.3
155.1
152.5
144.1
133.1
126.8
125.8
125.1
115.1
108.2
49.5
31.4
22.5
O
N
H
N
O
O
N
O
NH
O
O
210 200 190 180 170 160 150 140 130 120 110 100 90 80 70 60 50 40 30 20 10 0 -10
f1 (ppm)

[1]H and [13]C NMR of Entry **78**

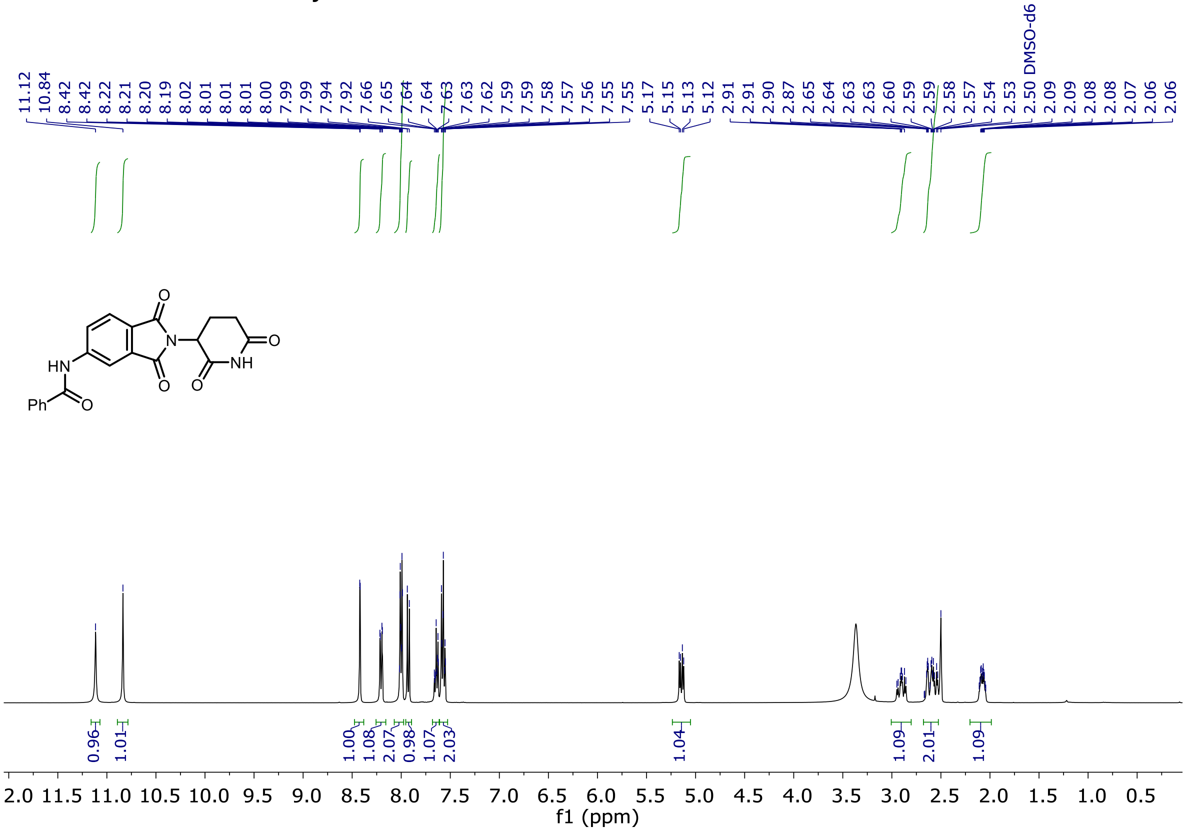

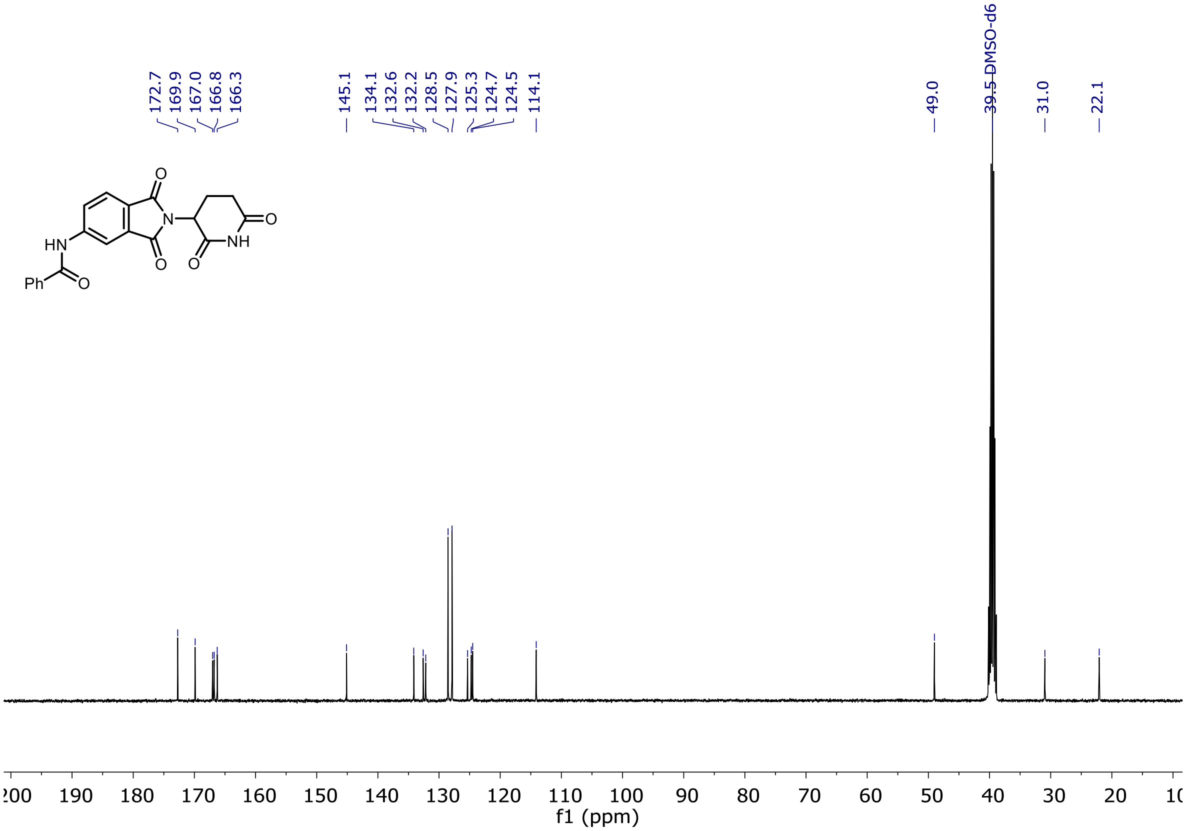

$^{1}$H and $^{13}$C NMR of Entry **79**

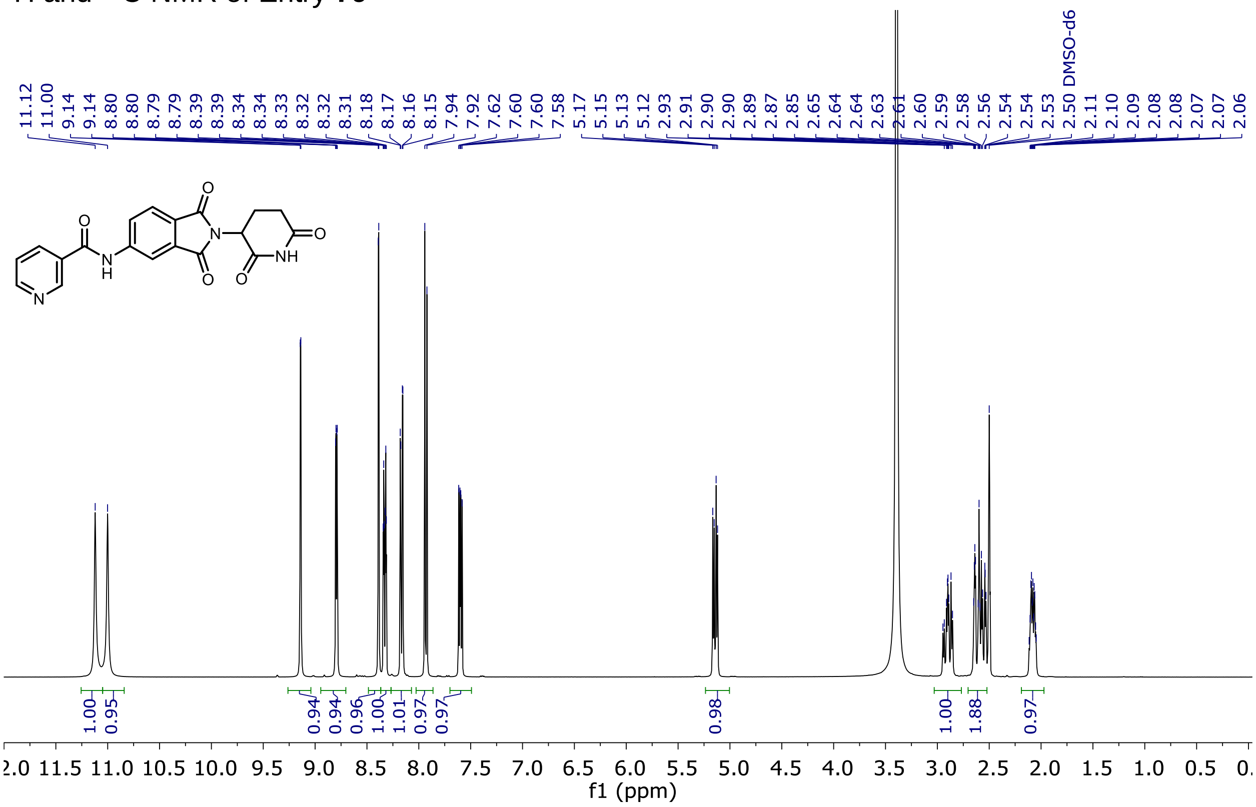

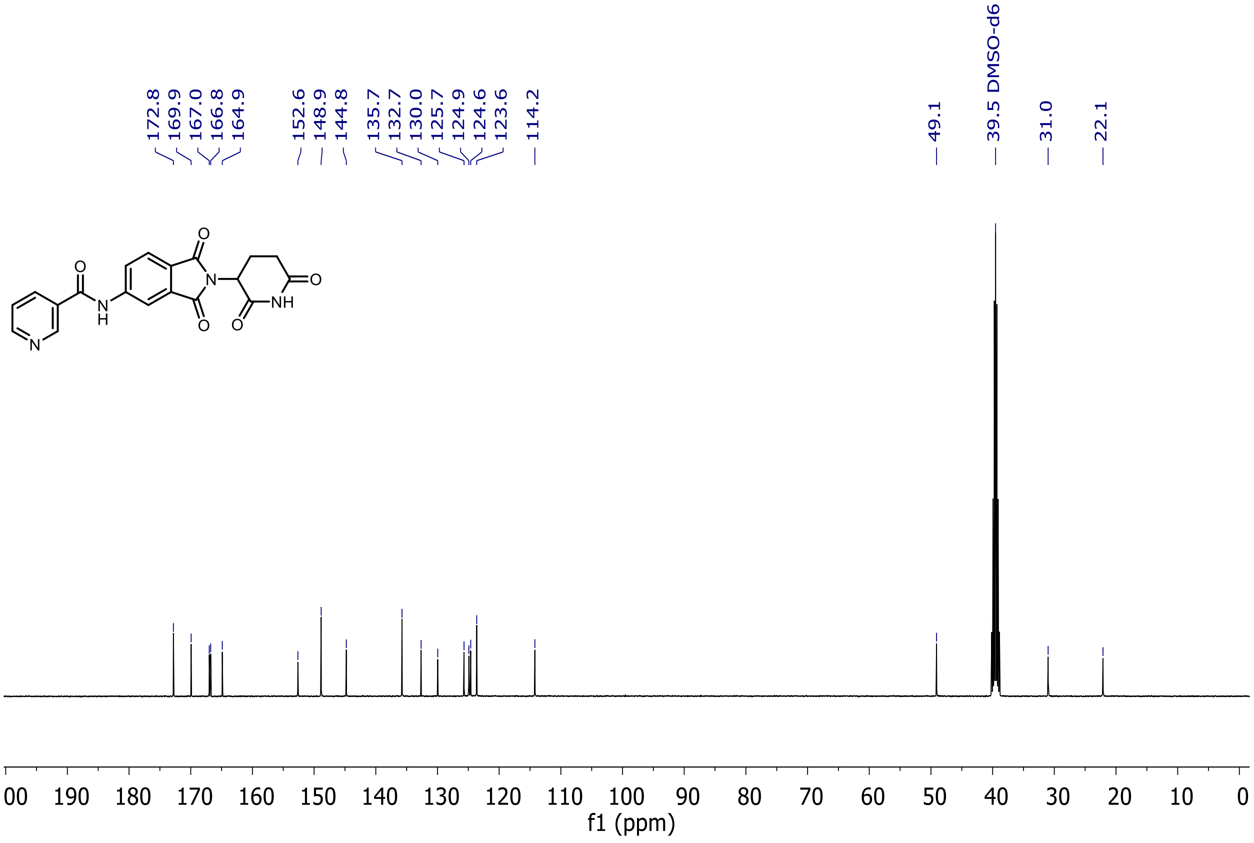

$^{1}$H and $^{13}$C NMR of Entry **80**

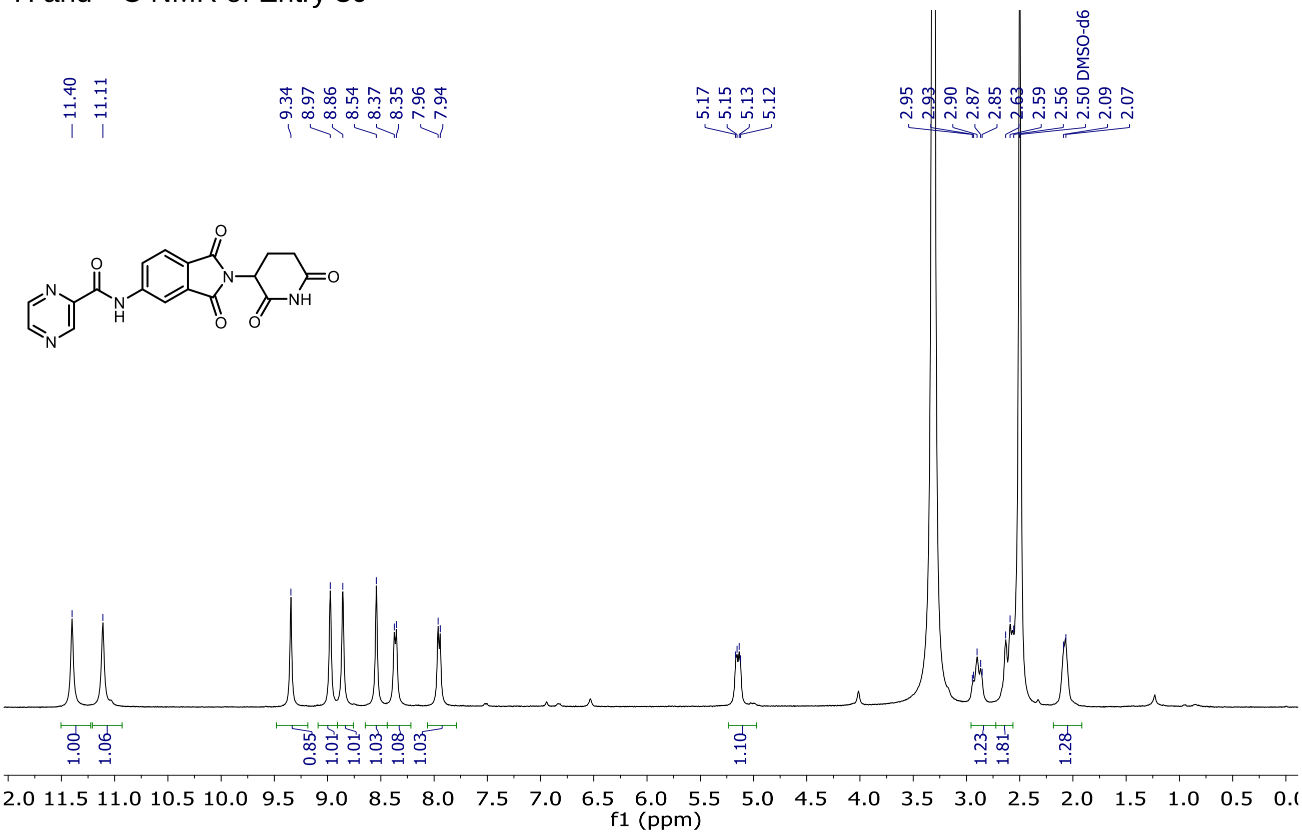

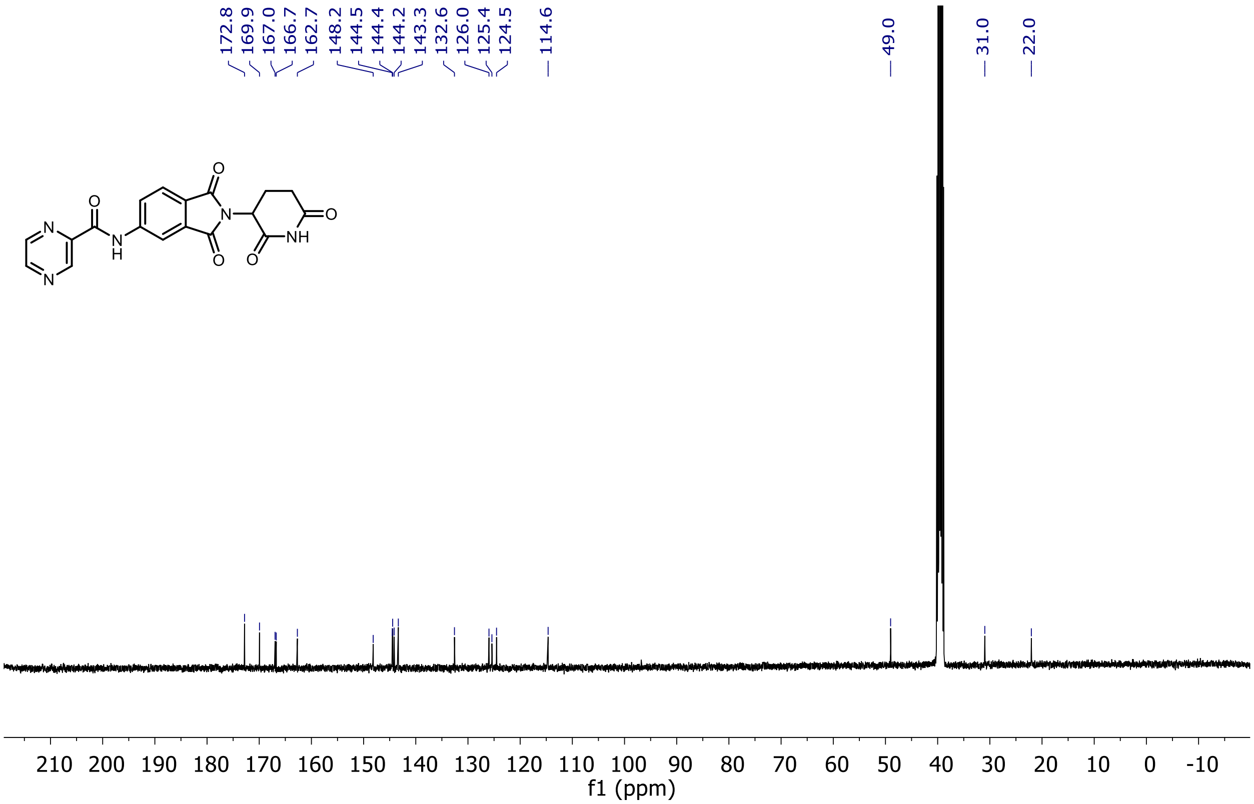

¹H and ¹³C NMR of Entry **81**

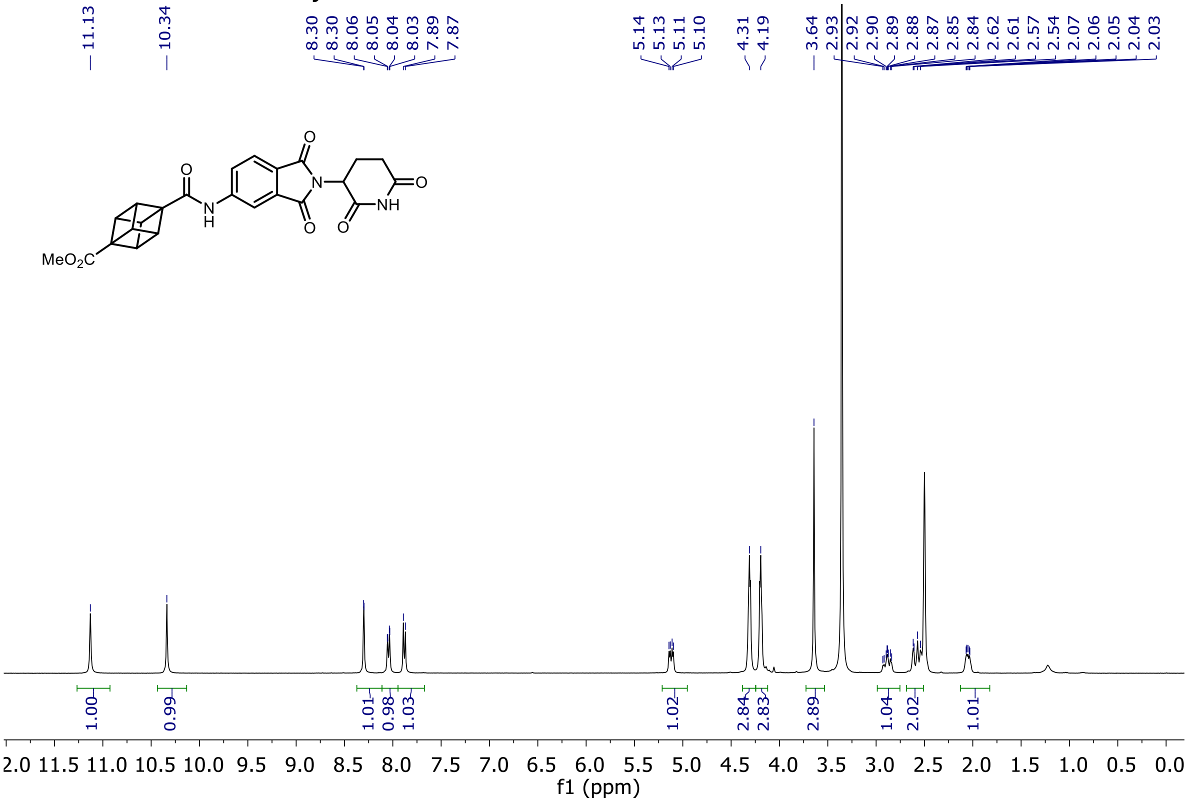

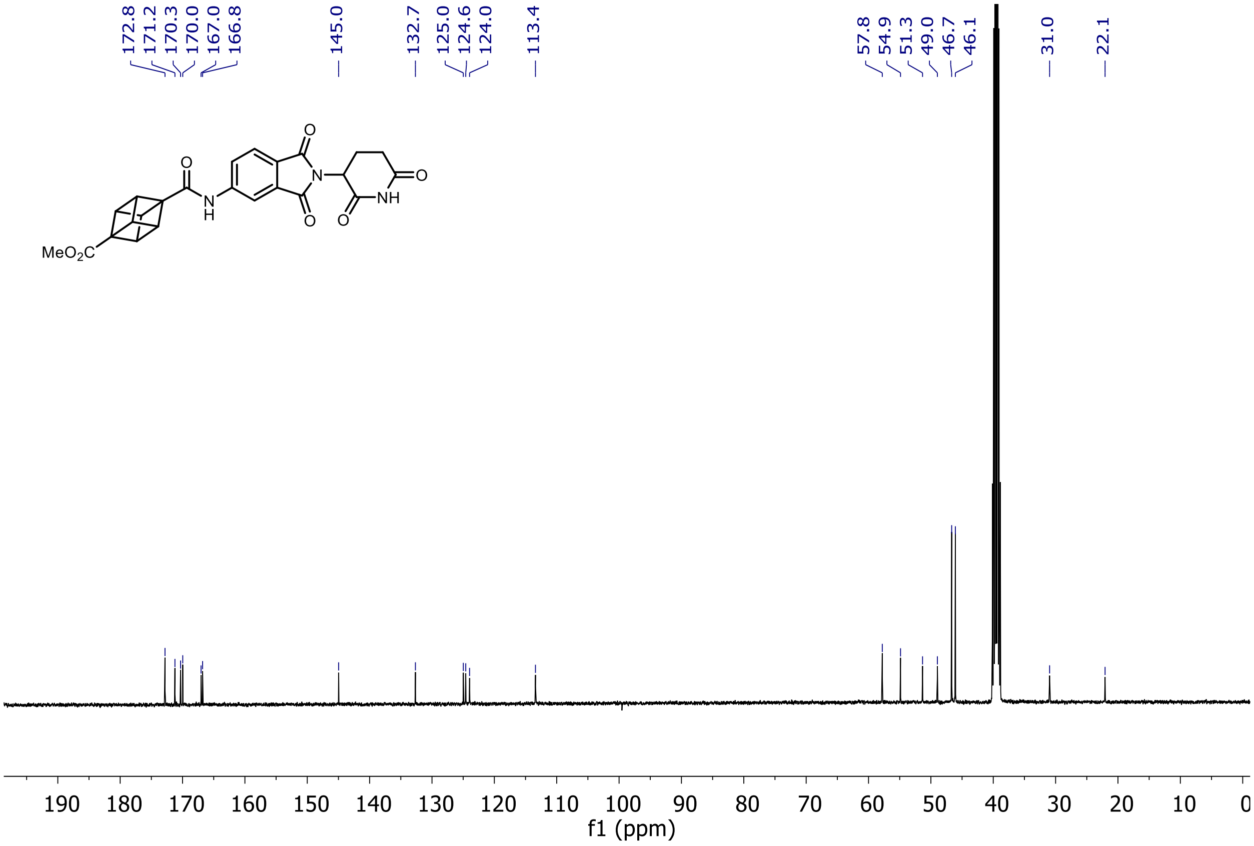

$^{1}H$ and $^{13}C$ NMR of Entry **16**

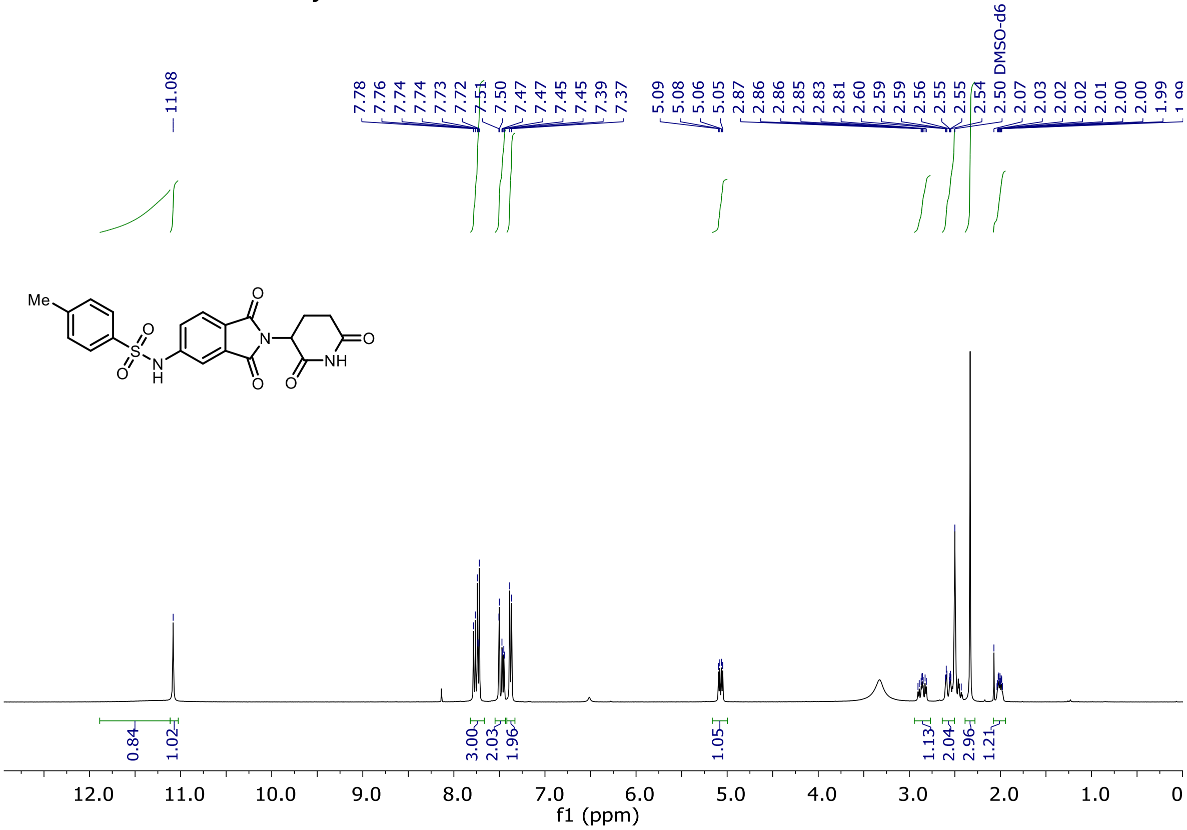

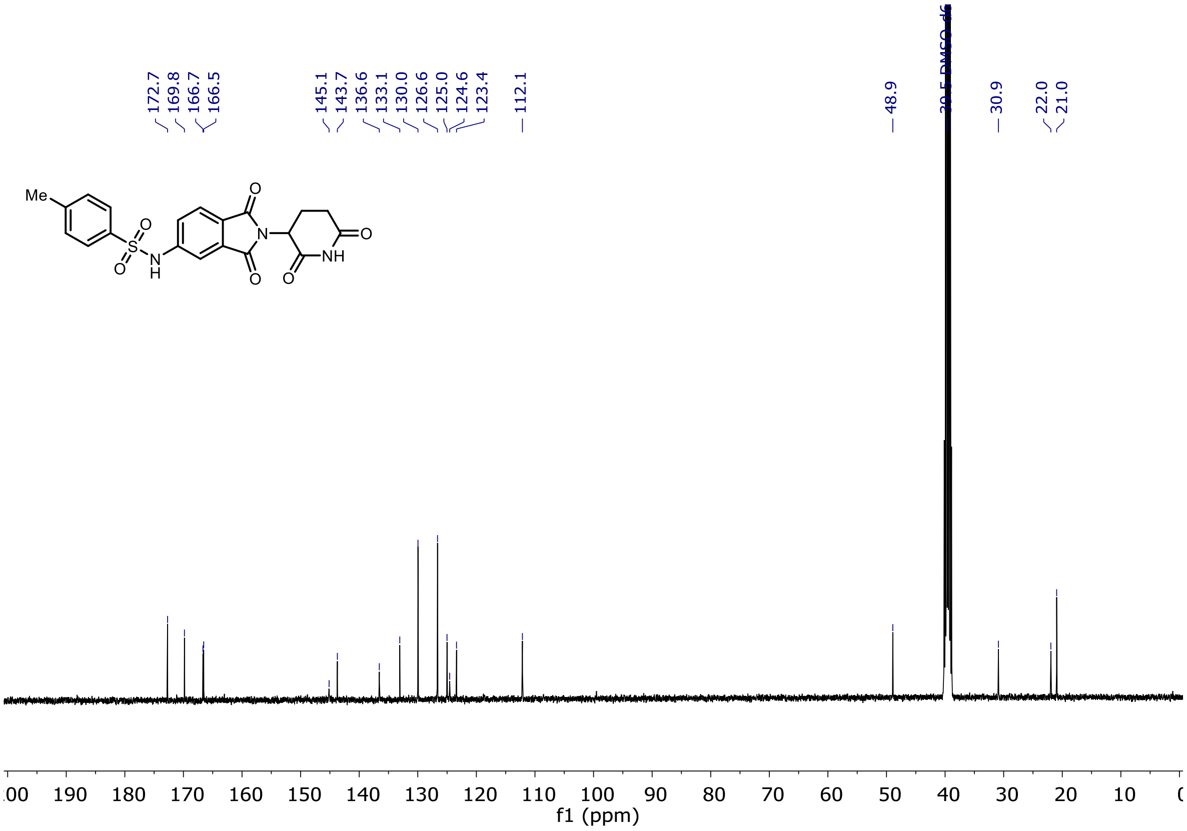

$^{1}H$ and $^{13}C$ NMR of Entry **7**

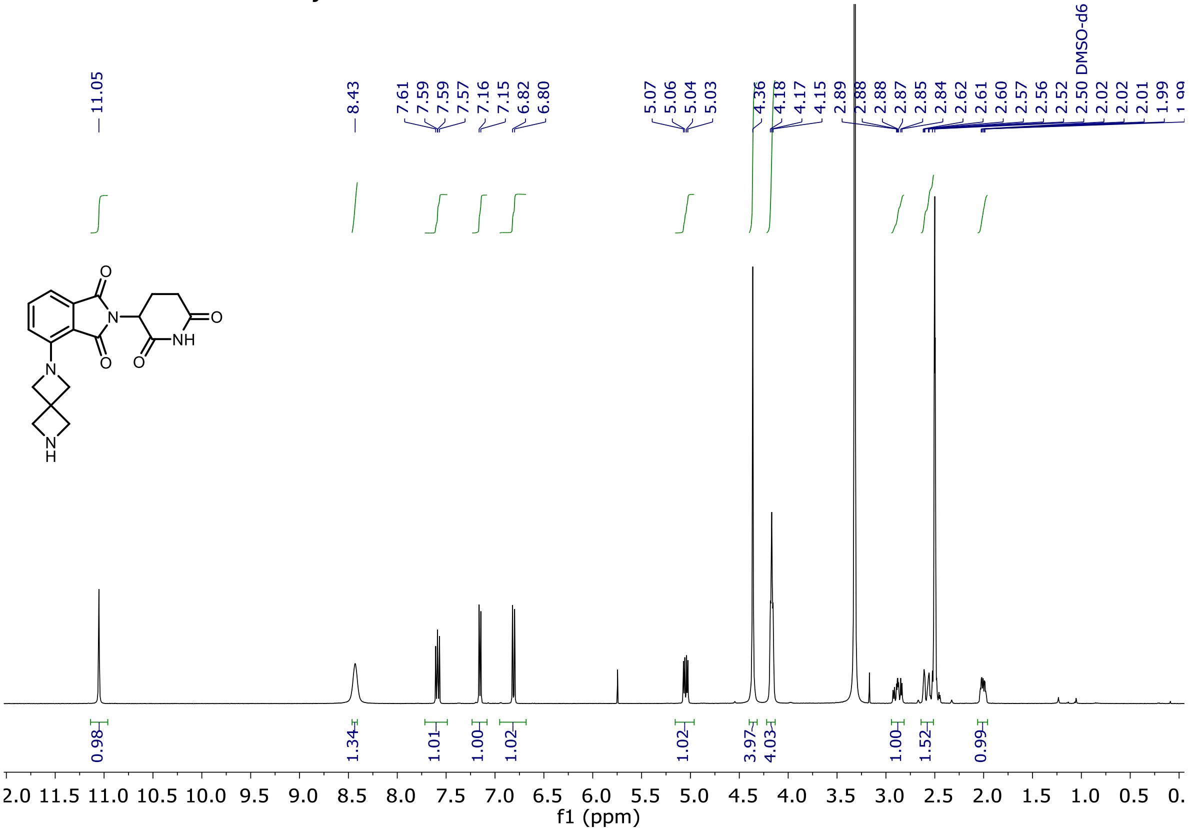

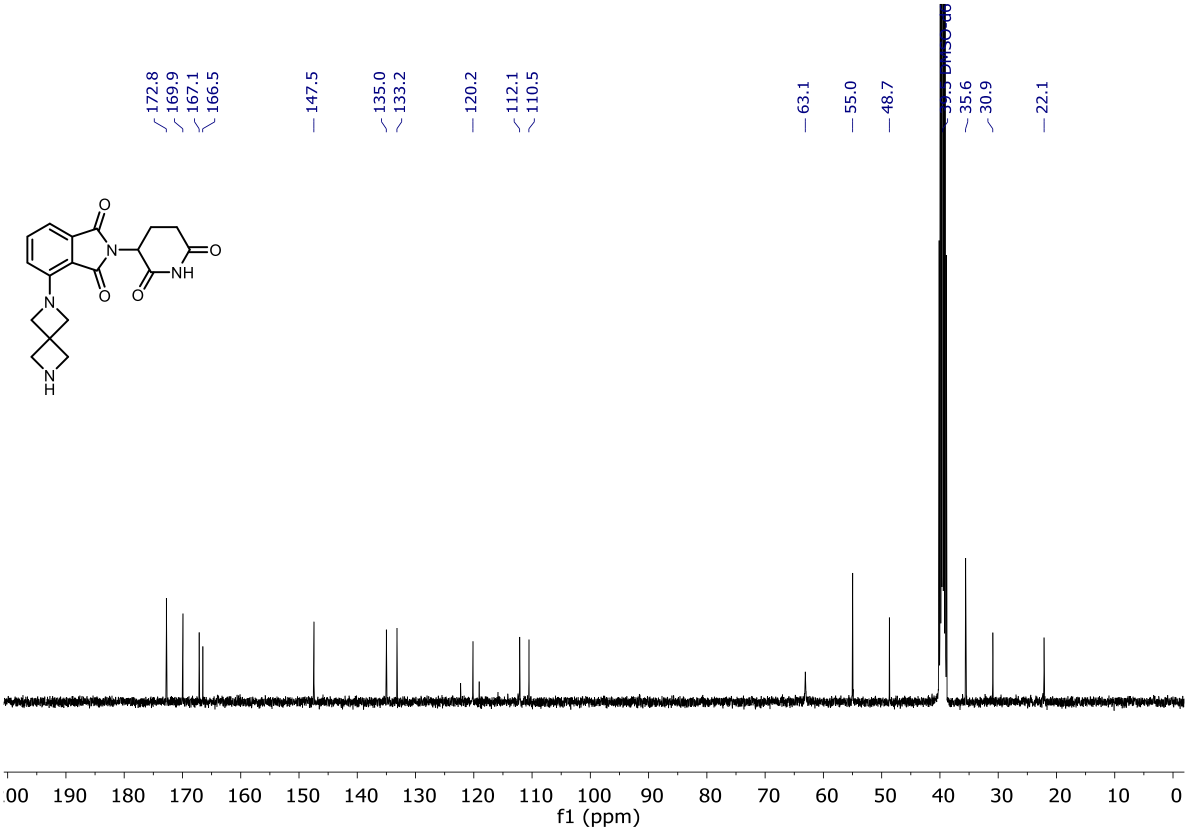

$^{1}H$ and $^{13}C$ NMR of Entry **14**

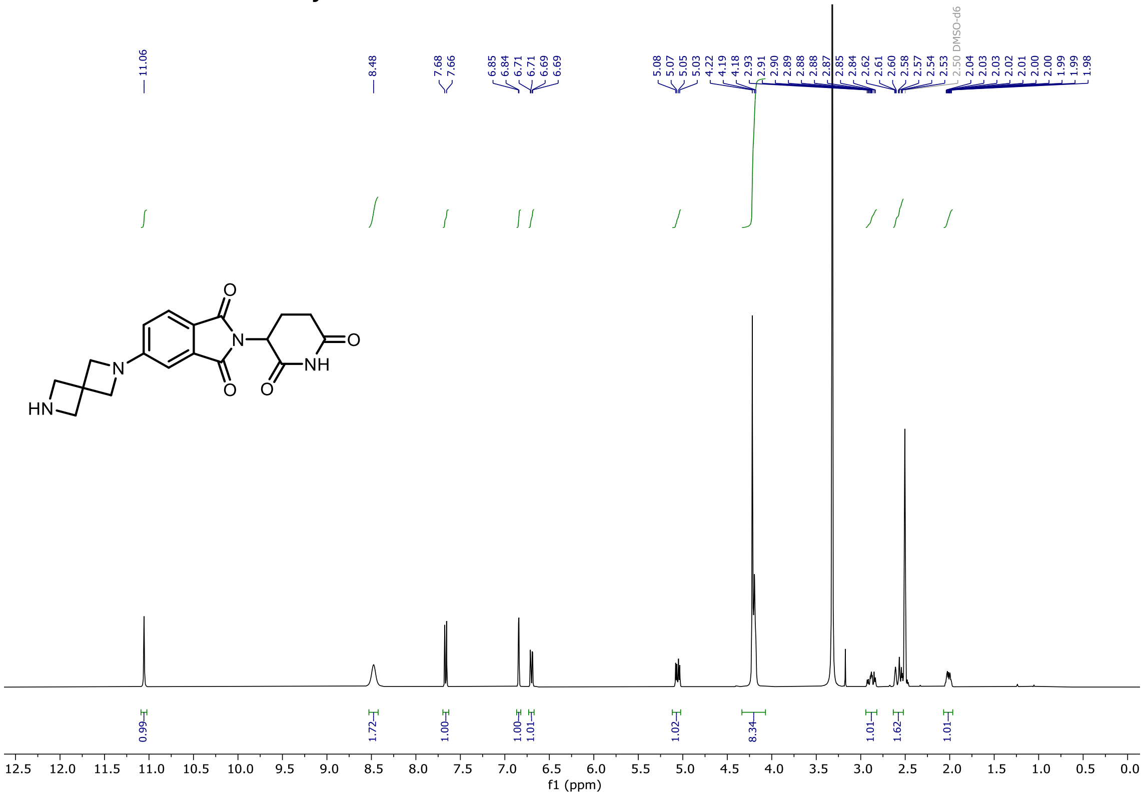

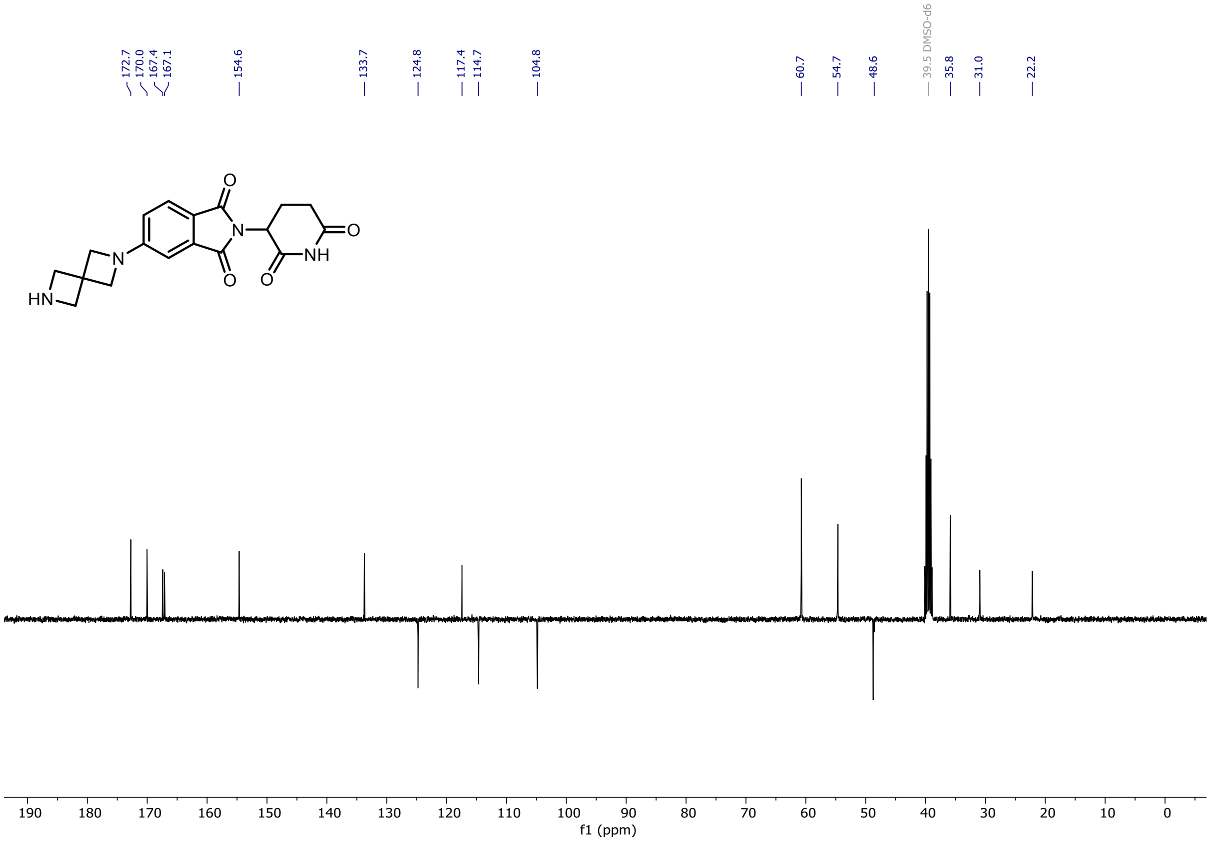

$^{1}H$ and $^{13}C$ NMR of **dALK-1**

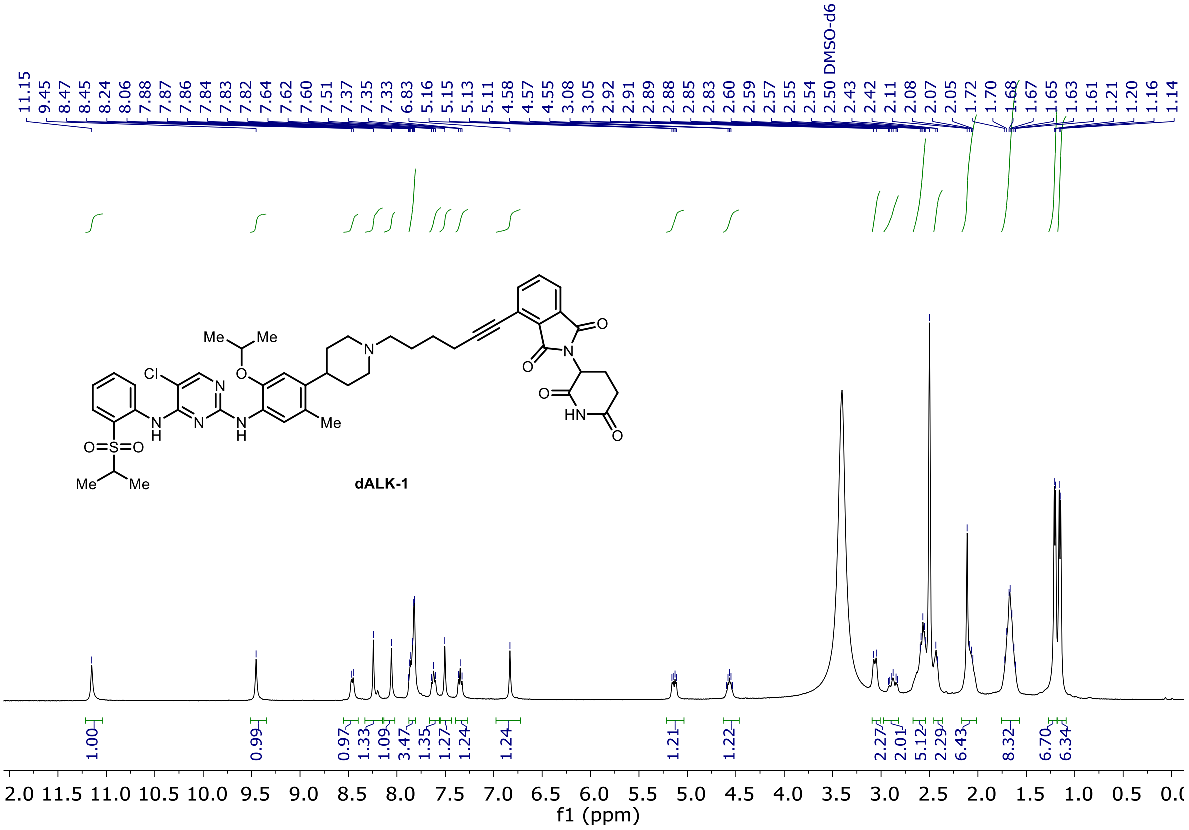

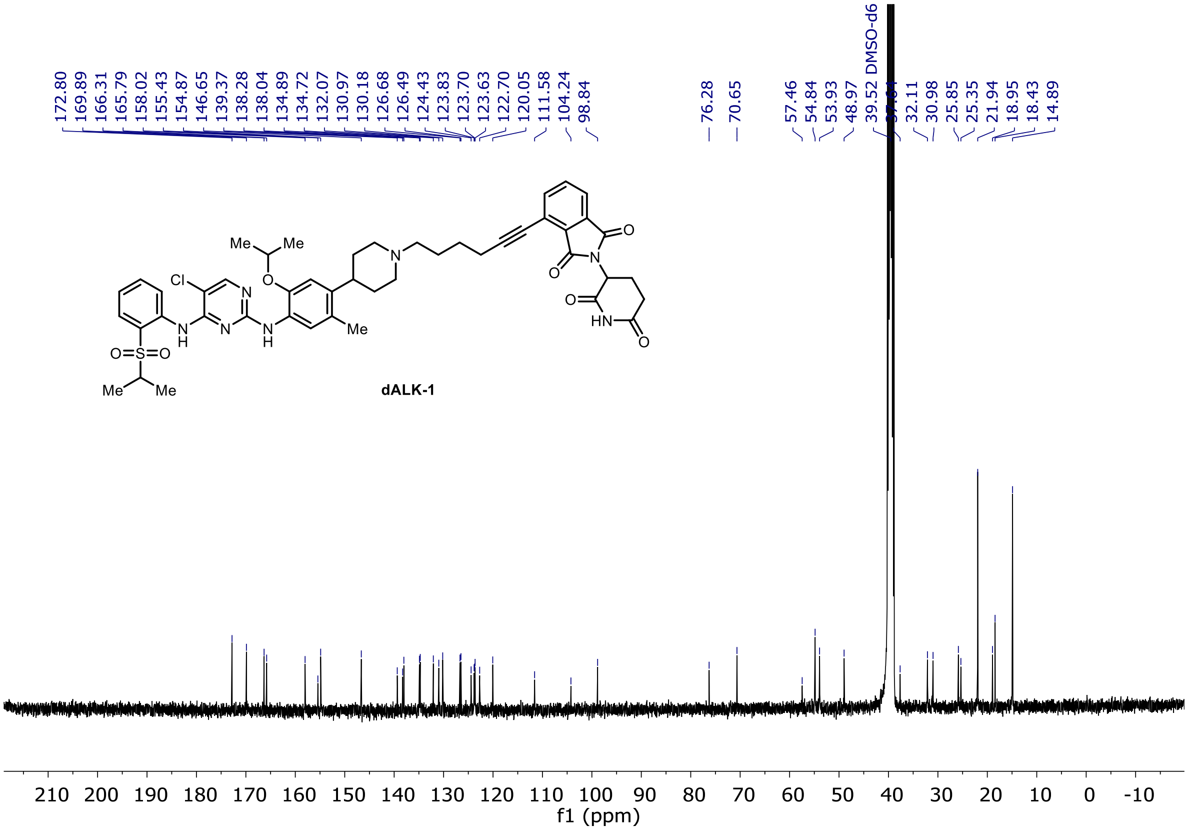

[1]H and [13]C NMR of **dALK-2**

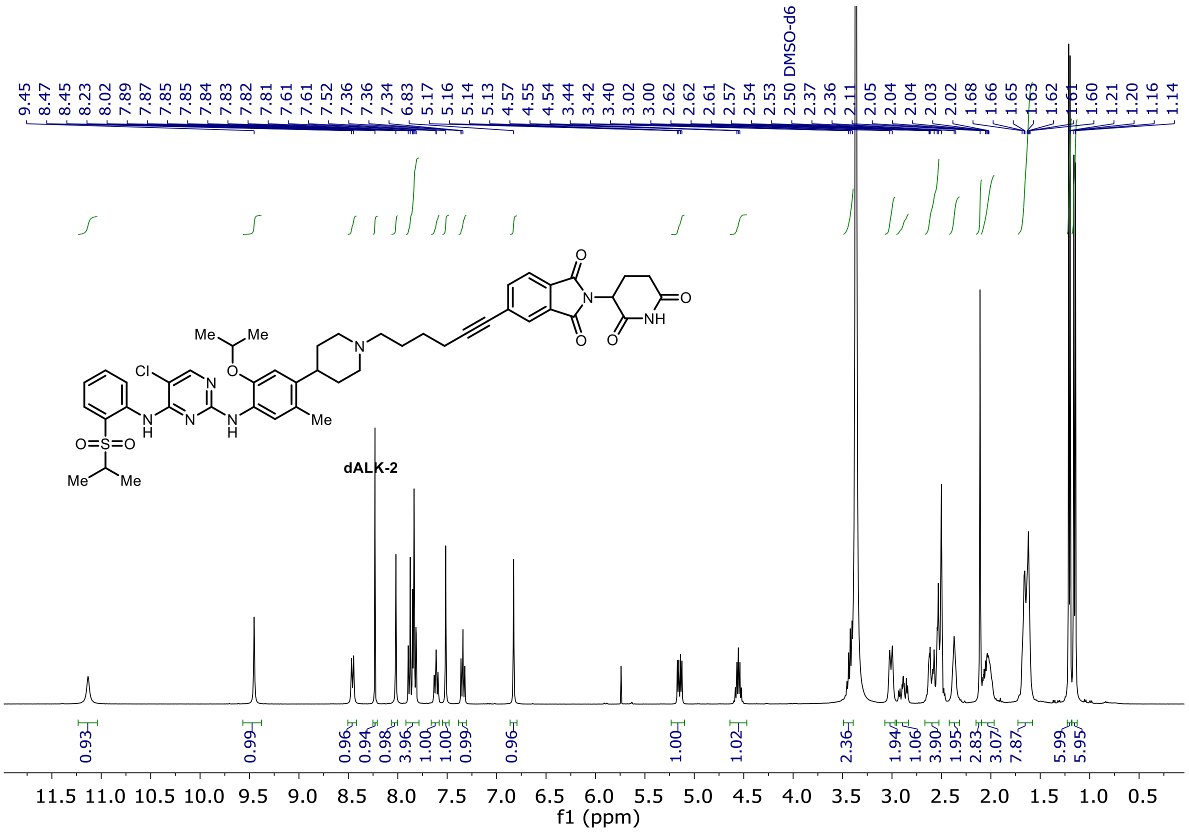

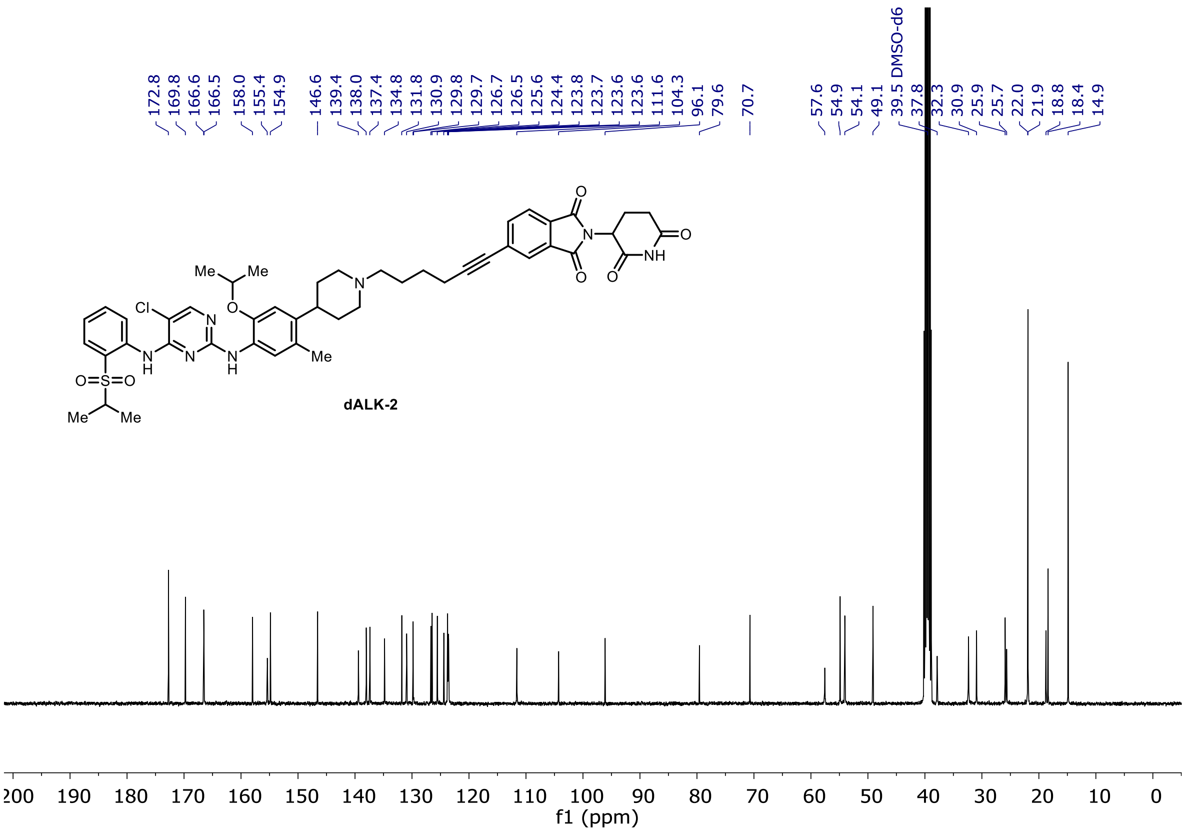

[1]H and [13]C NMR of **dALK-3**

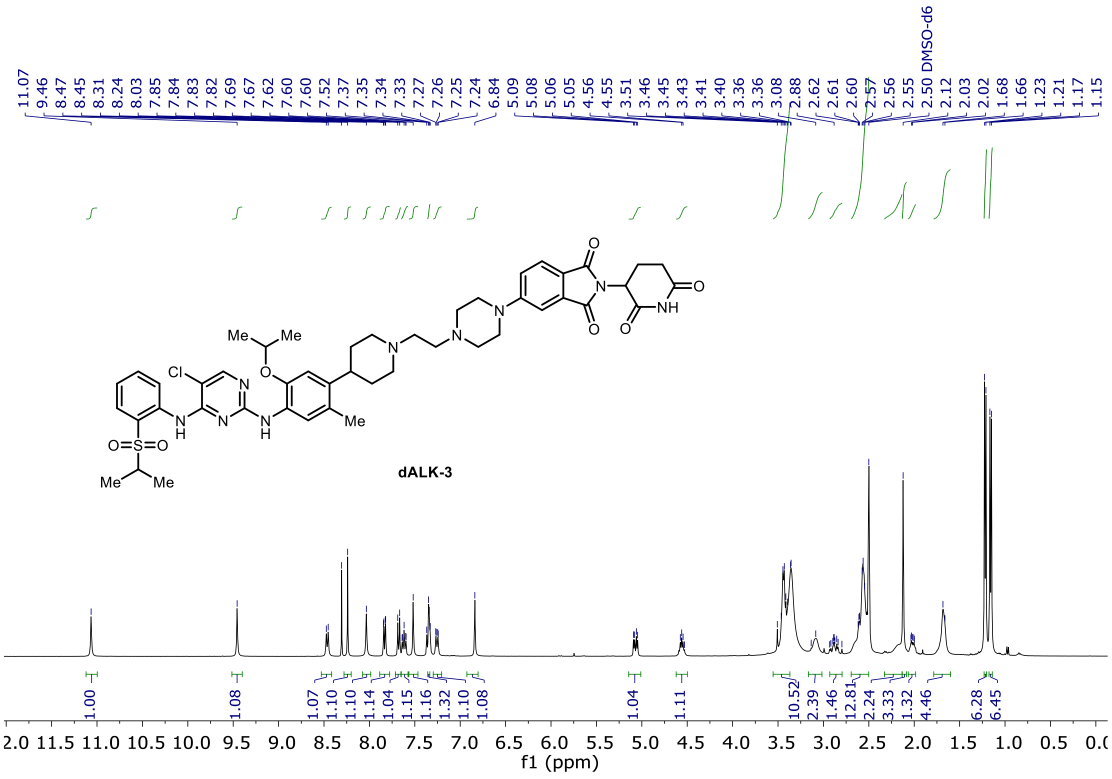

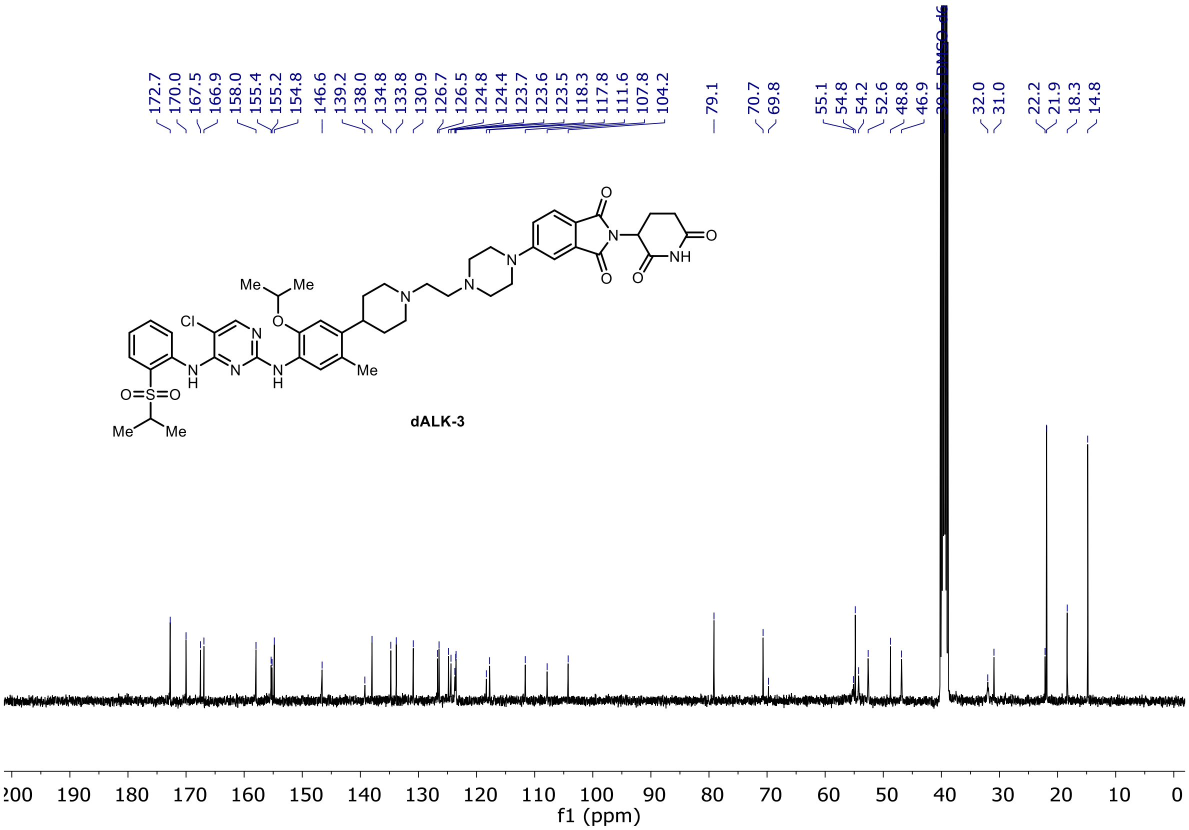

### [1]H and [13]C NMR of **dALK-4**

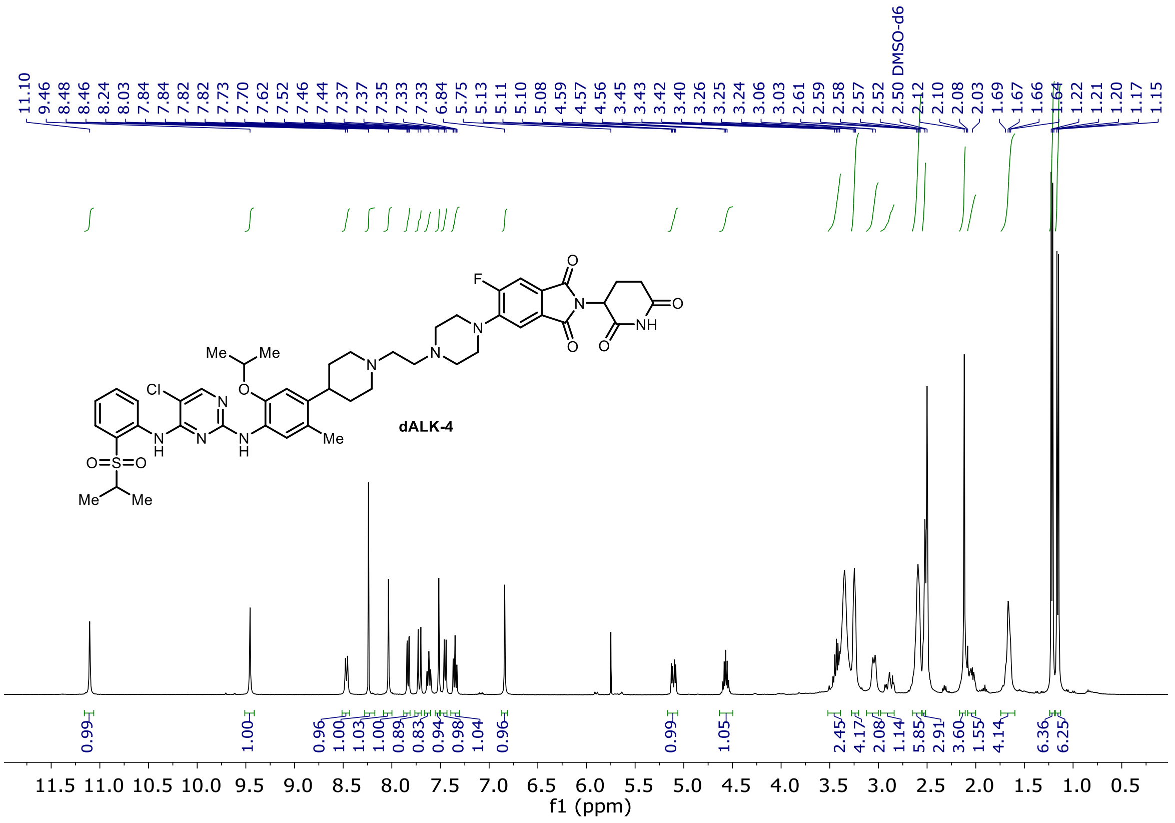

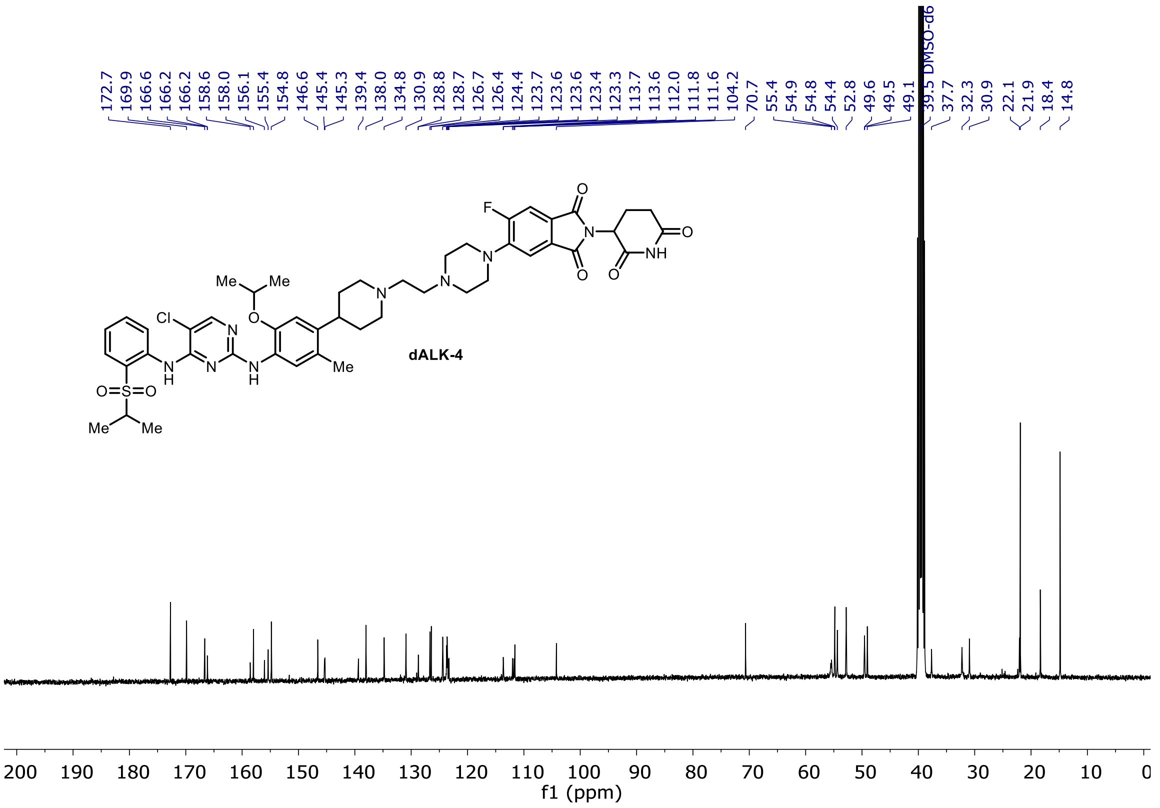

$^{19}$F NMR of **dALK-4**

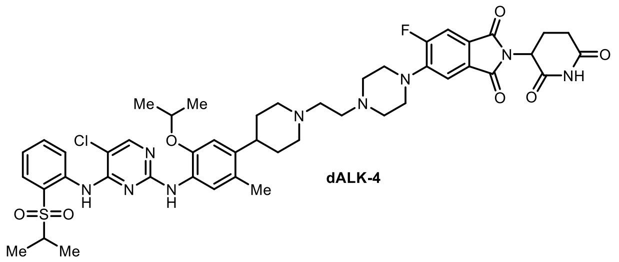

dALK-4

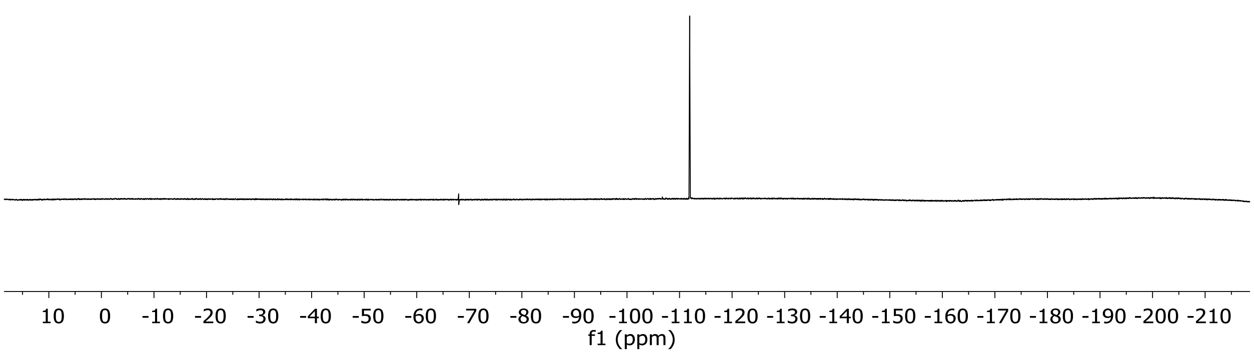

$^1H$ and $^{13}C$ NMR of **dALK-5**

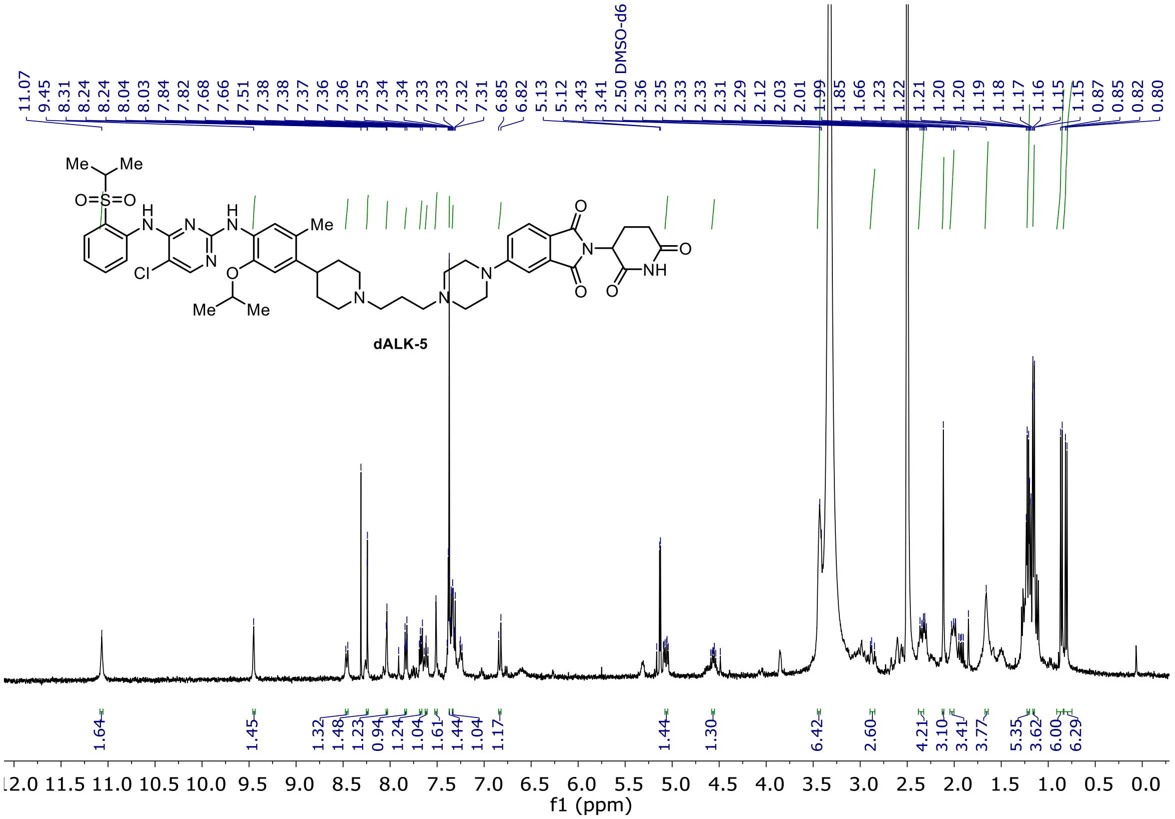

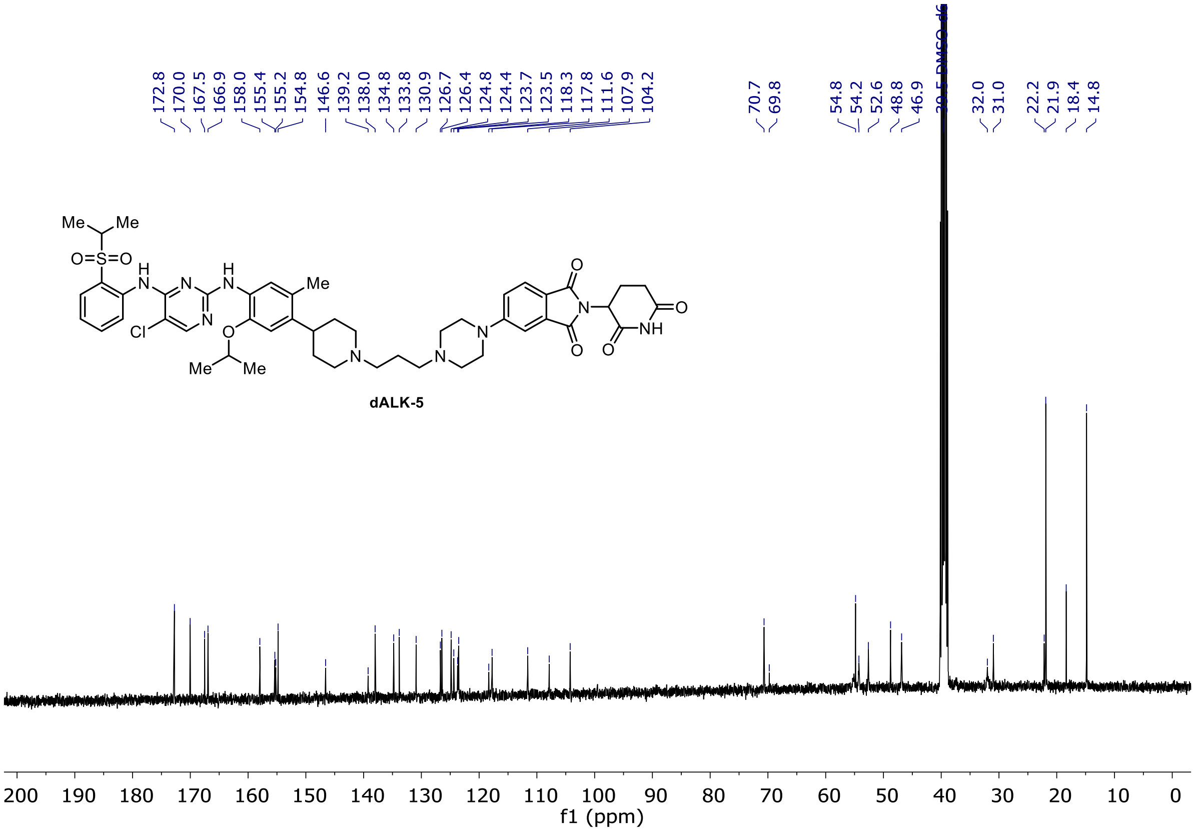

$^{1}H$ and $^{13}C$ NMR of **dALK-6**

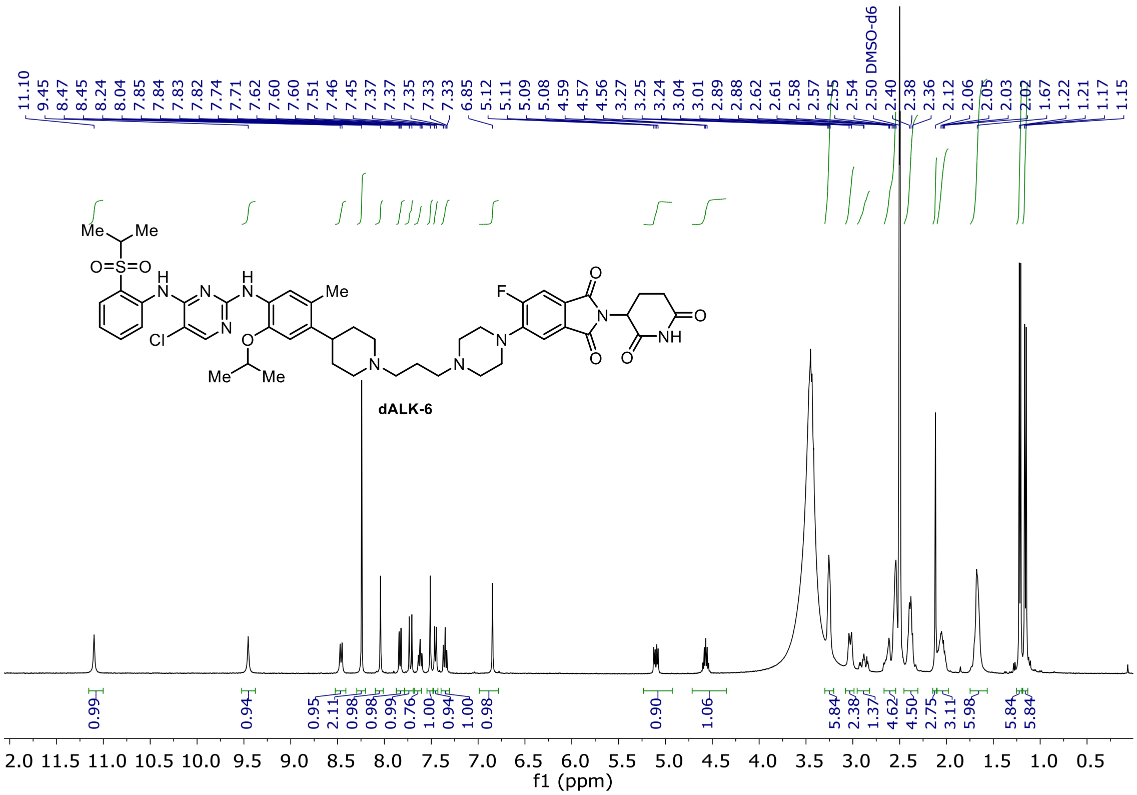

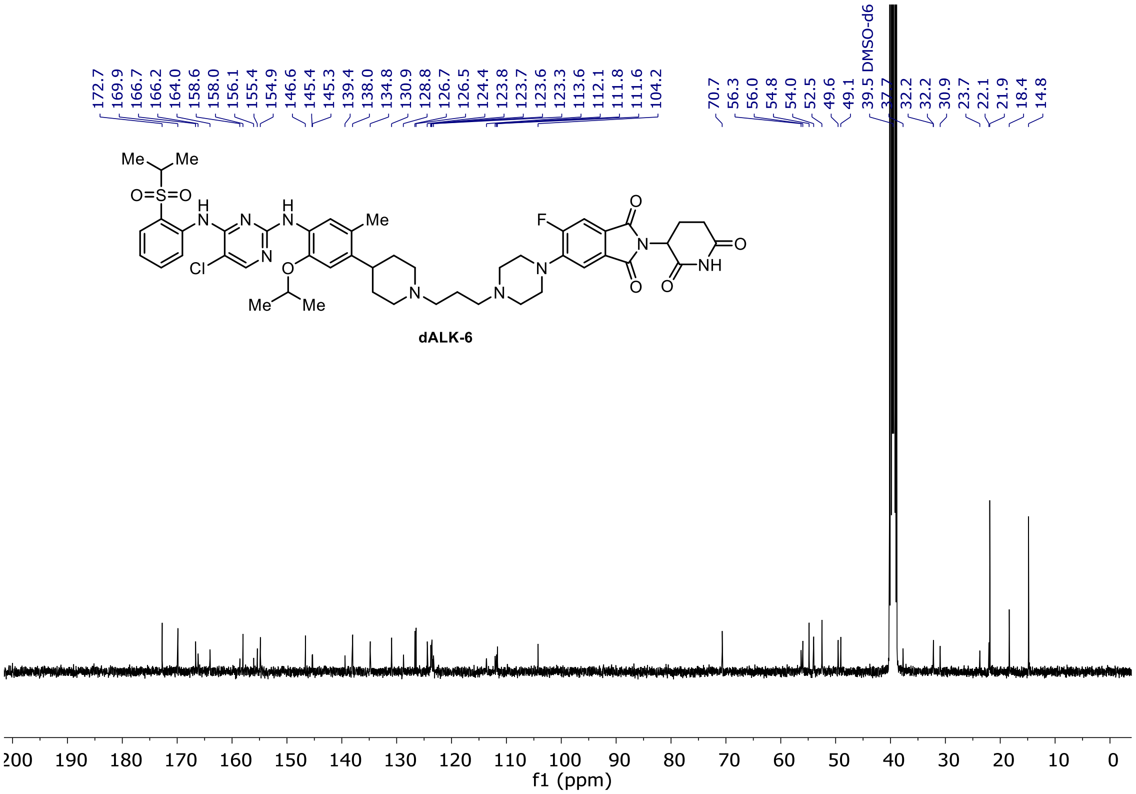

$^{19}$F NMR of **dALK-6**

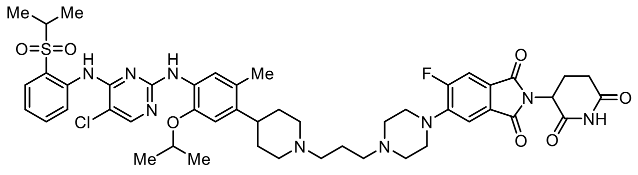

**dALK-6**

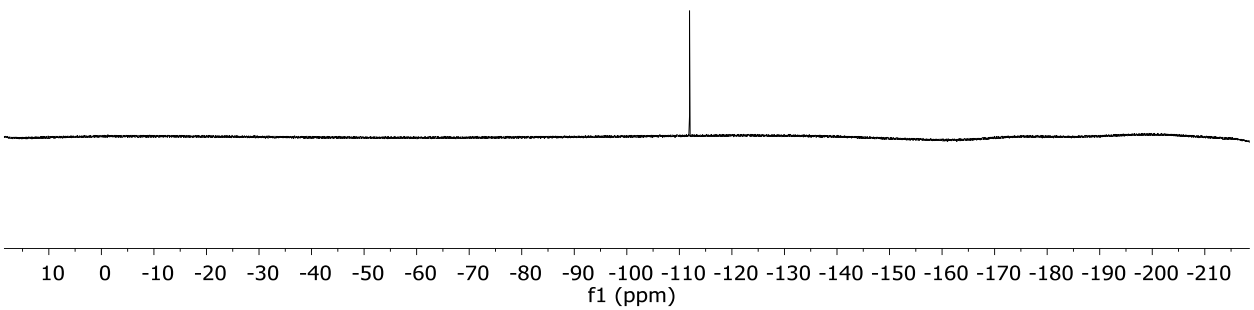

$^{1}$H and $^{13}$C NMR of **dALK-7**

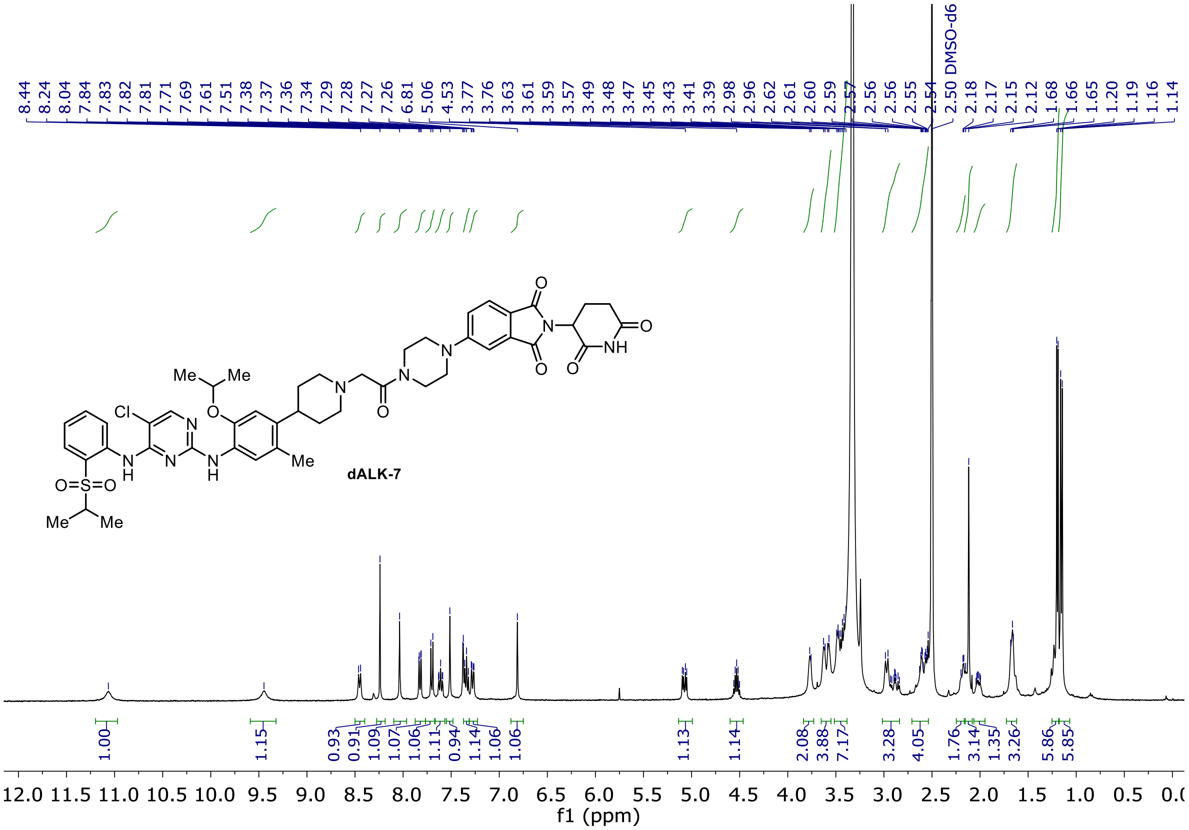

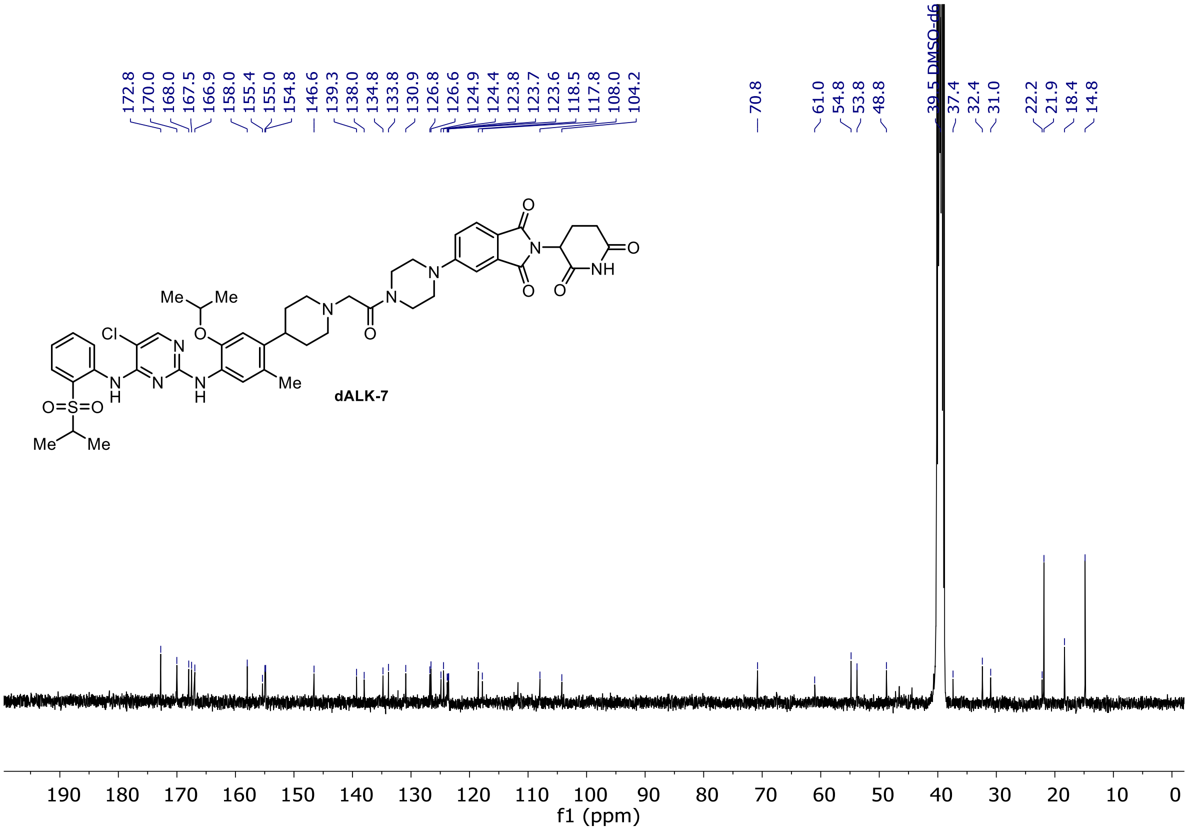

[1]H and [13]C NMR of **dALK-8**

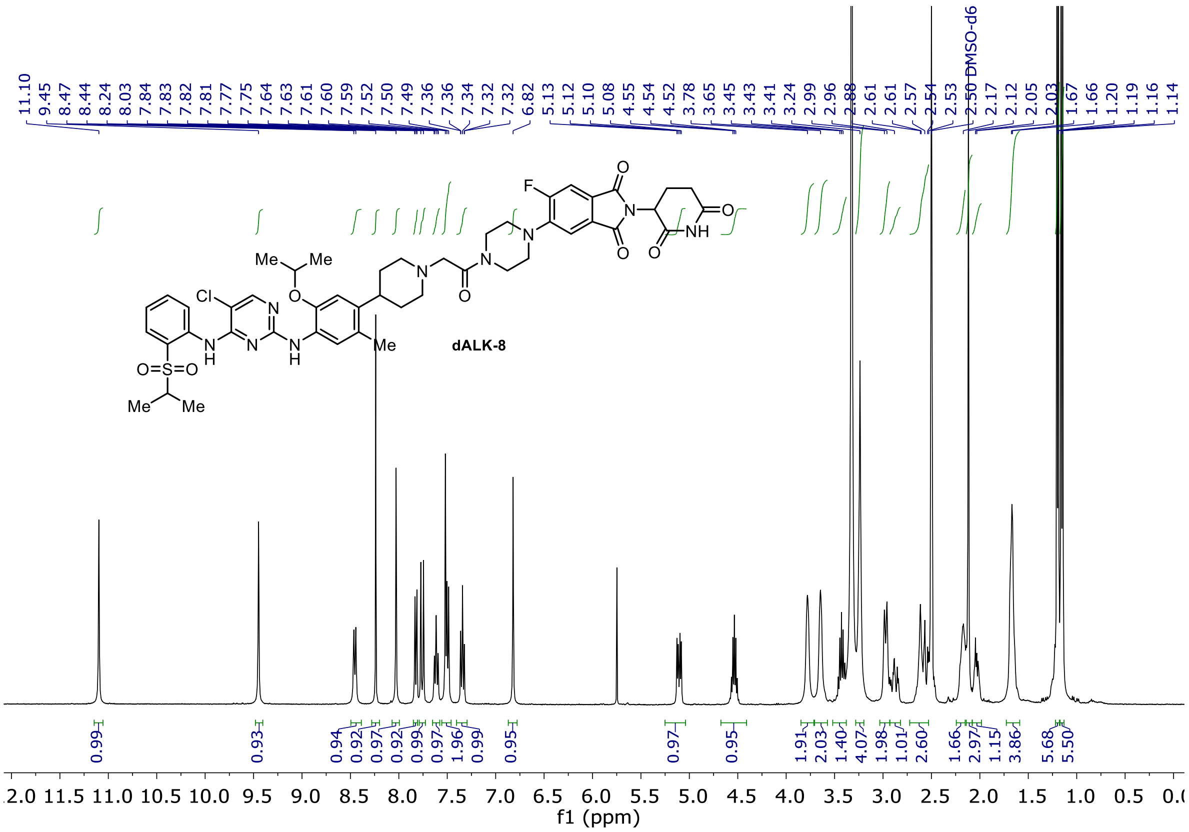

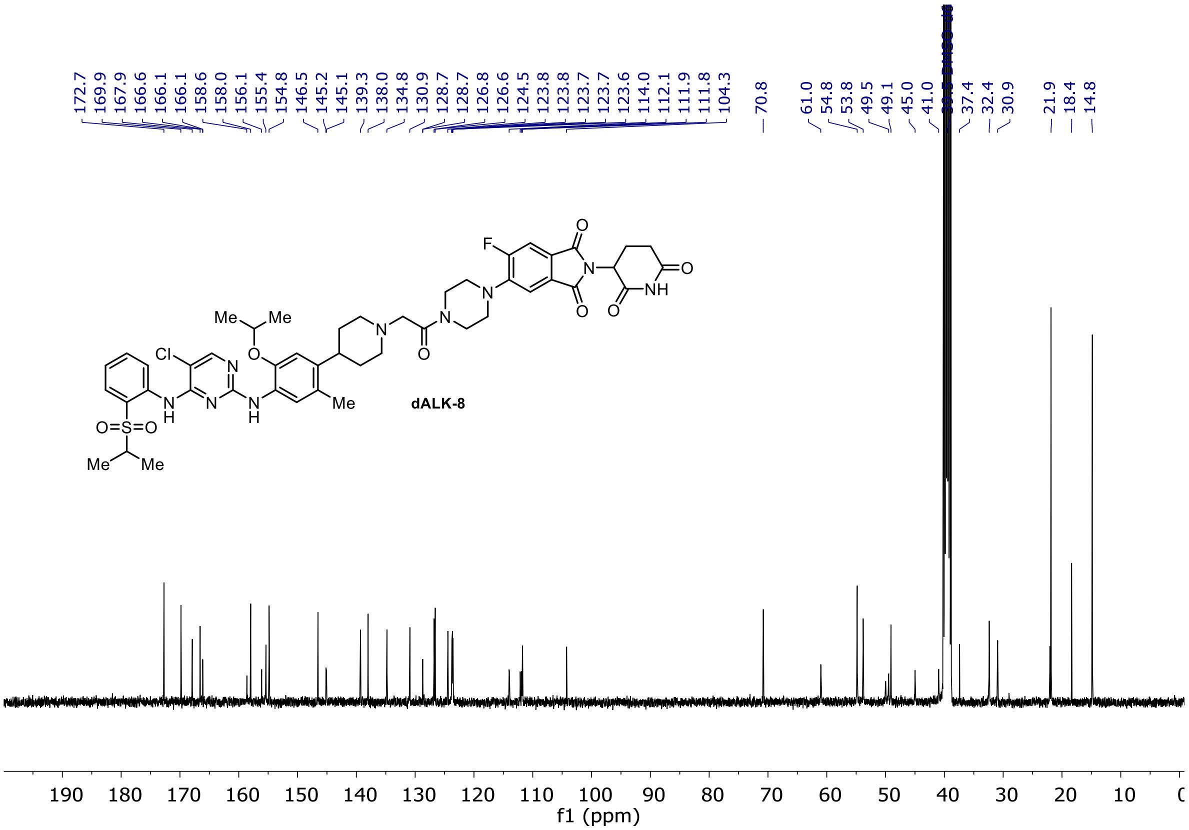

$^{19}$F NMR of **dALK-8**

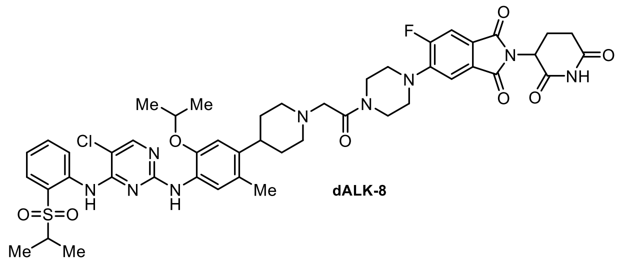

dALK-8

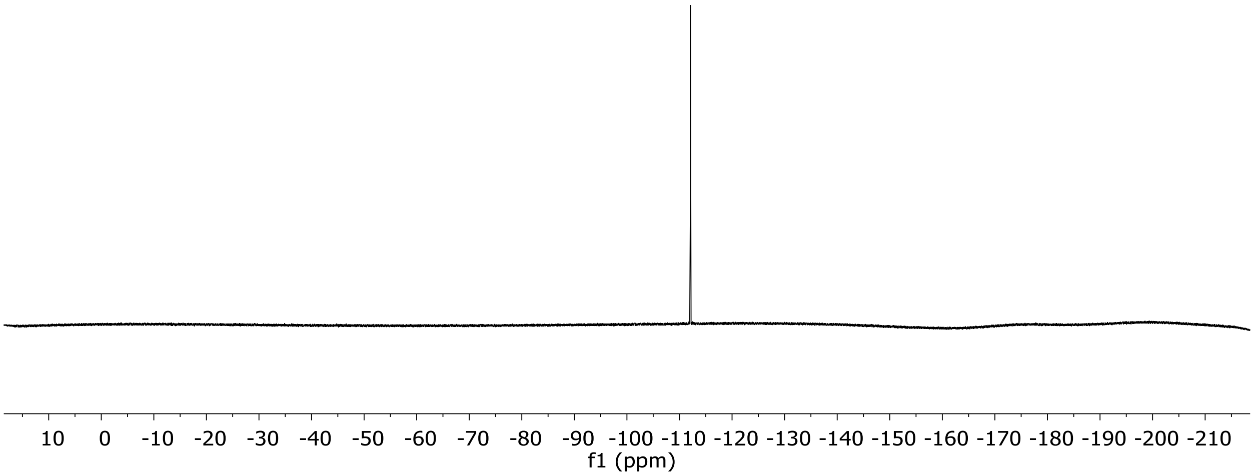

#### $^{1}H$ and $^{13}C$ NMR of **dALK-9**

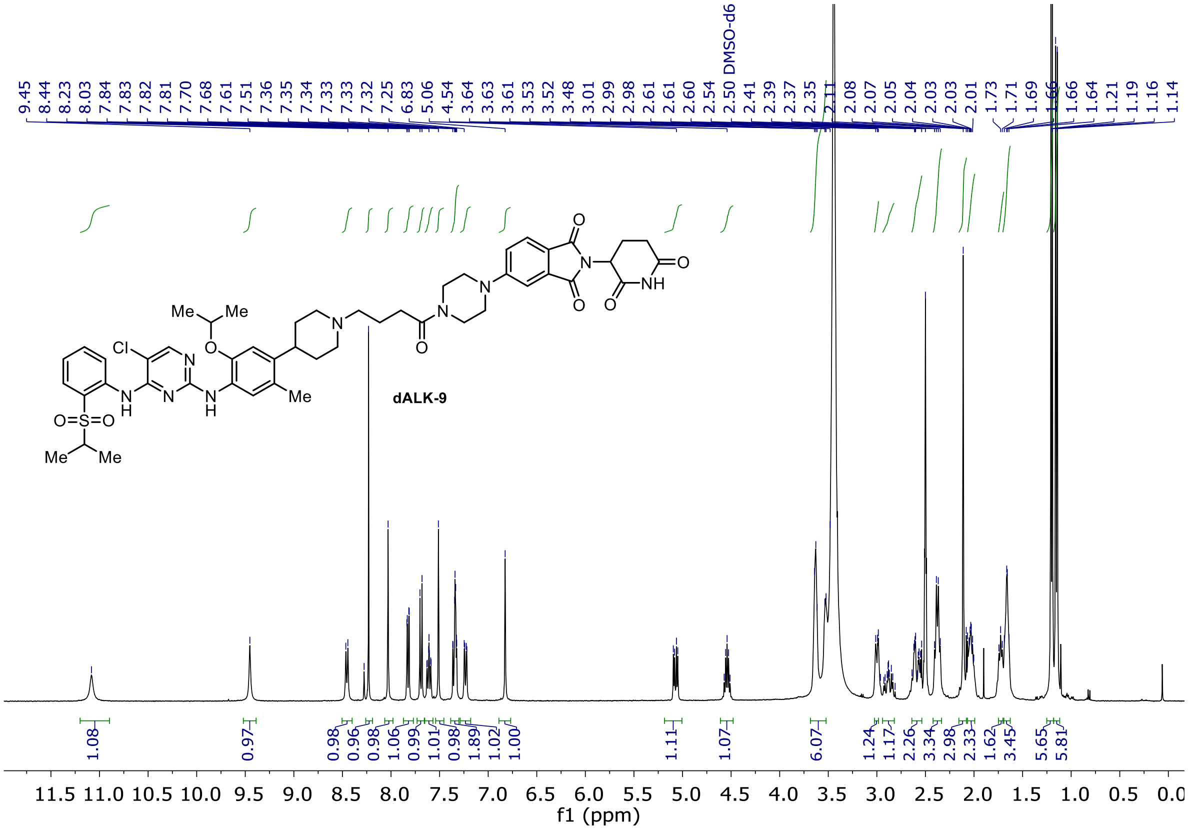

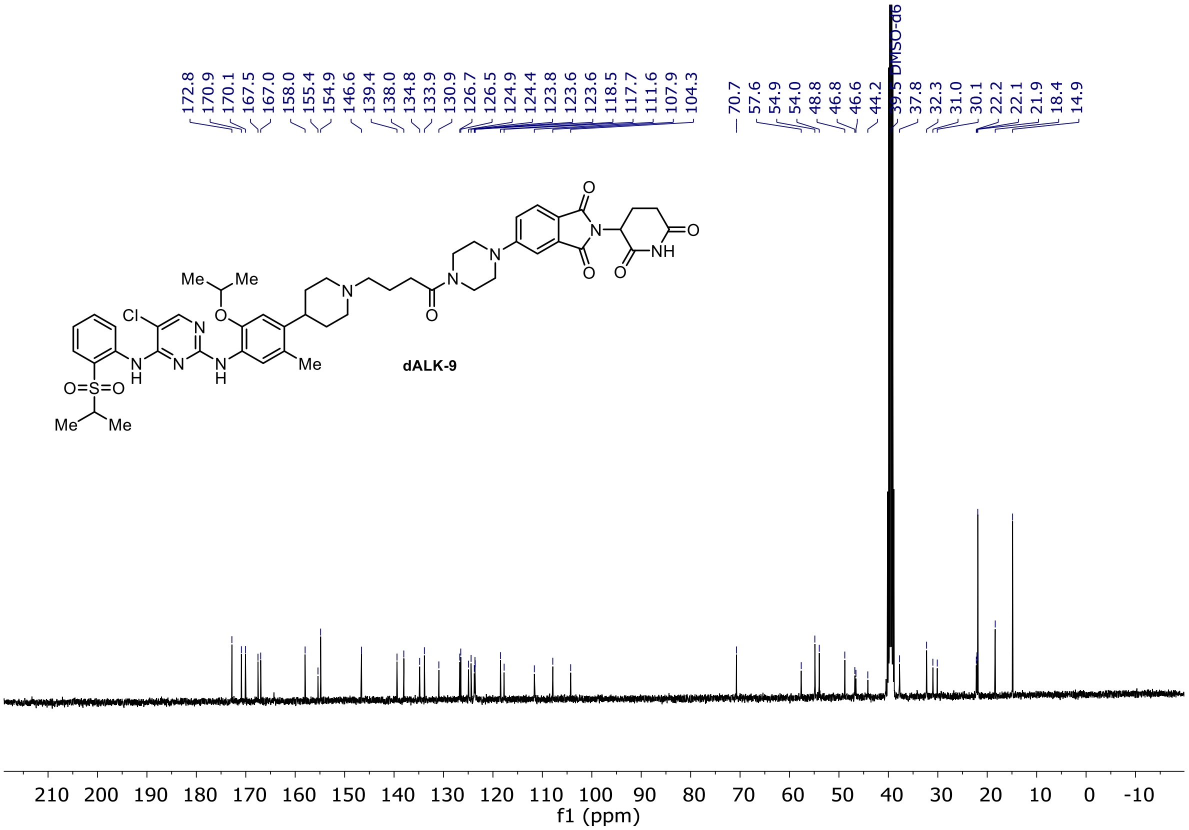

[1]H and [13]C NMR of **dALK-10**

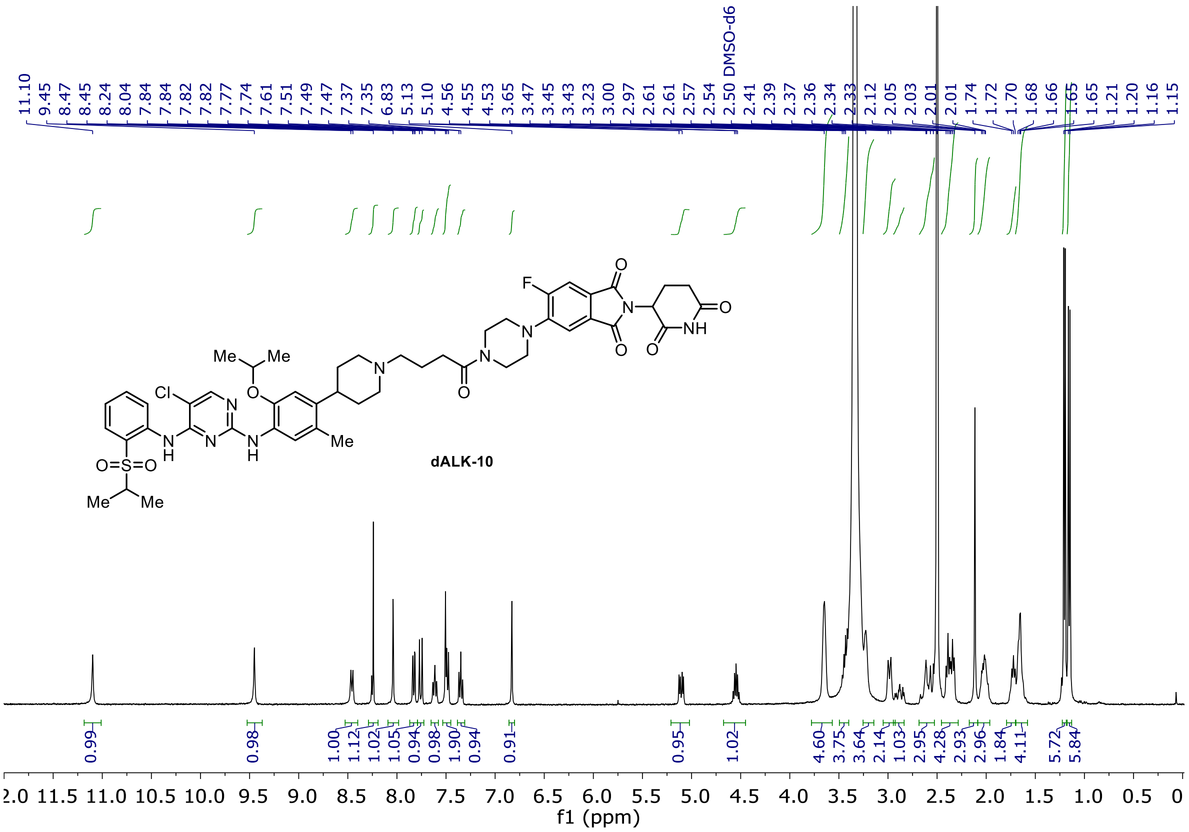

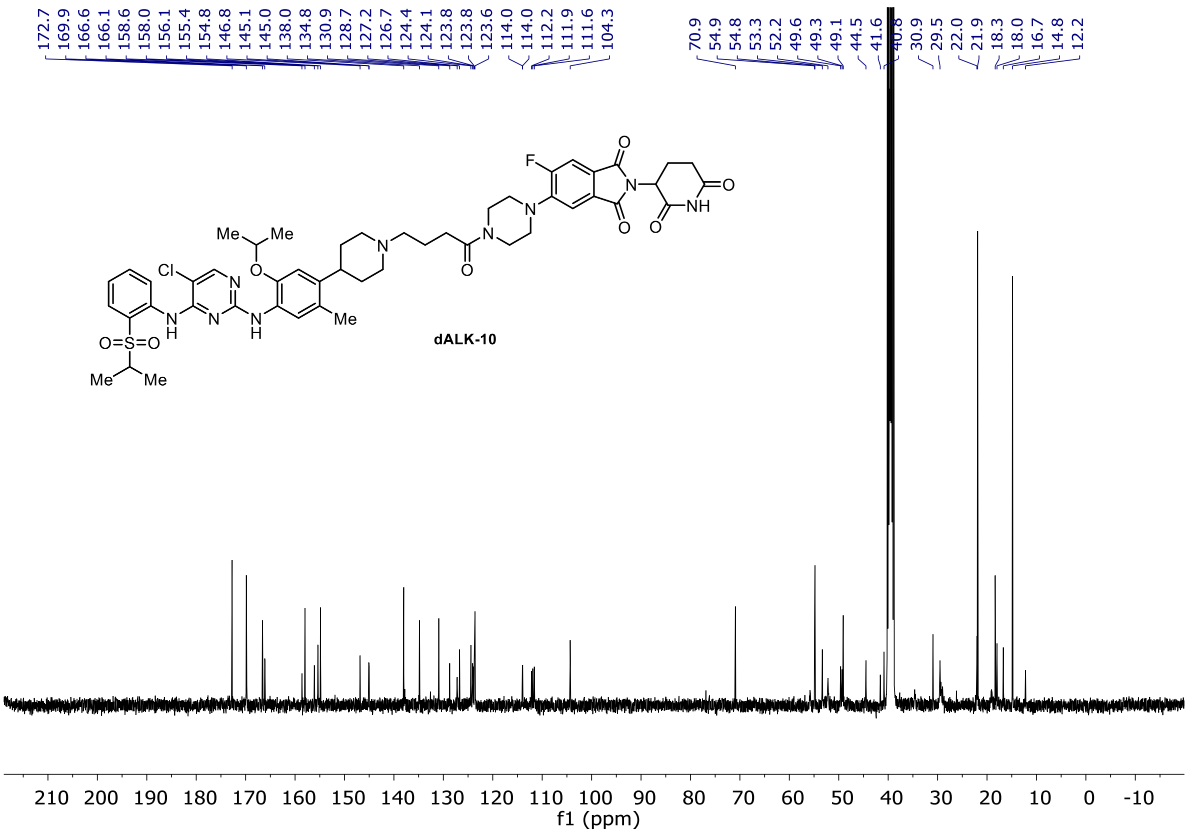

$^{19}$F NMR of **dALK-10**

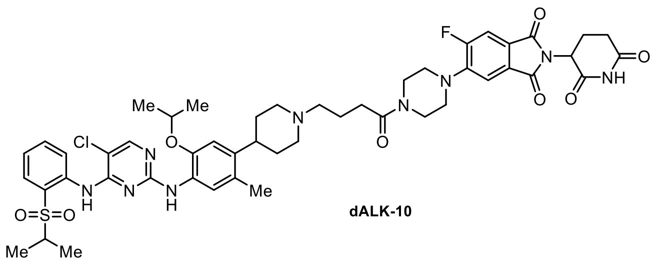

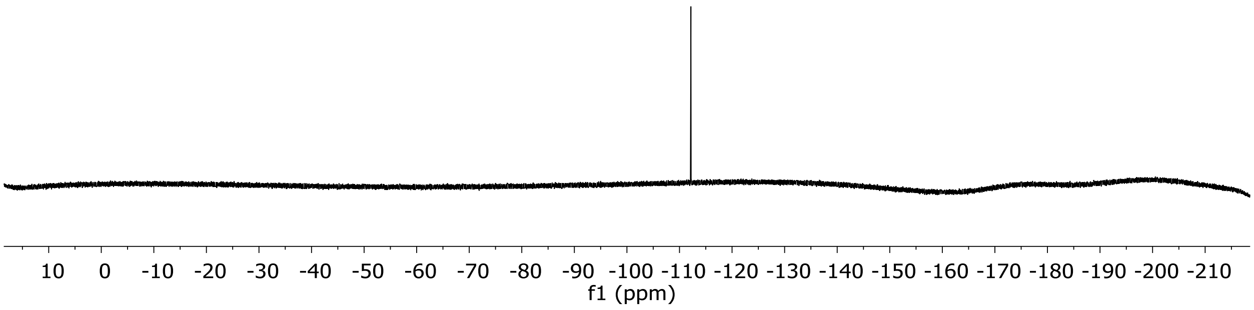

[1]H and [13]C NMR of **dALK-11**

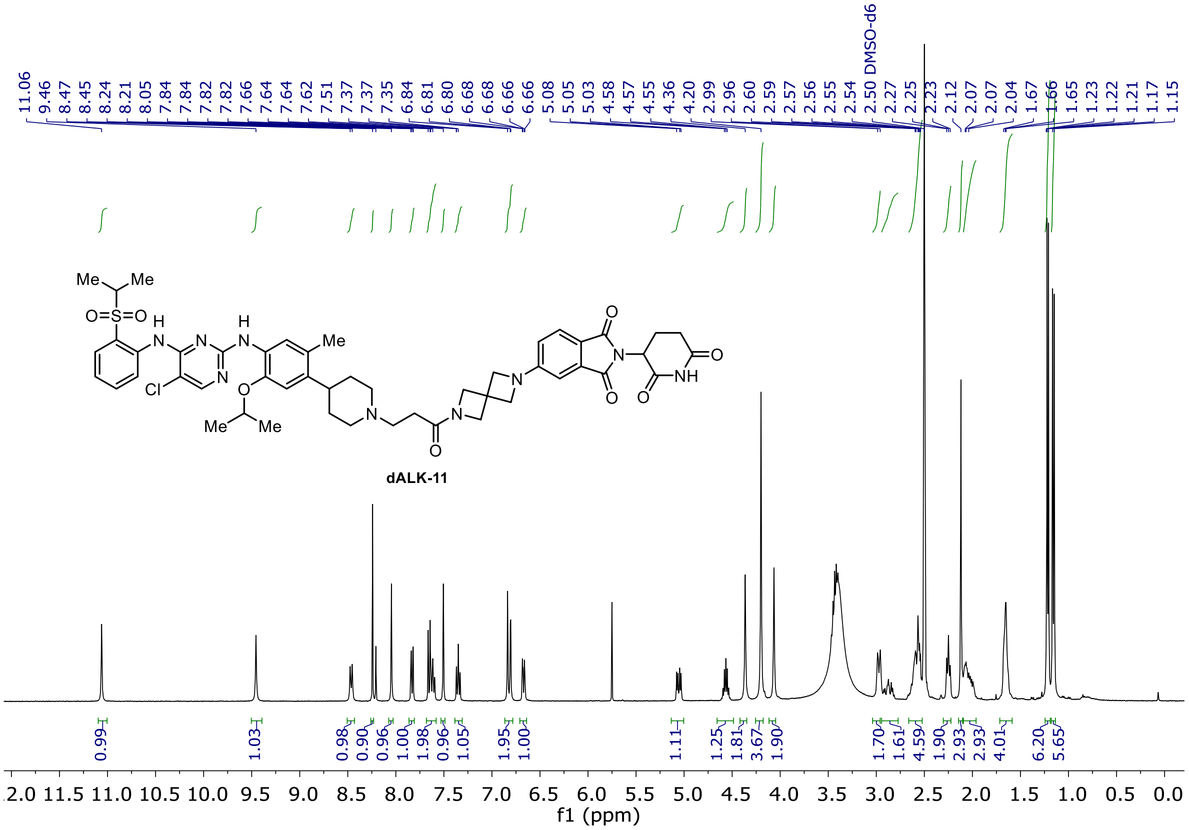

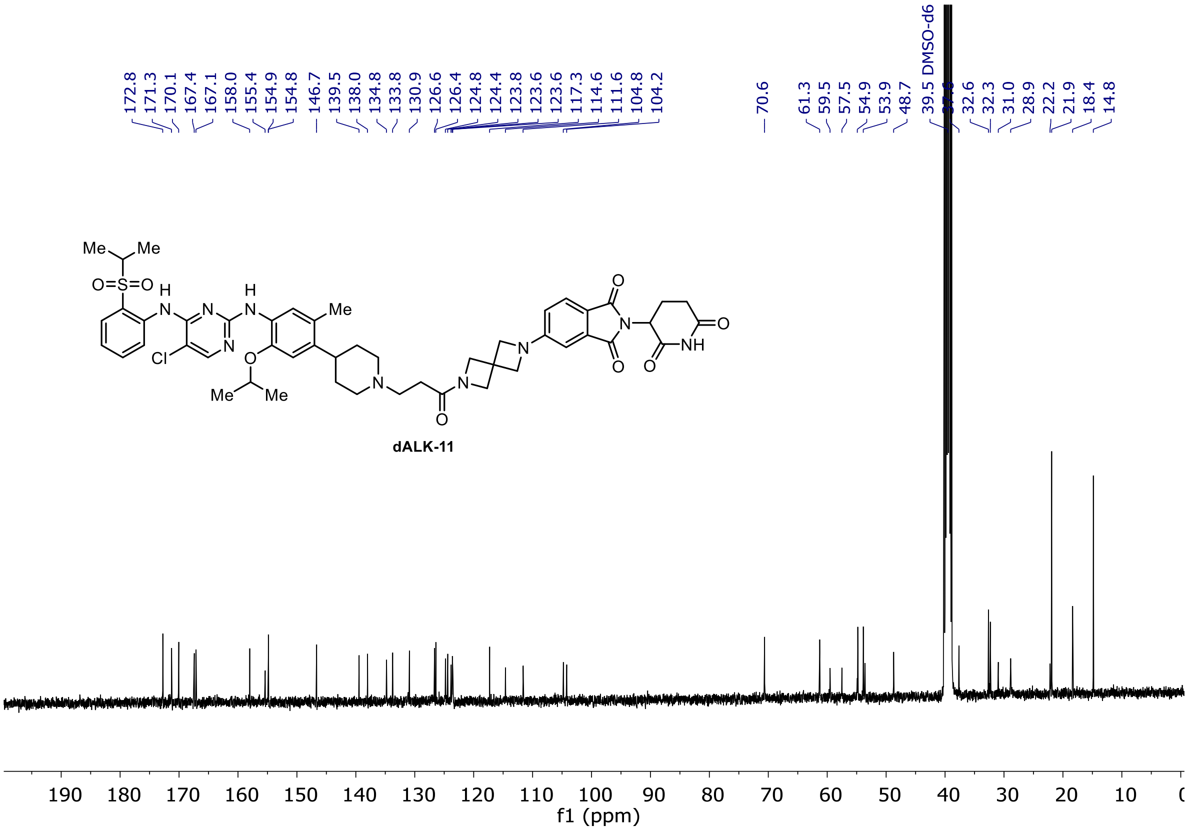

[1]H and [13]C NMR of **dALK-12**

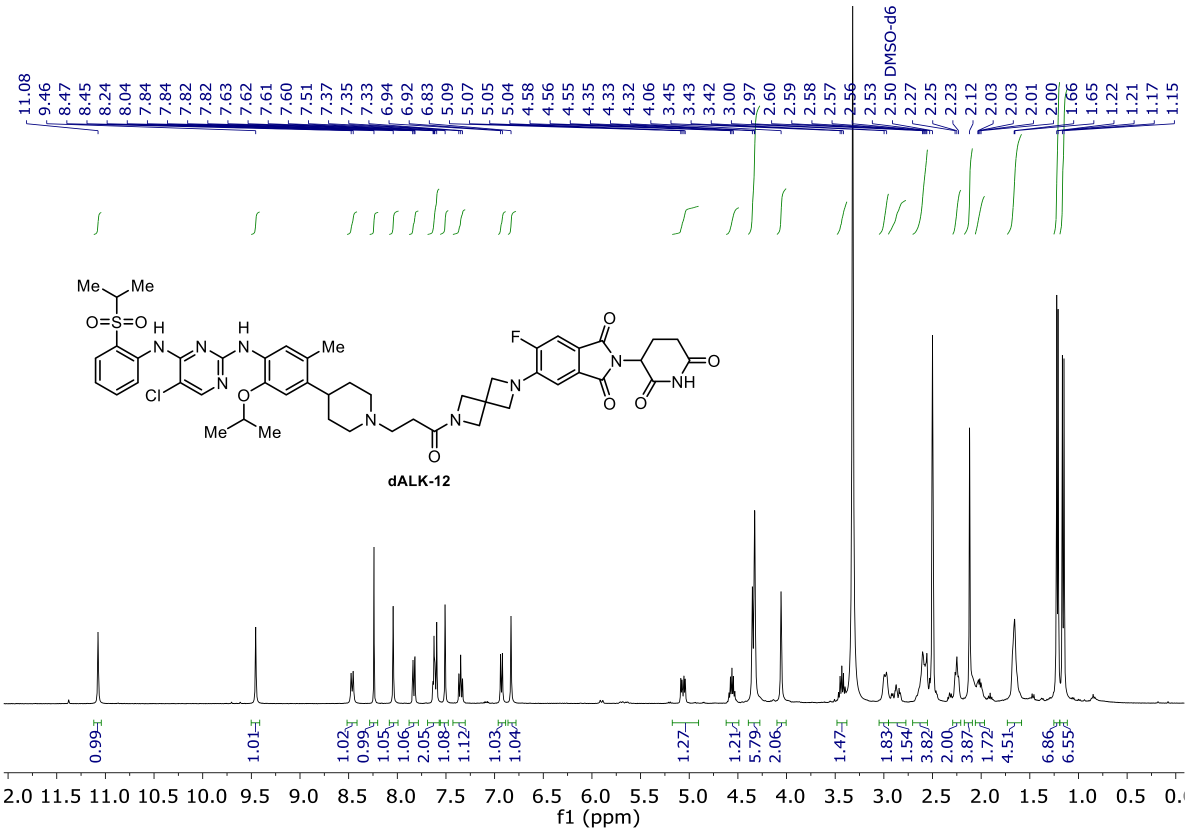

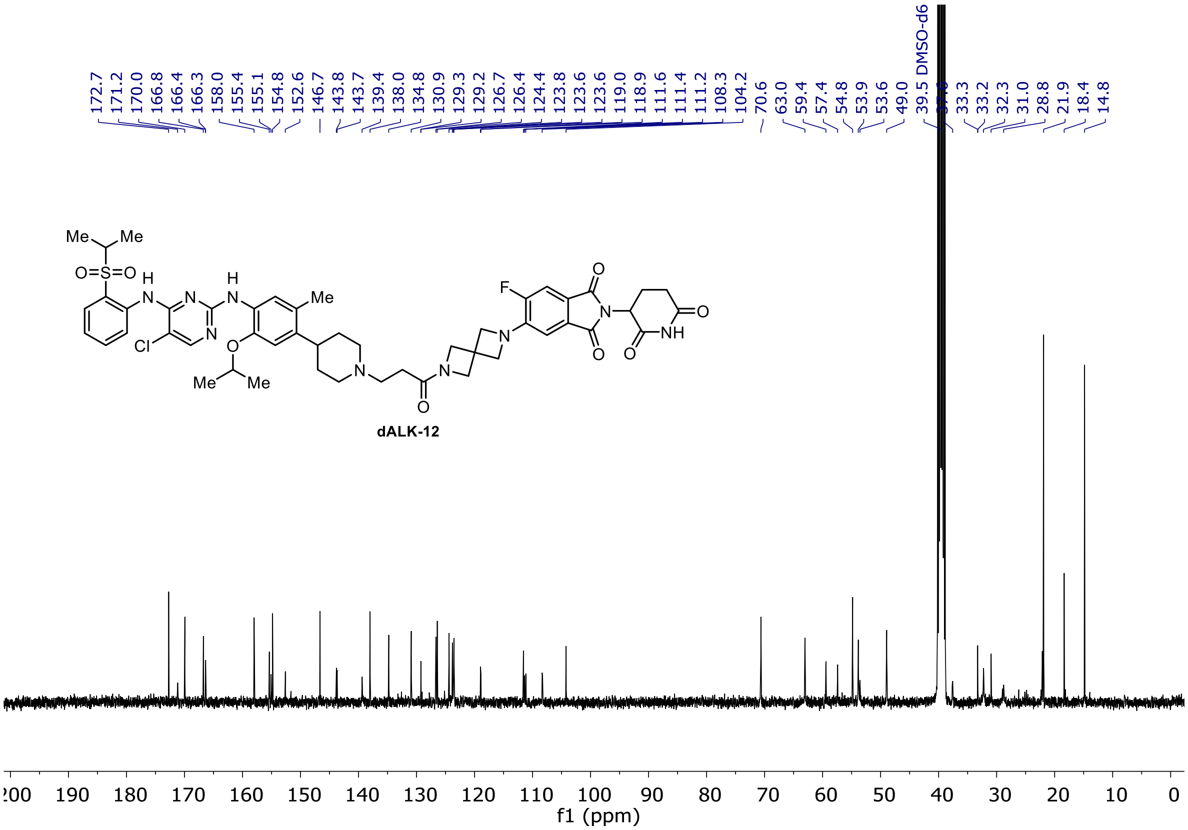

$^{19}F$ NMR of **dALK-12**

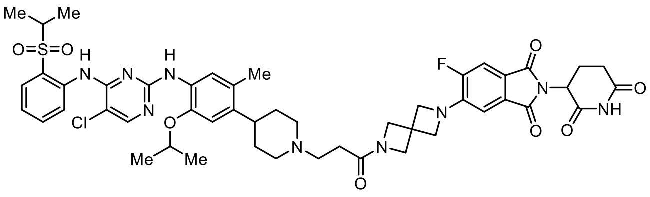

**dALK-12**

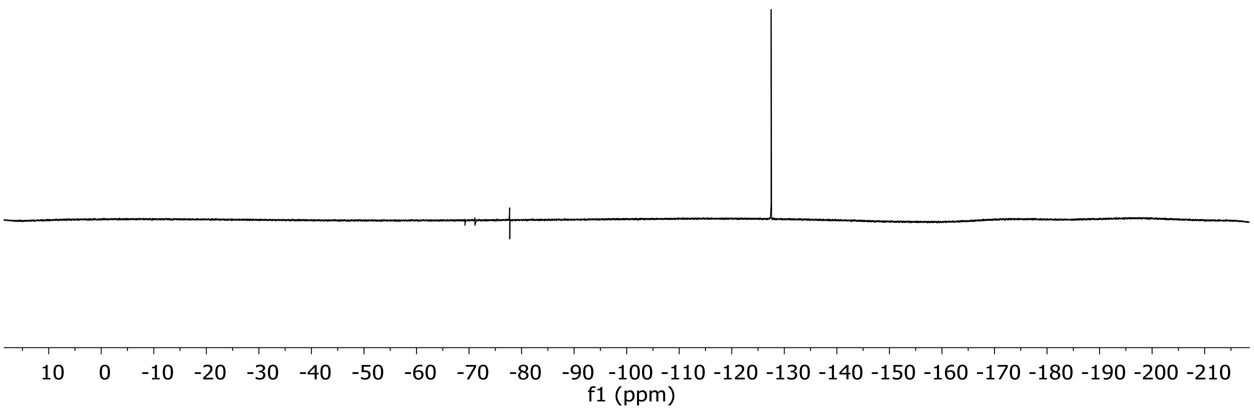

[1]H and [13]C NMR of 5-chloro-N2-(4-(1-(hex-5-yn-1-yl)piperidin-4-yl)-2-isopropoxy-5-methylphenyl)-N4-(2-(isopropylsulfonyl)phenyl)pyrimidine-2,4-diamine (**82**)

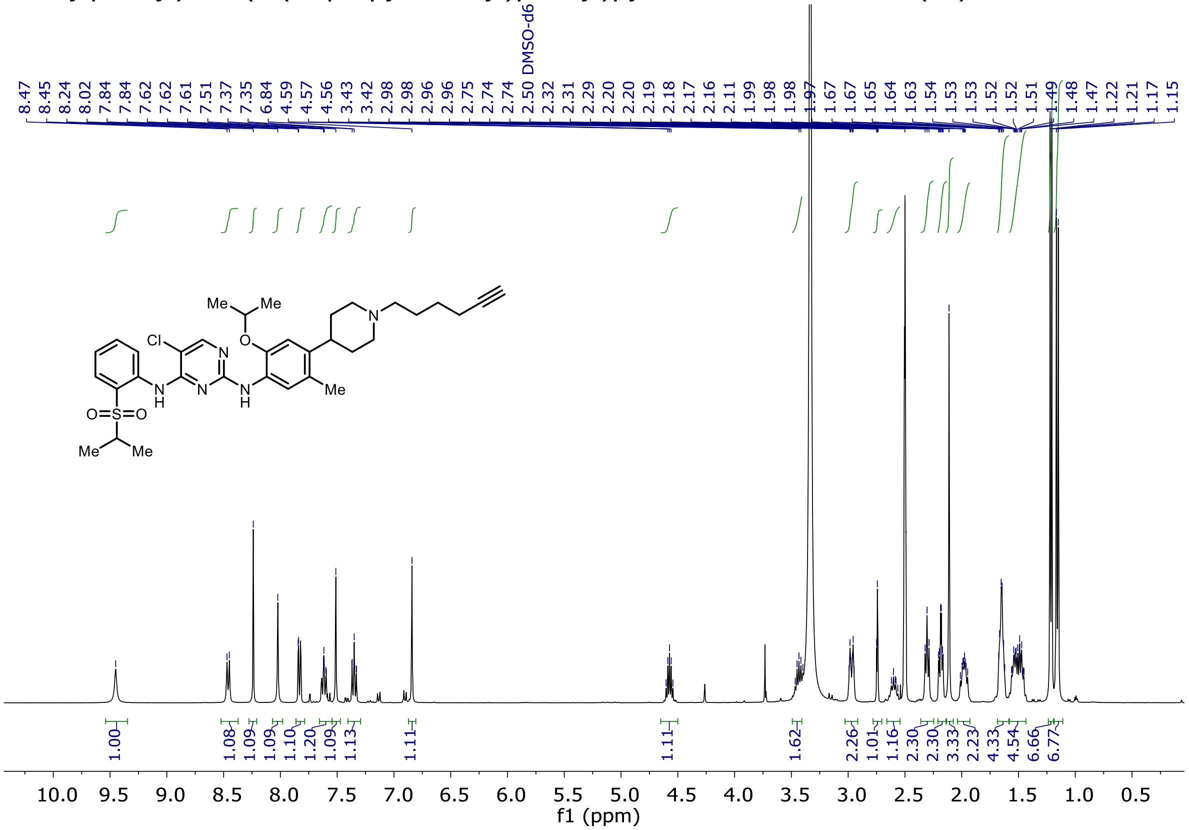

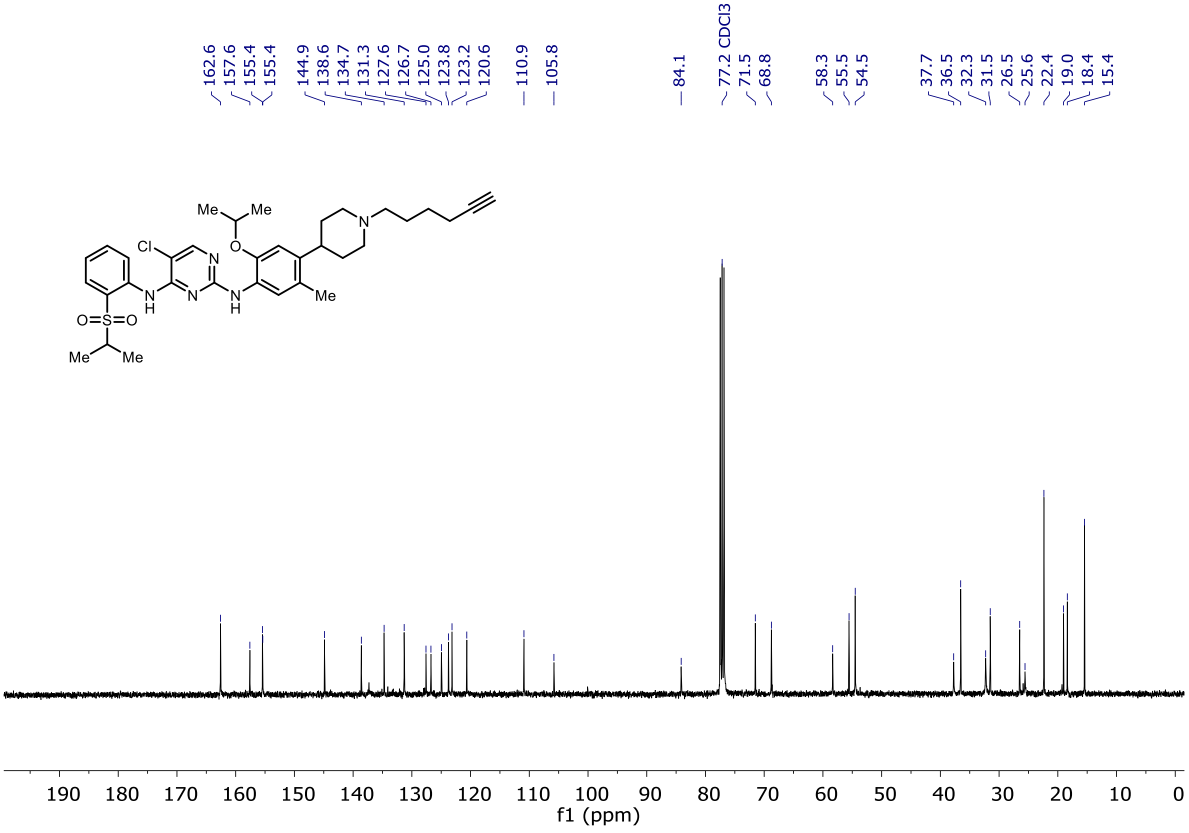

[1]H and [13]C NMR of 2-(4-(4-((5-chloro-4-((2-(isopropylsulfonyl)phenyl)amino)pyrimidin-2-yl)amino)-5-isopropoxy-2-methylphenyl)piperidin-1-yl)-*N*-methoxy-*N*-methylacetamide (**83**)

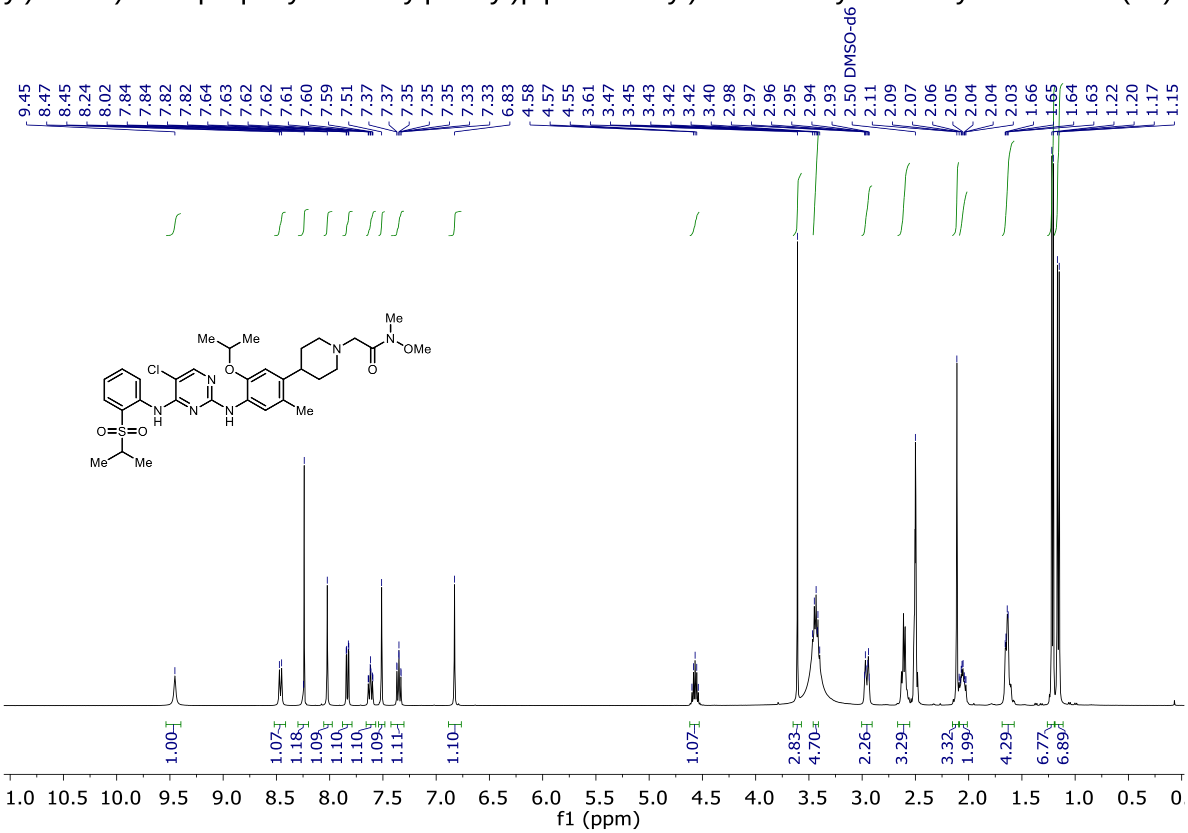

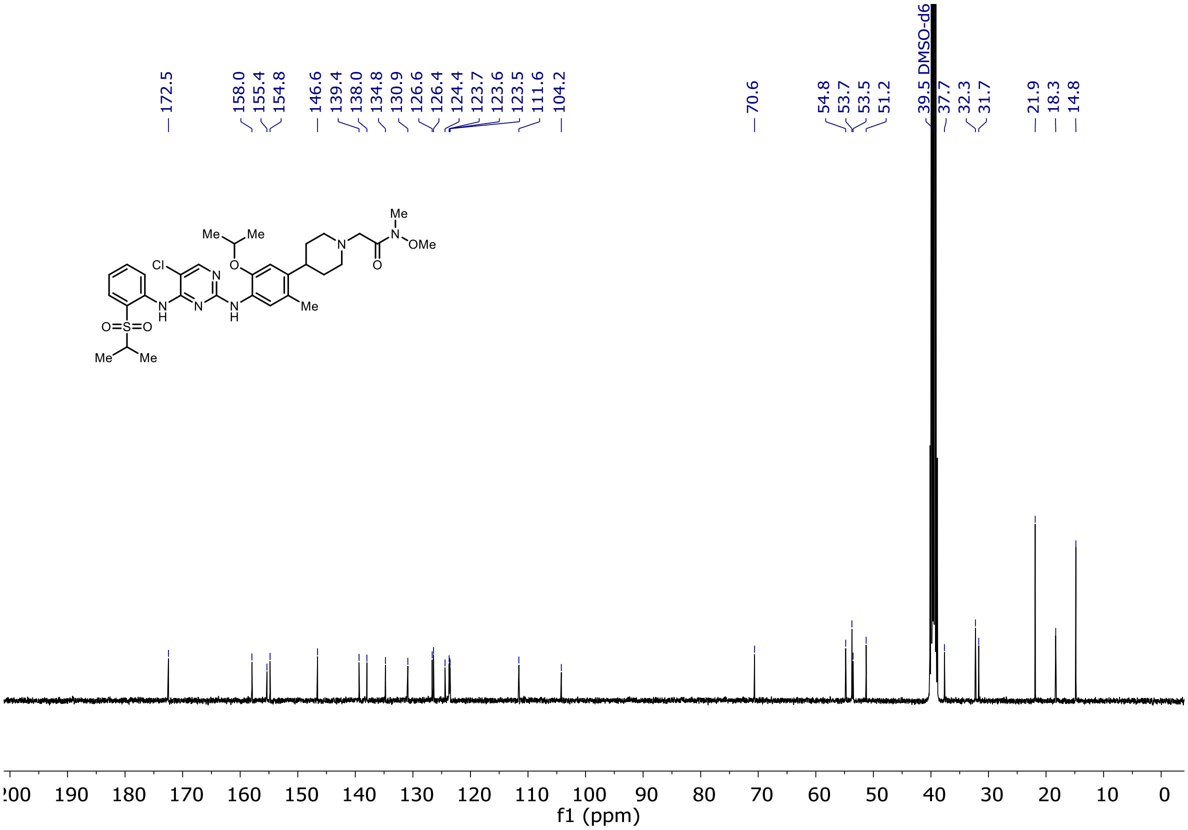

$^1H$ and $^{13}C$ NMR of 3-(4-(4-((5-chloro-4-((2-(isopropylsulfonyl)phenyl)amino)pyrimidin-2-yl)amino)-5-isopropoxy-2-methylphenyl)piperidin-1-yl)-*N*-methoxy-*N*-methylpropanamide (**86**)

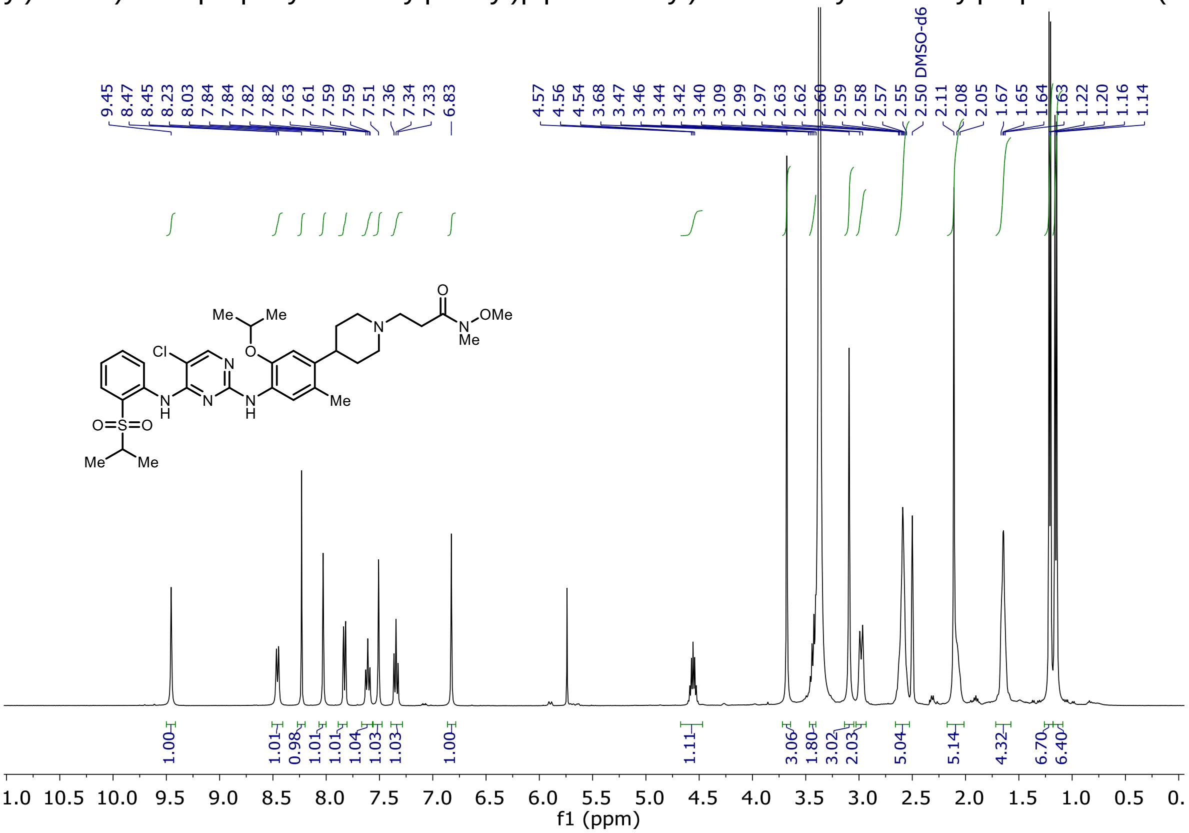

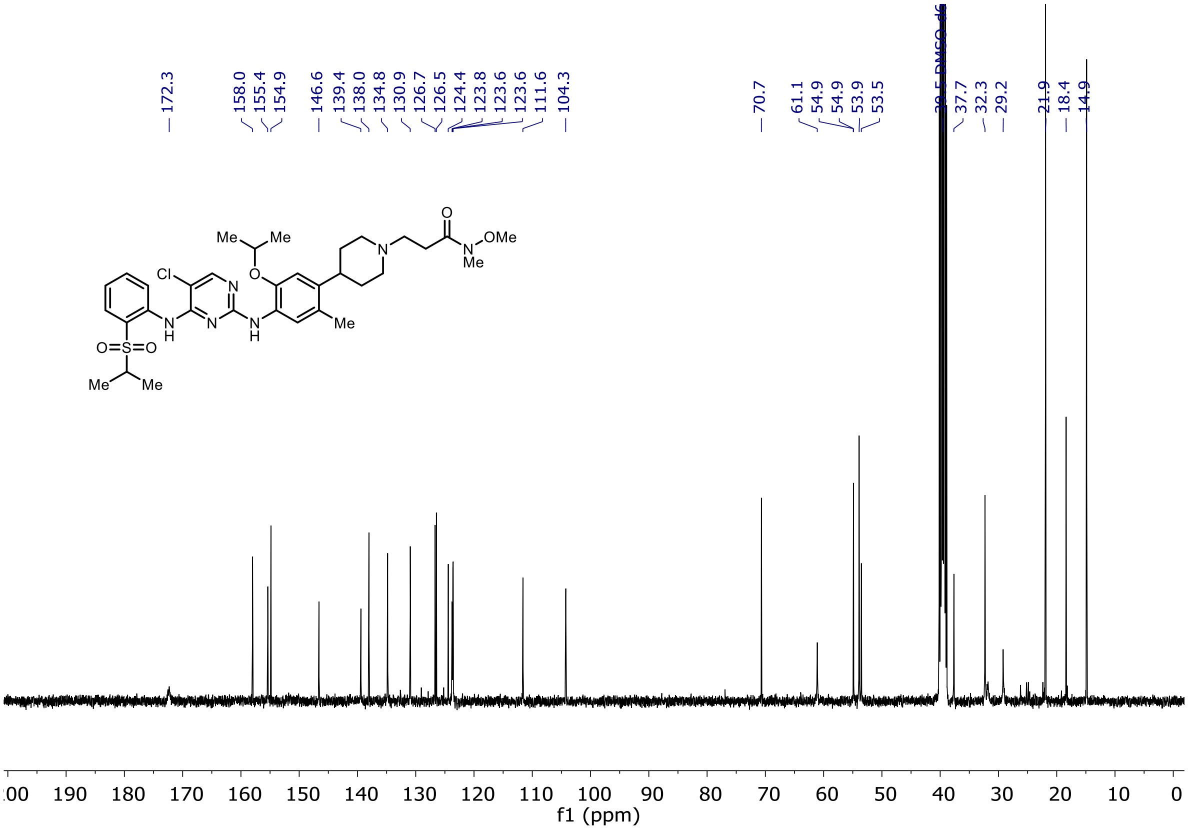

$^{1}H$ and $^{13}C$ NMR of *tert*-butyl 4-(4-(4-((5-chloro-4-((2-(isopropylsulfonyl)phenyl)amino)pyrimidin-2-yl)amino)-5-isopropoxy-2-methylphenyl)piperidin-1-yl)butanoate (**91**)

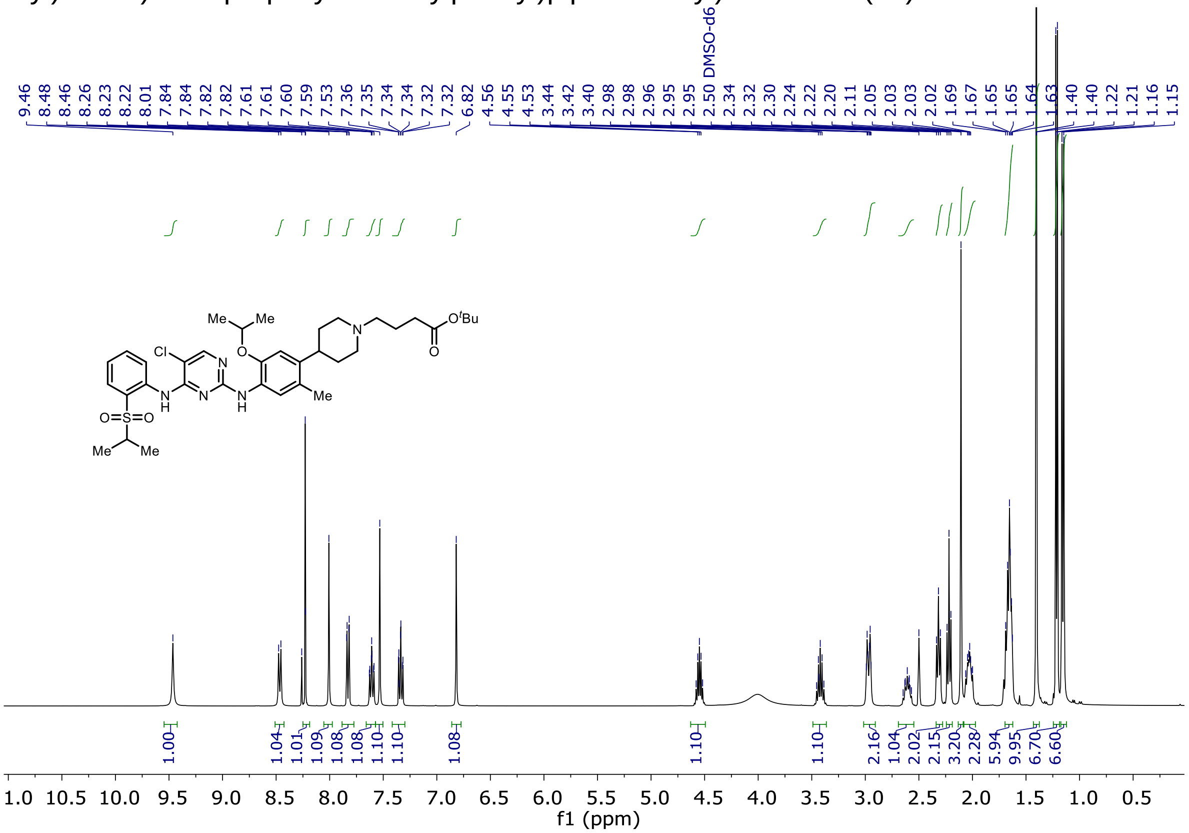

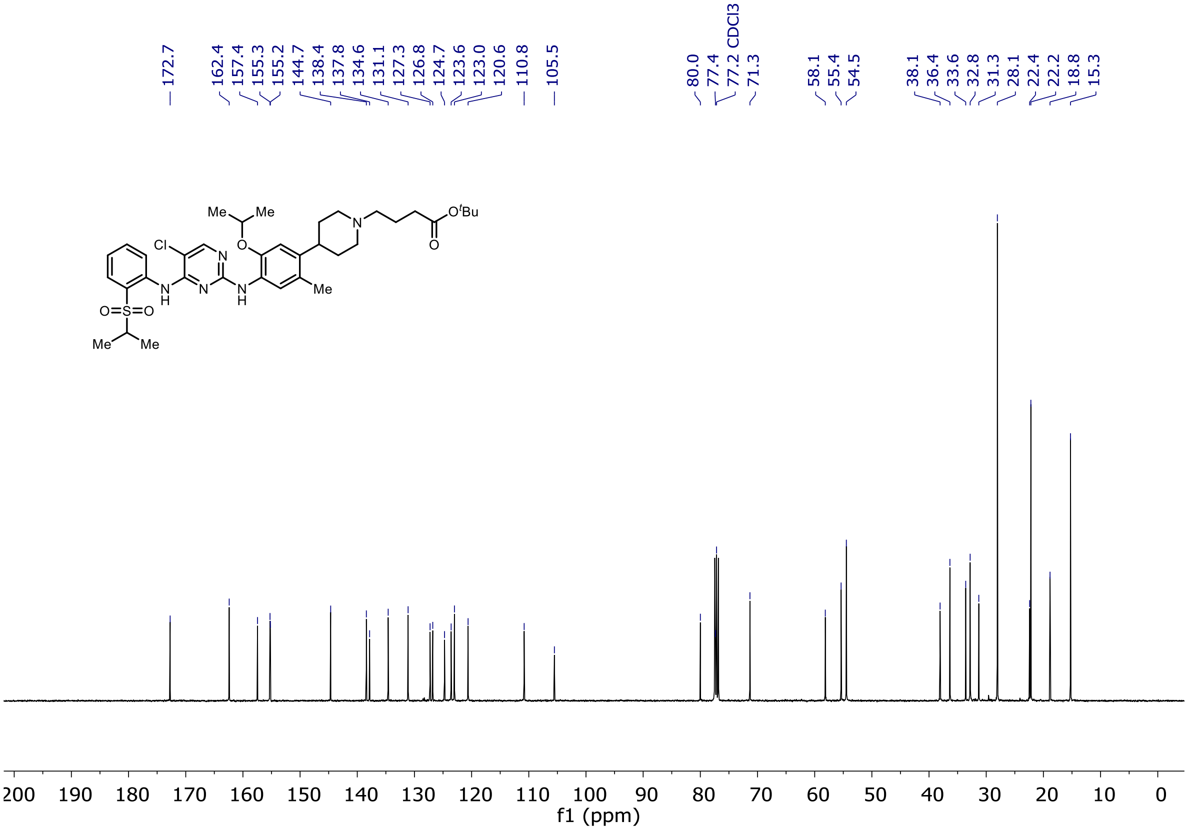

$^{1}H$ and $^{13}C$ NMR of methyl 3-(4-(4-((5-chloro-4-((2-(isopropylsulfonyl)phenyl)amino)pyrimidin-2-yl)amino)-5-isopropoxy-2-methylphenyl)piperidin-1-yl)propanoate (**85**)

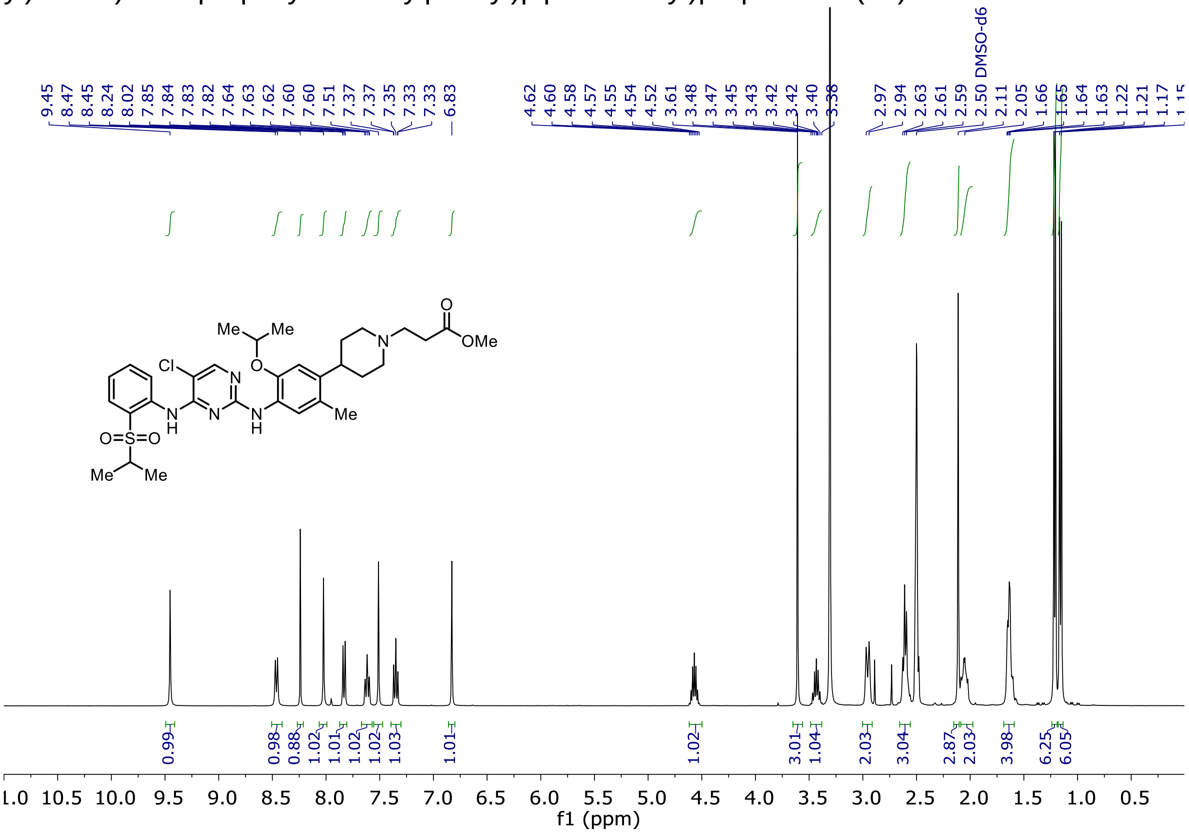

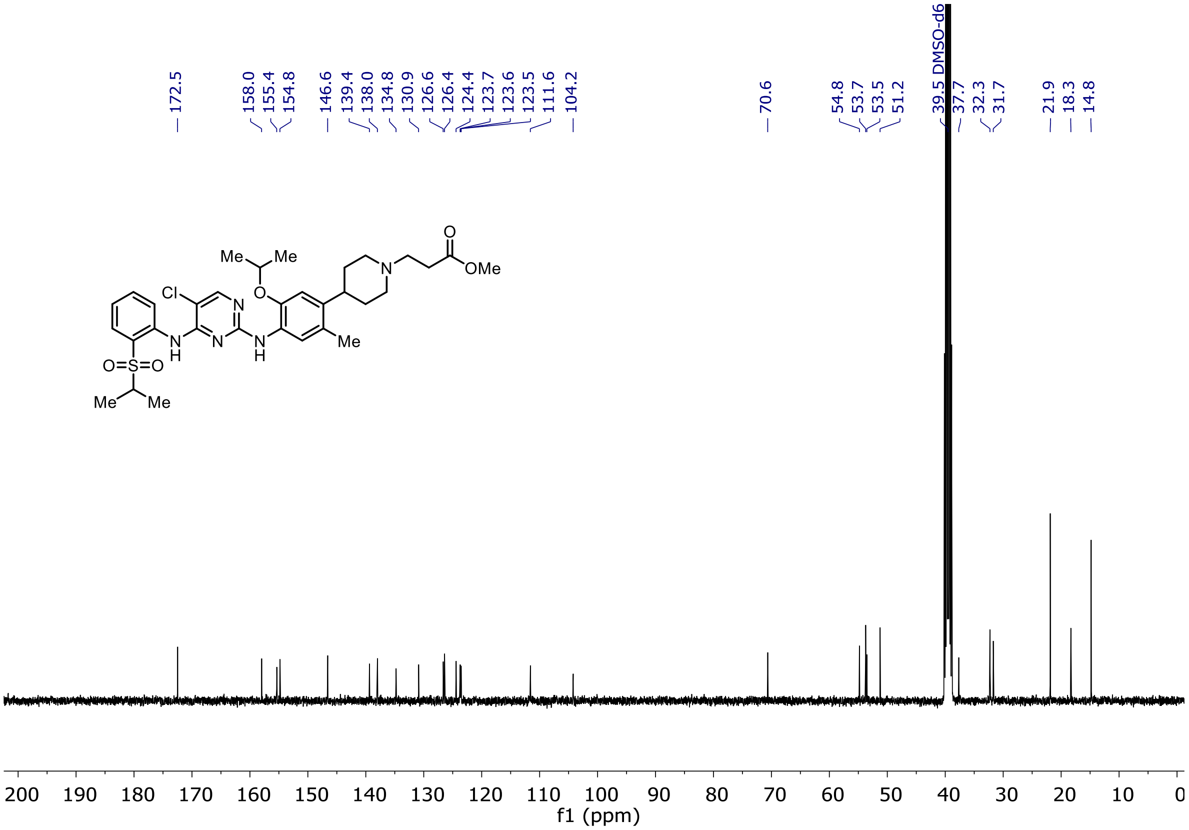

[1]H and [13]C NMR of 3-(4-(4-((5-chloro-4-((2-(isopropylsulfonyl)phenyl)amino)pyrimidin-2-yl)amino)-5-isopropoxy-2-methylphenyl)piperidin-1-yl)propanal (**87**)

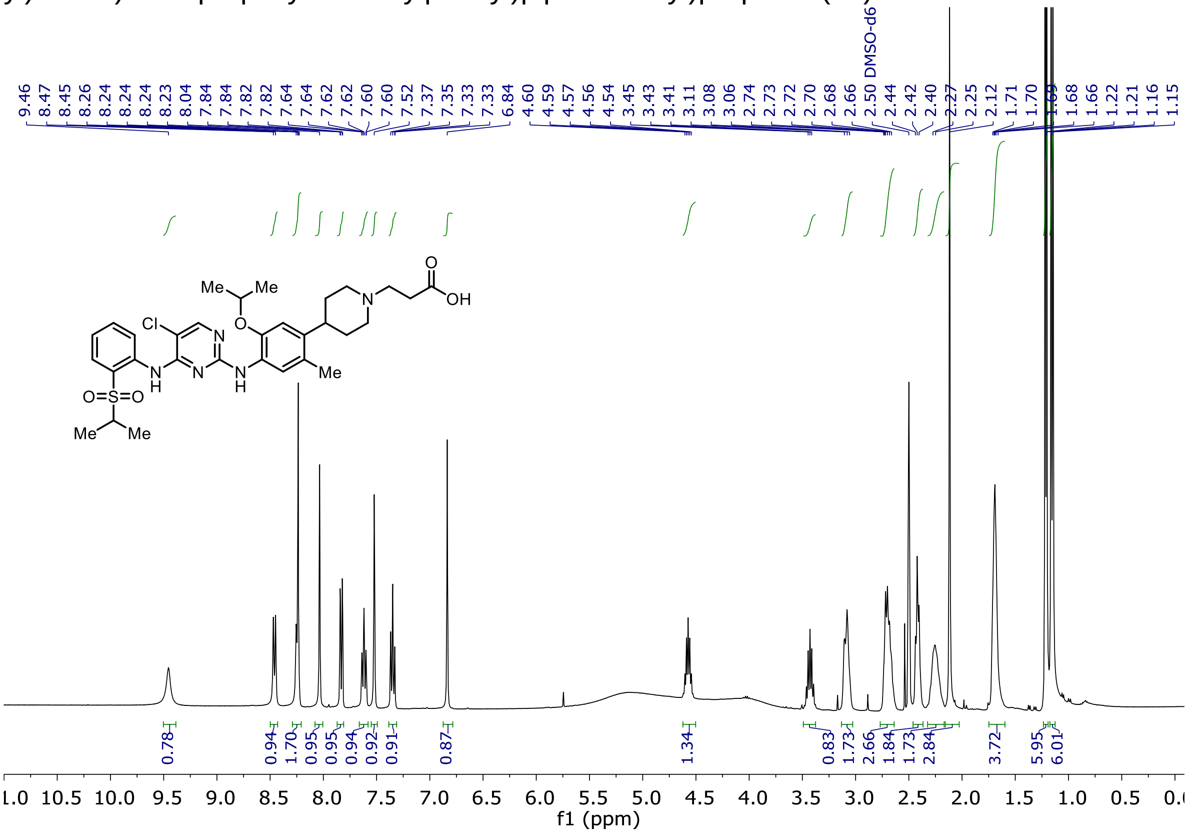

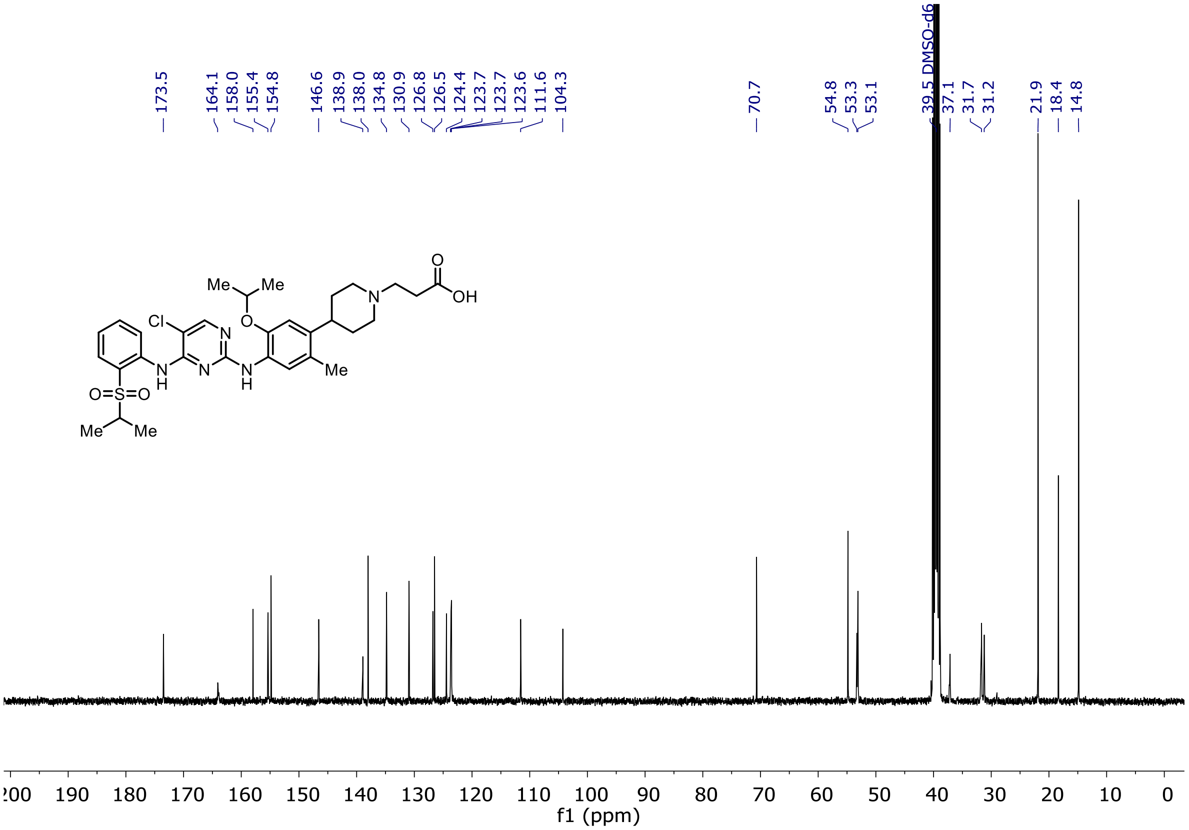

[1]H and [13]C NMR of *tert*-butyl (2-((2-(2,6-dioxopiperidin-3-yl)-1,3-dioxoisoindolin-5-yl)amino)ethyl)carbamate (**95**)

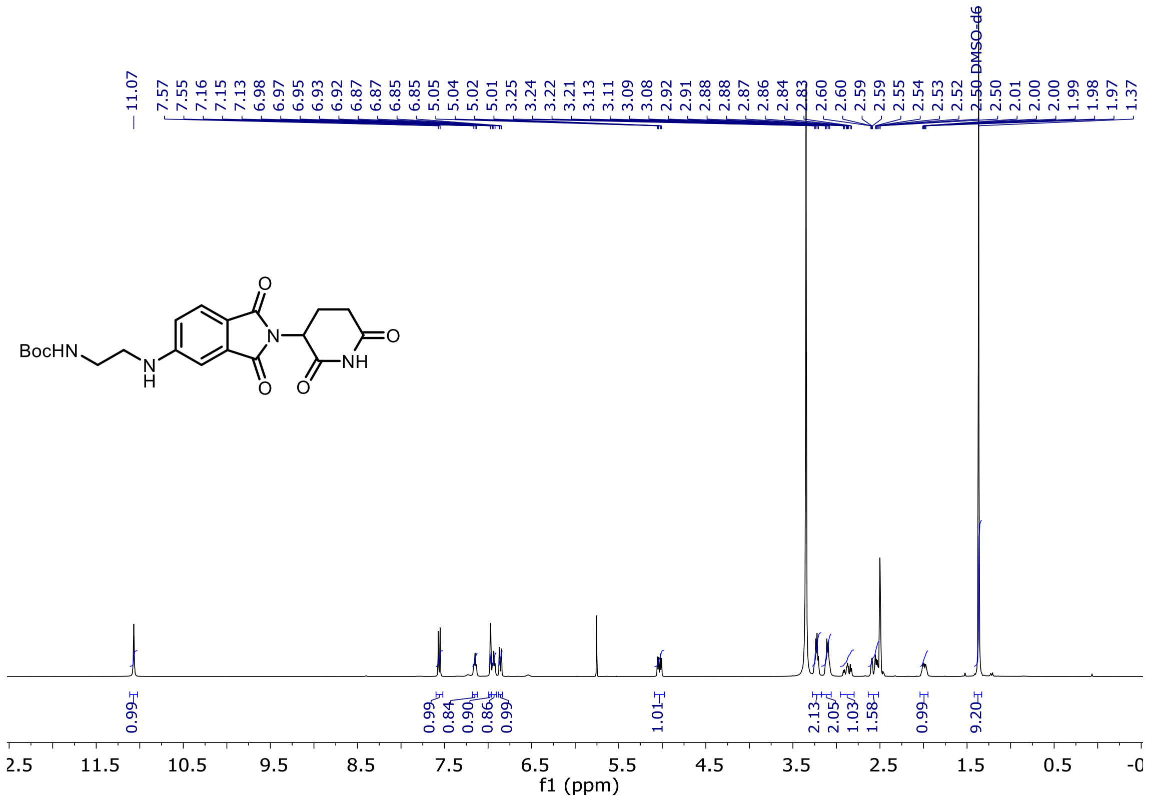

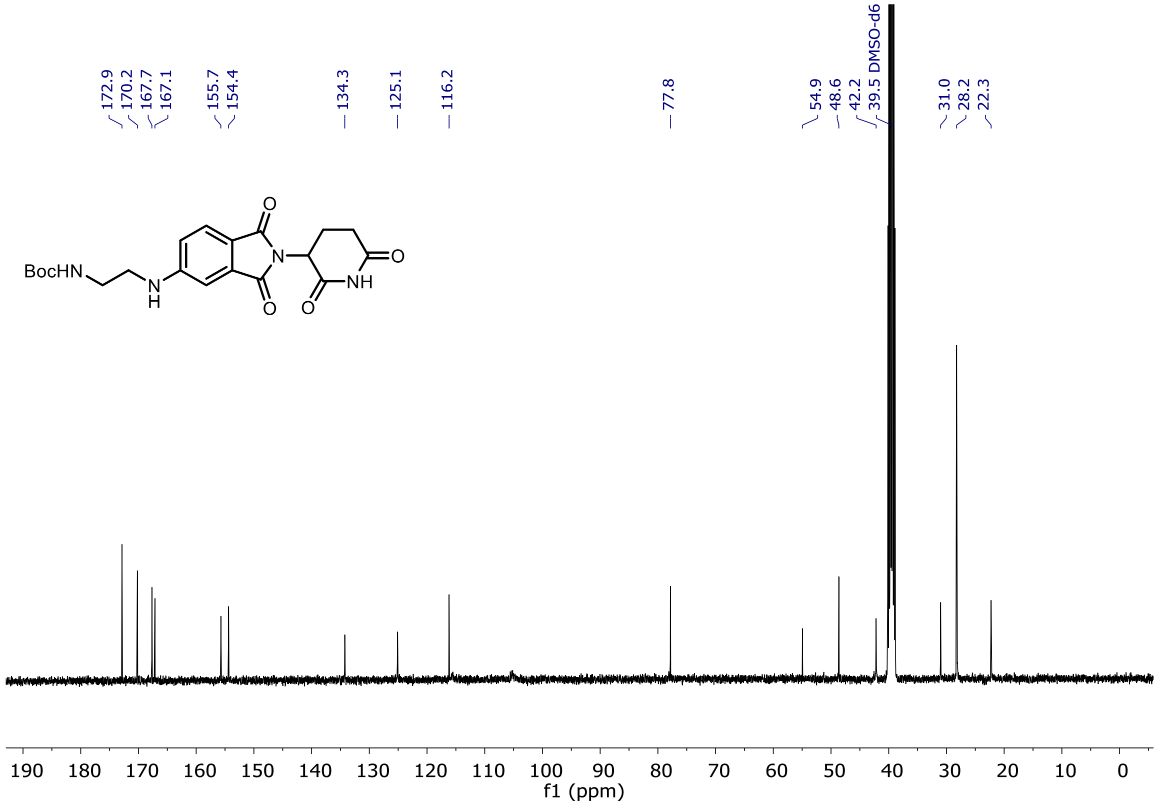

[1]H and [13]C NMR of **(5)-MS4078**

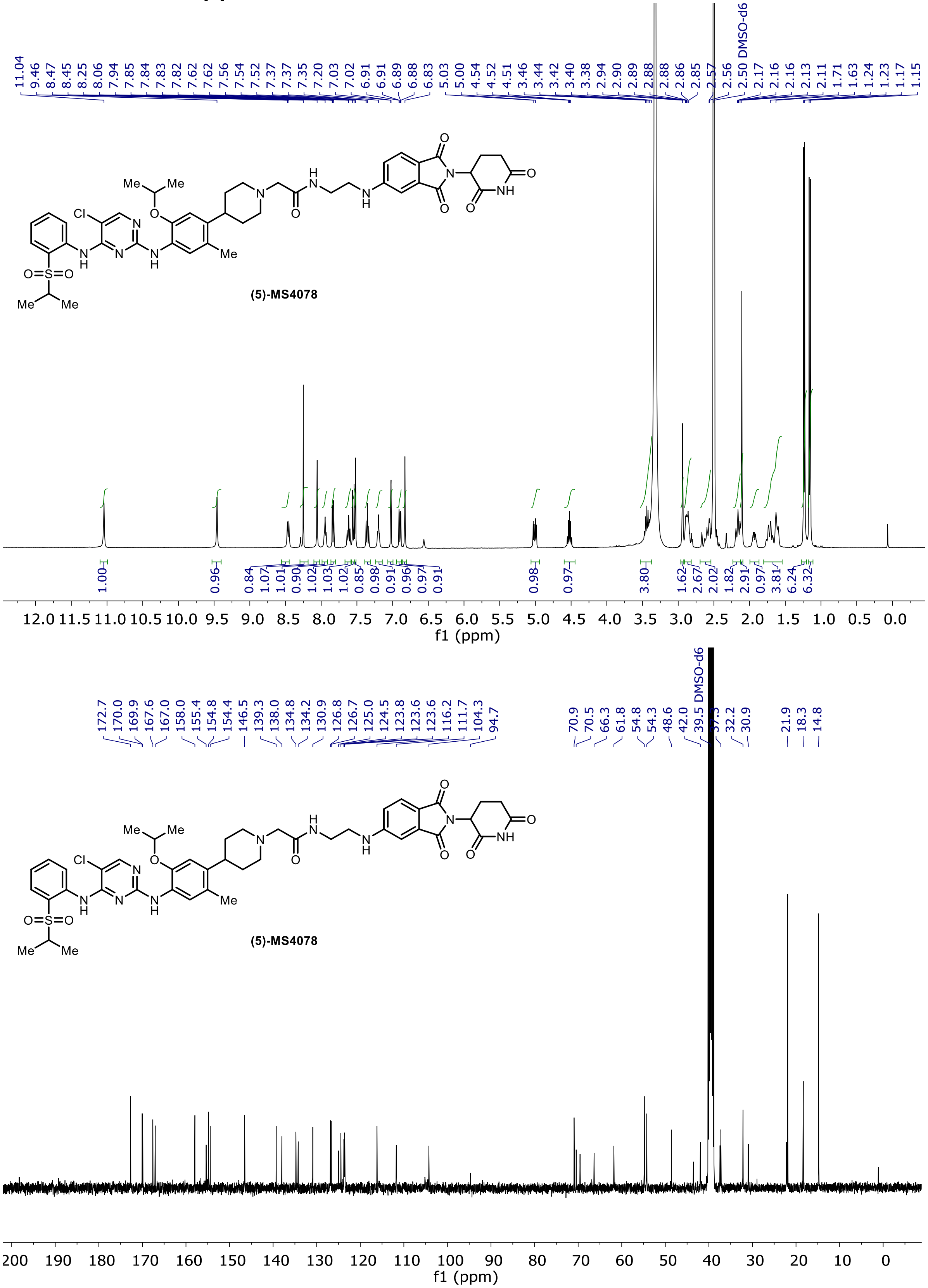
